# Tetracycline-Responsive Aptazymes Enable Tunable Expression of Self-Amplifying RNA

**DOI:** 10.64898/2026.11.10.765764

**Authors:** Owen Karmel, Herbert Fountain, Perry Katsarakes, Pratyusha Mandal, Daniel Liu, Alexander Pietraru, Yanru Cui

## Abstract

Self-amplifying RNA (saRNA) enables prolonged protein expression at substantially lower doses than conventional mRNA. Beyond vaccination, uncontrolled saRNA replication raises safety concerns. Overexpressing the gene of interest exhausts cellular resources and leads to immunogenic double-stranded saRNA. Here, we investigated Tet-On and Tet-Off tetracycline-responsive aptazymes to control saRNA amplification exogenously. Insertion of the aptazymes within the 3′ untranslated region (UTR) resulted in significantly less eGFP expression, while insertion in the subgenomic promoter region maintained expression. Additionally, we found that 33% N1-methyl-pseudouridine (m1Ψ) substitution maintained eGFP expression, while complete substitution dramatically reduced it. At physiologically relevant tetracycline doses, Tet-On and Tet-Off aptazymes lowered eGFP expression 1.5-fold and 2.1-fold, respectively, relative to control saRNA. These *in vitro* results demonstrate saRNA with tunable expression.

## Introduction

Self-amplifying RNA (saRNA) is a promising platform for RNA-based gene expression. Derived from the genome of an RNA-based virus, saRNA platforms typically replace a gene of interest (GOI) with viral capsid proteins, leaving the viral RNA-dependent RNA polymerase (RdRp) and untranslated regions identical to those of the wild-type virus. Self-amplification offers the advantages of requiring a 10-to 50-fold lower initial dose and providing up to a month of expression time^1^. The technology has progressed into the clinic with the approval of ARCT-154, an saRNA-based vaccine for the prevention of COVID-19^2^. The long expression time of saRNA offers unique advantages for broader gene therapy applications^3^, such as protein replacement therapy^4^ or the production of signaling peptides^5^. However, unlike in vaccines, the immunogenicity associated with saRNA in these applications is undesirable. Immunogenicity is primarily due to the presence of double-stranded RNA during replication^6^. At high concentrations, the RdRp can also directly affect cell viability due to its viral origin^6^. Furthermore, a high replication rate is cytotoxic due to overburdening of the host’s cellular machinery^6^. If it were possible to exogenously tune the amount of replication, cytotoxicity from each of these effects could be reduced, improving the safety of saRNA.

Aptazymes are RNA-based riboswitching devices that self-cleave in response to a small molecule, allowing RNA-based control of gene expression. Aptazyme-mediated control of alphaviruses has been explored^7,8^. However, these studies have focused on viral replication rather than application in saRNA platforms. Furthermore, they utilize high doses of theophylline, which are unsuited for clinical use. We focus on tetracycline, which is safe, well understood, and routinely used in clinical settings^9^. Meanwhile, others have regulated different parts of saRNA, such as the RdRp^10,11^ or GOI^12^, but have not focused on the genomic RNA strand, which could be a strategy to robustly stop replication.

Here we report Tet-On and Tet-Off platforms for direct conditional control of genomic saRNA using tetracycline aptazymes (Figure 1). After assessing various placements of the aptazyme, we demonstrate that expression decreases with both the K4 aptazyme (Tet-On) and Tc40 aptazyme (Tet-Off). This strategy addresses the challenge of controlling saRNA by directly regulating the genomic saRNA strand. By using a common saRNA system and focusing on doses of tetracycline readily achievable in the body^13^, our *in vitro* results suggest a translationally relevant method to control saRNA for use beyond vaccines.

**Figure 1.**
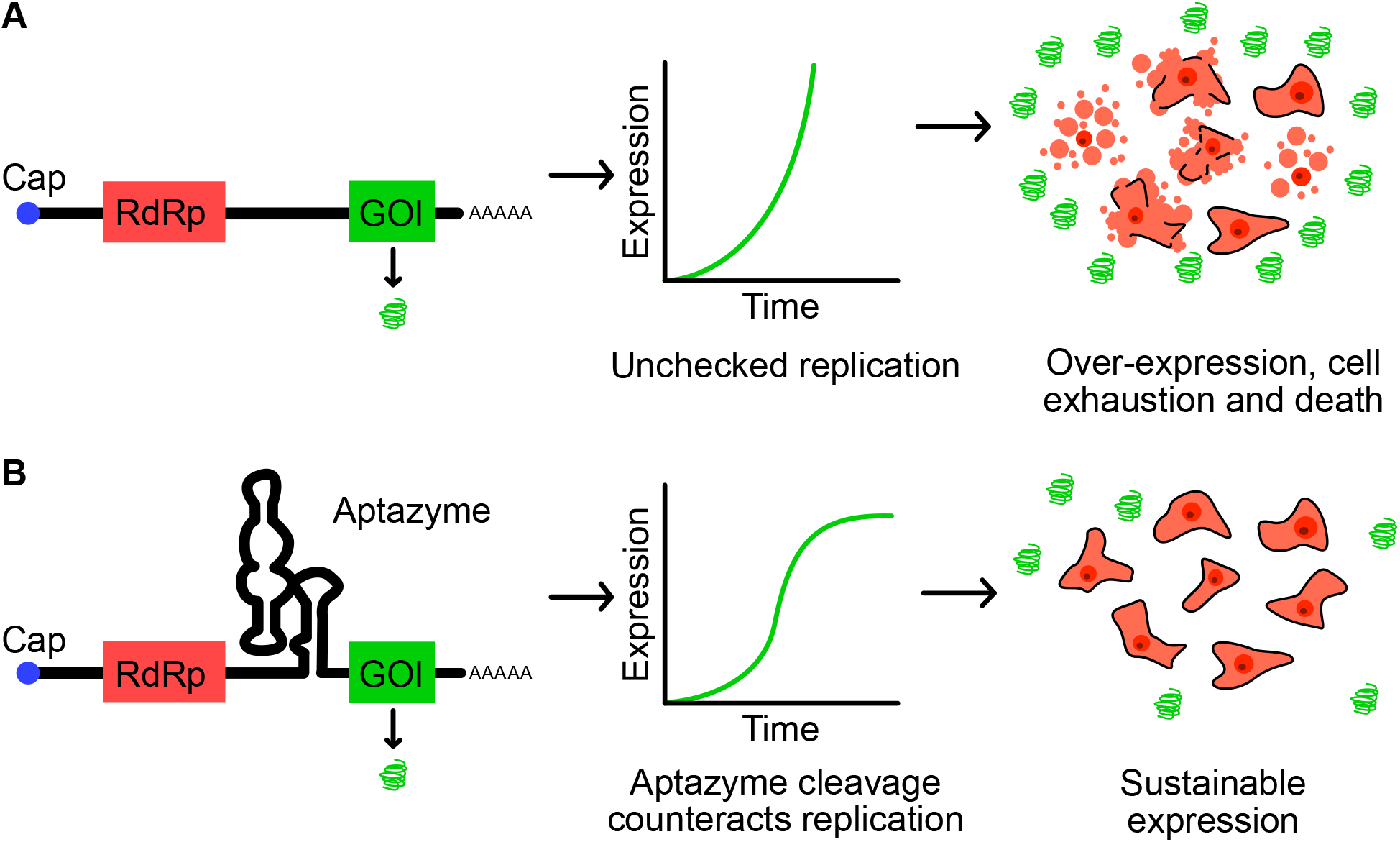
Aptazymes provide tunable control of saRNA amplification at the RNA level. Alphavirus-based saRNA consists of a 5′ cap, four nonstructural proteins collectively forming an RdRp complex, a gene of interest, and a poly(A) tail. (A) Replication occurs indefinitely until RdRp and GOI production exhaust cellular resources, resulting in cell death. (B) In regulated saRNA, aptazyme-mediated cleavage and degradation constrain saRNA copy number and, therefore, expression. Exogenous regulation of the aptazyme (via tetracycline) enables sustained expression of the gene of interest.

## Results and Discussion

We utilized the genome of the Venezuelan equine encephalitis virus, a platform often explored in translational studies and the same alphavirus used in ARCT-154^2^. In this saRNA system, four nonstructural proteins are encoded and assembled to form the RdRp complex. We engineered several parameters to increase the safety and controllability of saRNA. These design aspects included aptazyme type, aptazyme placement, and modified nucleotide content.

We chose an aptazyme (K4) from a class of tetracycline-responsive, hammerhead-ribozyme-based riboswitches that have enabled control of gene expression in mammalian cell culture^14^, *C. elegans*^15^, and mice^16^. The autocatalytic activity of these aptazymes is inactivated by tetracycline, preventing the RNA strand from being degraded (Figure 3A). Importantly, regulation occurs through self-cleavage without exogenous protein expression. To assess aptazyme function, we measured its degradation in a time-course experiment (Figure S1E). We confirmed that a physiologically relevant (10 µM) dose of tetracycline slowed aptazyme cleavage.

**Figure 2.**
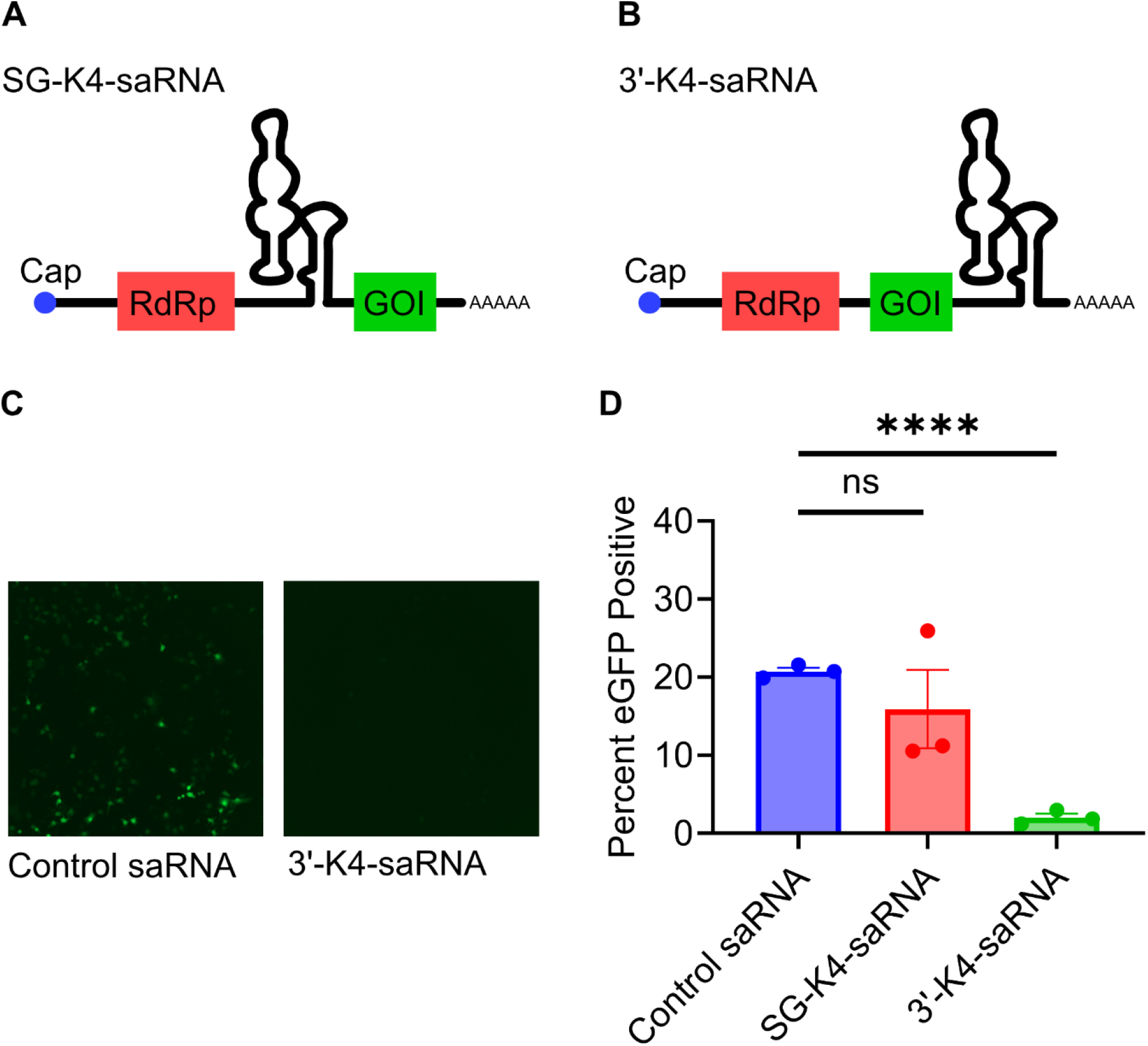
Aptazyme placement affects eGFP expression. (A, B) K4 was inserted in the subgenomic promoter region (SG-K4-saRNA; A) or in the 3′ UTR downstream of the GOI (3′-K4-saRNA; B). (C) Representative fluorescence images of cells transfected with control saRNA or 3′-K4-saRNA. (D) Flow cytometry showed markedly lower eGFP expression with 3′-K4-saRNA, whereas SG-K4-saRNA did not differ significantly from the control. Data are mean ± SEM from three biological replicates. ****p < 0.0001; ns, not significant.

**Figure 3.**
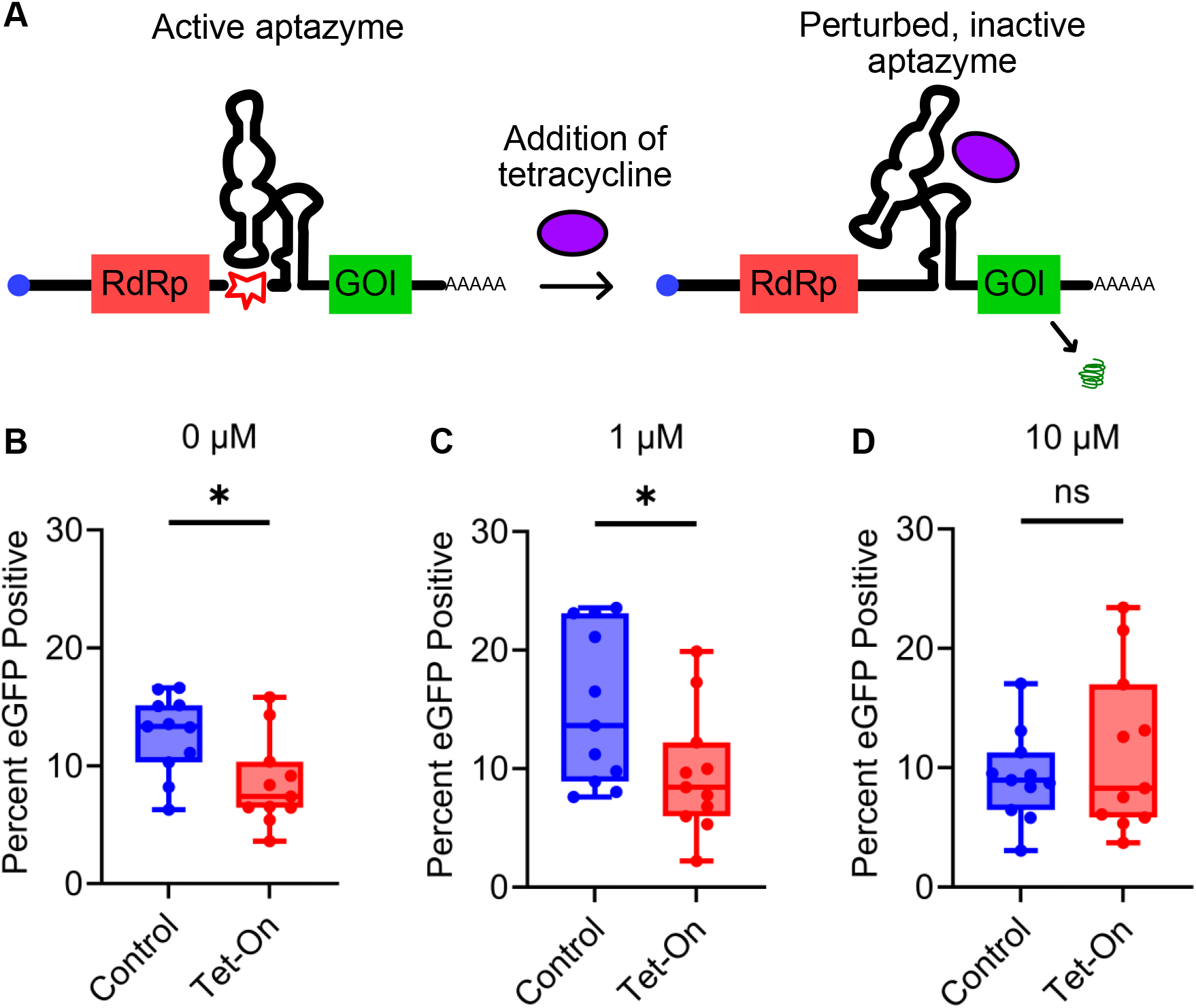
The Tet-On aptazyme modulates eGFP expression from saRNA. (A) K4 undergoes autocatalytic cleavage in the absence of tetracycline. Tetracycline disrupts the active aptazyme conformation, inhibiting cleavage and allowing GOI expression. (B, C) Tet-On saRNA showed significantly lower eGFP expression than control saRNA at 0 and 1 µM tetracycline, respectively. (D) At 10 µM, expression did not differ significantly from the control. Box-and-whisker plots show 11 biological replicates. *p < 0.05; ns, not significant.

We then assessed two locations for aptazyme insertion. In mRNA, the aptazyme can be placed in either the 5′ or 3′ untranslated region (UTR), with the 3′ location exhibiting a greater dynamic range in most studies^7,14^. Unlike in mRNA, alphavirus saRNA has a UTR between the RdRp and GOI, where the subgenomic promoter is present. We have not previously explored inserting the aptazyme in this region; therefore, we compared insertion at the 3′ end with insertion in the subgenomic promoter region. Fluorescence imaging of transfected cells revealed almost no expression from the 3′ UTR construct compared with control saRNA lacking the aptazyme (Figure 2C). Flow cytometry data confirmed this result and showed that the subgenomic UTR construct achieved expression comparable to the control saRNA (Figure 2D). For subsequent experiments, we inserted aptazymes into the subgenomic promoter region.

To mitigate immunogenicity, modified nucleotides such as N1-methyl-pseudouridine (m1Ψ) are often utilized^17^. As previously shown^18^, complete replacement of uridine with m1Ψ dramatically reduces eGFP expression (Figure S1D). However, we showed that 33% replacement with m1Ψ had no significant impact on eGFP expression compared with saRNA synthesized with 100% uridine (Figure S1D). Mitigating immunogenicity with partial m1Ψ substitution would be particularly important for non-vaccine uses of saRNA. We conducted gene expression experiments in Huh-7 cells, a liver-derived cell line, because most non-viral RNA vectors show significant liver tropism. Using flow cytometry, we confirmed that our control saRNA achieved expression comparable to commercial saRNA (Figure S1A). Interestingly, we observed that incorporating a short (120-nt) poly(A) tail directly into the DNA *in vitro* transcription (IVT) template was sufficient for high expression, while a poly(A) tail lengthened by poly(A) polymerase reduced expression (Figure S1B).

We then tested eGFP expression from Tet-On saRNA compared with the saRNA control without aptazyme. We assessed no tetracycline, 1 µM tetracycline, and 10 µM tetracycline, which are doses achievable *in vivo*. In contrast, 100 µM tetracycline caused a nonspecific decrease in expression (Figure S1C). Flow cytometry analysis of eGFP expression showed significant decreases of 1.5- and 1.6-fold in the 0 µM and 1 µM tetracycline groups (Figure 3B, C). However, no significant decrease in expression was observed at a concentration of 10 µM tetracycline (Figure 3D). These results demonstrate controlled saRNA expression in response to tetracycline.

Efficient Tet-Off aptazymes have been reported with high dynamic ranges^18^. Like Tet-On platforms, Tet-Off systems are based on hammerhead ribozymes and contain tetracycline-specific aptamer domains. In contrast to Tet-On, Tet-Off aptazymes cleave in the presence of tetracycline (Figure 4A). To confirm that saRNA regulation is possible using a Tet-Off system, we used the Tc40 Tet-Off aptazyme. This aptazyme had a previously reported dynamic range of 20-fold^18^.

**Figure 4.**
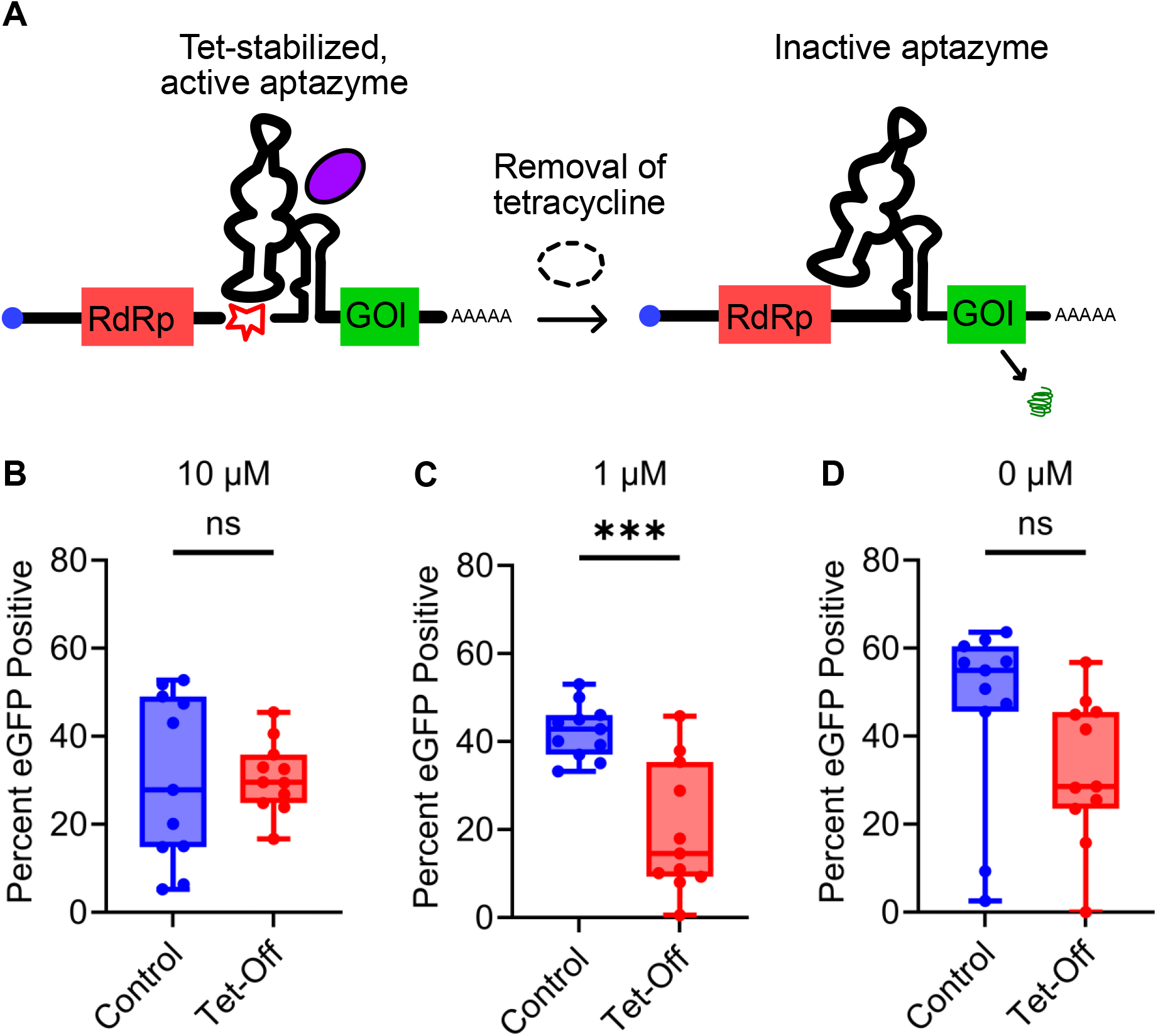
Tet-Off aptazyme regulation is effective at low tetracycline doses. (A) Tc40, a Tet-Off aptazyme, is stabilized and performs autocatalytic cleavage in the presence of tetracycline; removal of tetracycline disrupts the aptazyme, resulting in no cleavage. (B) Interestingly, at a high tetracycline dose, no significant knockdown was observed. This could be due to a nonsignificant (p = 0.1) decrease in expression seen in the control at high tetracycline doses (B, C, blue boxes), suggesting a small effect of tetracycline itself on eGFP expression. (C) At a moderate tetracycline dose, significant knockdown was observed, suggesting aptazyme cleavage reduced expression. (D) No significant knockdown was observed without tetracycline. Box-and-whisker plots show 11 biological replicates. ***p < 0.001; ns, not significant.

To investigate Tet-Off regulation of saRNA, Tc40 was cloned into our saRNA construct in the subgenomic promoter UTR. Flow cytometry revealed a significant decrease in expression with 1 µM tetracycline (Figure 4C); this decrease was not observed when tetracycline was absent (Figure 4D). The fold change of 2.1 is greater than what was observed for the Tet-On system; this was likely due to the 20-fold dynamic range of the Tc40 aptazyme in comparison to the 4-fold dynamic range of K4^14^.

Interestingly, no significant change in expression was observed in response to 10 µM tetracycline (Figure 4B). This may be explained by the nonspecific effect of tetracycline on eGFP expression. Expression decreased with increasing tetracycline levels in the control saRNA groups (Figure 4B, C, and D, blue boxes). When the tetracycline concentration was increased to 10 µM, a nonspecific decrease in eGFP was already occurring in both the control saRNA and Tc40-regulated saRNA. This overall decrease in expression may have masked a smaller tetracycline-specific response.

We investigated both Tet-On and Tet-Off aptazyme systems to regulate saRNA. We explored m1Ψ substitution and aptazyme placement, then tested the Tet-On system *in vitro* and achieved a one-third reduction in expression. We also achieved regulation using a Tet-Off aptazyme, which showed a larger reduction in expression. By showing that tetracycline-responsive aptazymes can tune gene expression in saRNA, our work identifies a potential avenue to improve saRNA safety in applications beyond vaccines.

## Methods

### Vector Construction

The saRNA plasmid was ordered from Addgene (plasmid #58977). Then, Gibson assembly was used to add a 120-nt poly(A) tail, non-essential sequences were removed to reduce the overall plasmid size and shorten the transcript length, and a CleanCap binding site was introduced to enable co-transcriptional capping. Commercial eGFP saRNA (Genscript) was used as a positive control.

The K4 aptazyme sequence was ordered as a gene from Integrated DNA Technologies. For the aptazyme degradation assay, restriction enzymes XhoI and EcoRI (NEB) were used for digestion and ligation into a longer backbone (GFP-expressing mRNA) more suitable for visualization on the gel. Gibson assembly^19^ was performed using NEB Gibson Assembly Master Mix for 1 hour at 50 °C at a 5:1 insert-to-backbone molar ratio according to the manufacturer’s protocol. Linearization of the saRNA backbone was performed using NdeI for insertion into the subgenomic promoter UTR and XbaI for insertion into the 3′ UTR. Tc40 was ordered as a gene fragment and was inserted into the subgenomic promoter UTR via Gibson assembly following backbone linearization using NdeI and AscI.

### *In Vitro* Transcription

Plasmids were linearized using MluI. Using an IVT kit (Takara Bio), 20 µL reactions were performed with 1 µg of linearized DNA, CleanCap AG (Trilink N-7113), and kit reagents, according to the manufacturer’s protocol. We replaced uridine with N1-methyl-pseudouridine (Trilink N-1081) at a 1:1 molar ratio, at 0%, 33%, or 100%. The solution was incubated at 37 °C for 2 hours, then incubated for an additional 15 minutes with RNase-free DNase I and purified using a 500 µg RNA spin column (NEB).

### *In Vitro* Aptazyme Degradation Assay

Aptazyme-containing RNA in the longer backbone was synthesized via IVT and then immediately placed on ice. Aqueous RNA was incubated at 37 °C at a concentration of 100 ng/µL with 2 mM manganese ions (a coenzyme of hammerhead ribozymes^20^) and with or without 1.125 µM tetracycline. Upon reaching a given timepoint, 800 ng of RNA (8 µL of solution) was mixed with RNA Loading Dye (Thermo Scientific) and kept on ice to stop the reaction. After all timepoints were completed, the solutions were heated at 70 °C for 15 minutes, then run on a 3% TBE gel.

### Cell Culture

Huh-7 cells were grown in complete DMEM (Thermo Fisher), supplemented with 10% fetal bovine serum (FBS), 1% Pen-Strep, and 10 µg/mL Fungin (Invitrogen), and incubated in a 5% CO_2_ humidified atmosphere at 37 °C. Cells were passaged every 3–4 days. Twenty-four hours before transfection, 96-well plates (Corning) were seeded with 10,000 live Huh-7 cells per well in 100 µL of medium. According to the manufacturer’s protocol, 0.3 µL of Lipofectamine 3000 reagent (Thermo Fisher) and 100 ng of RNA per well were used for transfection. Transfections were performed in OptiMEM without FBS. When applicable, tetracycline (aqueous, 2.5 mM) (Sigma-Aldrich) was added during transfection to a final concentration of 1, 10, or 100 µM.

### Fluorescence Imaging and Flow Cytometry

Twenty-four hours post-transfection, a single representative replicate per experimental group was Hoechst-stained and imaged using a multi-well plate reader for fluorescence detection (Cytation 7). Live-dead staining (eBioscience) was performed on the remaining biological replicates, which were then analyzed using a flow cytometer (Agilent NovoCyte Quanteon). After dead cells were excluded, the baseline of the eGFP gate was set against an untransfected control. To test for significance, an unpaired Welch’s t-test was conducted between samples while assuming sampling from normal distributions.

### Supporting Information

Additional flow cytometry data and TBE gel - effect of poly(A) tailing, excess tetracycline, and nucleotide substitutions on saRNA eGFP expression (PDF)

DNA sequences of saRNA and aptazyme plasmids, gene fragments and Gibson primers (XLSX)

Raw flow cytometry data (PDF)

## Supporting information

Sequence information

Raw flow data

## Acknowledgments

The authors would like to thank Rawlings Presidential Research Fellowship for generous sponsorship of conference travel. They would like to thank Dr. Wenchao Gu from the Jiang Lab for her feedback on the project and manuscript, as well as acknowledge the Robert Langer ‘70 Family and Friends Professorship. T7-VEE-GFP was a gift from Steven Dowdy (Addgene plasmid # 58977).

