## Supplemental for "Tetracycline-Responsive Aptazymes Enable Tunable Expression of Self-Amplifying RNA"

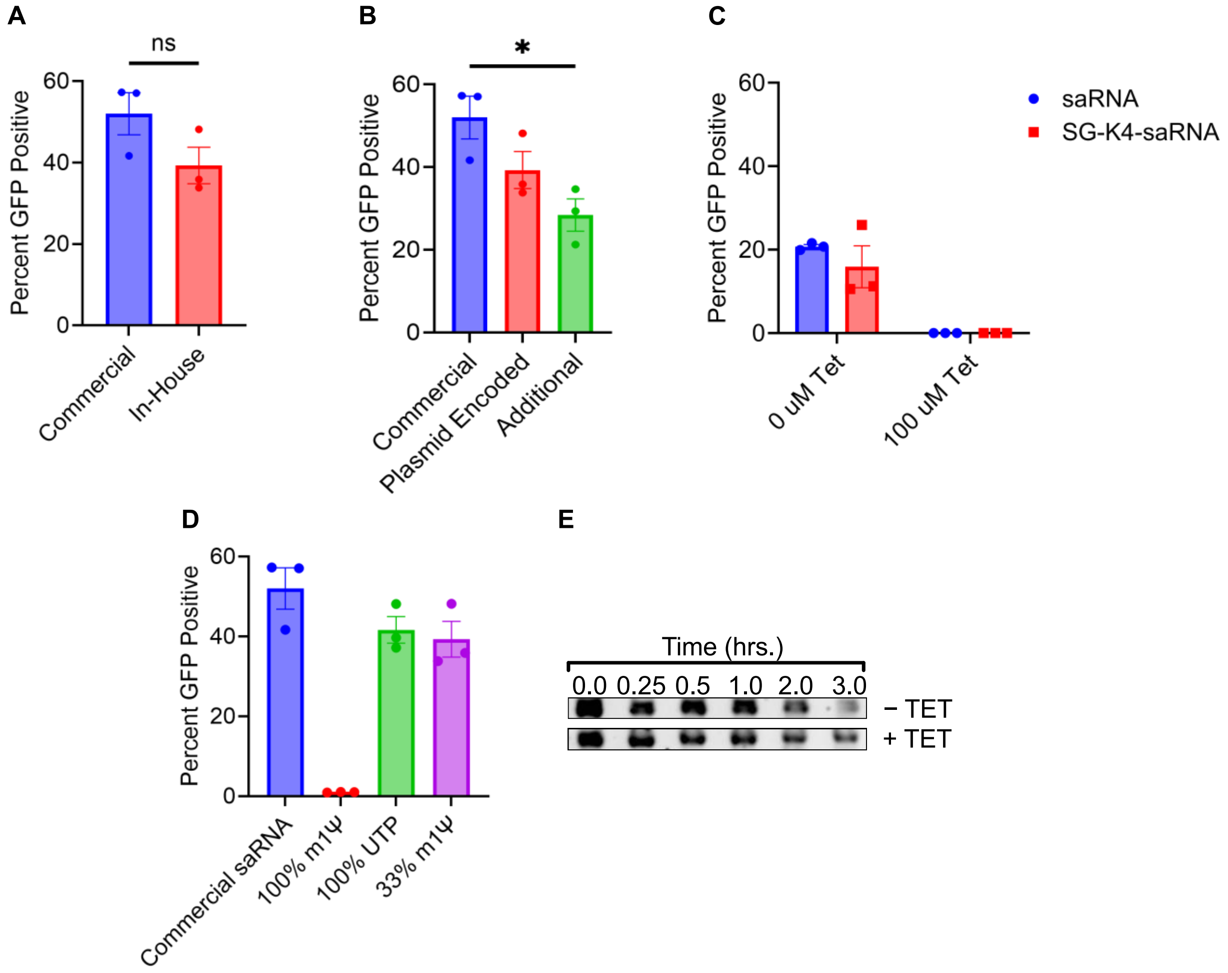


**Figure S1. Effect of poly(A) tailing, excess tetracycline, and nucleotide substitutions on saRNA eGFP expression.** (A) Our saRNA achieved comparable expression to commercial saRNA. (B) A short, plasmid-encoded poly(A) tail was sufficient for high eGFP expression; additional poly(A) tailing with poly(A) polymerase decreased expression. (C) 100 μM tetracycline greatly reduced eGFP expression in a nonspecific manner (note that these data were obtained with 0% m1Ψ saRNA with additional poly(A) tailing). (D) saRNA with 33% m1Ψ achieved comparable expression to commercial saRNA and saRNA with no modified nucleotides. In contrast, 100% m1Ψ compromised eGFP expression. (E) Visualization of *in vitro* aptazyme degradation on a 3% TBE gel with and without the addition of 1.125 μM tetracycline.
