## Supplementary material for "Tetracycline-Responsive Aptazymes Enable Tunable Expression of Self-Amplifying RNA": Raw flow data

### Report of Owen_Herb_TET_saRNA 3_11_26

File Name: Owen_Herb_TET_saRNA 3_11_26 - Analysis
Run Time: 3/11/2026 3:36 PM
Cytometer: NovoCyte Quanteon 621190610657
Software: NovoExpress 1.6.1

#### No Transfection

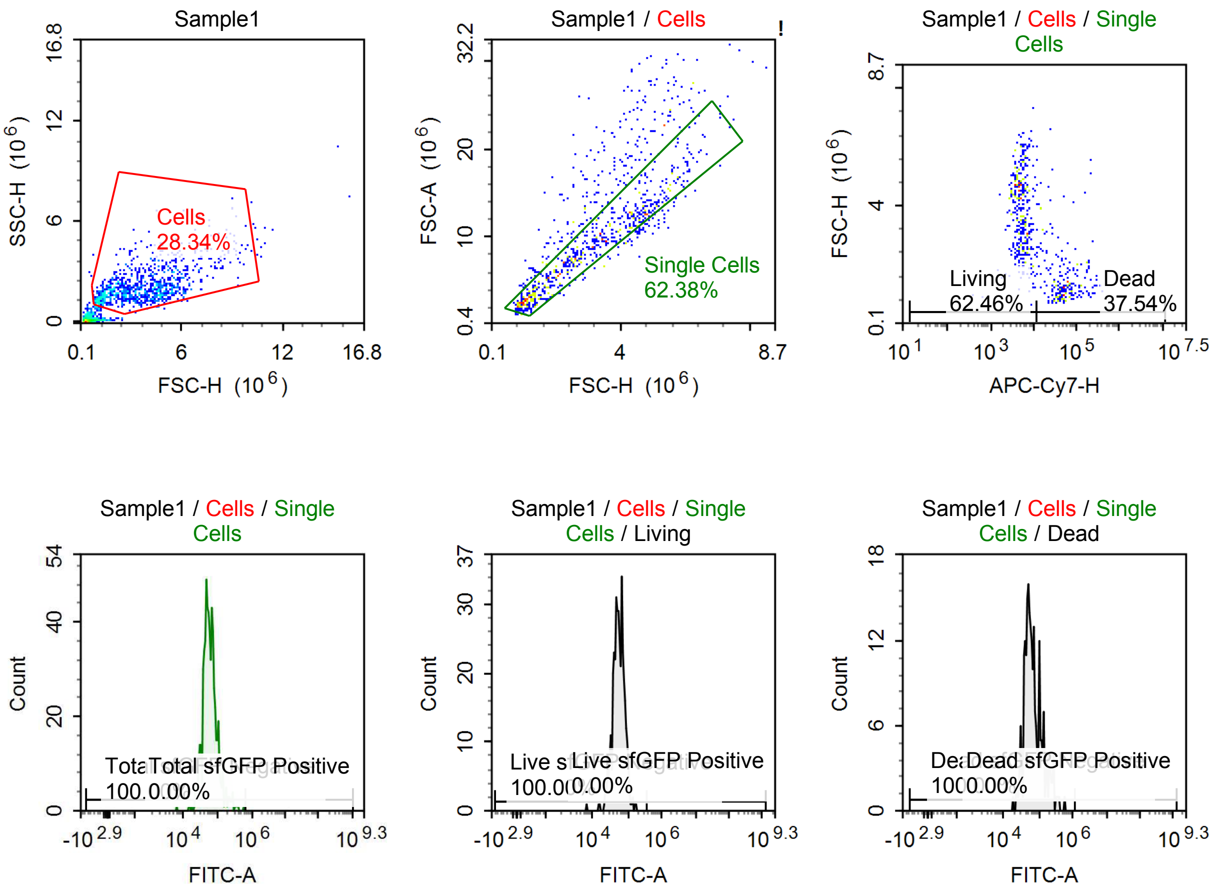


Sample Statistics of Sample1

| Gate | Count | % Parent | X | Y | Median X | Median Y |
|---|---|---|---|---|---|---|
| All | 3,405 | | | | | |
| Cells | 965 | 28.34 % | FSC-H | SSC-H | 3,508,347 | 1,913,246 |
| Single Cells | 602 | 62.38 % | FSC-H | FSC-A | 2,808,546 | 8,416,224 |
| Living | 376 | 62.46 % | APC-Cy7-H | FSC-H | 5,120 | 3,933,268 |
| Live sfGFP Negative | 376 | 100.00 % | FITC-A | Count | 54,753 | |
| Live sfGFP Positive | 0 | 0.00 % | FITC-A | Count | 0 | |
| Dead | 226 | 37.54 % | APC-Cy7-H | FSC-H | 48,326 | 1,304,638 |
| Dead sfGFP Negative | 226 | 100.00 % | FITC-A | Count | 60,801 | |
| Dead sfGFP Positive | 0 | 0.00 % | FITC-A | Count | 0 | |
| Total sfGFP Negative | 602 | 100.00 % | FITC-A | Count | 56,456 | |
| Total sfGFP Positive | 0 | 0.00 % | FITC-A | Count | 0 | |

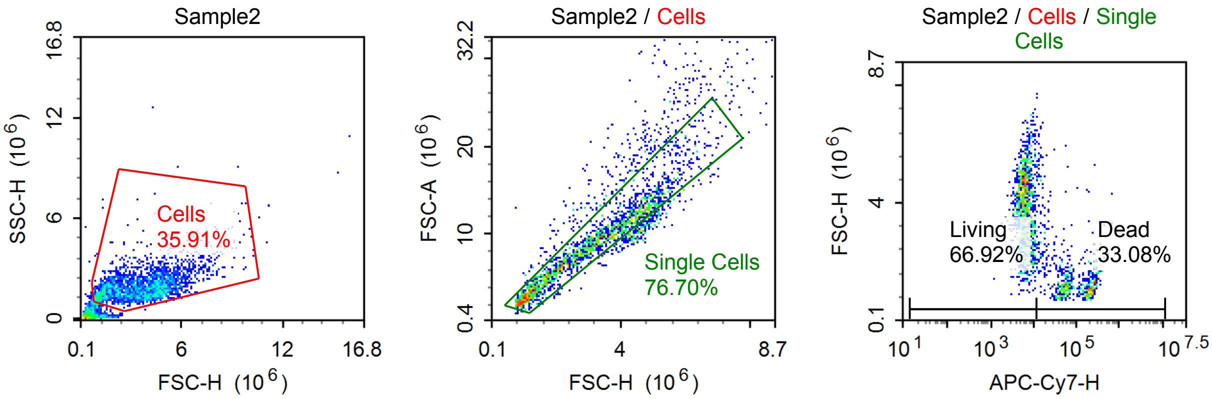


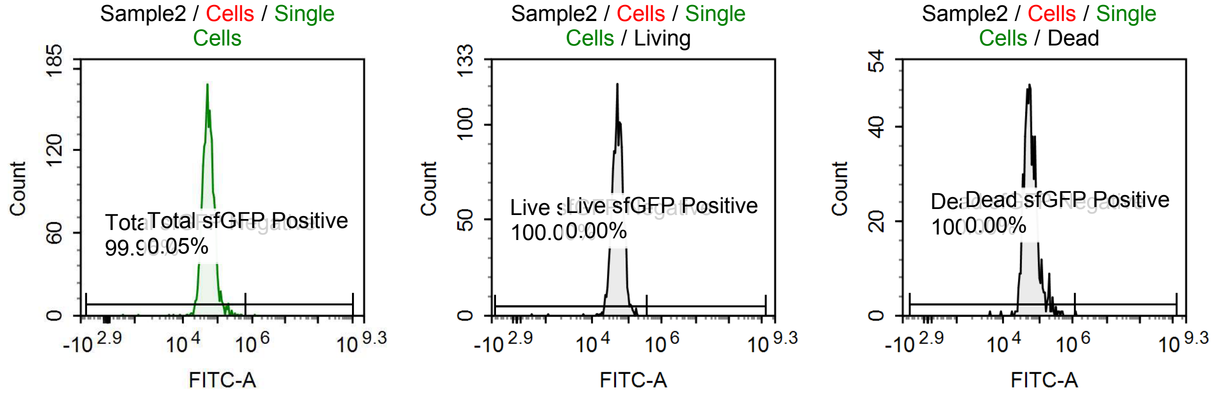


Sample Statistics of Sample2

| Gate | Count | % Parent | X | Y | Median X | Median Y |
|---|---|---|---|---|---|---|
| All | 7,134 | | | | | |
| Cells | 2,562 | 35.91 % | FSC-H | SSC-H | 3,557,061 | 1,843,180 |
| Single Cells | 1,965 | 76.70 % | FSC-H | FSC-A | 3,076,835 | 8,887,454 |
| Living | 1,315 | 66.92 % | APC-Cy7-H | FSC-H | 6,104 | 3,830,060 |
| Live sfGFP Negative | 1,315 | 100.00 % | FITC-A | Count | 51,244 | |
| Live sfGFP Positive | 0 | 0.00 % | FITC-A | Count | 0 | |
| Dead | 650 | 33.08 % | APC-Cy7-H | FSC-H | 64,178 | 1,308,988 |
| Dead sfGFP Negative | 650 | 100.00 % | FITC-A | Count | 55,174 | |
| Dead sfGFP Positive | 0 | 0.00 % | FITC-A | Count | 0 | |
| Total sfGFP Negative | 1,964 | 99.95 % | FITC-A | Count | 52,226 | |
| Total sfGFP Positive | 1 | 0.05 % | FITC-A | Count | 1,238,202 | |

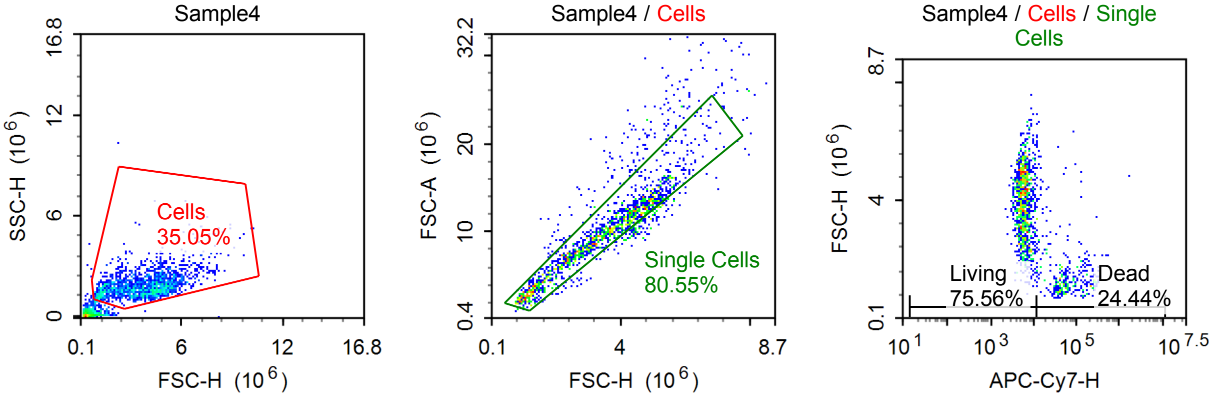

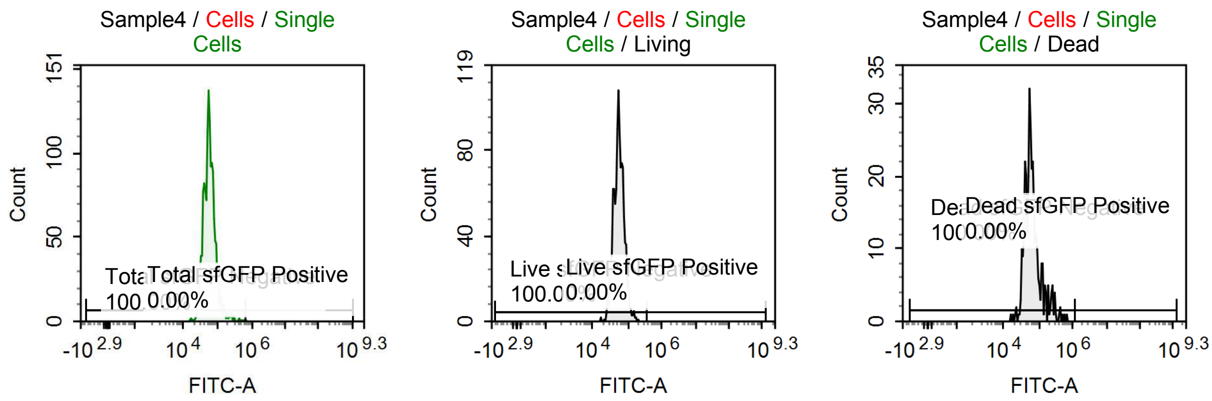


Sample Statistics of Sample4

| Gate | Count | % Parent | X | Y | Median X | Median Y |
|---|---|---|---|---|---|---|
| All | 4,884 | | | | | |
| Cells | 1,712 | 35.05 % | FSC-H | SSC-H | 3,815,447 | 1,762,727 |
| Single Cells | 1,379 | 80.55 % | FSC-H | FSC-A | 3,503,739 | 9,992,830 |
| Living | 1,042 | 75.56 % | APC-Cy7-H | FSC-H | 5,852 | 4,016,604 |
| Live sfGFP Negative | 1,042 | 100.00 % | FITC-A | Count | 54,742 | |
| Live sfGFP Positive | 0 | 0.00 % | FITC-A | Count | 0 | |
| Dead | 337 | 24.44 % | APC-Cy7-H | FSC-H | 54,471 | 1,310,279 |
| Dead sfGFP Negative | 337 | 100.00 % | FITC-A | Count | 58,787 | |
| Dead sfGFP Positive | 0 | 0.00 % | FITC-A | Count | 0 | |
| Total sfGFP Negative | 1,379 | 100.00 % | FITC-A | Count | 55,545 | |
| Total sfGFP Positive | 0 | 0.00 % | FITC-A | Count | 0 | |

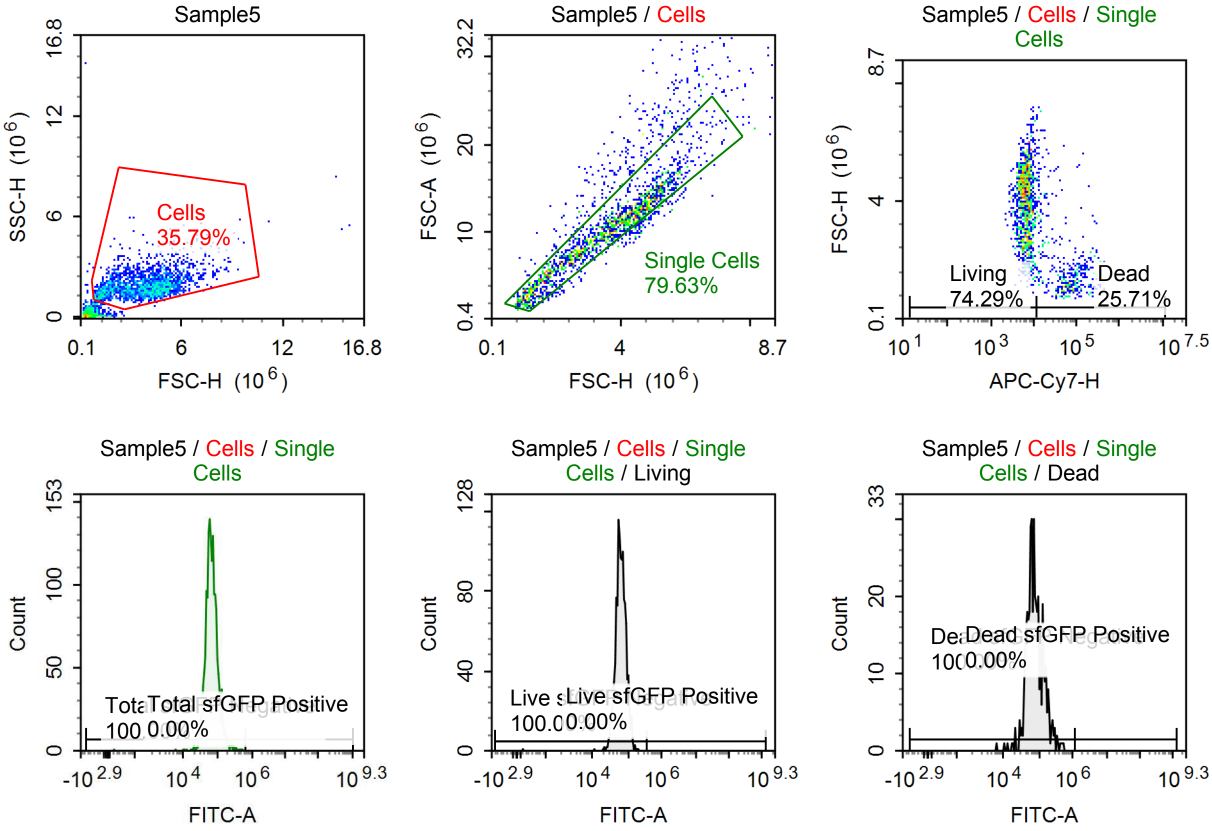

Sample Statistics of Sample5

| Gate | Count | % Parent | X | Y | Median X | Median Y |
|---|---|---|---|---|---|---|
| All | 5,596 | | | | | |
| Cells | 2,003 | 35.79 % | FSC-H | SSC-H | 3,842,502 | 1,838,610 |
| Single Cells | 1,595 | 79.63 % | FSC-H | FSC-A | 3,606,830 | 10,284,029 |
| Living | 1,185 | 74.29 % | APC-Cy7-H | FSC-H | 6,494 | 4,051,501 |
| Live sfGFP Negative | 1,185 | 100.00 % | FITC-A | Count | 62,206 | |
| Live sfGFP Positive | 0 | 0.00 % | FITC-A | Count | 0 | |
| Dead | 410 | 25.71 % | APC-Cy7-H | FSC-H | 59,038 | 1,550,417 |
| Dead sfGFP Negative | 410 | 100.00 % | FITC-A | Count | 72,293 | |
| Dead sfGFP Positive | 0 | 0.00 % | FITC-A | Count | 0 | |
| Total sfGFP Negative | 1,595 | 100.00 % | FITC-A | Count | 64,148 | |
| Total sfGFP Positive | 0 | 0.00 % | FITC-A | Count | 0 | |

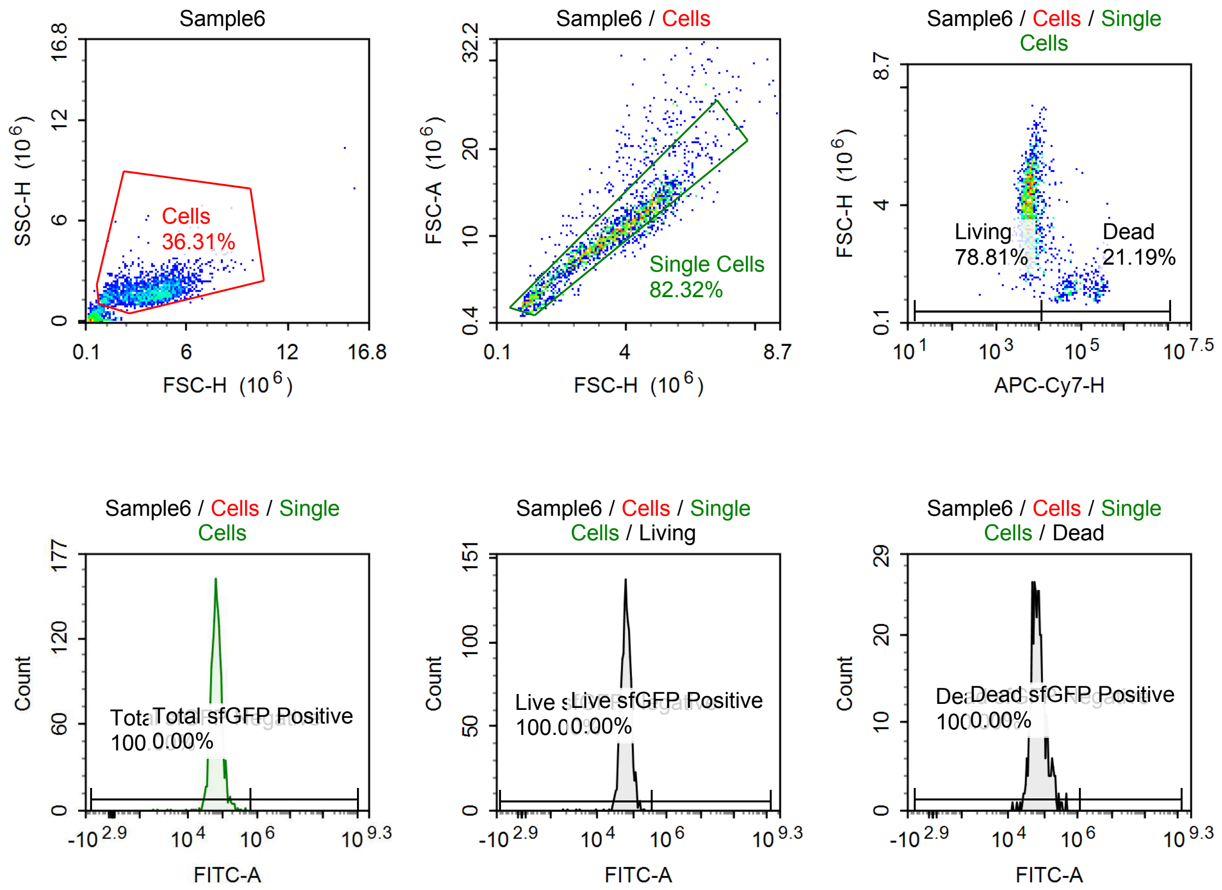

Sample Statistics of Sample6

| Gate | Count | % Parent | X | Y | Median X | Median Y |
|---|---|---|---|---|---|---|
| All | 5,794 | | | | | |
| Cells | 2,104 | 36.31 % | FSC-H | SSC-H | 3,827,725 | 1,745,340 |
| Single Cells | 1,732 | 82.32 % | FSC-H | FSC-A | 3,608,055 | 10,401,255 |
| Living | 1,365 | 78.81 % | APC-Cy7-H | FSC-H | 6,033 | 3,962,298 |
| Live sfGFP Negative | 1,365 | 100.00 % | FITC-A | Count | 63,444 | |
| Live sfGFP Positive | 0 | 0.00 % | FITC-A | Count | 0 | |
| Dead | 367 | 21.19 % | APC-Cy7-H | FSC-H | 56,392 | 1,366,433 |
| Dead sfGFP Negative | 367 | 100.00 % | FITC-A | Count | 66,292 | |
| Dead sfGFP Positive | 0 | 0.00 % | FITC-A | Count | 0 | |
| Total sfGFP Negative | 1,732 | 100.00 % | FITC-A | Count | 63,859 | |
| Total sfGFP Positive | 0 | 0.00 % | FITC-A | Count | 0 | |

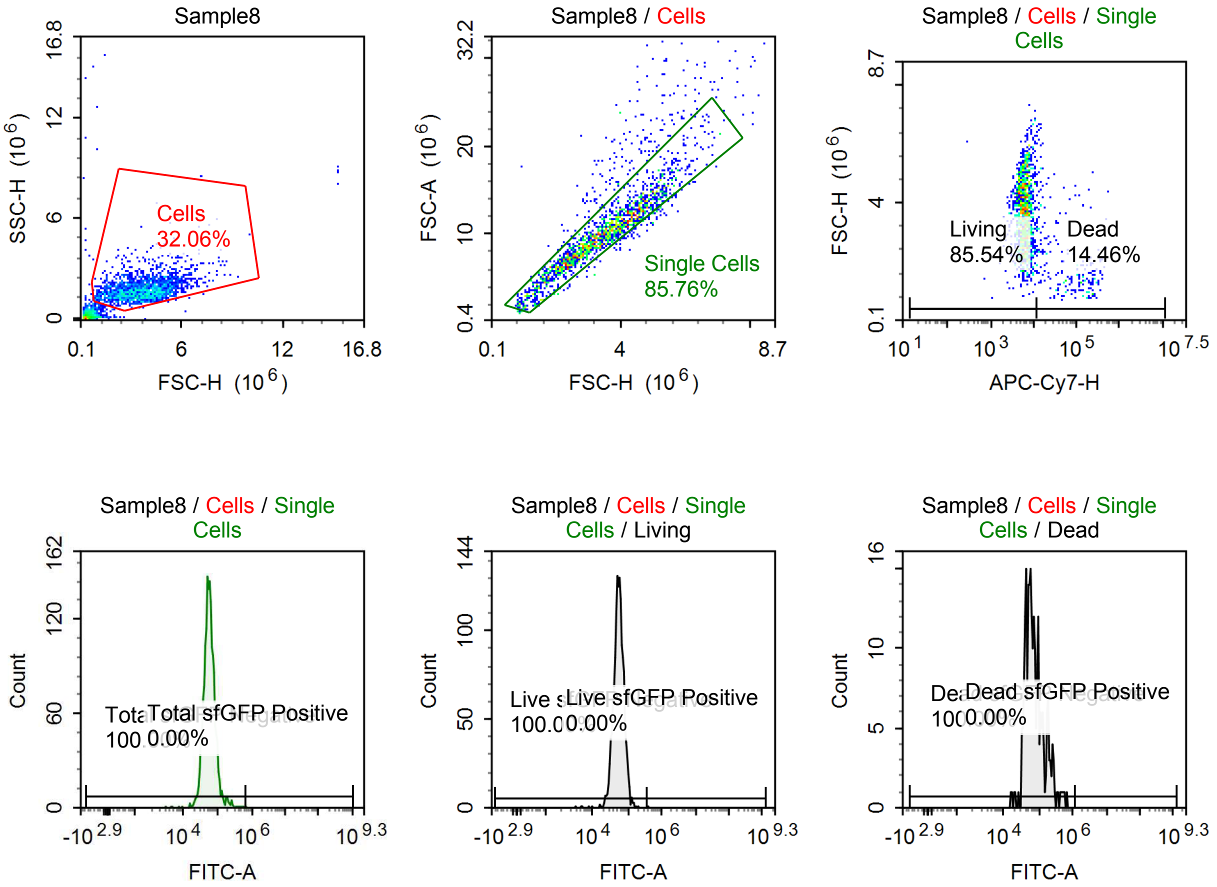

Sample Statistics of Sample8

| Gate | Count | % Parent | X | Y | Median X | Median Y |
|---|---|---|---|---|---|---|
| All | 5,936 | | | | | |
| Cells | 1,903 | 32.06 % | FSC-H | SSC-H | 3,625,208 | 1,678,777 |
| Single Cells | 1,632 | 85.76 % | FSC-H | FSC-A | 3,488,848 | 10,041,160 |
| Living | 1,396 | 85.54 % | APC-Cy7-H | FSC-H | 6,025 | 3,665,897 |
| Live sfGFP Negative | 1,396 | 100.00 % | FITC-A | Count | 54,569 | |
| Live sfGFP Positive | 0 | 0.00 % | FITC-A | Count | 0 | |
| Dead | 236 | 14.46 % | APC-Cy7-H | FSC-H | 74,174 | 1,586,360 |
| Dead sfGFP Negative | 236 | 100.00 % | FITC-A | Count | 66,418 | |
| Dead sfGFP Positive | 0 | 0.00 % | FITC-A | Count | 0 | |
| Total sfGFP Negative | 1,632 | 100.00 % | FITC-A | Count | 55,676 | |
| Total sfGFP Positive | 0 | 0.00 % | FITC-A | Count | 0 | |

#### Comercial saRNA

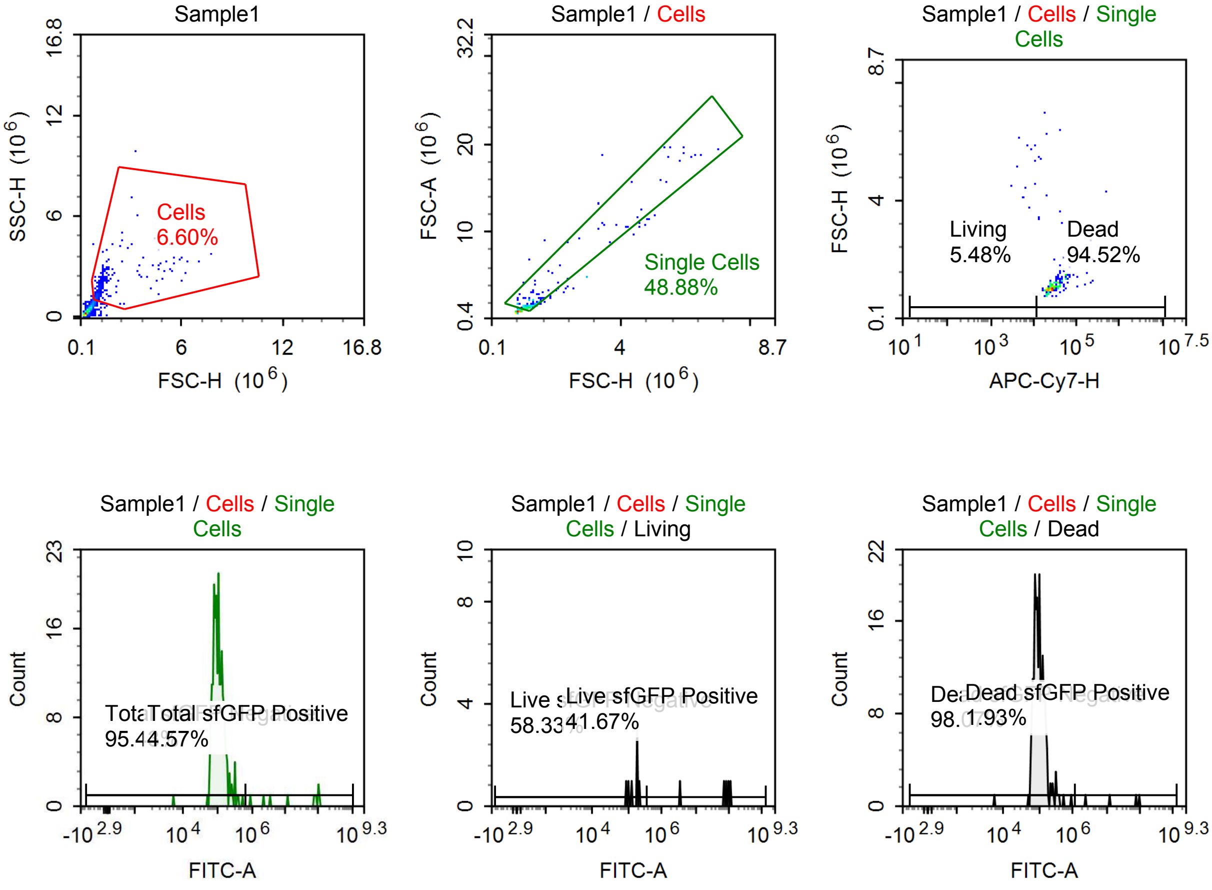

Sample Statistics of Sample1

| Gate | Count | % Parent | X | Y | Median X | Median Y |
|---|---|---|---|---|---|---|
| All | 6,788 | | | | | |
| Cells | 448 | 6.60 % | FSC-H | SSC-H | 944,358 | 1,376,343 |
| Single Cells | 219 | 48.88 % | FSC-H | FSC-A | 1,115,062 | 1,616,420 |
| Living | 12 | 5.48 % | APC-Cy7-H | FSC-H | 7,205 | 4,901,891 |
| Live sfGFP Negative | 7 | 58.33 % | FITC-A | Count | 184,330 | |
| Live sfGFP Positive | 5 | 41.67 % | FITC-A | Count | 82,877,336 | |
| Dead | 207 | 94.52 % | APC-Cy7-H | FSC-H | 26,780 | 1,100,218 |
| Dead sfGFP Negative | 203 | 98.07 % | FITC-A | Count | 97,225 | |
| Dead sfGFP Positive | 4 | 1.93 % | FITC-A | Count | 43,879,192 | |
| Total sfGFP Negative | 209 | 95.43 % | FITC-A | Count | 100,489 | |
| Total sfGFP Positive | 10 | 4.57 % | FITC-A | Count | 73,356,224 | |

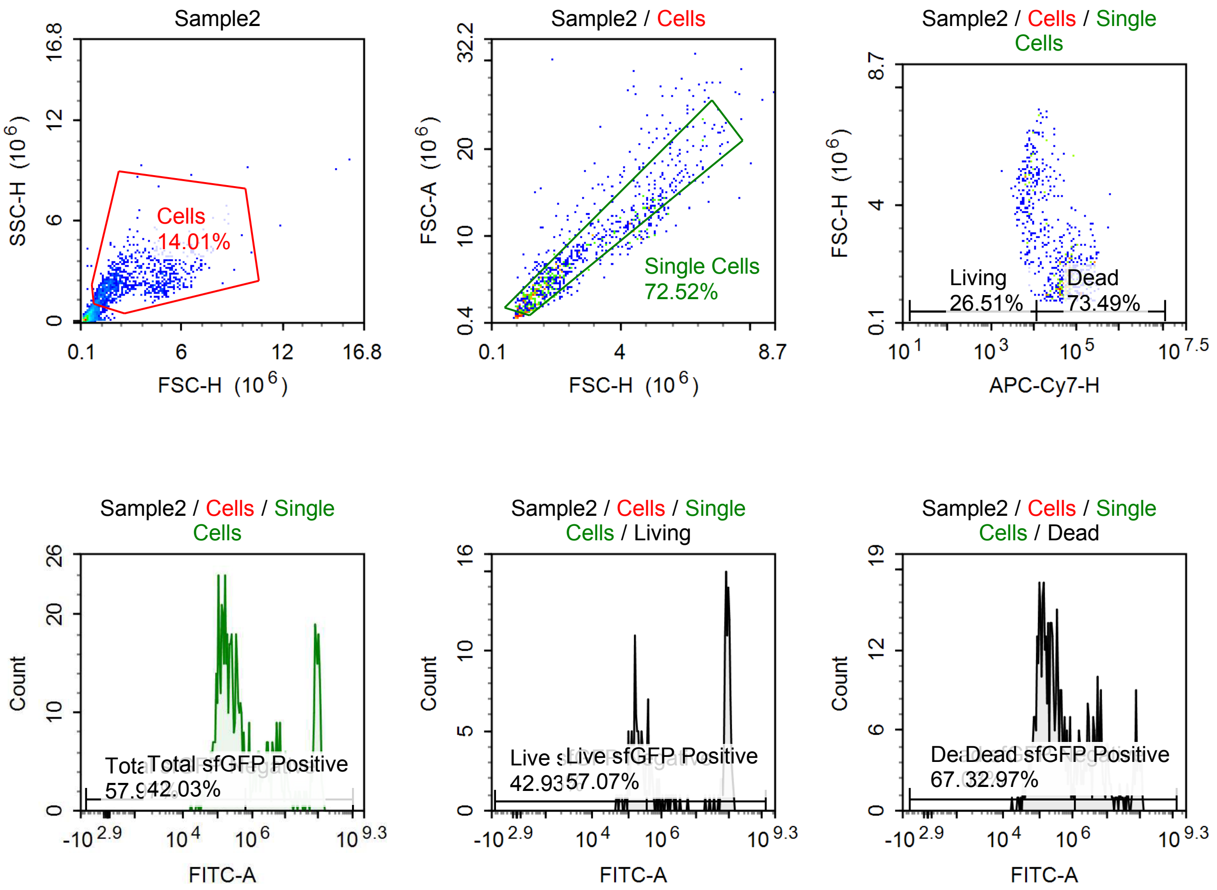

Sample Statistics of Sample2

| Gate | Count | % Parent | X | Y | Median X | Median Y |
|---|---|---|---|---|---|---|
| All | 7,353 | | | | | |
| Cells | 1,030 | 14.01 % | FSC-H | SSC-H | 2,185,967 | 2,201,542 |
| Single Cells | 747 | 72.52 % | FSC-H | FSC-A | 2,043,855 | 5,830,365 |
| Living | 198 | 26.51 % | APC-Cy7-H | FSC-H | 7,370 | 4,464,174 |
| Live sfGFP Negative | 85 | 42.93 % | FITC-A | Count | 177,775 | |
| Live sfGFP Positive | 113 | 57.07 % | FITC-A | Count | 83,599,096 | |
| Dead | 549 | 73.49 % | APC-Cy7-H | FSC-H | 55,898 | 1,674,573 |
| Dead sfGFP Negative | 368 | 67.03 % | FITC-A | Count | 180,659 | |
| Dead sfGFP Positive | 181 | 32.97 % | FITC-A | Count | 6,387,026 | |
| Total sfGFP Negative | 433 | 57.97 % | FITC-A | Count | 170,150 | |
| Total sfGFP Positive | 314 | 42.03 % | FITC-A | Count | 9,532,928 | |

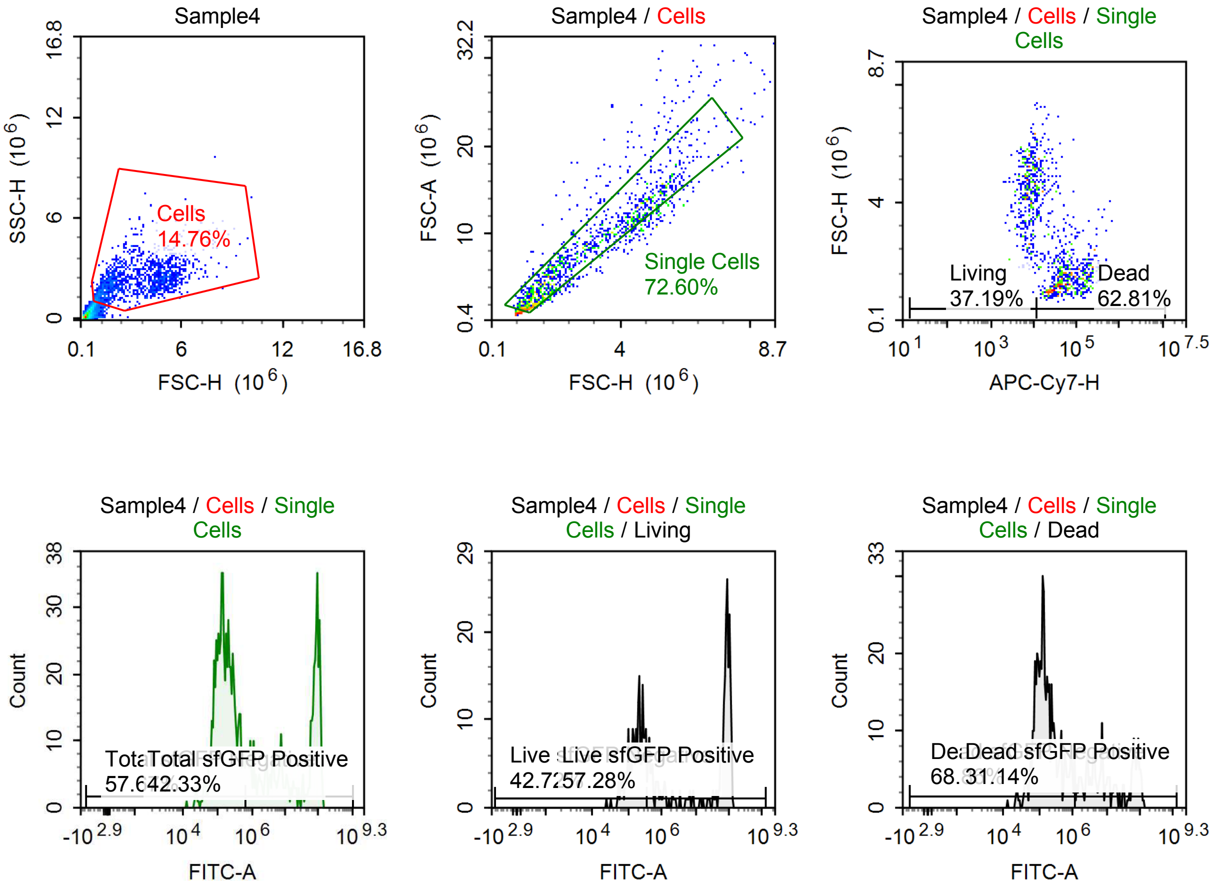

Sample Statistics of Sample4

| Gate | Count | % Parent | X | Y | Median X | Median Y |
|---|---|---|---|---|---|---|
| All | 10,163 | | | | | |
| Cells | 1,500 | 14.76 % | FSC-H | SSC-H | 2,491,301 | 2,085,273 |
| Single Cells | 1,089 | 72.60 % | FSC-H | FSC-A | 2,345,172 | 6,956,062 |
| Living | 405 | 37.19 % | APC-Cy7-H | FSC-H | 7,401 | 4,415,830 |
| Live sfGFP Negative | 173 | 42.72 % | FITC-A | Count | 191,965 | |
| Live sfGFP Positive | 232 | 57.28 % | FITC-A | Count | 78,173,496 | |
| Dead | 684 | 62.81 % | APC-Cy7-H | FSC-H | 45,154 | 1,608,597 |
| Dead sfGFP Negative | 471 | 68.86 % | FITC-A | Count | 142,928 | |
| Dead sfGFP Positive | 213 | 31.14 % | FITC-A | Count | 9,412,337 | |
| Total sfGFP Negative | 628 | 57.67 % | FITC-A | Count | 149,760 | |
| Total sfGFP Positive | 461 | 42.33 % | FITC-A | Count | 39,585,160 | |

#### p059 poly-a talied, 100% m1U

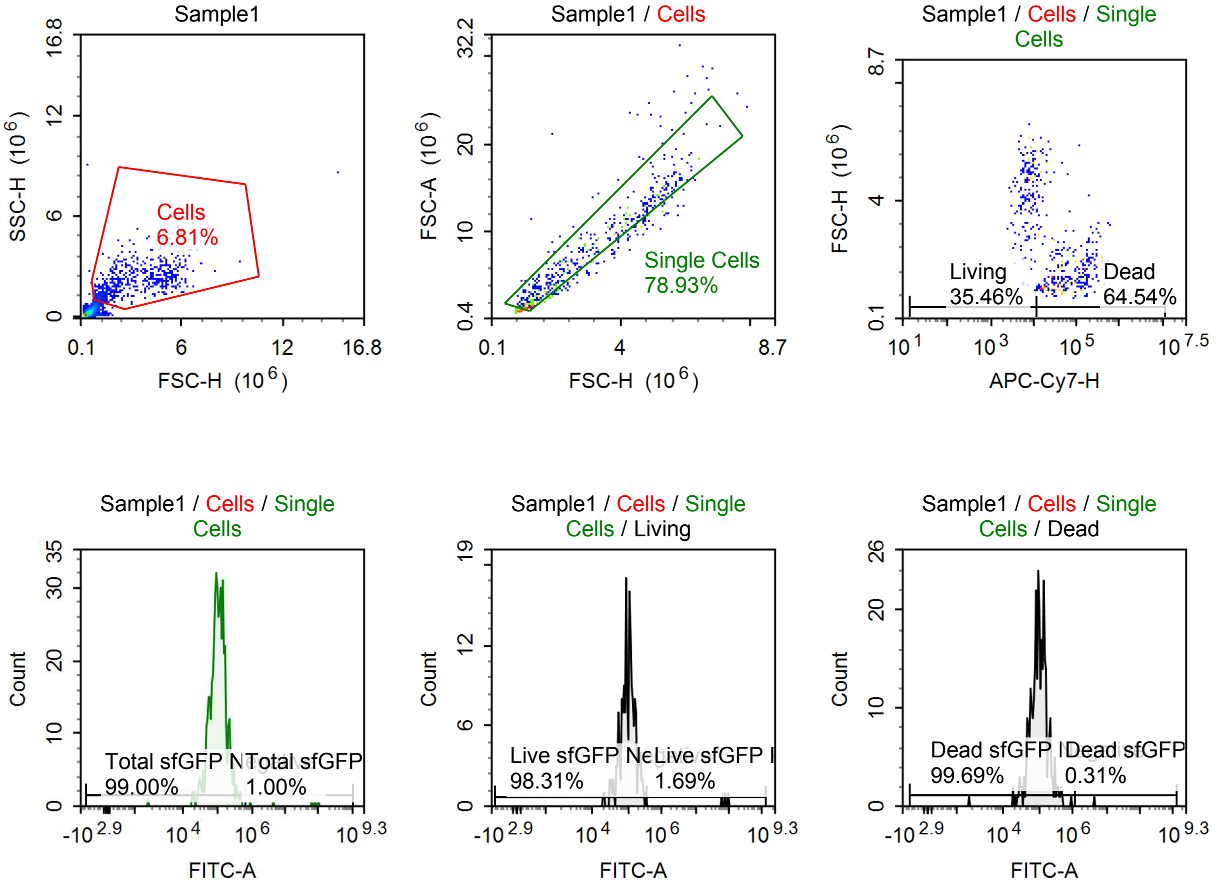

Sample Statistics of Sample1

| Gate | Count | % Parent | X | Y | Median X | Median Y |
|---|---|---|---|---|---|---|
| All | 9,337 | | | | | |
| Cells | 636 | 6.81 % | FSC-H | SSC-H | 2,236,105 | 1,936,213 |
| Single Cells | 502 | 78.93 % | FSC-H | FSC-A | 2,126,269 | 5,341,899 |
| Living | 178 | 35.46 % | APC-Cy7-H | FSC-H | 7,170 | 4,430,228 |
| Live sfGFP Negative | 175 | 98.31 % | FITC-A | Count | 104,515 | |
| Live sfGFP Positive | 3 | 1.69 % | FITC-A | Count | 75,428,488 | |
| Dead | 324 | 64.54 % | APC-Cy7-H | FSC-H | 50,724 | 1,486,946 |
| Dead sfGFP Negative | 323 | 99.69 % | FITC-A | Count | 100,815 | |
| Dead sfGFP Positive | 1 | 0.31 % | FITC-A | Count | 4,564,363 | |
| Total sfGFP Negative | 497 | 99.00 % | FITC-A | Count | 101,094 | |
| Total sfGFP Positive | 5 | 1.00 % | FITC-A | Count | 61,159,348 | |

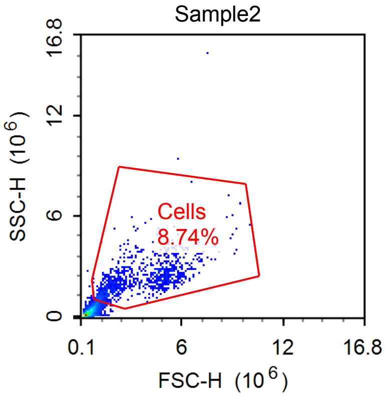


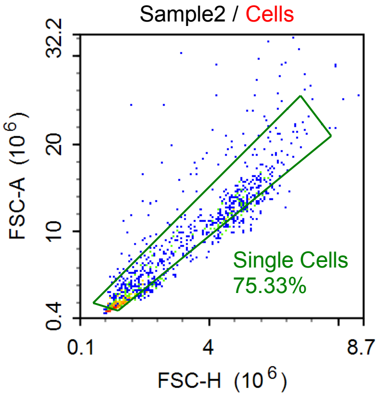


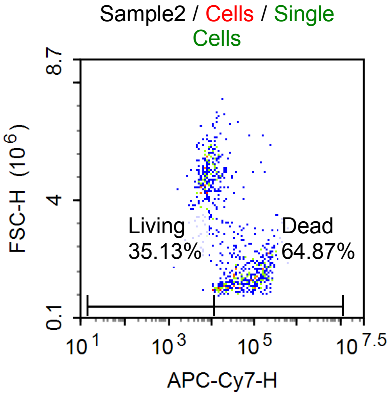


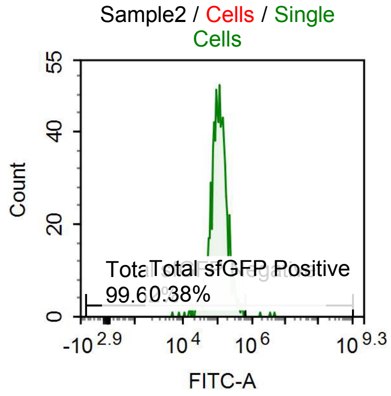


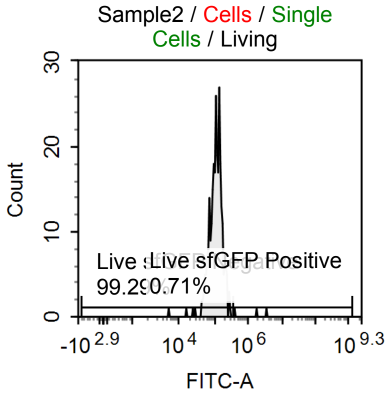


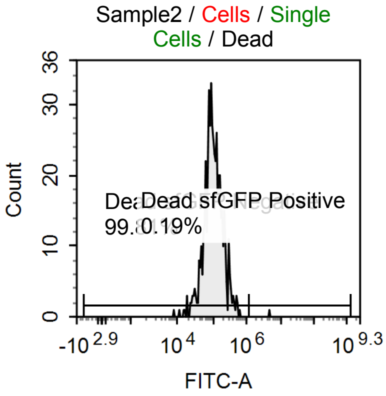

Sample Statistics of Sample2

| Gate | Count | % Parent | X | Y | Median X | Median Y |
|---|---|---|---|---|---|---|
| All | 12,106 | | | | | |
| Cells | 1,058 | 8.74 % | FSC-H | SSC-H | 2,534,202 | 2,056,785 |
| Single Cells | 797 | 75.33 % | FSC-H | FSC-A | 2,256,235 | 6,056,904 |
| Living | 280 | 35.13 % | APC-Cy7-H | FSC-H | 7,548 | 4,647,898 |
| Live sfGFP Negative | 278 | 99.29 % | FITC-A | Count | 113,671 | |
| Live sfGFP Positive | 2 | 0.71 % | FITC-A | Count | 2,593,014 | |
| Dead | 517 | 64.87 % | APC-Cy7-H | FSC-H | 51,840 | 1,560,267 |
| Dead sfGFP Negative | 516 | 99.81 % | FITC-A | Count | 100,027 | |
| Dead sfGFP Positive | 1 | 0.19 % | FITC-A | Count | 4,908,801 | |
| Total sfGFP Negative | 794 | 99.62 % | FITC-A | Count | 107,086 | |
| Total sfGFP Positive | 3 | 0.38 % | FITC-A | Count | 3,413,660 | |

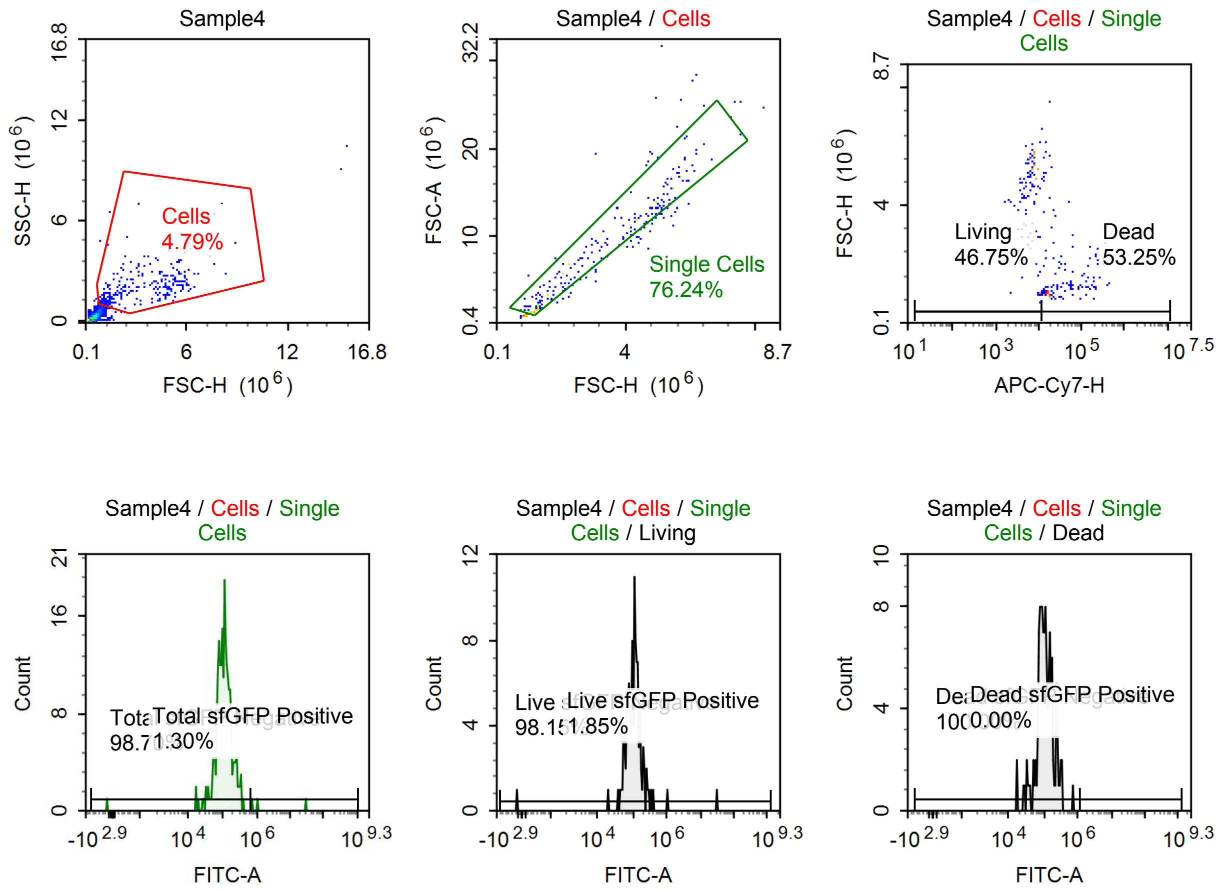

Sample Statistics of Sample4

| Gate | Count | % Parent | X | Y | Median X | Median Y |
|---|---|---|---|---|---|---|
| All | 6,329 | | | | | |
| Cells | 303 | 4.79 % | FSC-H | SSC-H | 2,509,835 | 1,794,281 |
| Single Cells | 231 | 76.24 % | FSC-H | FSC-A | 2,404,565 | 6,525,313 |
| Living | 108 | 46.75 % | APC-Cy7-H | FSC-H | 6,613 | 4,613,890 |
| Live sfGFP Negative | 106 | 98.15 % | FITC-A | Count | 110,222 | |
| Live sfGFP Positive | 2 | 1.85 % | FITC-A | Count | 15,633,802 | |
| Dead | 123 | 53.25 % | APC-Cy7-H | FSC-H | 30,410 | 1,333,017 |
| Dead sfGFP Negative | 123 | 100.00 % | FITC-A | Count | 106,839 | |
| Dead sfGFP Positive | 0 | 0.00 % | FITC-A | Count | 0 | |
| Total sfGFP Negative | 228 | 98.70 % | FITC-A | Count | 107,963 | |
| Total sfGFP Positive | 3 | 1.30 % | FITC-A | Count | 1,030,418 | |

#### p059 non poly-a tailed, 100% m1U

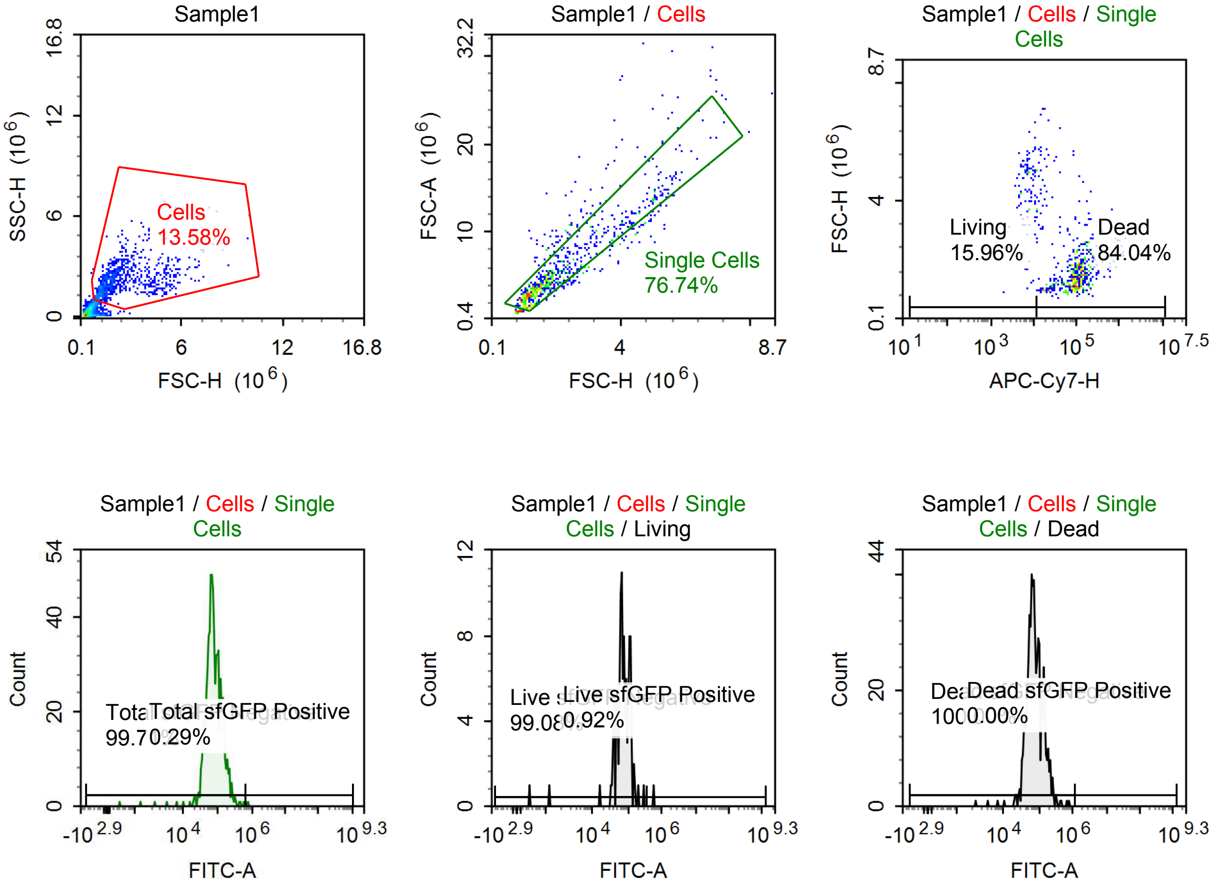

Sample Statistics of Sample1

| Gate | Count | % Parent | X | Y | Median X | Median Y |
|---|---|---|---|---|---|---|
| All | 6,553 | | | | | |
| Cells | 890 | 13.58 % | FSC-H | SSC-H | 1,782,704 | 2,021,040 |
| Single Cells | 683 | 76.74 % | FSC-H | FSC-A | 1,685,843 | 4,305,437 |
| Living | 109 | 15.96 % | APC-Cy7-H | FSC-H | 7,135 | 4,361,074 |
| Live sfGFP Negative | 108 | 99.08 % | FITC-A | Count | 72,141 | |
| Live sfGFP Positive | 1 | 0.92 % | FITC-A | Count | 565,557 | |
| Dead | 574 | 84.04 % | APC-Cy7-H | FSC-H | 96,034 | 1,500,948 |
| Dead sfGFP Negative | 574 | 100.00 % | FITC-A | Count | 73,082 | |
| Dead sfGFP Positive | 0 | 0.00 % | FITC-A | Count | 0 | |
| Total sfGFP Negative | 681 | 99.71 % | FITC-A | Count | 72,948 | |
| Total sfGFP Positive | 2 | 0.29 % | FITC-A | Count | 735,988 | |

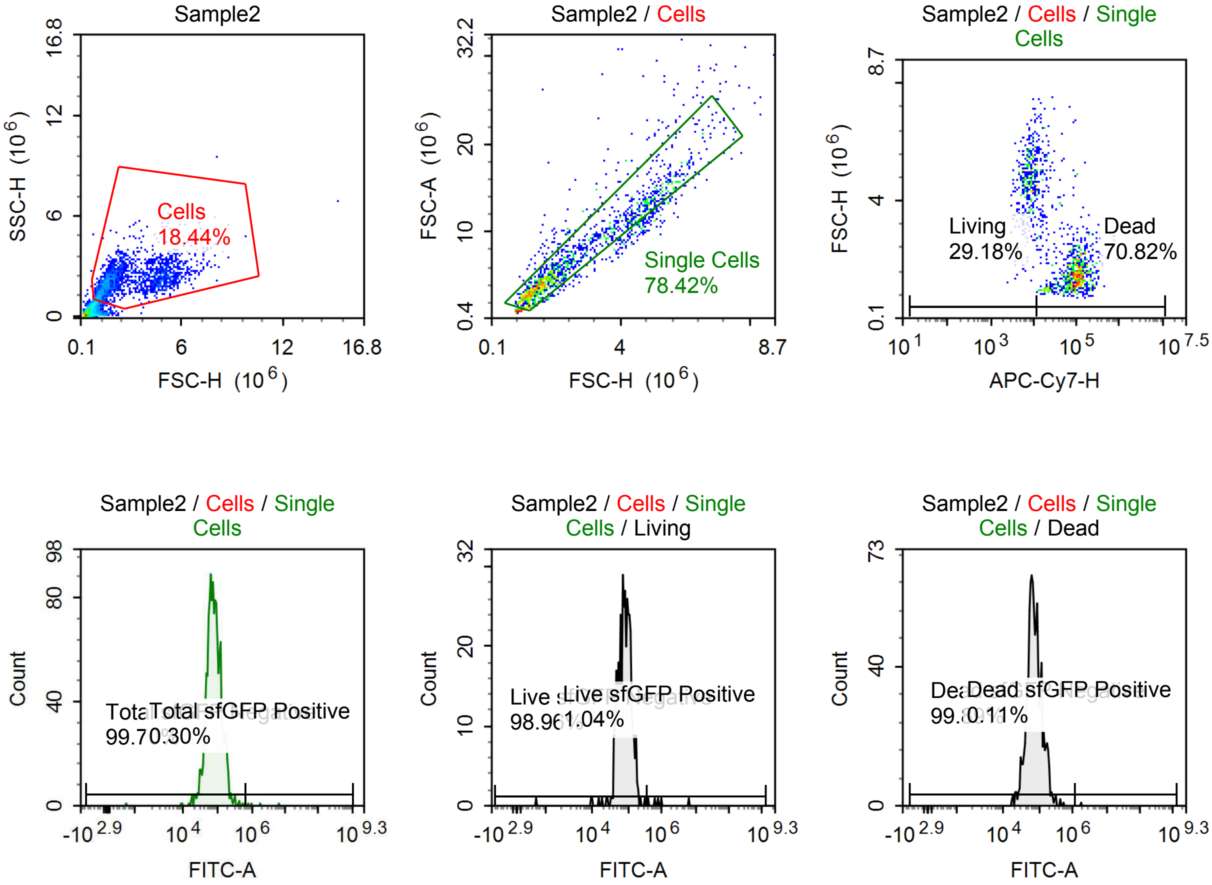

Sample Statistics of Sample2

| Gate | Count | % Parent | X | Y | Median X | Median Y |
|---|---|---|---|---|---|---|
| All | 9,150 | | | | | |
| Cells | 1,687 | 18.44 % | FSC-H | SSC-H | 2,134,404 | 2,147,726 |
| Single Cells | 1,323 | 78.42 % | FSC-H | FSC-A | 1,958,934 | 5,363,687 |
| Living | 386 | 29.18 % | APC-Cy7-H | FSC-H | 7,678 | 4,618,425 |
| Live sfGFP Negative | 382 | 98.96 % | FITC-A | Count | 81,585 | |
| Live sfGFP Positive | 4 | 1.04 % | FITC-A | Count | 883,240 | |
| Dead | 937 | 70.82 % | APC-Cy7-H | FSC-H | 87,664 | 1,561,024 |
| Dead sfGFP Negative | 936 | 99.89 % | FITC-A | Count | 71,661 | |
| Dead sfGFP Positive | 1 | 0.11 % | FITC-A | Count | 1,840,401 | |
| Total sfGFP Negative | 1,319 | 99.70 % | FITC-A | Count | 74,457 | |
| Total sfGFP Positive | 4 | 0.30 % | FITC-A | Count | 1,415,037 | |

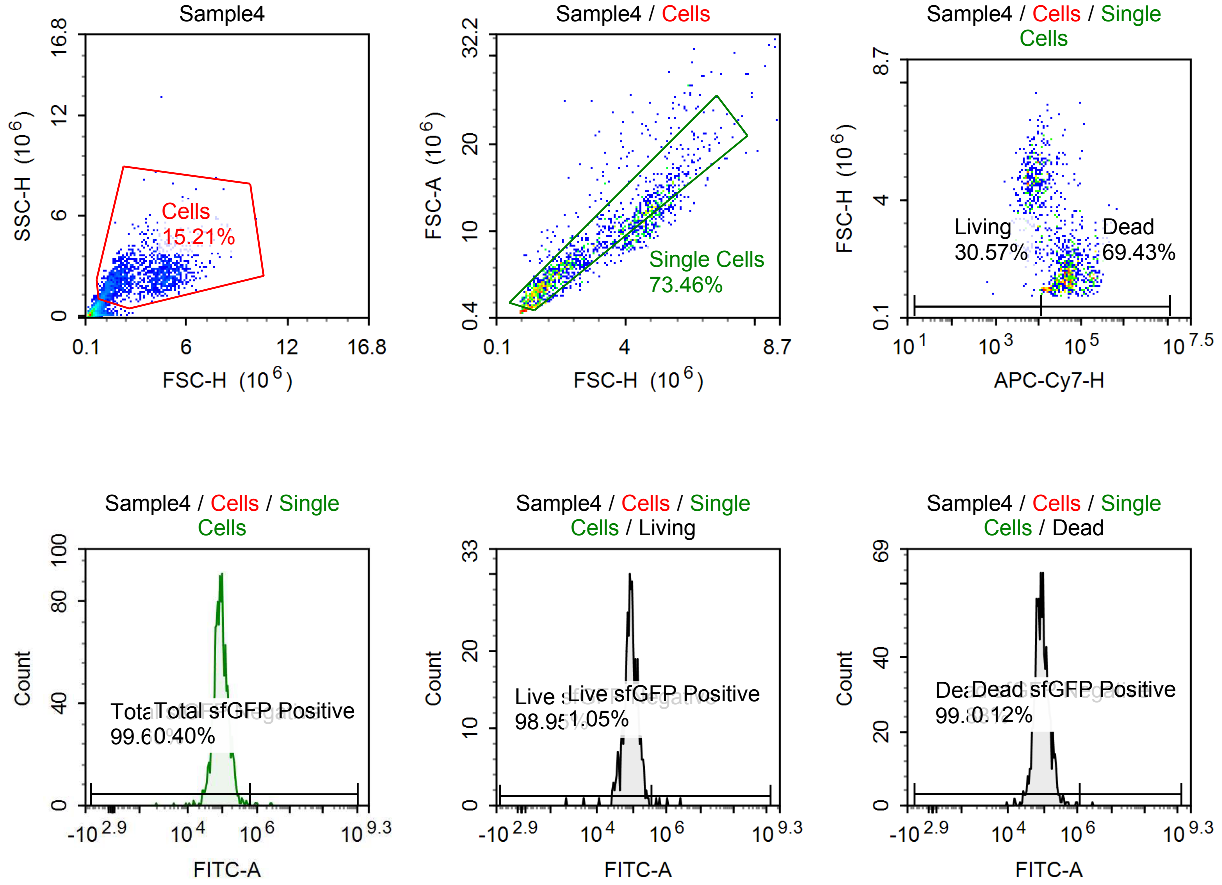

Sample Statistics of Sample4

| Gate | Count | % Parent | X | Y | Median X | Median Y |
|---|---|---|---|---|---|---|
| All | 11,124 | | | | | |
| Cells | 1,692 | 15.21 % | FSC-H | SSC-H | 2,405,466 | 2,153,868 |
| Single Cells | 1,243 | 73.46 % | FSC-H | FSC-A | 2,128,923 | 6,026,666 |
| Living | 380 | 30.57 % | APC-Cy7-H | FSC-H | 6,998 | 4,518,421 |
| Live sfGFP Negative | 376 | 98.95 % | FITC-A | Count | 88,671 | |
| Live sfGFP Positive | 4 | 1.05 % | FITC-A | Count | 850,020 | |
| Dead | 863 | 69.43 % | APC-Cy7-H | FSC-H | 45,186 | 1,601,562 |
| Dead sfGFP Negative | 862 | 99.88 % | FITC-A | Count | 86,608 | |
| Dead sfGFP Positive | 1 | 0.12 % | FITC-A | Count | 2,771,579 | |
| Total sfGFP Negative | 1,238 | 99.60 % | FITC-A | Count | 87,222 | |
| Total sfGFP Positive | 5 | 0.40 % | FITC-A | Count | 1,089,562 | |

#### p059, poly-a tailed, 33% m1U

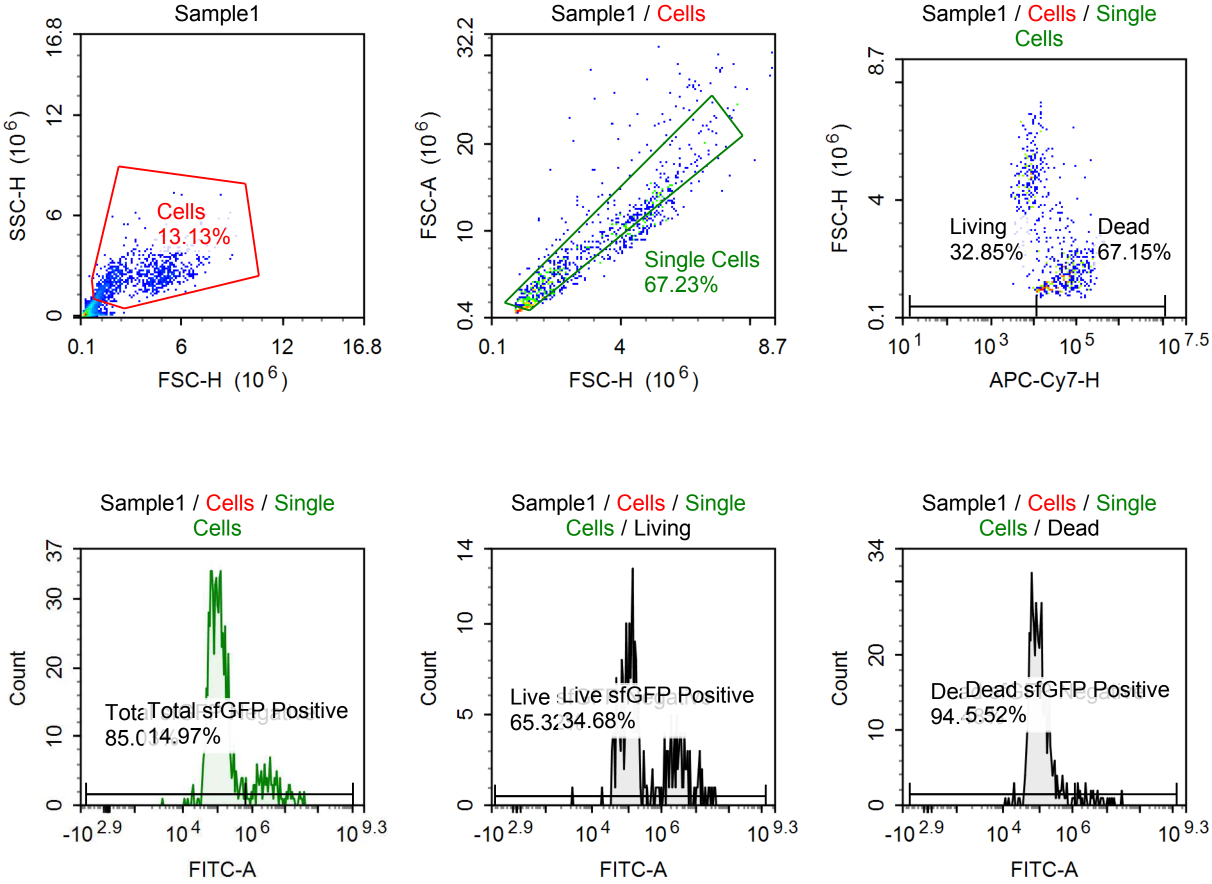

Sample Statistics of Sample1

| Gate | Count | % Parent | X | Y | Median X | Median Y |
|---|---|---|---|---|---|---|
| All | 8,552 | | | | | |
| Cells | 1,123 | 13.13 % | FSC-H | SSC-H | 2,244,346 | 2,098,226 |
| Single Cells | 755 | 67.23 % | FSC-H | FSC-A | 2,059,677 | 5,549,528 |
| Living | 248 | 32.85 % | APC-Cy7-H | FSC-H | 7,454 | 4,496,022 |
| Live sfGFP Negative | 162 | 65.32 % | FITC-A | Count | 108,861 | |
| Live sfGFP Positive | 86 | 34.68 % | FITC-A | Count | 3,134,354 | |
| Dead | 507 | 67.15 % | APC-Cy7-H | FSC-H | 46,674 | 1,564,910 |
| Dead sfGFP Negative | 479 | 94.48 % | FITC-A | Count | 92,401 | |
| Dead sfGFP Positive | 28 | 5.52 % | FITC-A | Count | 3,975,924 | |
| Total sfGFP Negative | 642 | 85.03 % | FITC-A | Count | 96,940 | |
| Total sfGFP Positive | 113 | 14.97 % | FITC-A | Count | 3,402,326 | |

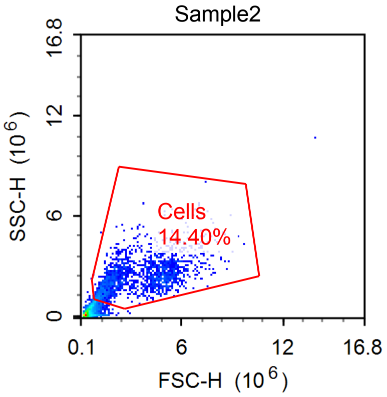


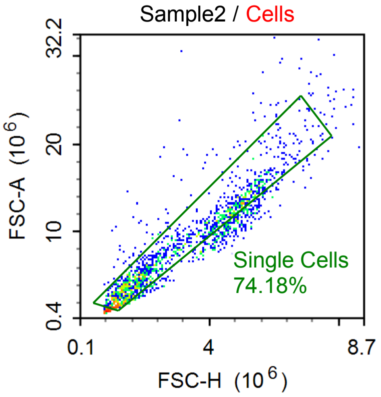


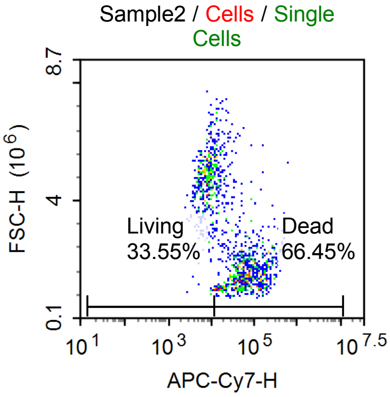


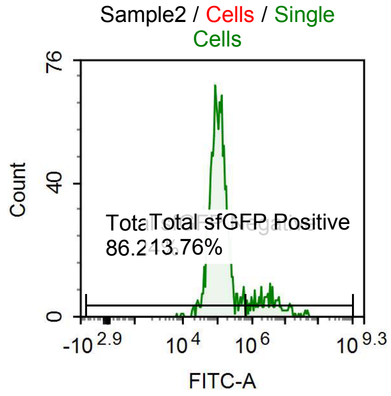


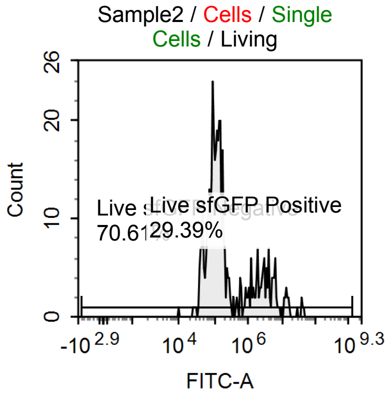


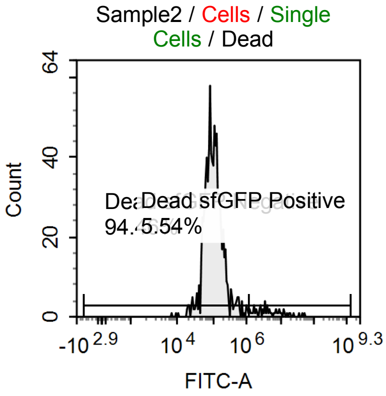

Sample Statistics of Sample2

| Gate | Count | % Parent | X | Y | Median X | Median Y |
|---|---|---|---|---|---|---|
| All | 12,722 | | | | | |
| Cells | 1,832 | 14.40 % | FSC-H | SSC-H | 2,606,954 | 2,207,510 |
| Single Cells | 1,359 | 74.18 % | FSC-H | FSC-A | 2,365,068 | 6,653,004 |
| Living | 456 | 33.55 % | APC-Cy7-H | FSC-H | 7,789 | 4,660,754 |
| Live sfGFP Negative | 322 | 70.61 % | FITC-A | Count | 106,003 | |
| Live sfGFP Positive | 134 | 29.39 % | FITC-A | Count | 2,885,991 | |
| Dead | 903 | 66.45 % | APC-Cy7-H | FSC-H | 51,276 | 1,724,960 |
| Dead sfGFP Negative | 853 | 94.46 % | FITC-A | Count | 101,096 | |
| Dead sfGFP Positive | 50 | 5.54 % | FITC-A | Count | 4,193,904 | |
| Total sfGFP Negative | 1,172 | 86.24 % | FITC-A | Count | 101,985 | |
| Total sfGFP Positive | 187 | 13.76 % | FITC-A | Count | 3,269,261 | |

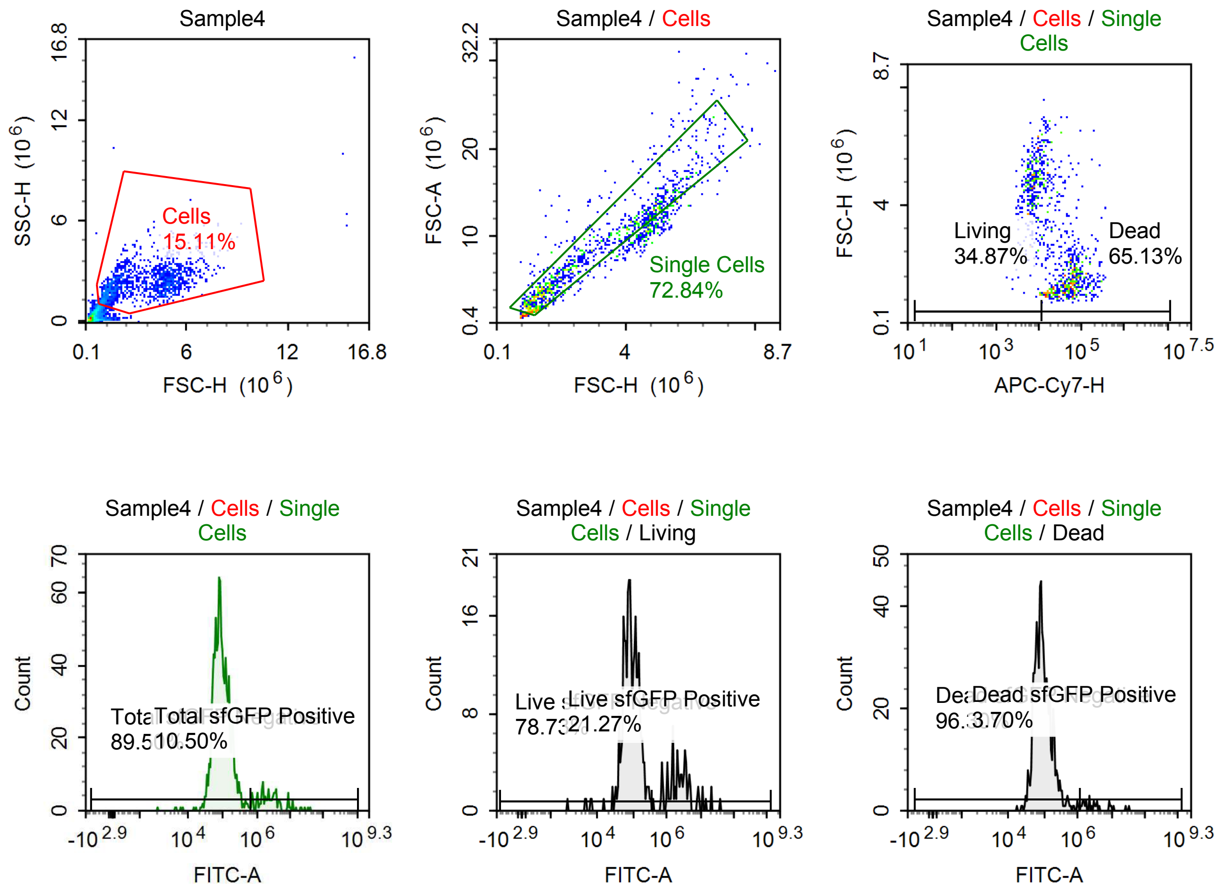

Sample Statistics of Sample4

| Gate | Count | % Parent | X | Y | Median X | Median Y |
|---|---|---|---|---|---|---|
| All | 9,430 | | | | | |
| Cells | 1,425 | 15.11 % | FSC-H | SSC-H | 2,703,226 | 2,109,696 |
| Single Cells | 1,038 | 72.84 % | FSC-H | FSC-A | 2,369,760 | 6,770,229 |
| Living | 362 | 34.87 % | APC-Cy7-H | FSC-H | 7,302 | 4,681,969 |
| Live sfGFP Negative | 285 | 78.73 % | FITC-A | Count | 85,316 | |
| Live sfGFP Positive | 77 | 21.27 % | FITC-A | Count | 2,431,541 | |
| Dead | 676 | 65.13 % | APC-Cy7-H | FSC-H | 45,156 | 1,602,141 |
| Dead sfGFP Negative | 651 | 96.30 % | FITC-A | Count | 85,996 | |
| Dead sfGFP Positive | 25 | 3.70 % | FITC-A | Count | 3,139,052 | |
| Total sfGFP Negative | 929 | 89.50 % | FITC-A | Count | 85,316 | |
| Total sfGFP Positive | 109 | 10.50 % | FITC-A | Count | 2,371,426 | |

#### p059 non poly-a tailed, 100% UTP

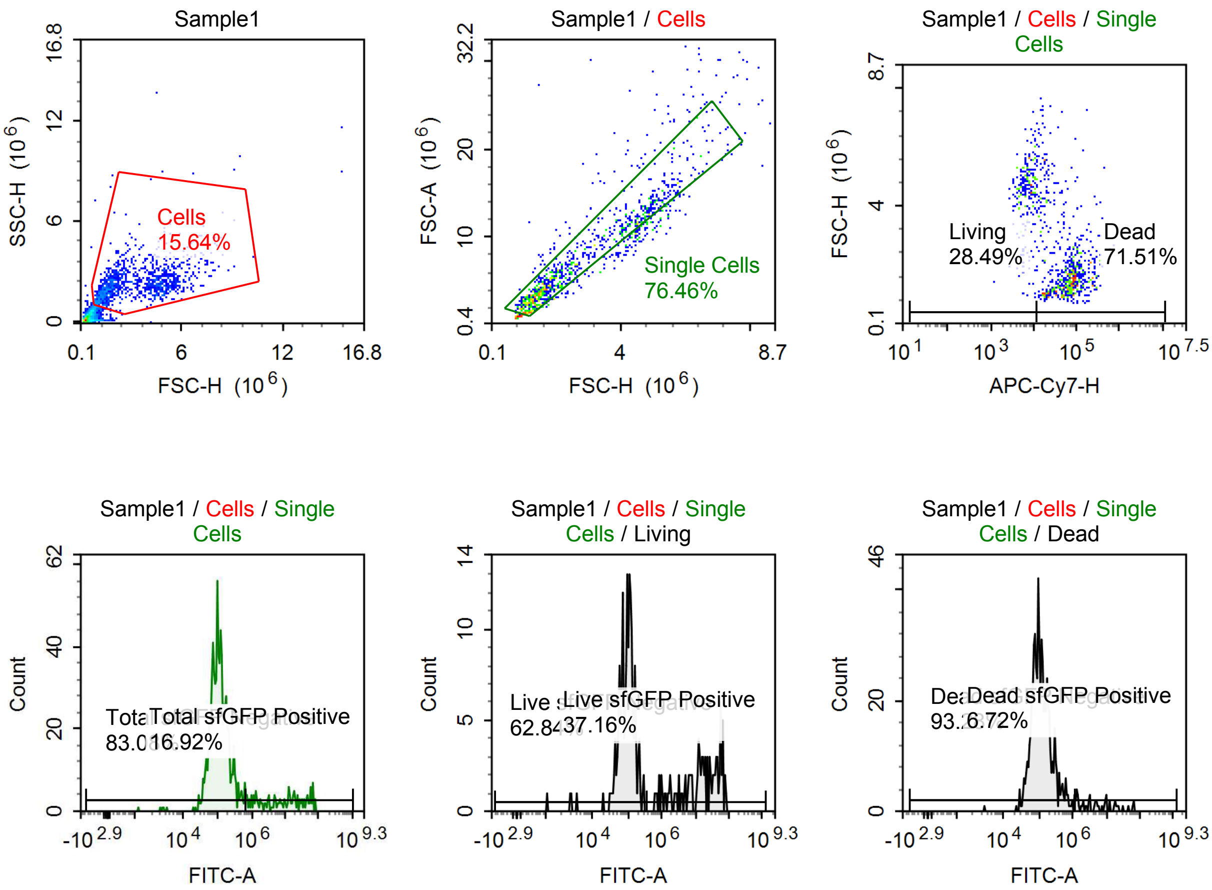

Sample Statistics of Sample1

| Gate | Count | % Parent | X | Y | Median X | Median Y |
|---|---|---|---|---|---|---|
| All | 7,662 | | | | | |
| Cells | 1,198 | 15.64 % | FSC-H | SSC-H | 2,265,921 | 2,096,899 |
| Single Cells | 916 | 76.46 % | FSC-H | FSC-A | 2,060,036 | 5,558,493 |
| Living | 261 | 28.49 % | APC-Cy7-H | FSC-H | 7,099 | 4,631,056 |
| Live sfGFP Negative | 164 | 62.84 % | FITC-A | Count | 96,482 | |
| Live sfGFP Positive | 97 | 37.16 % | FITC-A | Count | 16,361,052 | |
| Dead | 655 | 71.51 % | APC-Cy7-H | FSC-H | 65,088 | 1,623,751 |
| Dead sfGFP Negative | 611 | 93.28 % | FITC-A | Count | 110,129 | |
| Dead sfGFP Positive | 44 | 6.72 % | FITC-A | Count | 5,312,497 | |
| Total sfGFP Negative | 761 | 83.08 % | FITC-A | Count | 105,029 | |
| Total sfGFP Positive | 155 | 16.92 % | FITC-A | Count | 9,845,002 | |

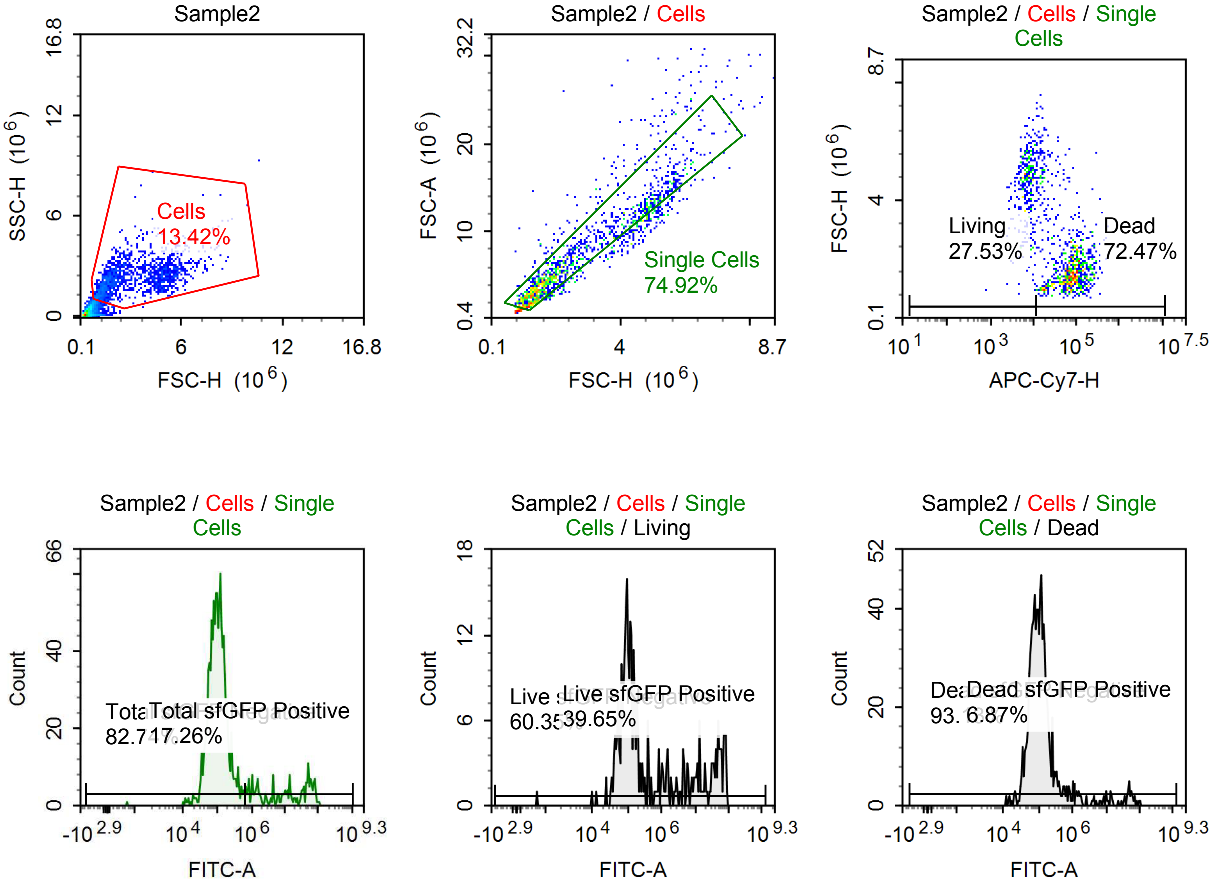

Sample Statistics of Sample2

| Gate | Count | % Parent | X | Y | Median X | Median Y |
|---|---|---|---|---|---|---|
| All | 12,390 | | | | | |
| Cells | 1,663 | 13.42 % | FSC-H | SSC-H | 2,182,498 | 2,170,522 |
| Single Cells | 1,246 | 74.92 % | FSC-H | FSC-A | 1,979,552 | 5,263,523 |
| Living | 343 | 27.53 % | APC-Cy7-H | FSC-H | 7,248 | 4,679,436 |
| Live sfGFP Negative | 207 | 60.35 % | FITC-A | Count | 99,766 | |
| Live sfGFP Positive | 136 | 39.65 % | FITC-A | Count | 13,522,322 | |
| Dead | 903 | 72.47 % | APC-Cy7-H | FSC-H | 74,575 | 1,614,339 |
| Dead sfGFP Negative | 841 | 93.13 % | FITC-A | Count | 100,138 | |
| Dead sfGFP Positive | 62 | 6.87 % | FITC-A | Count | 14,478,820 | |
| Total sfGFP Negative | 1,031 | 82.74 % | FITC-A | Count | 98,598 | |
| Total sfGFP Positive | 215 | 17.26 % | FITC-A | Count | 11,425,560 | |

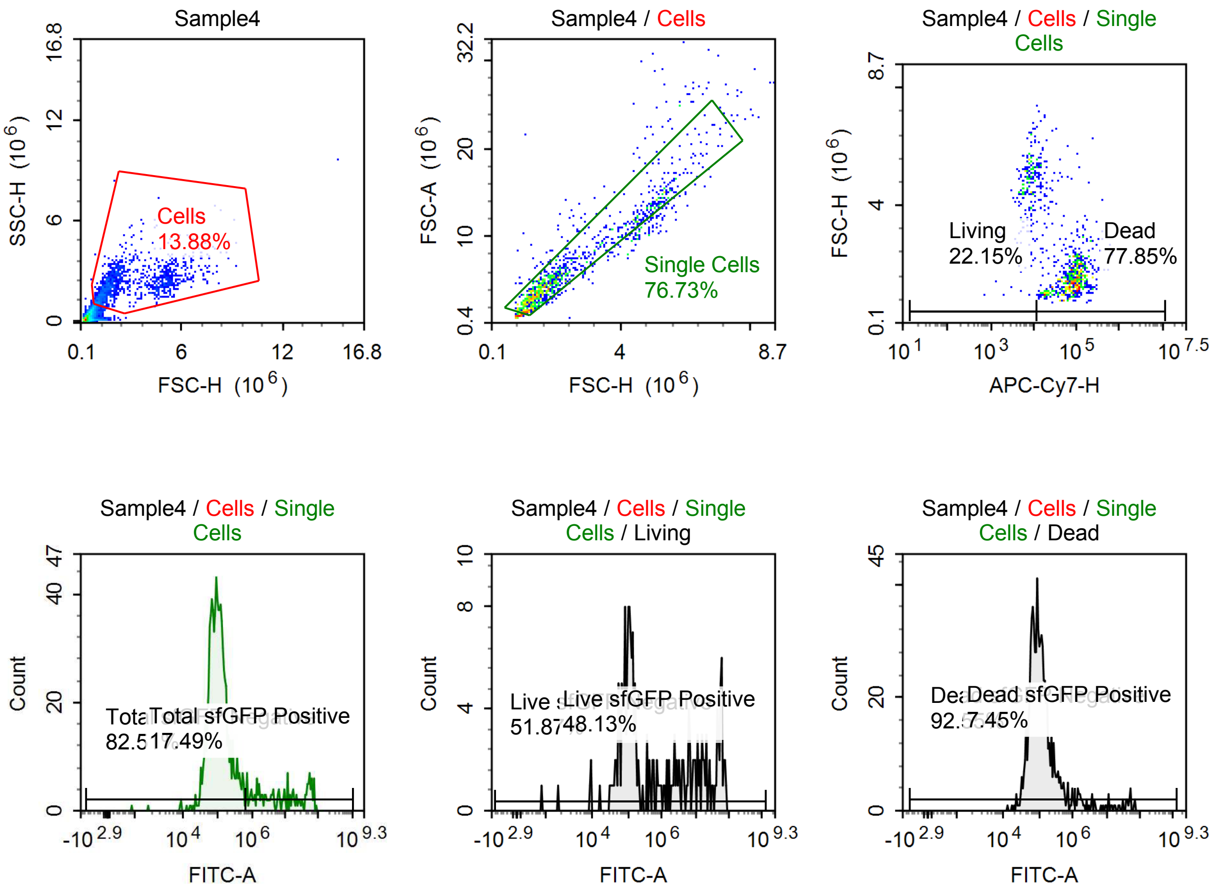

Sample Statistics of Sample4

| Gate | Count | % Parent | X | Y | Median X | Median Y |
|---|---|---|---|---|---|---|
| All | 9,068 | | | | | |
| Cells | 1,259 | 13.88 % | FSC-H | SSC-H | 1,860,147 | 2,108,464 |
| Single Cells | 966 | 76.73 % | FSC-H | FSC-A | 1,780,323 | 4,798,524 |
| Living | 214 | 22.15 % | APC-Cy7-H | FSC-H | 7,637 | 4,799,393 |
| Live sfGFP Negative | 111 | 51.87 % | FITC-A | Count | 100,205 | |
| Live sfGFP Positive | 103 | 48.13 % | FITC-A | Count | 10,099,129 | |
| Dead | 752 | 77.85 % | APC-Cy7-H | FSC-H | 79,920 | 1,528,222 |
| Dead sfGFP Negative | 696 | 92.55 % | FITC-A | Count | 97,315 | |
| Dead sfGFP Positive | 56 | 7.45 % | FITC-A | Count | 7,312,273 | |
| Total sfGFP Negative | 797 | 82.51 % | FITC-A | Count | 96,855 | |
| Total sfGFP Positive | 169 | 17.49 % | FITC-A | Count | 7,905,631 | |

#### p059 non poly-a tailed, 33% m1U

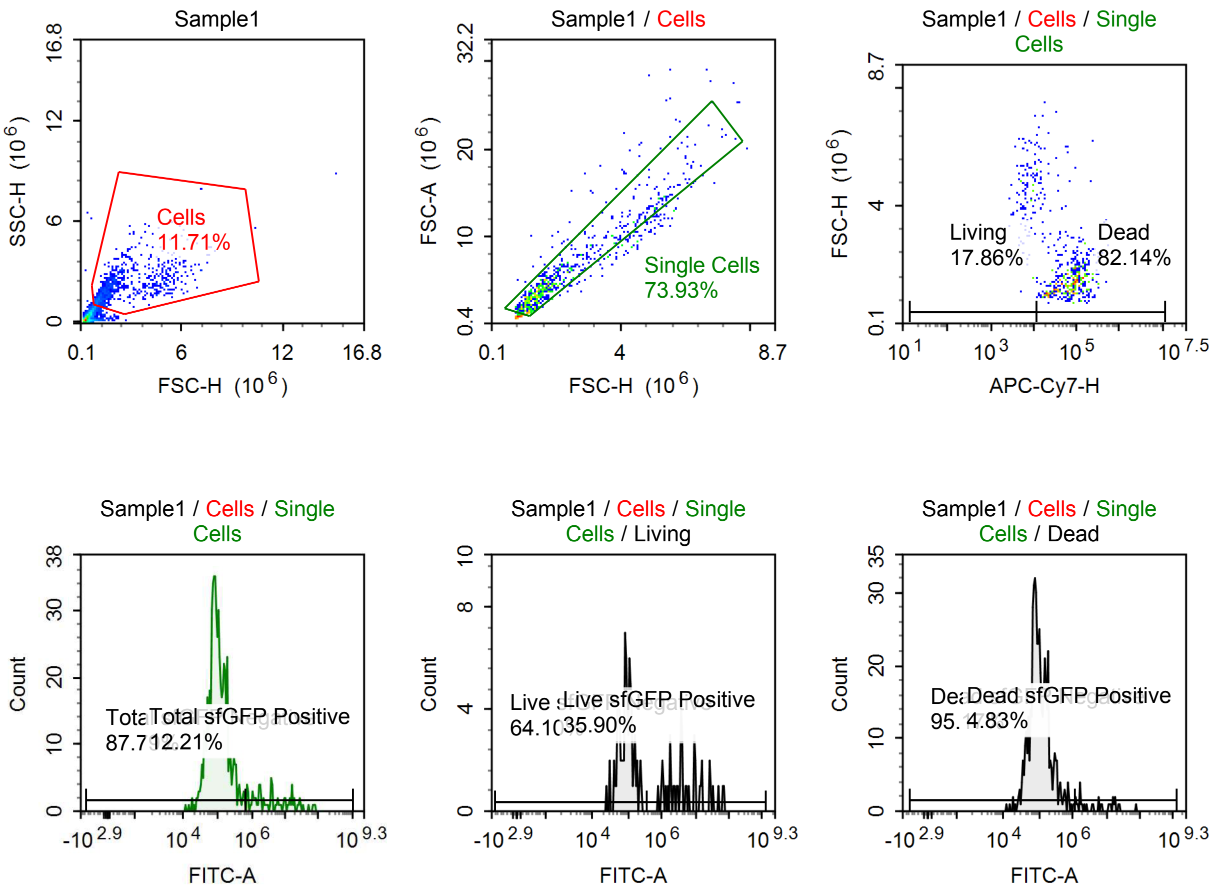

Sample Statistics of Sample1

| Gate | Count | % Parent | X | Y | Median X | Median Y |
|---|---|---|---|---|---|---|
| All | 7,566 | | | | | |
| Cells | 886 | 11.71 % | FSC-H | SSC-H | 1,642,376 | 1,897,702 |
| Single Cells | 655 | 73.93 % | FSC-H | FSC-A | 1,643,042 | 4,478,488 |
| Living | 117 | 17.86 % | APC-Cy7-H | FSC-H | 7,007 | 4,434,529 |
| Live sfGFP Negative | 75 | 64.10 % | FITC-A | Count | 88,946 | |
| Live sfGFP Positive | 42 | 35.90 % | FITC-A | Count | 4,114,328 | |
| Dead | 538 | 82.14 % | APC-Cy7-H | FSC-H | 73,353 | 1,485,464 |
| Dead sfGFP Negative | 512 | 95.17 % | FITC-A | Count | 95,839 | |
| Dead sfGFP Positive | 26 | 4.83 % | FITC-A | Count | 6,565,768 | |
| Total sfGFP Negative | 575 | 87.79 % | FITC-A | Count | 93,172 | |
| Total sfGFP Positive | 80 | 12.21 % | FITC-A | Count | 3,813,737 | |

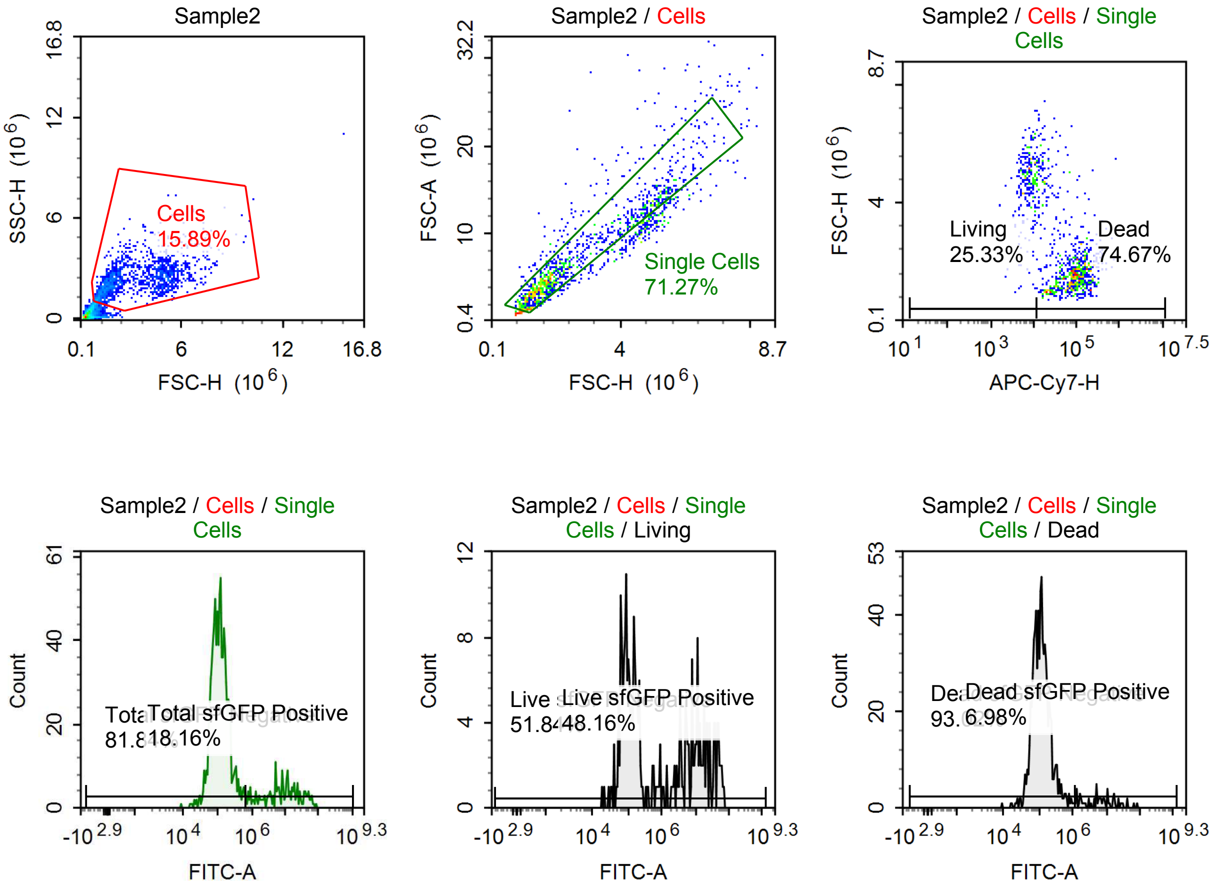

Sample Statistics of Sample2

| Gate | Count | % Parent | X | Y | Median X | Median Y |
|---|---|---|---|---|---|---|
| All | 9,485 | | | | | |
| Cells | 1,507 | 15.89 % | FSC-H | SSC-H | 2,229,384 | 2,240,419 |
| Single Cells | 1,074 | 71.27 % | FSC-H | FSC-A | 1,915,316 | 5,303,774 |
| Living | 272 | 25.33 % | APC-Cy7-H | FSC-H | 7,704 | 4,784,051 |
| Live sfGFP Negative | 141 | 51.84 % | FITC-A | Count | 100,337 | |
| Live sfGFP Positive | 131 | 48.16 % | FITC-A | Count | 10,832,287 | |
| Dead | 802 | 74.67 % | APC-Cy7-H | FSC-H | 81,650 | 1,638,312 |
| Dead sfGFP Negative | 746 | 93.02 % | FITC-A | Count | 114,103 | |
| Dead sfGFP Positive | 56 | 6.98 % | FITC-A | Count | 6,970,439 | |
| Total sfGFP Negative | 879 | 81.84 % | FITC-A | Count | 111,861 | |
| Total sfGFP Positive | 195 | 18.16 % | FITC-A | Count | 8,432,353 | |

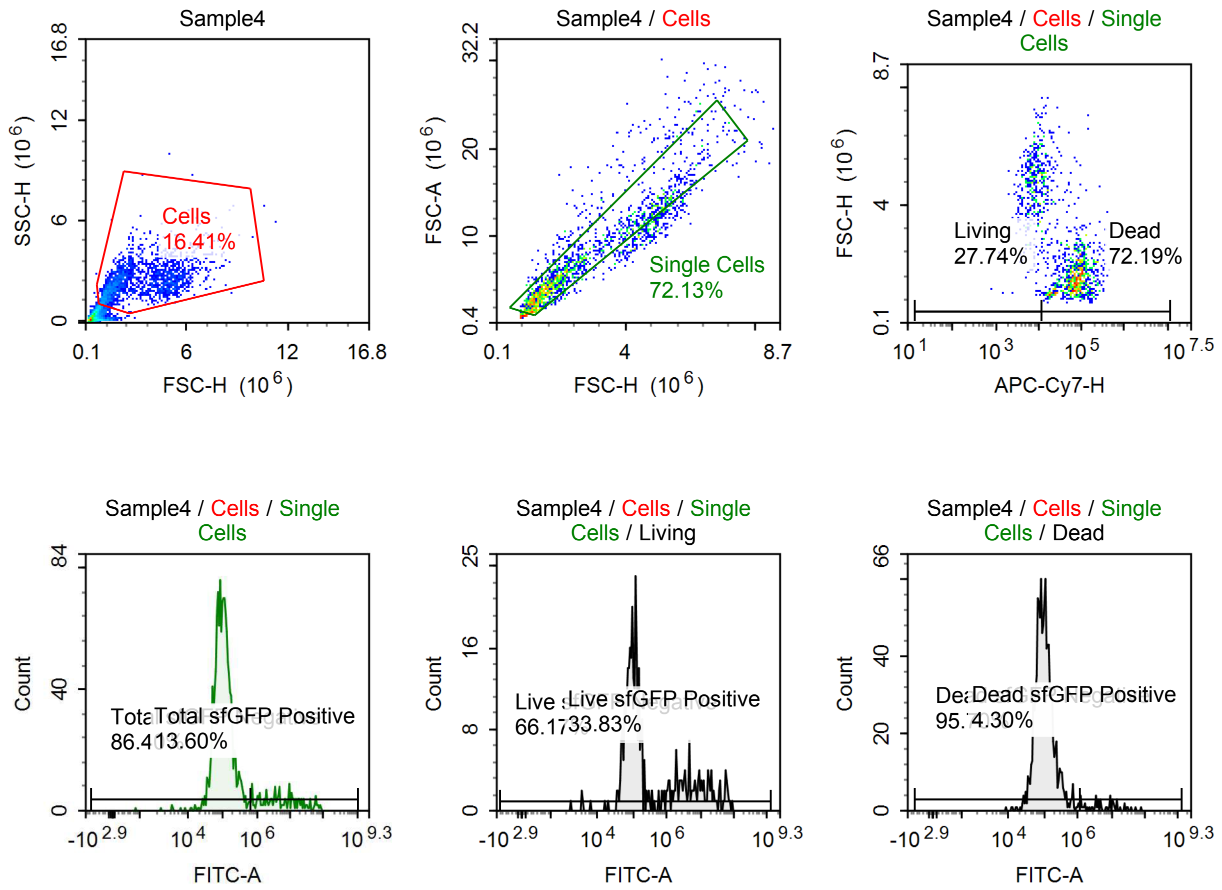

Sample Statistics of Sample4

| Gate | Count | % Parent | X | Y | Median X | Median Y |
|---|---|---|---|---|---|---|
| All | 12,243 | | | | | |
| Cells | 2,009 | 16.41 % | FSC-H | SSC-H | 2,243,192 | 2,213,668 |
| Single Cells | 1,449 | 72.13 % | FSC-H | FSC-A | 1,993,315 | 5,551,445 |
| Living | 402 | 27.74 % | APC-Cy7-H | FSC-H | 7,292 | 4,692,459 |
| Live sfGFP Negative | 266 | 66.17 % | FITC-A | Count | 95,616 | |
| Live sfGFP Positive | 136 | 33.83 % | FITC-A | Count | 5,806,482 | |
| Dead | 1,046 | 72.19 % | APC-Cy7-H | FSC-H | 68,229 | 1,619,636 |
| Dead sfGFP Negative | 1,001 | 95.70 % | FITC-A | Count | 99,402 | |
| Dead sfGFP Positive | 45 | 4.30 % | FITC-A | Count | 7,581,222 | |
| Total sfGFP Negative | 1,252 | 86.40 % | FITC-A | Count | 97,943 | |
| Total sfGFP Positive | 197 | 13.60 % | FITC-A | Count | 4,909,537 | |

#### p005 + 10 uM Tet

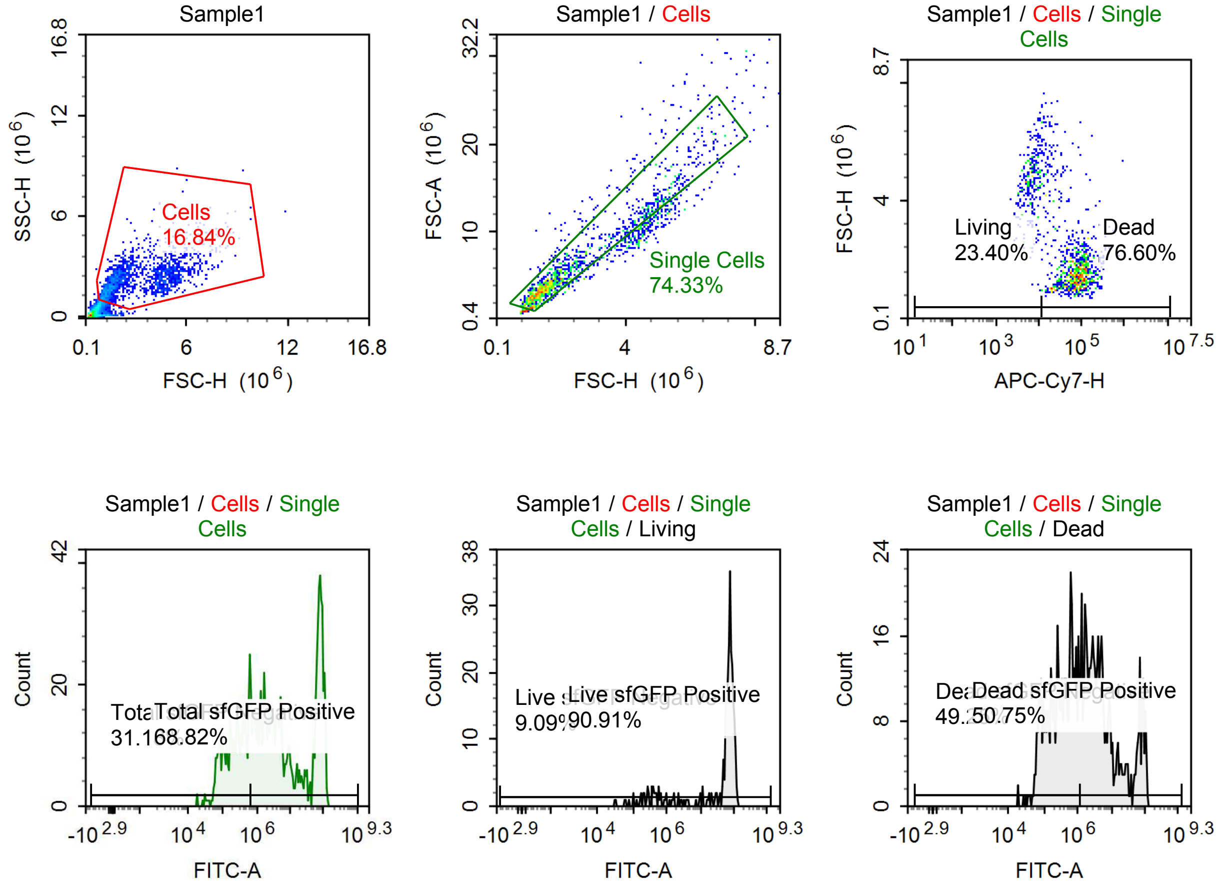

Sample Statistics of Sample1

| Gate | Count | % Parent | X | Y | Median X | Median Y |
|---|---|---|---|---|---|---|
| All | 9,760 | | | | | |
| Cells | 1,644 | 16.84 % | FSC-H | SSC-H | 2,145,470 | 2,153,647 |
| Single Cells | 1,222 | 74.33 % | FSC-H | FSC-A | 1,859,259 | 4,749,352 |
| Living | 286 | 23.40 % | APC-Cy7-H | FSC-H | 6,819 | 4,663,245 |
| Live sfGFP Negative | 26 | 9.09 % | FITC-A | Count | 206,877 | |
| Live sfGFP Positive | 260 | 90.91 % | FITC-A | Count | 71,820,720 | |
| Dead | 936 | 76.60 % | APC-Cy7-H | FSC-H | 71,538 | 1,559,634 |
| Dead sfGFP Negative | 461 | 49.25 % | FITC-A | Count | 340,488 | |
| Dead sfGFP Positive | 475 | 50.75 % | FITC-A | Count | 4,981,603 | |
| Total sfGFP Negative | 381 | 31.18 % | FITC-A | Count | 241,935 | |
| Total sfGFP Positive | 841 | 68.82 % | FITC-A | Count | 6,960,781 | |

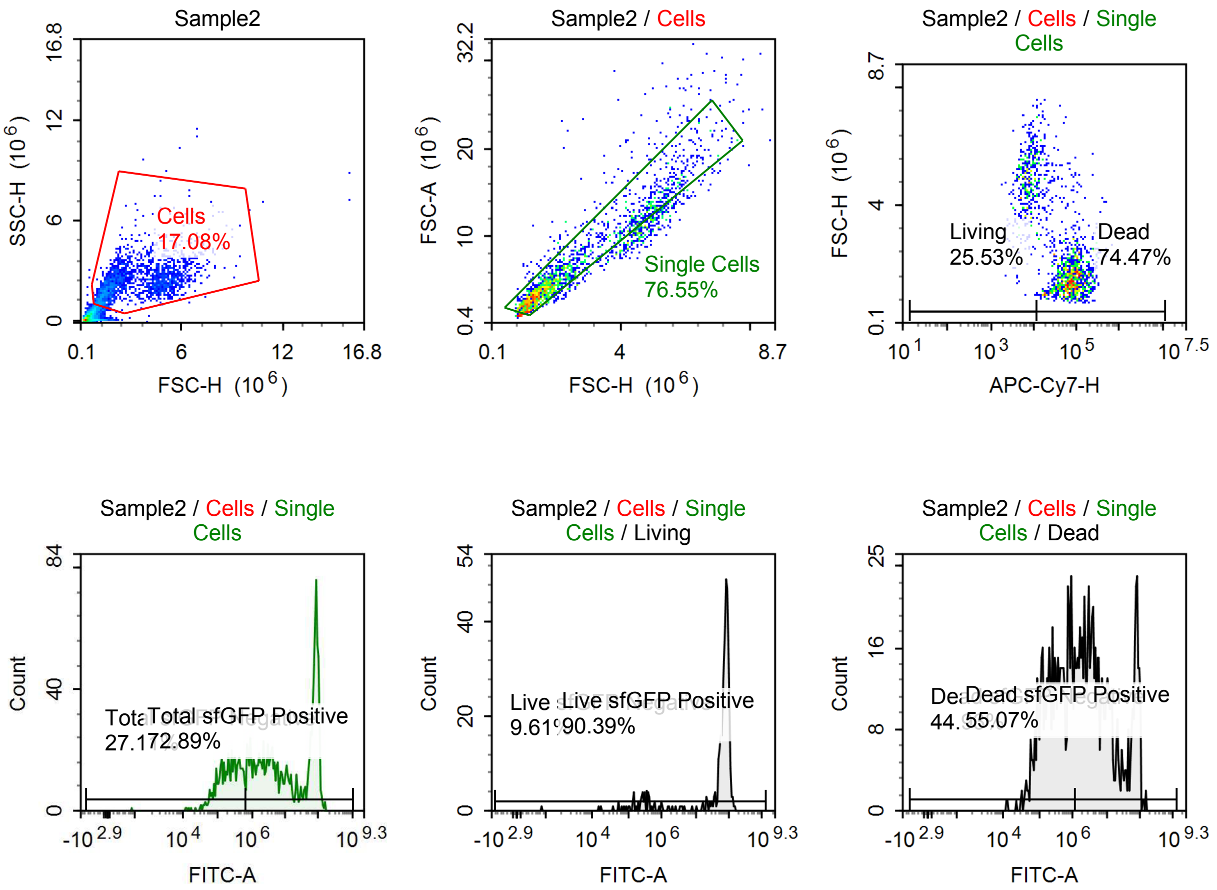

Sample Statistics of Sample2

| Gate | Count | % Parent | X | Y | Median X | Median Y |
|---|---|---|---|---|---|---|
| All | 12,161 | | | | | |
| Cells | 2,077 | 17.08 % | FSC-H | SSC-H | 2,374,543 | 2,212,683 |
| Single Cells | 1,590 | 76.55 % | FSC-H | FSC-A | 1,979,224 | 5,258,242 |
| Living | 406 | 25.53 % | APC-Cy7-H | FSC-H | 6,902 | 4,775,909 |
| Live sfGFP Negative | 39 | 9.61 % | FITC-A | Count | 199,994 | |
| Live sfGFP Positive | 367 | 90.39 % | FITC-A | Count | 78,110,448 | |
| Dead | 1,184 | 74.47 % | APC-Cy7-H | FSC-H | 69,664 | 1,640,438 |
| Dead sfGFP Negative | 532 | 44.93 % | FITC-A | Count | 327,516 | |
| Dead sfGFP Positive | 652 | 55.07 % | FITC-A | Count | 6,866,402 | |
| Total sfGFP Negative | 431 | 27.11 % | FITC-A | Count | 214,267 | |
| Total sfGFP Positive | 1,159 | 72.89 % | FITC-A | Count | 12,497,676 | |

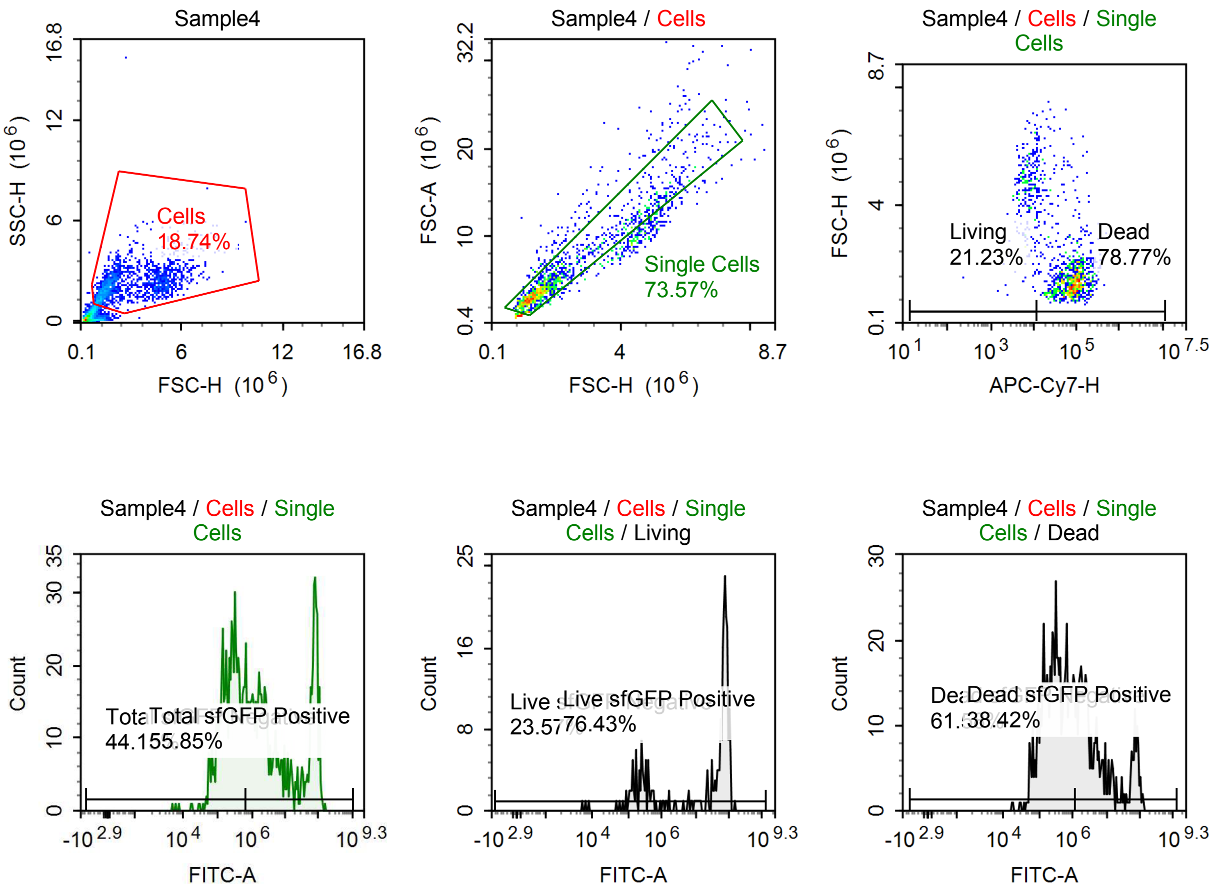

Sample Statistics of Sample4

| Gate | Count | % Parent | X | Y | Median X | Median Y |
|---|---|---|---|---|---|---|
| All | 8,988 | | | | | |
| Cells | 1,684 | 18.74 % | FSC-H | SSC-H | 1,919,254 | 2,094,932 |
| Single Cells | 1,239 | 73.57 % | FSC-H | FSC-A | 1,677,328 | 4,345,631 |
| Living | 263 | 21.23 % | APC-Cy7-H | FSC-H | 7,277 | 4,618,992 |
| Live sfGFP Negative | 62 | 23.57 % | FITC-A | Count | 220,203 | |
| Live sfGFP Positive | 201 | 76.43 % | FITC-A | Count | 70,105,536 | |
| Dead | 976 | 78.77 % | APC-Cy7-H | FSC-H | 79,780 | 1,492,799 |
| Dead sfGFP Negative | 601 | 61.58 % | FITC-A | Count | 310,008 | |
| Dead sfGFP Positive | 375 | 38.42 % | FITC-A | Count | 4,370,776 | |
| Total sfGFP Negative | 547 | 44.15 % | FITC-A | Count | 242,883 | |
| Total sfGFP Positive | 692 | 55.85 % | FITC-A | Count | 5,337,415 | |

#### p005 + 100 uM Tet

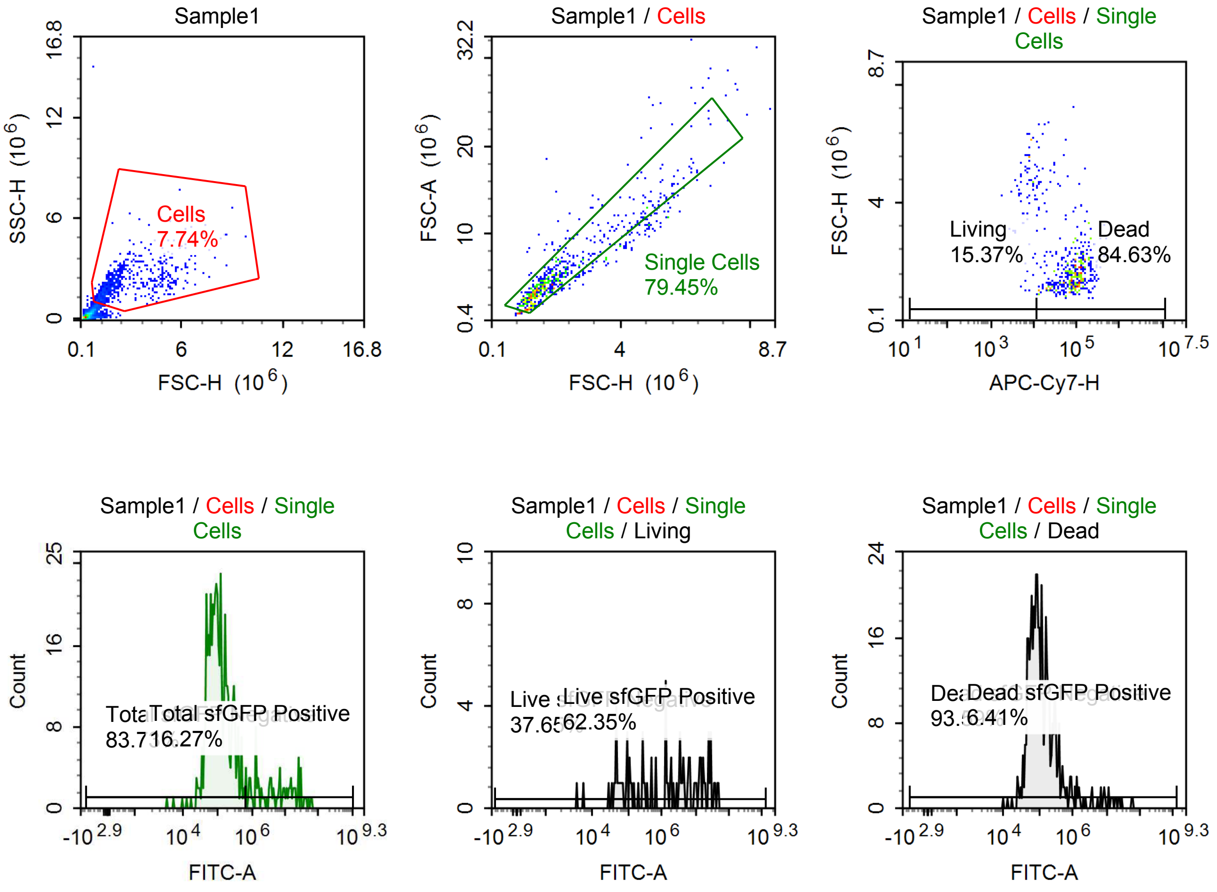

Sample Statistics of Sample1

| Gate | Count | % Parent | X | Y | Median X | Median Y |
|---|---|---|---|---|---|---|
| All | 8,996 | | | | | |
| Cells | 696 | 7.74 % | FSC-H | SSC-H | 1,861,795 | 1,978,867 |
| Single Cells | 553 | 79.45 % | FSC-H | FSC-A | 1,672,574 | 4,379,531 |
| Living | 85 | 15.37 % | APC-Cy7-H | FSC-H | 7,359 | 4,483,752 |
| Live sfGFP Negative | 32 | 37.65 % | FITC-A | Count | 80,354 | |
| Live sfGFP Positive | 53 | 62.35 % | FITC-A | Count | 6,311,553 | |
| Dead | 468 | 84.63 % | APC-Cy7-H | FSC-H | 83,619 | 1,571,914 |
| Dead sfGFP Negative | 438 | 93.59 % | FITC-A | Count | 97,950 | |
| Dead sfGFP Positive | 30 | 6.41 % | FITC-A | Count | 8,449,386 | |
| Total sfGFP Negative | 463 | 83.73 % | FITC-A | Count | 95,119 | |
| Total sfGFP Positive | 90 | 16.27 % | FITC-A | Count | 6,233,691 | |

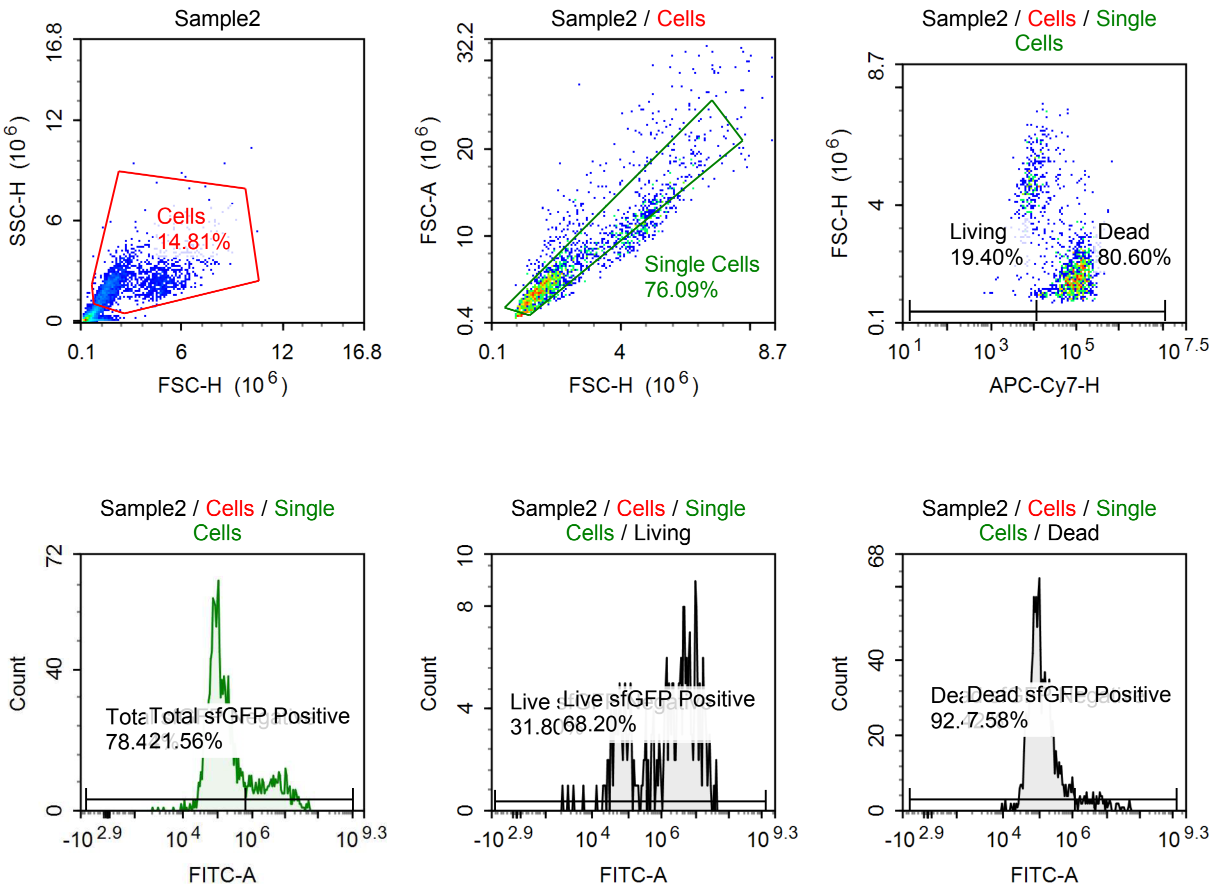

Sample Statistics of Sample2

| Gate | Count | % Parent | X | Y | Median X | Median Y |
|---|---|---|---|---|---|---|
| All | 13,949 | | | | | |
| Cells | 2,066 | 14.81 % | FSC-H | SSC-H | 2,058,836 | 2,190,080 |
| Single Cells | 1,572 | 76.09 % | FSC-H | FSC-A | 1,841,818 | 5,042,429 |
| Living | 305 | 19.40 % | APC-Cy7-H | FSC-H | 7,478 | 4,459,851 |
| Live sfGFP Negative | 97 | 31.80 % | FITC-A | Count | 68,067 | |
| Live sfGFP Positive | 208 | 68.20 % | FITC-A | Count | 4,766,847 | |
| Dead | 1,267 | 80.60 % | APC-Cy7-H | FSC-H | 89,386 | 1,670,479 |
| Dead sfGFP Negative | 1,171 | 92.42 % | FITC-A | Count | 102,823 | |
| Dead sfGFP Positive | 96 | 7.58 % | FITC-A | Count | 5,492,284 | |
| Total sfGFP Negative | 1,233 | 78.44 % | FITC-A | Count | 99,248 | |
| Total sfGFP Positive | 339 | 21.56 % | FITC-A | Count | 4,242,255 | |

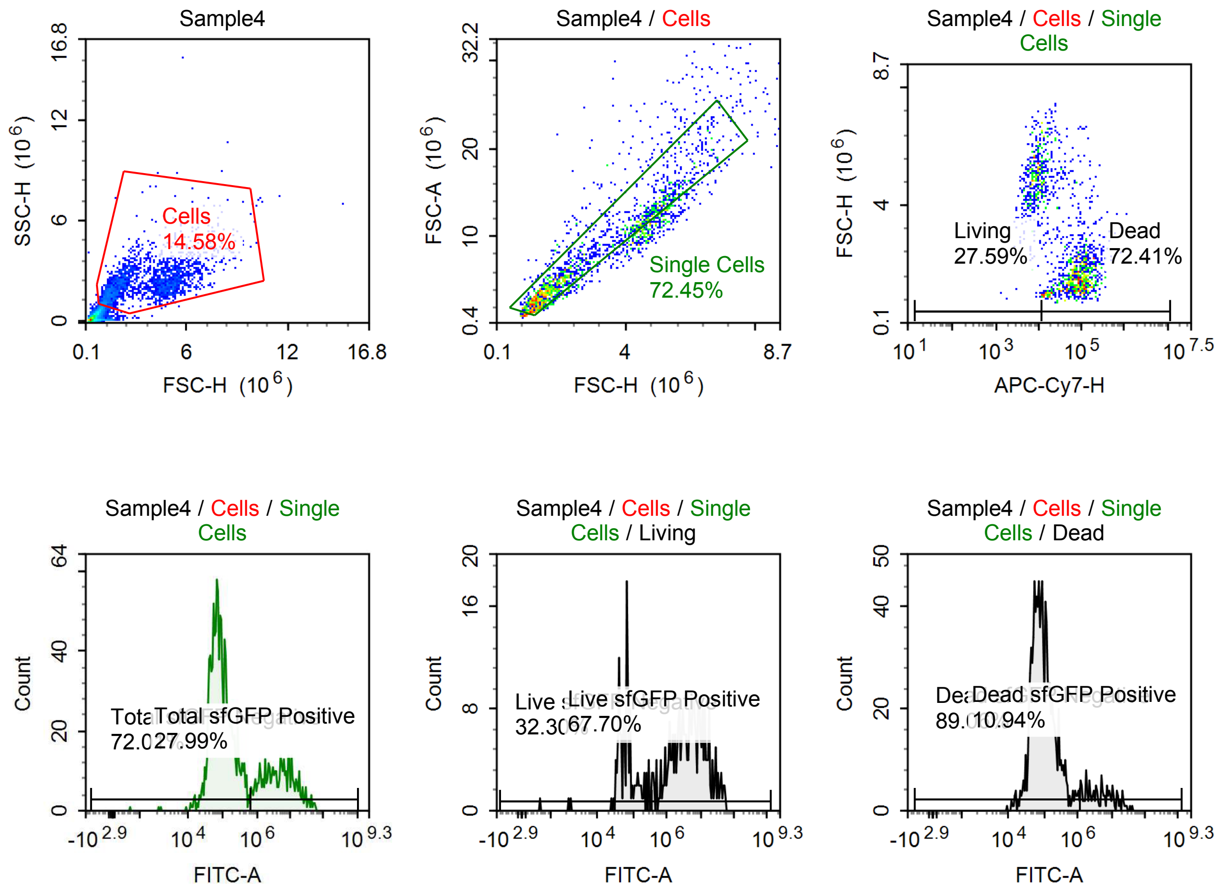

Sample Statistics of Sample4

| Gate | Count | % Parent | X | Y | Median X | Median Y |
|---|---|---|---|---|---|---|
| All | 14,345 | | | | | |
| Cells | 2,091 | 14.58 % | FSC-H | SSC-H | 3,043,987 | 2,183,868 |
| Single Cells | 1,515 | 72.45 % | FSC-H | FSC-A | 2,216,708 | 5,836,150 |
| Living | 418 | 27.59 % | APC-Cy7-H | FSC-H | 7,873 | 4,769,799 |
| Live sfGFP Negative | 135 | 32.30 % | FITC-A | Count | 67,298 | |
| Live sfGFP Positive | 283 | 67.70 % | FITC-A | Count | 5,242,210 | |
| Dead | 1,097 | 72.41 % | APC-Cy7-H | FSC-H | 75,297 | 1,746,243 |
| Dead sfGFP Negative | 977 | 89.06 % | FITC-A | Count | 86,023 | |
| Dead sfGFP Positive | 120 | 10.94 % | FITC-A | Count | 5,271,745 | |
| Total sfGFP Negative | 1,091 | 72.01 % | FITC-A | Count | 80,948 | |
| Total sfGFP Positive | 424 | 27.99 % | FITC-A | Count | 4,974,250 | |

## p005

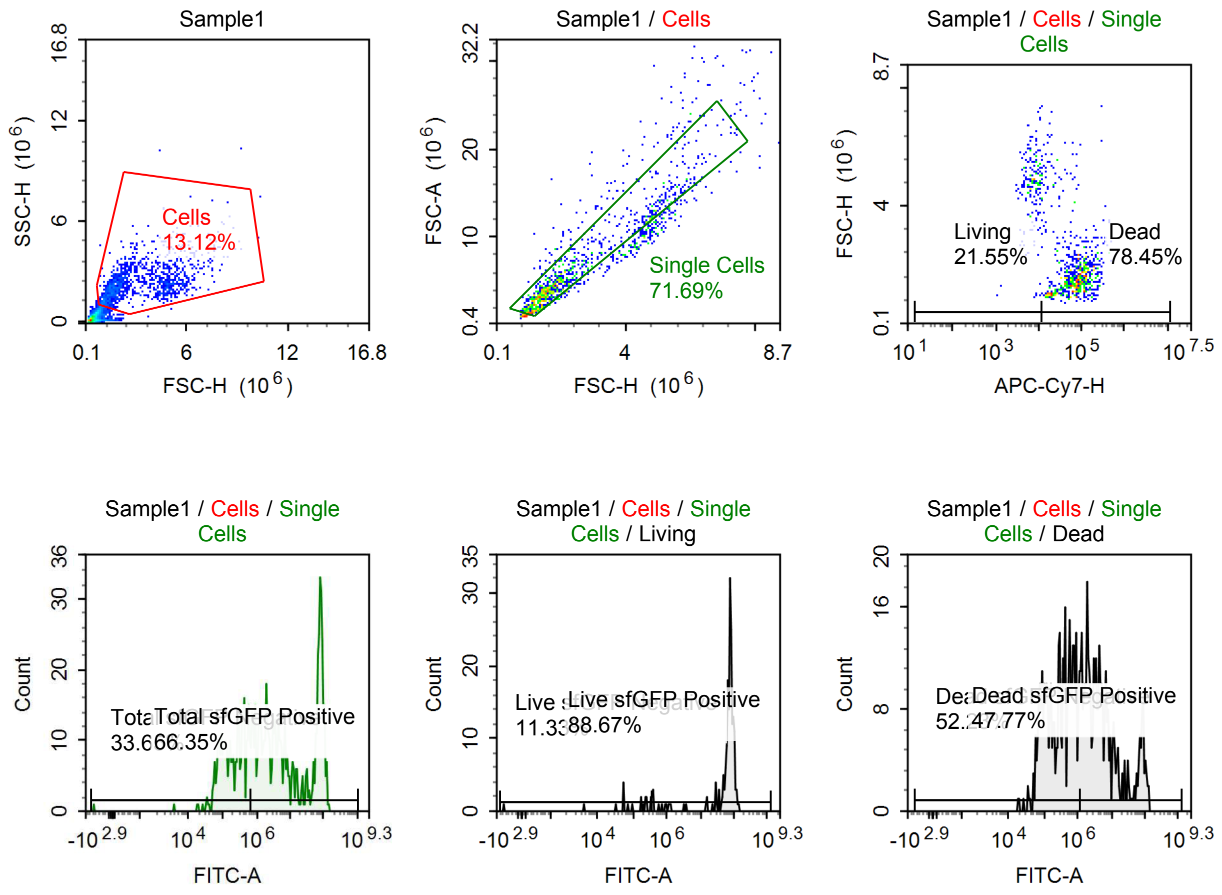

Sample Statistics of Sample1

| Gate | Count | % Parent | X | Y | Median X | Median Y |
|---|---|---|---|---|---|---|
| All | 10,017 | | | | | |
| Cells | 1,314 | 13.12 % | FSC-H | SSC-H | 2,082,155 | 2,120,415 |
| Single Cells | 942 | 71.69 % | FSC-H | FSC-A | 1,798,431 | 4,588,544 |
| Living | 203 | 21.55 % | APC-Cy7-H | FSC-H | 7,504 | 4,802,663 |
| Live sfGFP Negative | 23 | 11.33 % | FITC-A | Count | 198,733 | |
| Live sfGFP Positive | 180 | 88.67 % | FITC-A | Count | 75,811,256 | |
| Dead | 739 | 78.45 % | APC-Cy7-H | FSC-H | 75,785 | 1,563,462 |
| Dead sfGFP Negative | 386 | 52.23 % | FITC-A | Count | 311,693 | |
| Dead sfGFP Positive | 353 | 47.77 % | FITC-A | Count | 4,760,618 | |
| Total sfGFP Negative | 317 | 33.65 % | FITC-A | Count | 199,702 | |
| Total sfGFP Positive | 625 | 66.35 % | FITC-A | Count | 6,749,131 | |

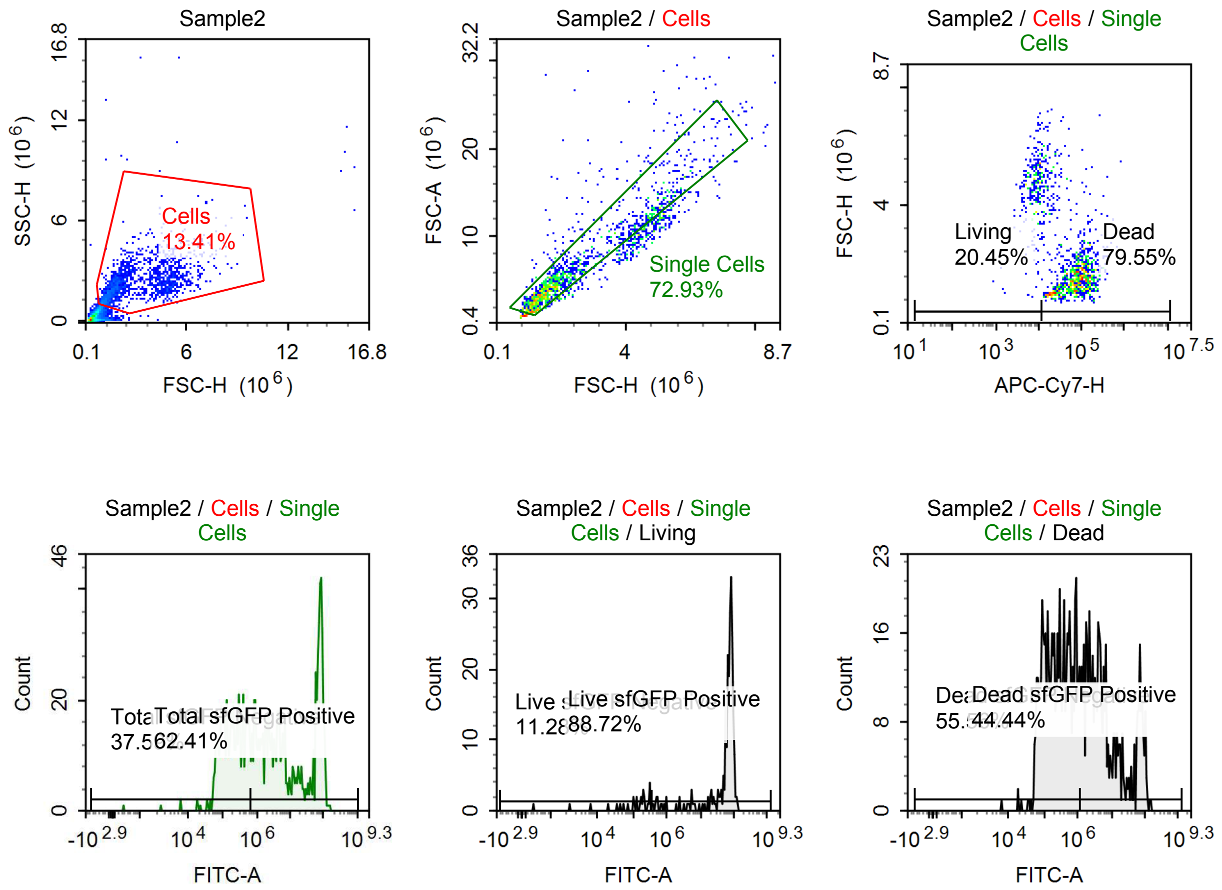

Sample Statistics of Sample2

| Gate | Count | % Parent | X | Y | Median X | Median Y |
|---|---|---|---|---|---|---|
| All | 13,299 | | | | | |
| Cells | 1,784 | 13.41 % | FSC-H | SSC-H | 2,013,684 | 2,070,146 |
| Single Cells | 1,301 | 72.93 % | FSC-H | FSC-A | 1,822,490 | 4,734,554 |
| Living | 266 | 20.45 % | APC-Cy7-H | FSC-H | 8,060 | 4,735,526 |
| Live sfGFP Negative | 30 | 11.28 % | FITC-A | Count | 175,280 | |
| Live sfGFP Positive | 236 | 88.72 % | FITC-A | Count | 73,413,256 | |
| Dead | 1,035 | 79.55 % | APC-Cy7-H | FSC-H | 72,016 | 1,590,153 |
| Dead sfGFP Negative | 575 | 55.56 % | FITC-A | Count | 263,300 | |
| Dead sfGFP Positive | 460 | 44.44 % | FITC-A | Count | 5,433,141 | |
| Total sfGFP Negative | 489 | 37.59 % | FITC-A | Count | 187,682 | |
| Total sfGFP Positive | 812 | 62.41 % | FITC-A | Count | 6,969,156 | |

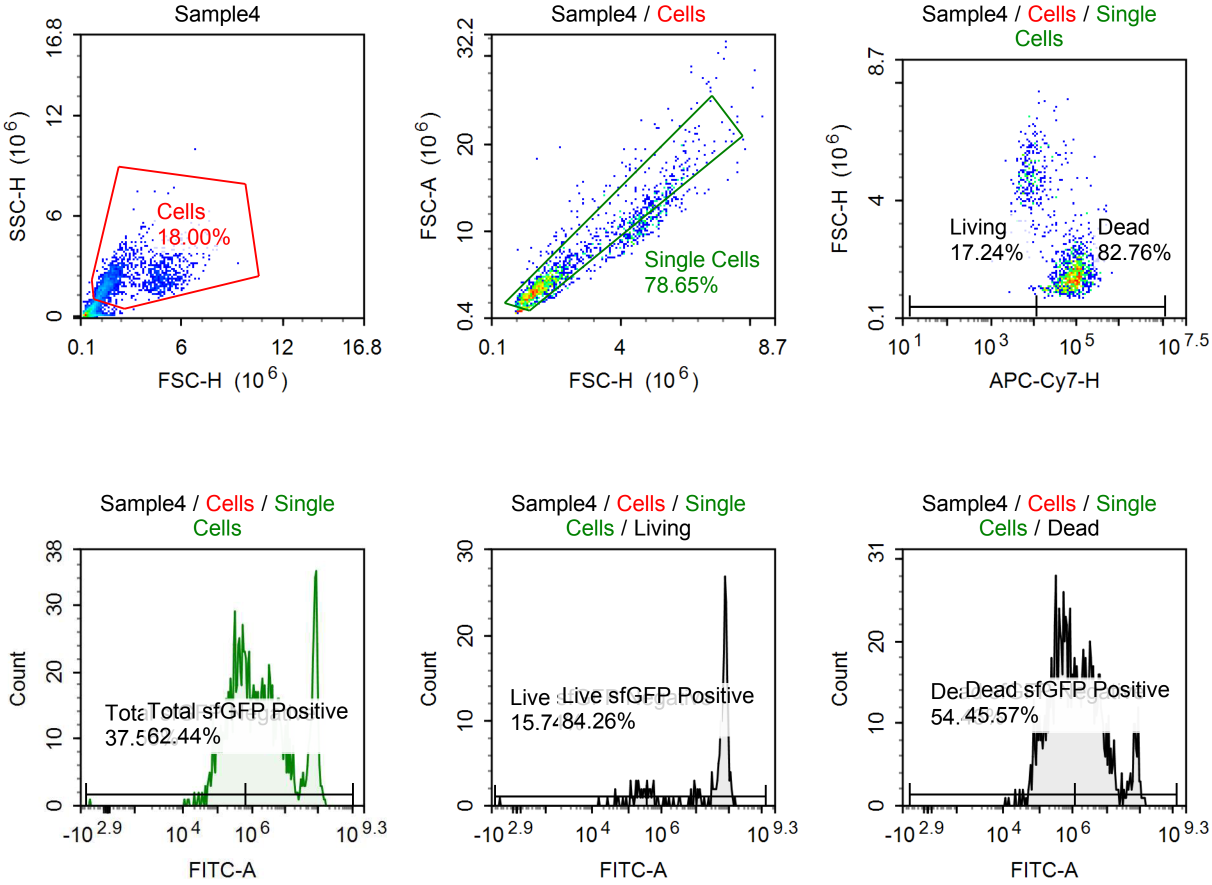

Sample Statistics of Sample4

| Gate | Count | % Parent | X | Y | Median X | Median Y |
|---|---|---|---|---|---|---|
| All | 9,630 | | | | | |
| Cells | 1,733 | 18.00 % | FSC-H | SSC-H | 1,830,575 | 2,057,359 |
| Single Cells | 1,363 | 78.65 % | FSC-H | FSC-A | 1,667,055 | 4,296,000 |
| Living | 235 | 17.24 % | APC-Cy7-H | FSC-H | 7,308 | 4,752,562 |
| Live sfGFP Negative | 37 | 15.74 % | FITC-A | Count | 188,430 | |
| Live sfGFP Positive | 198 | 84.26 % | FITC-A | Count | 71,224,880 | |
| Dead | 1,128 | 82.76 % | APC-Cy7-H | FSC-H | 78,943 | 1,517,743 |
| Dead sfGFP Negative | 614 | 54.43 % | FITC-A | Count | 387,081 | |
| Dead sfGFP Positive | 514 | 45.57 % | FITC-A | Count | 4,654,434 | |
| Total sfGFP Negative | 512 | 37.56 % | FITC-A | Count | 288,201 | |
| Total sfGFP Positive | 851 | 62.44 % | FITC-A | Count | 4,963,829 | |

#### p005 + 1 uM Tet

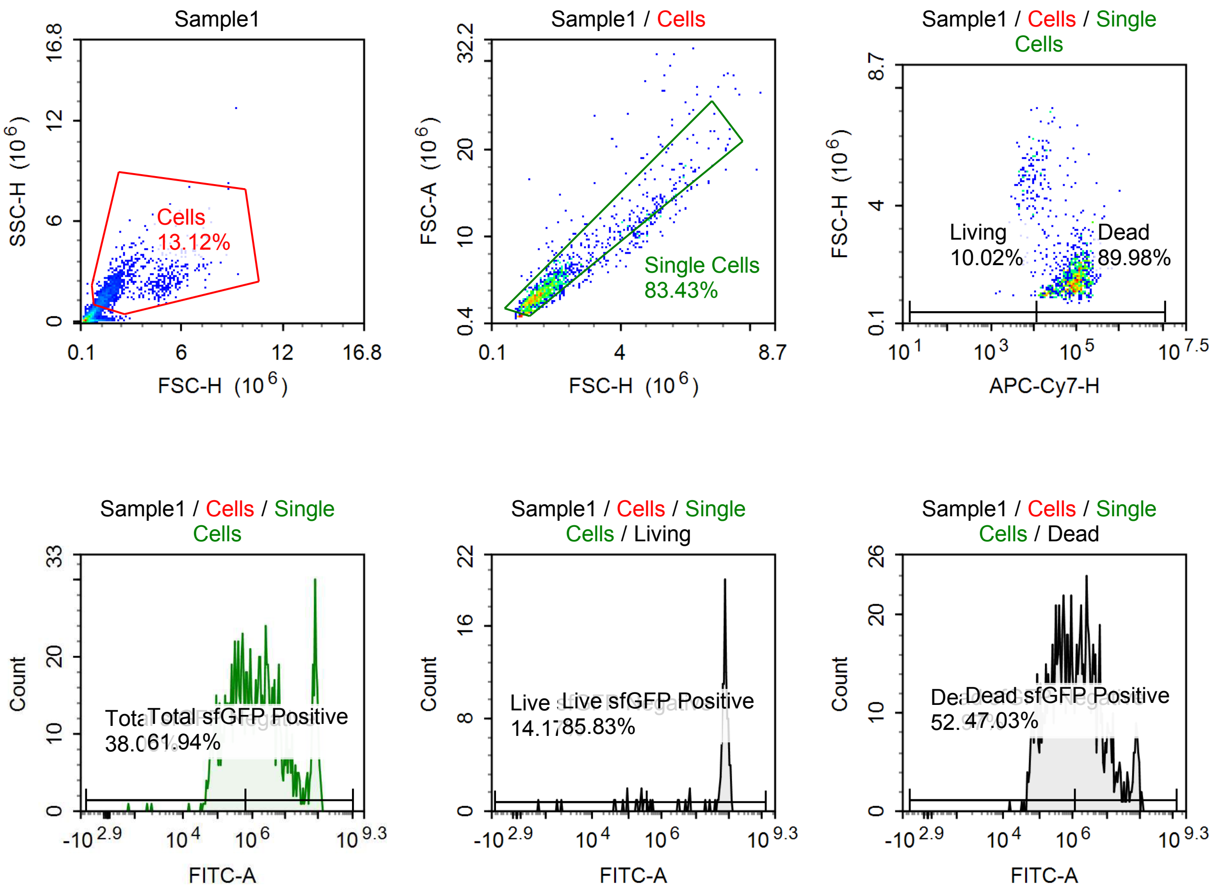

Sample Statistics of Sample1

| Gate | Count | % Parent | X | Y | Median X | Median Y |
|---|---|---|---|---|---|---|
| All | 10,948 | | | | | |
| Cells | 1,436 | 13.12 % | FSC-H | SSC-H | 1,657,089 | 1,791,870 |
| Single Cells | 1,198 | 83.43 % | FSC-H | FSC-A | 1,603,246 | 3,923,249 |
| Living | 120 | 10.02 % | APC-Cy7-H | FSC-H | 7,111 | 4,557,723 |
| Live sfGFP Negative | 17 | 14.17 % | FITC-A | Count | 180,582 | |
| Live sfGFP Positive | 103 | 85.83 % | FITC-A | Count | 75,845,472 | |
| Dead | 1,078 | 89.98 % | APC-Cy7-H | FSC-H | 85,744 | 1,518,264 |
| Dead sfGFP Negative | 571 | 52.97 % | FITC-A | Count | 339,908 | |
| Dead sfGFP Positive | 507 | 47.03 % | FITC-A | Count | 4,368,765 | |
| Total sfGFP Negative | 456 | 38.06 % | FITC-A | Count | 243,932 | |
| Total sfGFP Positive | 742 | 61.94 % | FITC-A | Count | 3,989,113 | |

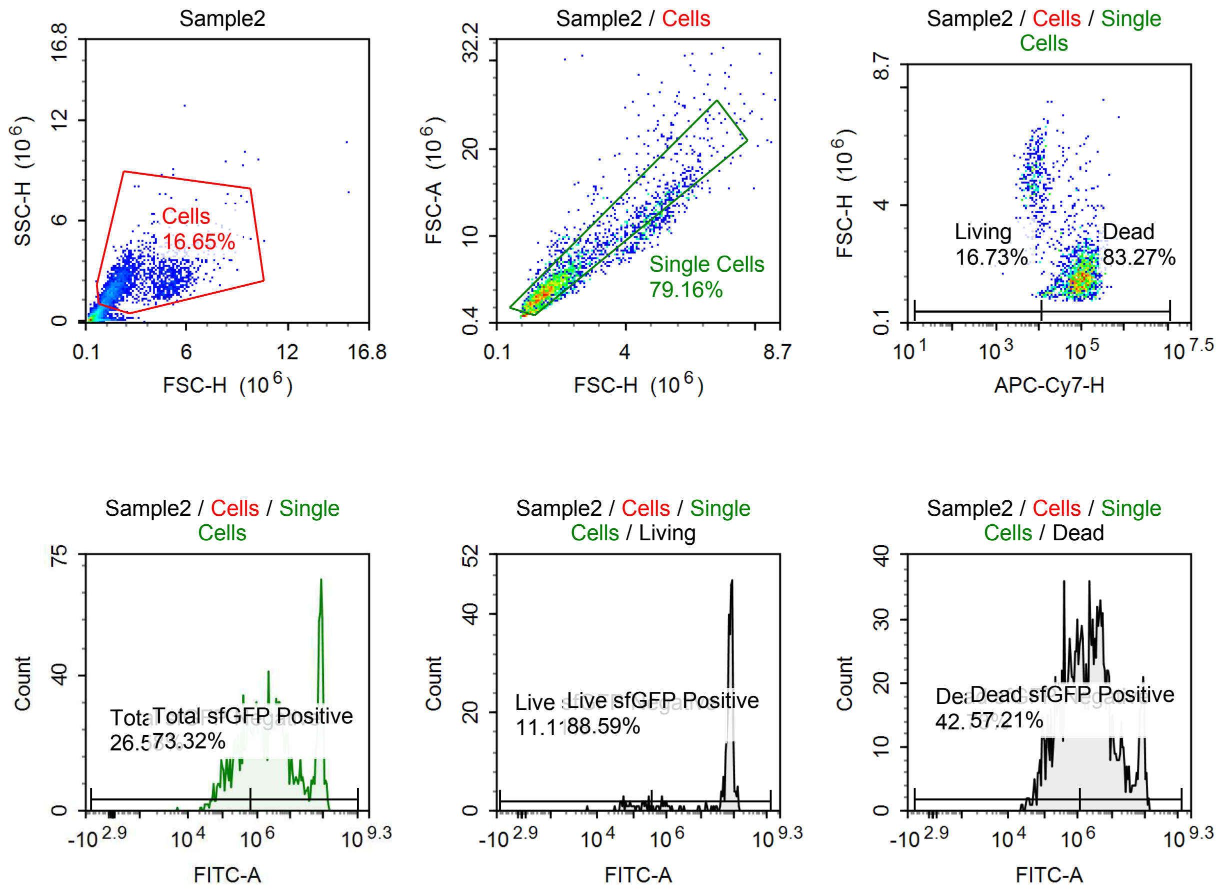

Sample Statistics of Sample2

| Gate | Count | % Parent | X | Y | Median X | Median Y |
|---|---|---|---|---|---|---|
| All | 15,098 | | | | | |
| Cells | 2,514 | 16.65 % | FSC-H | SSC-H | 1,923,021 | 2,080,108 |
| Single Cells | 1,990 | 79.16 % | FSC-H | FSC-A | 1,759,779 | 4,530,289 |
| Living | 333 | 16.73 % | APC-Cy7-H | FSC-H | 7,555 | 4,745,803 |
| Live sfGFP Negative | 37 | 11.11 % | FITC-A | Count | 93,849 | |
| Live sfGFP Positive | 295 | 88.59 % | FITC-A | Count | 76,981,856 | |
| Dead | 1,657 | 83.27 % | APC-Cy7-H | FSC-H | 90,748 | 1,625,003 |
| Dead sfGFP Negative | 708 | 42.73 % | FITC-A | Count | 426,301 | |
| Dead sfGFP Positive | 948 | 57.21 % | FITC-A | Count | 5,184,143 | |
| Total sfGFP Negative | 529 | 26.58 % | FITC-A | Count | 263,581 | |
| Total sfGFP Positive | 1,459 | 73.32 % | FITC-A | Count | 5,421,632 | |

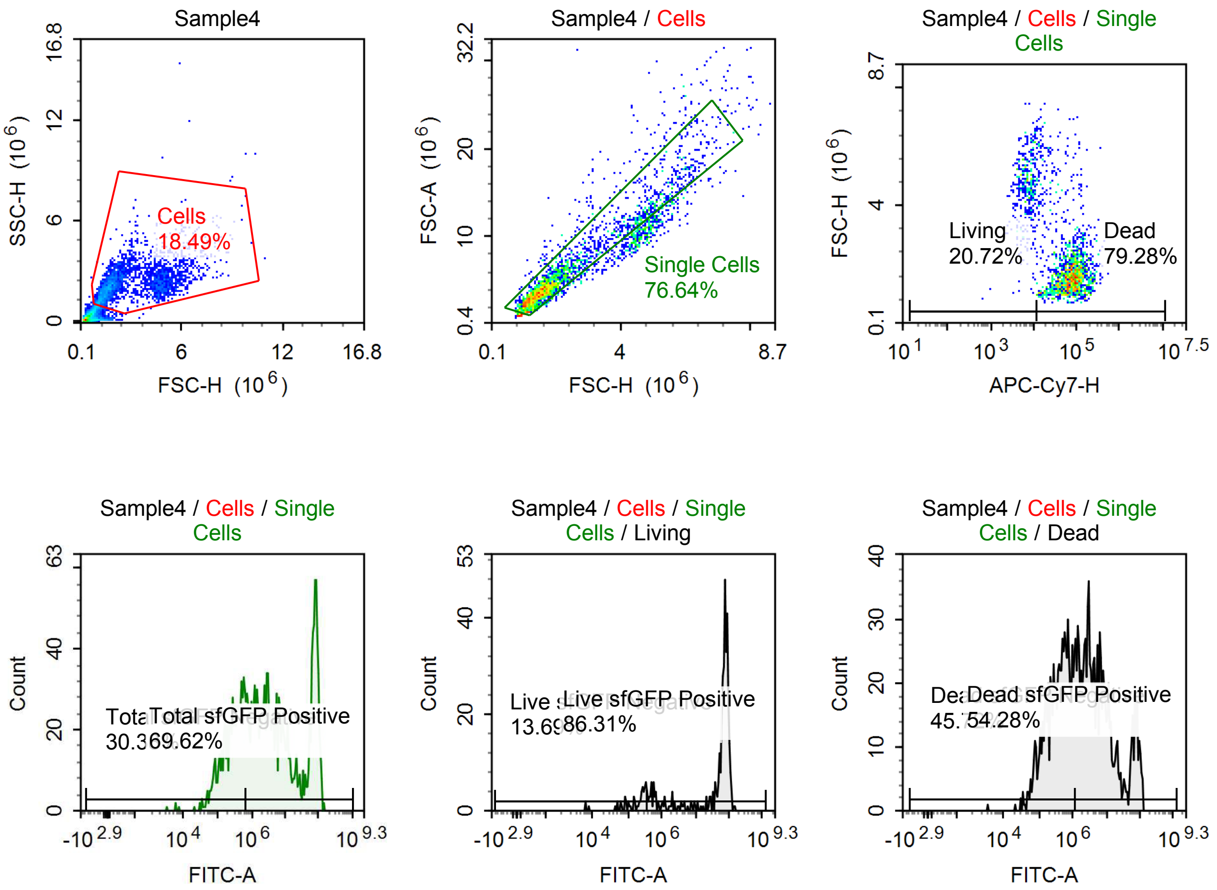

Sample Statistics of Sample4

| Gate | Count | % Parent | X | Y | Median X | Median Y |
|---|---|---|---|---|---|---|
| All | 14,676 | | | | | |
| Cells | 2,714 | 18.49 % | FSC-H | SSC-H | 2,119,628 | 2,185,568 |
| Single Cells | 2,080 | 76.64 % | FSC-H | FSC-A | 1,829,528 | 4,695,843 |
| Living | 431 | 20.72 % | APC-Cy7-H | FSC-H | 6,894 | 4,737,332 |
| Live sfGFP Negative | 59 | 13.69 % | FITC-A | Count | 249,143 | |
| Live sfGFP Positive | 372 | 86.31 % | FITC-A | Count | 71,891,008 | |
| Dead | 1,649 | 79.28 % | APC-Cy7-H | FSC-H | 72,618 | 1,622,334 |
| Dead sfGFP Negative | 754 | 45.72 % | FITC-A | Count | 404,920 | |
| Dead sfGFP Positive | 895 | 54.28 % | FITC-A | Count | 5,611,933 | |
| Total sfGFP Negative | 632 | 30.38 % | FITC-A | Count | 282,384 | |
| Total sfGFP Positive | 1,448 | 69.62 % | FITC-A | Count | 6,531,981 | |

#### p061 +10 uM Tet

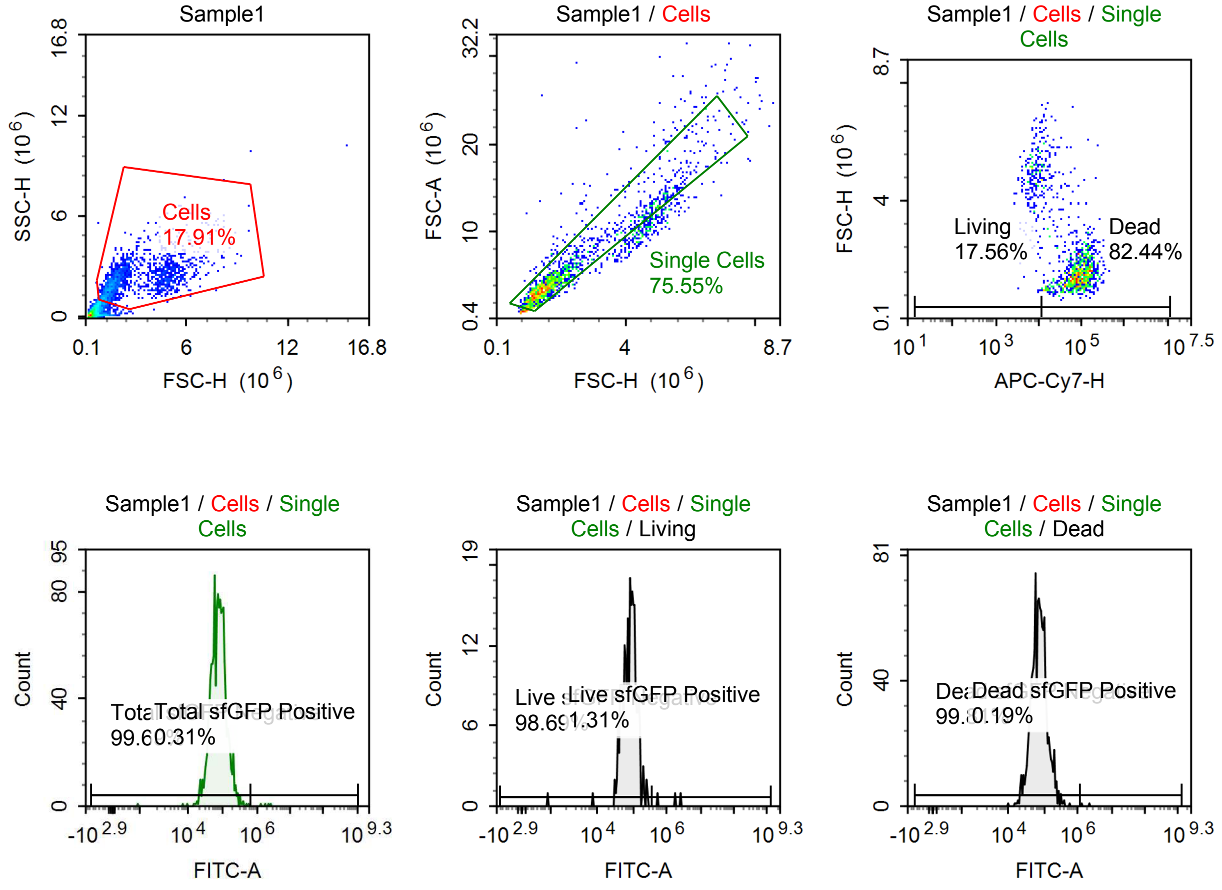

Sample Statistics of Sample1

| Gate | Count | % Parent | X | Y | Median X | Median Y |
|---|---|---|---|---|---|---|
| All | 9,637 | | | | | |
| Cells | 1,726 | 17.91 % | FSC-H | SSC-H | 1,903,663 | 2,095,401 |
| Single Cells | 1,304 | 75.55 % | FSC-H | FSC-A | 1,681,919 | 4,311,472 |
| Living | 229 | 17.56 % | APC-Cy7-H | FSC-H | 7,672 | 4,773,122 |
| Live sfGFP Negative | 226 | 98.69 % | FITC-A | Count | 84,656 | |
| Live sfGFP Positive | 3 | 1.31 % | FITC-A | Count | 1,763,407 | |
| Dead | 1,075 | 82.44 % | APC-Cy7-H | FSC-H | 79,414 | 1,537,484 |
| Dead sfGFP Negative | 1,073 | 99.81 % | FITC-A | Count | 73,204 | |
| Dead sfGFP Positive | 2 | 0.19 % | FITC-A | Count | 1,770,554 | |
| Total sfGFP Negative | 1,300 | 99.69 % | FITC-A | Count | 74,544 | |
| Total sfGFP Positive | 4 | 0.31 % | FITC-A | Count | 1,976,821 | |

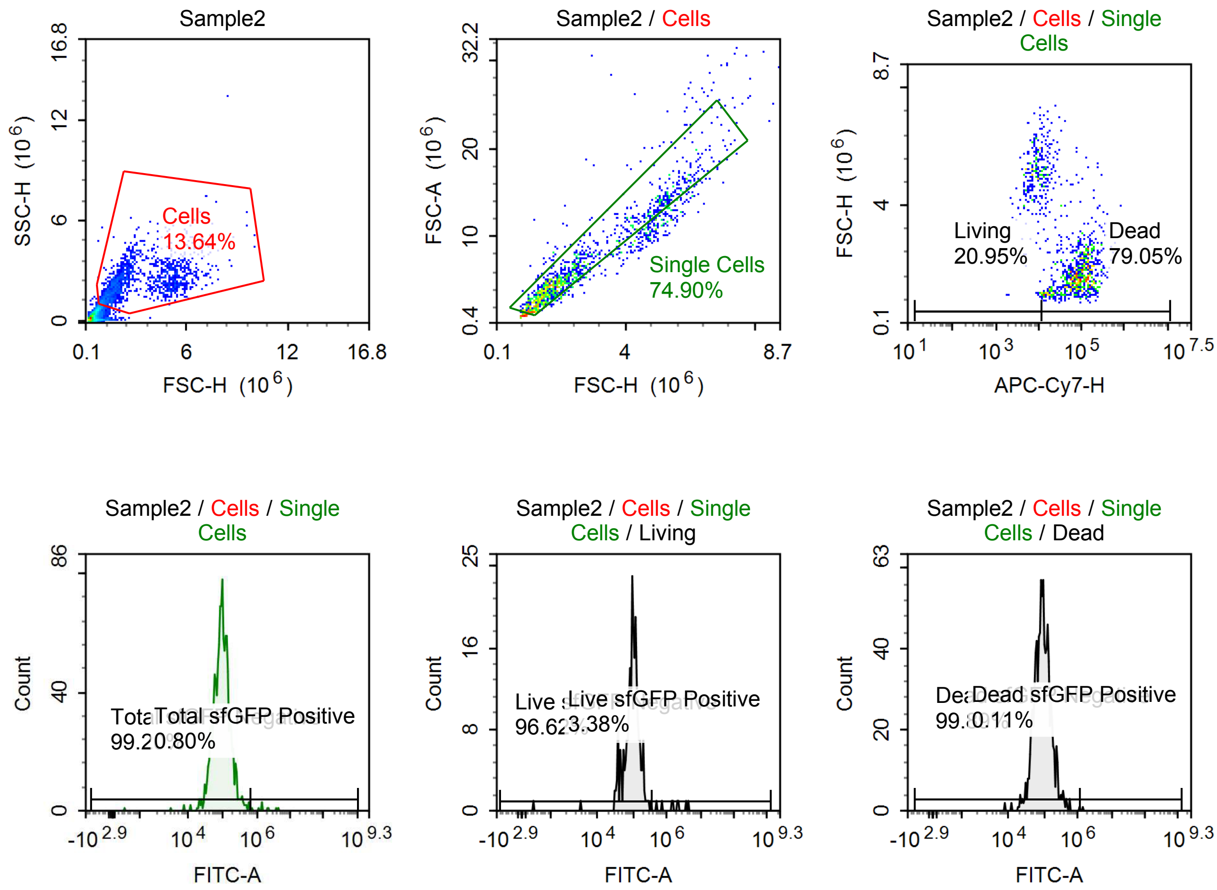

Sample Statistics of Sample2

| Gate | Count | % Parent | X | Y | Median X | Median Y |
|---|---|---|---|---|---|---|
| All | 11,073 | | | | | |
| Cells | 1,510 | 13.64 % | FSC-H | SSC-H | 2,088,438 | 2,198,113 |
| Single Cells | 1,131 | 74.90 % | FSC-H | FSC-A | 1,943,480 | 5,014,571 |
| Living | 237 | 20.95 % | APC-Cy7-H | FSC-H | 8,067 | 4,949,725 |
| Live sfGFP Negative | 229 | 96.62 % | FITC-A | Count | 98,709 | |
| Live sfGFP Positive | 8 | 3.38 % | FITC-A | Count | 1,782,320 | |
| Dead | 894 | 79.05 % | APC-Cy7-H | FSC-H | 78,402 | 1,721,161 |
| Dead sfGFP Negative | 893 | 99.89 % | FITC-A | Count | 90,460 | |
| Dead sfGFP Positive | 1 | 0.11 % | FITC-A | Count | 1,460,122 | |
| Total sfGFP Negative | 1,122 | 99.20 % | FITC-A | Count | 92,725 | |
| Total sfGFP Positive | 9 | 0.80 % | FITC-A | Count | 1,483,823 | |

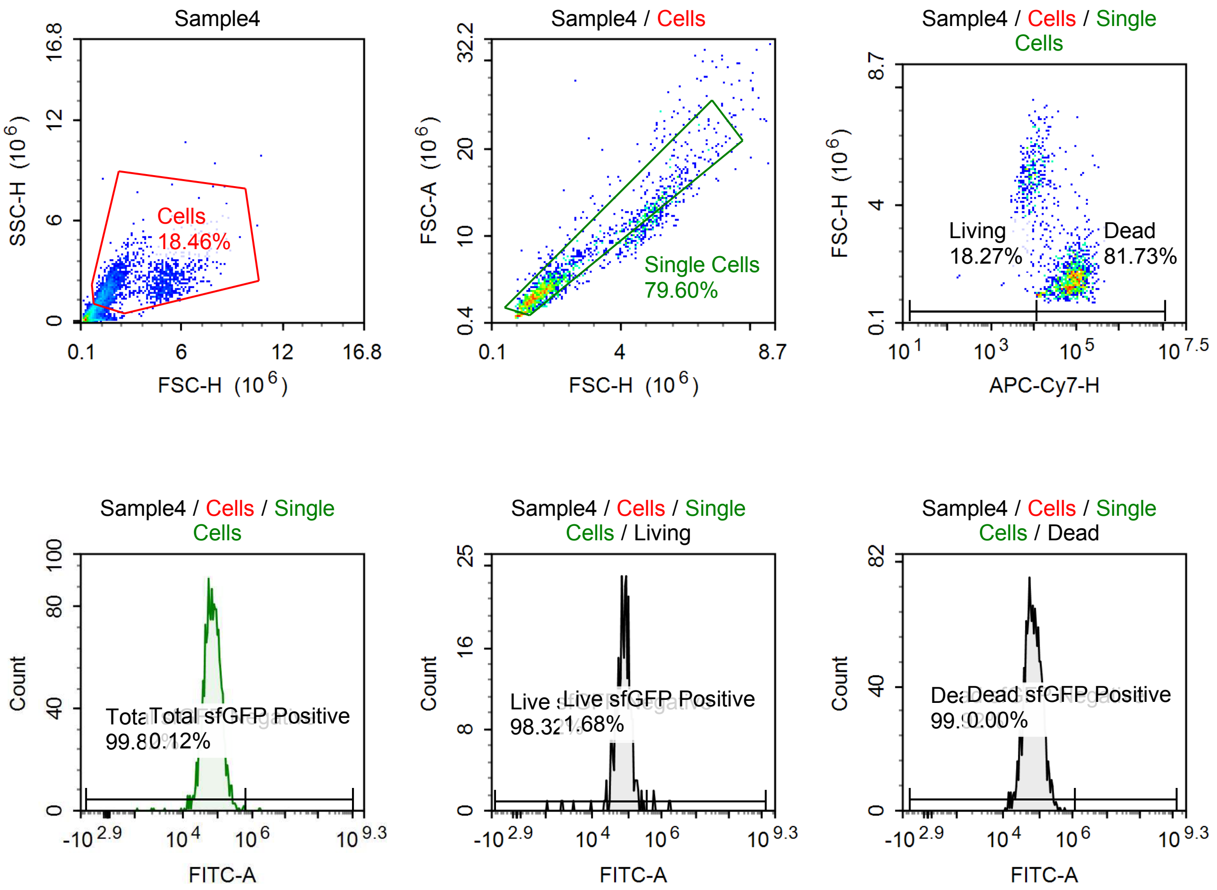

Sample Statistics of Sample4

| Gate | Count | % Parent | X | Y | Median X | Median Y |
|---|---|---|---|---|---|---|
| All | 11,097 | | | | | |
| Cells | 2,049 | 18.46 % | FSC-H | SSC-H | 1,800,433 | 1,950,915 |
| Single Cells | 1,631 | 79.60 % | FSC-H | FSC-A | 1,687,531 | 4,253,554 |
| Living | 298 | 18.27 % | APC-Cy7-H | FSC-H | 7,779 | 4,800,980 |
| Live sfGFP Negative | 293 | 98.32 % | FITC-A | Count | 75,653 | |
| Live sfGFP Positive | 5 | 1.68 % | FITC-A | Count | 605,404 | |
| Dead | 1,333 | 81.73 % | APC-Cy7-H | FSC-H | 73,068 | 1,556,706 |
| Dead sfGFP Negative | 1,332 | 99.92 % | FITC-A | Count | 66,691 | |
| Dead sfGFP Positive | 0 | 0.00 % | FITC-A | Count | 0 | |
| Total sfGFP Negative | 1,628 | 99.82 % | FITC-A | Count | 68,647 | |
| Total sfGFP Positive | 2 | 0.12 % | FITC-A | Count | 1,686,359 | |

#### p061 + 100 uM Tet

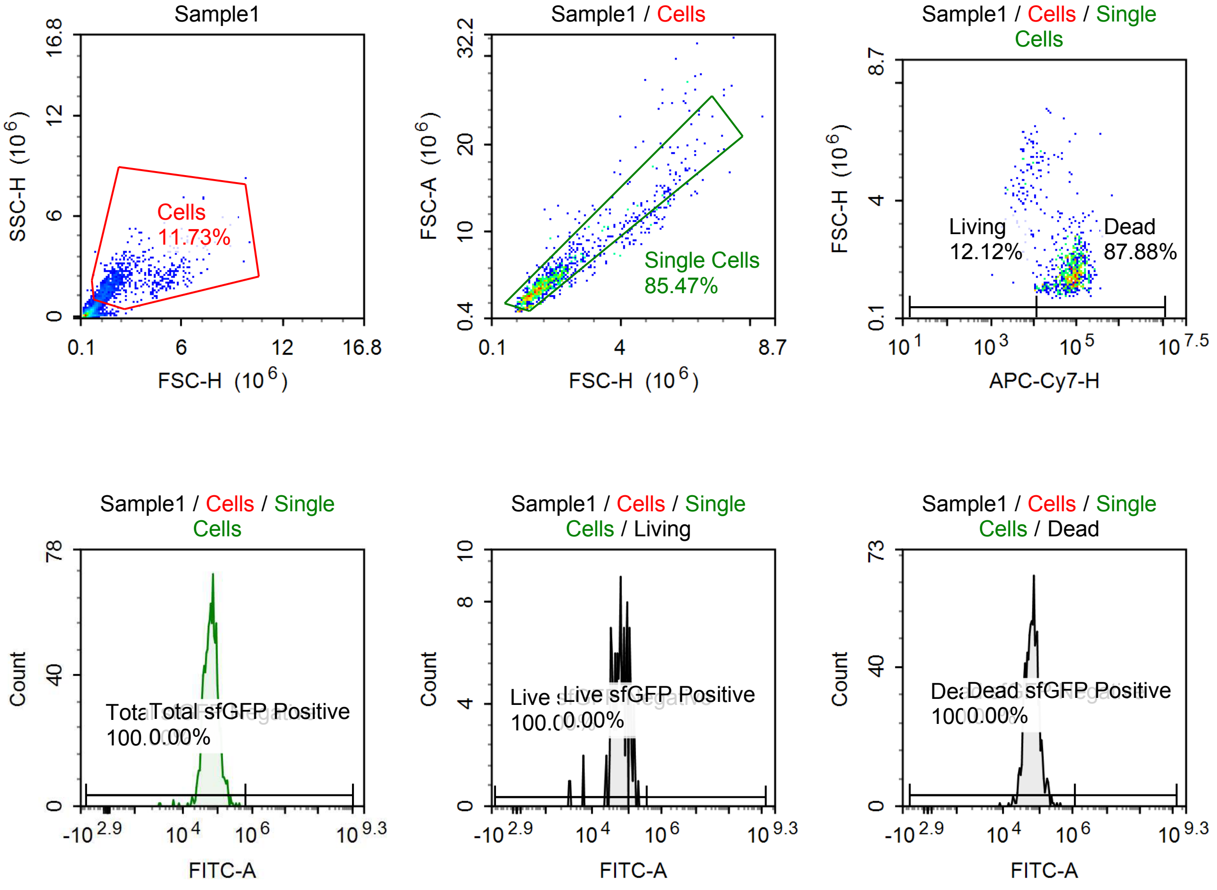

Sample Statistics of Sample1

| Gate | Count | % Parent | X | Y | Median X | Median Y |
|---|---|---|---|---|---|---|
| All | 9,626 | | | | | |
| Cells | 1,129 | 11.73 % | FSC-H | SSC-H | 1,709,489 | 1,817,422 |
| Single Cells | 965 | 85.47 % | FSC-H | FSC-A | 1,614,749 | 4,157,375 |
| Living | 117 | 12.12 % | APC-Cy7-H | FSC-H | 7,147 | 4,260,092 |
| Live sfGFP Negative | 117 | 100.00 % | FITC-A | Count | 62,689 | |
| Live sfGFP Positive | 0 | 0.00 % | FITC-A | Count | 0 | |
| Dead | 848 | 87.88 % | APC-Cy7-H | FSC-H | 78,260 | 1,542,945 |
| Dead sfGFP Negative | 848 | 100.00 % | FITC-A | Count | 63,579 | |
| Dead sfGFP Positive | 0 | 0.00 % | FITC-A | Count | 0 | |
| Total sfGFP Negative | 965 | 100.00 % | FITC-A | Count | 63,516 | |
| Total sfGFP Positive | 0 | 0.00 % | FITC-A | Count | 0 | |

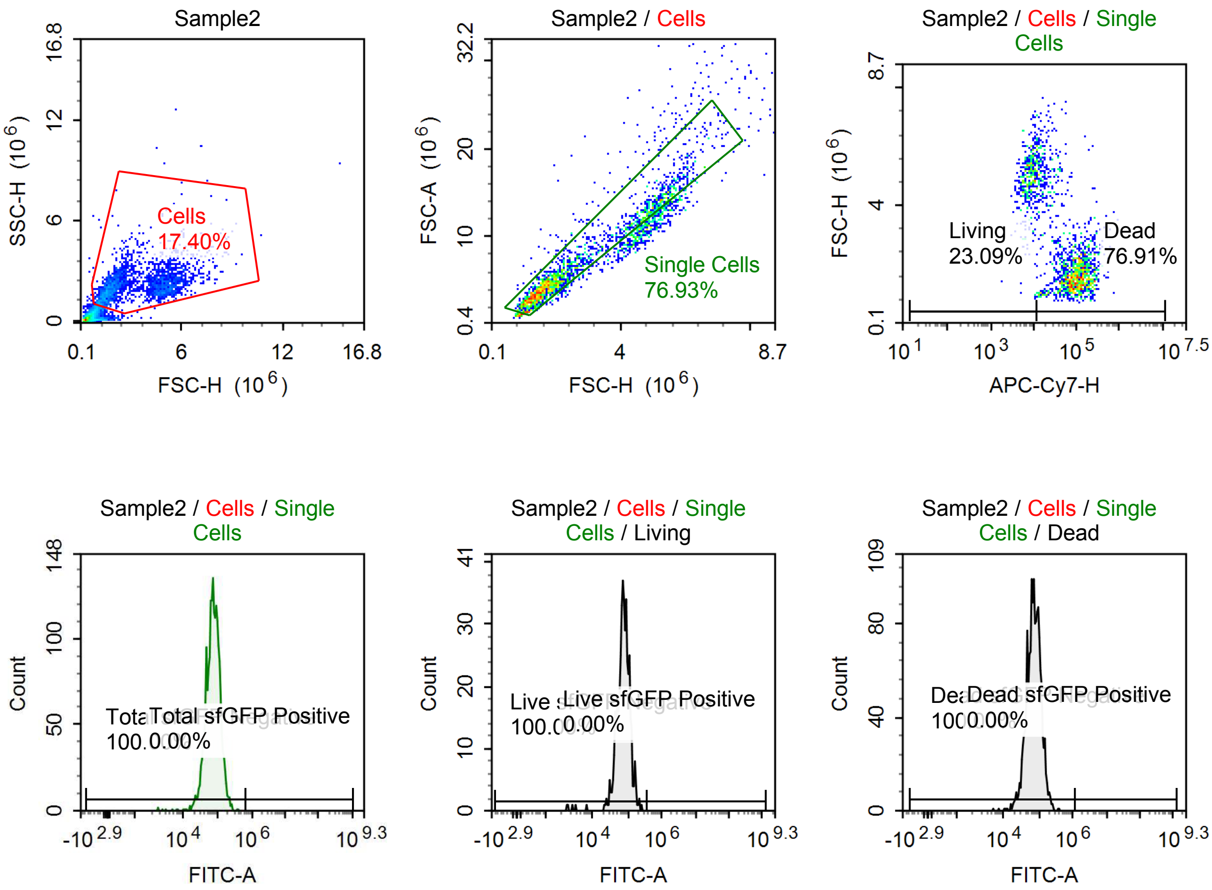

Sample Statistics of Sample2

| Gate | Count | % Parent | X | Y | Median X | Median Y |
|---|---|---|---|---|---|---|
| All | 14,076 | | | | | |
| Cells | 2,449 | 17.40 % | FSC-H | SSC-H | 2,630,008 | 2,087,196 |
| Single Cells | 1,884 | 76.93 % | FSC-H | FSC-A | 2,082,729 | 5,472,073 |
| Living | 435 | 23.09 % | APC-Cy7-H | FSC-H | 8,039 | 4,952,888 |
| Live sfGFP Negative | 435 | 100.00 % | FITC-A | Count | 75,973 | |
| Live sfGFP Positive | 0 | 0.00 % | FITC-A | Count | 0 | |
| Dead | 1,449 | 76.91 % | APC-Cy7-H | FSC-H | 85,153 | 1,766,125 |
| Dead sfGFP Negative | 1,449 | 100.00 % | FITC-A | Count | 72,034 | |
| Dead sfGFP Positive | 0 | 0.00 % | FITC-A | Count | 0 | |
| Total sfGFP Negative | 1,884 | 100.00 % | FITC-A | Count | 72,944 | |
| Total sfGFP Positive | 0 | 0.00 % | FITC-A | Count | 0 | |

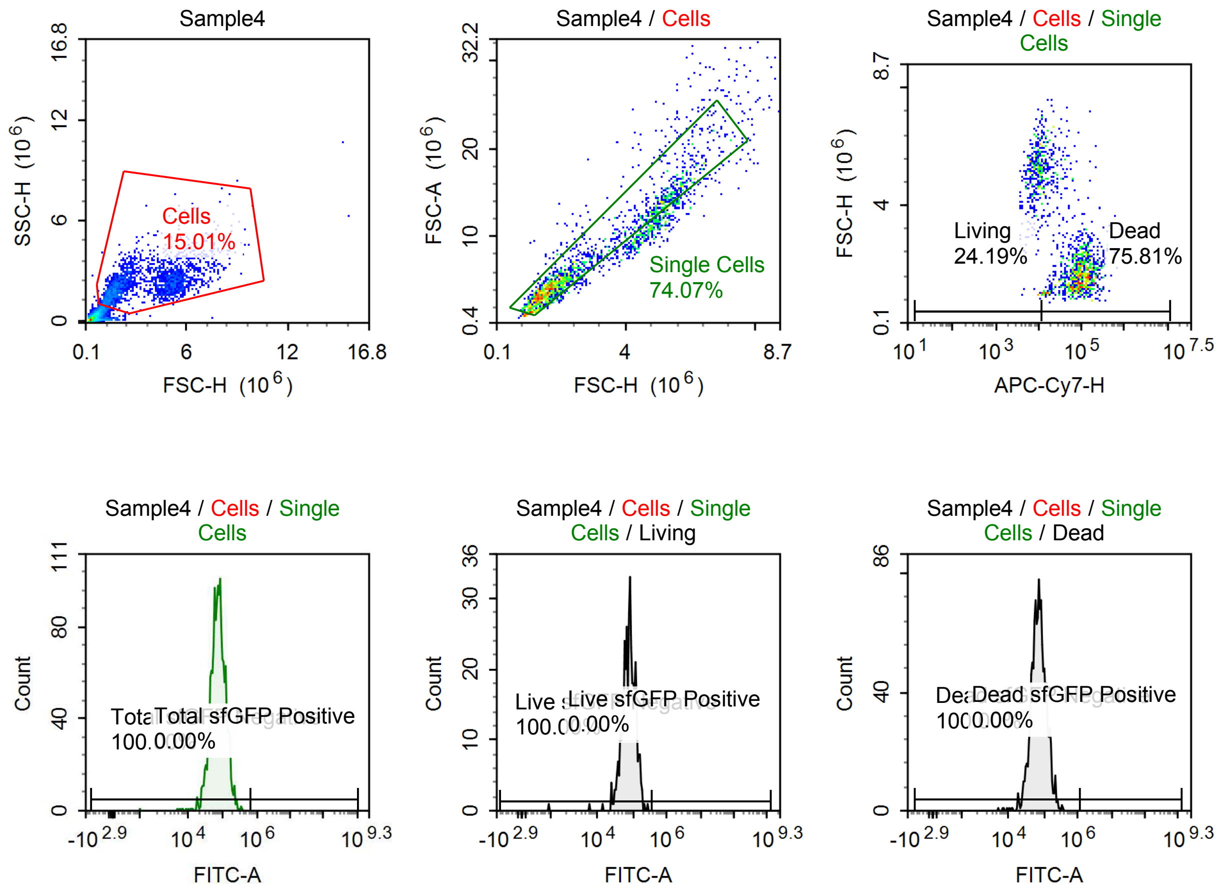

Sample Statistics of Sample4

| Gate | Count | % Parent | X | Y | Median X | Median Y |
|---|---|---|---|---|---|---|
| All | 13,641 | | | | | |
| └ Cells | 2,048 | 15.01 % | FSC-H | SSC-H | 2,718,616 | 2,184,229 |
| └ Single Cells | 1,517 | 74.07 % | FSC-H | FSC-A | 2,076,456 | 5,328,555 |
| ├ Living | 367 | 24.19 % | APC-Cy7-H | FSC-H | 8,470 | 5,118,948 |
| ├ Live sfGFP Negative | 367 | 100.00 % | FITC-A | Count | 78,291 | |
| └ Live sfGFP Positive | 0 | 0.00 % | FITC-A | Count | 0 | |
| ├ Dead | 1,150 | 75.81 % | APC-Cy7-H | FSC-H | 81,149 | 1,758,305 |
| ├ Dead sfGFP Negative | 1,150 | 100.00 % | FITC-A | Count | 71,181 | |
| └ Dead sfGFP Positive | 0 | 0.00 % | FITC-A | Count | 0 | |
| ├ Total sfGFP Negative | 1,517 | 100.00 % | FITC-A | Count | 72,790 | |
| └ Total sfGFP Positive | 0 | 0.00 % | FITC-A | Count | 0 | |

## p061

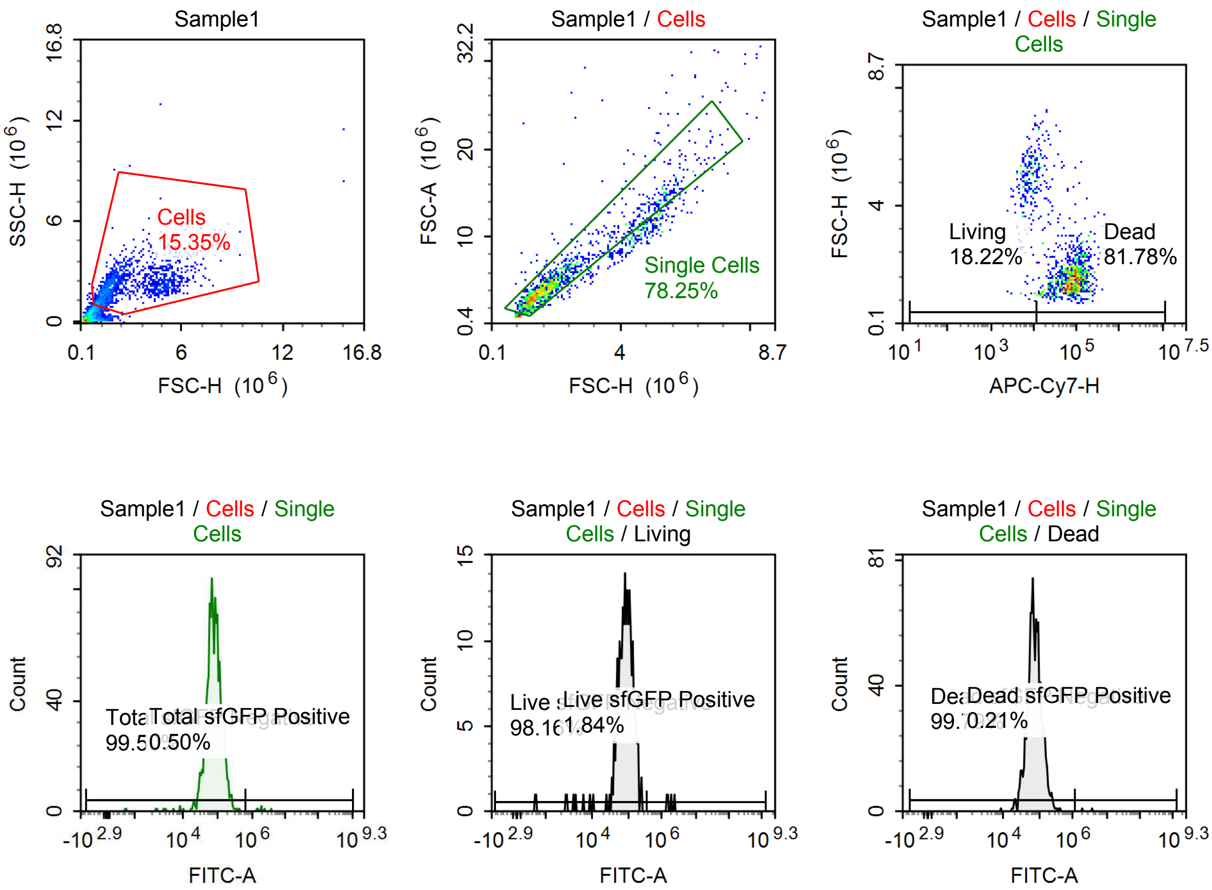

Sample Statistics of Sample1

| Gate | Count | % Parent | X | Y | Median X | Median Y |
|---|---|---|---|---|---|---|
| All | 9,914 | | | | | |
| Cells | 1,522 | 15.35 % | FSC-H | SSC-H | 1,840,930 | 2,068,938 |
| Single Cells | 1,191 | 78.25 % | FSC-H | FSC-A | 1,716,715 | 4,503,718 |
| Living | 217 | 18.22 % | APC-Cy7-H | FSC-H | 7,334 | 4,747,368 |
| Live sfGFP Negative | 213 | 98.16 % | FITC-A | Count | 86,372 | |
| Live sfGFP Positive | 4 | 1.84 % | FITC-A | Count | 1,739,934 | |
| Dead | 974 | 81.78 % | APC-Cy7-H | FSC-H | 76,300 | 1,576,727 |
| Dead sfGFP Negative | 972 | 99.79 % | FITC-A | Count | 73,148 | |
| Dead sfGFP Positive | 2 | 0.21 % | FITC-A | Count | 2,964,587 | |
| Total sfGFP Negative | 1,185 | 99.50 % | FITC-A | Count | 75,446 | |
| Total sfGFP Positive | 6 | 0.50 % | FITC-A | Count | 1,934,993 | |

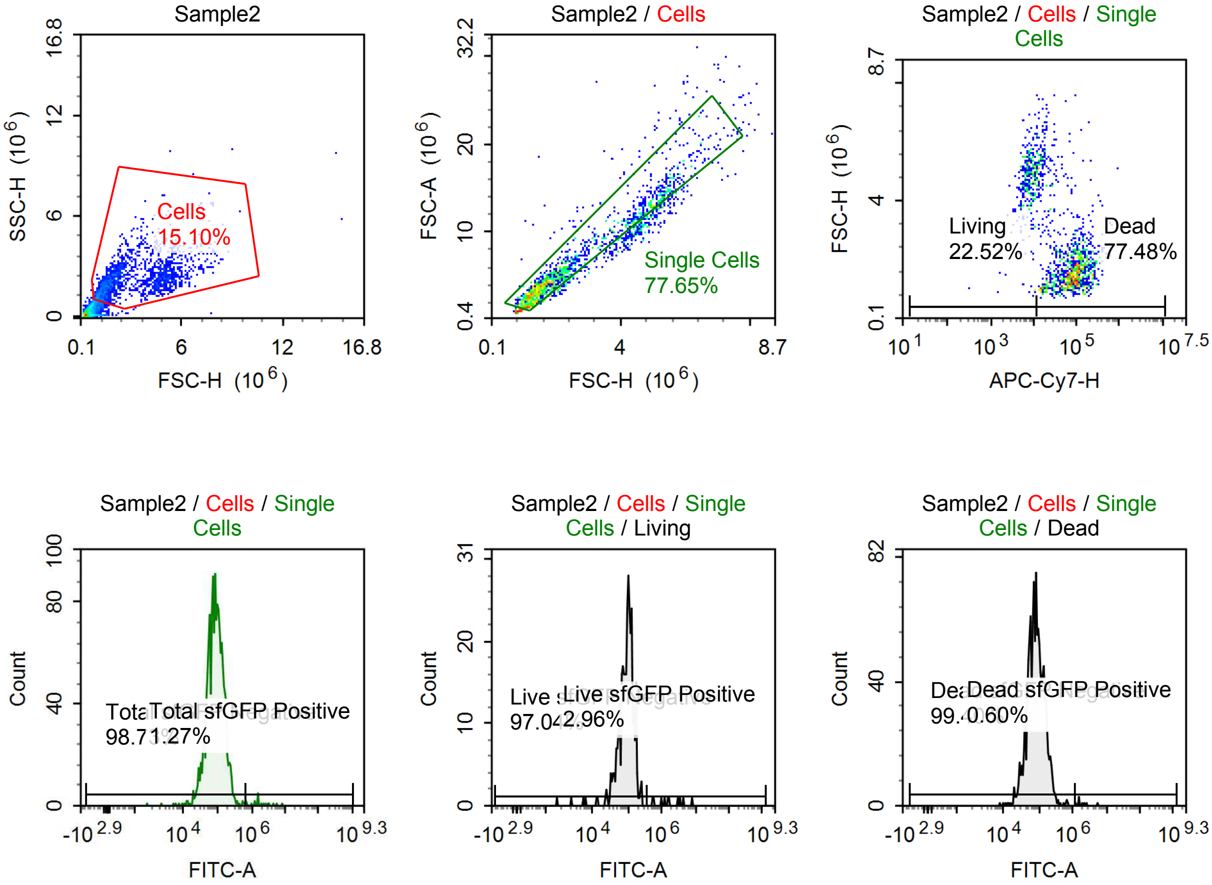

Sample Statistics of Sample2

| Gate | Count | % Parent | X | Y | Median X | Median Y |
|---|---|---|---|---|---|---|
| All | 12,798 | | | | | |
| Cells | 1,933 | 15.10 % | FSC-H | SSC-H | 2,139,981 | 2,059,416 |
| Single Cells | 1,501 | 77.65 % | FSC-H | FSC-A | 1,981,349 | 4,948,060 |
| Living | 338 | 22.52 % | APC-Cy7-H | FSC-H | 7,850 | 4,691,357 |
| Live sfGFP Negative | 328 | 97.04 % | FITC-A | Count | 98,143 | |
| Live sfGFP Positive | 10 | 2.96 % | FITC-A | Count | 1,573,372 | |
| Dead | 1,163 | 77.48 % | APC-Cy7-H | FSC-H | 74,671 | 1,678,122 |
| Dead sfGFP Negative | 1,156 | 99.40 % | FITC-A | Count | 81,581 | |
| Dead sfGFP Positive | 7 | 0.60 % | FITC-A | Count | 1,877,908 | |
| Total sfGFP Negative | 1,482 | 98.73 % | FITC-A | Count | 84,841 | |
| Total sfGFP Positive | 19 | 1.27 % | FITC-A | Count | 1,565,558 | |

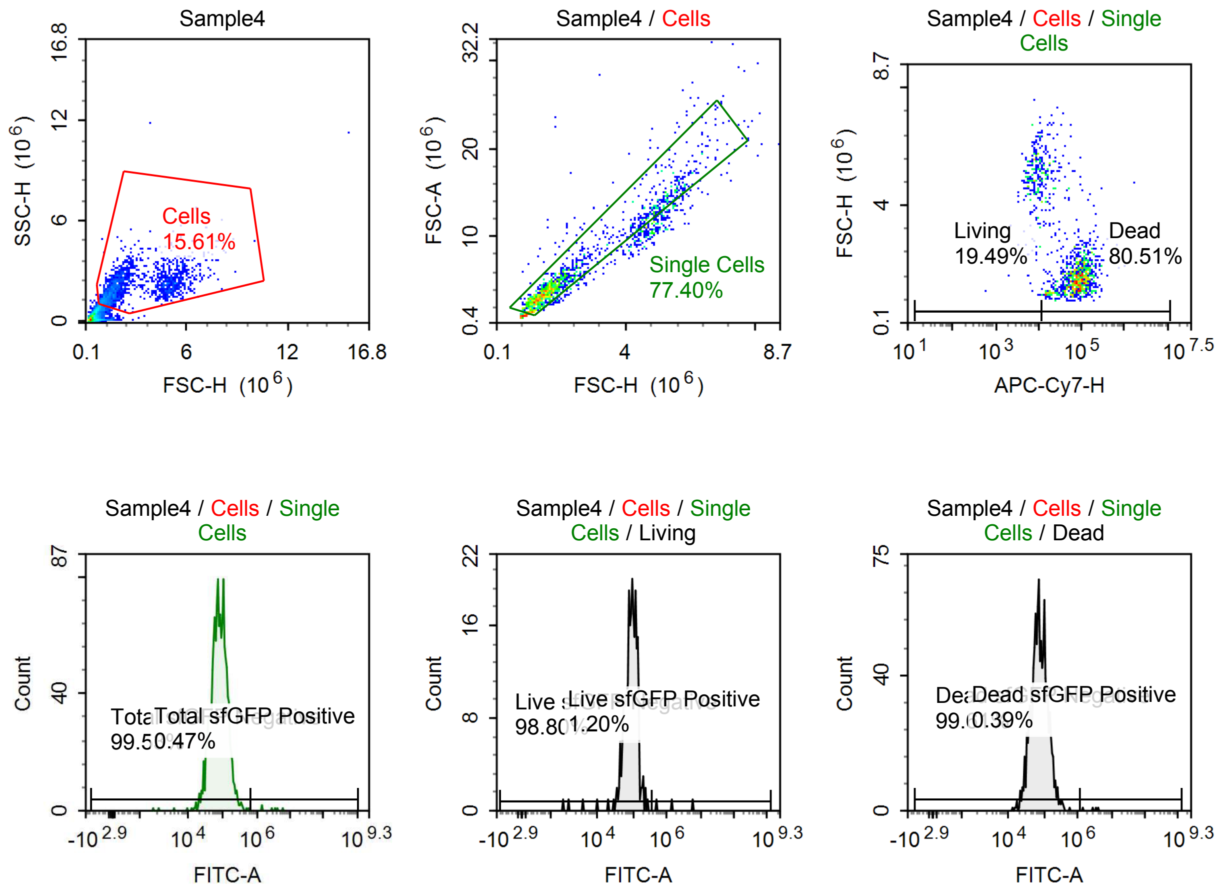

Sample Statistics of Sample4

| Gate | Count | % Parent | X | Y | Median X | Median Y |
|---|---|---|---|---|---|---|
| All | 10,662 | | | | | |
| Cells | 1,664 | 15.61 % | FSC-H | SSC-H | 1,868,142 | 1,980,056 |
| Single Cells | 1,288 | 77.40 % | FSC-H | FSC-A | 1,737,729 | 4,420,523 |
| Living | 251 | 19.49 % | APC-Cy7-H | FSC-H | 8,111 | 4,903,332 |
| Live sfGFP Negative | 248 | 98.80 % | FITC-A | Count | 90,418 | |
| Live sfGFP Positive | 3 | 1.20 % | FITC-A | Count | 1,438,194 | |
| Dead | 1,037 | 80.51 % | APC-Cy7-H | FSC-H | 76,990 | 1,593,346 |
| Dead sfGFP Negative | 1,033 | 99.61 % | FITC-A | Count | 73,534 | |
| Dead sfGFP Positive | 4 | 0.39 % | FITC-A | Count | 3,103,785 | |
| Total sfGFP Negative | 1,282 | 99.53 % | FITC-A | Count | 77,826 | |
| Total sfGFP Positive | 6 | 0.47 % | FITC-A | Count | 3,103,785 | |

#### p061 + 1 uM Tet

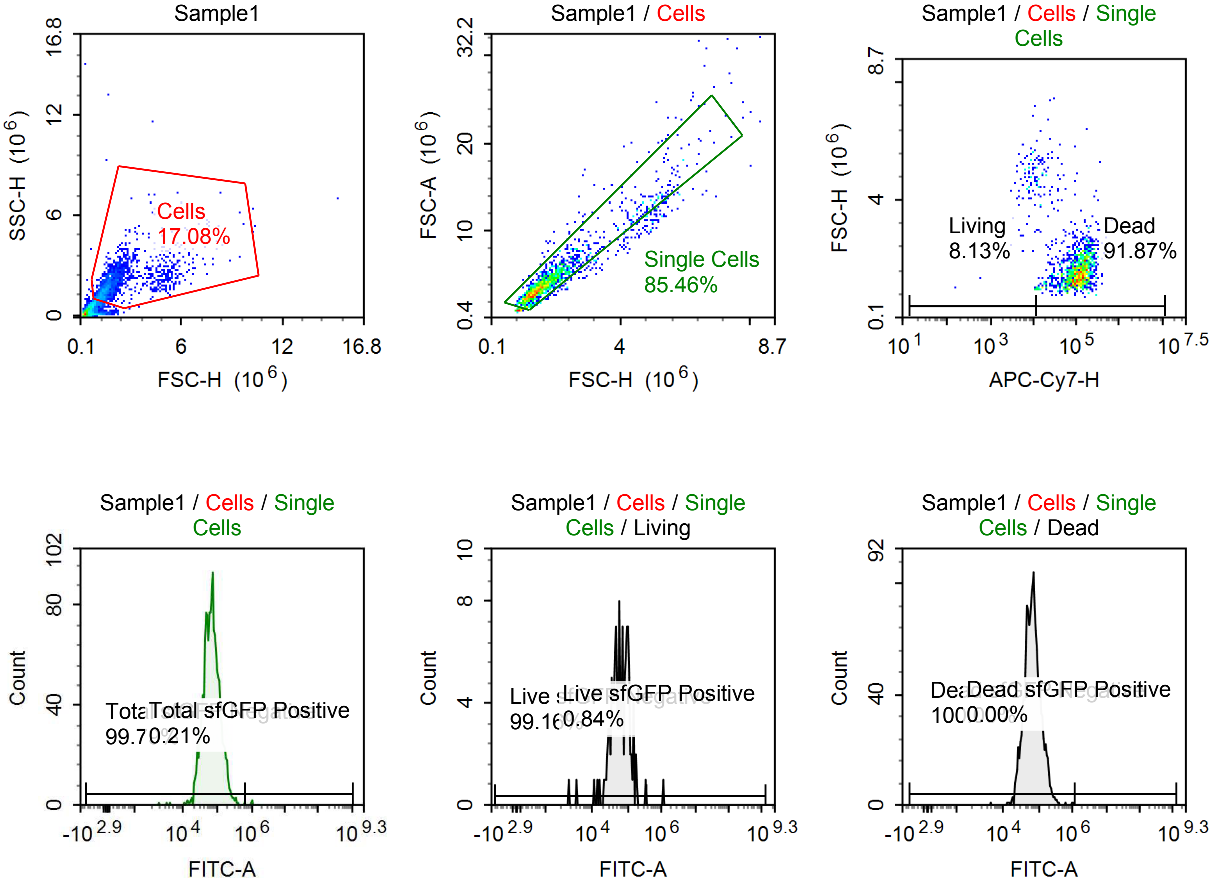

Sample Statistics of Sample1

| Gate | Count | % Parent | X | Y | Median X | Median Y |
|---|---|---|---|---|---|---|
| All | 10,023 | | | | | |
| Cells | 1,712 | 17.08 % | FSC-H | SSC-H | 1,631,826 | 1,784,774 |
| Single Cells | 1,463 | 85.46 % | FSC-H | FSC-A | 1,585,222 | 3,955,628 |
| Living | 119 | 8.13 % | APC-Cy7-H | FSC-H | 8,614 | 4,674,370 |
| Live sfGFP Negative | 118 | 99.16 % | FITC-A | Count | 60,331 | |
| Live sfGFP Positive | 1 | 0.84 % | FITC-A | Count | 1,099,257 | |
| Dead | 1,344 | 91.87 % | APC-Cy7-H | FSC-H | 110,592 | 1,551,354 |
| Dead sfGFP Negative | 1,344 | 100.00 % | FITC-A | Count | 65,806 | |
| Dead sfGFP Positive | 0 | 0.00 % | FITC-A | Count | 0 | |
| Total sfGFP Negative | 1,460 | 99.79 % | FITC-A | Count | 65,678 | |
| Total sfGFP Positive | 3 | 0.21 % | FITC-A | Count | 1,022,058 | |

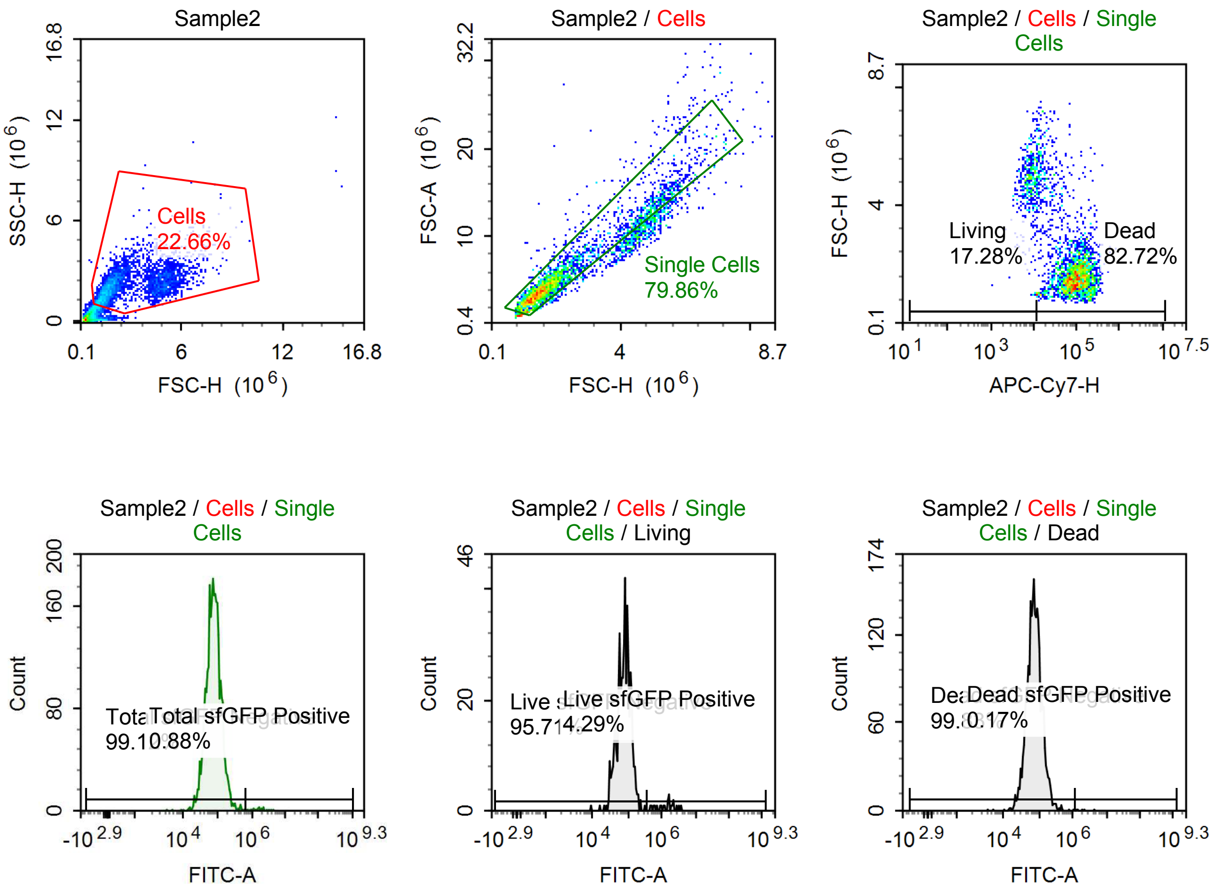

Sample Statistics of Sample2

| Gate | Count | % Parent | X | Y | Median X | Median Y |
|---|---|---|---|---|---|---|
| All | 15,663 | | | | | |
| Cells | 3,550 | 22.66 % | FSC-H | SSC-H | 1,990,749 | 2,078,174 |
| Single Cells | 2,835 | 79.86 % | FSC-H | FSC-A | 1,795,947 | 4,707,609 |
| Living | 490 | 17.28 % | APC-Cy7-H | FSC-H | 8,348 | 4,804,567 |
| Live sfGFP Negative | 469 | 95.71 % | FITC-A | Count | 80,213 | |
| Live sfGFP Positive | 21 | 4.29 % | FITC-A | Count | 1,535,116 | |
| Dead | 2,345 | 82.72 % | APC-Cy7-H | FSC-H | 88,162 | 1,640,282 |
| Dead sfGFP Negative | 2,341 | 99.83 % | FITC-A | Count | 73,567 | |
| Dead sfGFP Positive | 4 | 0.17 % | FITC-A | Count | 2,103,919 | |
| Total sfGFP Negative | 2,810 | 99.12 % | FITC-A | Count | 74,524 | |
| Total sfGFP Positive | 25 | 0.88 % | FITC-A | Count | 1,612,988 | |

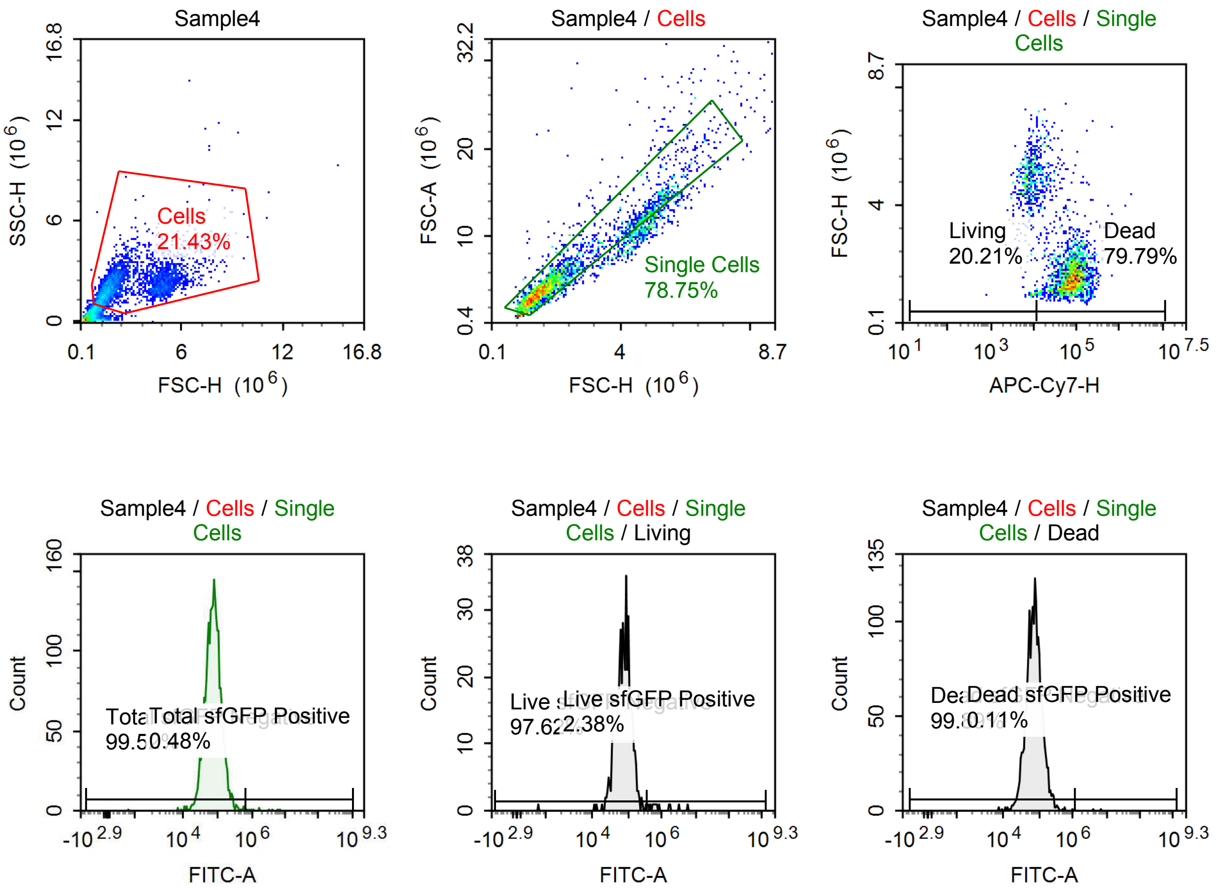

Sample Statistics of Sample4

| Gate | Count | % Parent | X | Y | Median X | Median Y |
|---|---|---|---|---|---|---|
| All | 13,546 | | | | | |
| Cells | 2,903 | 21.43 % | FSC-H | SSC-H | 2,040,724 | 2,075,279 |
| Single Cells | 2,286 | 78.75 % | FSC-H | FSC-A | 1,786,963 | 4,614,020 |
| Living | 462 | 20.21 % | APC-Cy7-H | FSC-H | 7,839 | 4,759,388 |
| Live sfGFP Negative | 451 | 97.62 % | FITC-A | Count | 75,758 | |
| Live sfGFP Positive | 11 | 2.38 % | FITC-A | Count | 719,378 | |
| Dead | 1,824 | 79.79 % | APC-Cy7-H | FSC-H | 74,710 | 1,614,778 |
| Dead sfGFP Negative | 1,822 | 99.89 % | FITC-A | Count | 73,102 | |
| Dead sfGFP Positive | 2 | 0.11 % | FITC-A | Count | 5,404,541 | |
| Total sfGFP Negative | 2,275 | 99.52 % | FITC-A | Count | 73,833 | |
| Total sfGFP Positive | 11 | 0.48 % | FITC-A | Count | 1,713,401 | |

#### p060 + 10 uM Tet

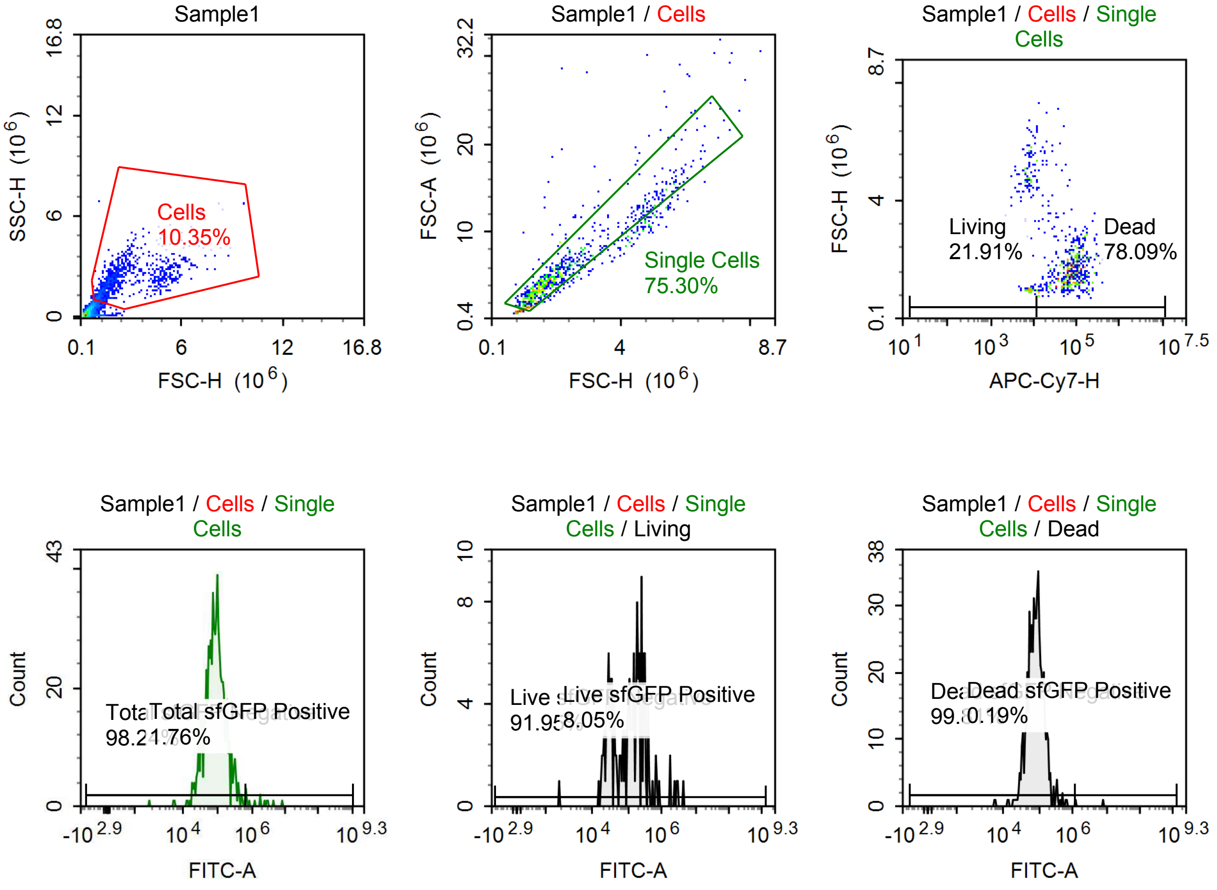

Sample Statistics of Sample1

| Gate | Count | % Parent | X | Y | Median X | Median Y |
|---|---|---|---|---|---|---|
| All | 8,728 | | | | | |
| Cells | 903 | 10.35 % | FSC-H | SSC-H | 1,880,545 | 2,127,820 |
| Single Cells | 680 | 75.30 % | FSC-H | FSC-A | 1,797,718 | 4,529,028 |
| Living | 149 | 21.91 % | APC-Cy7-H | FSC-H | 7,788 | 4,596,796 |
| Live sfGFP Negative | 137 | 91.95 % | FITC-A | Count | 109,670 | |
| Live sfGFP Positive | 12 | 8.05 % | FITC-A | Count | 1,325,858 | |
| Dead | 531 | 78.09 % | APC-Cy7-H | FSC-H | 72,861 | 1,666,990 |
| Dead sfGFP Negative | 530 | 99.81 % | FITC-A | Count | 80,712 | |
| Dead sfGFP Positive | 1 | 0.19 % | FITC-A | Count | 7,831,356 | |
| Total sfGFP Negative | 668 | 98.24 % | FITC-A | Count | 83,851 | |
| Total sfGFP Positive | 12 | 1.76 % | FITC-A | Count | 1,824,636 | |

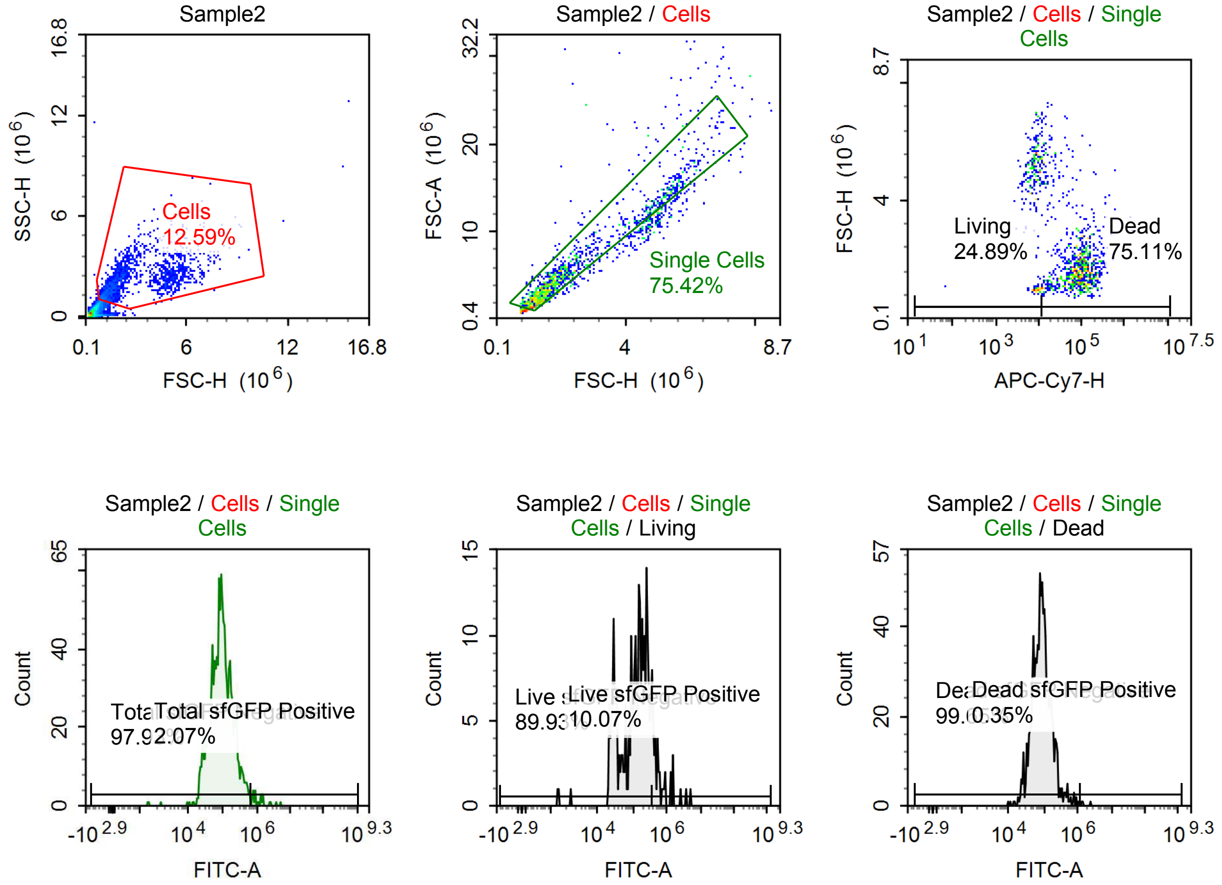

Sample Statistics of Sample2

| Gate | Count | % Parent | X | Y | Median X | Median Y |
|---|---|---|---|---|---|---|
| All | 12,180 | | | | | |
| Cells | 1,534 | 12.59 % | FSC-H | SSC-H | 2,064,887 | 2,118,915 |
| Single Cells | 1,157 | 75.42 % | FSC-H | FSC-A | 1,899,547 | 4,931,681 |
| Living | 288 | 24.89 % | APC-Cy7-H | FSC-H | 8,181 | 4,916,010 |
| Live sfGFP Negative | 259 | 89.93 % | FITC-A | Count | 131,839 | |
| Live sfGFP Positive | 29 | 10.07 % | FITC-A | Count | 620,785 | |
| Dead | 869 | 75.11 % | APC-Cy7-H | FSC-H | 83,256 | 1,704,689 |
| Dead sfGFP Negative | 866 | 99.65 % | FITC-A | Count | 85,591 | |
| Dead sfGFP Positive | 3 | 0.35 % | FITC-A | Count | 1,422,933 | |
| Total sfGFP Negative | 1,133 | 97.93 % | FITC-A | Count | 90,769 | |
| Total sfGFP Positive | 24 | 2.07 % | FITC-A | Count | 1,092,603 | |

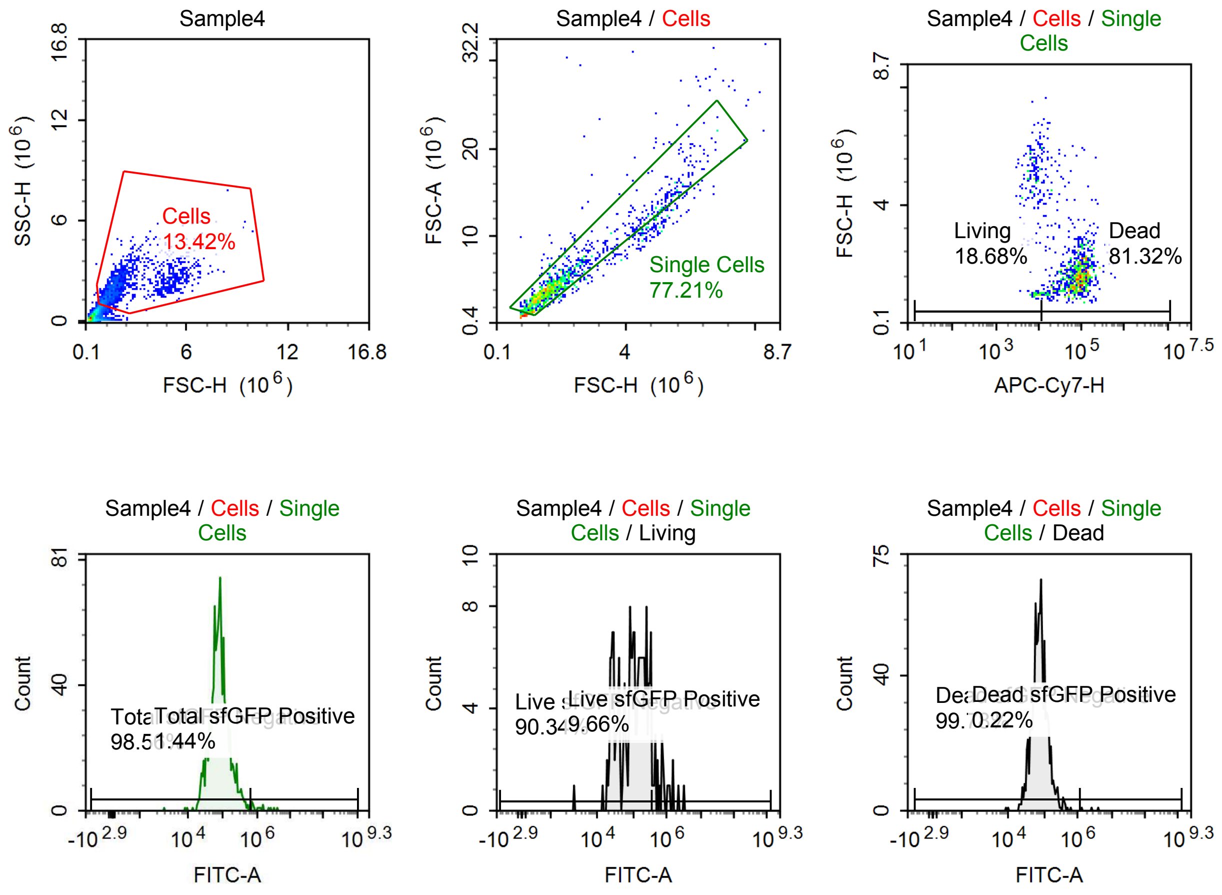

Sample Statistics of Sample4

| Gate | Count | % Parent | X | Y | Median X | Median Y |
|---|---|---|---|---|---|---|
| All | 10,696 | | | | | |
| Cells | 1,435 | 13.42 % | FSC-H | SSC-H | 1,757,019 | 2,024,190 |
| Single Cells | 1,108 | 77.21 % | FSC-H | FSC-A | 1,722,112 | 4,373,614 |
| Living | 207 | 18.68 % | APC-Cy7-H | FSC-H | 7,853 | 4,812,216 |
| Live sfGFP Negative | 187 | 90.34 % | FITC-A | Count | 101,248 | |
| Live sfGFP Positive | 20 | 9.66 % | FITC-A | Count | 753,448 | |
| Dead | 901 | 81.32 % | APC-Cy7-H | FSC-H | 80,289 | 1,624,786 |
| Dead sfGFP Negative | 899 | 99.78 % | FITC-A | Count | 75,549 | |
| Dead sfGFP Positive | 2 | 0.22 % | FITC-A | Count | 3,300,210 | |
| Total sfGFP Negative | 1,092 | 98.56 % | FITC-A | Count | 77,142 | |
| Total sfGFP Positive | 16 | 1.44 % | FITC-A | Count | 1,174,424 | |

#### p060 + 100 uM Tet

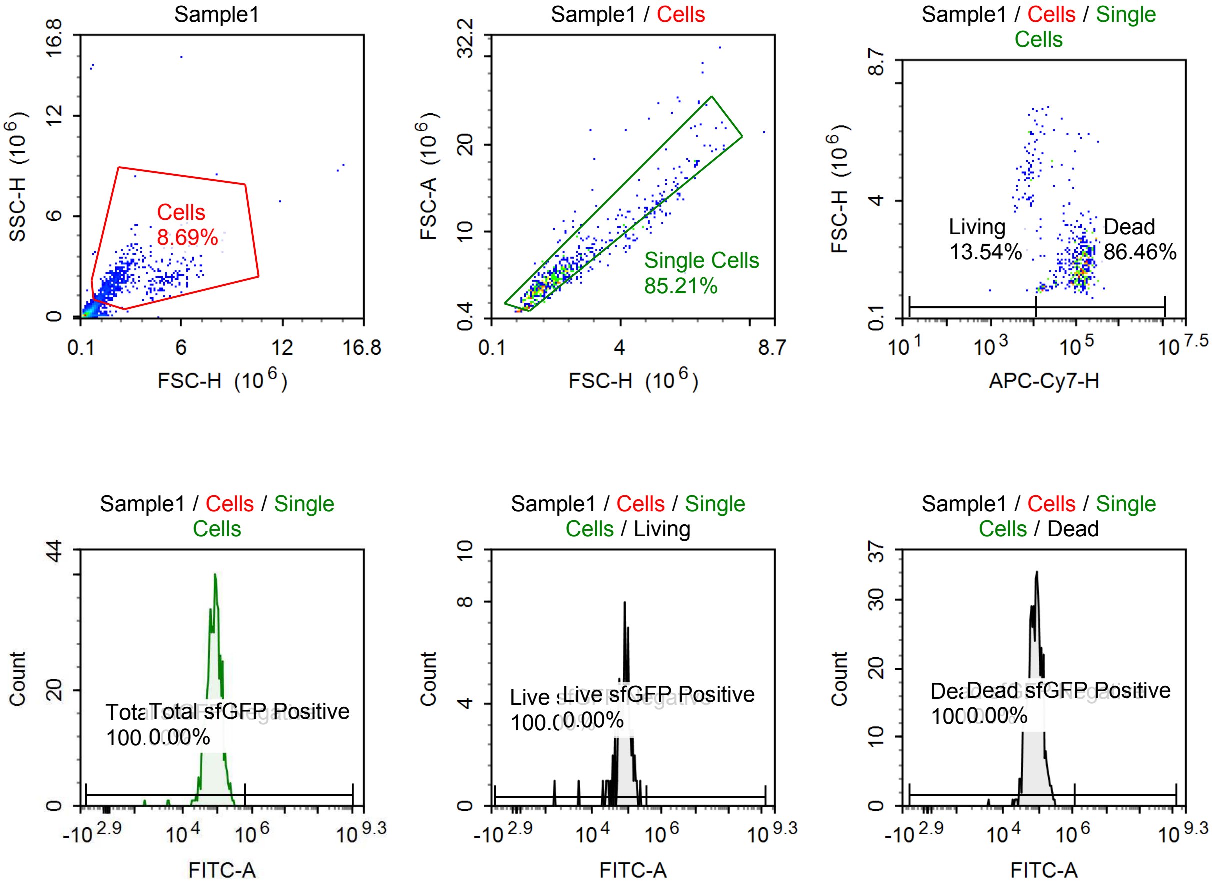

Sample Statistics of Sample1

| Gate | Count | % Parent | X | Y | Median X | Median Y |
|---|---|---|---|---|---|---|
| All | 7,781 | | | | | |
| Cells | 676 | 8.69 % | FSC-H | SSC-H | 2,003,969 | 2,023,233 |
| Single Cells | 576 | 85.21 % | FSC-H | FSC-A | 1,934,191 | 4,853,801 |
| Living | 78 | 13.54 % | APC-Cy7-H | FSC-H | 7,874 | 4,861,067 |
| Live sfGFP Negative | 78 | 100.00 % | FITC-A | Count | 81,394 | |
| Live sfGFP Positive | 0 | 0.00 % | FITC-A | Count | 0 | |
| Dead | 498 | 86.46 % | APC-Cy7-H | FSC-H | 113,575 | 1,792,579 |
| Dead sfGFP Negative | 498 | 100.00 % | FITC-A | Count | 79,894 | |
| Dead sfGFP Positive | 0 | 0.00 % | FITC-A | Count | 0 | |
| Total sfGFP Negative | 576 | 100.00 % | FITC-A | Count | 80,054 | |
| Total sfGFP Positive | 0 | 0.00 % | FITC-A | Count | 0 | |

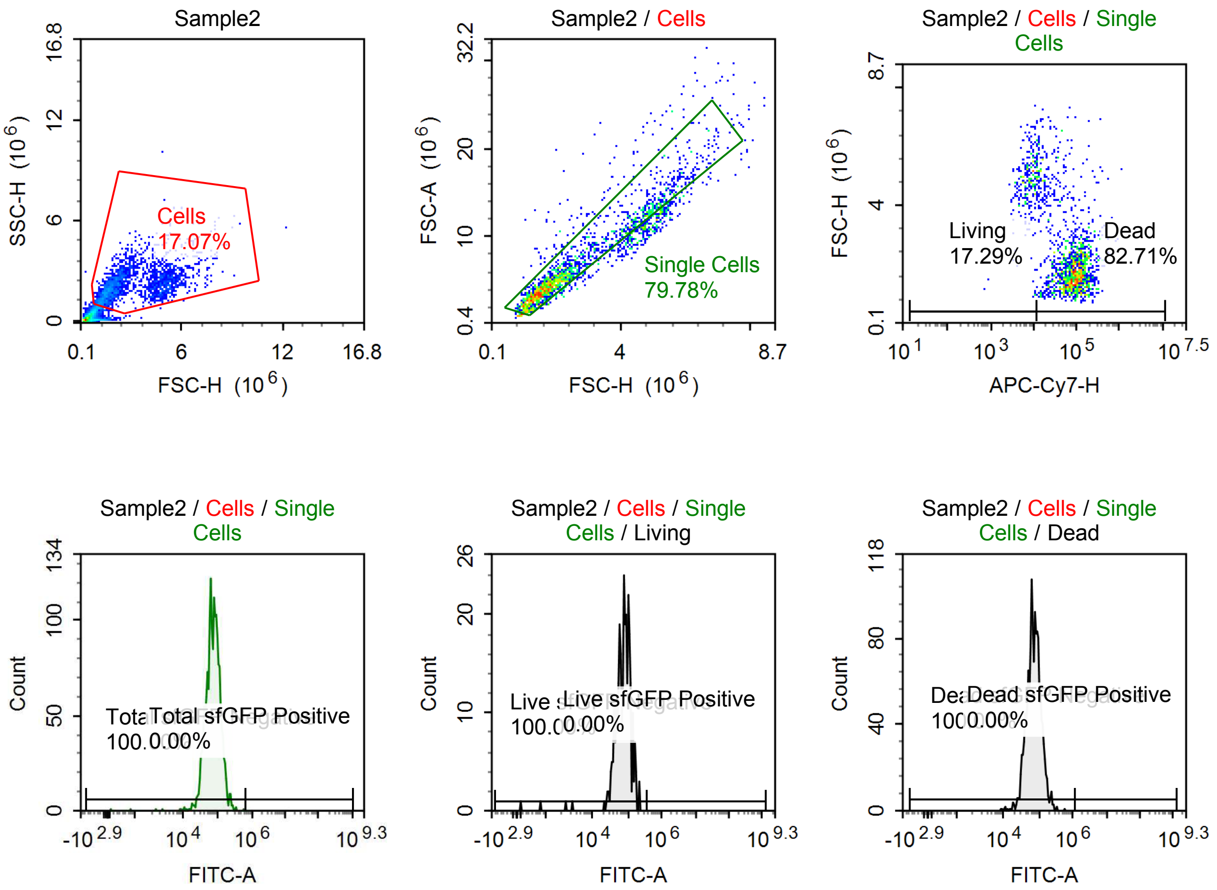

Sample Statistics of Sample2

| Gate | Count | % Parent | X | Y | Median X | Median Y |
|---|---|---|---|---|---|---|
| All | 12,316 | | | | | |
| Cells | 2,102 | 17.07 % | FSC-H | SSC-H | 2,311,630 | 2,131,377 |
| Single Cells | 1,677 | 79.78 % | FSC-H | FSC-A | 2,052,900 | 5,228,795 |
| Living | 290 | 17.29 % | APC-Cy7-H | FSC-H | 7,786 | 4,838,998 |
| Live sfGFP Negative | 290 | 100.00 % | FITC-A | Count | 78,105 | |
| Live sfGFP Positive | 0 | 0.00 % | FITC-A | Count | 0 | |
| Dead | 1,387 | 82.71 % | APC-Cy7-H | FSC-H | 81,766 | 1,804,431 |
| Dead sfGFP Negative | 1,387 | 100.00 % | FITC-A | Count | 71,372 | |
| Dead sfGFP Positive | 0 | 0.00 % | FITC-A | Count | 0 | |
| Total sfGFP Negative | 1,677 | 100.00 % | FITC-A | Count | 72,918 | |
| Total sfGFP Positive | 0 | 0.00 % | FITC-A | Count | 0 | |

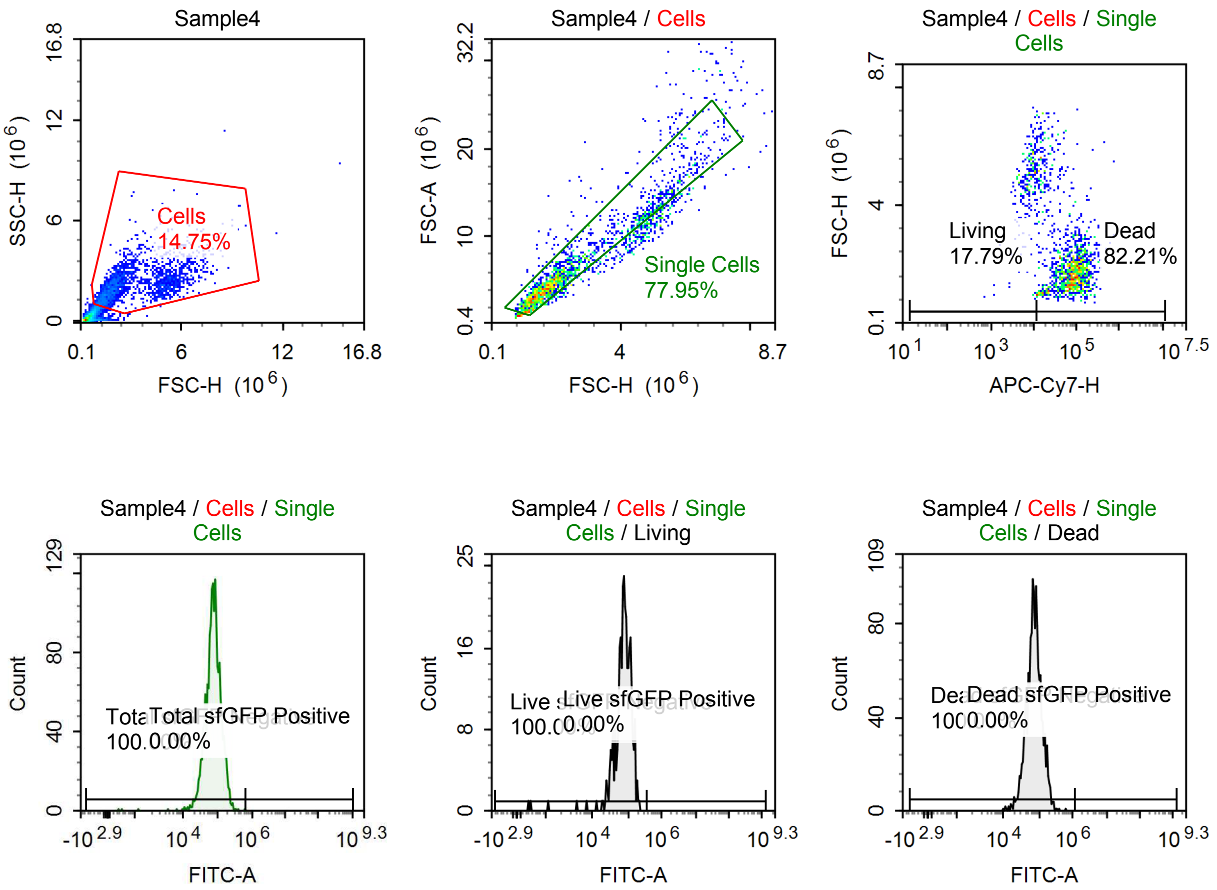

Sample Statistics of Sample4

| Gate | Count | % Parent | X | Y | Median X | Median Y |
|---|---|---|---|---|---|---|
| All | 14,664 | | | | | |
| Cells | 2,163 | 14.75 % | FSC-H | SSC-H | 2,223,460 | 2,121,176 |
| Single Cells | 1,686 | 77.95 % | FSC-H | FSC-A | 1,983,028 | 5,178,644 |
| Living | 300 | 17.79 % | APC-Cy7-H | FSC-H | 8,190 | 4,849,035 |
| Live sfGFP Negative | 300 | 100.00 % | FITC-A | Count | 78,541 | |
| Live sfGFP Positive | 0 | 0.00 % | FITC-A | Count | 0 | |
| Dead | 1,386 | 82.21 % | APC-Cy7-H | FSC-H | 75,209 | 1,829,643 |
| Dead sfGFP Negative | 1,386 | 100.00 % | FITC-A | Count | 73,289 | |
| Dead sfGFP Positive | 0 | 0.00 % | FITC-A | Count | 0 | |
| Total sfGFP Negative | 1,686 | 100.00 % | FITC-A | Count | 74,108 | |
| Total sfGFP Positive | 0 | 0.00 % | FITC-A | Count | 0 | |

## p060

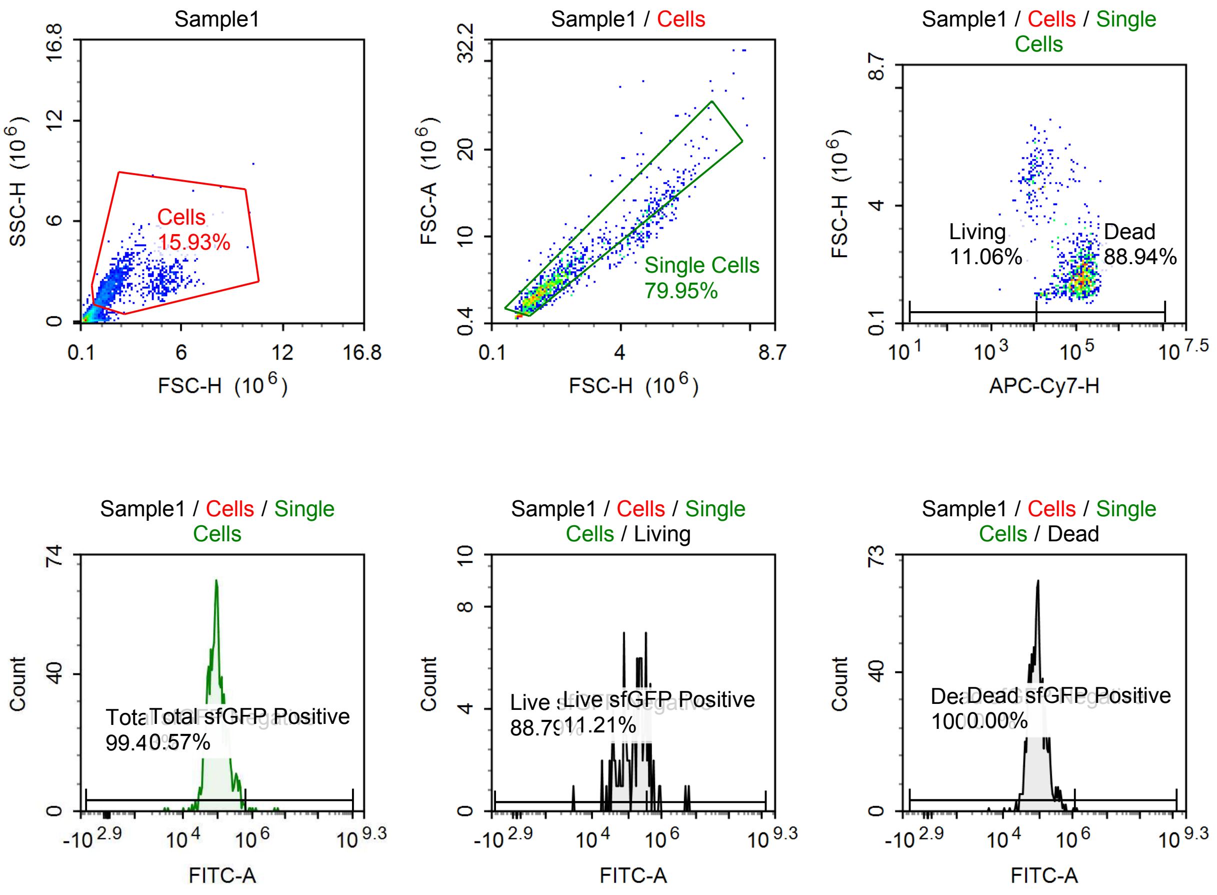

Sample Statistics of Sample1

| Gate | Count | % Parent | X | Y | Median X | Median Y |
|---|---|---|---|---|---|---|
| All | 8,234 | | | | | |
| Cells | 1,312 | 15.93 % | FSC-H | SSC-H | 1,794,108 | 2,060,895 |
| Single Cells | 1,049 | 79.95 % | FSC-H | FSC-A | 1,716,934 | 4,441,339 |
| Living | 116 | 11.06 % | APC-Cy7-H | FSC-H | 8,514 | 4,858,272 |
| Live sfGFP Negative | 103 | 88.79 % | FITC-A | Count | 159,451 | |
| Live sfGFP Positive | 13 | 11.21 % | FITC-A | Count | 533,810 | |
| Dead | 933 | 88.94 % | APC-Cy7-H | FSC-H | 115,860 | 1,640,380 |
| Dead sfGFP Negative | 933 | 100.00 % | FITC-A | Count | 88,472 | |
| Dead sfGFP Positive | 0 | 0.00 % | FITC-A | Count | 0 | |
| Total sfGFP Negative | 1,043 | 99.43 % | FITC-A | Count | 90,072 | |
| Total sfGFP Positive | 6 | 0.57 % | FITC-A | Count | 1,163,379 | |

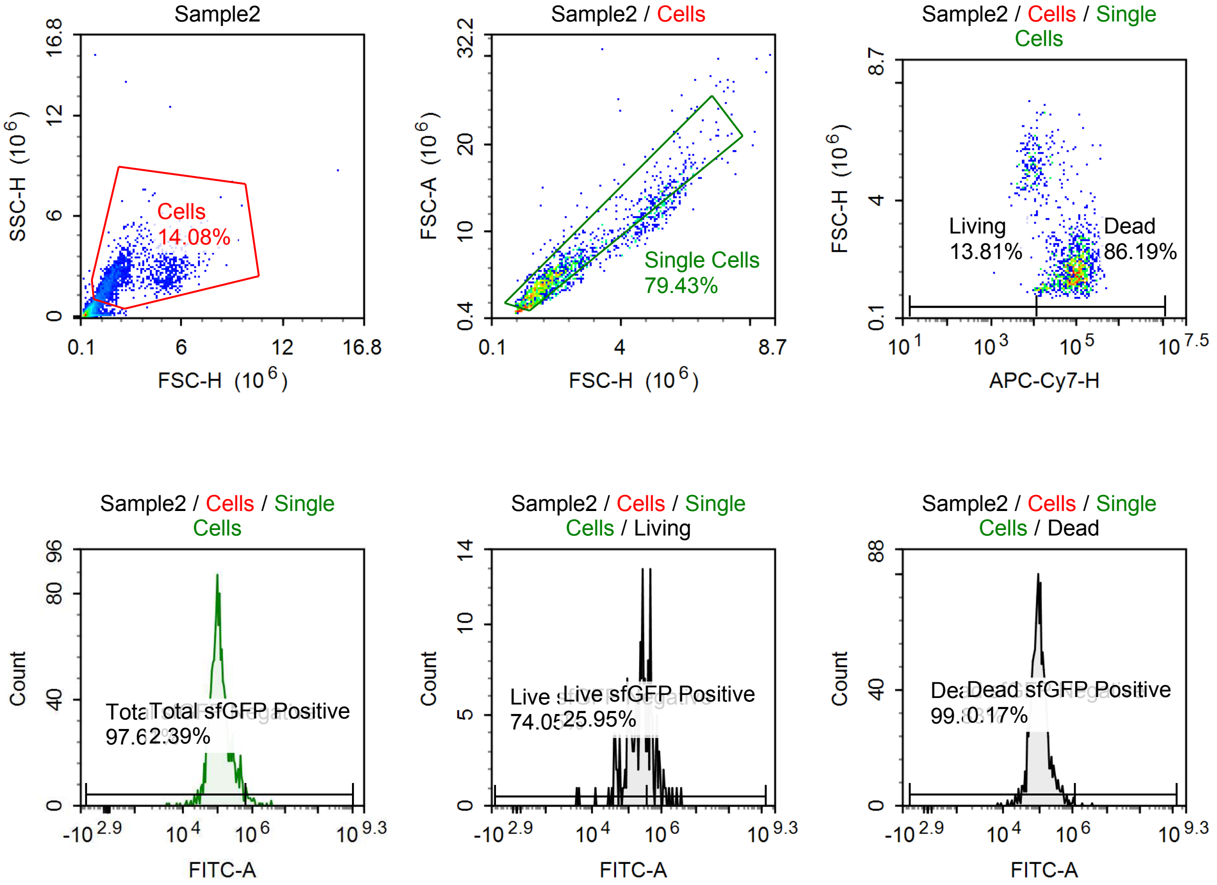

Sample Statistics of Sample2

| Gate | Count | % Parent | X | Y | Median X | Median Y |
|---|---|---|---|---|---|---|
| All | 11,981 | | | | | |
| Cells | 1,687 | 14.08 % | FSC-H | SSC-H | 1,926,733 | 2,188,223 |
| Single Cells | 1,340 | 79.43 % | FSC-H | FSC-A | 1,810,265 | 4,705,097 |
| Living | 185 | 13.81 % | APC-Cy7-H | FSC-H | 7,933 | 4,993,088 |
| Live sfGFP Negative | 137 | 74.05 % | FITC-A | Count | 184,027 | |
| Live sfGFP Positive | 48 | 25.95 % | FITC-A | Count | 553,077 | |
| Dead | 1,155 | 86.19 % | APC-Cy7-H | FSC-H | 85,140 | 1,697,143 |
| Dead sfGFP Negative | 1,153 | 99.83 % | FITC-A | Count | 99,543 | |
| Dead sfGFP Positive | 2 | 0.17 % | FITC-A | Count | 2,777,240 | |
| Total sfGFP Negative | 1,308 | 97.61 % | FITC-A | Count | 103,677 | |
| Total sfGFP Positive | 32 | 2.39 % | FITC-A | Count | 934,167 | |

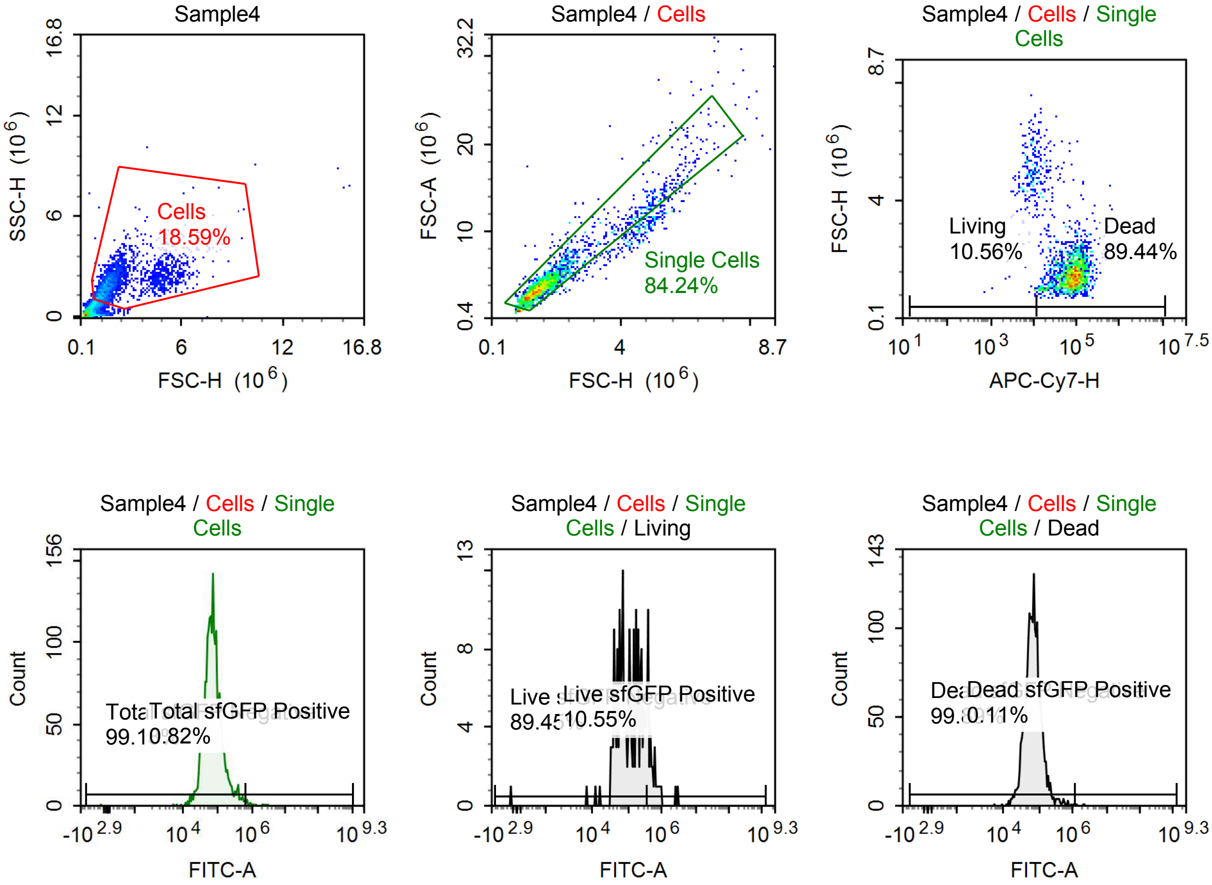

Sample Statistics of Sample4

| Gate | Count | % Parent | X | Y | Median X | Median Y |
|---|---|---|---|---|---|---|
| All | 13,177 | | | | | |
| Cells | 2,450 | 18.59 % | FSC-H | SSC-H | 1,617,012 | 1,891,219 |
| Single Cells | 2,064 | 84.24 % | FSC-H | FSC-A | 1,574,978 | 3,939,061 |
| Living | 218 | 10.56 % | APC-Cy7-H | FSC-H | 8,314 | 4,766,979 |
| Live sfGFP Negative | 195 | 89.45 % | FITC-A | Count | 109,663 | |
| Live sfGFP Positive | 23 | 10.55 % | FITC-A | Count | 582,841 | |
| Dead | 1,846 | 89.44 % | APC-Cy7-H | FSC-H | 81,220 | 1,514,203 |
| Dead sfGFP Negative | 1,844 | 99.89 % | FITC-A | Count | 67,304 | |
| Dead sfGFP Positive | 2 | 0.11 % | FITC-A | Count | 2,149,488 | |
| Total sfGFP Negative | 2,047 | 99.18 % | FITC-A | Count | 69,455 | |
| Total sfGFP Positive | 17 | 0.82 % | FITC-A | Count | 904,468 | |

#### p060 + 1 uM Tet

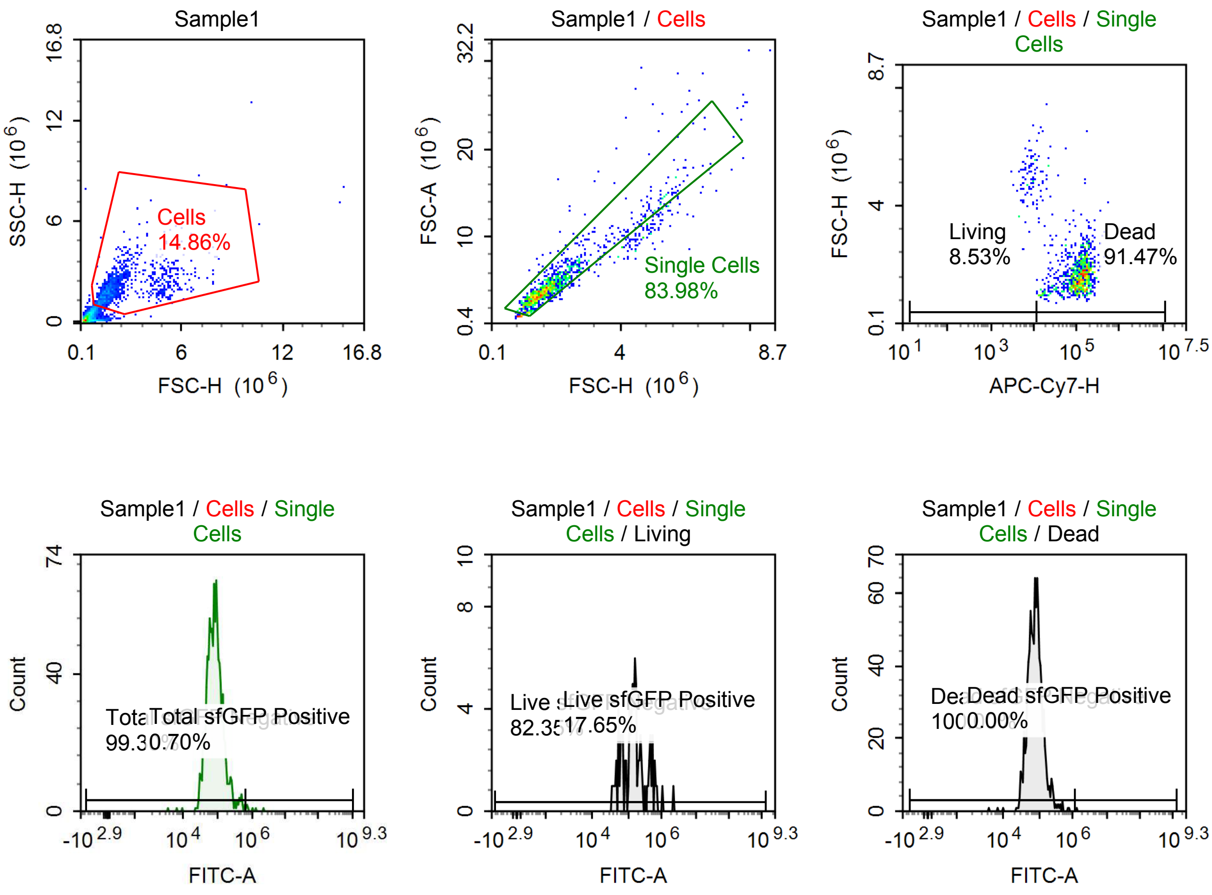

Sample Statistics of Sample1

| Gate | Count | % Parent | X | Y | Median X | Median Y |
|---|---|---|---|---|---|---|
| All | 7,980 | | | | | |
| Cells | 1,186 | 14.86 % | FSC-H | SSC-H | 1,759,862 | 1,917,813 |
| Single Cells | 996 | 83.98 % | FSC-H | FSC-A | 1,686,985 | 4,177,923 |
| Living | 85 | 8.53 % | APC-Cy7-H | FSC-H | 7,884 | 4,860,701 |
| Live sfGFP Negative | 70 | 82.35 % | FITC-A | Count | 133,074 | |
| Live sfGFP Positive | 15 | 17.65 % | FITC-A | Count | 552,977 | |
| Dead | 911 | 91.47 % | APC-Cy7-H | FSC-H | 117,860 | 1,637,369 |
| Dead sfGFP Negative | 911 | 100.00 % | FITC-A | Count | 78,140 | |
| Dead sfGFP Positive | 0 | 0.00 % | FITC-A | Count | 0 | |
| Total sfGFP Negative | 989 | 99.30 % | FITC-A | Count | 80,036 | |
| Total sfGFP Positive | 7 | 0.70 % | FITC-A | Count | 792,001 | |

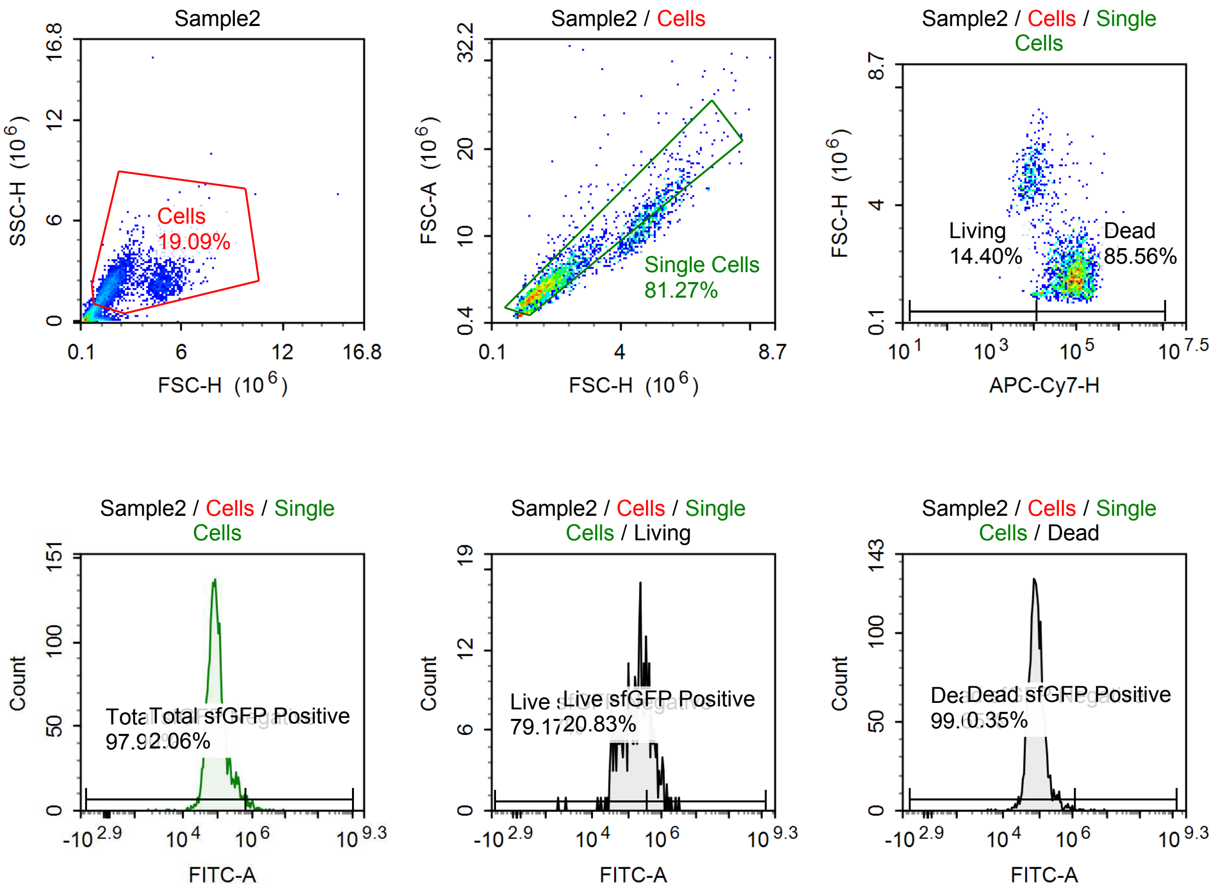

Sample Statistics of Sample2

| Gate | Count | % Parent | X | Y | Median X | Median Y |
|---|---|---|---|---|---|---|
| All | 15,041 | | | | | |
| Cells | 2,872 | 19.09 % | FSC-H | SSC-H | 1,886,985 | 2,110,229 |
| Single Cells | 2,334 | 81.27 % | FSC-H | FSC-A | 1,775,423 | 4,631,396 |
| Living | 336 | 14.40 % | APC-Cy7-H | FSC-H | 7,882 | 4,803,772 |
| Live sfGFP Negative | 266 | 79.17 % | FITC-A | Count | 147,513 | |
| Live sfGFP Positive | 70 | 20.83 % | FITC-A | Count | 600,683 | |
| Dead | 1,997 | 85.56 % | APC-Cy7-H | FSC-H | 81,506 | 1,663,843 |
| Dead sfGFP Negative | 1,990 | 99.65 % | FITC-A | Count | 82,526 | |
| Dead sfGFP Positive | 7 | 0.35 % | FITC-A | Count | 2,112,077 | |
| Total sfGFP Negative | 2,285 | 97.90 % | FITC-A | Count | 85,400 | |
| Total sfGFP Positive | 48 | 2.06 % | FITC-A | Count | 1,009,446 | |

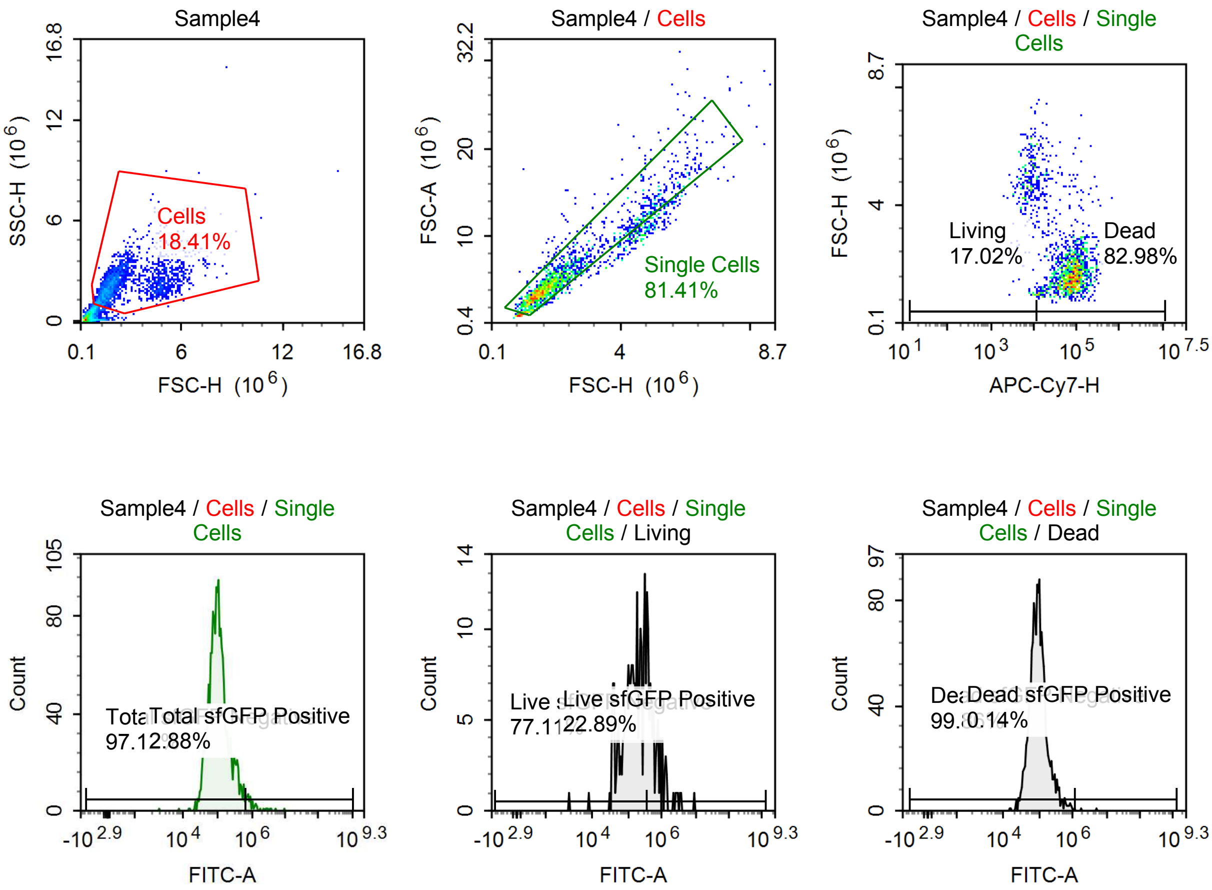

Sample Statistics of Sample4

| Gate | Count | % Parent | X | Y | Median X | Median Y |
|---|---|---|---|---|---|---|
| All | 11,135 | | | | | |
| Cells | 2,050 | 18.41 % | FSC-H | SSC-H | 1,911,459 | 2,133,912 |
| Single Cells | 1,669 | 81.41 % | FSC-H | FSC-A | 1,819,484 | 4,664,995 |
| Living | 284 | 17.02 % | APC-Cy7-H | FSC-H | 8,096 | 4,697,458 |
| Live sfGFP Negative | 219 | 77.11 % | FITC-A | Count | 153,527 | |
| Live sfGFP Positive | 65 | 22.89 % | FITC-A | Count | 688,639 | |
| Dead | 1,385 | 82.98 % | APC-Cy7-H | FSC-H | 75,231 | 1,693,734 |
| Dead sfGFP Negative | 1,383 | 99.86 % | FITC-A | Count | 93,782 | |
| Dead sfGFP Positive | 2 | 0.14 % | FITC-A | Count | 3,501,713 | |
| Total sfGFP Negative | 1,621 | 97.12 % | FITC-A | Count | 98,551 | |
| Total sfGFP Positive | 48 | 2.88 % | FITC-A | Count | 918,523 | |

#### p059 + 10 uM Tet

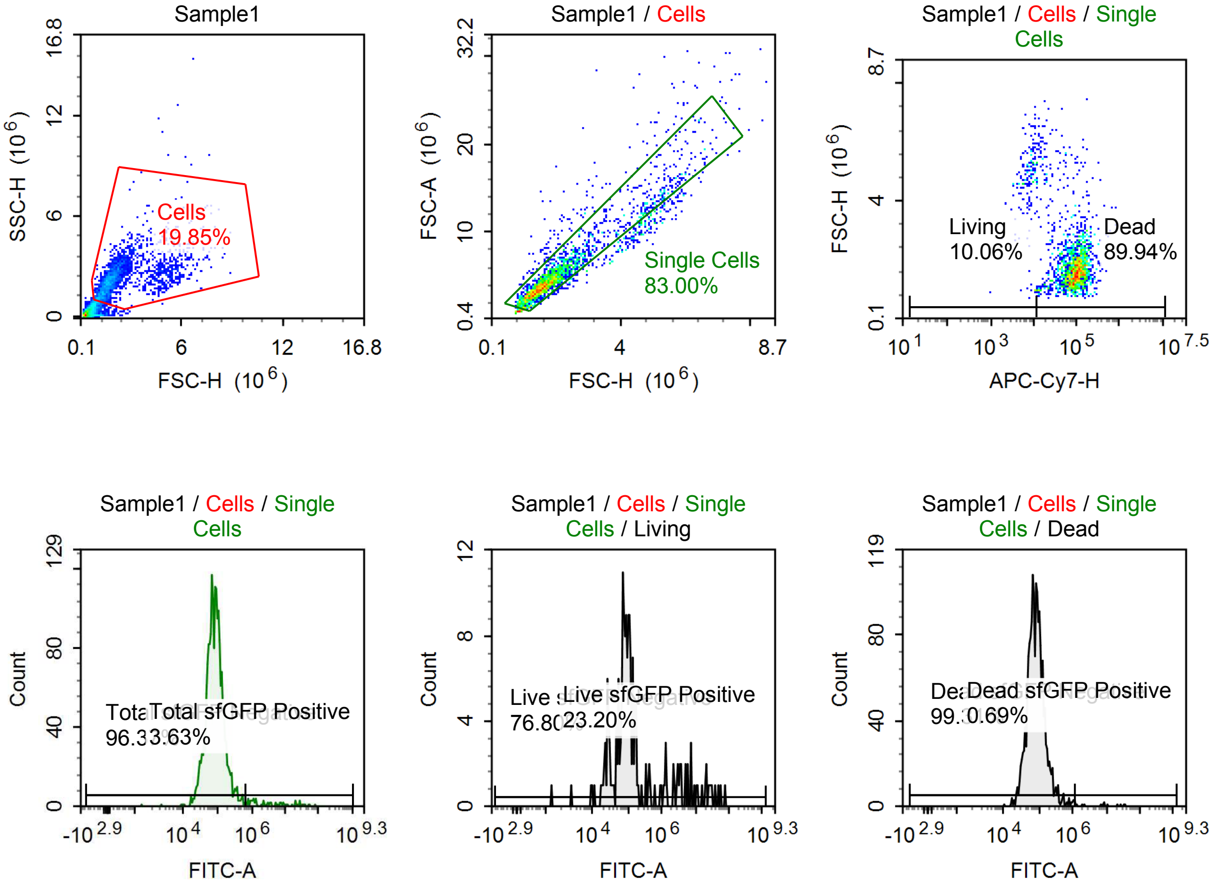

Sample Statistics of Sample1

| Gate | Count | % Parent | X | Y | Median X | Median Y |
|---|---|---|---|---|---|---|
| All | 11,705 | | | | | |
| Cells | 2,323 | 19.85 % | FSC-H | SSC-H | 1,835,132 | 2,249,901 |
| Single Cells | 1,928 | 83.00 % | FSC-H | FSC-A | 1,742,164 | 4,423,712 |
| Living | 194 | 10.06 % | APC-Cy7-H | FSC-H | 8,441 | 4,761,743 |
| Live sfGFP Negative | 149 | 76.80 % | FITC-A | Count | 82,220 | |
| Live sfGFP Positive | 45 | 23.20 % | FITC-A | Count | 3,803,483 | |
| Dead | 1,734 | 89.94 % | APC-Cy7-H | FSC-H | 92,127 | 1,671,618 |
| Dead sfGFP Negative | 1,722 | 99.31 % | FITC-A | Count | 81,315 | |
| Dead sfGFP Positive | 12 | 0.69 % | FITC-A | Count | 8,665,620 | |
| Total sfGFP Negative | 1,858 | 96.37 % | FITC-A | Count | 81,070 | |
| Total sfGFP Positive | 70 | 3.63 % | FITC-A | Count | 3,354,498 | |

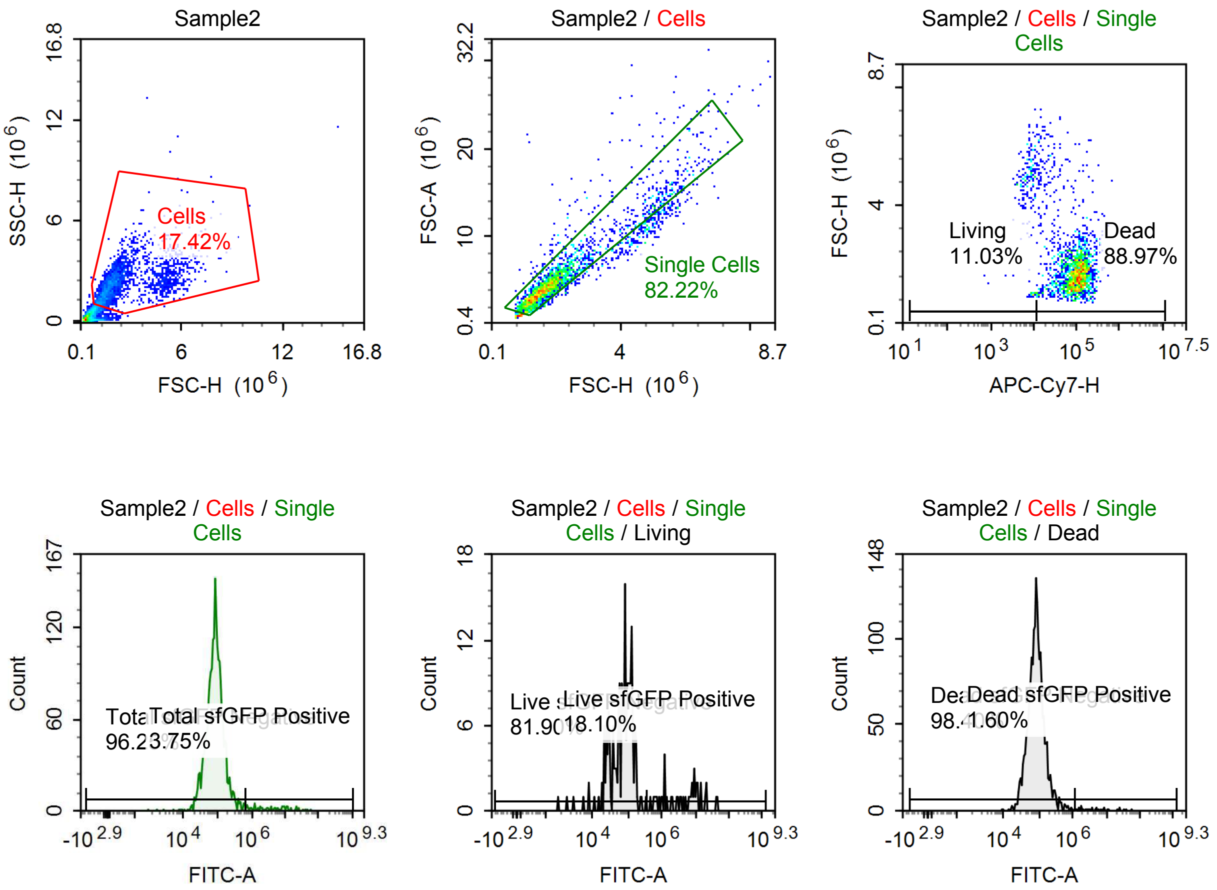

Sample Statistics of Sample2

| Gate | Count | % Parent | X | Y | Median X | Median Y |
|---|---|---|---|---|---|---|
| All | 14,693 | | | | | |
| Cells | 2,559 | 17.42 % | FSC-H | SSC-H | 1,840,452 | 2,179,900 |
| Single Cells | 2,104 | 82.22 % | FSC-H | FSC-A | 1,775,472 | 4,494,791 |
| Living | 232 | 11.03 % | APC-Cy7-H | FSC-H | 8,146 | 4,798,404 |
| Live sfGFP Negative | 190 | 81.90 % | FITC-A | Count | 77,460 | |
| Live sfGFP Positive | 42 | 18.10 % | FITC-A | Count | 5,694,399 | |
| Dead | 1,872 | 88.97 % | APC-Cy7-H | FSC-H | 98,844 | 1,715,235 |
| Dead sfGFP Negative | 1,842 | 98.40 % | FITC-A | Count | 83,768 | |
| Dead sfGFP Positive | 30 | 1.60 % | FITC-A | Count | 4,840,047 | |
| Total sfGFP Negative | 2,025 | 96.25 % | FITC-A | Count | 83,235 | |
| Total sfGFP Positive | 79 | 3.75 % | FITC-A | Count | 3,744,668 | |

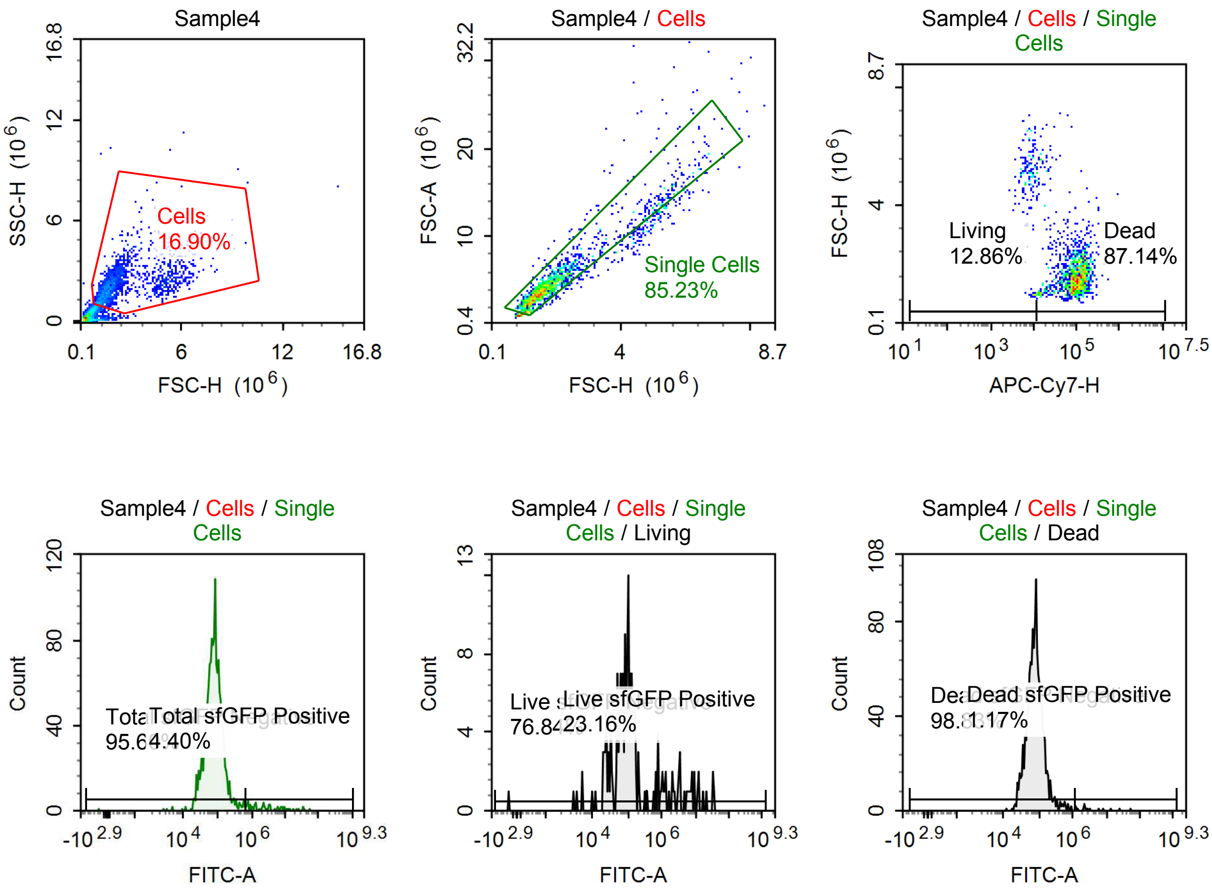

Sample Statistics of Sample4

| Gate | Count | % Parent | X | Y | Median X | Median Y |
|---|---|---|---|---|---|---|
| All | 10,255 | | | | | |
| Cells | 1,733 | 16.90 % | FSC-H | SSC-H | 1,723,507 | 2,060,354 |
| Single Cells | 1,477 | 85.23 % | FSC-H | FSC-A | 1,653,884 | 4,242,349 |
| Living | 190 | 12.86 % | APC-Cy7-H | FSC-H | 8,050 | 5,006,651 |
| Live sfGFP Negative | 146 | 76.84 % | FITC-A | Count | 80,617 | |
| Live sfGFP Positive | 44 | 23.16 % | FITC-A | Count | 3,273,657 | |
| Dead | 1,287 | 87.14 % | APC-Cy7-H | FSC-H | 96,832 | 1,584,491 |
| Dead sfGFP Negative | 1,272 | 98.83 % | FITC-A | Count | 77,359 | |
| Dead sfGFP Positive | 15 | 1.17 % | FITC-A | Count | 2,689,206 | |
| Total sfGFP Negative | 1,412 | 95.60 % | FITC-A | Count | 77,327 | |
| Total sfGFP Positive | 65 | 4.40 % | FITC-A | Count | 2,455,250 | |

#### p059 + 100 uM Tet

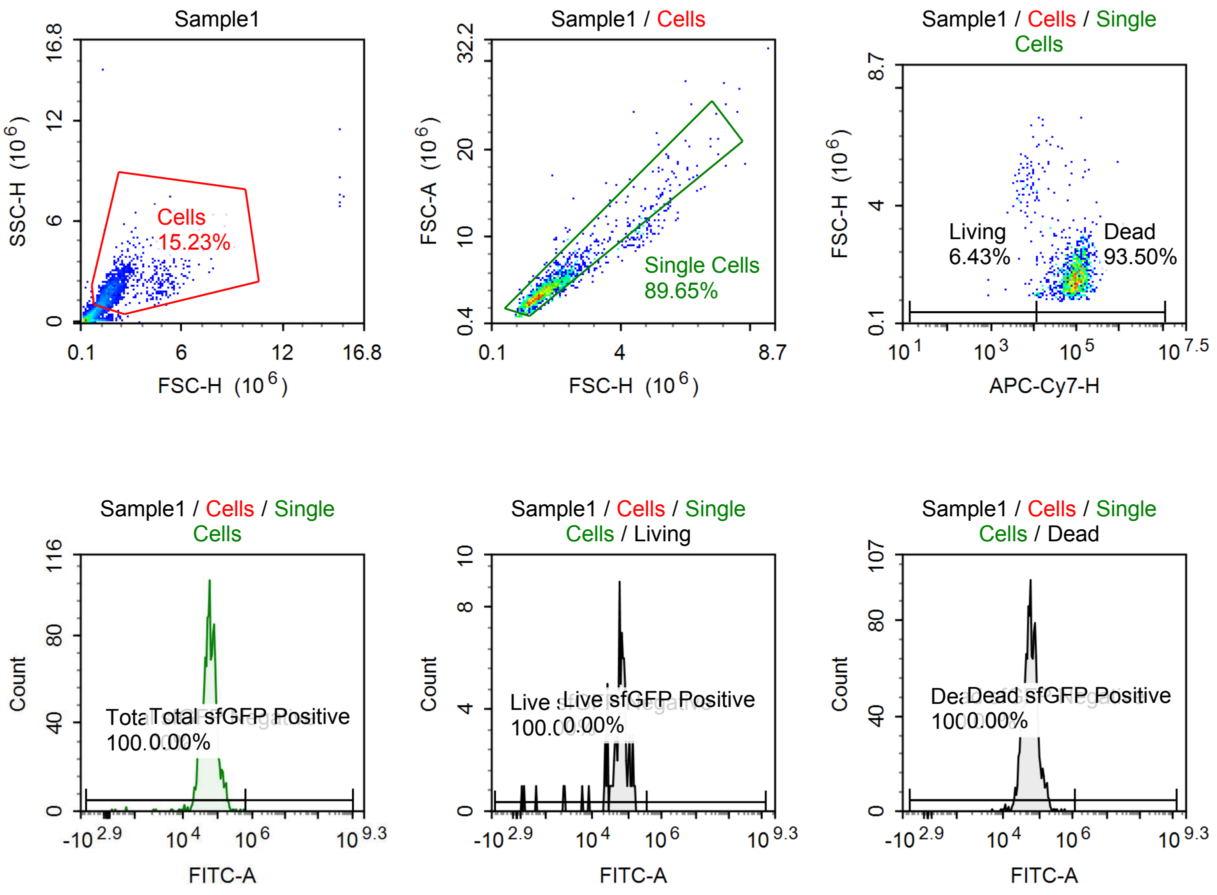

Sample Statistics of Sample1

| Gate | Count | % Parent | X | Y | Median X | Median Y |
|---|---|---|---|---|---|---|
| All | 10,027 | | | | | |
| Cells | 1,527 | 15.23 % | FSC-H | SSC-H | 1,695,626 | 1,781,595 |
| Single Cells | 1,369 | 89.65 % | FSC-H | FSC-A | 1,648,450 | 4,001,227 |
| Living | 88 | 6.43 % | APC-Cy7-H | FSC-H | 6,909 | 4,386,665 |
| Live sfGFP Negative | 88 | 100.00 % | FITC-A | Count | 60,112 | |
| Live sfGFP Positive | 0 | 0.00 % | FITC-A | Count | 0 | |
| Dead | 1,280 | 93.50 % | APC-Cy7-H | FSC-H | 93,905 | 1,612,458 |
| Dead sfGFP Negative | 1,280 | 100.00 % | FITC-A | Count | 56,962 | |
| Dead sfGFP Positive | 0 | 0.00 % | FITC-A | Count | 0 | |
| Total sfGFP Negative | 1,369 | 100.00 % | FITC-A | Count | 57,079 | |
| Total sfGFP Positive | 0 | 0.00 % | FITC-A | Count | 0 | |

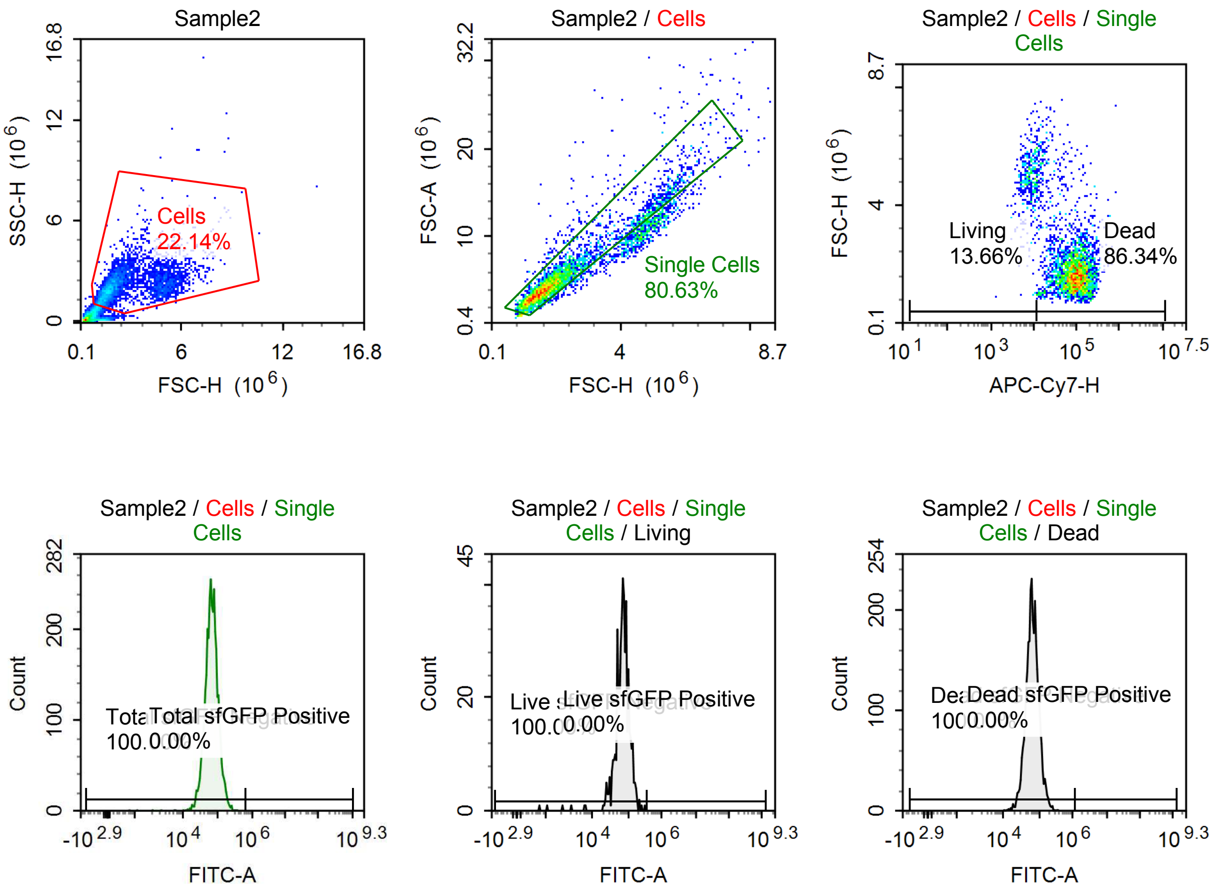

Sample Statistics of Sample2

| Gate | Count | % Parent | X | Y | Median X | Median Y |
|---|---|---|---|---|---|---|
| All | 18,742 | | | | | |
| Cells | 4,150 | 22.14 % | FSC-H | SSC-H | 2,024,790 | 2,063,361 |
| Single Cells | 3,346 | 80.63 % | FSC-H | FSC-A | 1,831,385 | 4,655,524 |
| Living | 457 | 13.66 % | APC-Cy7-H | FSC-H | 8,030 | 4,853,820 |
| Live sfGFP Negative | 457 | 100.00 % | FITC-A | Count | 72,690 | |
| Live sfGFP Positive | 0 | 0.00 % | FITC-A | Count | 0 | |
| Dead | 2,889 | 86.34 % | APC-Cy7-H | FSC-H | 88,167 | 1,712,040 |
| Dead sfGFP Negative | 2,889 | 100.00 % | FITC-A | Count | 64,176 | |
| Dead sfGFP Positive | 0 | 0.00 % | FITC-A | Count | 0 | |
| Total sfGFP Negative | 3,346 | 100.00 % | FITC-A | Count | 65,517 | |
| Total sfGFP Positive | 0 | 0.00 % | FITC-A | Count | 0 | |

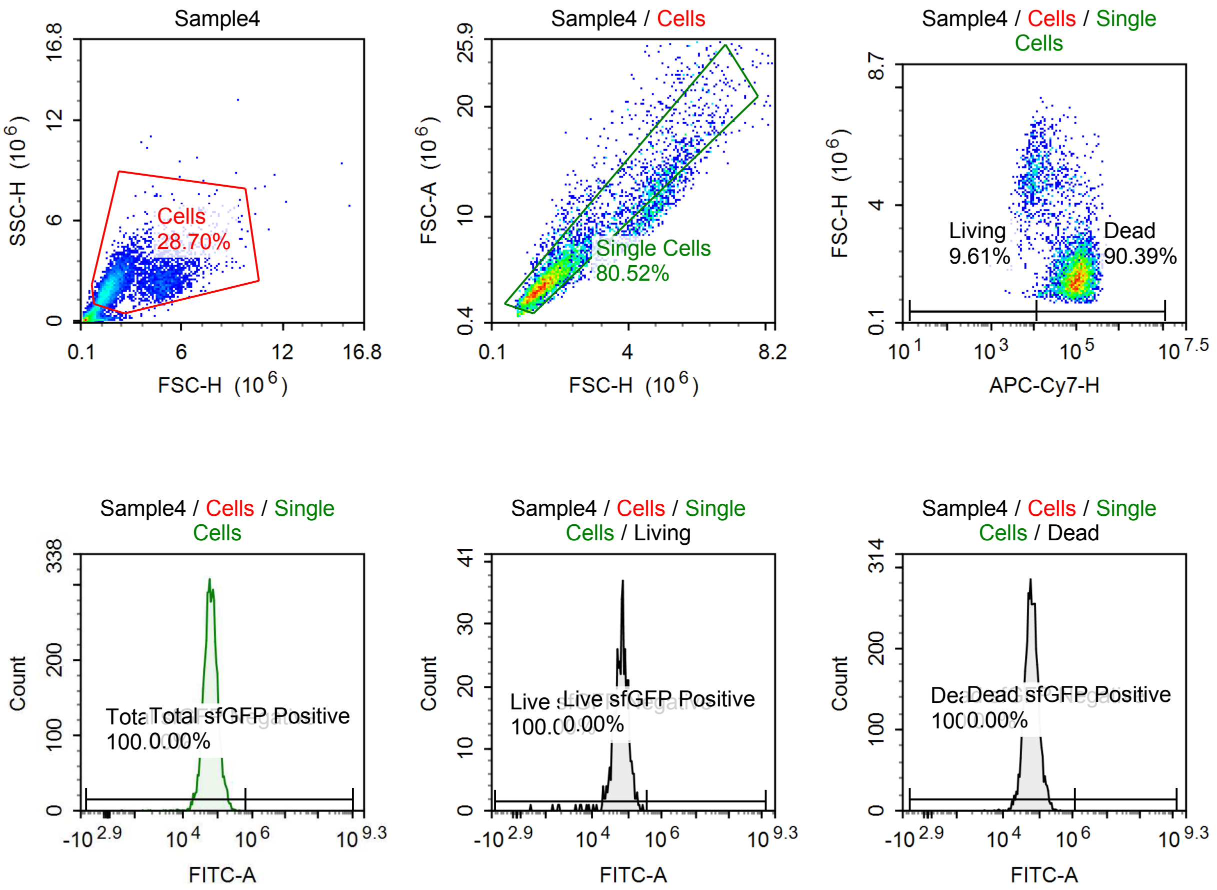

Sample Statistics of Sample4

| Gate | Count | % Parent | X | Y | Median X | Median Y |
|---|---|---|---|---|---|---|
| All | 19,766 | | | | | |
| Cells | 5,673 | 28.70 % | FSC-H | SSC-H | 1,918,826 | 2,164,354 |
| Single Cells | 4,568 | 80.52 % | FSC-H | FSC-A | 1,746,417 | 4,447,635 |
| Living | 439 | 9.61 % | APC-Cy7-H | FSC-H | 8,200 | 4,757,239 |
| Live sfGFP Negative | 439 | 100.00 % | FITC-A | Count | 65,559 | |
| Live sfGFP Positive | 0 | 0.00 % | FITC-A | Count | 0 | |
| Dead | 4,129 | 90.39 % | APC-Cy7-H | FSC-H | 92,435 | 1,673,105 |
| Dead sfGFP Negative | 4,129 | 100.00 % | FITC-A | Count | 60,839 | |
| Dead sfGFP Positive | 0 | 0.00 % | FITC-A | Count | 0 | |
| Total sfGFP Negative | 4,568 | 100.00 % | FITC-A | Count | 61,308 | |
| Total sfGFP Positive | 0 | 0.00 % | FITC-A | Count | 0 | |

## p059

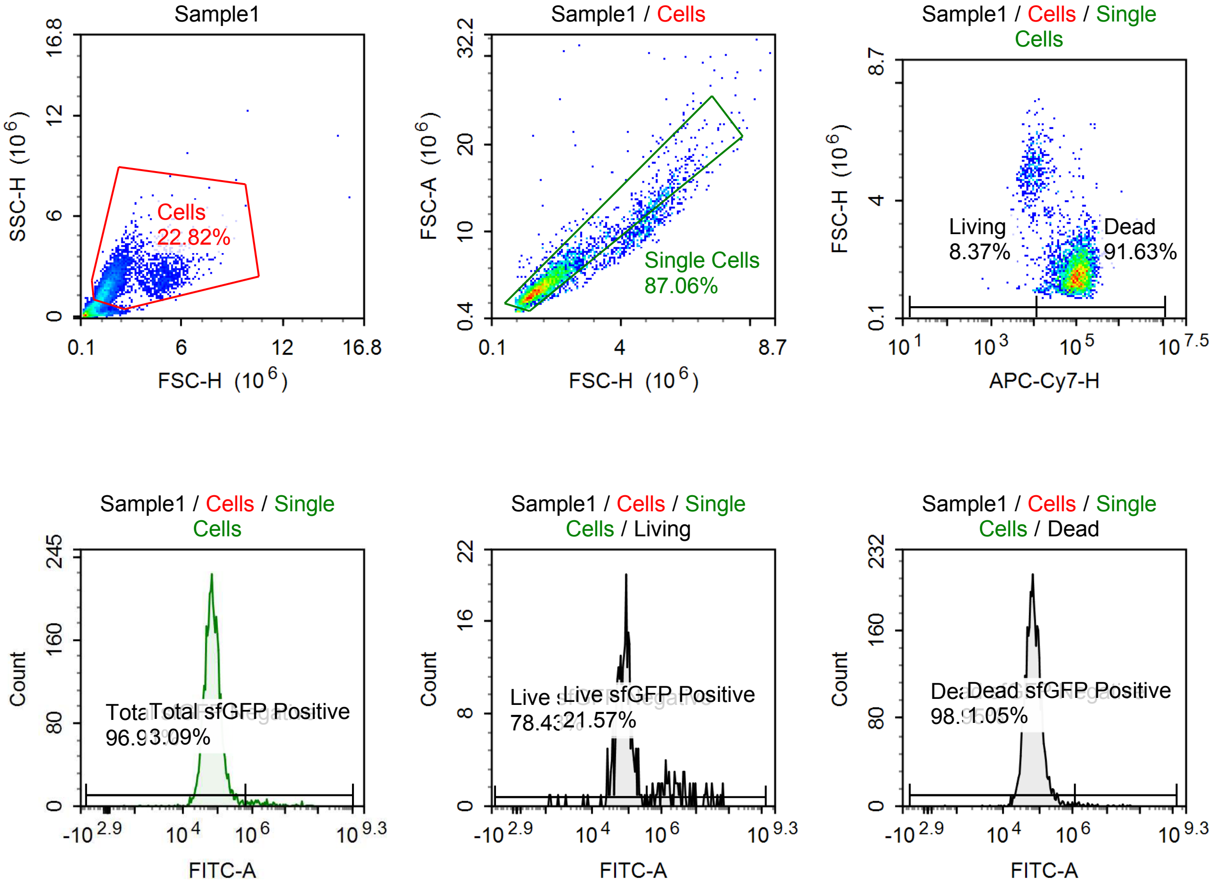

Sample Statistics of Sample1

| Gate | Count | % Parent | X | Y | Median X | Median Y |
|---|---|---|---|---|---|---|
| All | 18,394 | | | | | |
| Cells | 4,197 | 22.82 % | FSC-H | SSC-H | 1,682,791 | 1,920,581 |
| Single Cells | 3,654 | 87.06 % | FSC-H | FSC-A | 1,606,929 | 4,018,609 |
| Living | 306 | 8.37 % | APC-Cy7-H | FSC-H | 8,155 | 4,816,316 |
| Live sfGFP Negative | 240 | 78.43 % | FITC-A | Count | 77,512 | |
| Live sfGFP Positive | 66 | 21.57 % | FITC-A | Count | 2,564,600 | |
| Dead | 3,348 | 91.63 % | APC-Cy7-H | FSC-H | 99,169 | 1,556,671 |
| Dead sfGFP Negative | 3,313 | 98.95 % | FITC-A | Count | 67,712 | |
| Dead sfGFP Positive | 35 | 1.05 % | FITC-A | Count | 3,058,821 | |
| Total sfGFP Negative | 3,541 | 96.91 % | FITC-A | Count | 68,099 | |
| Total sfGFP Positive | 113 | 3.09 % | FITC-A | Count | 2,392,065 | |

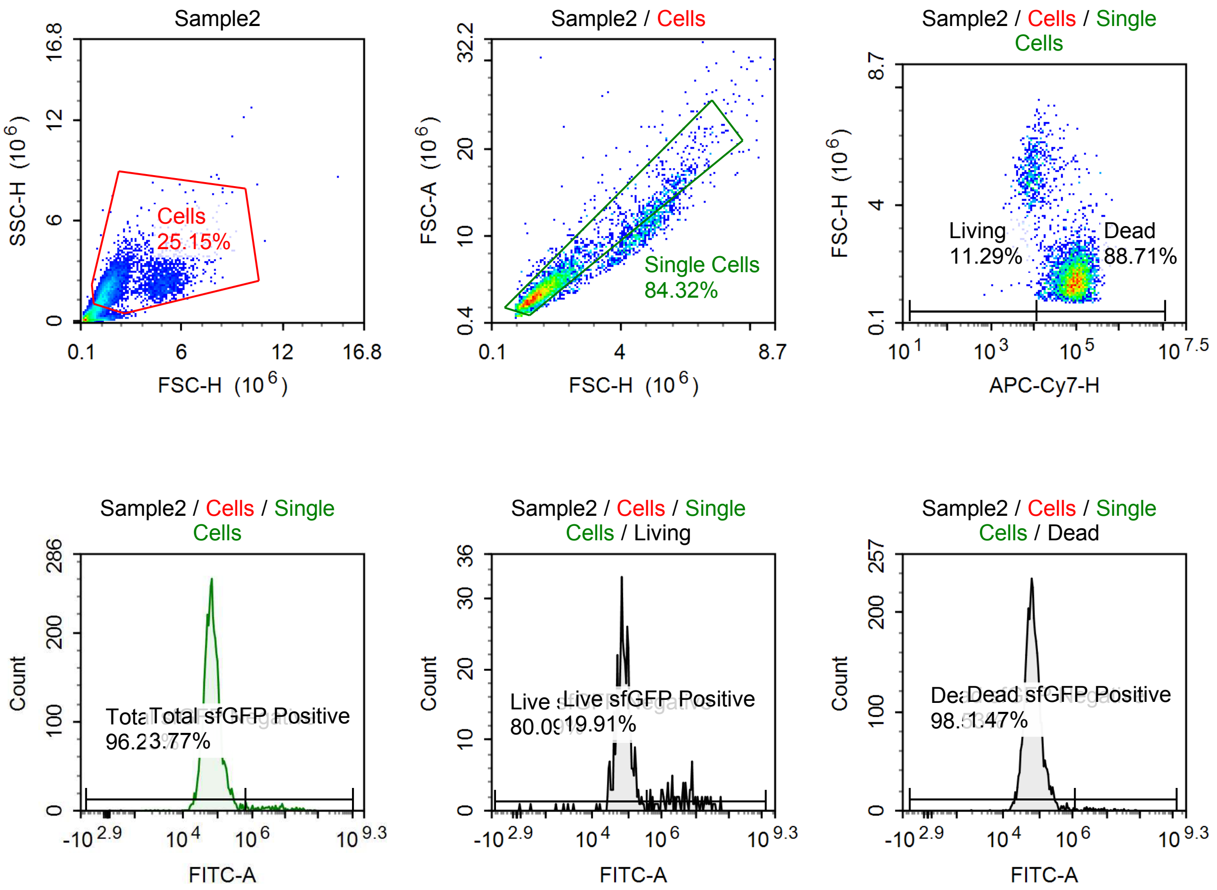

Sample Statistics of Sample2

| Gate | Count | % Parent | X | Y | Median X | Median Y |
|---|---|---|---|---|---|---|
| All | 19,502 | | | | | |
| Cells | 4,905 | 25.15 % | FSC-H | SSC-H | 1,727,851 | 1,988,363 |
| Single Cells | 4,136 | 84.32 % | FSC-H | FSC-A | 1,608,813 | 4,009,735 |
| Living | 467 | 11.29 % | APC-Cy7-H | FSC-H | 7,789 | 4,868,162 |
| Live sfGFP Negative | 374 | 80.09 % | FITC-A | Count | 73,777 | |
| Live sfGFP Positive | 93 | 19.91 % | FITC-A | Count | 4,531,137 | |
| Dead | 3,669 | 88.71 % | APC-Cy7-H | FSC-H | 85,096 | 1,523,328 |
| Dead sfGFP Negative | 3,615 | 98.53 % | FITC-A | Count | 64,788 | |
| Dead sfGFP Positive | 54 | 1.47 % | FITC-A | Count | 5,534,638 | |
| Total sfGFP Negative | 3,980 | 96.23 % | FITC-A | Count | 65,463 | |
| Total sfGFP Positive | 156 | 3.77 % | FITC-A | Count | 4,386,168 | |

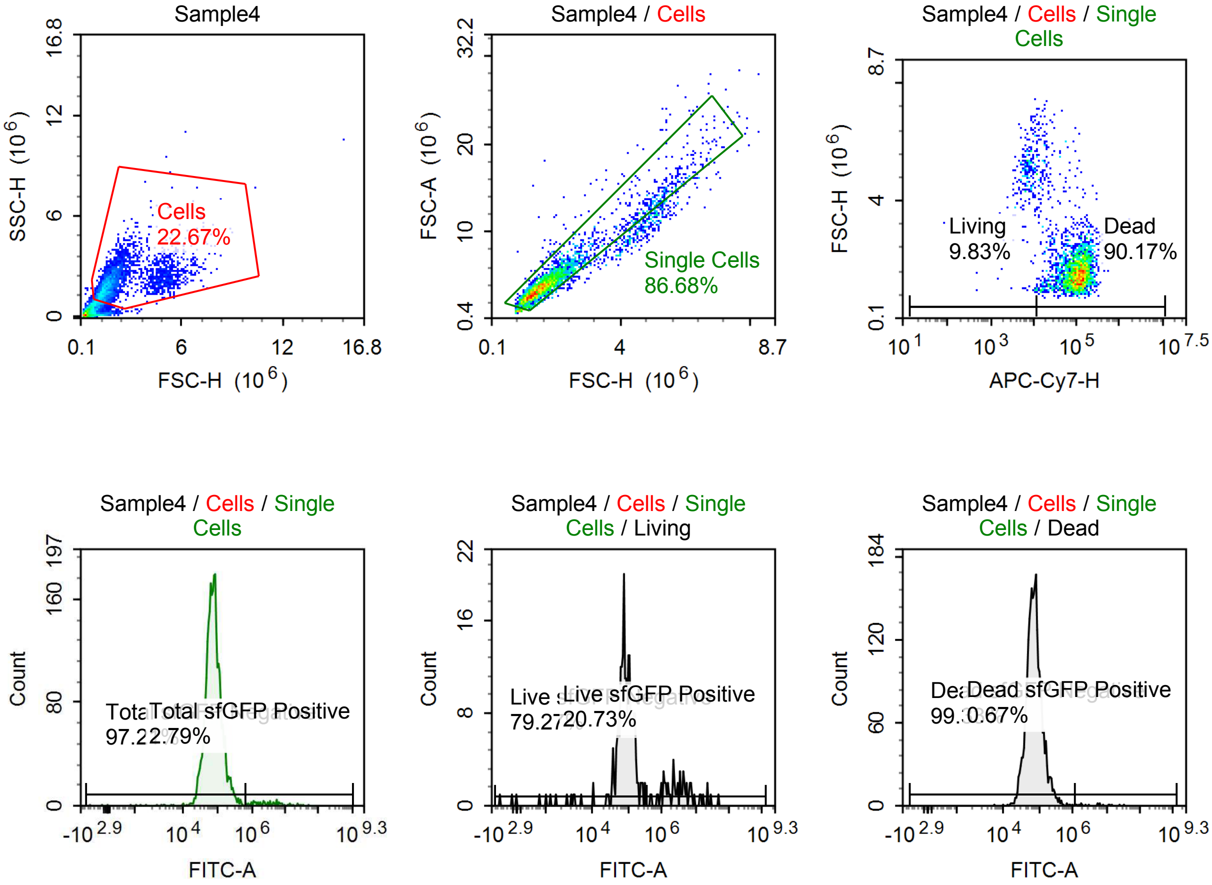

Sample Statistics of Sample4

| Gate | Count | % Parent | X | Y | Median X | Median Y |
|---|---|---|---|---|---|---|
| All | 14,241 | | | | | |
| Cells | 3,228 | 22.67 % | FSC-H | SSC-H | 1,699,334 | 1,964,629 |
| Single Cells | 2,798 | 86.68 % | FSC-H | FSC-A | 1,628,552 | 4,040,780 |
| Living | 275 | 9.83 % | APC-Cy7-H | FSC-H | 7,445 | 4,972,978 |
| Live sfGFP Negative | 218 | 79.27 % | FITC-A | Count | 80,606 | |
| Live sfGFP Positive | 57 | 20.73 % | FITC-A | Count | 3,014,812 | |
| Dead | 2,523 | 90.17 % | APC-Cy7-H | FSC-H | 101,701 | 1,560,428 |
| Dead sfGFP Negative | 2,506 | 99.33 % | FITC-A | Count | 72,331 | |
| Dead sfGFP Positive | 17 | 0.67 % | FITC-A | Count | 5,769,047 | |
| Total sfGFP Negative | 2,720 | 97.21 % | FITC-A | Count | 73,337 | |
| Total sfGFP Positive | 78 | 2.79 % | FITC-A | Count | 3,049,141 | |

#### p059 + 1 uM Tet

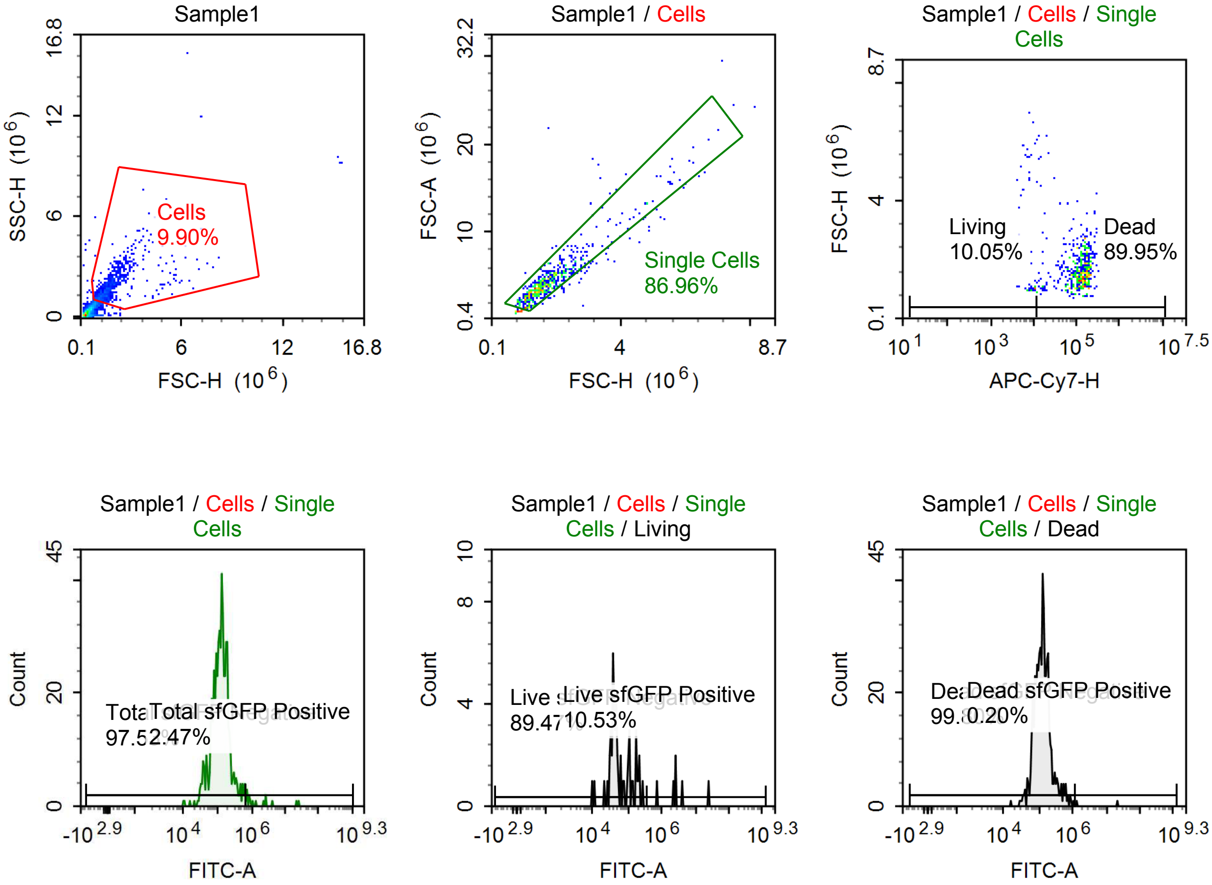

Sample Statistics of Sample1

| Gate | Count | % Parent | X | Y | Median X | Median Y |
|---|---|---|---|---|---|---|
| All | 6,588 | | | | | |
| Cells | 652 | 9.90 % | FSC-H | SSC-H | 1,583,140 | 1,703,861 |
| Single Cells | 567 | 86.96 % | FSC-H | FSC-A | 1,582,969 | 3,985,626 |
| Living | 57 | 10.05 % | APC-Cy7-H | FSC-H | 7,933 | 1,337,084 |
| Live sfGFP Negative | 51 | 89.47 % | FITC-A | Count | 46,787 | |
| Live sfGFP Positive | 6 | 10.53 % | FITC-A | Count | 2,615,035 | |
| Dead | 510 | 89.95 % | APC-Cy7-H | FSC-H | 128,396 | 1,584,986 |
| Dead sfGFP Negative | 509 | 99.80 % | FITC-A | Count | 127,335 | |
| Dead sfGFP Positive | 1 | 0.20 % | FITC-A | Count | 21,836,930 | |
| Total sfGFP Negative | 553 | 97.53 % | FITC-A | Count | 122,163 | |
| Total sfGFP Positive | 14 | 2.47 % | FITC-A | Count | 1,189,044 | |

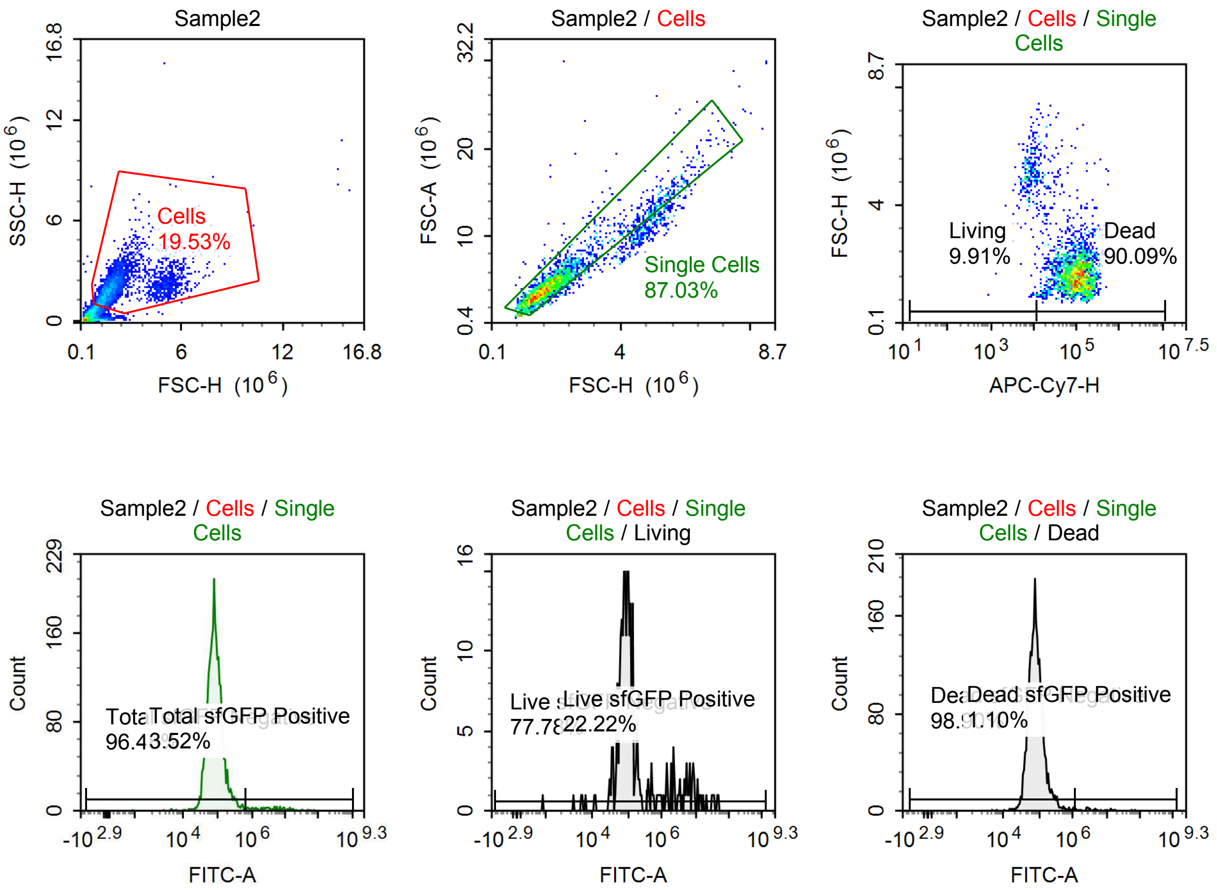

Sample Statistics of Sample2

| Gate | Count | % Parent | X | Y | Median X | Median Y |
|---|---|---|---|---|---|---|
| All | 16,030 | | | | | |
| Cells | 3,131 | 19.53 % | FSC-H | SSC-H | 1,806,629 | 1,963,008 |
| Single Cells | 2,725 | 87.03 % | FSC-H | FSC-A | 1,730,290 | 4,327,121 |
| Living | 270 | 9.91 % | APC-Cy7-H | FSC-H | 8,212 | 4,993,359 |
| Live sfGFP Negative | 210 | 77.78 % | FITC-A | Count | 85,877 | |
| Live sfGFP Positive | 60 | 22.22 % | FITC-A | Count | 3,875,599 | |
| Dead | 2,455 | 90.09 % | APC-Cy7-H | FSC-H | 96,974 | 1,664,094 |
| Dead sfGFP Negative | 2,428 | 98.90 % | FITC-A | Count | 79,414 | |
| Dead sfGFP Positive | 27 | 1.10 % | FITC-A | Count | 4,466,340 | |
| Total sfGFP Negative | 2,629 | 96.48 % | FITC-A | Count | 79,691 | |
| Total sfGFP Positive | 96 | 3.52 % | FITC-A | Count | 3,318,827 | |

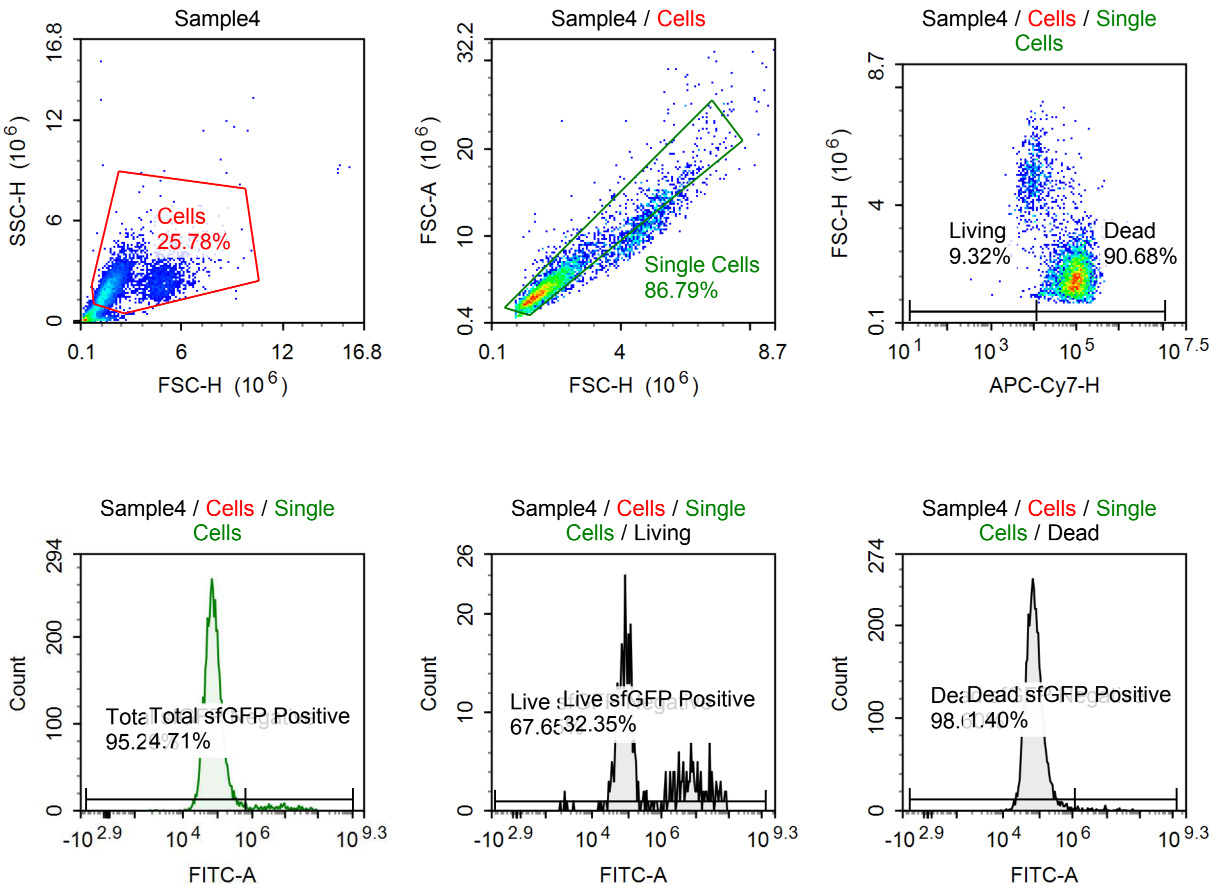

Sample Statistics of Sample4

| Gate | Count | % Parent | X | Y | Median X | Median Y |
|---|---|---|---|---|---|---|
| All | 21,051 | | | | | |
| Cells | 5,428 | 25.78 % | FSC-H | SSC-H | 1,726,221 | 2,007,316 |
| Single Cells | 4,711 | 86.79 % | FSC-H | FSC-A | 1,633,184 | 4,133,591 |
| Living | 439 | 9.32 % | APC-Cy7-H | FSC-H | 8,135 | 4,810,179 |
| Live sfGFP Negative | 297 | 67.65 % | FITC-A | Count | 82,072 | |
| Live sfGFP Positive | 142 | 32.35 % | FITC-A | Count | 7,542,460 | |
| Dead | 4,272 | 90.68 % | APC-Cy7-H | FSC-H | 87,192 | 1,570,977 |
| Dead sfGFP Negative | 4,212 | 98.60 % | FITC-A | Count | 71,373 | |
| Dead sfGFP Positive | 60 | 1.40 % | FITC-A | Count | 5,780,530 | |
| Total sfGFP Negative | 4,489 | 95.29 % | FITC-A | Count | 71,730 | |
| Total sfGFP Positive | 222 | 4.71 % | FITC-A | Count | 6,373,531 | |

### Report of Herb_Owen TET Aptazyme Second

File Name: Herb_Owen TET Aptazyme Second Attempt 3_27_26 Run Time: 3/27/2026 10:57 AM

Cytometer: NovoCyte Quanteon 621190610657 Software: NovoExpress 1.6.1

#### p059 mRNA [100 ng]

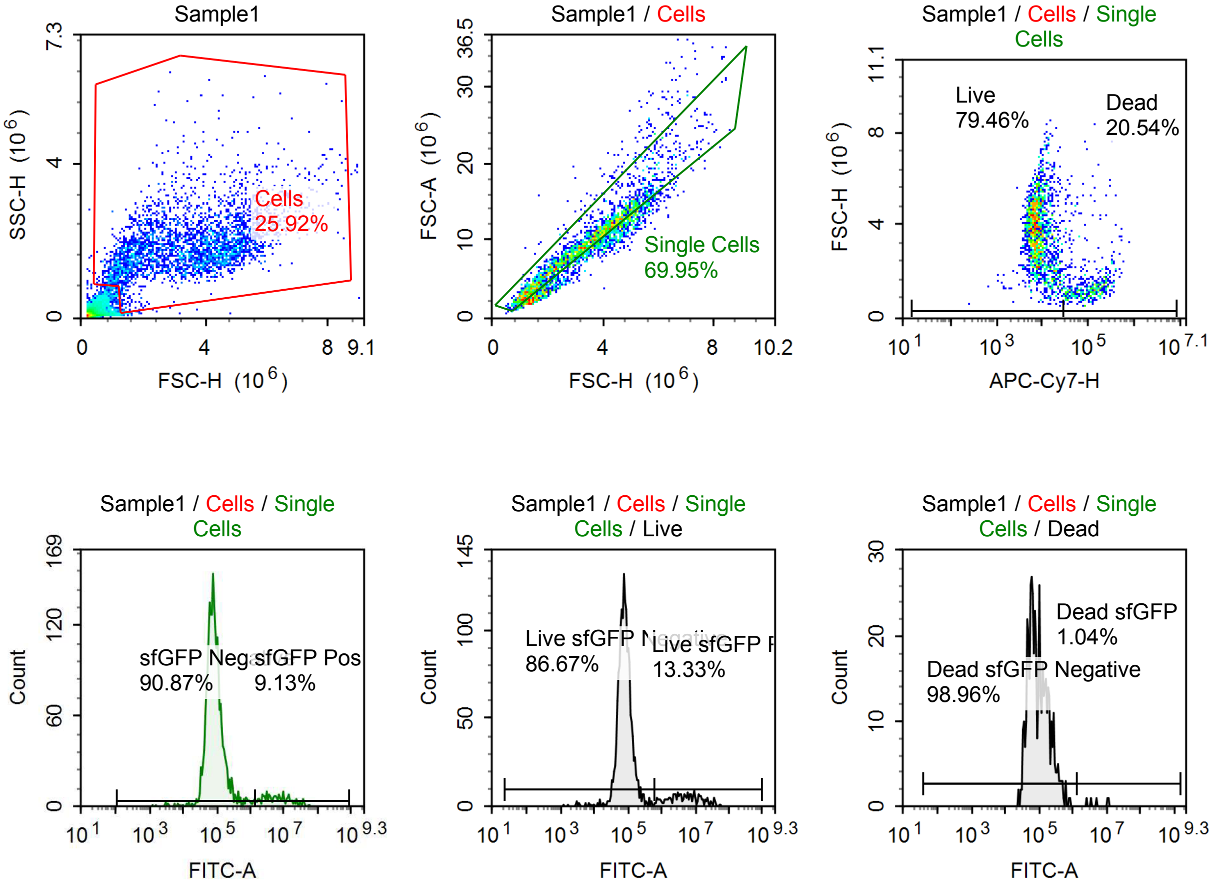


Sample Statistics of Sample1

| Gate | Count | % Parent | X | Y | Median X | Median Y |
|---|---|---|---|---|---|---|
| All | 12,863 | | | | | |
| Cells | 3,334 | 25.92 % | FSC-H | SSC-H | 3,418,338 | 1,786,455 |
| Single Cells | 2,332 | 69.95 % | FSC-H | FSC-A | 3,270,850 | 9,354,888 |
| Live | 1,853 | 79.46 % | APC-Cy7-H | FSC-H | 7,697 | 3,680,644 |
| Live sfGFP Negative | 1,606 | 86.67 % | FITC-A | Count | 78,812 | |
| Live sfGFP Positive | 247 | 13.33 % | FITC-A | Count | 4,911,307 | |
| Dead | 479 | 20.54 % | APC-Cy7-H | FSC-H | 110,166 | 1,324,854 |
| Dead sfGFP Negative | 474 | 98.96 % | FITC-A | Count | 84,615 | |
| Dead sfGFP Positive | 5 | 1.04 % | FITC-A | Count | 5,007,589 | |
| sfGFP Negative | 2,119 | 90.87 % | FITC-A | Count | 80,746 | |
| sfGFP Positive | 213 | 9.13 % | FITC-A | Count | 6,137,635 | |

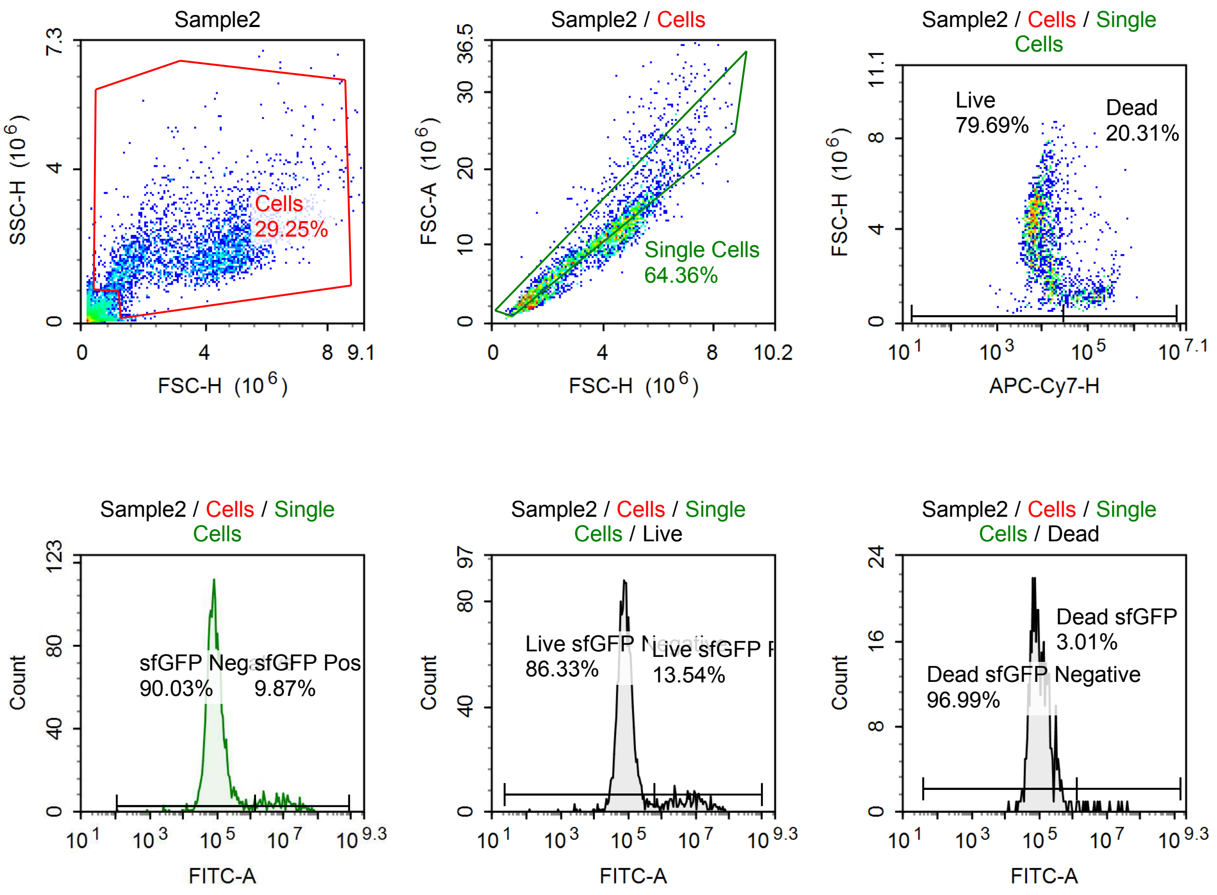


Sample Statistics of Sample2

| Gate | Count | % Parent | X | Y | Median X | Median Y |
|---|---|---|---|---|---|---|
| All | 10,438 | | | | | |
| Cells | 3,053 | 29.25 % | FSC-H | SSC-H | 3,879,912 | 1,811,085 |
| Single Cells | 1,965 | 64.36 % | FSC-H | FSC-A | 3,506,661 | 9,964,322 |
| Live | 1,566 | 79.69 % | APC-Cy7-H | FSC-H | 8,476 | 3,939,926 |
| Live sfGFP Negative | 1,352 | 86.33 % | FITC-A | Count | 82,297 | |
| Live sfGFP Positive | 212 | 13.54 % | FITC-A | Count | 6,338,961 | |
| Dead | 399 | 20.31 % | APC-Cy7-H | FSC-H | 103,223 | 1,379,677 |
| Dead sfGFP Negative | 387 | 96.99 % | FITC-A | Count | 97,512 | |
| Dead sfGFP Positive | 12 | 3.01 % | FITC-A | Count | 4,800,572 | |
| sfGFP Negative | 1,769 | 90.03 % | FITC-A | Count | 85,224 | |
| sfGFP Positive | 194 | 9.87 % | FITC-A | Count | 8,004,019 | |

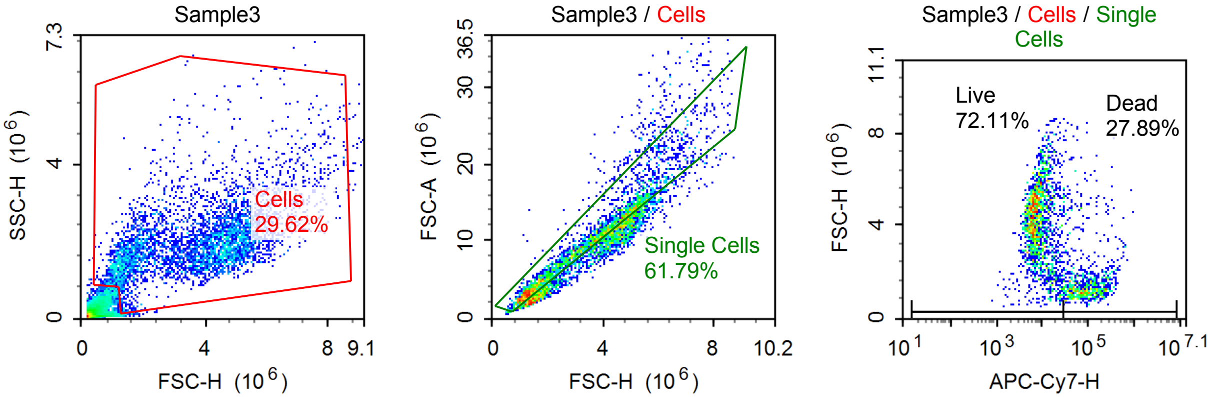

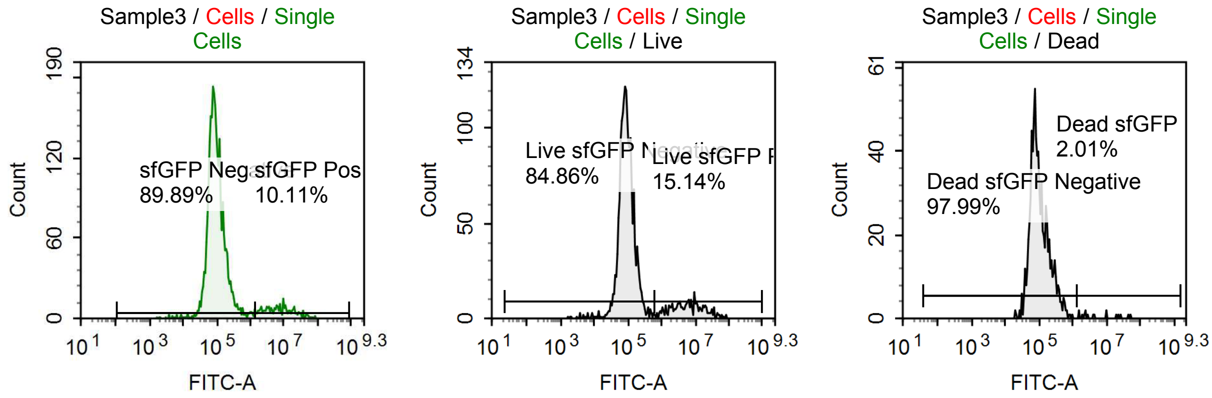


Sample Statistics of Sample3

| Gate | Count | % Parent | X | Y | Median X | Median Y |
|---|---|---|---|---|---|---|
| All | 15,612 | | | | | |
| Cells | 4,625 | 29.62 % | FSC-H | SSC-H | 3,913,393 | 1,883,159 |
| Single Cells | 2,858 | 61.79 % | FSC-H | FSC-A | 3,547,041 | 10,100,480 |
| Live | 2,061 | 72.11 % | APC-Cy7-H | FSC-H | 7,952 | 4,180,852 |
| Live sfGFP Negative | 1,749 | 84.86 % | FITC-A | Count | 88,766 | |
| Live sfGFP Positive | 312 | 15.14 % | FITC-A | Count | 6,223,671 | |
| Dead | 797 | 27.89 % | APC-Cy7-H | FSC-H | 89,350 | 1,375,322 |
| Dead sfGFP Negative | 781 | 97.99 % | FITC-A | Count | 91,086 | |
| Dead sfGFP Positive | 16 | 2.01 % | FITC-A | Count | 4,304,054 | |
| sfGFP Negative | 2,569 | 89.89 % | FITC-A | Count | 90,244 | |
| sfGFP Positive | 289 | 10.11 % | FITC-A | Count | 7,325,173 | |

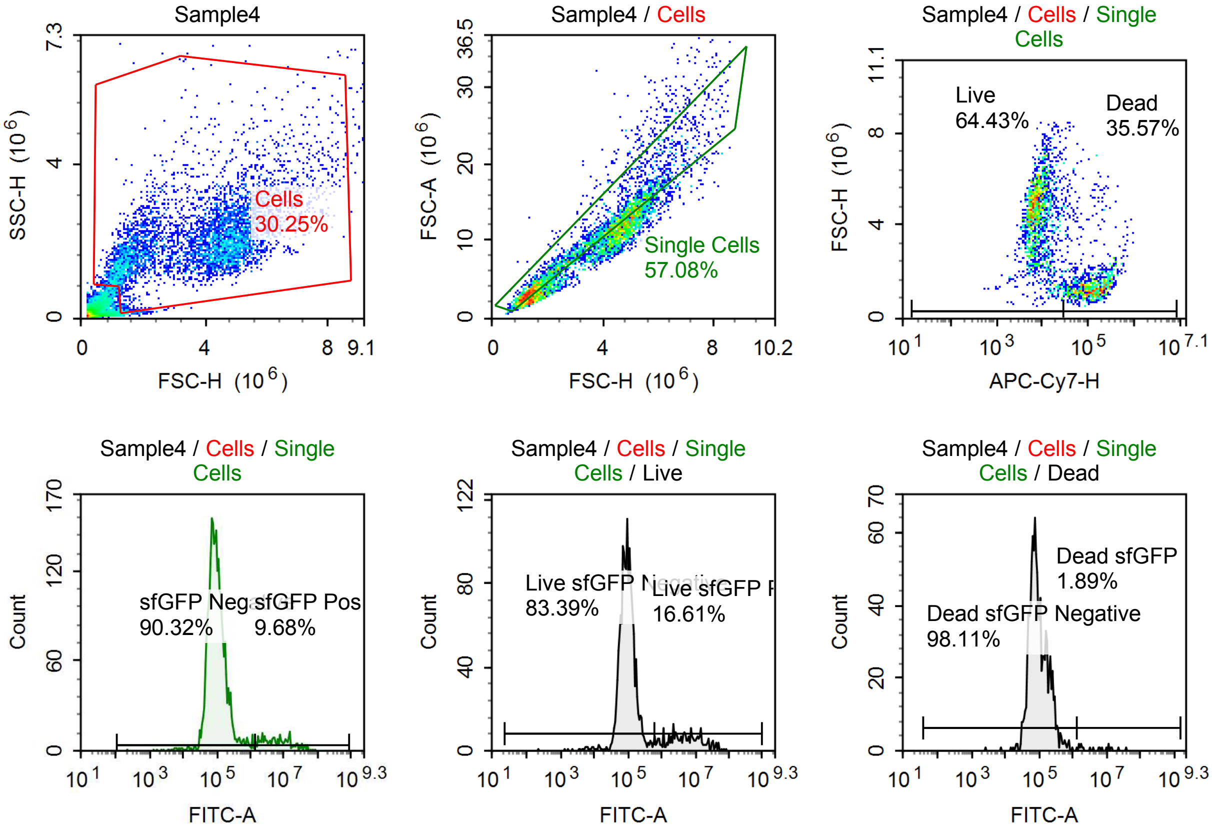

Sample Statistics of Sample4

| Gate | Count | % Parent | X | Y | Median X | Median Y |
|---|---|---|---|---|---|---|
| All | 16,397 | | | | | |
| Cells | 4,960 | 30.25 % | FSC-H | SSC-H | 4,143,060 | 1,913,154 |
| Single Cells | 2,831 | 57.08 % | FSC-H | FSC-A | 3,445,039 | 9,874,781 |
| Live | 1,824 | 64.43 % | APC-Cy7-H | FSC-H | 7,797 | 4,519,877 |
| Live sfGFP Negative | 1,521 | 83.39 % | FITC-A | Count | 94,538 | |
| Live sfGFP Positive | 303 | 16.61 % | FITC-A | Count | 5,021,314 | |
| Dead | 1,007 | 35.57 % | APC-Cy7-H | FSC-H | 115,908 | 1,381,749 |
| Dead sfGFP Negative | 988 | 98.11 % | FITC-A | Count | 87,403 | |
| Dead sfGFP Positive | 19 | 1.89 % | FITC-A | Count | 5,665,973 | |
| sfGFP Negative | 2,557 | 90.32 % | FITC-A | Count | 93,112 | |
| sfGFP Positive | 274 | 9.68 % | FITC-A | Count | 6,689,175 | |

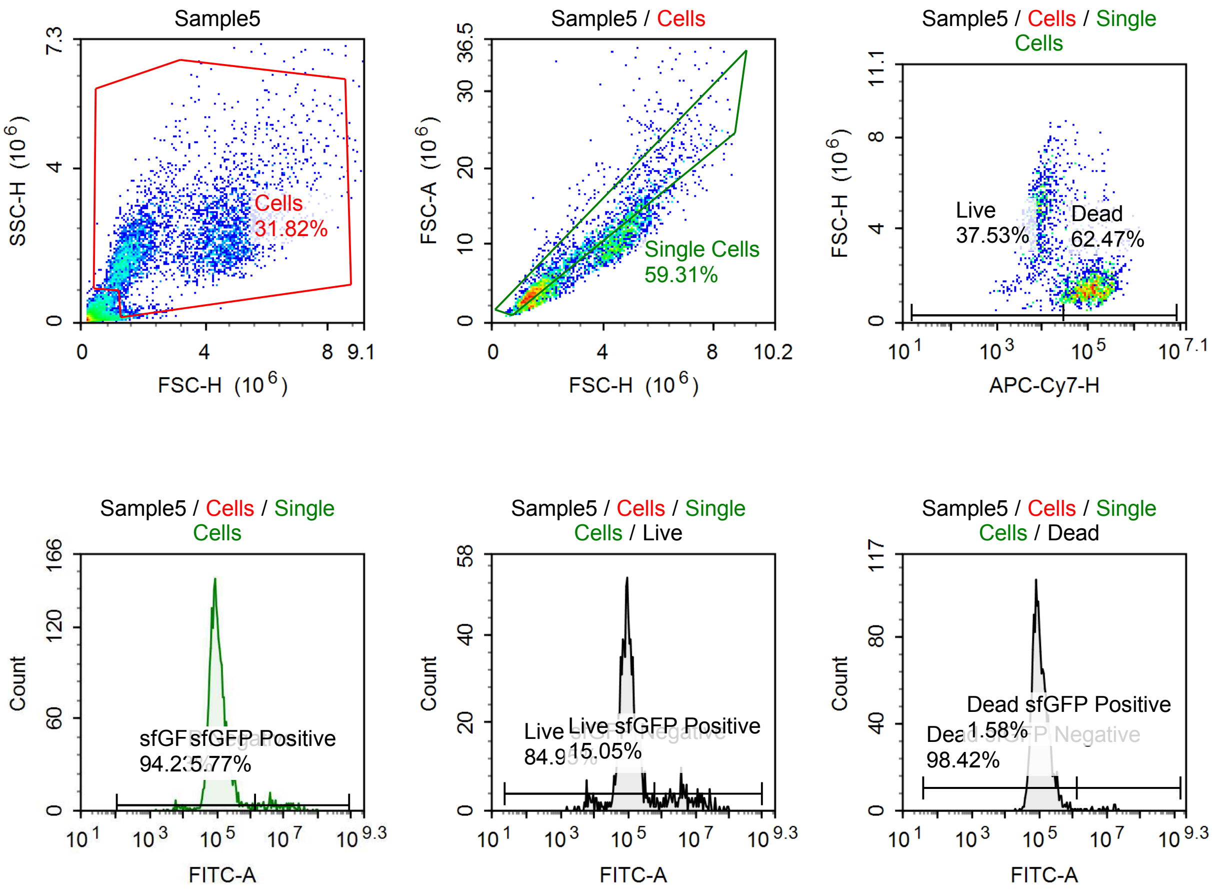

Sample Statistics of Sample5

| Gate | Count | % Parent | X | Y | Median X | Median Y |
|---|---|---|---|---|---|---|
| All | 12,852 | | | | | |
| Cells | 4,089 | 31.82 % | FSC-H | SSC-H | 2,636,904 | 1,983,598 |
| Single Cells | 2,425 | 59.31 % | FSC-H | FSC-A | 1,814,007 | 5,149,901 |
| Live | 910 | 37.53 % | APC-Cy7-H | FSC-H | 8,933 | 4,300,268 |
| Live sfGFP Negative | 773 | 84.95 % | FITC-A | Count | 95,500 | |
| Live sfGFP Positive | 137 | 15.05 % | FITC-A | Count | 4,543,510 | |
| Dead | 1,515 | 62.47 % | APC-Cy7-H | FSC-H | 112,709 | 1,478,356 |
| Dead sfGFP Negative | 1,491 | 98.42 % | FITC-A | Count | 100,597 | |
| Dead sfGFP Positive | 24 | 1.58 % | FITC-A | Count | 7,857,137 | |
| sfGFP Negative | 2,285 | 94.23 % | FITC-A | Count | 99,010 | |
| sfGFP Positive | 140 | 5.77 % | FITC-A | Count | 5,700,365 | |

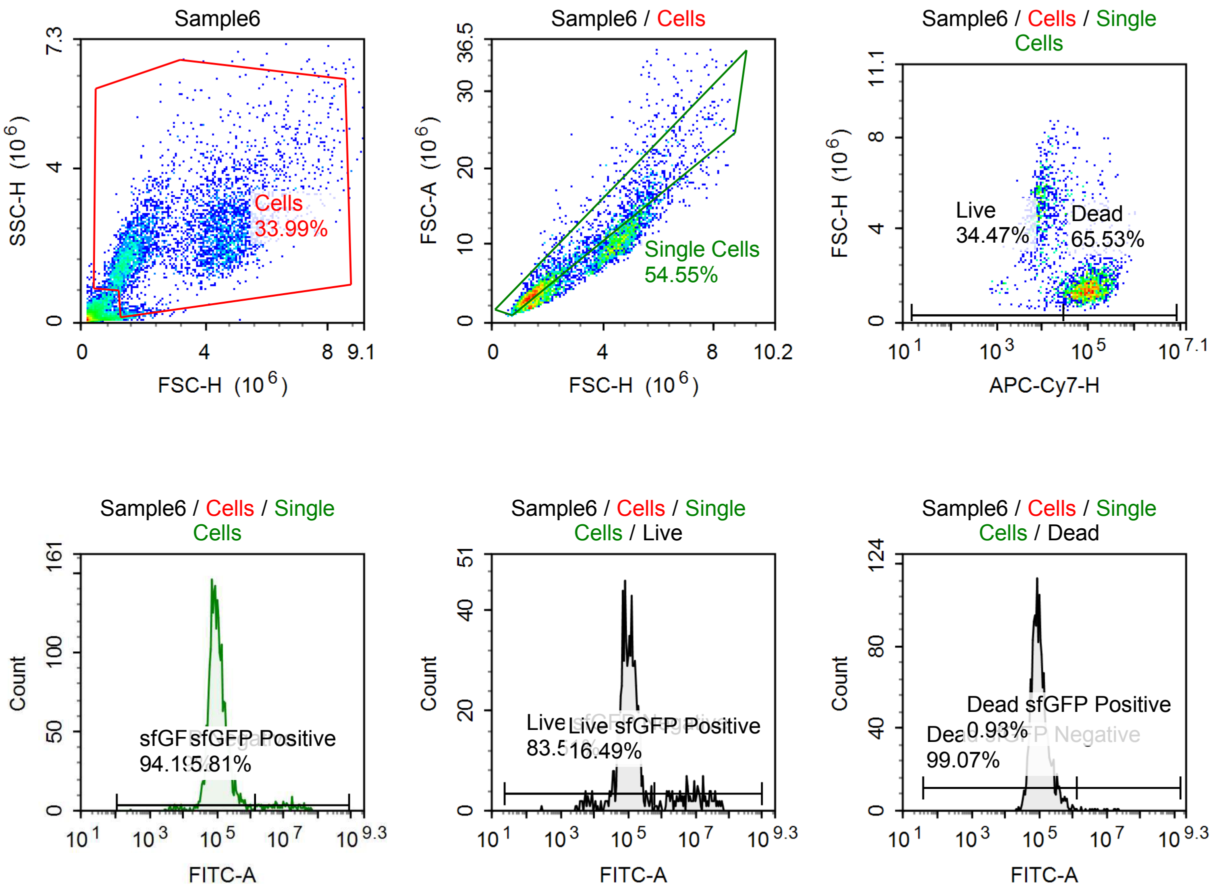

Sample Statistics of Sample6

| Gate | Count | % Parent | X | Y | Median X | Median Y |
|---|---|---|---|---|---|---|
| All | 13,284 | | | | | |
| Cells | 4,515 | 33.99 % | FSC-H | SSC-H | 2,755,861 | 2,039,483 |
| Single Cells | 2,463 | 54.55 % | FSC-H | FSC-A | 1,740,876 | 4,783,328 |
| Live | 849 | 34.47 % | APC-Cy7-H | FSC-H | 9,627 | 4,506,512 |
| Live sfGFP Negative | 709 | 83.51 % | FITC-A | Count | 101,352 | |
| Live sfGFP Positive | 140 | 16.49 % | FITC-A | Count | 8,329,657 | |
| Dead | 1,614 | 65.53 % | APC-Cy7-H | FSC-H | 101,871 | 1,488,792 |
| Dead sfGFP Negative | 1,599 | 99.07 % | FITC-A | Count | 96,916 | |
| Dead sfGFP Positive | 15 | 0.93 % | FITC-A | Count | 5,298,641 | |
| sfGFP Negative | 2,320 | 94.19 % | FITC-A | Count | 98,461 | |
| sfGFP Positive | 143 | 5.81 % | FITC-A | Count | 9,395,766 | |

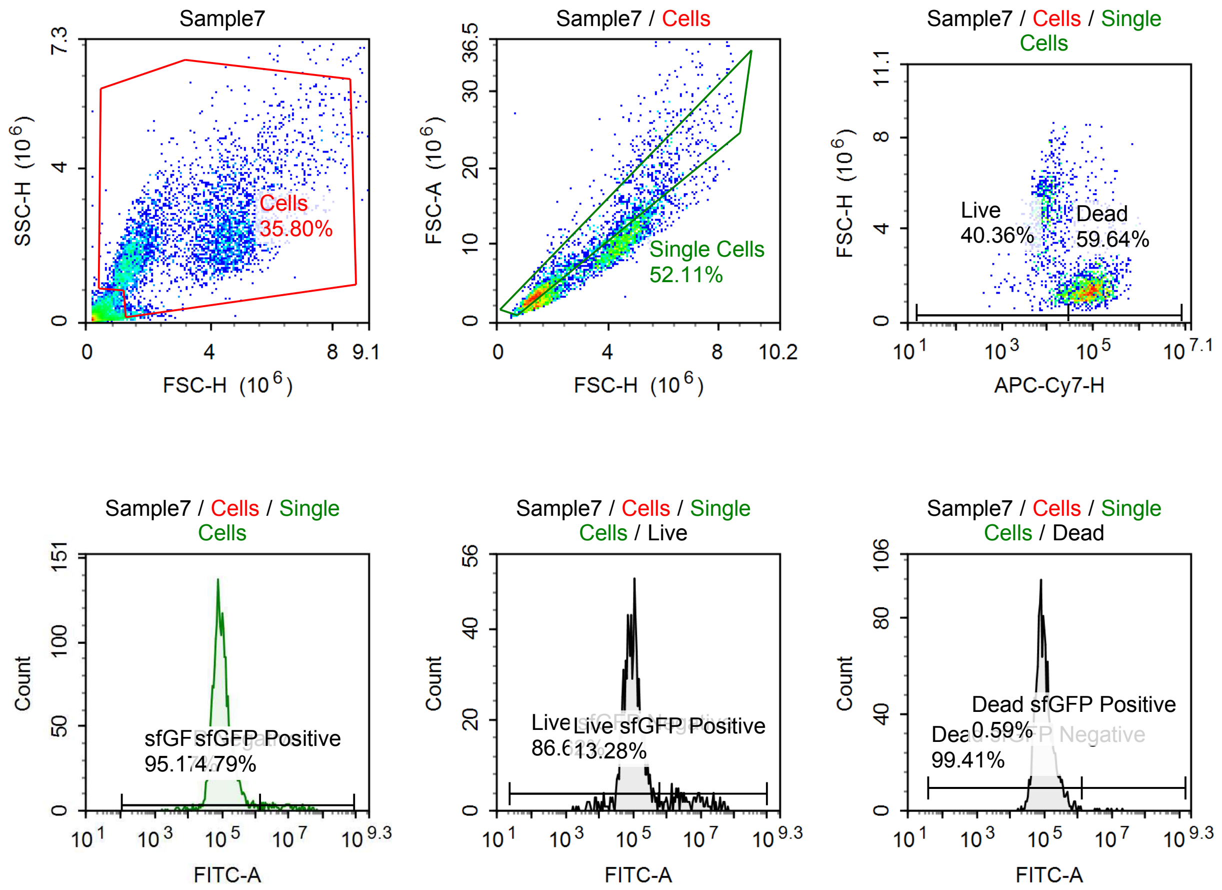

Sample Statistics of Sample7

| Gate | Count | % Parent | X | Y | Median X | Median Y |
|---|---|---|---|---|---|---|
| All | 12,205 | | | | | |
| Cells | 4,370 | 35.80 % | FSC-H | SSC-H | 3,101,802 | 2,032,149 |
| Single Cells | 2,277 | 52.11 % | FSC-H | FSC-A | 1,723,048 | 4,660,251 |
| Live | 919 | 40.36 % | APC-Cy7-H | FSC-H | 10,231 | 4,273,580 |
| Live sfGFP Negative | 796 | 86.62 % | FITC-A | Count | 93,767 | |
| Live sfGFP Positive | 122 | 13.28 % | FITC-A | Count | 4,294,995 | |
| Dead | 1,358 | 59.64 % | APC-Cy7-H | FSC-H | 88,023 | 1,441,887 |
| Dead sfGFP Negative | 1,350 | 99.41 % | FITC-A | Count | 92,495 | |
| Dead sfGFP Positive | 8 | 0.59 % | FITC-A | Count | 7,759,381 | |
| sfGFP Negative | 2,167 | 95.17 % | FITC-A | Count | 93,738 | |
| sfGFP Positive | 109 | 4.79 % | FITC-A | Count | 7,205,552 | |

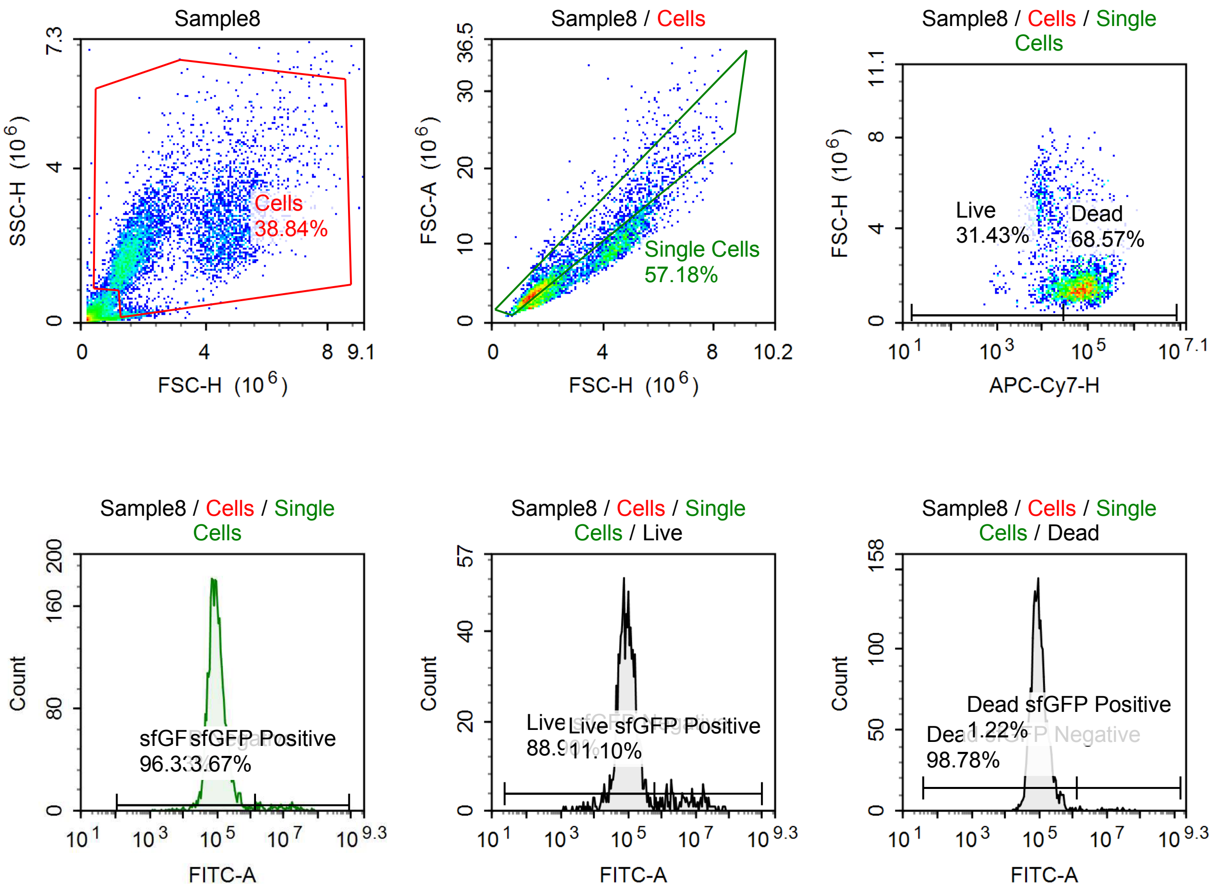

Sample Statistics of Sample8

| Gate | Count | % Parent | X | Y | Median X | Median Y |
|---|---|---|---|---|---|---|
| All | 14,585 | | | | | |
| Cells | 5,665 | 38.84 % | FSC-H | SSC-H | 2,050,278 | 2,011,427 |
| Single Cells | 3,239 | 57.18 % | FSC-H | FSC-A | 1,611,413 | 4,226,895 |
| Live | 1,018 | 31.43 % | APC-Cy7-H | FSC-H | 13,326 | 2,483,542 |
| Live sfGFP Negative | 905 | 88.90 % | FITC-A | Count | 91,243 | |
| Live sfGFP Positive | 113 | 11.10 % | FITC-A | Count | 5,363,308 | |
| Dead | 2,221 | 68.57 % | APC-Cy7-H | FSC-H | 79,516 | 1,508,697 |
| Dead sfGFP Negative | 2,194 | 98.78 % | FITC-A | Count | 94,550 | |
| Dead sfGFP Positive | 27 | 1.22 % | FITC-A | Count | 11,927,610 | |
| sfGFP Negative | 3,120 | 96.33 % | FITC-A | Count | 93,829 | |
| sfGFP Positive | 119 | 3.67 % | FITC-A | Count | 8,320,900 | |

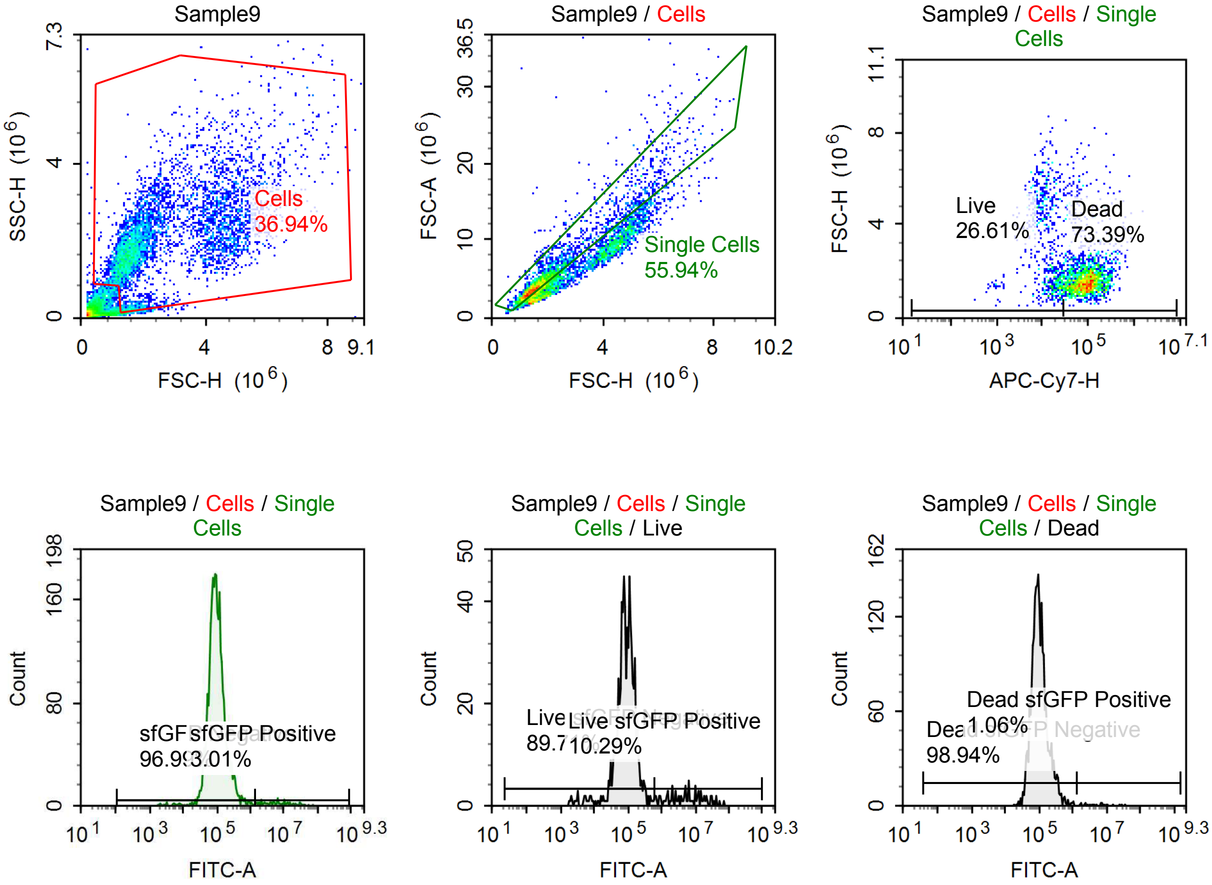

Sample Statistics of Sample9

| Gate | Count | % Parent | X | Y | Median X | Median Y |
|---|---|---|---|---|---|---|
| All | 14,317 | | | | | |
| Cells | 5,288 | 36.94 % | FSC-H | SSC-H | 1,976,198 | 1,910,397 |
| Single Cells | 2,958 | 55.94 % | FSC-H | FSC-A | 1,622,980 | 4,197,661 |
| Live | 787 | 26.61 % | APC-Cy7-H | FSC-H | 13,538 | 2,440,705 |
| Live sfGFP Negative | 706 | 89.71 % | FITC-A | Count | 88,095 | |
| Live sfGFP Positive | 81 | 10.29 % | FITC-A | Count | 5,481,060 | |
| Dead | 2,171 | 73.39 % | APC-Cy7-H | FSC-H | 91,552 | 1,559,436 |
| Dead sfGFP Negative | 2,148 | 98.94 % | FITC-A | Count | 96,798 | |
| Dead sfGFP Positive | 23 | 1.06 % | FITC-A | Count | 4,916,780 | |
| sfGFP Negative | 2,869 | 96.99 % | FITC-A | Count | 95,648 | |
| sfGFP Positive | 89 | 3.01 % | FITC-A | Count | 6,605,445 | |

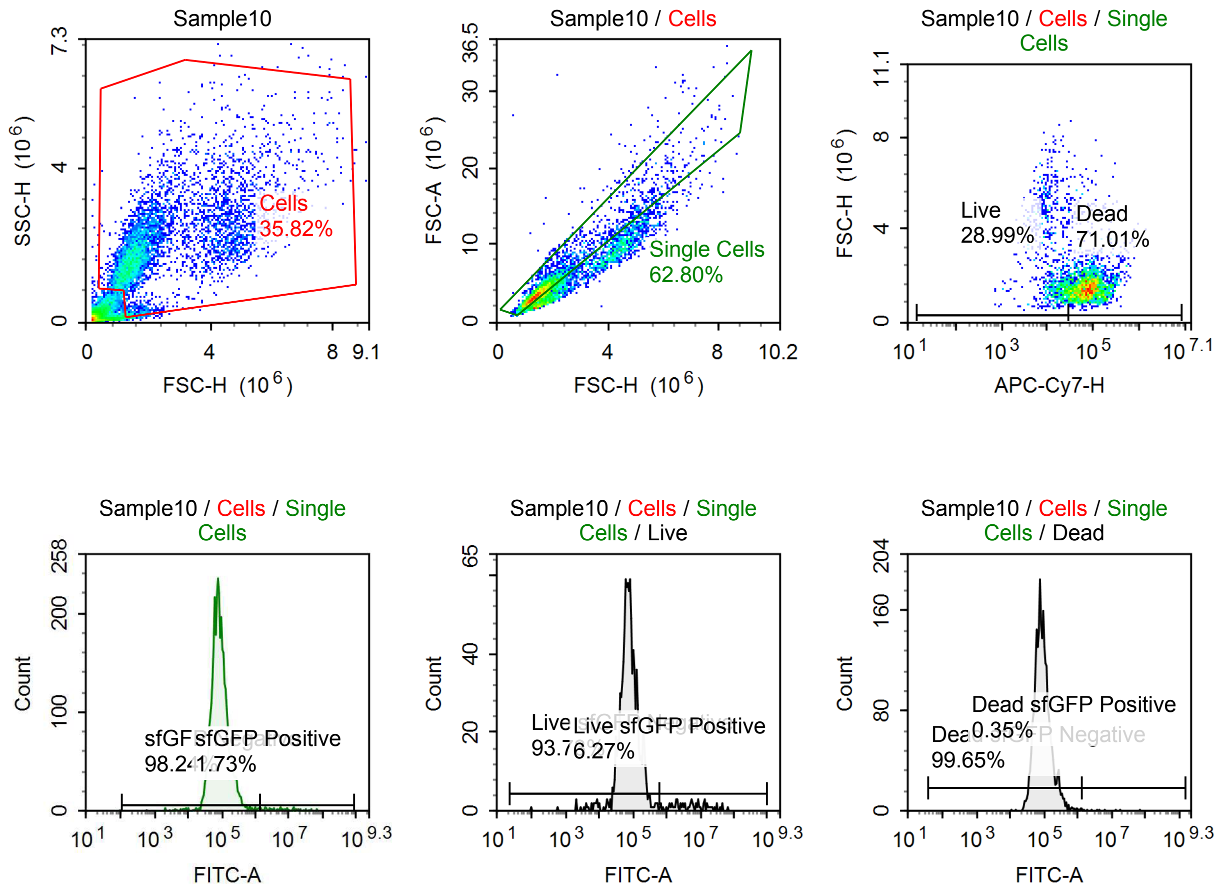

Sample Statistics of Sample10

| Gate | Count | % Parent | X | Y | Median X | Median Y |
|---|---|---|---|---|---|---|
| All | 15,903 | | | | | |
| Cells | 5,696 | 35.82 % | FSC-H | SSC-H | 1,753,175 | 1,851,059 |
| Single Cells | 3,577 | 62.80 % | FSC-H | FSC-A | 1,527,338 | 3,937,869 |
| Live | 1,037 | 28.99 % | APC-Cy7-H | FSC-H | 14,348 | 1,786,297 |
| Live sfGFP Negative | 972 | 93.73 % | FITC-A | Count | 75,880 | |
| Live sfGFP Positive | 65 | 6.27 % | FITC-A | Count | 5,499,774 | |
| Dead | 2,540 | 71.01 % | APC-Cy7-H | FSC-H | 77,592 | 1,476,257 |
| Dead sfGFP Negative | 2,531 | 99.65 % | FITC-A | Count | 84,926 | |
| Dead sfGFP Positive | 9 | 0.35 % | FITC-A | Count | 4,347,979 | |
| sfGFP Negative | 3,514 | 98.24 % | FITC-A | Count | 82,398 | |
| sfGFP Positive | 62 | 1.73 % | FITC-A | Count | 6,356,365 | |

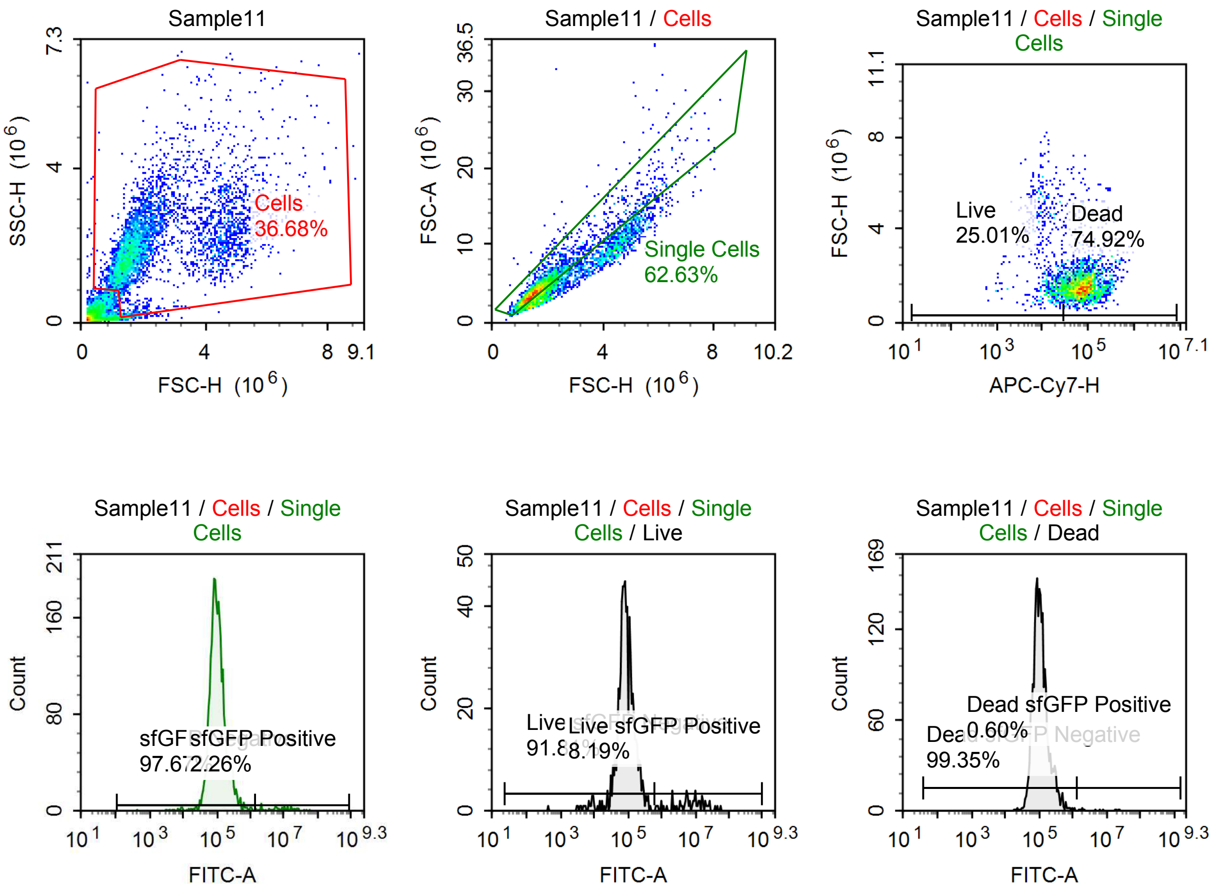

Sample Statistics of Sample11

| Gate | Count | % Parent | X | Y | Median X | Median Y |
|---|---|---|---|---|---|---|
| All | 12,532 | | | | | |
| Cells | 4,597 | 36.68 % | FSC-H | SSC-H | 1,800,110 | 1,934,550 |
| Single Cells | 2,879 | 62.63 % | FSC-H | FSC-A | 1,573,610 | 4,110,585 |
| Live | 720 | 25.01 % | APC-Cy7-H | FSC-H | 15,526 | 1,800,582 |
| Live sfGFP Negative | 661 | 91.81 % | FITC-A | Count | 84,851 | |
| Live sfGFP Positive | 59 | 8.19 % | FITC-A | Count | 8,153,970 | |
| Dead | 2,157 | 74.92 % | APC-Cy7-H | FSC-H | 78,219 | 1,533,593 |
| Dead sfGFP Negative | 2,143 | 99.35 % | FITC-A | Count | 104,864 | |
| Dead sfGFP Positive | 13 | 0.60 % | FITC-A | Count | 4,530,115 | |
| sfGFP Negative | 2,812 | 97.67 % | FITC-A | Count | 100,010 | |
| sfGFP Positive | 65 | 2.26 % | FITC-A | Count | 8,587,649 | |

#### p059 mRNA [100 ng] + TET [1 uM]

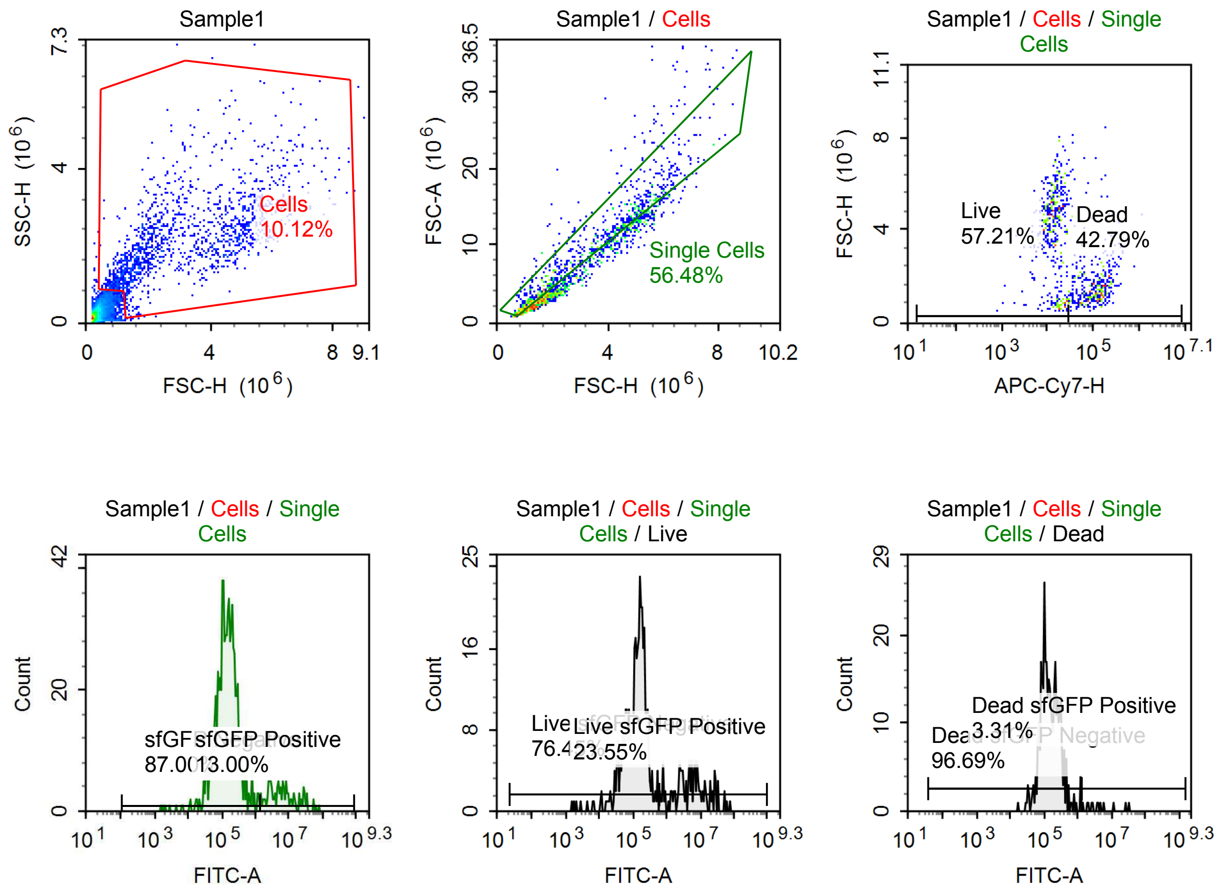

Sample Statistics of Sample1

| Gate | Count | % Parent | X | Y | Median X | Median Y |
|---|---|---|---|---|---|---|
| All | 14,796 | | | | | |
| Cells | 1,498 | 10.12 % | FSC-H | SSC-H | 2,606,639 | 2,063,246 |
| Single Cells | 846 | 56.48 % | FSC-H | FSC-A | 2,633,609 | 7,786,895 |
| Live | 484 | 57.21 % | APC-Cy7-H | FSC-H | 13,662 | 4,357,177 |
| Live sfGFP Negative | 370 | 76.45 % | FITC-A | Count | 134,017 | |
| Live sfGFP Positive | 114 | 23.55 % | FITC-A | Count | 5,485,342 | |
| Dead | 362 | 42.79 % | APC-Cy7-H | FSC-H | 118,897 | 1,581,604 |
| Dead sfGFP Negative | 350 | 96.69 % | FITC-A | Count | 137,796 | |
| Dead sfGFP Positive | 12 | 3.31 % | FITC-A | Count | 3,959,733 | |
| sfGFP Negative | 736 | 87.00 % | FITC-A | Count | 139,030 | |
| sfGFP Positive | 110 | 13.00 % | FITC-A | Count | 6,370,041 | |

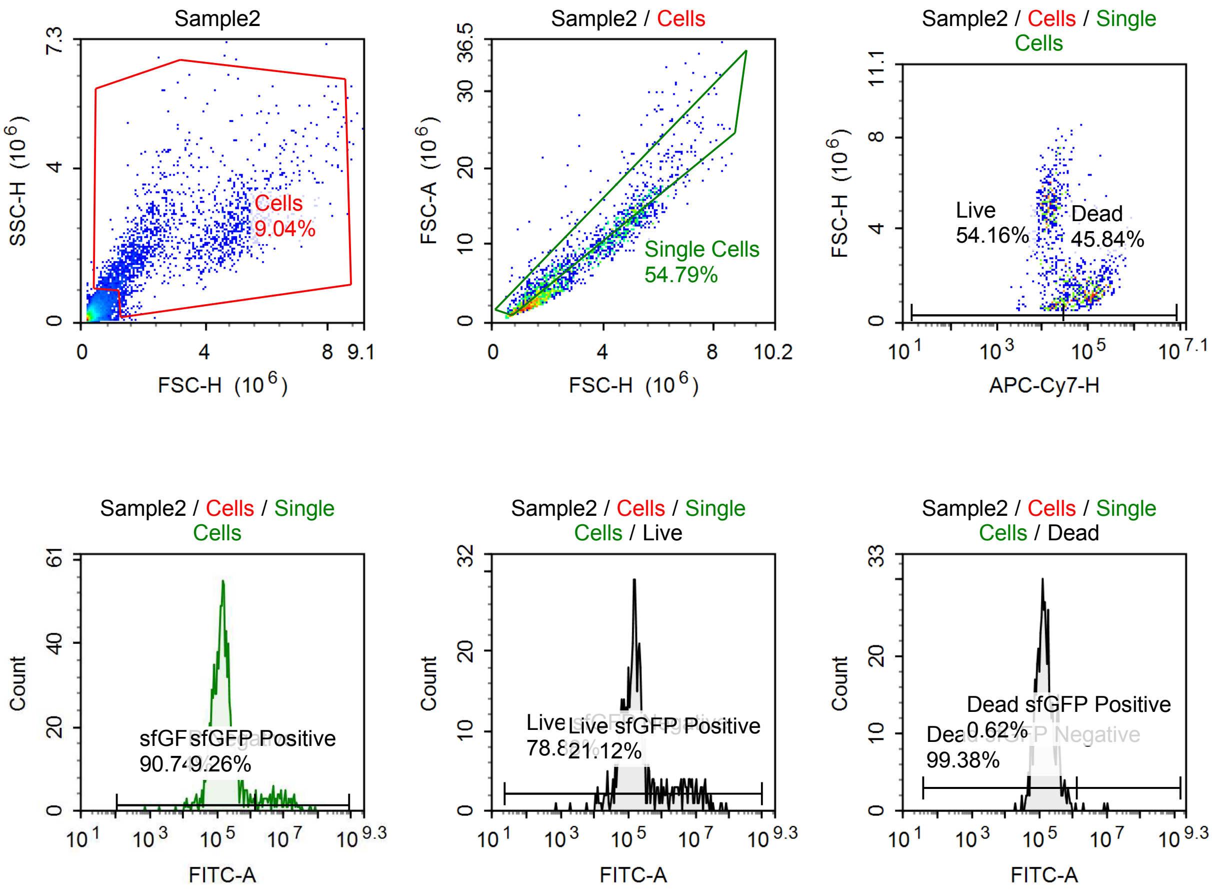

Sample Statistics of Sample2

| Gate | Count | % Parent | X | Y | Median X | Median Y |
|---|---|---|---|---|---|---|
| All | 21,372 | | | | | |
| Cells | 1,931 | 9.04 % | FSC-H | SSC-H | 2,361,355 | 1,987,715 |
| Single Cells | 1,058 | 54.79 % | FSC-H | FSC-A | 2,178,515 | 6,263,531 |
| Live | 573 | 54.16 % | APC-Cy7-H | FSC-H | 14,489 | 4,260,781 |
| Live sfGFP Negative | 452 | 78.88 % | FITC-A | Count | 139,448 | |
| Live sfGFP Positive | 121 | 21.12 % | FITC-A | Count | 4,837,531 | |
| Dead | 485 | 45.84 % | APC-Cy7-H | FSC-H | 96,452 | 1,440,434 |
| Dead sfGFP Negative | 482 | 99.38 % | FITC-A | Count | 142,014 | |
| Dead sfGFP Positive | 3 | 0.62 % | FITC-A | Count | 8,355,424 | |
| sfGFP Negative | 960 | 90.74 % | FITC-A | Count | 143,389 | |
| sfGFP Positive | 98 | 9.26 % | FITC-A | Count | 6,685,743 | |

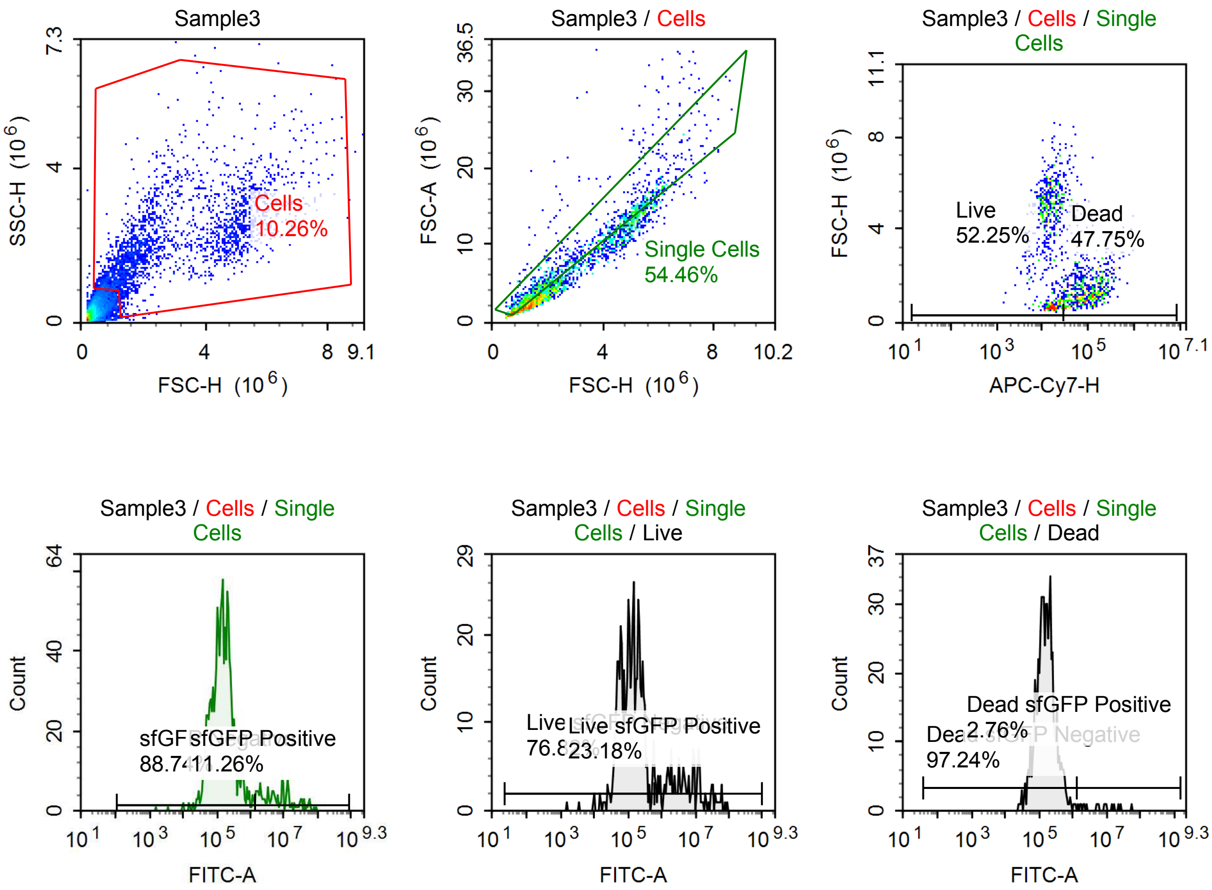

Sample Statistics of Sample3

| Gate | Count | % Parent | X | Y | Median X | Median Y |
|---|---|---|---|---|---|---|
| All | 23,045 | | | | | |
| Cells | 2,365 | 10.26 % | FSC-H | SSC-H | 2,229,069 | 1,982,859 |
| Single Cells | 1,288 | 54.46 % | FSC-H | FSC-A | 1,922,774 | 5,335,078 |
| Live | 673 | 52.25 % | APC-Cy7-H | FSC-H | 14,307 | 4,141,715 |
| Live sfGFP Negative | 517 | 76.82 % | FITC-A | Count | 123,198 | |
| Live sfGFP Positive | 156 | 23.18 % | FITC-A | Count | 4,714,463 | |
| Dead | 615 | 47.75 % | APC-Cy7-H | FSC-H | 92,142 | 1,431,436 |
| Dead sfGFP Negative | 598 | 97.24 % | FITC-A | Count | 154,805 | |
| Dead sfGFP Positive | 17 | 2.76 % | FITC-A | Count | 5,717,252 | |
| sfGFP Negative | 1,143 | 88.74 % | FITC-A | Count | 143,644 | |
| sfGFP Positive | 145 | 11.26 % | FITC-A | Count | 6,591,267 | |

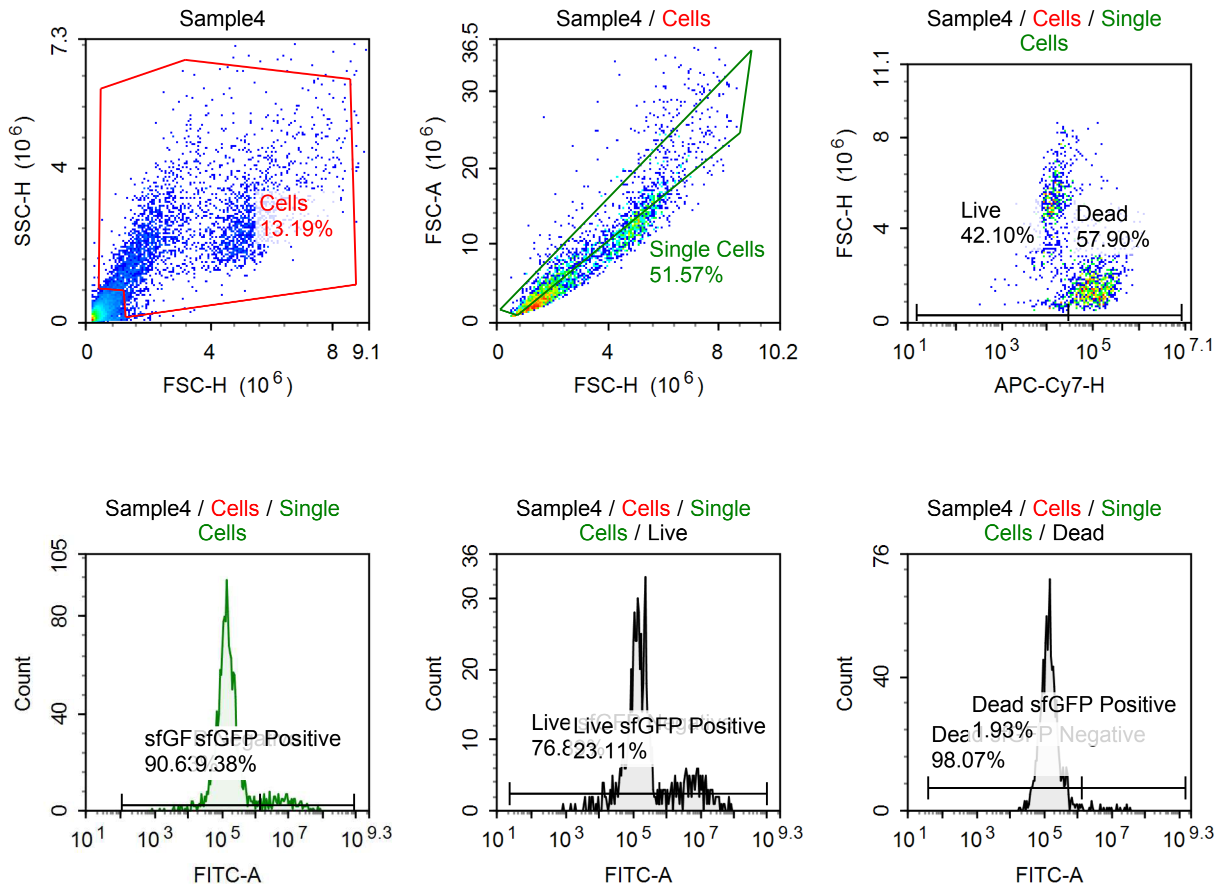

Sample Statistics of Sample4

| Gate | Count | % Parent | X | Y | Median X | Median Y |
|---|---|---|---|---|---|---|
| All | 24,945 | | | | | |
| Cells | 3,289 | 13.19 % | FSC-H | SSC-H | 2,581,409 | 2,120,134 |
| Single Cells | 1,696 | 51.57 % | FSC-H | FSC-A | 2,137,725 | 6,224,110 |
| Live | 714 | 42.10 % | APC-Cy7-H | FSC-H | 13,154 | 4,801,872 |
| Live sfGFP Negative | 549 | 76.89 % | FITC-A | Count | 137,239 | |
| Live sfGFP Positive | 165 | 23.11 % | FITC-A | Count | 5,821,768 | |
| Dead | 982 | 57.90 % | APC-Cy7-H | FSC-H | 99,156 | 1,587,302 |
| Dead sfGFP Negative | 963 | 98.07 % | FITC-A | Count | 137,534 | |
| Dead sfGFP Positive | 19 | 1.93 % | FITC-A | Count | 7,076,098 | |
| sfGFP Negative | 1,537 | 90.63 % | FITC-A | Count | 138,517 | |
| sfGFP Positive | 159 | 9.38 % | FITC-A | Count | 7,414,022 | |

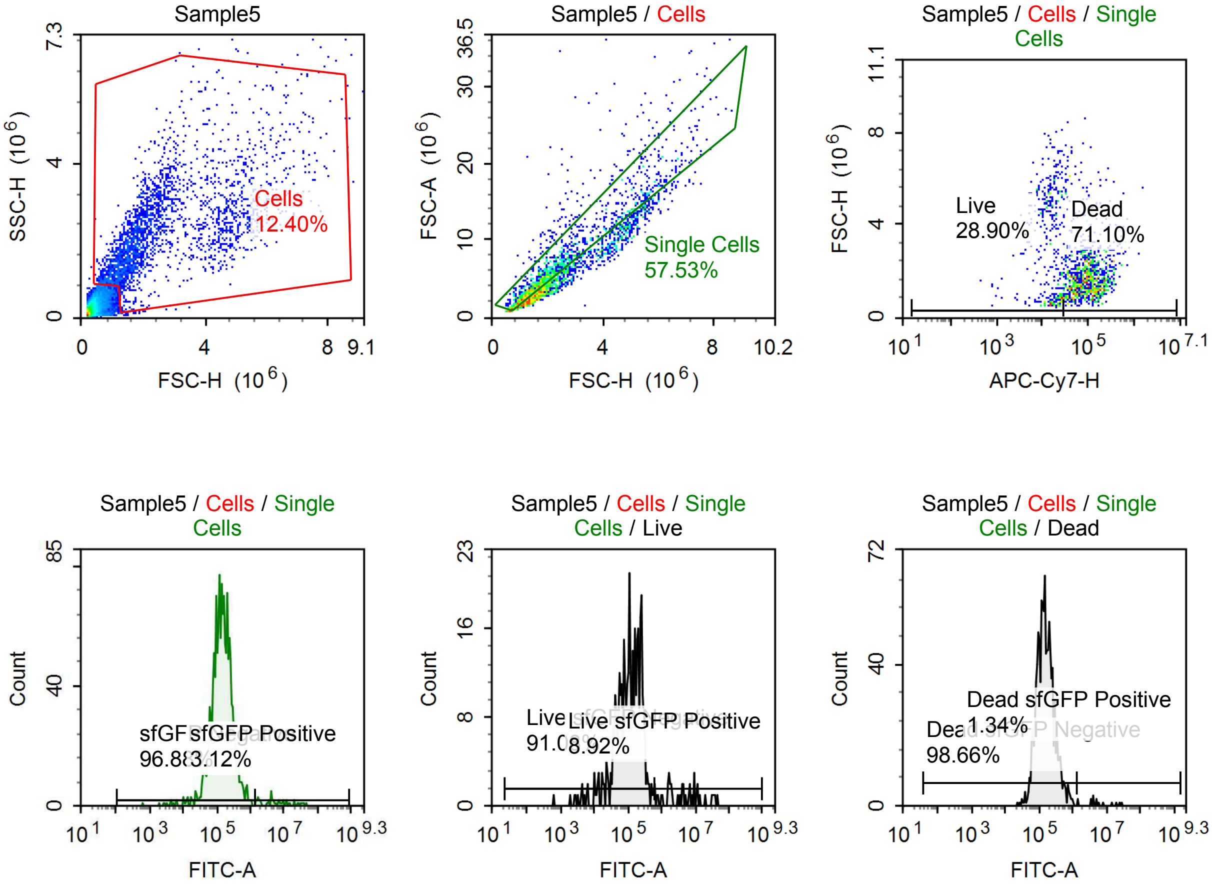

Sample Statistics of Sample5

| Gate | Count | % Parent | X | Y | Median X | Median Y |
|---|---|---|---|---|---|---|
| All | 20,654 | | | | | |
| Cells | 2,562 | 12.40 % | FSC-H | SSC-H | 2,003,440 | 2,076,767 |
| Single Cells | 1,474 | 57.53 % | FSC-H | FSC-A | 1,738,463 | 4,744,911 |
| Live | 426 | 28.90 % | APC-Cy7-H | FSC-H | 15,726 | 3,240,560 |
| Live sfGFP Negative | 388 | 91.08 % | FITC-A | Count | 111,147 | |
| Live sfGFP Positive | 38 | 8.92 % | FITC-A | Count | 4,489,199 | |
| Dead | 1,048 | 71.10 % | APC-Cy7-H | FSC-H | 94,475 | 1,647,843 |
| Dead sfGFP Negative | 1,034 | 98.66 % | FITC-A | Count | 147,685 | |
| Dead sfGFP Positive | 14 | 1.34 % | FITC-A | Count | 4,982,726 | |
| sfGFP Negative | 1,428 | 96.88 % | FITC-A | Count | 141,883 | |
| sfGFP Positive | 46 | 3.12 % | FITC-A | Count | 5,328,311 | |

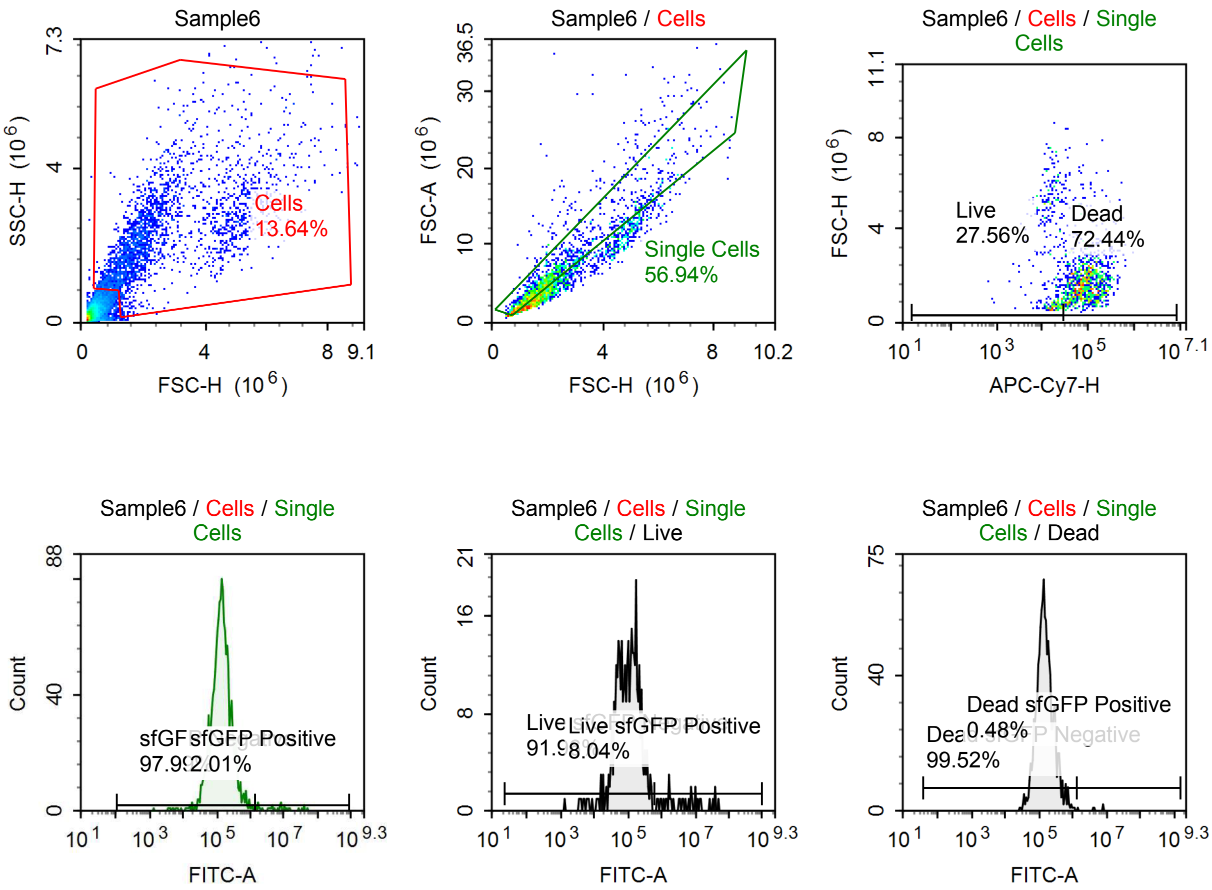

Sample Statistics of Sample6

| Gate | Count | % Parent | X | Y | Median X | Median Y |
|---|---|---|---|---|---|---|
| All | 18,596 | | | | | |
| Cells | 2,536 | 13.64 % | FSC-H | SSC-H | 1,990,415 | 2,124,025 |
| Single Cells | 1,444 | 56.94 % | FSC-H | FSC-A | 1,731,763 | 4,769,100 |
| Live | 398 | 27.56 % | APC-Cy7-H | FSC-H | 16,230 | 2,272,138 |
| Live sfGFP Negative | 366 | 91.96 % | FITC-A | Count | 99,034 | |
| Live sfGFP Positive | 32 | 8.04 % | FITC-A | Count | 4,311,082 | |
| Dead | 1,046 | 72.44 % | APC-Cy7-H | FSC-H | 89,000 | 1,701,429 |
| Dead sfGFP Negative | 1,041 | 99.52 % | FITC-A | Count | 143,032 | |
| Dead sfGFP Positive | 5 | 0.48 % | FITC-A | Count | 4,387,567 | |
| sfGFP Negative | 1,415 | 97.99 % | FITC-A | Count | 136,637 | |
| sfGFP Positive | 29 | 2.01 % | FITC-A | Count | 7,359,505 | |

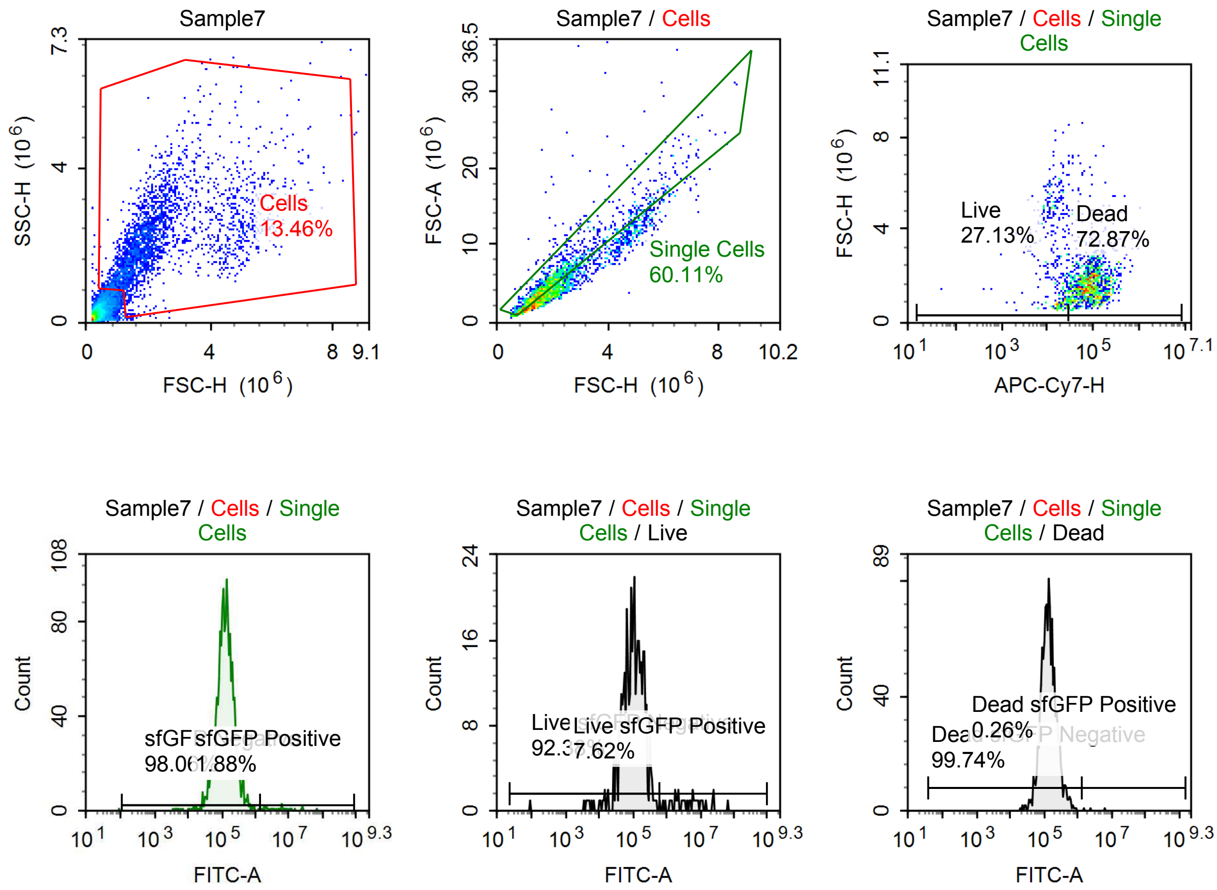

Sample Statistics of Sample7

| Gate | Count | % Parent | X | Y | Median X | Median Y |
|---|---|---|---|---|---|---|
| All | 19,727 | | | | | |
| Cells | 2,655 | 13.46 % | FSC-H | SSC-H | 1,901,604 | 2,030,995 |
| Single Cells | 1,596 | 60.11 % | FSC-H | FSC-A | 1,707,041 | 4,520,864 |
| Live | 433 | 27.13 % | APC-Cy7-H | FSC-H | 16,240 | 2,225,203 |
| Live sfGFP Negative | 400 | 92.38 % | FITC-A | Count | 102,613 | |
| Live sfGFP Positive | 33 | 7.62 % | FITC-A | Count | 4,741,506 | |
| Dead | 1,163 | 72.87 % | APC-Cy7-H | FSC-H | 89,063 | 1,643,928 |
| Dead sfGFP Negative | 1,160 | 99.74 % | FITC-A | Count | 134,234 | |
| Dead sfGFP Positive | 3 | 0.26 % | FITC-A | Count | 2,414,825 | |
| sfGFP Negative | 1,565 | 98.06 % | FITC-A | Count | 127,395 | |
| sfGFP Positive | 30 | 1.88 % | FITC-A | Count | 6,030,777 | |

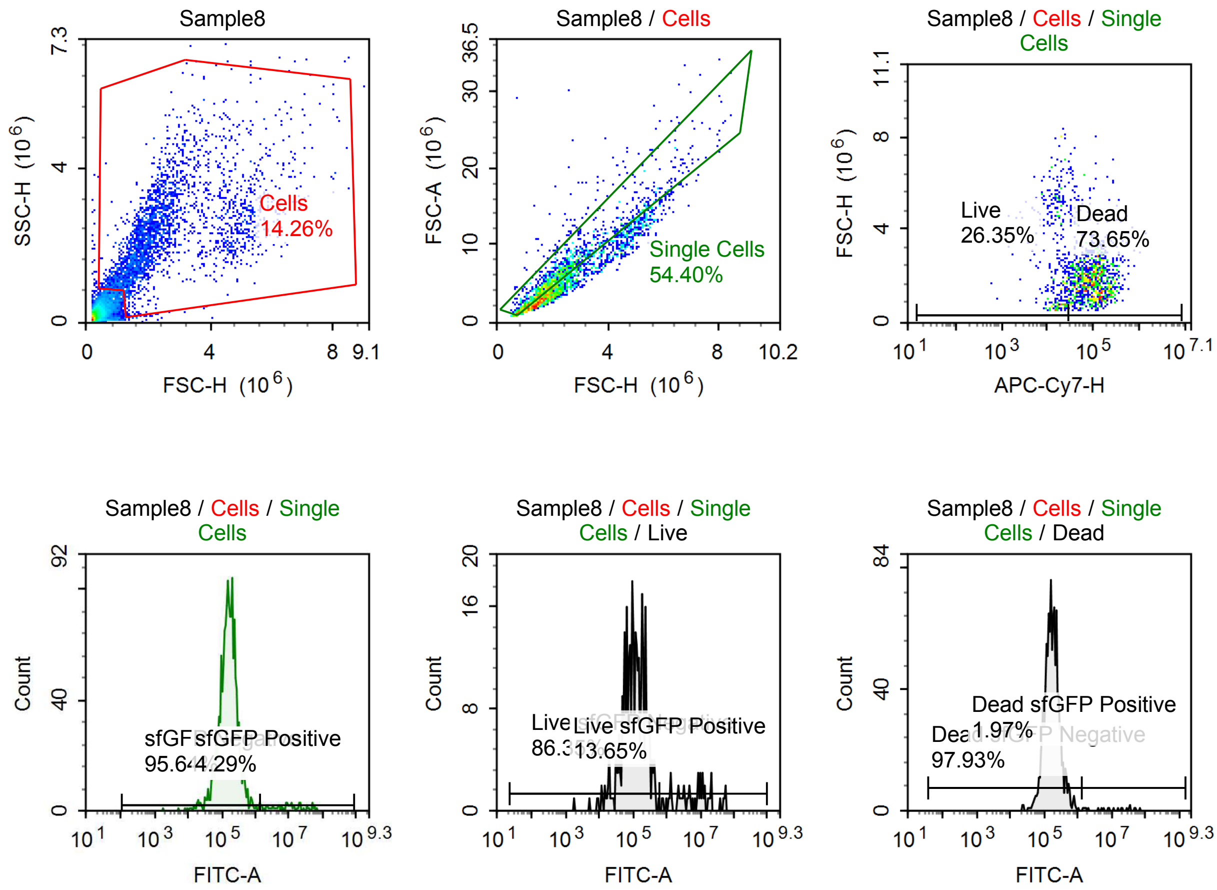

Sample Statistics of Sample8

| Gate | Count | % Parent | X | Y | Median X | Median Y |
|---|---|---|---|---|---|---|
| All | 18,645 | | | | | |
| Cells | 2,658 | 14.26 % | FSC-H | SSC-H | 2,100,578 | 2,164,245 |
| Single Cells | 1,446 | 54.40 % | FSC-H | FSC-A | 1,923,336 | 5,340,372 |
| Live | 381 | 26.35 % | APC-Cy7-H | FSC-H | 16,835 | 2,494,955 |
| Live sfGFP Negative | 329 | 86.35 % | FITC-A | Count | 105,451 | |
| Live sfGFP Positive | 52 | 13.65 % | FITC-A | Count | 7,578,843 | |
| Dead | 1,065 | 73.65 % | APC-Cy7-H | FSC-H | 98,974 | 1,861,796 |
| Dead sfGFP Negative | 1,043 | 97.93 % | FITC-A | Count | 167,916 | |
| Dead sfGFP Positive | 21 | 1.97 % | FITC-A | Count | 11,613,558 | |
| sfGFP Negative | 1,383 | 95.64 % | FITC-A | Count | 159,094 | |
| sfGFP Positive | 62 | 4.29 % | FITC-A | Count | 9,886,854 | |

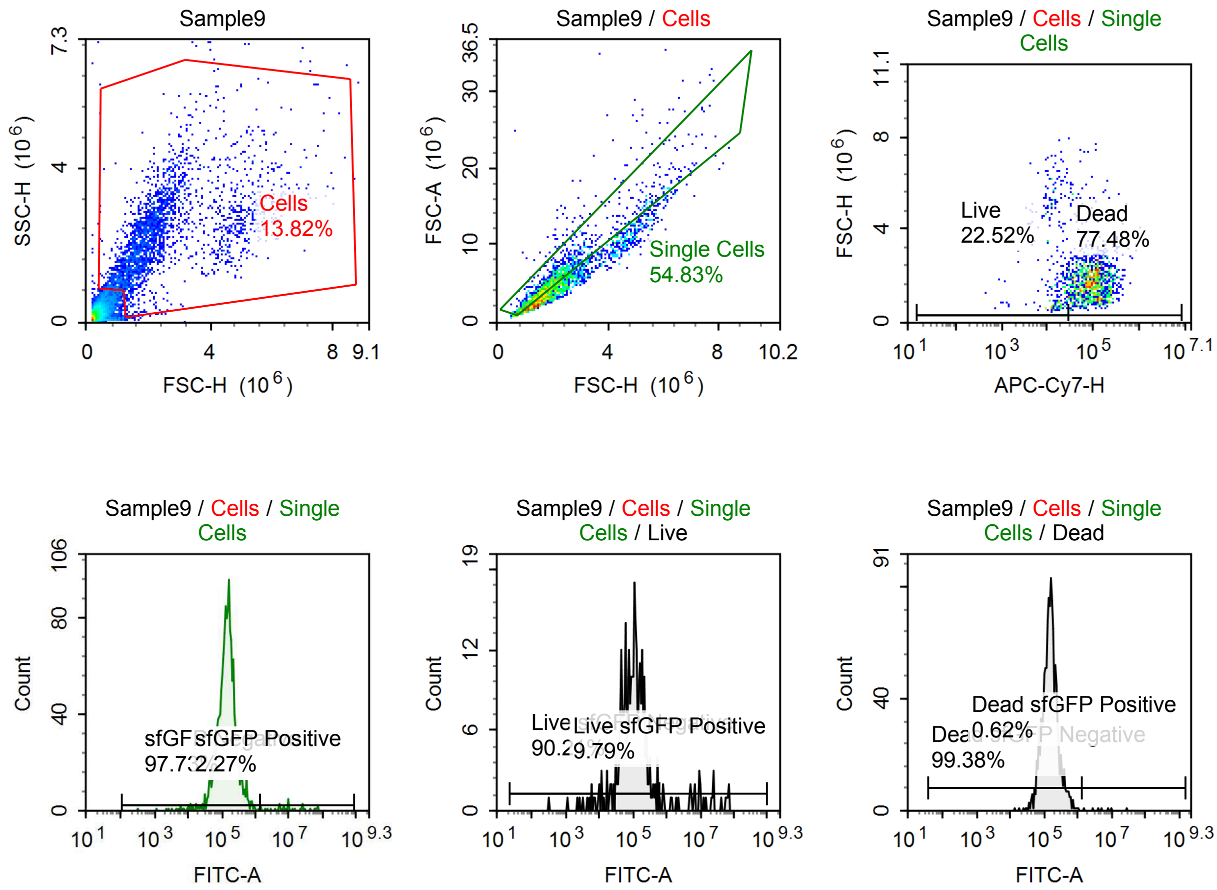

Sample Statistics of Sample9

| Gate | Count | % Parent | X | Y | Median X | Median Y |
|---|---|---|---|---|---|---|
| All | 19,156 | | | | | |
| Cells | 2,648 | 13.82 % | FSC-H | SSC-H | 2,002,356 | 2,128,326 |
| Single Cells | 1,452 | 54.83 % | FSC-H | FSC-A | 1,761,924 | 4,640,533 |
| Live | 327 | 22.52 % | APC-Cy7-H | FSC-H | 15,721 | 1,908,422 |
| Live sfGFP Negative | 295 | 90.21 % | FITC-A | Count | 99,726 | |
| Live sfGFP Positive | 32 | 9.79 % | FITC-A | Count | 9,551,002 | |
| Dead | 1,125 | 77.48 % | APC-Cy7-H | FSC-H | 96,808 | 1,737,897 |
| Dead sfGFP Negative | 1,118 | 99.38 % | FITC-A | Count | 151,660 | |
| Dead sfGFP Positive | 7 | 0.62 % | FITC-A | Count | 5,015,103 | |
| sfGFP Negative | 1,419 | 97.73 % | FITC-A | Count | 142,891 | |
| sfGFP Positive | 33 | 2.27 % | FITC-A | Count | 9,785,818 | |

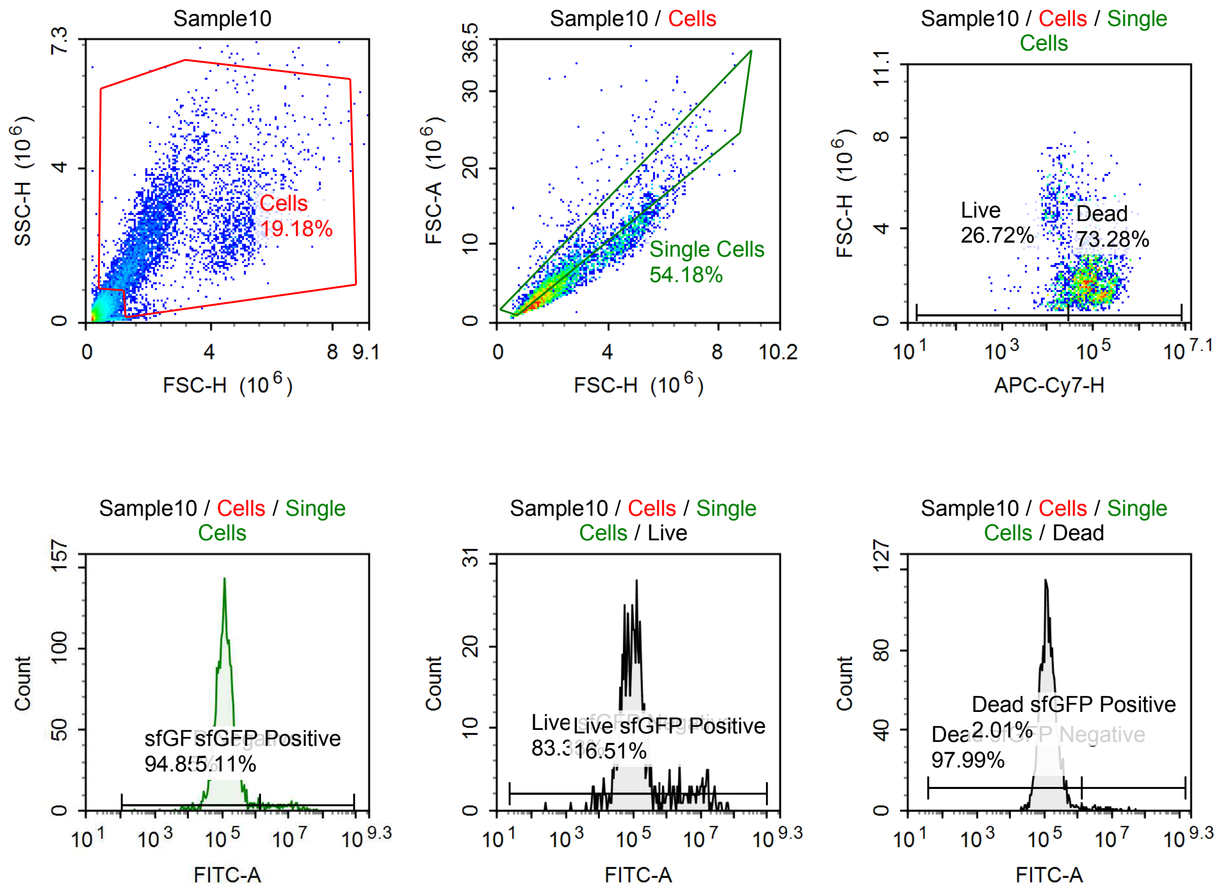

Sample Statistics of Sample10

| Gate | Count | % Parent | X | Y | Median X | Median Y |
|---|---|---|---|---|---|---|
| All | 23,559 | | | | | |
| Cells | 4,518 | 19.18 % | FSC-H | SSC-H | 2,059,844 | 2,107,653 |
| Single Cells | 2,448 | 54.18 % | FSC-H | FSC-A | 1,801,889 | 4,795,033 |
| Live | 654 | 26.72 % | APC-Cy7-H | FSC-H | 16,726 | 2,448,418 |
| Live sfGFP Negative | 545 | 83.33 % | FITC-A | Count | 93,590 | |
| Live sfGFP Positive | 108 | 16.51 % | FITC-A | Count | 5,814,489 | |
| Dead | 1,794 | 73.28 % | APC-Cy7-H | FSC-H | 87,879 | 1,737,315 |
| Dead sfGFP Negative | 1,758 | 97.99 % | FITC-A | Count | 126,888 | |
| Dead sfGFP Positive | 36 | 2.01 % | FITC-A | Count | 5,136,222 | |
| sfGFP Negative | 2,322 | 94.85 % | FITC-A | Count | 122,676 | |
| sfGFP Positive | 125 | 5.11 % | FITC-A | Count | 6,849,712 | |

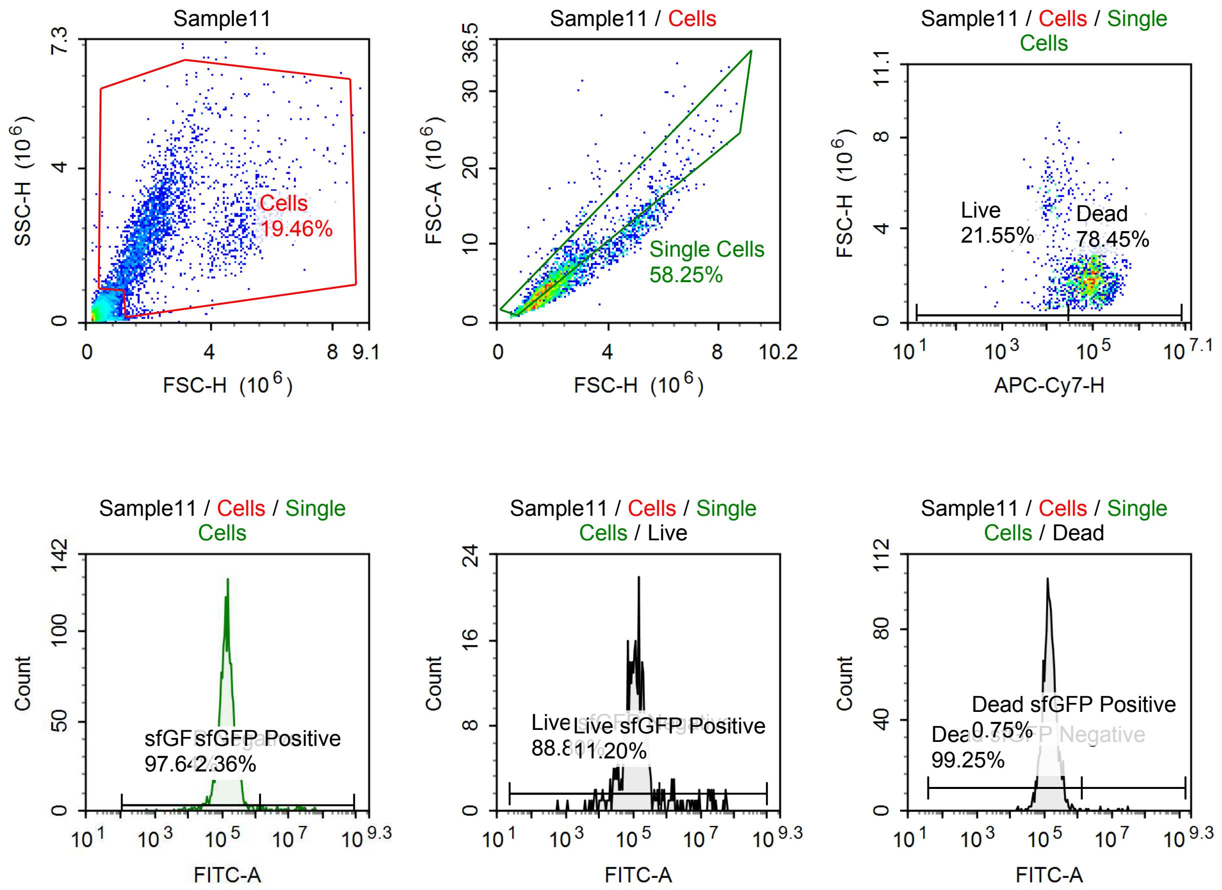

Sample Statistics of Sample11

| Gate | Count | % Parent | X | Y | Median X | Median Y |
|---|---|---|---|---|---|---|
| All | 14,976 | | | | | |
| Cells | 2,915 | 19.46 % | FSC-H | SSC-H | 2,089,099 | 2,302,912 |
| Single Cells | 1,698 | 58.25 % | FSC-H | FSC-A | 1,882,274 | 5,095,742 |
| Live | 366 | 21.55 % | APC-Cy7-H | FSC-H | 15,000 | 2,406,266 |
| Live sfGFP Negative | 325 | 88.80 % | FITC-A | Count | 111,593 | |
| Live sfGFP Positive | 41 | 11.20 % | FITC-A | Count | 4,795,918 | |
| Dead | 1,332 | 78.45 % | APC-Cy7-H | FSC-H | 93,115 | 1,846,294 |
| Dead sfGFP Negative | 1,322 | 99.25 % | FITC-A | Count | 142,964 | |
| Dead sfGFP Positive | 10 | 0.75 % | FITC-A | Count | 11,501,122 | |
| sfGFP Negative | 1,658 | 97.64 % | FITC-A | Count | 138,813 | |
| sfGFP Positive | 40 | 2.36 % | FITC-A | Count | 9,370,266 | |

#### p059 mRNA [100 ng] + TET [10 uM]

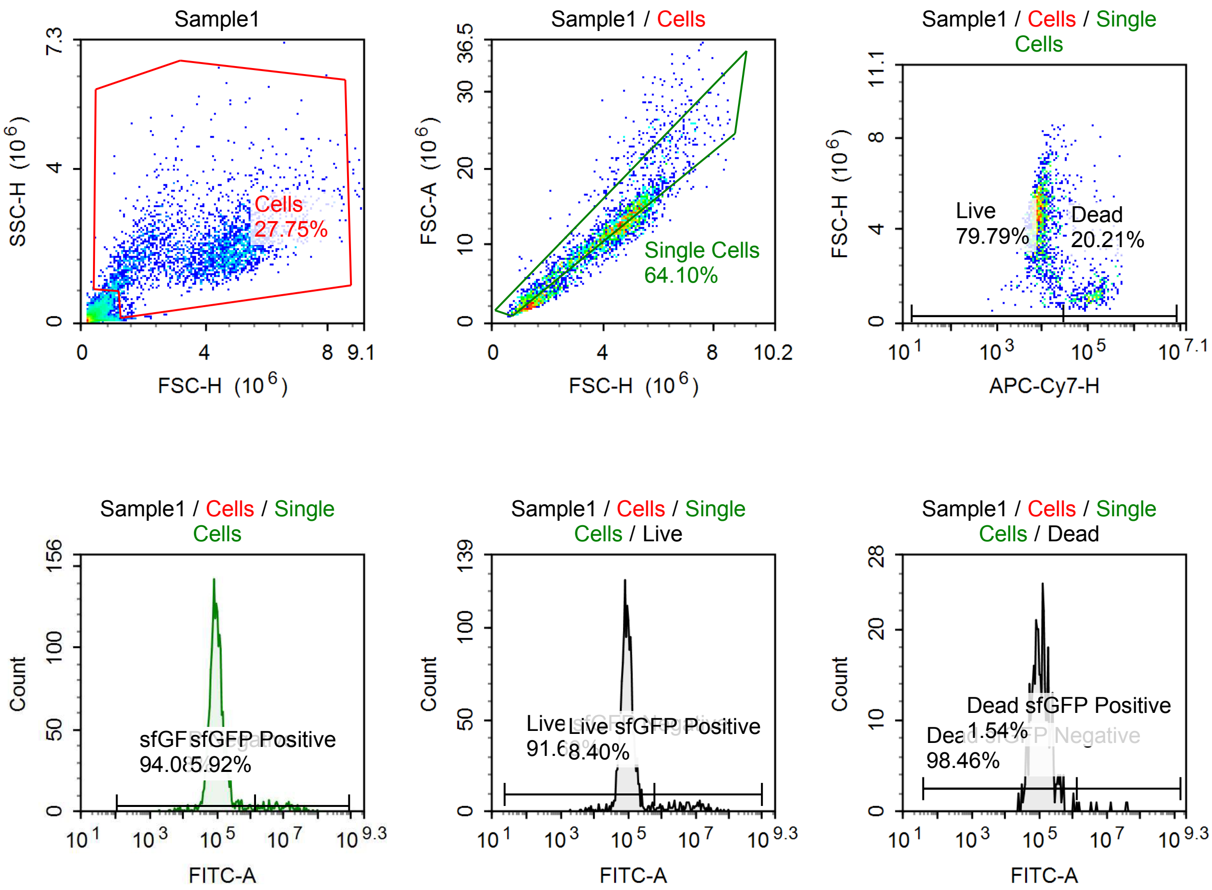

Sample Statistics of Sample1

| Gate | Count | % Parent | X | Y | Median X | Median Y |
|---|---|---|---|---|---|---|
| All | 10,821 | | | | | |
| Cells | 3,003 | 27.75 % | FSC-H | SSC-H | 4,193,729 | 1,852,379 |
| Single Cells | 1,925 | 64.10 % | FSC-H | FSC-A | 4,144,412 | 11,923,613 |
| Live | 1,536 | 79.79 % | APC-Cy7-H | FSC-H | 8,472 | 4,499,424 |
| Live sfGFP Negative | 1,407 | 91.60 % | FITC-A | Count | 93,187 | |
| Live sfGFP Positive | 129 | 8.40 % | FITC-A | Count | 5,912,794 | |
| Dead | 389 | 20.21 % | APC-Cy7-H | FSC-H | 121,544 | 1,345,753 |
| Dead sfGFP Negative | 383 | 98.46 % | FITC-A | Count | 109,824 | |
| Dead sfGFP Positive | 6 | 1.54 % | FITC-A | Count | 9,267,321 | |
| sfGFP Negative | 1,811 | 94.08 % | FITC-A | Count | 95,589 | |
| sfGFP Positive | 114 | 5.92 % | FITC-A | Count | 7,016,149 | |

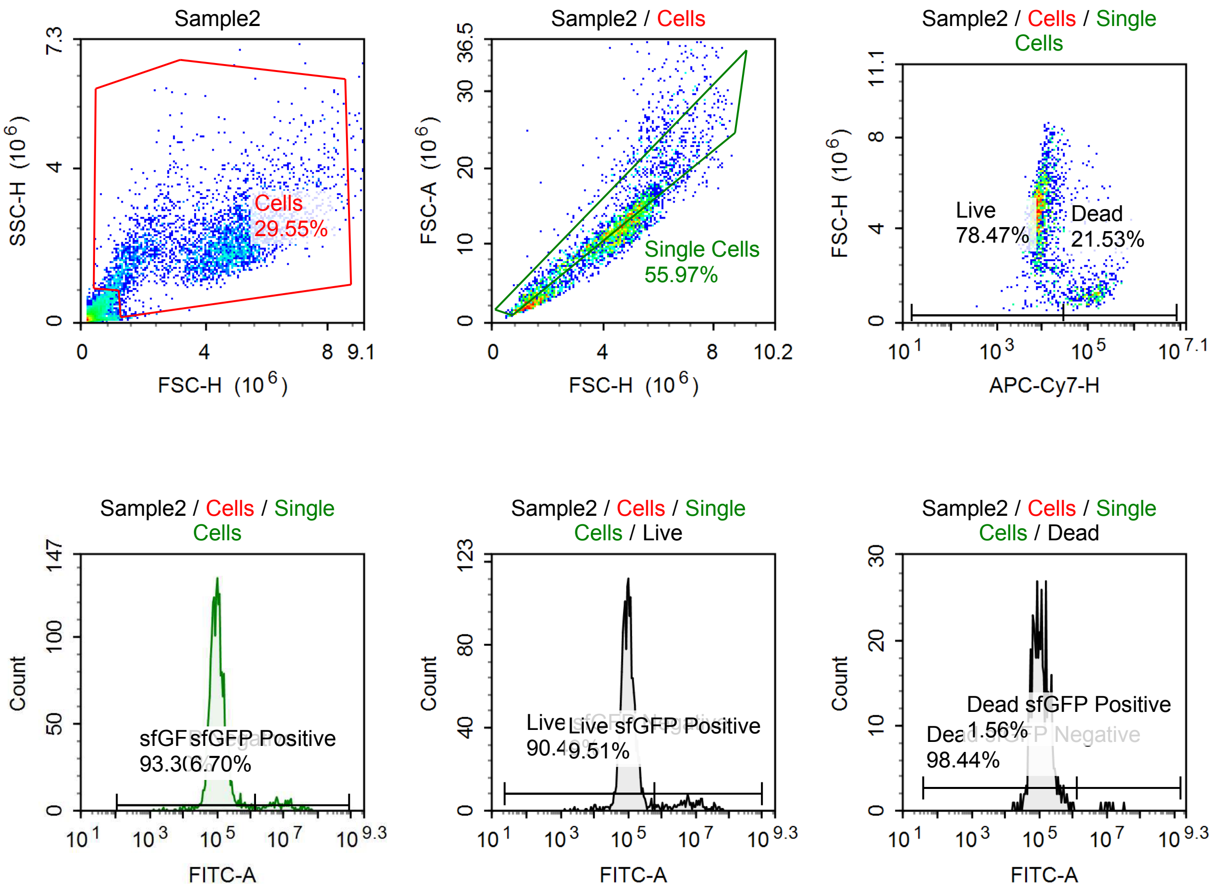

Sample Statistics of Sample2

| Gate | Count | % Parent | X | Y | Median X | Median Y |
|---|---|---|---|---|---|---|
| All | 12,635 | | | | | |
| Cells | 3,734 | 29.55 % | FSC-H | SSC-H | 4,411,761 | 1,964,809 |
| Single Cells | 2,090 | 55.97 % | FSC-H | FSC-A | 4,294,353 | 12,094,862 |
| Live | 1,640 | 78.47 % | APC-Cy7-H | FSC-H | 8,613 | 4,698,252 |
| Live sfGFP Negative | 1,484 | 90.49 % | FITC-A | Count | 99,191 | |
| Live sfGFP Positive | 156 | 9.51 % | FITC-A | Count | 6,191,044 | |
| Dead | 450 | 21.53 % | APC-Cy7-H | FSC-H | 115,864 | 1,370,697 |
| Dead sfGFP Negative | 443 | 98.44 % | FITC-A | Count | 108,390 | |
| Dead sfGFP Positive | 7 | 1.56 % | FITC-A | Count | 11,394,405 | |
| sfGFP Negative | 1,950 | 93.30 % | FITC-A | Count | 101,999 | |
| sfGFP Positive | 140 | 6.70 % | FITC-A | Count | 7,984,583 | |

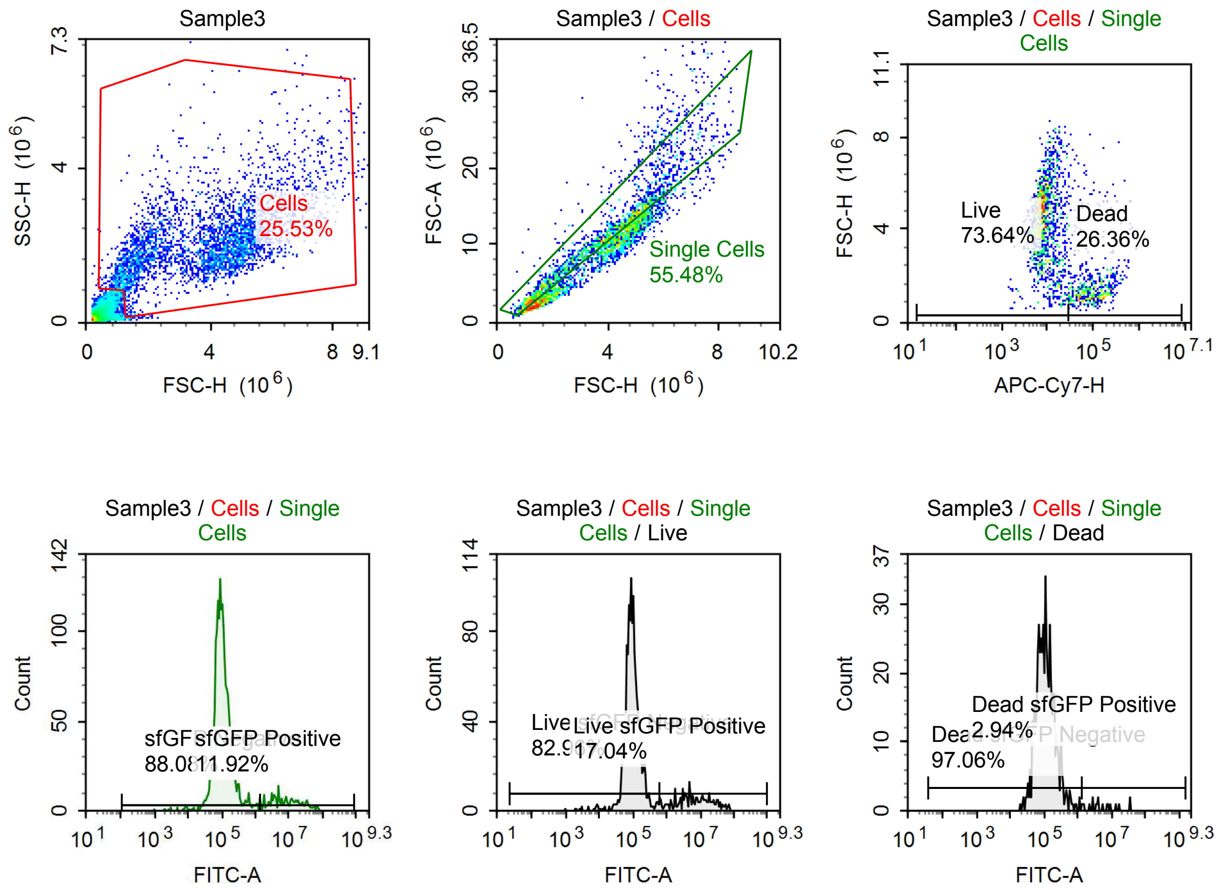

Sample Statistics of Sample3

| Gate | Count | % Parent | X | Y | Median X | Median Y |
|---|---|---|---|---|---|---|
| All | 14,571 | | | | | |
| Cells | 3,720 | 25.53 % | FSC-H | SSC-H | 4,065,825 | 1,982,418 |
| Single Cells | 2,064 | 55.48 % | FSC-H | FSC-A | 3,820,670 | 10,818,302 |
| Live | 1,520 | 73.64 % | APC-Cy7-H | FSC-H | 8,171 | 4,432,926 |
| Live sfGFP Negative | 1,261 | 82.96 % | FITC-A | Count | 93,770 | |
| Live sfGFP Positive | 259 | 17.04 % | FITC-A | Count | 6,632,680 | |
| Dead | 544 | 26.36 % | APC-Cy7-H | FSC-H | 93,386 | 1,402,968 |
| Dead sfGFP Negative | 528 | 97.06 % | FITC-A | Count | 108,184 | |
| Dead sfGFP Positive | 16 | 2.94 % | FITC-A | Count | 4,815,224 | |
| sfGFP Negative | 1,818 | 88.08 % | FITC-A | Count | 98,185 | |
| sfGFP Positive | 246 | 11.92 % | FITC-A | Count | 7,998,266 | |

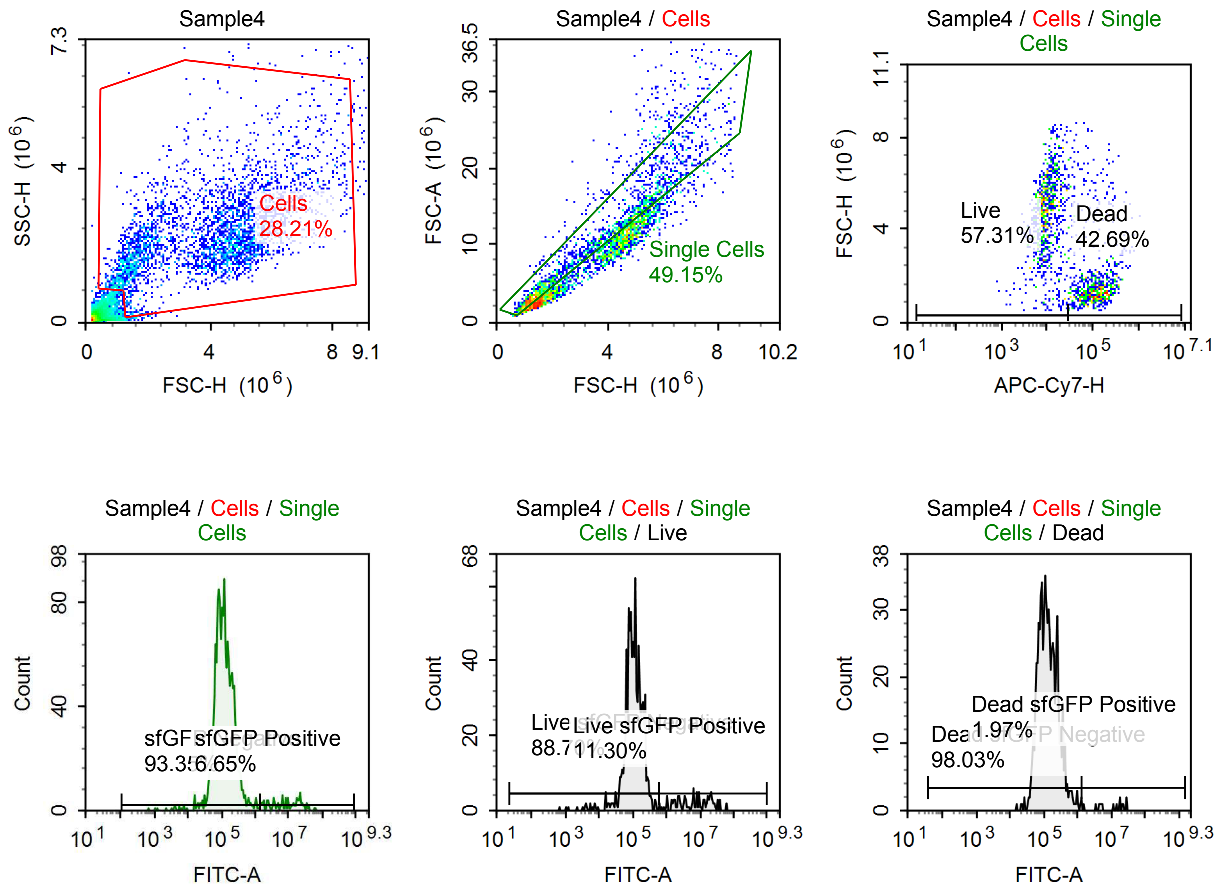

Sample Statistics of Sample4

| Gate | Count | % Parent | X | Y | Median X | Median Y |
|---|---|---|---|---|---|---|
| All | 12,032 | | | | | |
| Cells | 3,394 | 28.21 % | FSC-H | SSC-H | 4,133,035 | 2,111,166 |
| Single Cells | 1,668 | 49.15 % | FSC-H | FSC-A | 3,465,598 | 10,205,595 |
| Live | 956 | 57.31 % | APC-Cy7-H | FSC-H | 9,424 | 4,840,827 |
| Live sfGFP Negative | 848 | 88.70 % | FITC-A | Count | 107,402 | |
| Live sfGFP Positive | 108 | 11.30 % | FITC-A | Count | 7,695,671 | |
| Dead | 712 | 42.69 % | APC-Cy7-H | FSC-H | 112,439 | 1,468,702 |
| Dead sfGFP Negative | 698 | 98.03 % | FITC-A | Count | 125,252 | |
| Dead sfGFP Positive | 14 | 1.97 % | FITC-A | Count | 12,998,542 | |
| sfGFP Negative | 1,557 | 93.35 % | FITC-A | Count | 115,570 | |
| sfGFP Positive | 111 | 6.65 % | FITC-A | Count | 9,325,064 | |

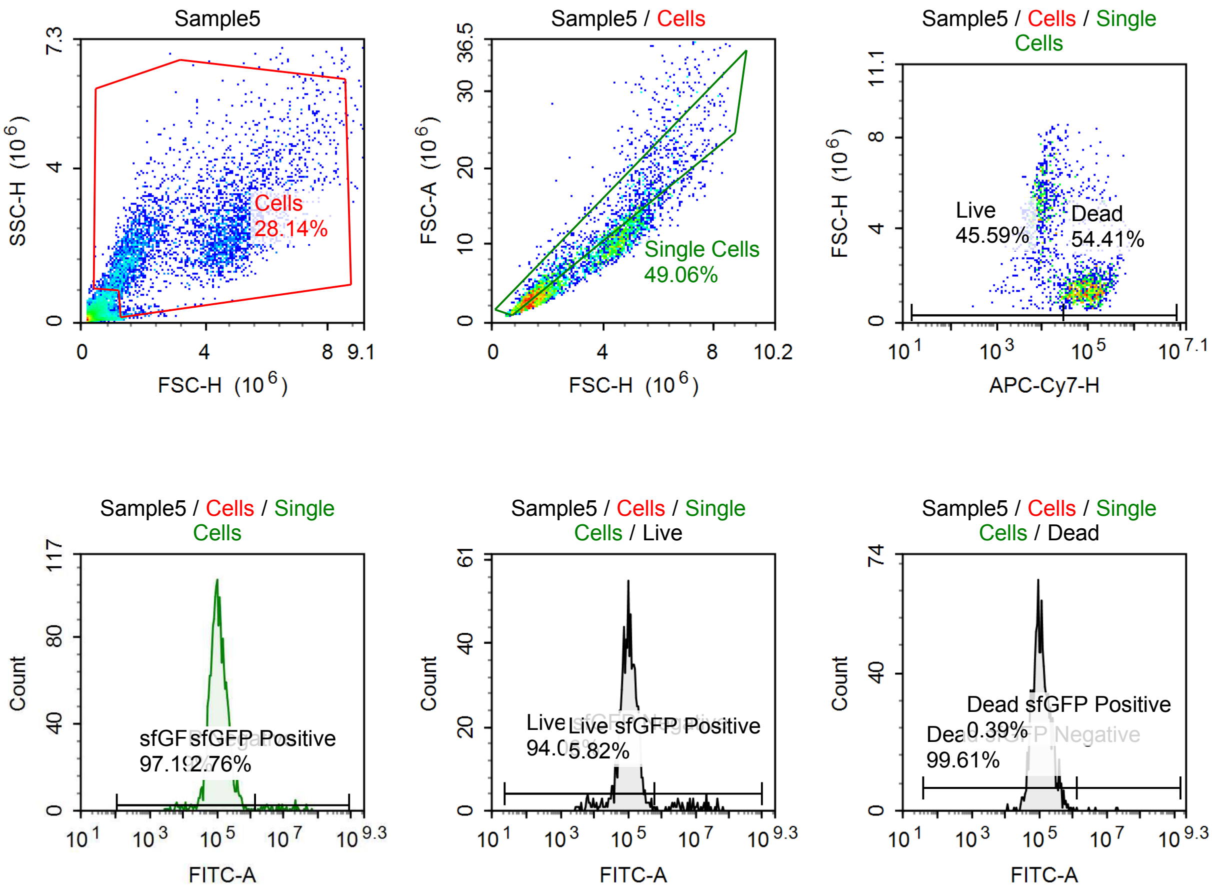

Sample Statistics of Sample5

| Gate | Count | % Parent | X | Y | Median X | Median Y |
|---|---|---|---|---|---|---|
| All | 13,648 | | | | | |
| Cells | 3,840 | 28.14 % | FSC-H | SSC-H | 3,404,537 | 2,122,632 |
| Single Cells | 1,884 | 49.06 % | FSC-H | FSC-A | 2,029,831 | 5,832,629 |
| Live | 859 | 45.59 % | APC-Cy7-H | FSC-H | 9,670 | 4,500,193 |
| Live sfGFP Negative | 808 | 94.06 % | FITC-A | Count | 104,448 | |
| Live sfGFP Positive | 50 | 5.82 % | FITC-A | Count | 7,678,457 | |
| Dead | 1,025 | 54.41 % | APC-Cy7-H | FSC-H | 99,663 | 1,498,009 |
| Dead sfGFP Negative | 1,021 | 99.61 % | FITC-A | Count | 112,439 | |
| Dead sfGFP Positive | 4 | 0.39 % | FITC-A | Count | 12,255,294 | |
| sfGFP Negative | 1,831 | 97.19 % | FITC-A | Count | 107,320 | |
| sfGFP Positive | 52 | 2.76 % | FITC-A | Count | 8,331,650 | |

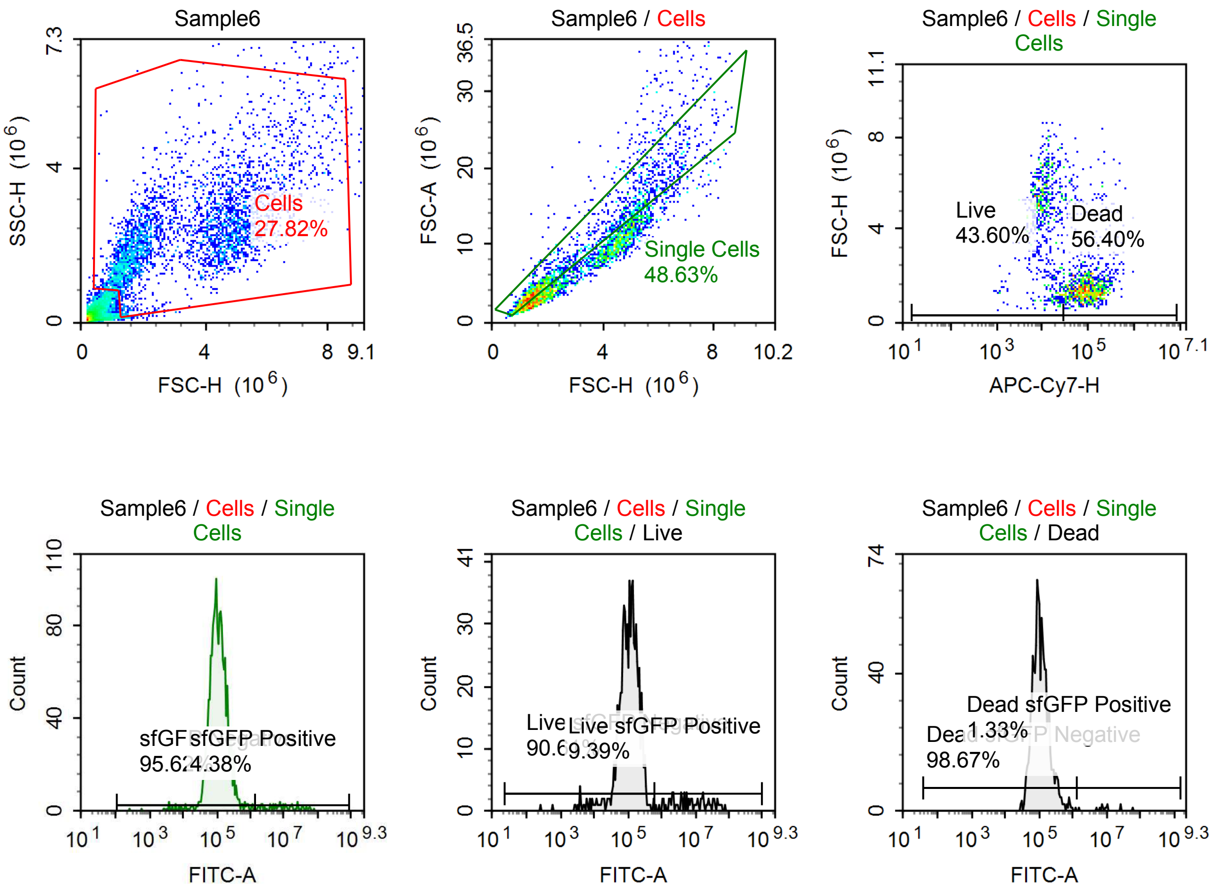

Sample Statistics of Sample6

| Gate | Count | % Parent | X | Y | Median X | Median Y |
|---|---|---|---|---|---|---|
| All | 12,817 | | | | | |
| Cells | 3,566 | 27.82 % | FSC-H | SSC-H | 3,182,381 | 2,177,327 |
| Single Cells | 1,734 | 48.63 % | FSC-H | FSC-A | 1,883,401 | 5,232,712 |
| Live | 756 | 43.60 % | APC-Cy7-H | FSC-H | 10,702 | 4,798,844 |
| Live sfGFP Negative | 685 | 90.61 % | FITC-A | Count | 110,695 | |
| Live sfGFP Positive | 71 | 9.39 % | FITC-A | Count | 7,344,119 | |
| Dead | 978 | 56.40 % | APC-Cy7-H | FSC-H | 94,646 | 1,449,916 |
| Dead sfGFP Negative | 965 | 98.67 % | FITC-A | Count | 104,564 | |
| Dead sfGFP Positive | 13 | 1.33 % | FITC-A | Count | 9,202,019 | |
| sfGFP Negative | 1,658 | 95.62 % | FITC-A | Count | 107,526 | |
| sfGFP Positive | 76 | 4.38 % | FITC-A | Count | 9,367,979 | |

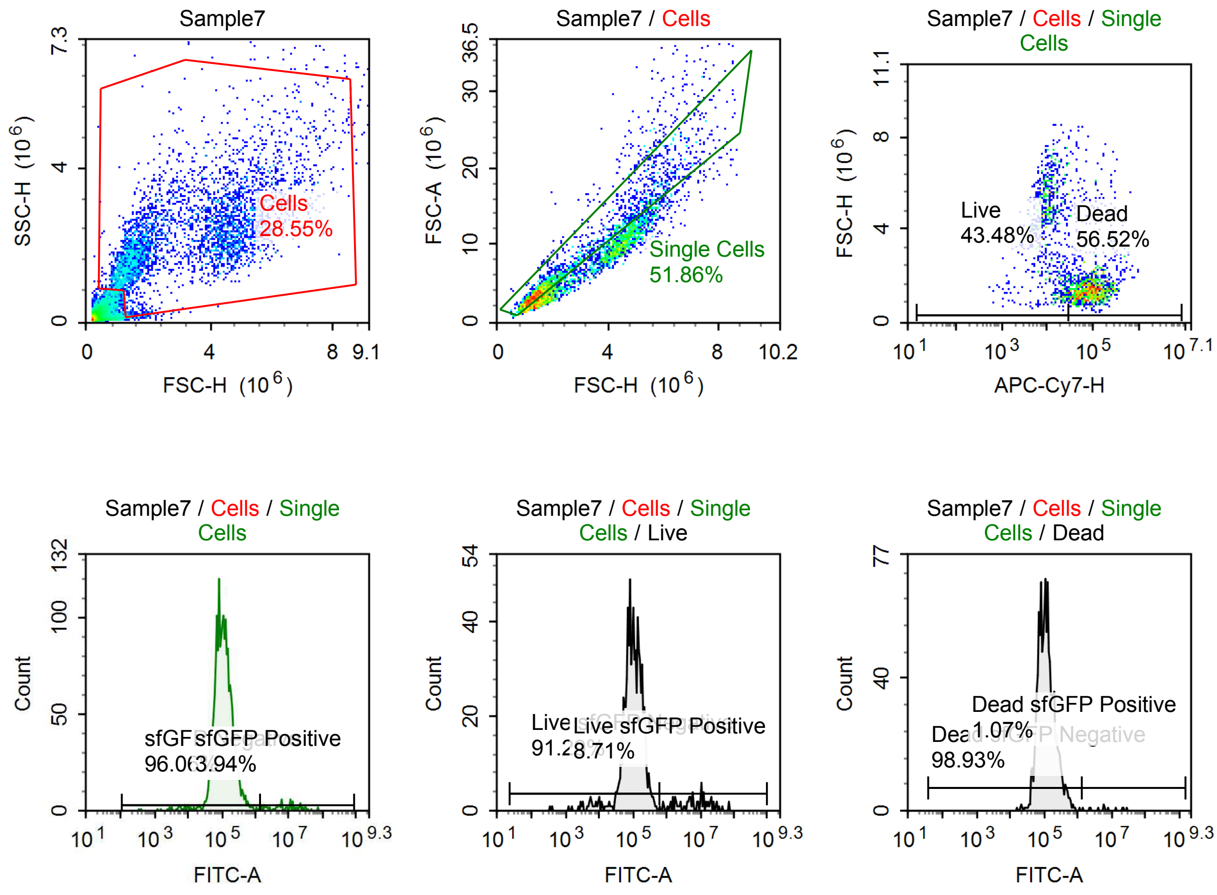

Sample Statistics of Sample7

| Gate | Count | % Parent | X | Y | Median X | Median Y |
|---|---|---|---|---|---|---|
| All | 13,374 | | | | | |
| Cells | 3,818 | 28.55 % | FSC-H | SSC-H | 3,059,433 | 2,141,464 |
| Single Cells | 1,980 | 51.86 % | FSC-H | FSC-A | 1,808,225 | 5,002,837 |
| Live | 861 | 43.48 % | APC-Cy7-H | FSC-H | 10,547 | 4,312,359 |
| Live sfGFP Negative | 786 | 91.29 % | FITC-A | Count | 100,800 | |
| Live sfGFP Positive | 75 | 8.71 % | FITC-A | Count | 6,289,923 | |
| Dead | 1,119 | 56.52 % | APC-Cy7-H | FSC-H | 89,504 | 1,451,620 |
| Dead sfGFP Negative | 1,107 | 98.93 % | FITC-A | Count | 111,622 | |
| Dead sfGFP Positive | 12 | 1.07 % | FITC-A | Count | 6,055,924 | |
| sfGFP Negative | 1,902 | 96.06 % | FITC-A | Count | 108,080 | |
| sfGFP Positive | 78 | 3.94 % | FITC-A | Count | 8,238,823 | |

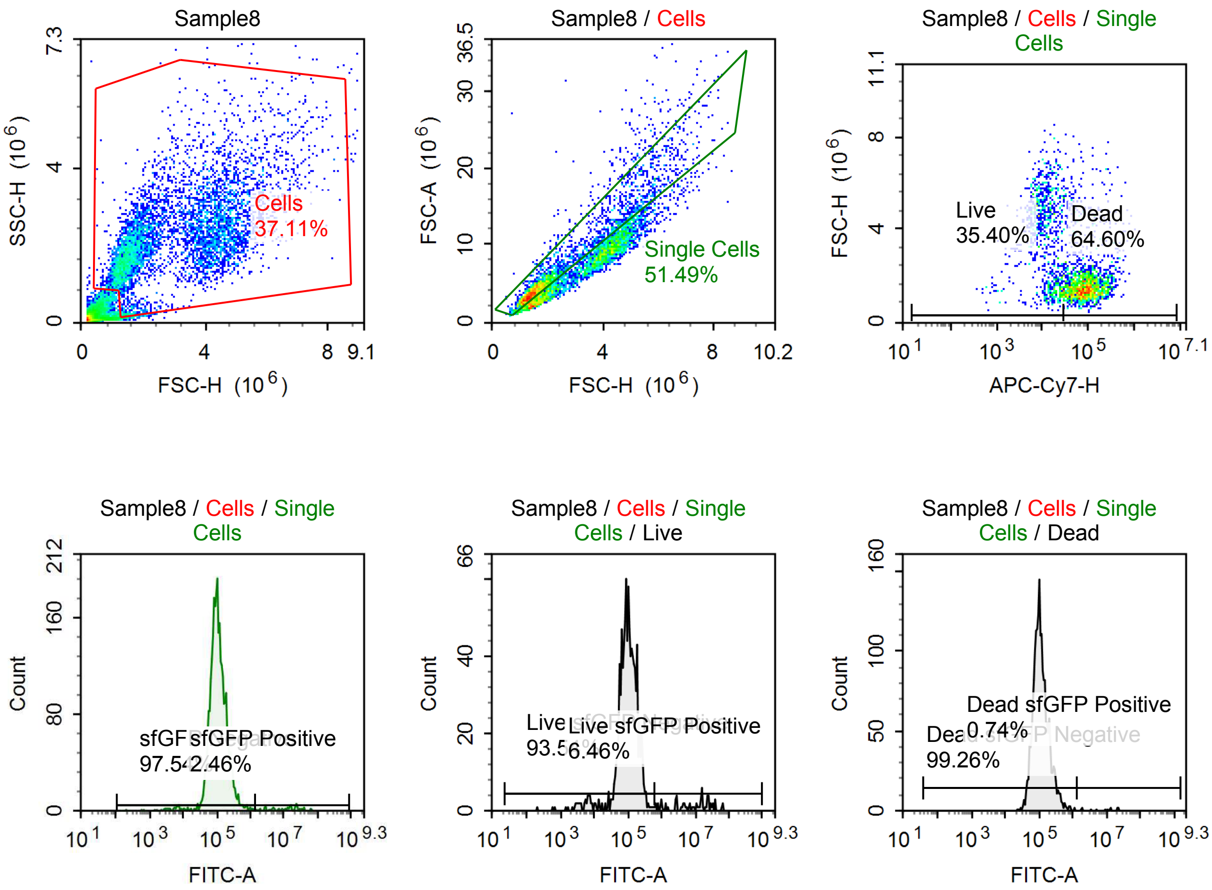

Sample Statistics of Sample8

| Gate | Count | % Parent | X | Y | Median X | Median Y |
|---|---|---|---|---|---|---|
| All | 15,332 | | | | | |
| Cells | 5,689 | 37.11 % | FSC-H | SSC-H | 2,490,435 | 2,087,856 |
| Single Cells | 2,929 | 51.49 % | FSC-H | FSC-A | 1,682,243 | 4,675,649 |
| Live | 1,037 | 35.40 % | APC-Cy7-H | FSC-H | 12,438 | 3,795,746 |
| Live sfGFP Negative | 970 | 93.54 % | FITC-A | Count | 101,253 | |
| Live sfGFP Positive | 67 | 6.46 % | FITC-A | Count | 11,685,778 | |
| Dead | 1,892 | 64.60 % | APC-Cy7-H | FSC-H | 86,123 | 1,493,844 |
| Dead sfGFP Negative | 1,878 | 99.26 % | FITC-A | Count | 105,431 | |
| Dead sfGFP Positive | 14 | 0.74 % | FITC-A | Count | 11,448,590 | |
| sfGFP Negative | 2,857 | 97.54 % | FITC-A | Count | 104,208 | |
| sfGFP Positive | 72 | 2.46 % | FITC-A | Count | 13,534,360 | |

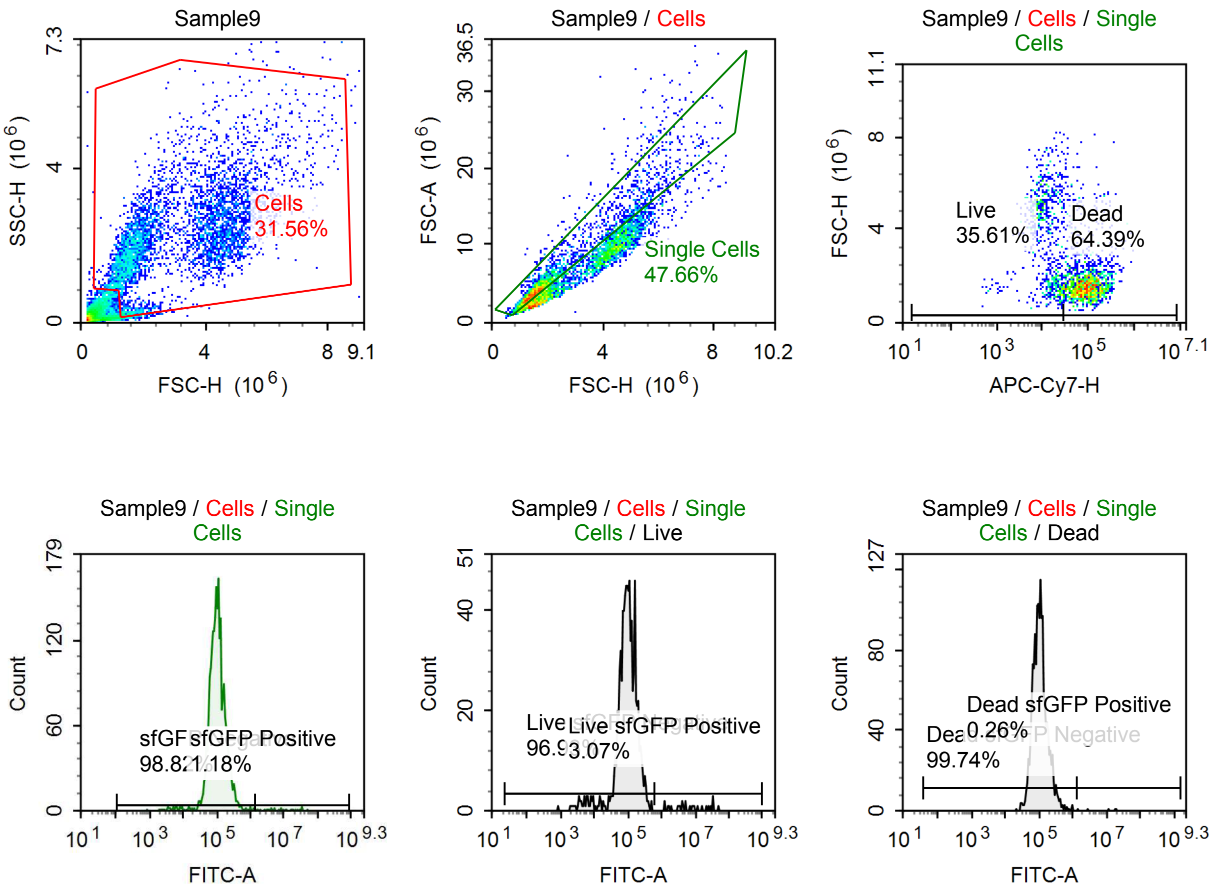

Sample Statistics of Sample9

| Gate | Count | % Parent | X | Y | Median X | Median Y |
|---|---|---|---|---|---|---|
| All | 15,797 | | | | | |
| Cells | 4,985 | 31.56 % | FSC-H | SSC-H | 2,387,649 | 2,112,833 |
| Single Cells | 2,376 | 47.66 % | FSC-H | FSC-A | 1,738,630 | 4,712,505 |
| Live | 846 | 35.61 % | APC-Cy7-H | FSC-H | 12,562 | 3,990,674 |
| Live sfGFP Negative | 820 | 96.93 % | FITC-A | Count | 97,578 | |
| Live sfGFP Positive | 26 | 3.07 % | FITC-A | Count | 6,201,850 | |
| Dead | 1,530 | 64.39 % | APC-Cy7-H | FSC-H | 90,759 | 1,568,011 |
| Dead sfGFP Negative | 1,526 | 99.74 % | FITC-A | Count | 104,252 | |
| Dead sfGFP Positive | 4 | 0.26 % | FITC-A | Count | 14,329,597 | |
| sfGFP Negative | 2,348 | 98.82 % | FITC-A | Count | 101,728 | |
| sfGFP Positive | 28 | 1.18 % | FITC-A | Count | 8,356,382 | |

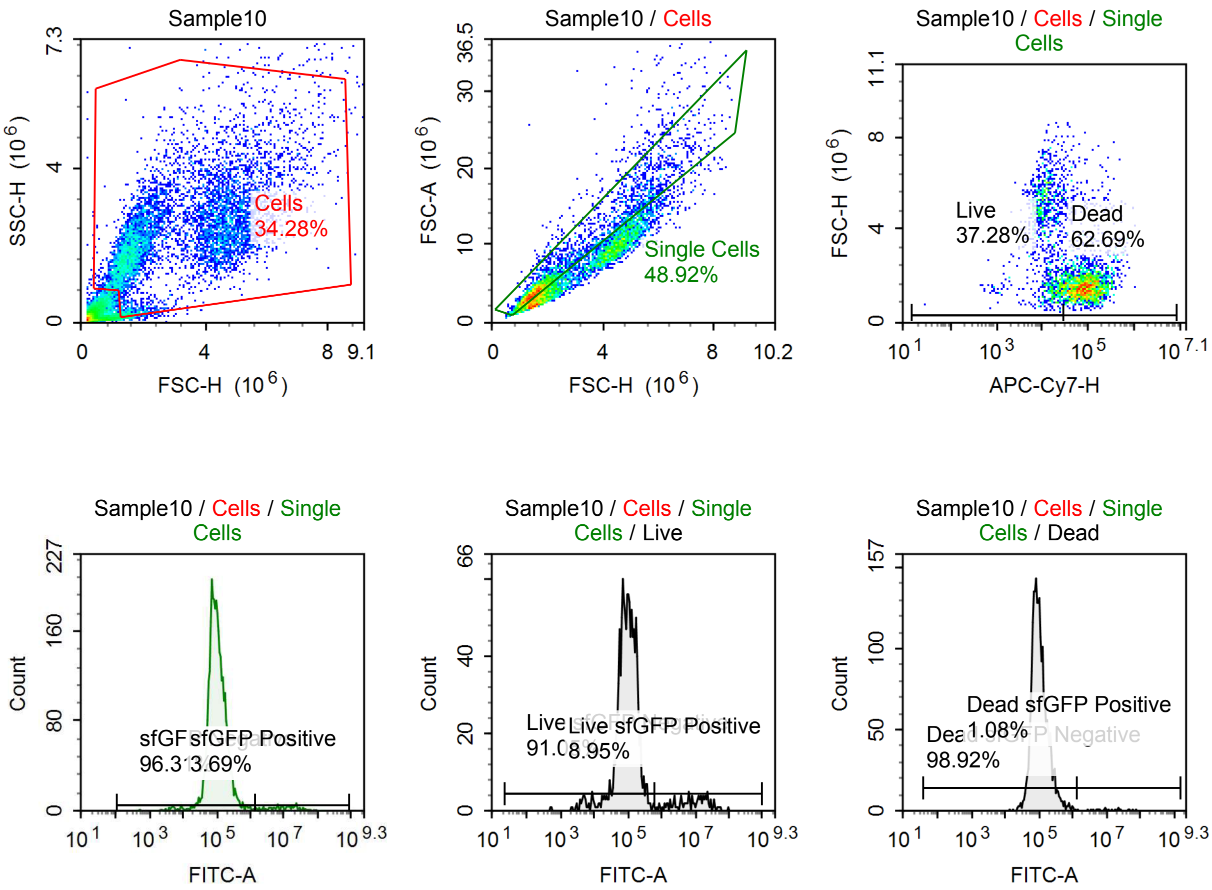

Sample Statistics of Sample10

| Gate | Count | % Parent | X | Y | Median X | Median Y |
|---|---|---|---|---|---|---|
| All | 20,184 | | | | | |
| Cells | 6,919 | 34.28 % | FSC-H | SSC-H | 2,405,291 | 2,168,471 |
| Single Cells | 3,385 | 48.92 % | FSC-H | FSC-A | 1,738,053 | 4,680,221 |
| Live | 1,262 | 37.28 % | APC-Cy7-H | FSC-H | 12,857 | 3,843,684 |
| Live sfGFP Negative | 1,149 | 91.05 % | FITC-A | Count | 96,222 | |
| Live sfGFP Positive | 113 | 8.95 % | FITC-A | Count | 8,875,283 | |
| Dead | 2,122 | 62.69 % | APC-Cy7-H | FSC-H | 87,163 | 1,572,397 |
| Dead sfGFP Negative | 2,099 | 98.92 % | FITC-A | Count | 96,228 | |
| Dead sfGFP Positive | 23 | 1.08 % | FITC-A | Count | 11,770,720 | |
| sfGFP Negative | 3,260 | 96.31 % | FITC-A | Count | 96,352 | |
| sfGFP Positive | 125 | 3.69 % | FITC-A | Count | 11,575,137 | |

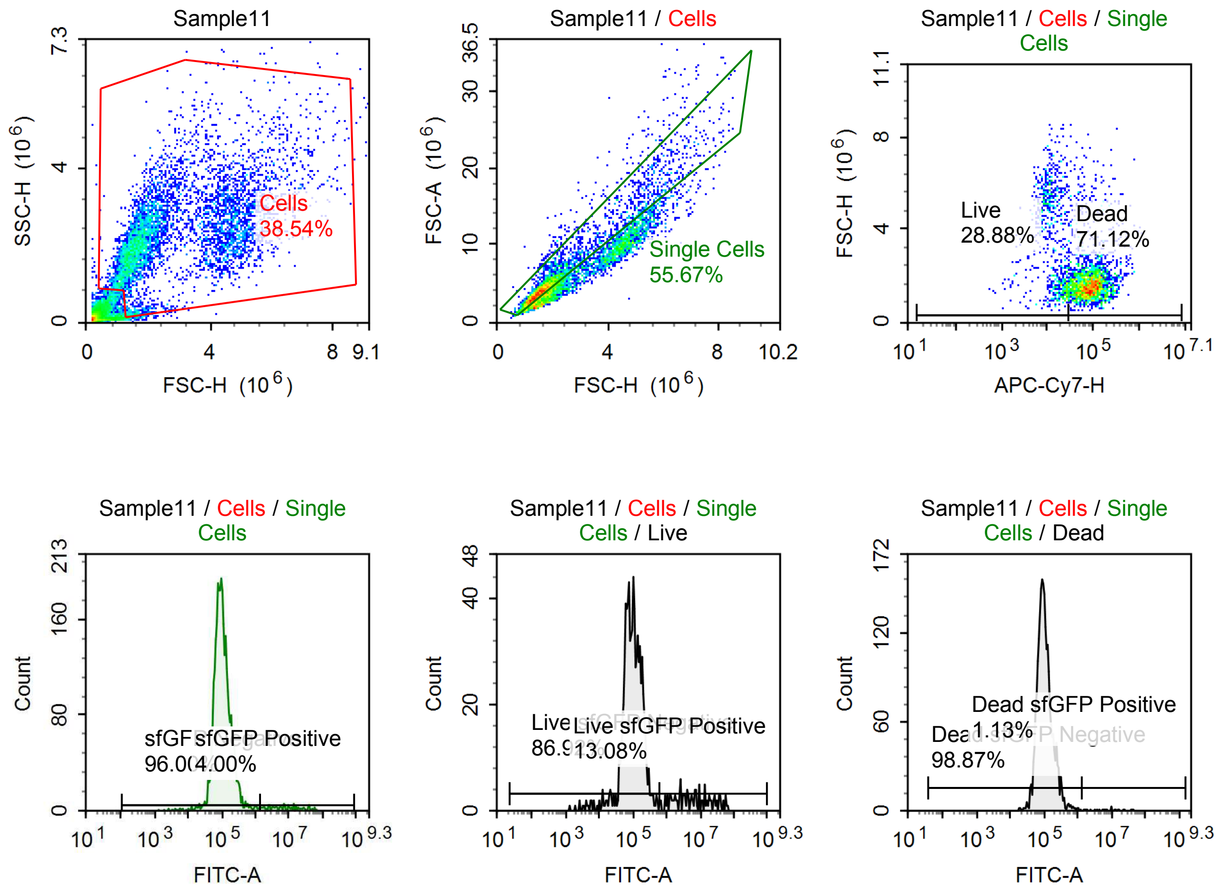

Sample Statistics of Sample11

| Gate | Count | % Parent | X | Y | Median X | Median Y |
|---|---|---|---|---|---|---|
| All | 14,557 | | | | | |
| Cells | 5,610 | 38.54 % | FSC-H | SSC-H | 2,113,428 | 2,195,097 |
| Single Cells | 3,123 | 55.67 % | FSC-H | FSC-A | 1,696,523 | 4,529,917 |
| Live | 902 | 28.88 % | APC-Cy7-H | FSC-H | 14,066 | 3,188,194 |
| Live sfGFP Negative | 784 | 86.92 % | FITC-A | Count | 97,174 | |
| Live sfGFP Positive | 118 | 13.08 % | FITC-A | Count | 5,794,189 | |
| Dead | 2,221 | 71.12 % | APC-Cy7-H | FSC-H | 85,149 | 1,593,833 |
| Dead sfGFP Negative | 2,196 | 98.87 % | FITC-A | Count | 98,314 | |
| Dead sfGFP Positive | 25 | 1.13 % | FITC-A | Count | 7,150,848 | |
| sfGFP Negative | 2,998 | 96.00 % | FITC-A | Count | 98,442 | |
| sfGFP Positive | 125 | 4.00 % | FITC-A | Count | 7,382,575 | |

#### p060 mRNA [100 ng]

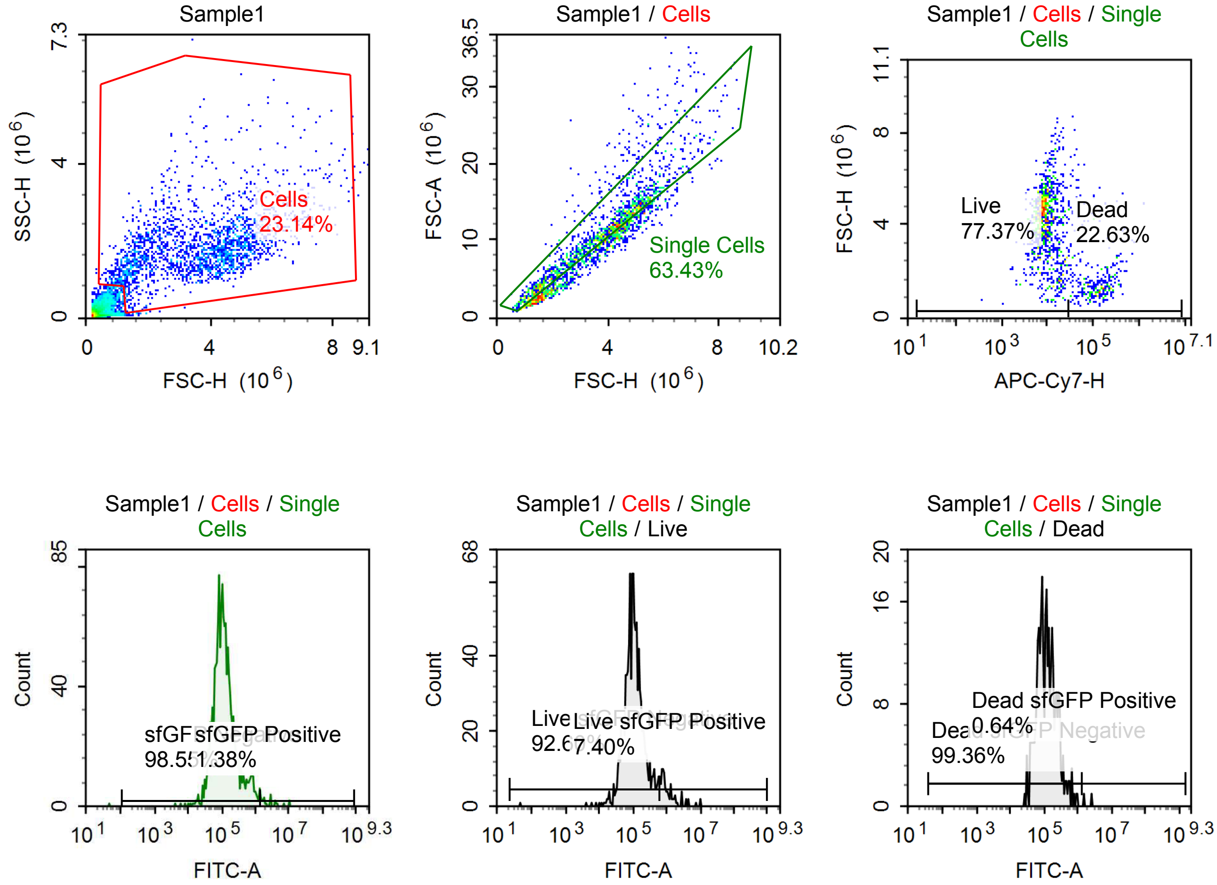

Sample Statistics of Sample1

| Gate | Count | % Parent | X | Y | Median X | Median Y |
|---|---|---|---|---|---|---|
| All | 9,396 | | | | | |
| Cells | 2,174 | 23.14 % | FSC-H | SSC-H | 3,748,654 | 1,755,889 |
| Single Cells | 1,379 | 63.43 % | FSC-H | FSC-A | 3,784,102 | 10,790,472 |
| Live | 1,067 | 77.37 % | APC-Cy7-H | FSC-H | 8,609 | 4,239,601 |
| Live sfGFP Negative | 988 | 92.60 % | FITC-A | Count | 102,929 | |
| Live sfGFP Positive | 79 | 7.40 % | FITC-A | Count | 996,849 | |
| Dead | 312 | 22.63 % | APC-Cy7-H | FSC-H | 118,817 | 1,452,952 |
| Dead sfGFP Negative | 310 | 99.36 % | FITC-A | Count | 115,312 | |
| Dead sfGFP Positive | 2 | 0.64 % | FITC-A | Count | 2,051,888 | |
| sfGFP Negative | 1,359 | 98.55 % | FITC-A | Count | 108,772 | |
| sfGFP Positive | 19 | 1.38 % | FITC-A | Count | 2,796,700 | |

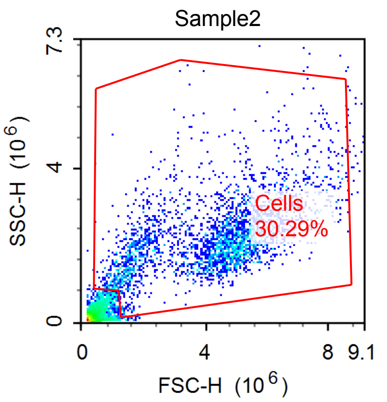


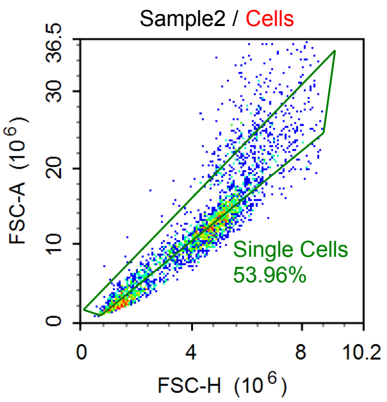


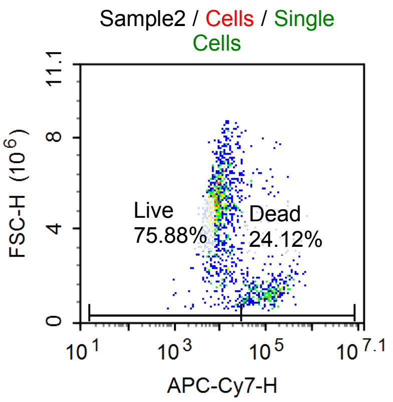


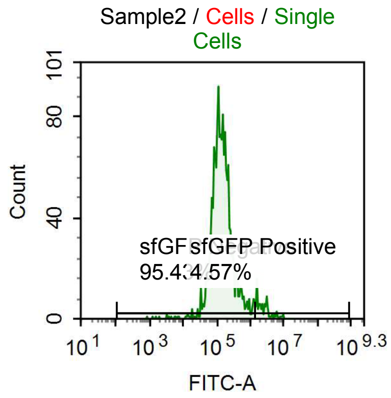


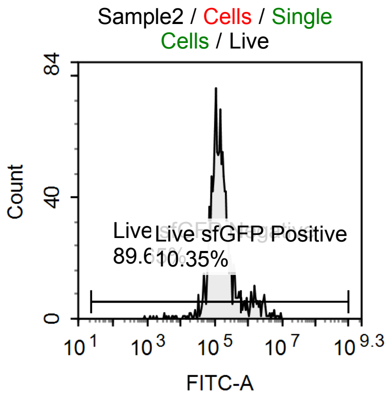


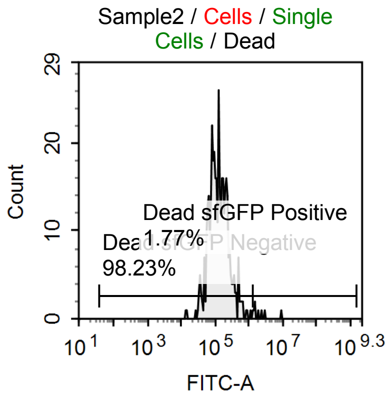

Sample Statistics of Sample2

| Gate | Count | % Parent | X | Y | Median X | Median Y |
|---|---|---|---|---|---|---|
| All | 10,046 | | | | | |
| Cells | 3,043 | 30.29 % | FSC-H | SSC-H | 4,564,913 | 2,067,470 |
| Single Cells | 1,642 | 53.96 % | FSC-H | FSC-A | 4,461,825 | 12,488,315 |
| Live | 1,246 | 75.88 % | APC-Cy7-H | FSC-H | 8,573 | 4,864,119 |
| Live sfGFP Negative | 1,117 | 89.65 % | FITC-A | Count | 134,354 | |
| Live sfGFP Positive | 129 | 10.35 % | FITC-A | Count | 1,635,180 | |
| Dead | 396 | 24.12 % | APC-Cy7-H | FSC-H | 109,877 | 1,478,412 |
| Dead sfGFP Negative | 389 | 98.23 % | FITC-A | Count | 124,775 | |
| Dead sfGFP Positive | 7 | 1.77 % | FITC-A | Count | 1,843,773 | |
| sfGFP Negative | 1,567 | 95.43 % | FITC-A | Count | 135,904 | |
| sfGFP Positive | 75 | 4.57 % | FITC-A | Count | 2,687,419 | |

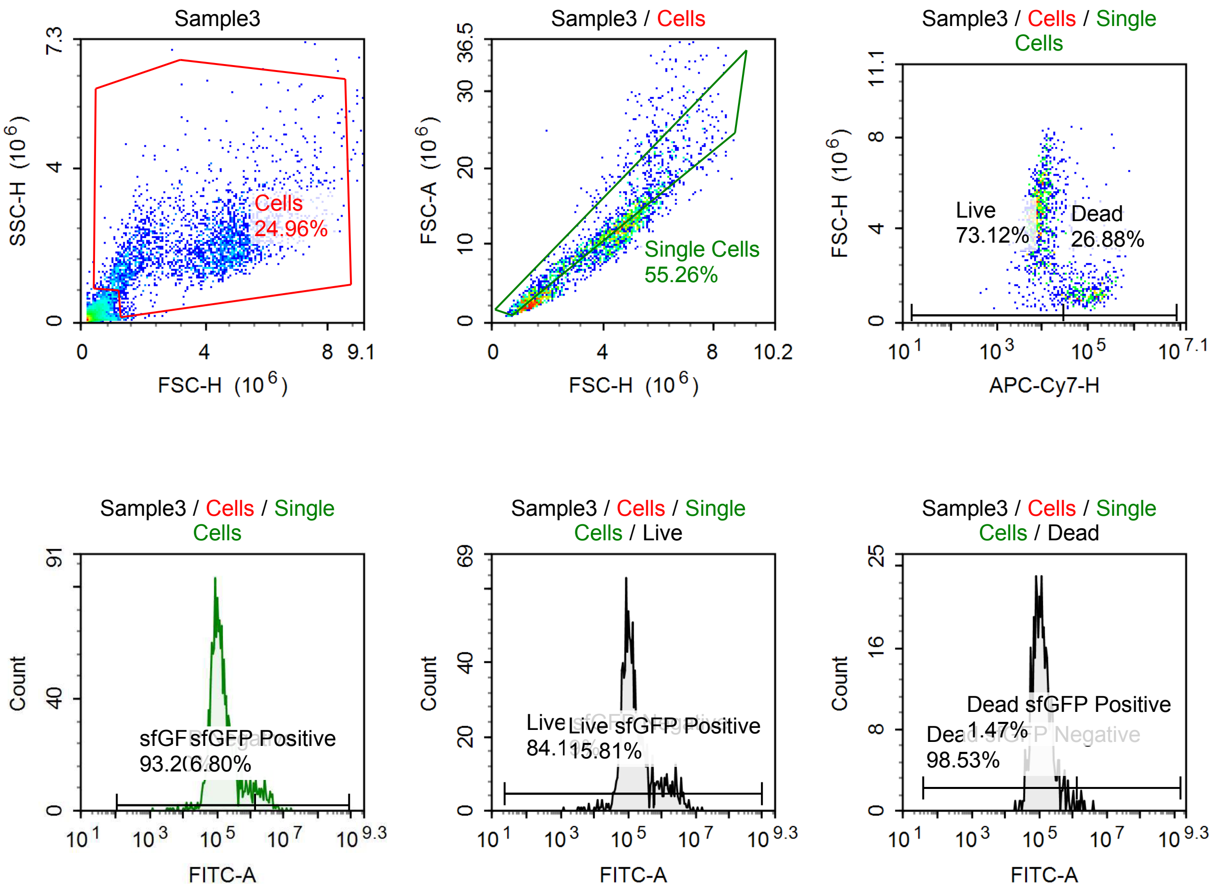

Sample Statistics of Sample3

| Gate | Count | % Parent | X | Y | Median X | Median Y |
|---|---|---|---|---|---|---|
| All | 10,979 | | | | | |
| Cells | 2,740 | 24.96 % | FSC-H | SSC-H | 4,141,788 | 1,941,813 |
| Single Cells | 1,514 | 55.26 % | FSC-H | FSC-A | 3,885,539 | 11,111,754 |
| Live | 1,107 | 73.12 % | APC-Cy7-H | FSC-H | 8,393 | 4,565,014 |
| Live sfGFP Negative | 932 | 84.19 % | FITC-A | Count | 116,242 | |
| Live sfGFP Positive | 175 | 15.81 % | FITC-A | Count | 1,757,015 | |
| Dead | 407 | 26.88 % | APC-Cy7-H | FSC-H | 97,372 | 1,364,947 |
| Dead sfGFP Negative | 401 | 98.53 % | FITC-A | Count | 113,444 | |
| Dead sfGFP Positive | 6 | 1.47 % | FITC-A | Count | 1,863,985 | |
| sfGFP Negative | 1,411 | 93.20 % | FITC-A | Count | 119,215 | |
| sfGFP Positive | 103 | 6.80 % | FITC-A | Count | 2,693,329 | |

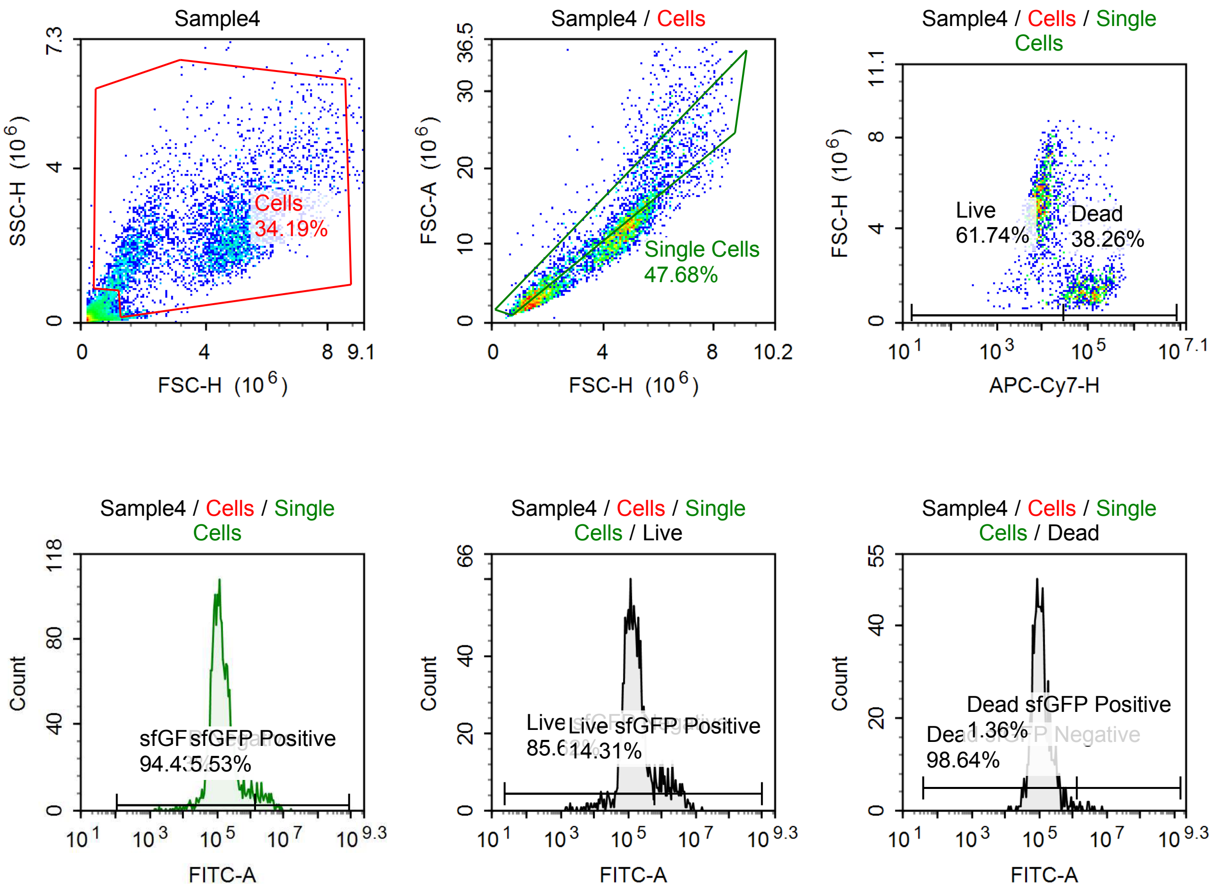

Sample Statistics of Sample4

| Gate | Count | % Parent | X | Y | Median X | Median Y |
|---|---|---|---|---|---|---|
| All | 12,988 | | | | | |
| Cells | 4,440 | 34.19 % | FSC-H | SSC-H | 4,367,120 | 2,181,143 |
| Single Cells | 2,117 | 47.68 % | FSC-H | FSC-A | 4,120,282 | 11,532,250 |
| Live | 1,307 | 61.74 % | APC-Cy7-H | FSC-H | 9,120 | 4,960,278 |
| Live sfGFP Negative | 1,119 | 85.62 % | FITC-A | Count | 134,601 | |
| Live sfGFP Positive | 187 | 14.31 % | FITC-A | Count | 1,803,909 | |
| Dead | 810 | 38.26 % | APC-Cy7-H | FSC-H | 106,685 | 1,542,876 |
| Dead sfGFP Negative | 799 | 98.64 % | FITC-A | Count | 105,681 | |
| Dead sfGFP Positive | 11 | 1.36 % | FITC-A | Count | 2,789,507 | |
| sfGFP Negative | 1,999 | 94.43 % | FITC-A | Count | 124,415 | |
| sfGFP Positive | 117 | 5.53 % | FITC-A | Count | 2,789,507 | |

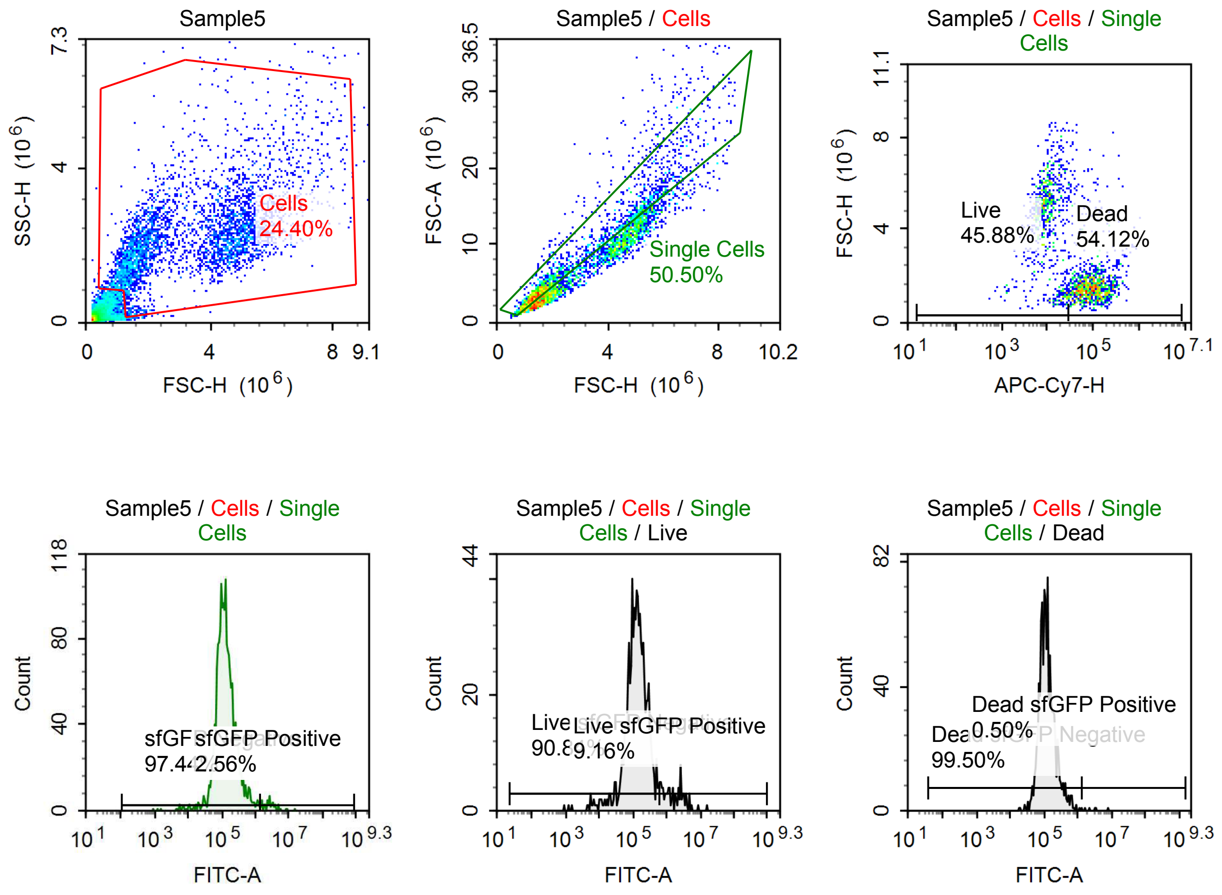

Sample Statistics of Sample5

| Gate | Count | % Parent | X | Y | Median X | Median Y |
|---|---|---|---|---|---|---|
| All | 14,877 | | | | | |
| Cells | 3,630 | 24.40 % | FSC-H | SSC-H | 3,104,793 | 2,105,700 |
| Single Cells | 1,833 | 50.50 % | FSC-H | FSC-A | 1,967,857 | 5,588,767 |
| Live | 841 | 45.88 % | APC-Cy7-H | FSC-H | 9,975 | 4,752,450 |
| Live sfGFP Negative | 764 | 90.84 % | FITC-A | Count | 130,974 | |
| Live sfGFP Positive | 77 | 9.16 % | FITC-A | Count | 1,763,190 | |
| Dead | 992 | 54.12 % | APC-Cy7-H | FSC-H | 92,131 | 1,507,504 |
| Dead sfGFP Negative | 987 | 99.50 % | FITC-A | Count | 115,169 | |
| Dead sfGFP Positive | 5 | 0.50 % | FITC-A | Count | 3,301,723 | |
| sfGFP Negative | 1,786 | 97.44 % | FITC-A | Count | 122,943 | |
| sfGFP Positive | 47 | 2.56 % | FITC-A | Count | 2,714,111 | |

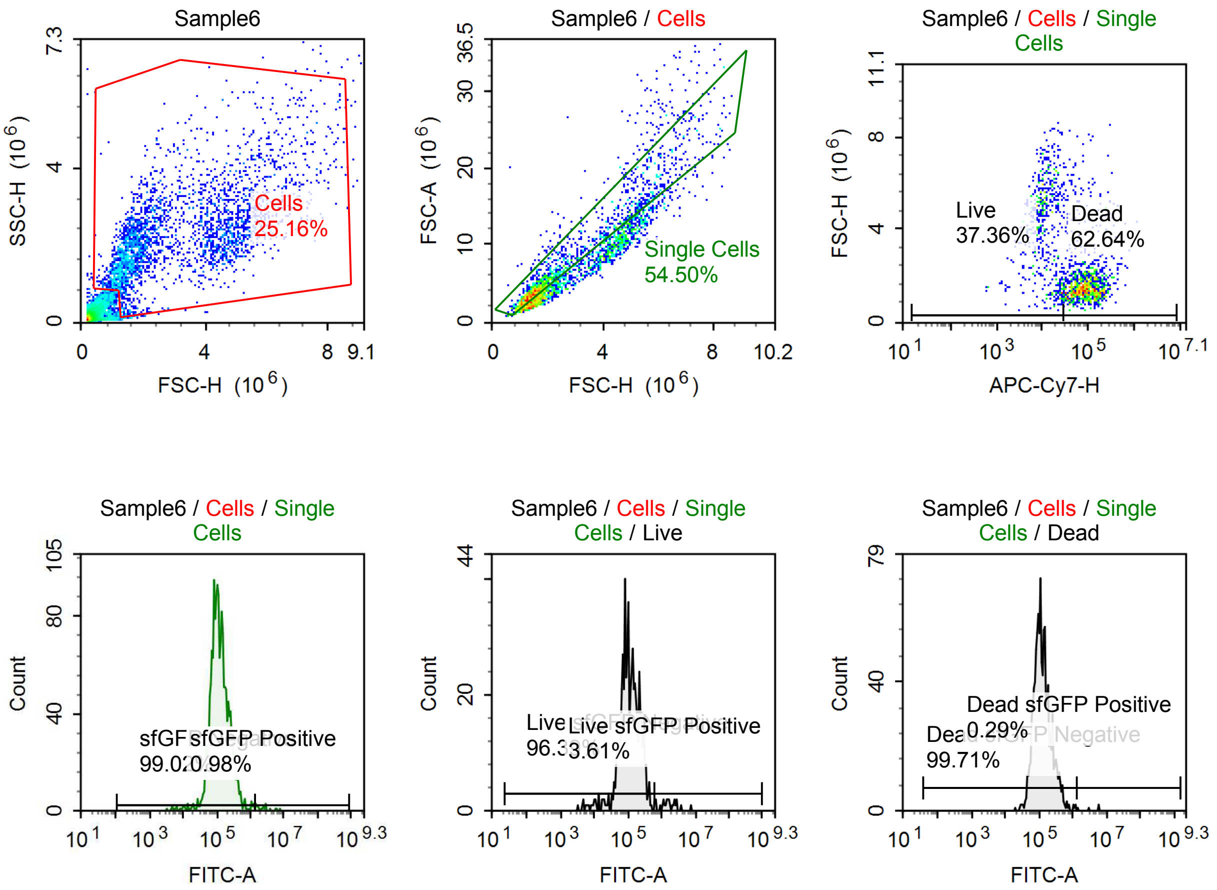

Sample Statistics of Sample6

| Gate | Count | % Parent | X | Y | Median X | Median Y |
|---|---|---|---|---|---|---|
| All | 11,890 | | | | | |
| Cells | 2,991 | 25.16 % | FSC-H | SSC-H | 2,293,000 | 2,099,009 |
| Single Cells | 1,630 | 54.50 % | FSC-H | FSC-A | 1,750,282 | 4,840,834 |
| Live | 609 | 37.36 % | APC-Cy7-H | FSC-H | 11,492 | 4,226,453 |
| Live sfGFP Negative | 587 | 96.39 % | FITC-A | Count | 108,804 | |
| Live sfGFP Positive | 22 | 3.61 % | FITC-A | Count | 1,669,167 | |
| Dead | 1,021 | 62.64 % | APC-Cy7-H | FSC-H | 87,560 | 1,484,681 |
| Dead sfGFP Negative | 1,018 | 99.71 % | FITC-A | Count | 115,801 | |
| Dead sfGFP Positive | 3 | 0.29 % | FITC-A | Count | 5,739,480 | |
| sfGFP Negative | 1,614 | 99.02 % | FITC-A | Count | 114,730 | |
| sfGFP Positive | 16 | 0.98 % | FITC-A | Count | 2,572,509 | |

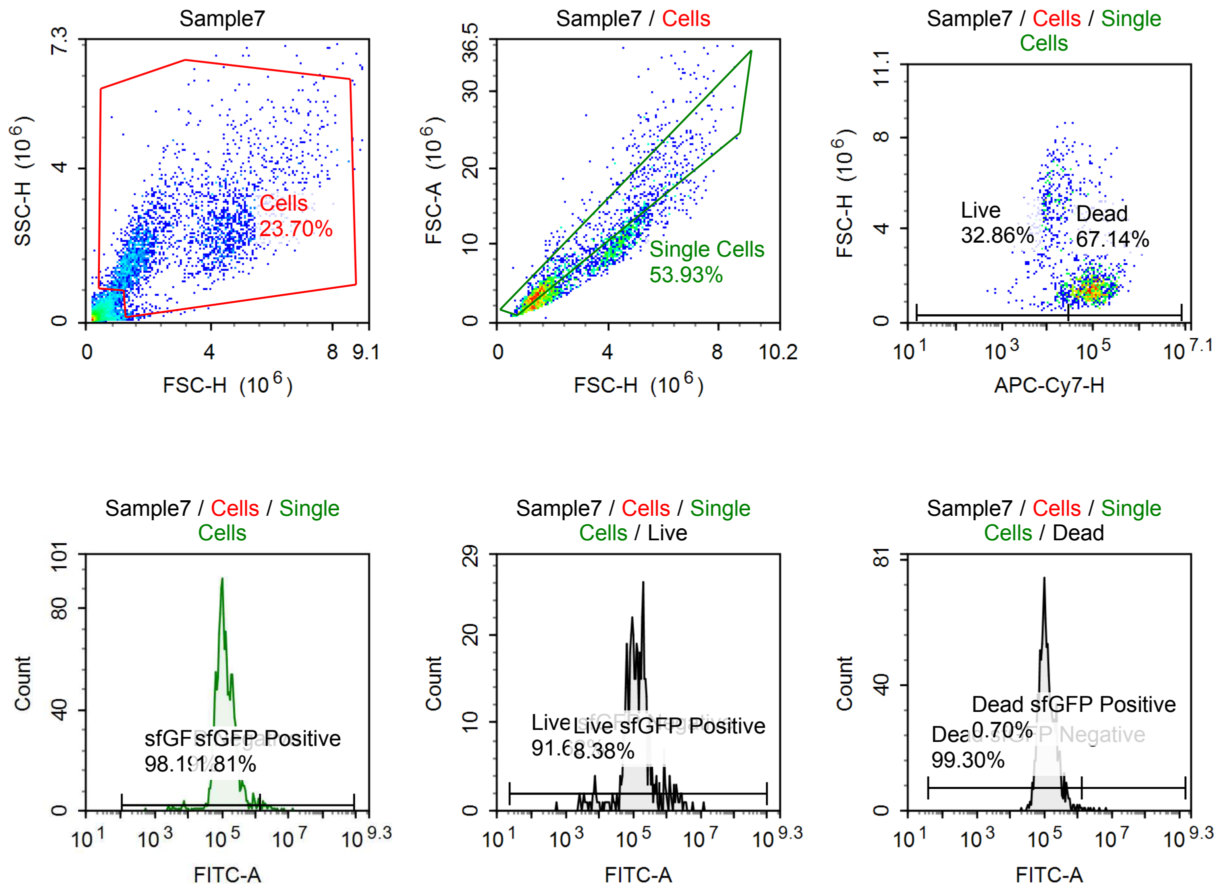

Sample Statistics of Sample7

| Gate | Count | % Parent | X | Y | Median X | Median Y |
|---|---|---|---|---|---|---|
| All | 11,641 | | | | | |
| Cells | 2,759 | 23.70 % | FSC-H | SSC-H | 2,220,837 | 2,037,956 |
| Single Cells | 1,488 | 53.93 % | FSC-H | FSC-A | 1,659,765 | 4,594,359 |
| Live | 489 | 32.86 % | APC-Cy7-H | FSC-H | 11,894 | 4,273,126 |
| Live sfGFP Negative | 448 | 91.62 % | FITC-A | Count | 118,490 | |
| Live sfGFP Positive | 41 | 8.38 % | FITC-A | Count | 1,485,317 | |
| Dead | 999 | 67.14 % | APC-Cy7-H | FSC-H | 92,472 | 1,486,120 |
| Dead sfGFP Negative | 992 | 99.30 % | FITC-A | Count | 113,424 | |
| Dead sfGFP Positive | 7 | 0.70 % | FITC-A | Count | 2,425,438 | |
| sfGFP Negative | 1,461 | 98.19 % | FITC-A | Count | 116,016 | |
| sfGFP Positive | 27 | 1.81 % | FITC-A | Count | 2,379,155 | |

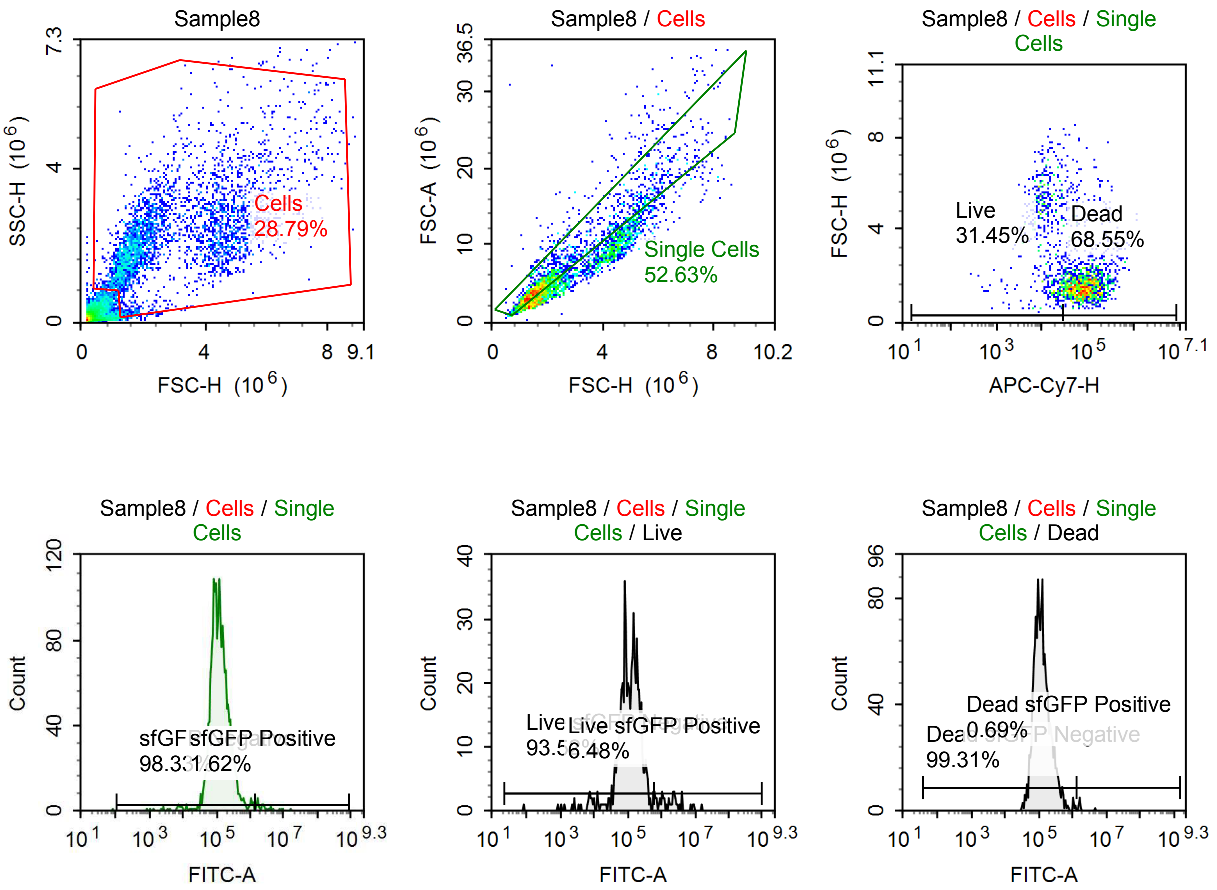

Sample Statistics of Sample8

| Gate | Count | % Parent | X | Y | Median X | Median Y |
|---|---|---|---|---|---|---|
| All | 12,634 | | | | | |
| Cells | 3,637 | 28.79 % | FSC-H | SSC-H | 2,237,182 | 2,106,317 |
| Single Cells | 1,914 | 52.63 % | FSC-H | FSC-A | 1,757,901 | 4,851,957 |
| Live | 602 | 31.45 % | APC-Cy7-H | FSC-H | 13,126 | 3,945,882 |
| Live sfGFP Negative | 563 | 93.52 % | FITC-A | Count | 115,899 | |
| Live sfGFP Positive | 39 | 6.48 % | FITC-A | Count | 1,980,066 | |
| Dead | 1,312 | 68.55 % | APC-Cy7-H | FSC-H | 85,850 | 1,607,389 |
| Dead sfGFP Negative | 1,303 | 99.31 % | FITC-A | Count | 118,183 | |
| Dead sfGFP Positive | 9 | 0.69 % | FITC-A | Count | 1,648,854 | |
| sfGFP Negative | 1,882 | 98.33 % | FITC-A | Count | 118,335 | |
| sfGFP Positive | 31 | 1.62 % | FITC-A | Count | 2,431,258 | |

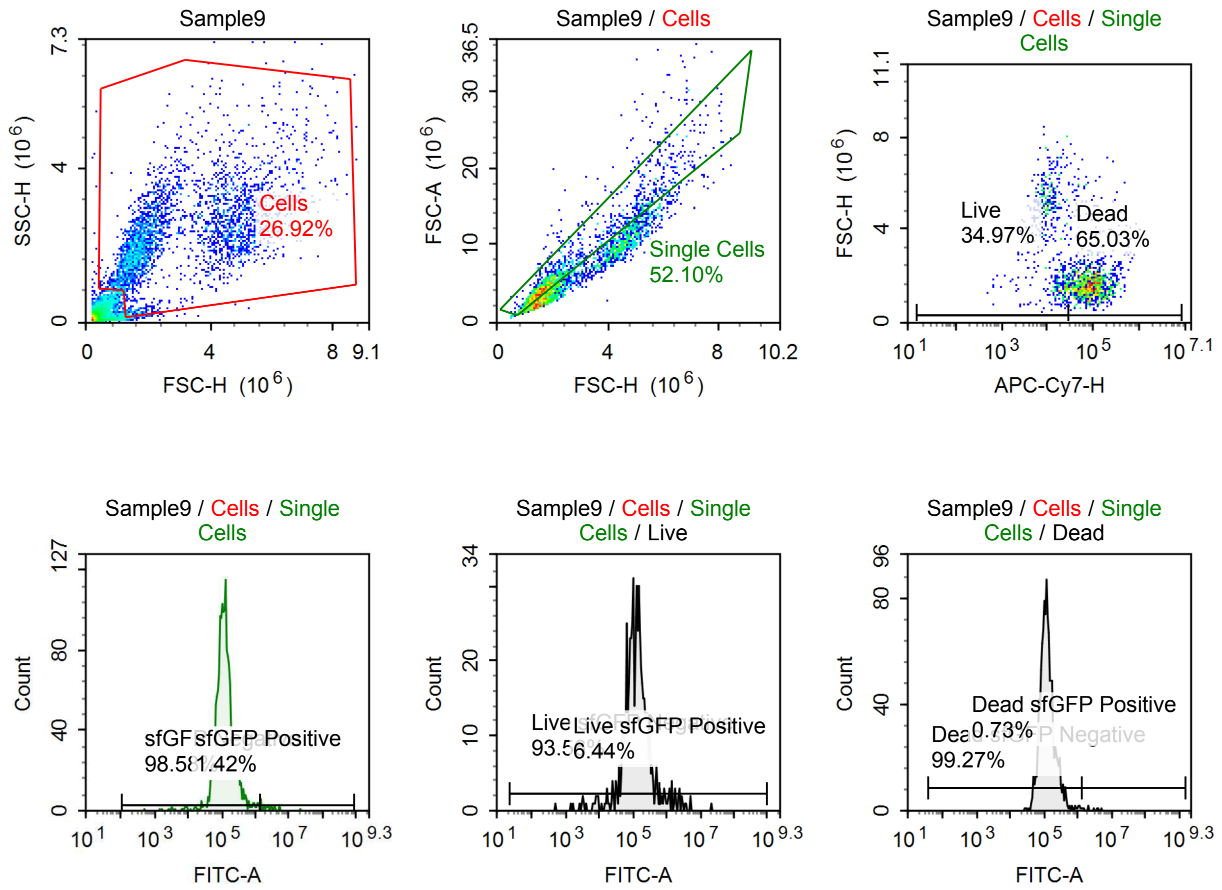

Sample Statistics of Sample9

| Gate | Count | % Parent | X | Y | Median X | Median Y |
|---|---|---|---|---|---|---|
| All | 12,028 | | | | | |
| Cells | 3,238 | 26.92 % | FSC-H | SSC-H | 2,150,868 | 2,141,569 |
| Single Cells | 1,687 | 52.10 % | FSC-H | FSC-A | 1,780,254 | 4,898,377 |
| Live | 590 | 34.97 % | APC-Cy7-H | FSC-H | 12,897 | 3,809,560 |
| Live sfGFP Negative | 552 | 93.56 % | FITC-A | Count | 118,614 | |
| Live sfGFP Positive | 38 | 6.44 % | FITC-A | Count | 1,489,602 | |
| Dead | 1,097 | 65.03 % | APC-Cy7-H | FSC-H | 88,438 | 1,640,070 |
| Dead sfGFP Negative | 1,089 | 99.27 % | FITC-A | Count | 118,986 | |
| Dead sfGFP Positive | 8 | 0.73 % | FITC-A | Count | 2,603,595 | |
| sfGFP Negative | 1,663 | 98.58 % | FITC-A | Count | 119,616 | |
| sfGFP Positive | 24 | 1.42 % | FITC-A | Count | 2,620,450 | |

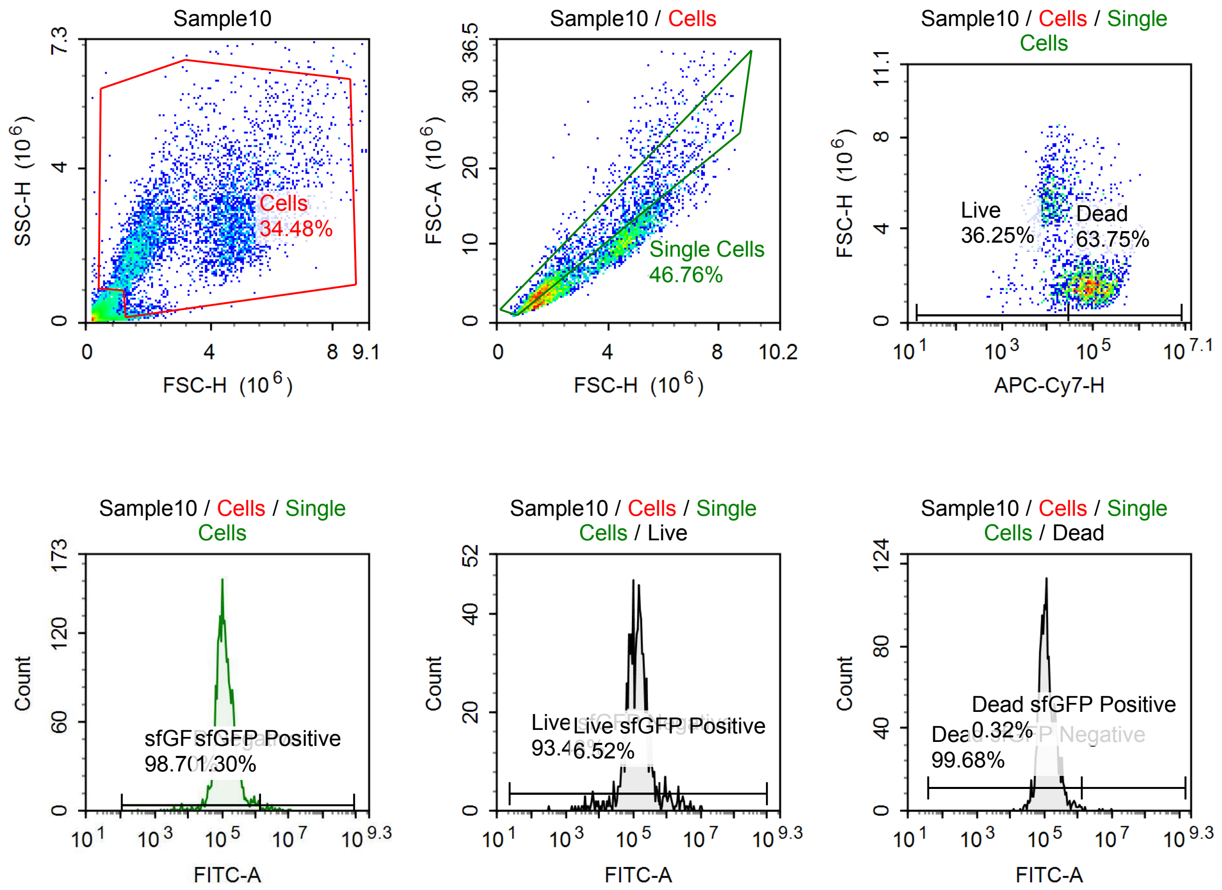

Sample Statistics of Sample10

| Gate | Count | % Parent | X | Y | Median X | Median Y |
|---|---|---|---|---|---|---|
| All | 15,227 | | | | | |
| Cells | 5,250 | 34.48 % | FSC-H | SSC-H | 2,731,715 | 2,278,009 |
| Single Cells | 2,455 | 46.76 % | FSC-H | FSC-A | 1,886,903 | 5,198,752 |
| Live | 890 | 36.25 % | APC-Cy7-H | FSC-H | 13,099 | 4,492,855 |
| Live sfGFP Negative | 832 | 93.48 % | FITC-A | Count | 129,190 | |
| Live sfGFP Positive | 58 | 6.52 % | FITC-A | Count | 1,366,912 | |
| Dead | 1,565 | 63.75 % | APC-Cy7-H | FSC-H | 89,444 | 1,668,288 |
| Dead sfGFP Negative | 1,560 | 99.68 % | FITC-A | Count | 114,806 | |
| Dead sfGFP Positive | 5 | 0.32 % | FITC-A | Count | 4,389,953 | |
| sfGFP Negative | 2,423 | 98.70 % | FITC-A | Count | 118,242 | |
| sfGFP Positive | 32 | 1.30 % | FITC-A | Count | 2,944,414 | |

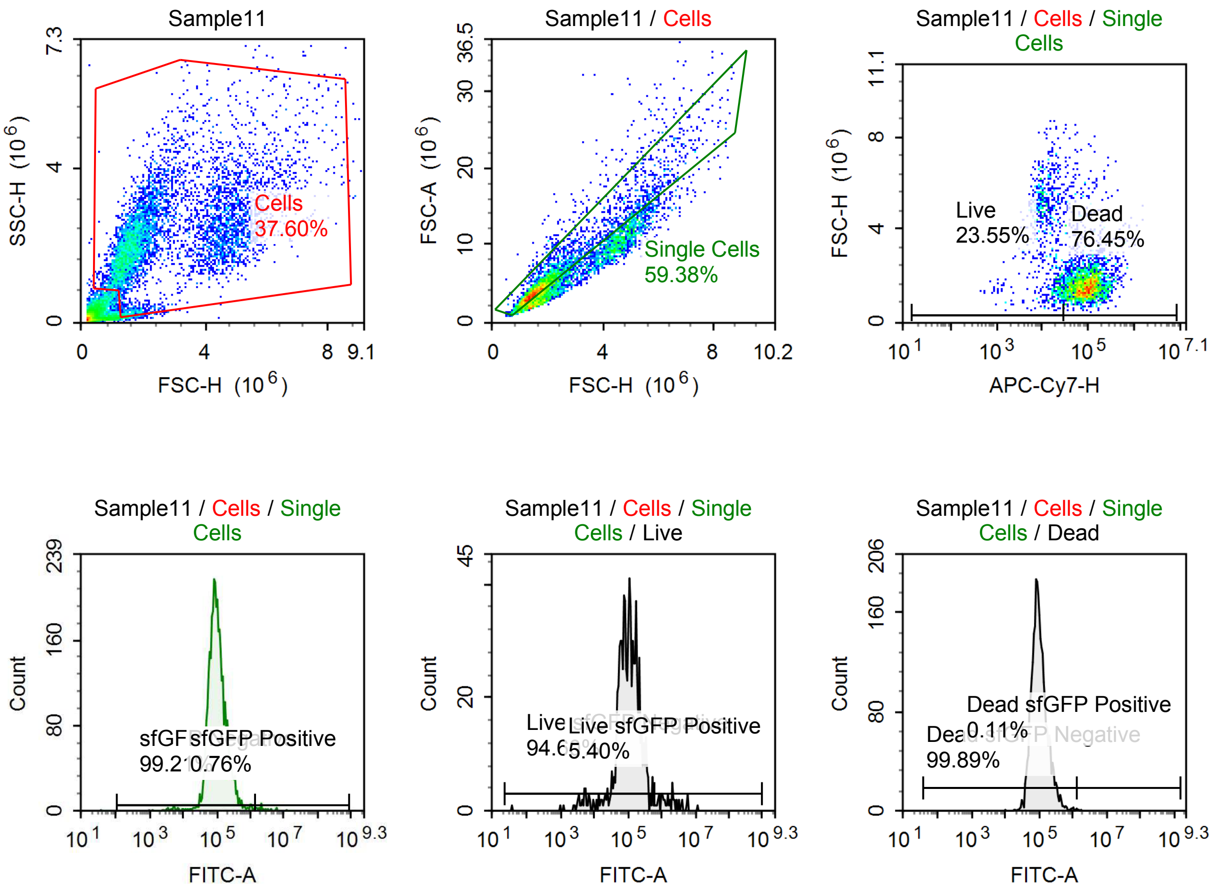

Sample Statistics of Sample11

| Gate | Count | % Parent | X | Y | Median X | Median Y |
|---|---|---|---|---|---|---|
| All | 15,843 | | | | | |
| Cells | 5,957 | 37.60 % | FSC-H | SSC-H | 1,983,838 | 2,134,143 |
| Single Cells | 3,537 | 59.38 % | FSC-H | FSC-A | 1,673,154 | 4,405,004 |
| Live | 833 | 23.55 % | APC-Cy7-H | FSC-H | 13,431 | 3,875,949 |
| Live sfGFP Negative | 788 | 94.60 % | FITC-A | Count | 104,886 | |
| Live sfGFP Positive | 45 | 5.40 % | FITC-A | Count | 1,832,173 | |
| Dead | 2,704 | 76.45 % | APC-Cy7-H | FSC-H | 92,476 | 1,579,461 |
| Dead sfGFP Negative | 2,701 | 99.89 % | FITC-A | Count | 96,036 | |
| Dead sfGFP Positive | 3 | 0.11 % | FITC-A | Count | 1,477,133 | |
| sfGFP Negative | 3,509 | 99.21 % | FITC-A | Count | 97,929 | |
| sfGFP Positive | 27 | 0.76 % | FITC-A | Count | 2,363,944 | |

#### p060 mRNA [100 ng] + TET [1 uM]

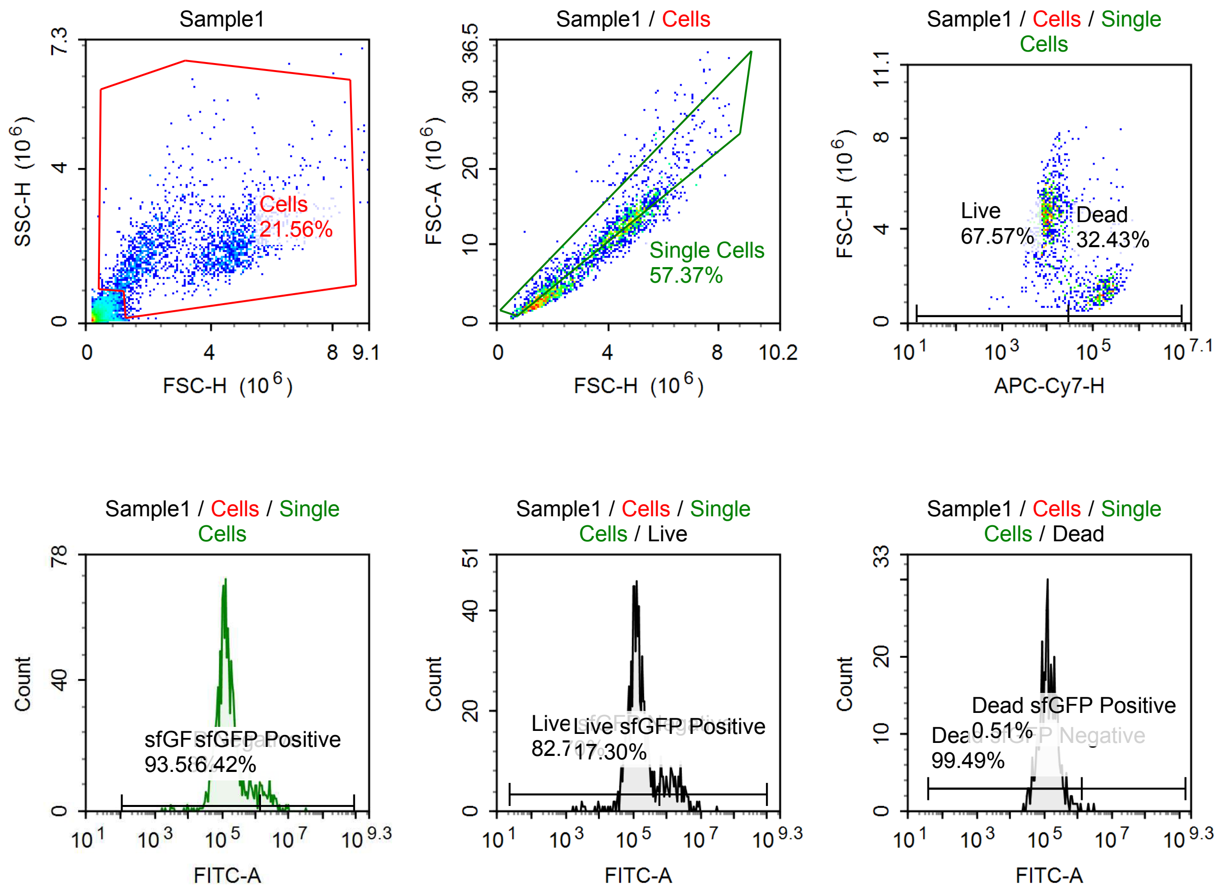

Sample Statistics of Sample1

| Gate | Count | % Parent | X | Y | Median X | Median Y |
|---|---|---|---|---|---|---|
| All | 9,826 | | | | | |
| Cells | 2,118 | 21.56 % | FSC-H | SSC-H | 3,616,204 | 1,903,652 |
| Single Cells | 1,215 | 57.37 % | FSC-H | FSC-A | 3,932,905 | 11,189,145 |
| Live | 821 | 67.57 % | APC-Cy7-H | FSC-H | 10,210 | 4,532,233 |
| Live sfGFP Negative | 679 | 82.70 % | FITC-A | Count | 128,402 | |
| Live sfGFP Positive | 142 | 17.30 % | FITC-A | Count | 1,622,640 | |
| Dead | 394 | 32.43 % | APC-Cy7-H | FSC-H | 135,775 | 1,502,402 |
| Dead sfGFP Negative | 392 | 99.49 % | FITC-A | Count | 129,395 | |
| Dead sfGFP Positive | 2 | 0.51 % | FITC-A | Count | 2,510,489 | |
| sfGFP Negative | 1,137 | 93.58 % | FITC-A | Count | 131,962 | |
| sfGFP Positive | 78 | 6.42 % | FITC-A | Count | 2,616,092 | |

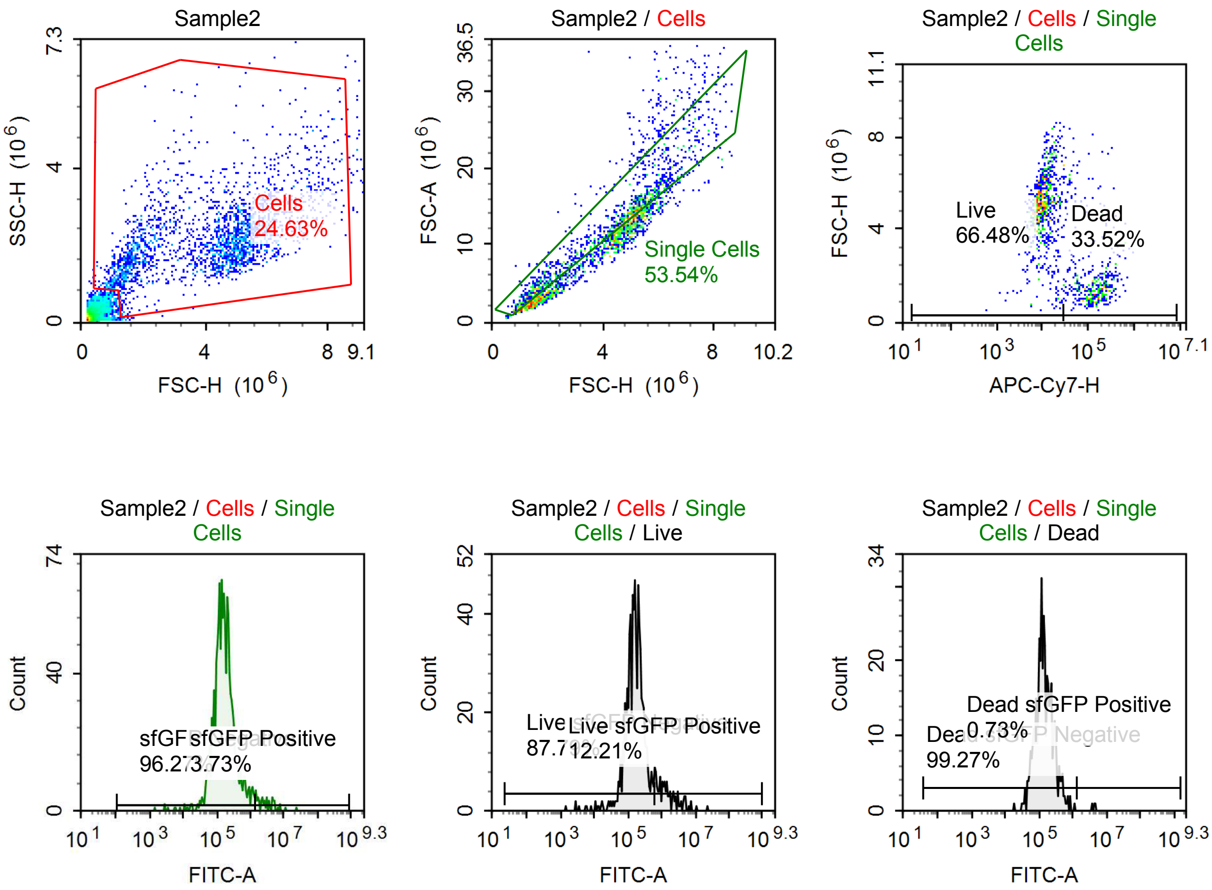

Sample Statistics of Sample2

| Gate | Count | % Parent | X | Y | Median X | Median Y |
|---|---|---|---|---|---|---|
| All | 9,343 | | | | | |
| Cells | 2,301 | 24.63 % | FSC-H | SSC-H | 4,548,276 | 2,085,336 |
| Single Cells | 1,232 | 53.54 % | FSC-H | FSC-A | 4,516,451 | 12,710,190 |
| Live | 819 | 66.48 % | APC-Cy7-H | FSC-H | 10,204 | 5,082,380 |
| Live sfGFP Negative | 719 | 87.79 % | FITC-A | Count | 160,020 | |
| Live sfGFP Positive | 100 | 12.21 % | FITC-A | Count | 1,380,876 | |
| Dead | 413 | 33.52 % | APC-Cy7-H | FSC-H | 136,792 | 1,554,543 |
| Dead sfGFP Negative | 410 | 99.27 % | FITC-A | Count | 135,623 | |
| Dead sfGFP Positive | 3 | 0.73 % | FITC-A | Count | 4,421,418 | |
| sfGFP Negative | 1,186 | 96.27 % | FITC-A | Count | 158,654 | |
| sfGFP Positive | 46 | 3.73 % | FITC-A | Count | 2,952,417 | |

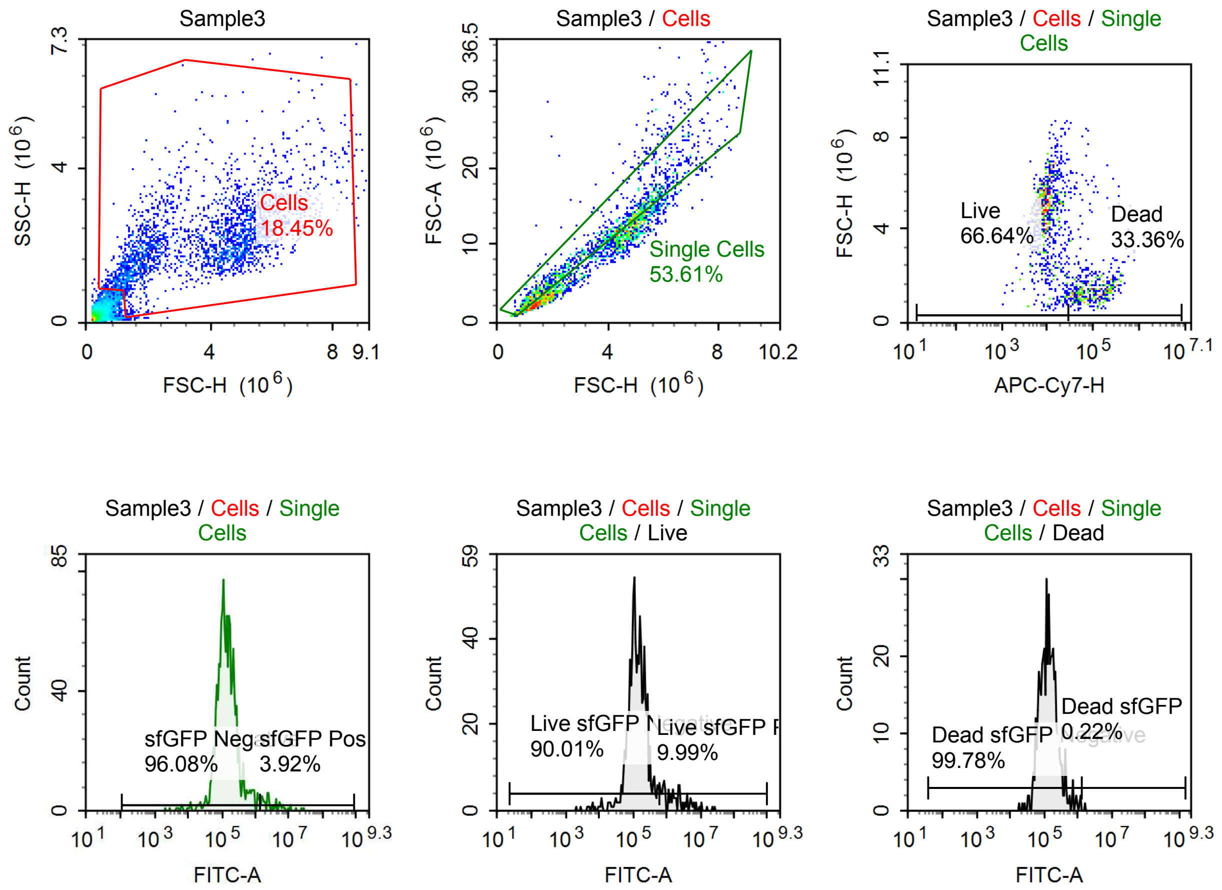

Sample Statistics of Sample3

| Gate | Count | % Parent | X | Y | Median X | Median Y |
|---|---|---|---|---|---|---|
| All | 13,667 | | | | | |
| Cells | 2,522 | 18.45 % | FSC-H | SSC-H | 4,003,029 | 2,006,488 |
| Single Cells | 1,352 | 53.61 % | FSC-H | FSC-A | 3,777,786 | 10,707,472 |
| Live | 901 | 66.64 % | APC-Cy7-H | FSC-H | 9,405 | 4,699,481 |
| Live sfGFP Negative | 811 | 90.01 % | FITC-A | Count | 130,405 | |
| Live sfGFP Positive | 90 | 9.99 % | FITC-A | Count | 1,769,188 | |
| Dead | 451 | 33.36 % | APC-Cy7-H | FSC-H | 92,542 | 1,456,560 |
| Dead sfGFP Negative | 450 | 99.78 % | FITC-A | Count | 133,021 | |
| Dead sfGFP Positive | 1 | 0.22 % | FITC-A | Count | 1,653,645 | |
| sfGFP Negative | 1,299 | 96.08 % | FITC-A | Count | 134,853 | |
| sfGFP Positive | 53 | 3.92 % | FITC-A | Count | 2,693,590 | |

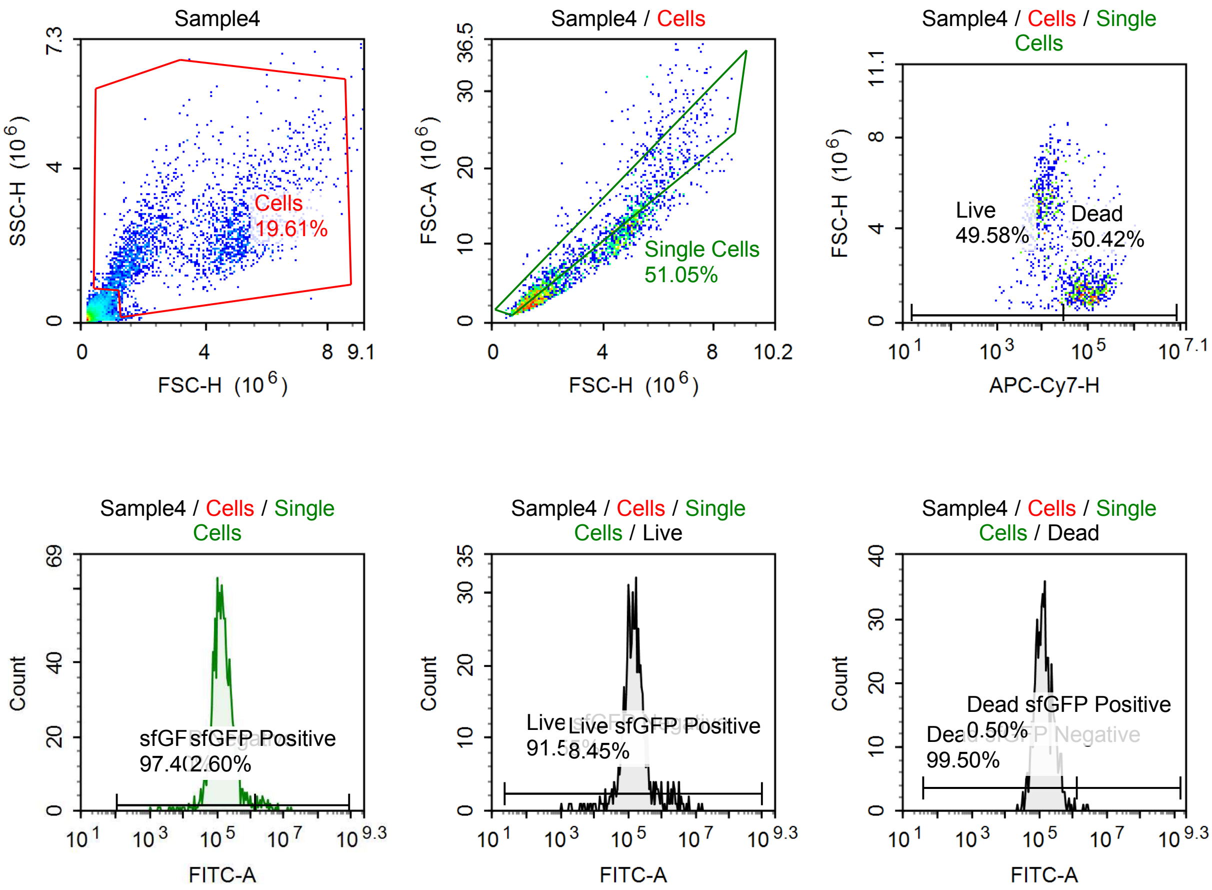

Sample Statistics of Sample4

| Gate | Count | % Parent | X | Y | Median X | Median Y |
|---|---|---|---|---|---|---|
| All | 11,930 | | | | | |
| Cells | 2,339 | 19.61 % | FSC-H | SSC-H | 3,382,350 | 2,048,012 |
| Single Cells | 1,194 | 51.05 % | FSC-H | FSC-A | 2,505,430 | 7,134,361 |
| Live | 592 | 49.58 % | APC-Cy7-H | FSC-H | 10,438 | 4,839,092 |
| Live sfGFP Negative | 542 | 91.55 % | FITC-A | Count | 141,373 | |
| Live sfGFP Positive | 50 | 8.45 % | FITC-A | Count | 1,897,888 | |
| Dead | 602 | 50.42 % | APC-Cy7-H | FSC-H | 97,187 | 1,487,324 |
| Dead sfGFP Negative | 599 | 99.50 % | FITC-A | Count | 133,015 | |
| Dead sfGFP Positive | 3 | 0.50 % | FITC-A | Count | 2,563,535 | |
| sfGFP Negative | 1,163 | 97.40 % | FITC-A | Count | 140,379 | |
| sfGFP Positive | 31 | 2.60 % | FITC-A | Count | 2,652,683 | |

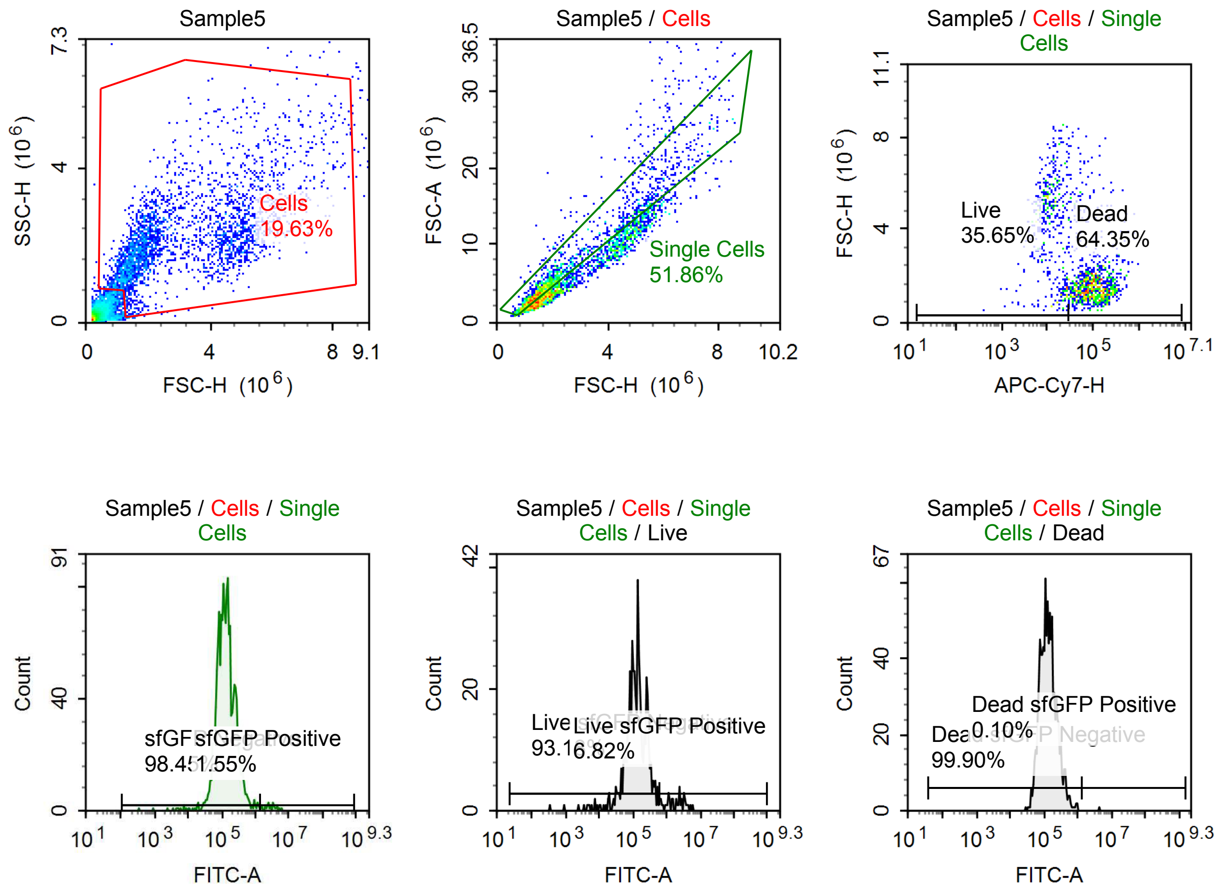

Sample Statistics of Sample5

| Gate | Count | % Parent | X | Y | Median X | Median Y |
|---|---|---|---|---|---|---|
| All | 14,548 | | | | | |
| Cells | 2,856 | 19.63 % | FSC-H | SSC-H | 2,267,572 | 2,060,907 |
| Single Cells | 1,481 | 51.86 % | FSC-H | FSC-A | 1,822,440 | 5,013,898 |
| Live | 528 | 35.65 % | APC-Cy7-H | FSC-H | 12,064 | 4,532,332 |
| Live sfGFP Negative | 492 | 93.18 % | FITC-A | Count | 128,118 | |
| Live sfGFP Positive | 36 | 6.82 % | FITC-A | Count | 2,178,750 | |
| Dead | 953 | 64.35 % | APC-Cy7-H | FSC-H | 97,054 | 1,529,983 |
| Dead sfGFP Negative | 952 | 99.90 % | FITC-A | Count | 128,698 | |
| Dead sfGFP Positive | 1 | 0.10 % | FITC-A | Count | 4,353,932 | |
| sfGFP Negative | 1,458 | 98.45 % | FITC-A | Count | 129,363 | |
| sfGFP Positive | 23 | 1.55 % | FITC-A | Count | 2,863,902 | |

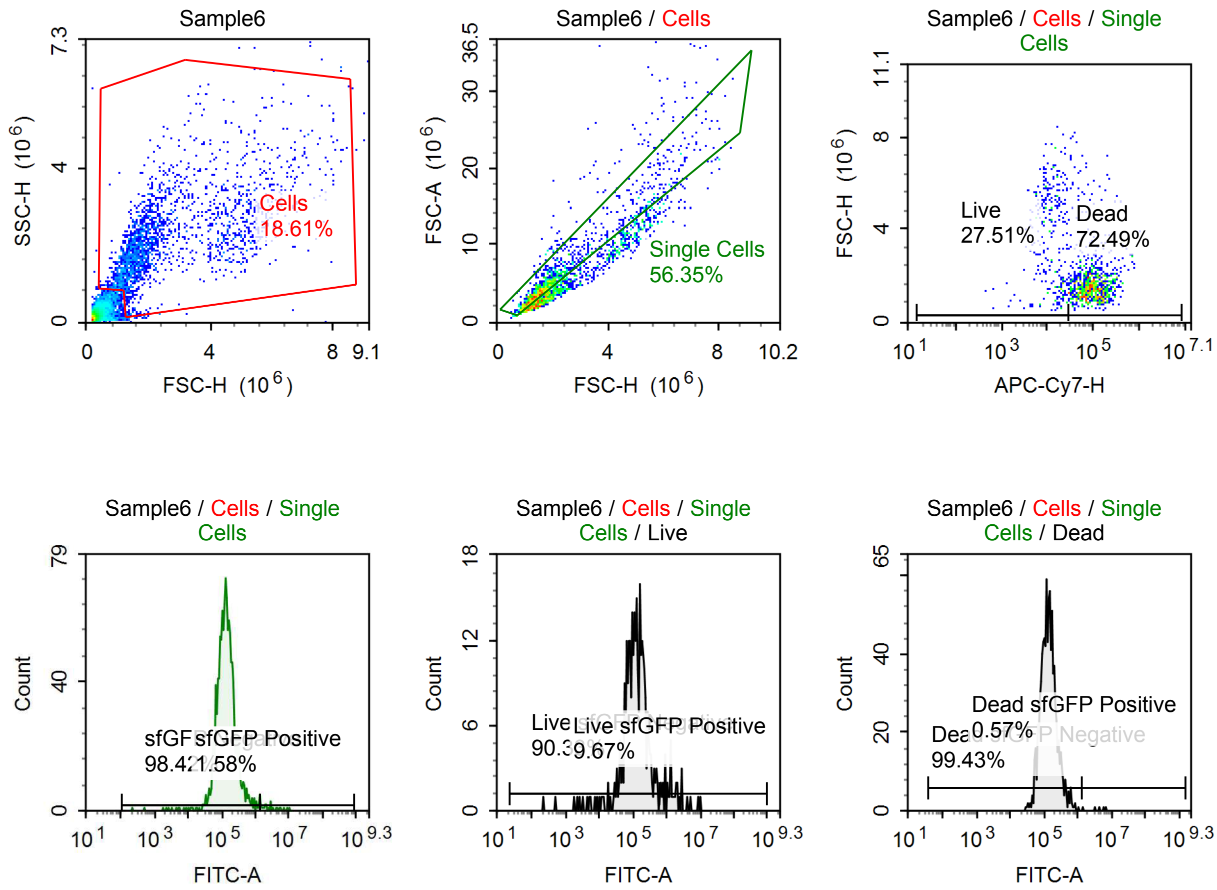

Sample Statistics of Sample6

| Gate | Count | % Parent | X | Y | Median X | Median Y |
|---|---|---|---|---|---|---|
| All | 11,475 | | | | | |
| Cells | 2,135 | 18.61 % | FSC-H | SSC-H | 1,919,969 | 2,019,565 |
| Single Cells | 1,203 | 56.35 % | FSC-H | FSC-A | 1,650,922 | 4,453,020 |
| Live | 331 | 27.51 % | APC-Cy7-H | FSC-H | 13,241 | 3,780,952 |
| Live sfGFP Negative | 299 | 90.33 % | FITC-A | Count | 117,102 | |
| Live sfGFP Positive | 32 | 9.67 % | FITC-A | Count | 1,391,297 | |
| Dead | 872 | 72.49 % | APC-Cy7-H | FSC-H | 89,685 | 1,502,724 |
| Dead sfGFP Negative | 867 | 99.43 % | FITC-A | Count | 133,988 | |
| Dead sfGFP Positive | 5 | 0.57 % | FITC-A | Count | 4,500,736 | |
| sfGFP Negative | 1,184 | 98.42 % | FITC-A | Count | 131,671 | |
| sfGFP Positive | 19 | 1.58 % | FITC-A | Count | 2,841,054 | |

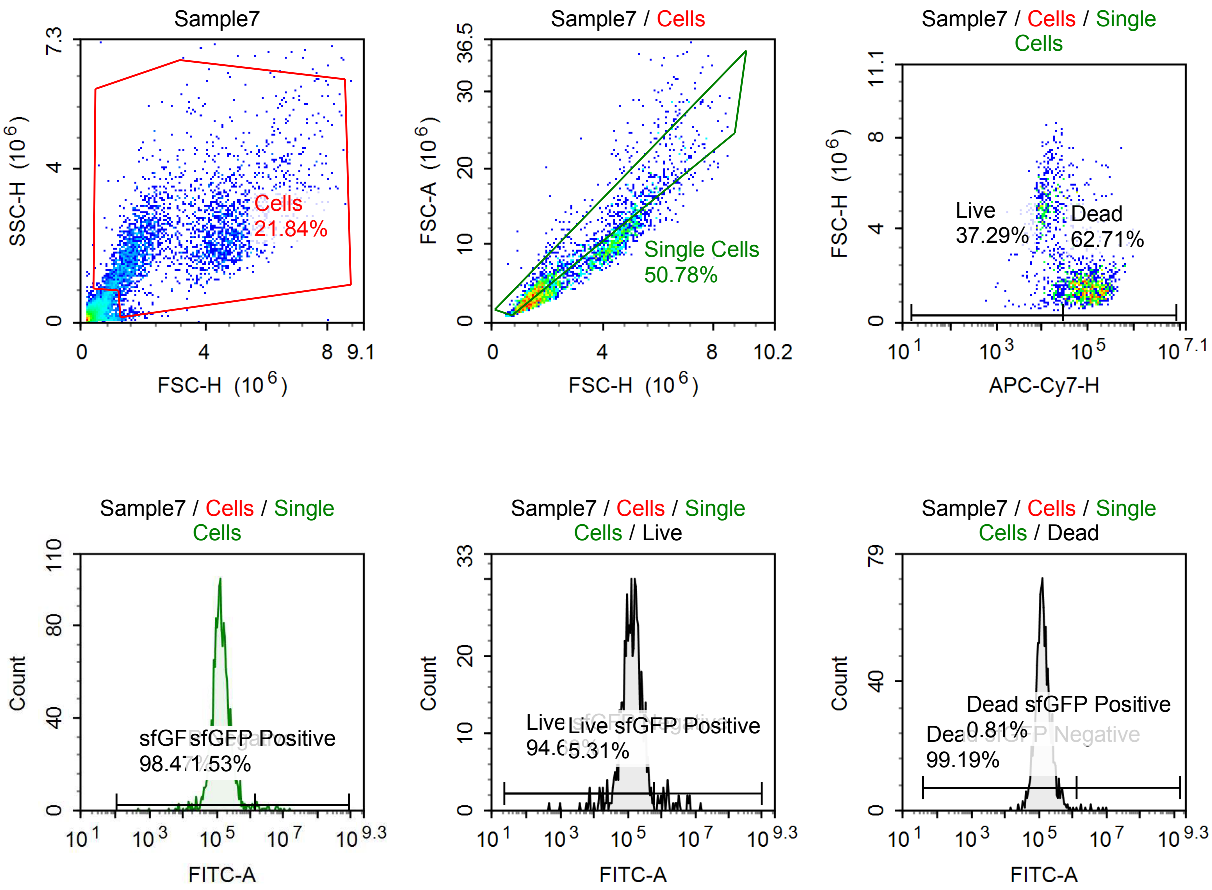

Sample Statistics of Sample7

| Gate | Count | % Parent | X | Y | Median X | Median Y |
|---|---|---|---|---|---|---|
| All | 14,120 | | | | | |
| Cells | 3,084 | 21.84 % | FSC-H | SSC-H | 2,401,664 | 2,018,002 |
| Single Cells | 1,566 | 50.78 % | FSC-H | FSC-A | 1,765,293 | 4,953,357 |
| Live | 584 | 37.29 % | APC-Cy7-H | FSC-H | 12,507 | 4,426,433 |
| Live sfGFP Negative | 553 | 94.69 % | FITC-A | Count | 133,158 | |
| Live sfGFP Positive | 31 | 5.31 % | FITC-A | Count | 1,563,314 | |
| Dead | 982 | 62.71 % | APC-Cy7-H | FSC-H | 101,059 | 1,526,969 |
| Dead sfGFP Negative | 974 | 99.19 % | FITC-A | Count | 128,858 | |
| Dead sfGFP Positive | 8 | 0.81 % | FITC-A | Count | 3,483,568 | |
| sfGFP Negative | 1,542 | 98.47 % | FITC-A | Count | 130,819 | |
| sfGFP Positive | 24 | 1.53 % | FITC-A | Count | 3,322,897 | |

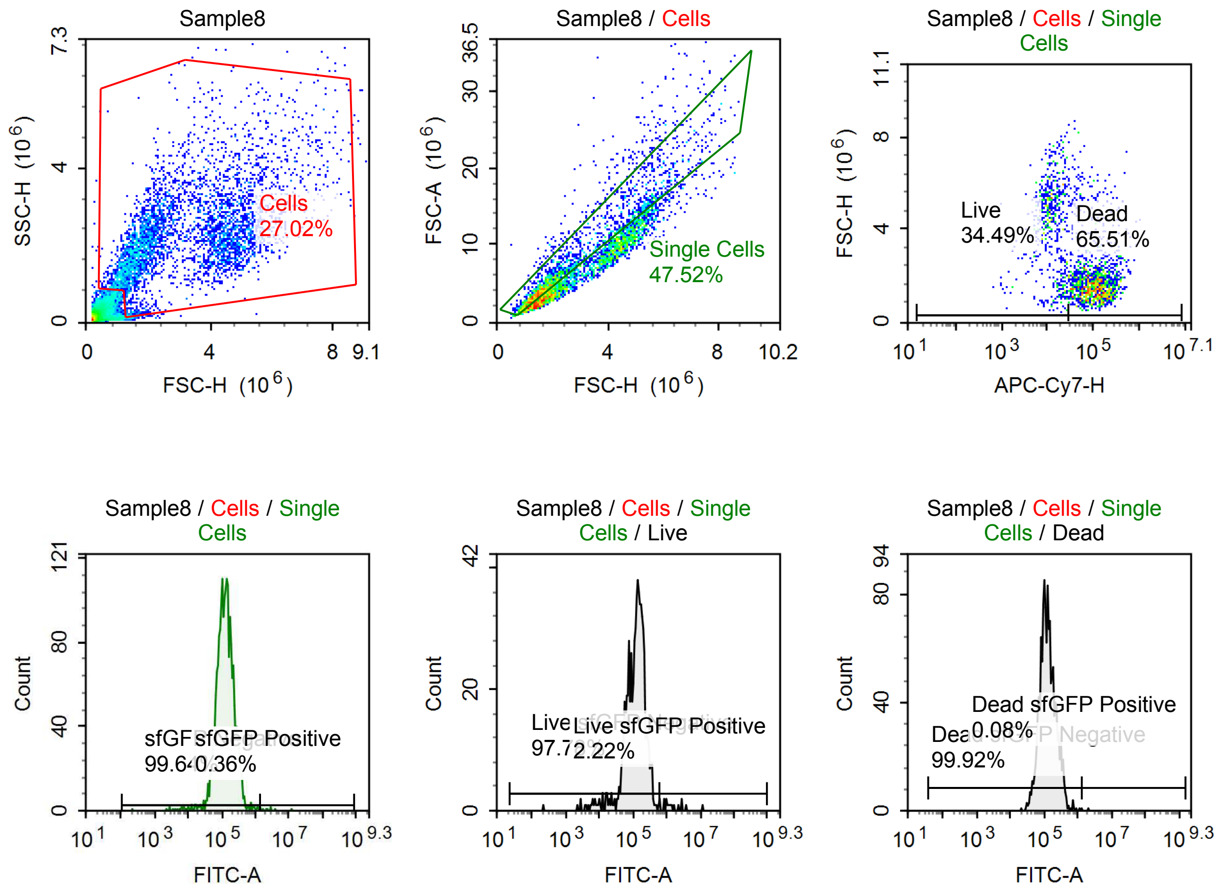

Sample Statistics of Sample8

| Gate | Count | % Parent | X | Y | Median X | Median Y |
|---|---|---|---|---|---|---|
| All | 15,291 | | | | | |
| Cells | 4,131 | 27.02 % | FSC-H | SSC-H | 2,716,400 | 2,136,761 |
| Single Cells | 1,963 | 47.52 % | FSC-H | FSC-A | 1,955,596 | 5,564,606 |
| Live | 677 | 34.49 % | APC-Cy7-H | FSC-H | 12,295 | 4,486,998 |
| Live sfGFP Negative | 662 | 97.78 % | FITC-A | Count | 129,390 | |
| Live sfGFP Positive | 15 | 2.22 % | FITC-A | Count | 1,372,440 | |
| Dead | 1,286 | 65.51 % | APC-Cy7-H | FSC-H | 107,675 | 1,610,611 |
| Dead sfGFP Negative | 1,285 | 99.92 % | FITC-A | Count | 124,849 | |
| Dead sfGFP Positive | 1 | 0.08 % | FITC-A | Count | 2,085,663 | |
| sfGFP Negative | 1,956 | 99.64 % | FITC-A | Count | 126,008 | |
| sfGFP Positive | 7 | 0.36 % | FITC-A | Count | 2,800,284 | |

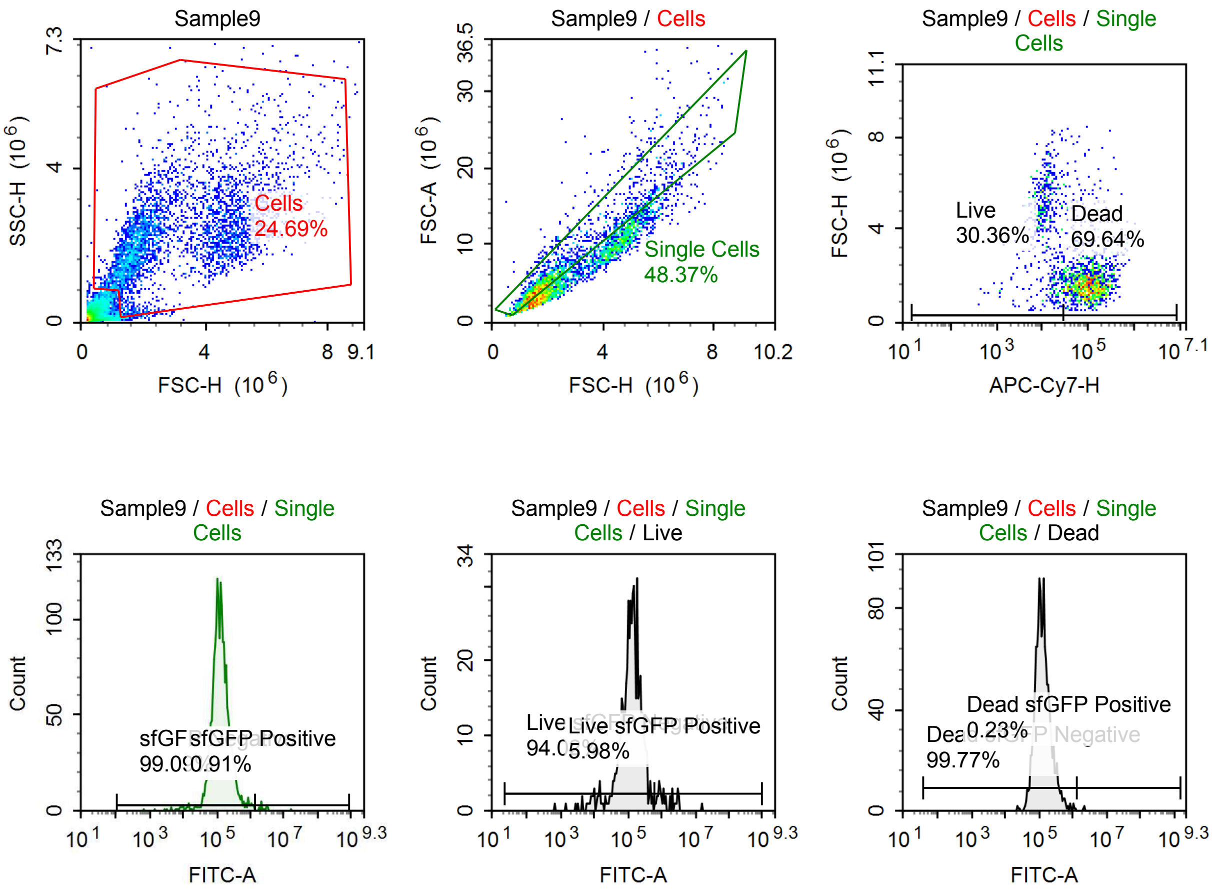

Sample Statistics of Sample9

| Gate | Count | % Parent | X | Y | Median X | Median Y |
|---|---|---|---|---|---|---|
| All | 15,688 | | | | | |
| Cells | 3,874 | 24.69 % | FSC-H | SSC-H | 2,333,080 | 2,163,926 |
| Single Cells | 1,874 | 48.37 % | FSC-H | FSC-A | 1,855,247 | 5,062,630 |
| Live | 569 | 30.36 % | APC-Cy7-H | FSC-H | 12,266 | 4,350,528 |
| Live sfGFP Negative | 535 | 94.02 % | FITC-A | Count | 128,120 | |
| Live sfGFP Positive | 34 | 5.98 % | FITC-A | Count | 1,192,942 | |
| Dead | 1,305 | 69.64 % | APC-Cy7-H | FSC-H | 102,444 | 1,647,728 |
| Dead sfGFP Negative | 1,302 | 99.77 % | FITC-A | Count | 124,291 | |
| Dead sfGFP Positive | 3 | 0.23 % | FITC-A | Count | 2,161,132 | |
| sfGFP Negative | 1,857 | 99.09 % | FITC-A | Count | 126,779 | |
| sfGFP Positive | 17 | 0.91 % | FITC-A | Count | 2,361,864 | |

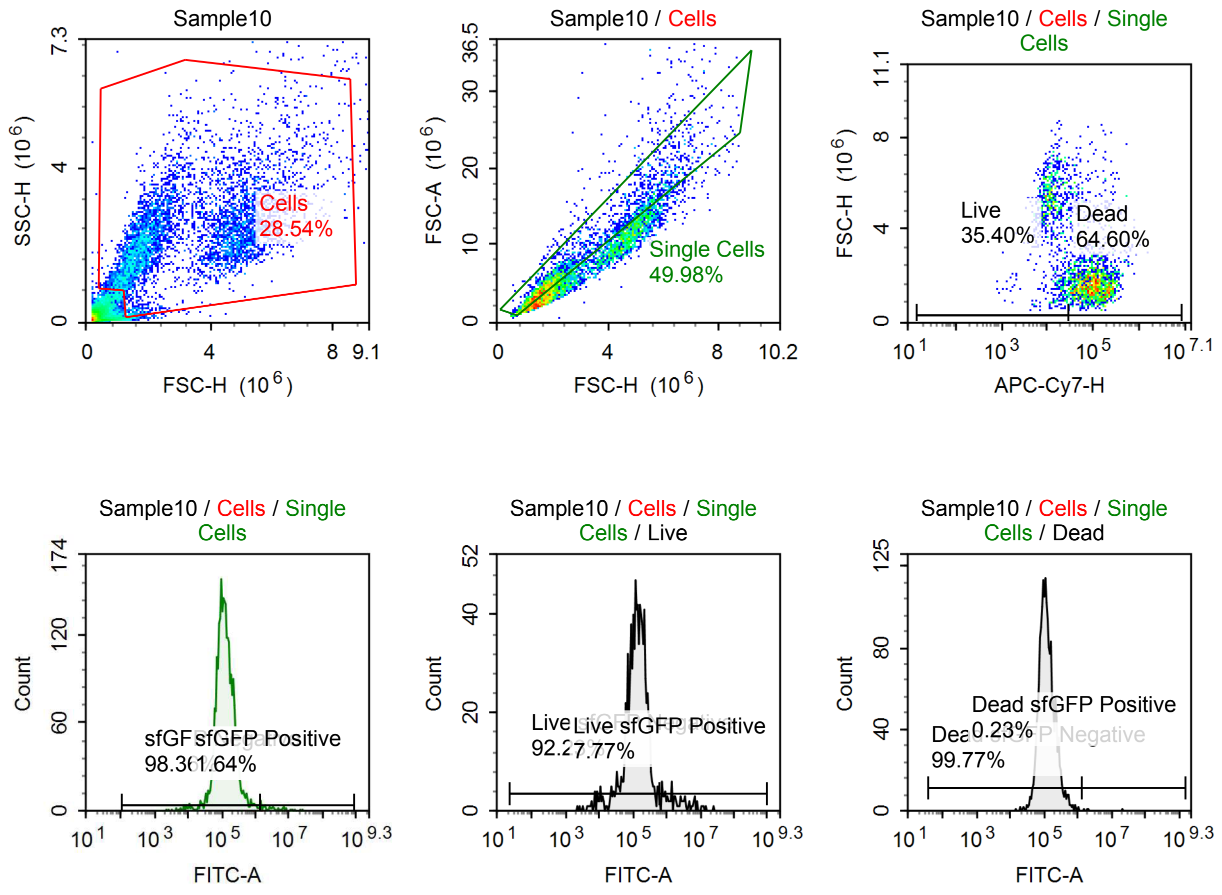

Sample Statistics of Sample10

| Gate | Count | % Parent | X | Y | Median X | Median Y |
|---|---|---|---|---|---|---|
| All | 18,848 | | | | | |
| Cells | 5,380 | 28.54 % | FSC-H | SSC-H | 2,583,960 | 2,175,512 |
| Single Cells | 2,689 | 49.98 % | FSC-H | FSC-A | 1,976,841 | 5,522,745 |
| Live | 952 | 35.40 % | APC-Cy7-H | FSC-H | 12,969 | 4,731,396 |
| Live sfGFP Negative | 878 | 92.23 % | FITC-A | Count | 130,371 | |
| Live sfGFP Positive | 74 | 7.77 % | FITC-A | Count | 1,795,311 | |
| Dead | 1,737 | 64.60 % | APC-Cy7-H | FSC-H | 98,624 | 1,687,893 |
| Dead sfGFP Negative | 1,733 | 99.77 % | FITC-A | Count | 114,921 | |
| Dead sfGFP Positive | 4 | 0.23 % | FITC-A | Count | 1,955,546 | |
| sfGFP Negative | 2,645 | 98.36 % | FITC-A | Count | 120,547 | |
| sfGFP Positive | 44 | 1.64 % | FITC-A | Count | 3,406,159 | |

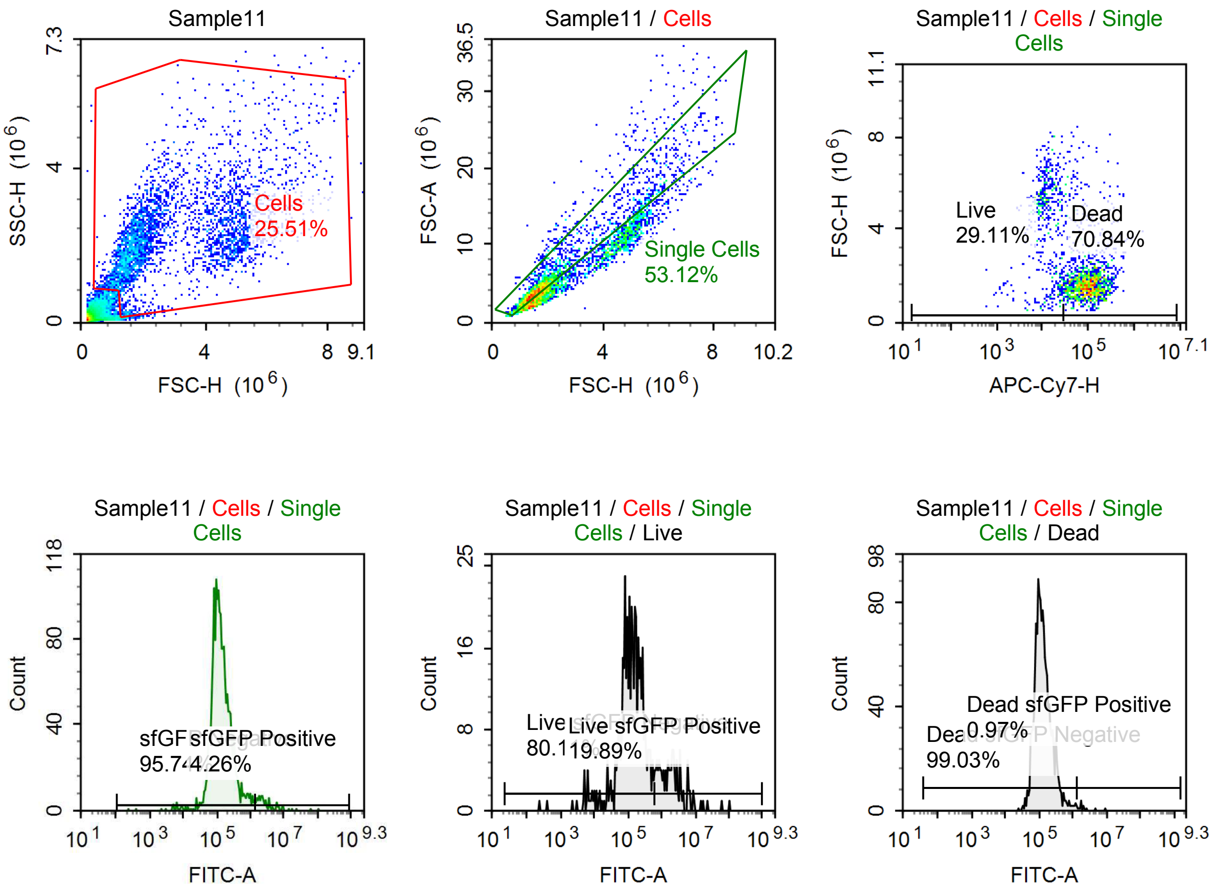

Sample Statistics of Sample11

| Gate | Count | % Parent | X | Y | Median X | Median Y |
|---|---|---|---|---|---|---|
| All | 14,024 | | | | | |
| Cells | 3,577 | 25.51 % | FSC-H | SSC-H | 2,218,348 | 2,137,472 |
| Single Cells | 1,900 | 53.12 % | FSC-H | FSC-A | 1,785,034 | 4,776,427 |
| Live | 553 | 29.11 % | APC-Cy7-H | FSC-H | 12,582 | 4,687,129 |
| Live sfGFP Negative | 443 | 80.11 % | FITC-A | Count | 127,967 | |
| Live sfGFP Positive | 110 | 19.89 % | FITC-A | Count | 1,957,620 | |
| Dead | 1,346 | 70.84 % | APC-Cy7-H | FSC-H | 98,580 | 1,637,965 |
| Dead sfGFP Negative | 1,333 | 99.03 % | FITC-A | Count | 121,401 | |
| Dead sfGFP Positive | 13 | 0.97 % | FITC-A | Count | 1,780,635 | |
| sfGFP Negative | 1,819 | 95.74 % | FITC-A | Count | 124,455 | |
| sfGFP Positive | 81 | 4.26 % | FITC-A | Count | 2,630,361 | |

#### p060 mRNA [100 ng] + TET [10 uM]

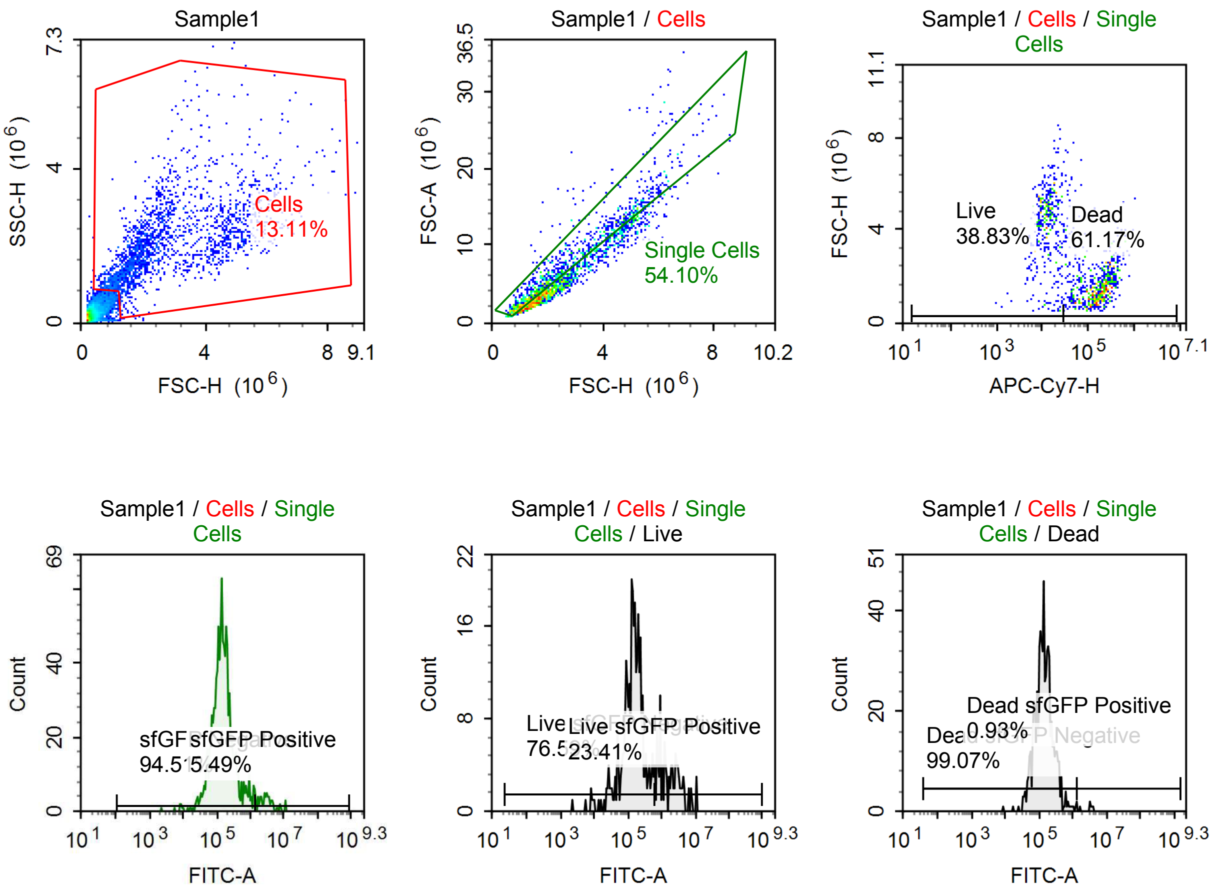

Sample Statistics of Sample1

| Gate | Count | % Parent | X | Y | Median X | Median Y |
|---|---|---|---|---|---|---|
| All | 14,894 | | | | | |
| Cells | 1,952 | 13.11 % | FSC-H | SSC-H | 2,257,766 | 1,961,131 |
| Single Cells | 1,056 | 54.10 % | FSC-H | FSC-A | 2,063,413 | 5,949,742 |
| Live | 410 | 38.83 % | APC-Cy7-H | FSC-H | 12,362 | 4,449,794 |
| Live sfGFP Negative | 314 | 76.59 % | FITC-A | Count | 145,749 | |
| Live sfGFP Positive | 96 | 23.41 % | FITC-A | Count | 1,795,926 | |
| Dead | 646 | 61.17 % | APC-Cy7-H | FSC-H | 163,436 | 1,518,501 |
| Dead sfGFP Negative | 640 | 99.07 % | FITC-A | Count | 137,575 | |
| Dead sfGFP Positive | 6 | 0.93 % | FITC-A | Count | 3,353,049 | |
| sfGFP Negative | 998 | 94.51 % | FITC-A | Count | 143,619 | |
| sfGFP Positive | 58 | 5.49 % | FITC-A | Count | 3,092,047 | |

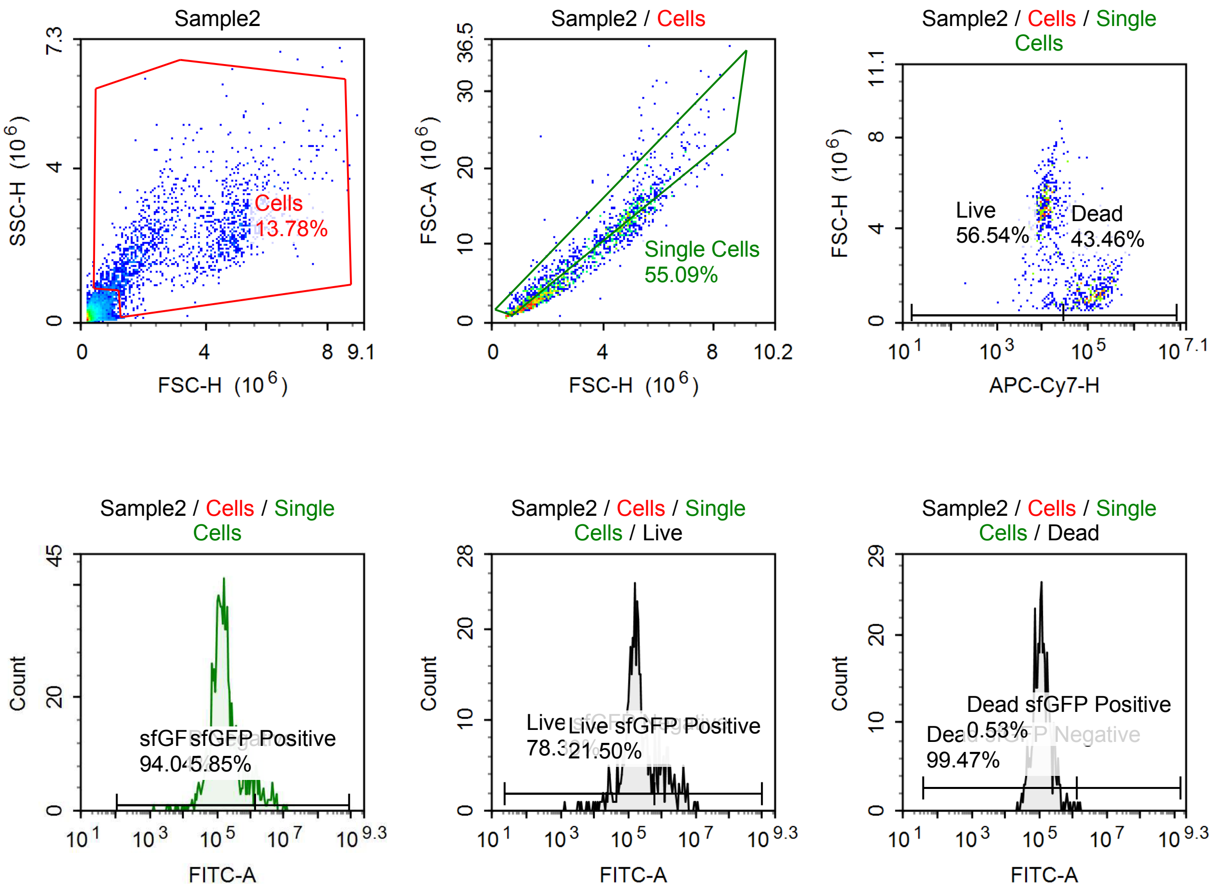

Sample Statistics of Sample2

| Gate | Count | % Parent | X | Y | Median X | Median Y |
|---|---|---|---|---|---|---|
| All | 11,485 | | | | | |
| Cells | 1,583 | 13.78 % | FSC-H | SSC-H | 2,794,288 | 2,049,793 |
| Single Cells | 872 | 55.09 % | FSC-H | FSC-A | 2,939,446 | 8,567,150 |
| Live | 493 | 56.54 % | APC-Cy7-H | FSC-H | 11,157 | 4,713,691 |
| Live sfGFP Negative | 386 | 78.30 % | FITC-A | Count | 161,313 | |
| Live sfGFP Positive | 106 | 21.50 % | FITC-A | Count | 1,522,771 | |
| Dead | 379 | 43.46 % | APC-Cy7-H | FSC-H | 118,772 | 1,407,847 |
| Dead sfGFP Negative | 377 | 99.47 % | FITC-A | Count | 116,694 | |
| Dead sfGFP Positive | 2 | 0.53 % | FITC-A | Count | 1,531,440 | |
| sfGFP Negative | 820 | 94.04 % | FITC-A | Count | 144,300 | |
| sfGFP Positive | 51 | 5.85 % | FITC-A | Count | 3,175,045 | |

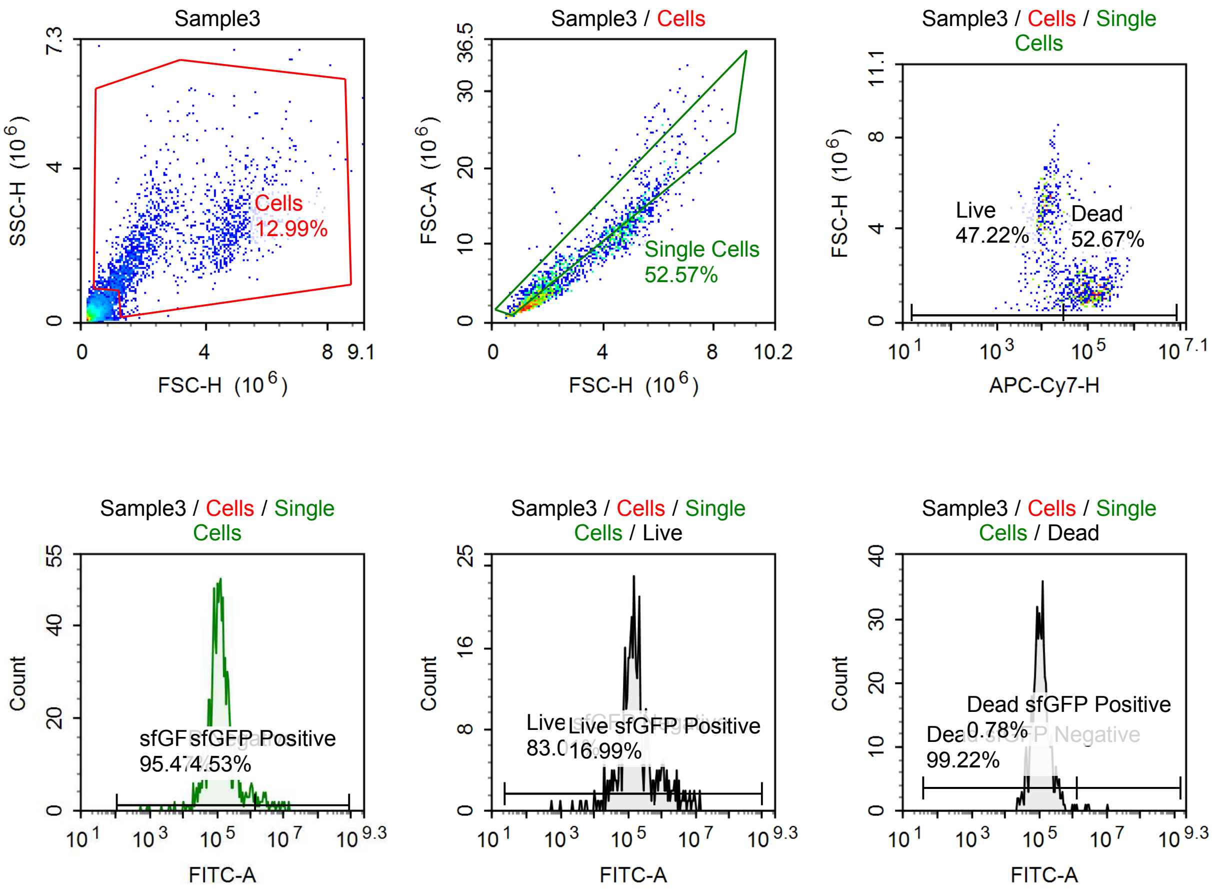

Sample Statistics of Sample3

| Gate | Count | % Parent | X | Y | Median X | Median Y |
|---|---|---|---|---|---|---|
| All | 14,238 | | | | | |
| Cells | 1,849 | 12.99 % | FSC-H | SSC-H | 2,211,757 | 1,979,017 |
| Single Cells | 972 | 52.57 % | FSC-H | FSC-A | 2,025,221 | 5,757,033 |
| Live | 459 | 47.22 % | APC-Cy7-H | FSC-H | 11,791 | 4,392,634 |
| Live sfGFP Negative | 381 | 83.01 % | FITC-A | Count | 134,832 | |
| Live sfGFP Positive | 78 | 16.99 % | FITC-A | Count | 1,614,240 | |
| Dead | 512 | 52.67 % | APC-Cy7-H | FSC-H | 104,089 | 1,414,940 |
| Dead sfGFP Negative | 508 | 99.22 % | FITC-A | Count | 108,791 | |
| Dead sfGFP Positive | 4 | 0.78 % | FITC-A | Count | 2,802,142 | |
| sfGFP Negative | 928 | 95.47 % | FITC-A | Count | 122,363 | |
| sfGFP Positive | 44 | 4.53 % | FITC-A | Count | 3,052,315 | |

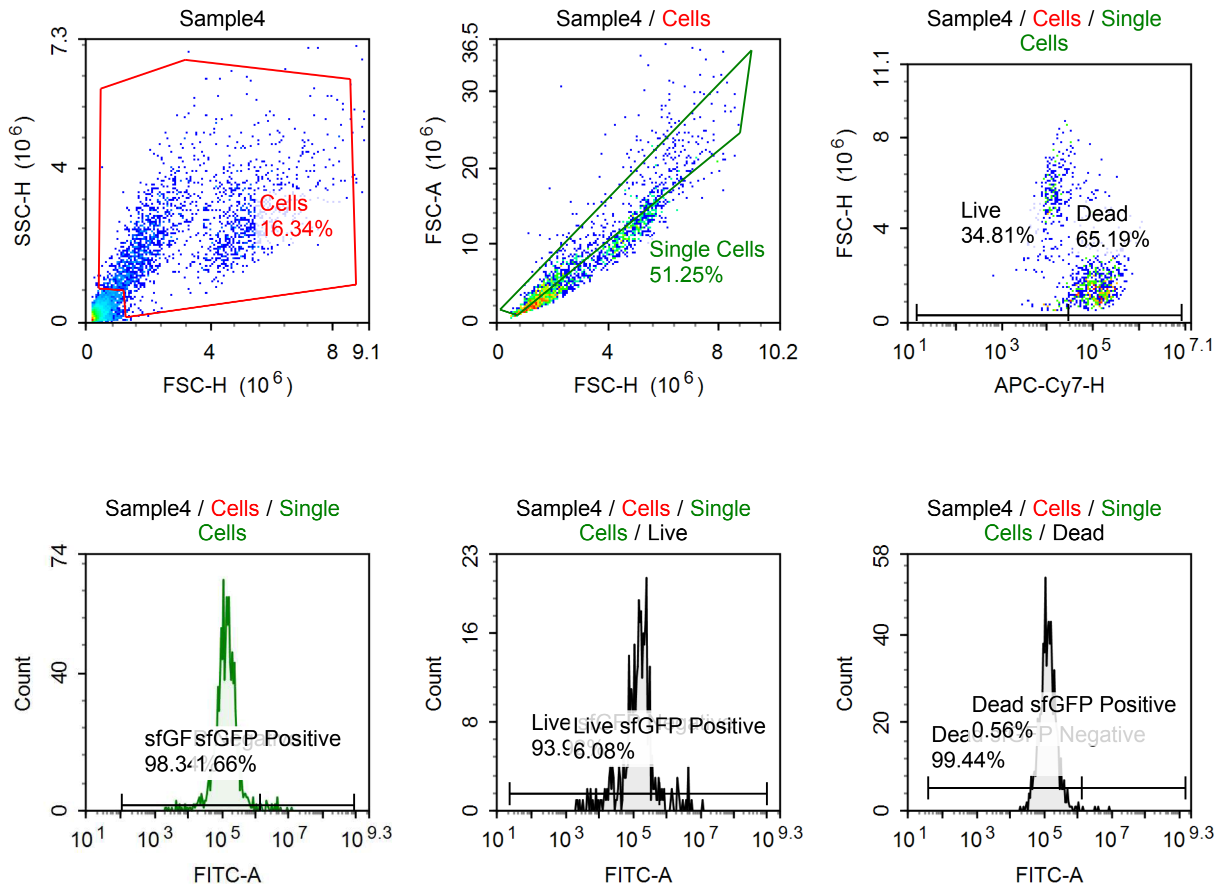

Sample Statistics of Sample4

| Gate | Count | % Parent | X | Y | Median X | Median Y |
|---|---|---|---|---|---|---|
| All | 12,971 | | | | | |
| Cells | 2,119 | 16.34 % | FSC-H | SSC-H | 2,467,819 | 2,189,385 |
| Single Cells | 1,086 | 51.25 % | FSC-H | FSC-A | 1,984,846 | 5,569,283 |
| Live | 378 | 34.81 % | APC-Cy7-H | FSC-H | 12,842 | 4,961,071 |
| Live sfGFP Negative | 355 | 93.92 % | FITC-A | Count | 149,475 | |
| Live sfGFP Positive | 23 | 6.08 % | FITC-A | Count | 2,157,805 | |
| Dead | 708 | 65.19 % | APC-Cy7-H | FSC-H | 112,325 | 1,601,681 |
| Dead sfGFP Negative | 704 | 99.44 % | FITC-A | Count | 129,378 | |
| Dead sfGFP Positive | 4 | 0.56 % | FITC-A | Count | 3,508,222 | |
| sfGFP Negative | 1,068 | 98.34 % | FITC-A | Count | 136,022 | |
| sfGFP Positive | 18 | 1.66 % | FITC-A | Count | 3,424,577 | |

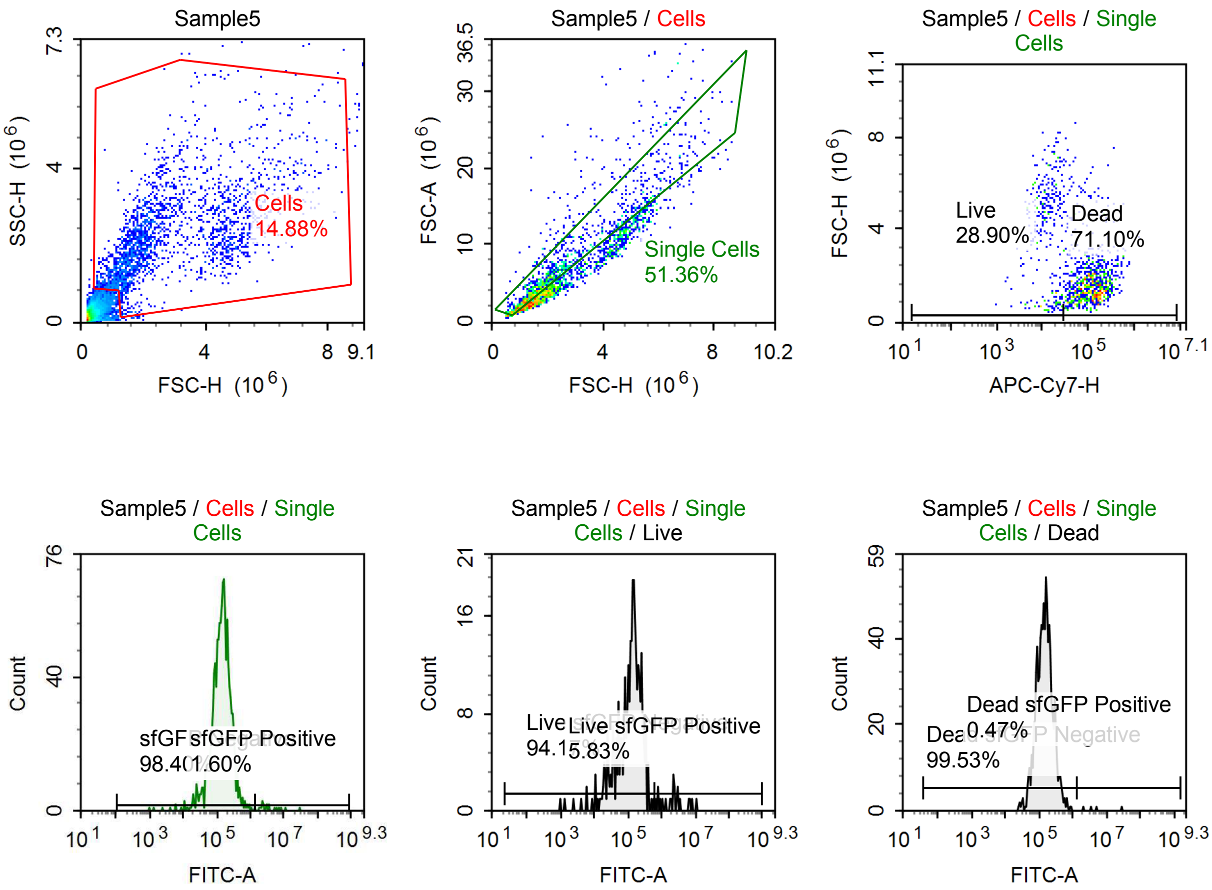

Sample Statistics of Sample5

| Gate | Count | % Parent | X | Y | Median X | Median Y |
|---|---|---|---|---|---|---|
| All | 15,534 | | | | | |
| Cells | 2,311 | 14.88 % | FSC-H | SSC-H | 2,119,006 | 2,061,336 |
| Single Cells | 1,187 | 51.36 % | FSC-H | FSC-A | 1,772,602 | 4,887,220 |
| Live | 343 | 28.90 % | APC-Cy7-H | FSC-H | 13,437 | 4,588,240 |
| Live sfGFP Negative | 323 | 94.17 % | FITC-A | Count | 130,635 | |
| Live sfGFP Positive | 20 | 5.83 % | FITC-A | Count | 2,362,784 | |
| Dead | 844 | 71.10 % | APC-Cy7-H | FSC-H | 115,884 | 1,572,562 |
| Dead sfGFP Negative | 840 | 99.53 % | FITC-A | Count | 147,619 | |
| Dead sfGFP Positive | 4 | 0.47 % | FITC-A | Count | 4,279,060 | |
| sfGFP Negative | 1,168 | 98.40 % | FITC-A | Count | 141,881 | |
| sfGFP Positive | 19 | 1.60 % | FITC-A | Count | 2,978,629 | |

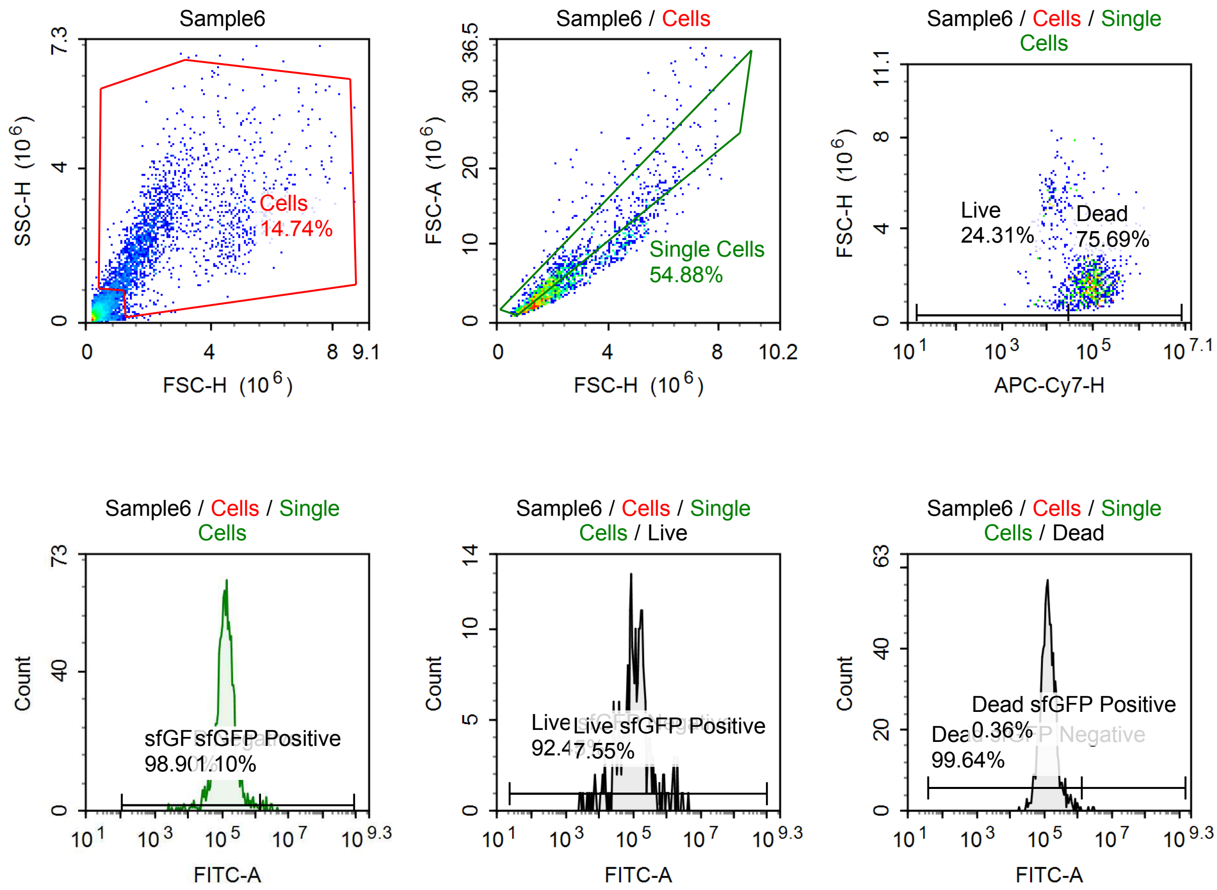

Sample Statistics of Sample6

| Gate | Count | % Parent | X | Y | Median X | Median Y |
|---|---|---|---|---|---|---|
| All | 13,469 | | | | | |
| Cells | 1,986 | 14.74 % | FSC-H | SSC-H | 1,988,133 | 2,111,019 |
| Single Cells | 1,090 | 54.88 % | FSC-H | FSC-A | 1,784,067 | 4,782,148 |
| Live | 265 | 24.31 % | APC-Cy7-H | FSC-H | 13,971 | 3,553,038 |
| Live sfGFP Negative | 245 | 92.45 % | FITC-A | Count | 102,068 | |
| Live sfGFP Positive | 20 | 7.55 % | FITC-A | Count | 1,539,131 | |
| Dead | 825 | 75.69 % | APC-Cy7-H | FSC-H | 96,614 | 1,700,193 |
| Dead sfGFP Negative | 822 | 99.64 % | FITC-A | Count | 130,499 | |
| Dead sfGFP Positive | 3 | 0.36 % | FITC-A | Count | 2,812,989 | |
| sfGFP Negative | 1,078 | 98.90 % | FITC-A | Count | 127,155 | |
| sfGFP Positive | 12 | 1.10 % | FITC-A | Count | 2,147,345 | |

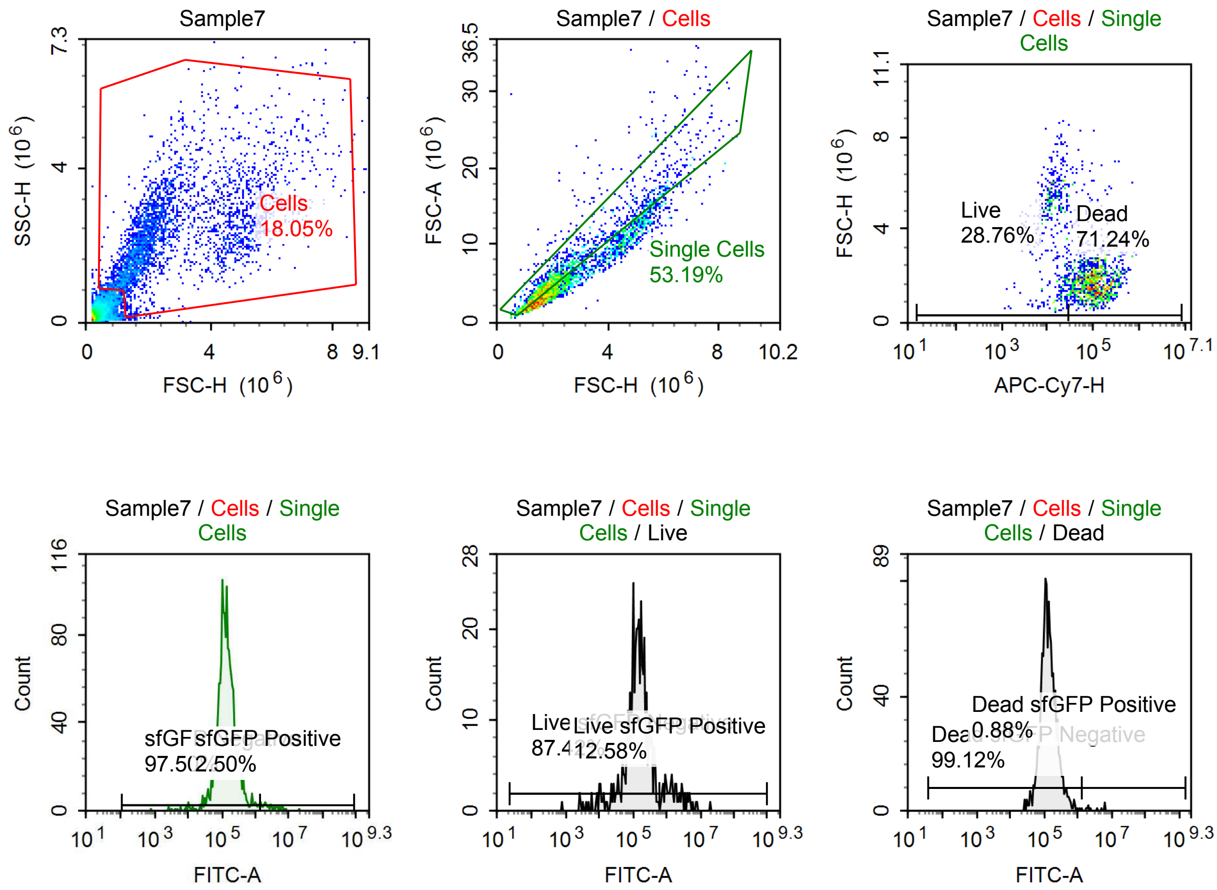

Sample Statistics of Sample7

| Gate | Count | % Parent | X | Y | Median X | Median Y |
|---|---|---|---|---|---|---|
| All | 16,699 | | | | | |
| Cells | 3,014 | 18.05 % | FSC-H | SSC-H | 2,190,256 | 2,208,951 |
| Single Cells | 1,603 | 53.19 % | FSC-H | FSC-A | 1,908,873 | 5,312,310 |
| Live | 461 | 28.76 % | APC-Cy7-H | FSC-H | 15,015 | 4,465,498 |
| Live sfGFP Negative | 403 | 87.42 % | FITC-A | Count | 135,645 | |
| Live sfGFP Positive | 58 | 12.58 % | FITC-A | Count | 1,637,985 | |
| Dead | 1,142 | 71.24 % | APC-Cy7-H | FSC-H | 102,476 | 1,736,803 |
| Dead sfGFP Negative | 1,132 | 99.12 % | FITC-A | Count | 130,535 | |
| Dead sfGFP Positive | 10 | 0.88 % | FITC-A | Count | 3,098,456 | |
| sfGFP Negative | 1,563 | 97.50 % | FITC-A | Count | 133,597 | |
| sfGFP Positive | 40 | 2.50 % | FITC-A | Count | 2,855,051 | |

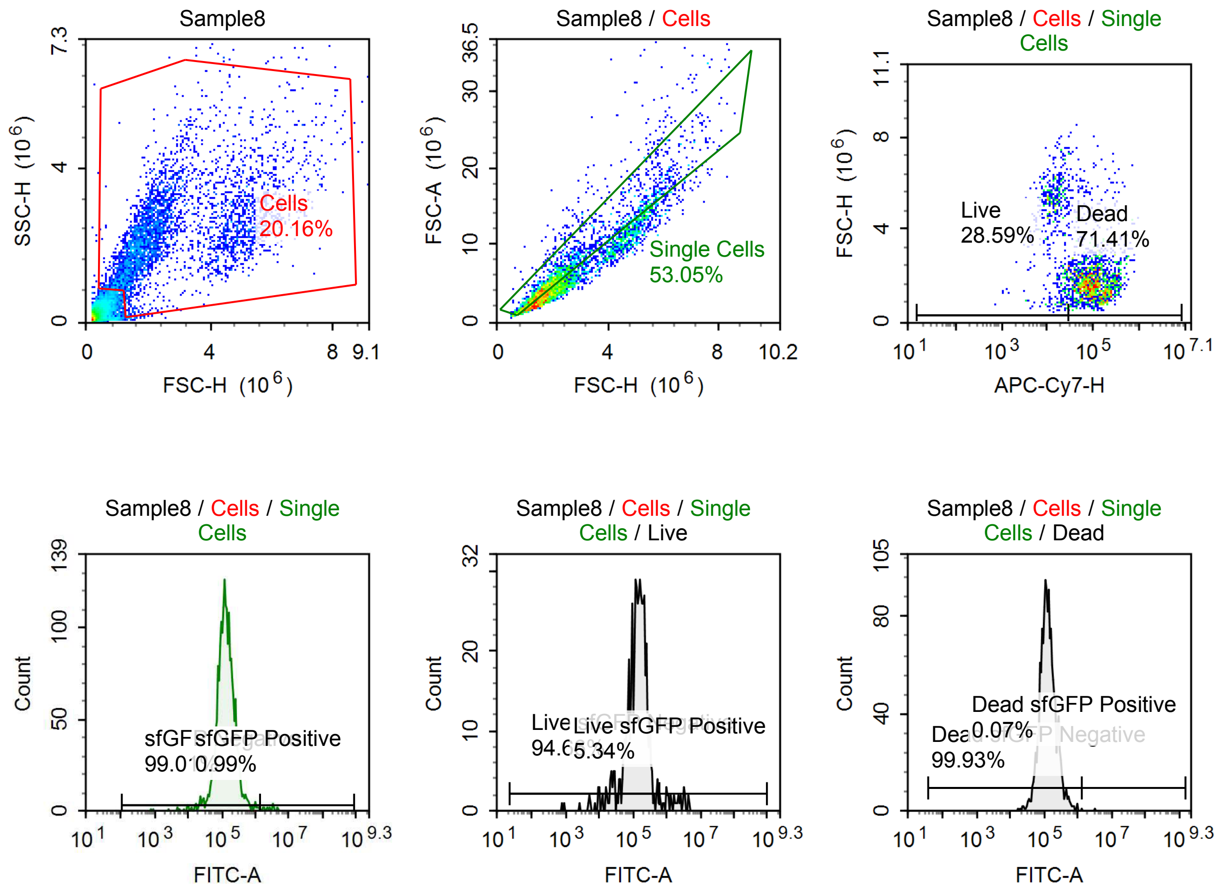

Sample Statistics of Sample8

| Gate | Count | % Parent | X | Y | Median X | Median Y |
|---|---|---|---|---|---|---|
| All | 18,975 | | | | | |
| Cells | 3,825 | 20.16 % | FSC-H | SSC-H | 2,242,040 | 2,239,017 |
| Single Cells | 2,029 | 53.05 % | FSC-H | FSC-A | 1,964,367 | 5,298,882 |
| Live | 580 | 28.59 % | APC-Cy7-H | FSC-H | 15,324 | 4,637,168 |
| Live sfGFP Negative | 549 | 94.66 % | FITC-A | Count | 136,177 | |
| Live sfGFP Positive | 31 | 5.34 % | FITC-A | Count | 1,880,206 | |
| Dead | 1,449 | 71.41 % | APC-Cy7-H | FSC-H | 101,312 | 1,741,600 |
| Dead sfGFP Negative | 1,448 | 99.93 % | FITC-A | Count | 125,658 | |
| Dead sfGFP Positive | 1 | 0.07 % | FITC-A | Count | 3,225,982 | |
| sfGFP Negative | 2,009 | 99.01 % | FITC-A | Count | 128,555 | |
| sfGFP Positive | 20 | 0.99 % | FITC-A | Count | 2,848,019 | |

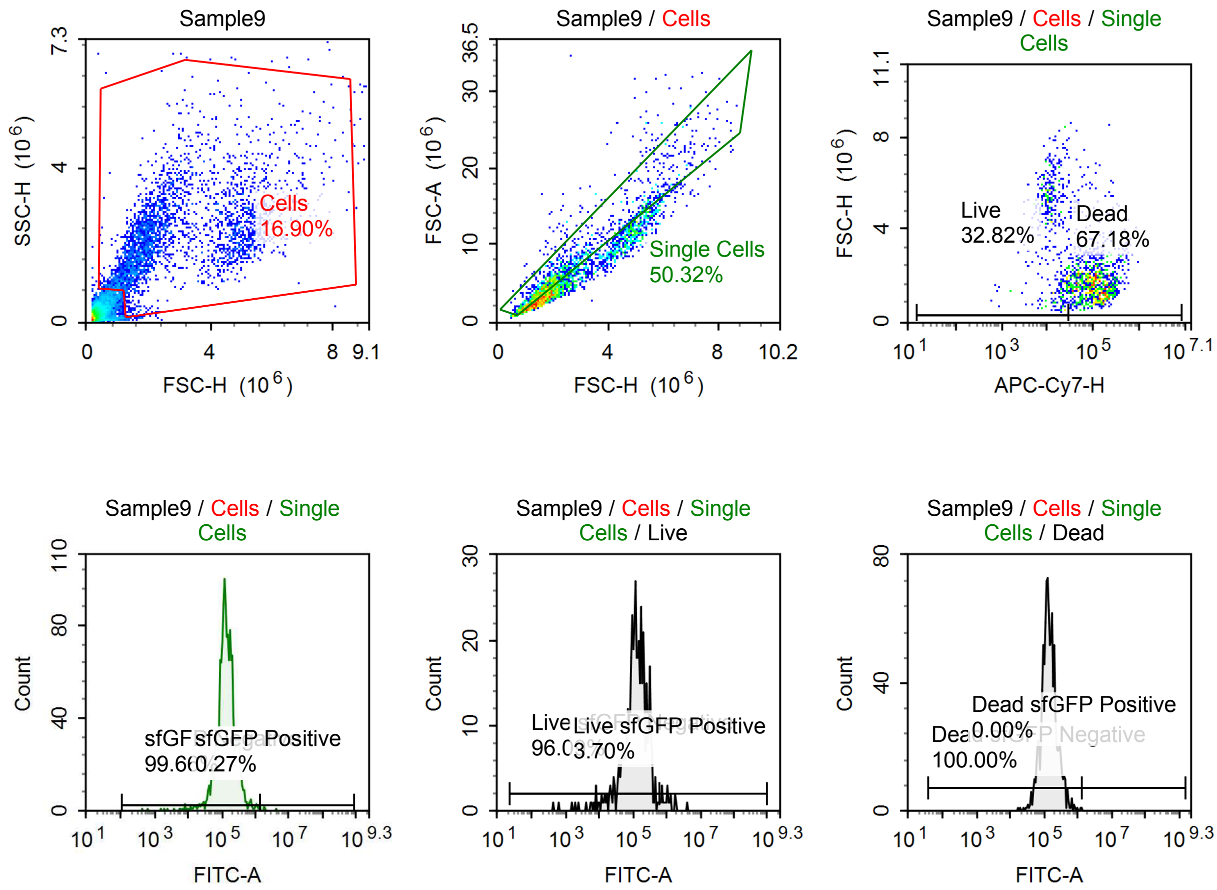

Sample Statistics of Sample9

| Gate | Count | % Parent | X | Y | Median X | Median Y |
|---|---|---|---|---|---|---|
| All | 17,417 | | | | | |
| Cells | 2,943 | 16.90 % | FSC-H | SSC-H | 2,230,673 | 2,155,028 |
| Single Cells | 1,481 | 50.32 % | FSC-H | FSC-A | 1,892,921 | 5,238,941 |
| Live | 486 | 32.82 % | APC-Cy7-H | FSC-H | 13,276 | 4,348,550 |
| Live sfGFP Negative | 467 | 96.09 % | FITC-A | Count | 124,281 | |
| Live sfGFP Positive | 18 | 3.70 % | FITC-A | Count | 1,095,712 | |
| Dead | 995 | 67.18 % | APC-Cy7-H | FSC-H | 104,955 | 1,706,825 |
| Dead sfGFP Negative | 995 | 100.00 % | FITC-A | Count | 135,308 | |
| Dead sfGFP Positive | 0 | 0.00 % | FITC-A | Count | 0 | |
| sfGFP Negative | 1,476 | 99.66 % | FITC-A | Count | 133,601 | |
| sfGFP Positive | 4 | 0.27 % | FITC-A | Count | 1,912,733 | |

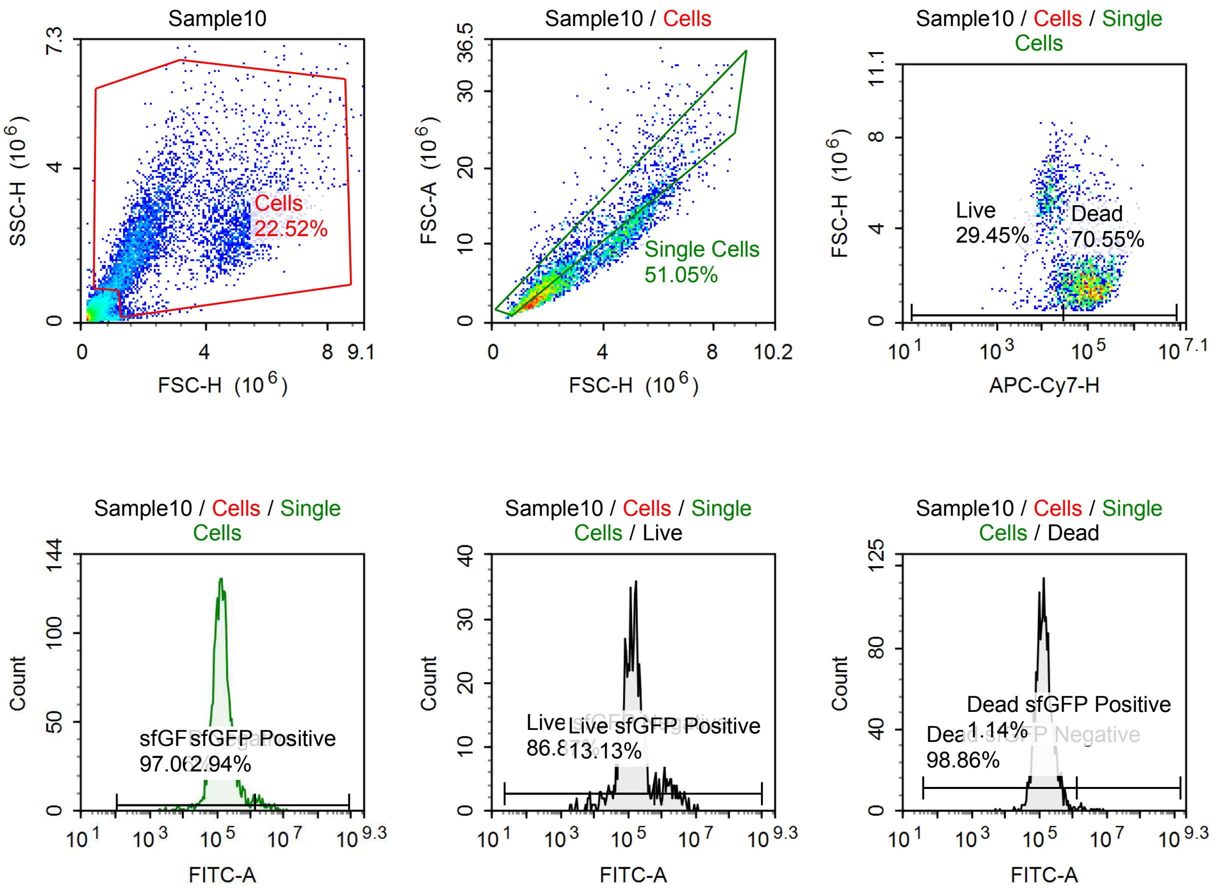

Sample Statistics of Sample10

| Gate | Count | % Parent | X | Y | Median X | Median Y |
|---|---|---|---|---|---|---|
| All | 21,590 | | | | | |
| Cells | 4,862 | 22.52 % | FSC-H | SSC-H | 2,308,835 | 2,186,062 |
| Single Cells | 2,482 | 51.05 % | FSC-H | FSC-A | 1,924,723 | 5,344,966 |
| Live | 731 | 29.45 % | APC-Cy7-H | FSC-H | 14,912 | 4,735,125 |
| Live sfGFP Negative | 635 | 86.87 % | FITC-A | Count | 130,677 | |
| Live sfGFP Positive | 96 | 13.13 % | FITC-A | Count | 1,742,514 | |
| Dead | 1,751 | 70.55 % | APC-Cy7-H | FSC-H | 104,694 | 1,711,390 |
| Dead sfGFP Negative | 1,731 | 98.86 % | FITC-A | Count | 136,566 | |
| Dead sfGFP Positive | 20 | 1.14 % | FITC-A | Count | 2,150,594 | |
| sfGFP Negative | 2,409 | 97.06 % | FITC-A | Count | 137,069 | |
| sfGFP Positive | 73 | 2.94 % | FITC-A | Count | 2,485,614 | |

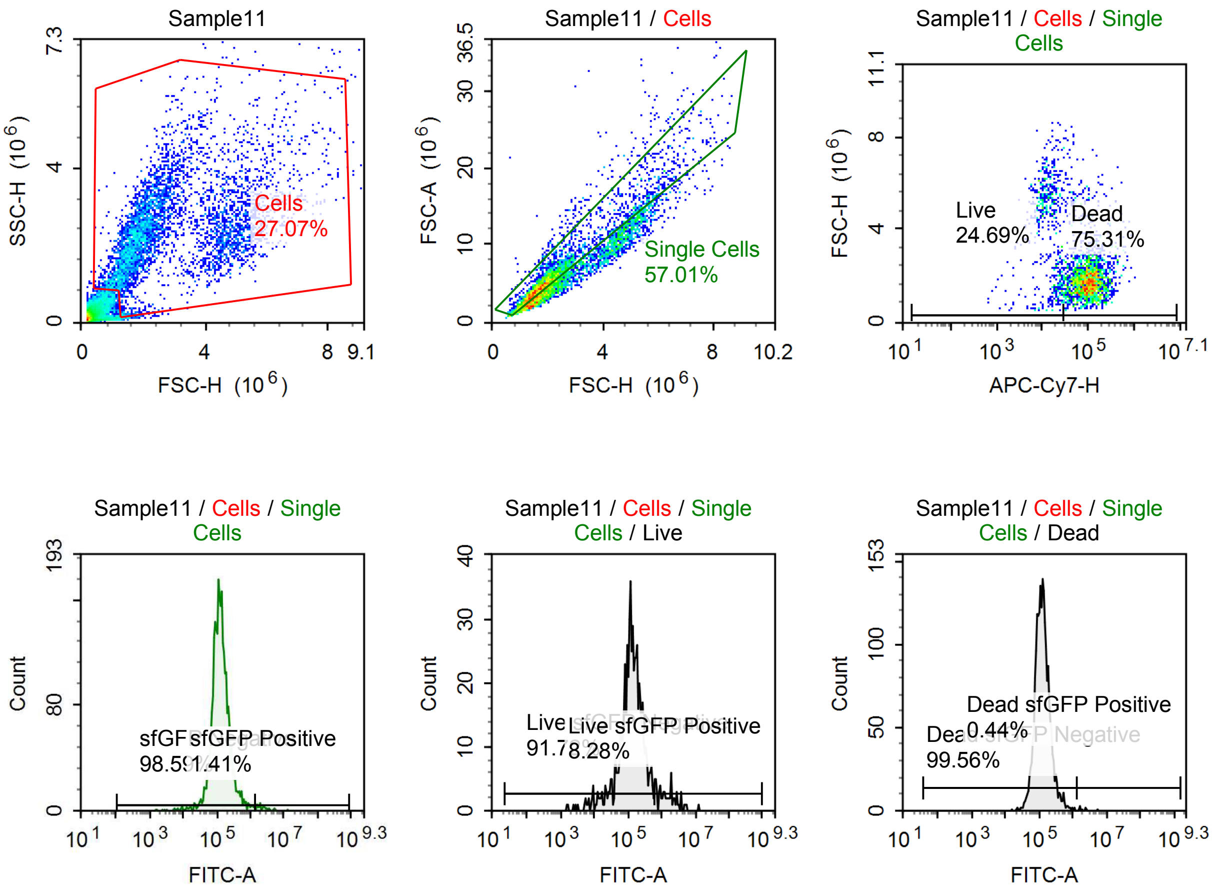

Sample Statistics of Sample11

| Gate | Count | % Parent | X | Y | Median X | Median Y |
|---|---|---|---|---|---|---|
| All | 17,427 | | | | | |
| Cells | 4,717 | 27.07 % | FSC-H | SSC-H | 2,231,164 | 2,308,305 |
| Single Cells | 2,689 | 57.01 % | FSC-H | FSC-A | 1,917,270 | 5,335,408 |
| Live | 664 | 24.69 % | APC-Cy7-H | FSC-H | 14,605 | 4,598,144 |
| Live sfGFP Negative | 609 | 91.72 % | FITC-A | Count | 127,471 | |
| Live sfGFP Positive | 55 | 8.28 % | FITC-A | Count | 1,641,784 | |
| Dead | 2,025 | 75.31 % | APC-Cy7-H | FSC-H | 98,785 | 1,780,420 |
| Dead sfGFP Negative | 2,016 | 99.56 % | FITC-A | Count | 123,579 | |
| Dead sfGFP Positive | 9 | 0.44 % | FITC-A | Count | 1,968,294 | |
| sfGFP Negative | 2,651 | 98.59 % | FITC-A | Count | 125,067 | |
| sfGFP Positive | 38 | 1.41 % | FITC-A | Count | 2,314,046 | |

#### p005 mRNA [100 ng]

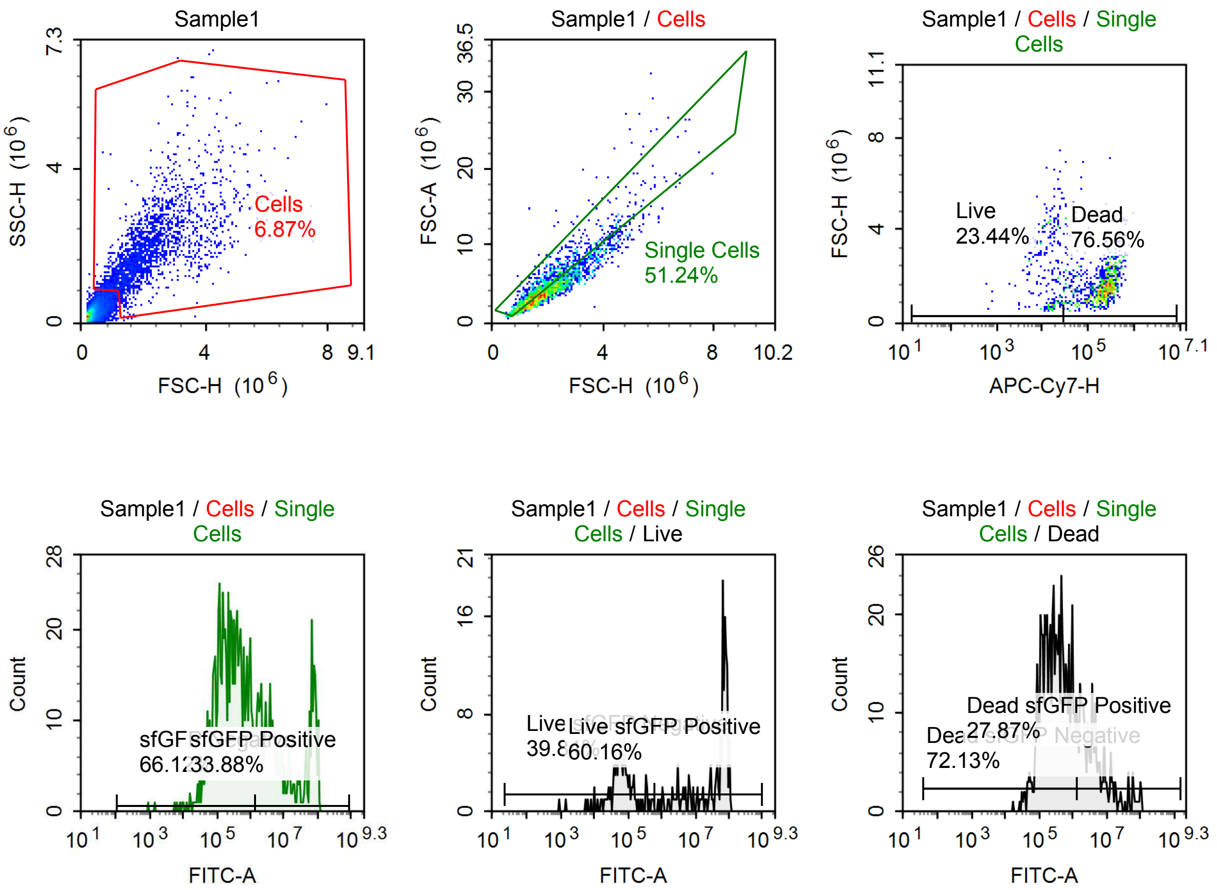

Sample Statistics of Sample1

| Gate | Count | % Parent | X | Y | Median X | Median Y |
|---|---|---|---|---|---|---|
| All | 31,040 | | | | | |
| Cells | 2,131 | 6.87 % | FSC-H | SSC-H | 1,917,821 | 1,753,100 |
| Single Cells | 1,092 | 51.24 % | FSC-H | FSC-A | 1,602,339 | 4,103,261 |
| Live | 256 | 23.44 % | APC-Cy7-H | FSC-H | 14,760 | 2,375,070 |
| Live sfGFP Negative | 102 | 39.84 % | FITC-A | Count | 64,068 | |
| Live sfGFP Positive | 154 | 60.16 % | FITC-A | Count | 64,711,284 | |
| Dead | 836 | 76.56 % | APC-Cy7-H | FSC-H | 220,568 | 1,544,339 |
| Dead sfGFP Negative | 603 | 72.13 % | FITC-A | Count | 280,966 | |
| Dead sfGFP Positive | 233 | 27.87 % | FITC-A | Count | 3,883,921 | |
| sfGFP Negative | 722 | 66.12 % | FITC-A | Count | 254,699 | |
| sfGFP Positive | 370 | 33.88 % | FITC-A | Count | 8,320,205 | |

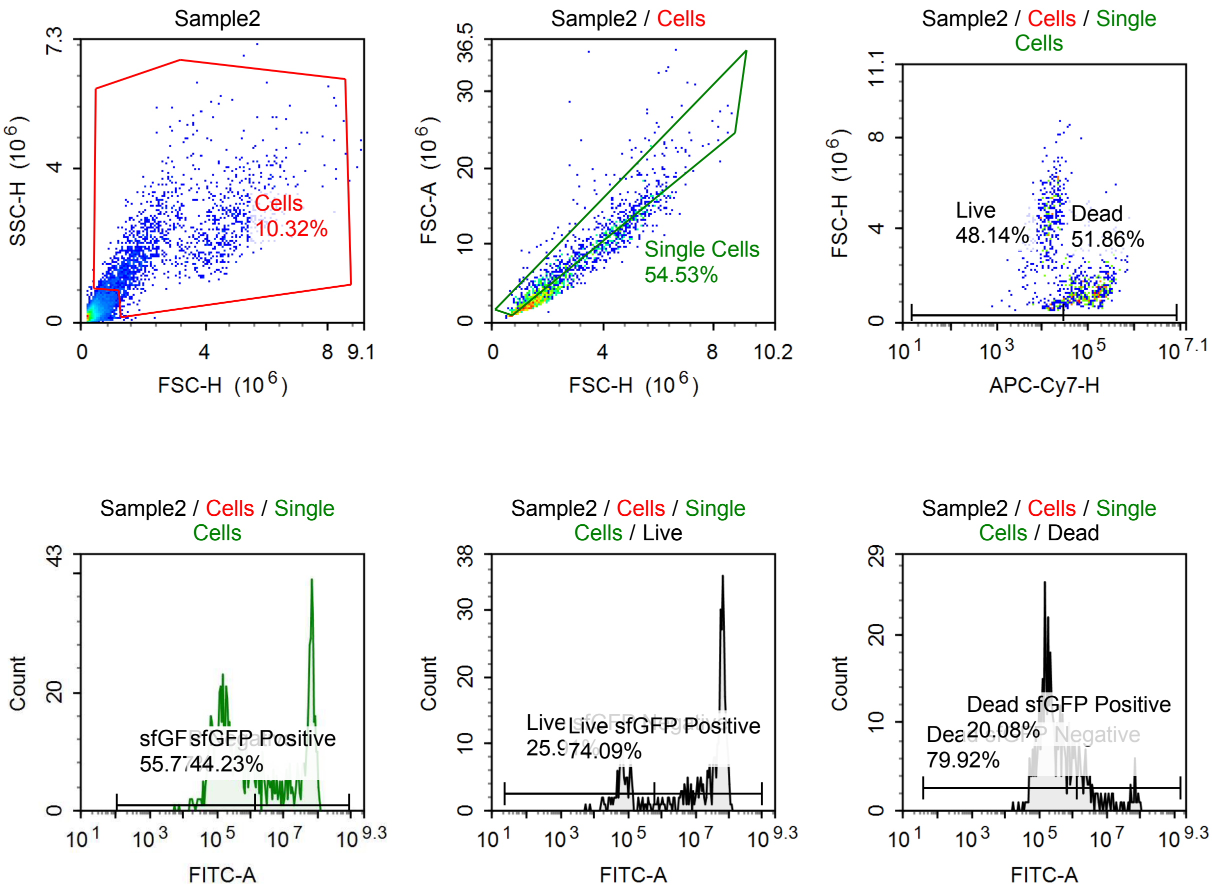

Sample Statistics of Sample2

| Gate | Count | % Parent | X | Y | Median X | Median Y |
|---|---|---|---|---|---|---|
| All | 17,234 | | | | | |
| Cells | 1,779 | 10.32 % | FSC-H | SSC-H | 2,089,418 | 1,829,500 |
| Single Cells | 970 | 54.53 % | FSC-H | FSC-A | 1,925,899 | 5,397,248 |
| Live | 467 | 48.14 % | APC-Cy7-H | FSC-H | 13,616 | 4,039,696 |
| Live sfGFP Negative | 121 | 25.91 % | FITC-A | Count | 71,248 | |
| Live sfGFP Positive | 346 | 74.09 % | FITC-A | Count | 55,135,624 | |
| Dead | 503 | 51.86 % | APC-Cy7-H | FSC-H | 109,289 | 1,426,106 |
| Dead sfGFP Negative | 402 | 79.92 % | FITC-A | Count | 208,320 | |
| Dead sfGFP Positive | 101 | 20.08 % | FITC-A | Count | 3,334,701 | |
| sfGFP Negative | 541 | 55.77 % | FITC-A | Count | 175,168 | |
| sfGFP Positive | 429 | 44.23 % | FITC-A | Count | 49,033,568 | |

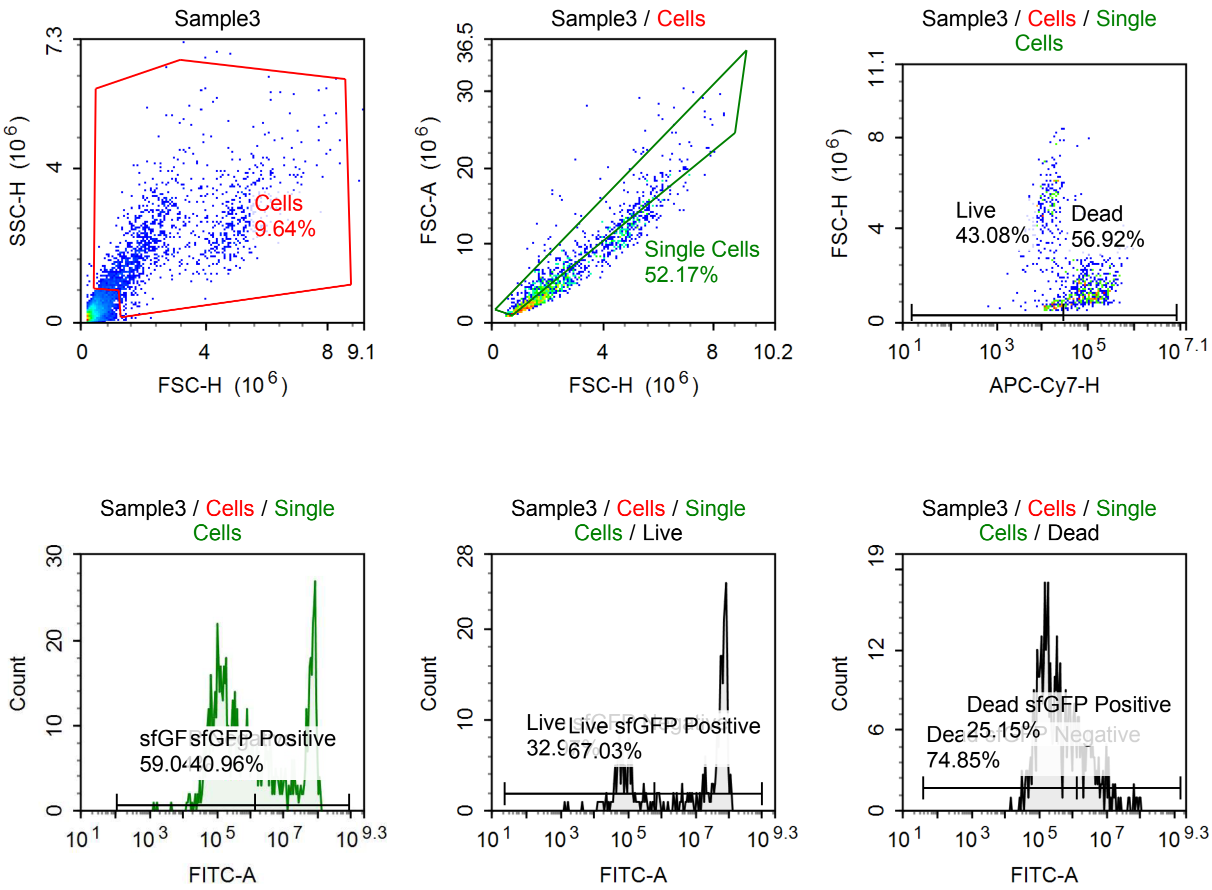

Sample Statistics of Sample3

| Gate | Count | % Parent | X | Y | Median X | Median Y |
|---|---|---|---|---|---|---|
| All | 16,940 | | | | | |
| Cells | 1,633 | 9.64 % | FSC-H | SSC-H | 2,022,224 | 1,889,203 |
| Single Cells | 852 | 52.17 % | FSC-H | FSC-A | 1,681,472 | 4,480,792 |
| Live | 367 | 43.08 % | APC-Cy7-H | FSC-H | 15,015 | 4,030,232 |
| Live sfGFP Negative | 121 | 32.97 % | FITC-A | Count | 72,015 | |
| Live sfGFP Positive | 246 | 67.03 % | FITC-A | Count | 61,735,516 | |
| Dead | 485 | 56.92 % | APC-Cy7-H | FSC-H | 108,886 | 1,401,461 |
| Dead sfGFP Negative | 363 | 74.85 % | FITC-A | Count | 192,430 | |
| Dead sfGFP Positive | 122 | 25.15 % | FITC-A | Count | 3,586,479 | |
| sfGFP Negative | 503 | 59.04 % | FITC-A | Count | 163,049 | |
| sfGFP Positive | 349 | 40.96 % | FITC-A | Count | 46,966,016 | |

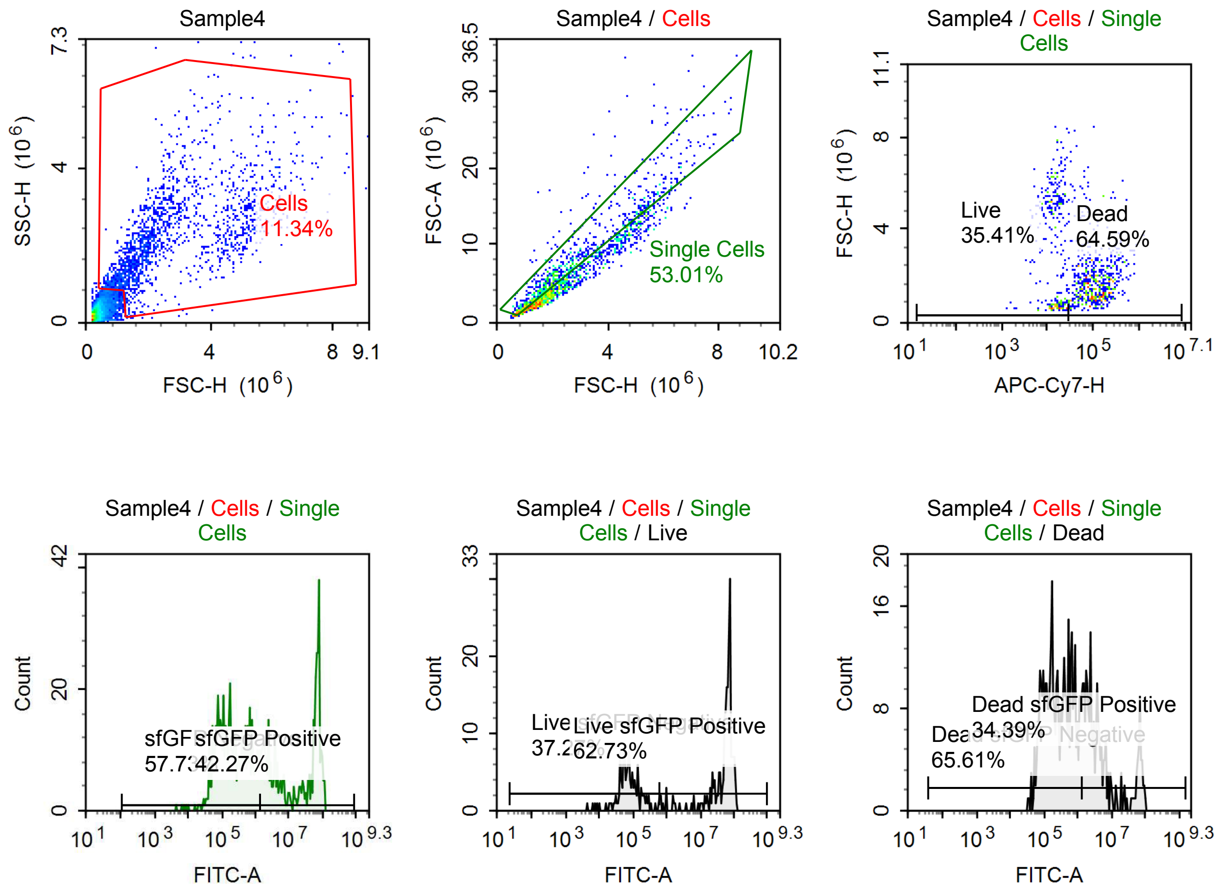

Sample Statistics of Sample4

| Gate | Count | % Parent | X | Y | Median X | Median Y |
| --- | --- | --- | --- | --- | --- | --- |
| All | 15,498 | | | | | |
| Cells | 1,758 | 11.34 % | FSC-H | SSC-H | 2,080,779 | 2,055,052 |
| Single Cells | 932 | 53.01 % | FSC-H | FSC-A | 1,857,845 | 5,228,513 |
| Live | 330 | 35.41 % | APC-Cy7-H | FSC-H | 14,912 | 4,162,818 |
| Live sfGFP Negative | 123 | 37.27 % | FITC-A | Count | 73,747 | |
| Live sfGFP Positive | 207 | 62.73 % | FITC-A | Count | 63,200,124 | |
| Dead | 602 | 64.59 % | APC-Cy7-H | FSC-H | 107,608 | 1,706,401 |
| Dead sfGFP Negative | 395 | 65.61 % | FITC-A | Count | 258,148 | |
| Dead sfGFP Positive | 207 | 34.39 % | FITC-A | Count | 3,726,803 | |
| sfGFP Negative | 538 | 57.73 % | FITC-A | Count | 185,549 | |
| sfGFP Positive | 394 | 42.27 % | FITC-A | Count | 36,854,852 | |

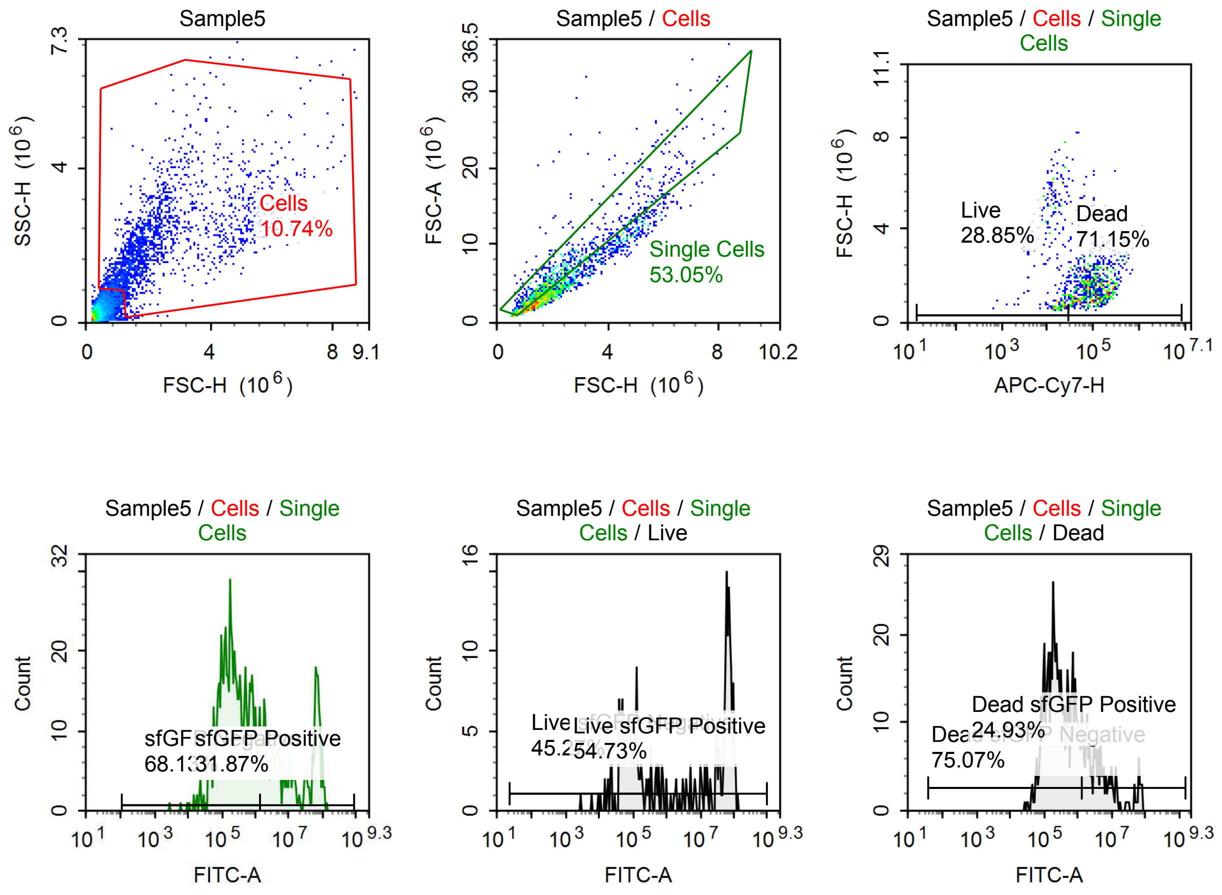

Sample Statistics of Sample5

| Gate | Count | % Parent | X | Y | Median X | Median Y |
|---|---|---|---|---|---|---|
| All | 18,005 | | | | | |
| Cells | 1,934 | 10.74 % | FSC-H | SSC-H | 1,837,613 | 1,956,595 |
| Single Cells | 1,026 | 53.05 % | FSC-H | FSC-A | 1,597,139 | 4,251,410 |
| Live | 296 | 28.85 % | APC-Cy7-H | FSC-H | 15,248 | 2,720,696 |
| Live sfGFP Negative | 134 | 45.27 % | FITC-A | Count | 79,651 | |
| Live sfGFP Positive | 162 | 54.73 % | FITC-A | Count | 58,268,264 | |
| Dead | 730 | 71.15 % | APC-Cy7-H | FSC-H | 102,181 | 1,548,128 |
| Dead sfGFP Negative | 548 | 75.07 % | FITC-A | Count | 244,527 | |
| Dead sfGFP Positive | 182 | 24.93 % | FITC-A | Count | 4,138,683 | |
| sfGFP Negative | 699 | 68.13 % | FITC-A | Count | 208,582 | |
| sfGFP Positive | 327 | 31.87 % | FITC-A | Count | 12,897,050 | |

#### p059 mRNA [10 ng]

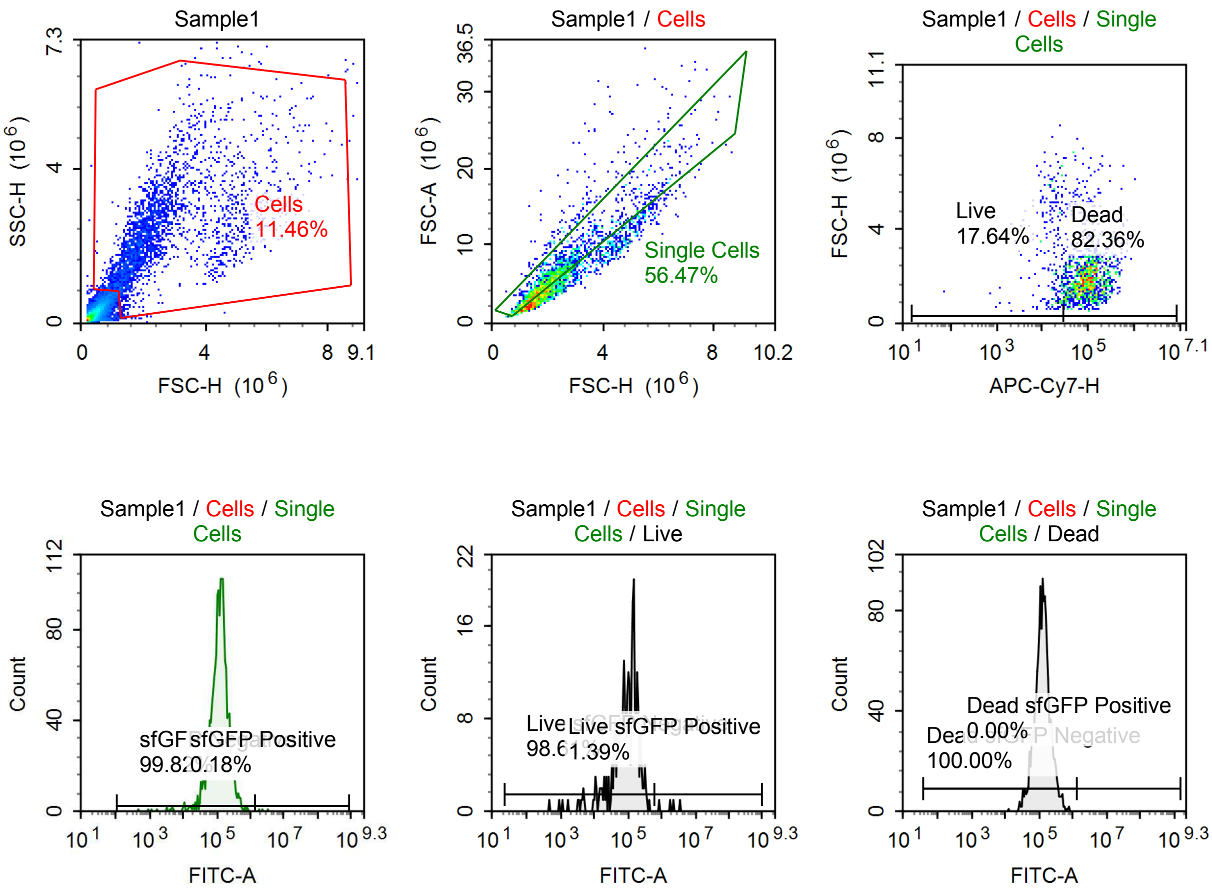

Sample Statistics of Sample1

| Gate | Count | % Parent | X | Y | Median X | Median Y |
|---|---|---|---|---|---|---|
| All | 25,143 | | | | | |
| Cells | 2,881 | 11.46 % | FSC-H | SSC-H | 2,096,112 | 2,168,445 |
| Single Cells | 1,627 | 56.47 % | FSC-H | FSC-A | 1,928,296 | 5,456,163 |
| Live | 287 | 17.64 % | APC-Cy7-H | FSC-H | 18,686 | 2,447,127 |
| Live sfGFP Negative | 283 | 98.61 % | FITC-A | Count | 98,624 | |
| Live sfGFP Positive | 4 | 1.39 % | FITC-A | Count | 1,994,782 | |
| Dead | 1,340 | 82.36 % | APC-Cy7-H | FSC-H | 97,417 | 1,884,335 |
| Dead sfGFP Negative | 1,340 | 100.00 % | FITC-A | Count | 126,340 | |
| Dead sfGFP Positive | 0 | 0.00 % | FITC-A | Count | 0 | |
| sfGFP Negative | 1,624 | 99.82 % | FITC-A | Count | 124,258 | |
| sfGFP Positive | 3 | 0.18 % | FITC-A | Count | 2,229,131 | |

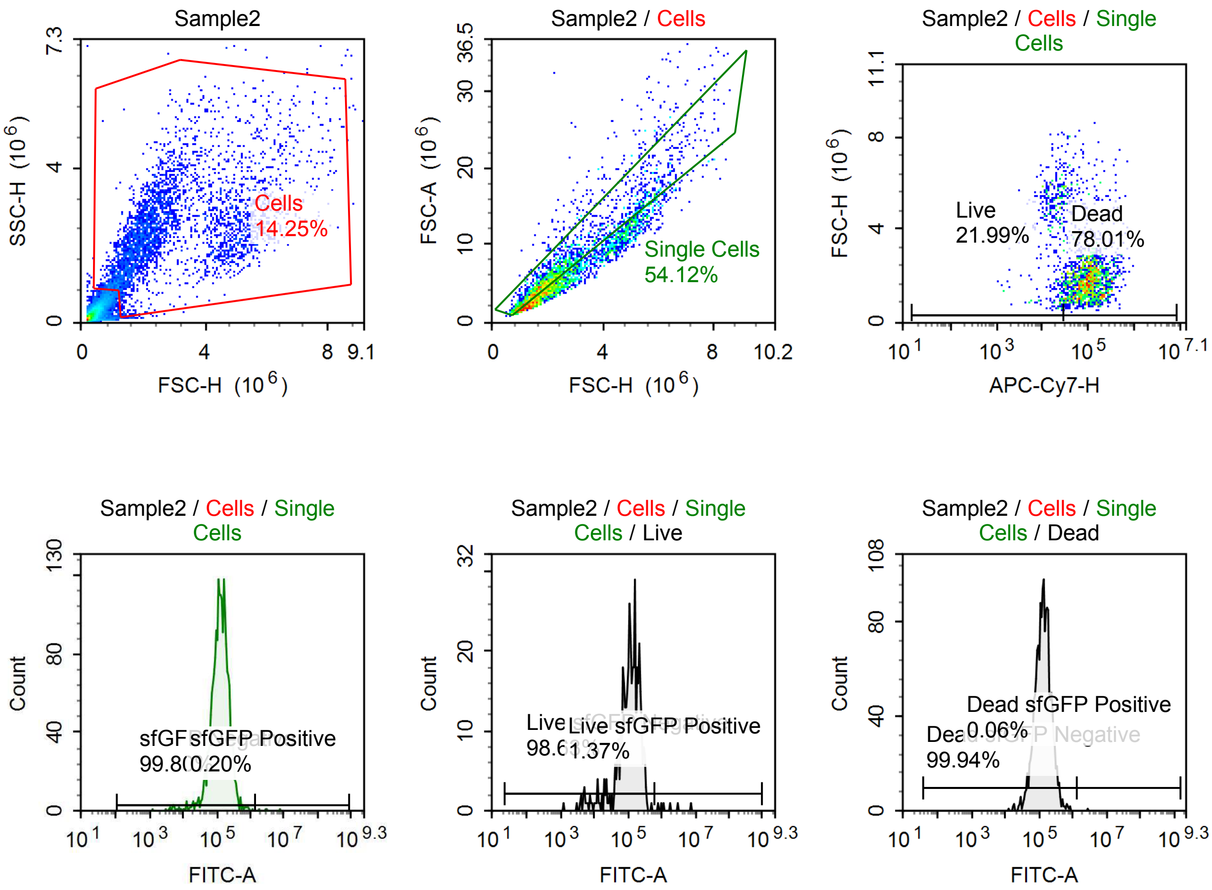

Sample Statistics of Sample2

| Gate | Count | % Parent | X | Y | Median X | Median Y |
|---|---|---|---|---|---|---|
| All | 25,837 | | | | | |
| Cells | 3,681 | 14.25 % | FSC-H | SSC-H | 2,322,599 | 2,260,966 |
| Single Cells | 1,992 | 54.12 % | FSC-H | FSC-A | 2,014,807 | 5,545,348 |
| Live | 438 | 21.99 % | APC-Cy7-H | FSC-H | 19,125 | 4,480,161 |
| Live sfGFP Negative | 432 | 98.63 % | FITC-A | Count | 116,876 | |
| Live sfGFP Positive | 6 | 1.37 % | FITC-A | Count | 1,506,904 | |
| Dead | 1,554 | 78.01 % | APC-Cy7-H | FSC-H | 101,520 | 1,882,600 |
| Dead sfGFP Negative | 1,553 | 99.94 % | FITC-A | Count | 131,538 | |
| Dead sfGFP Positive | 1 | 0.06 % | FITC-A | Count | 2,804,840 | |
| sfGFP Negative | 1,988 | 99.80 % | FITC-A | Count | 128,274 | |
| sfGFP Positive | 4 | 0.20 % | FITC-A | Count | 2,972,818 | |

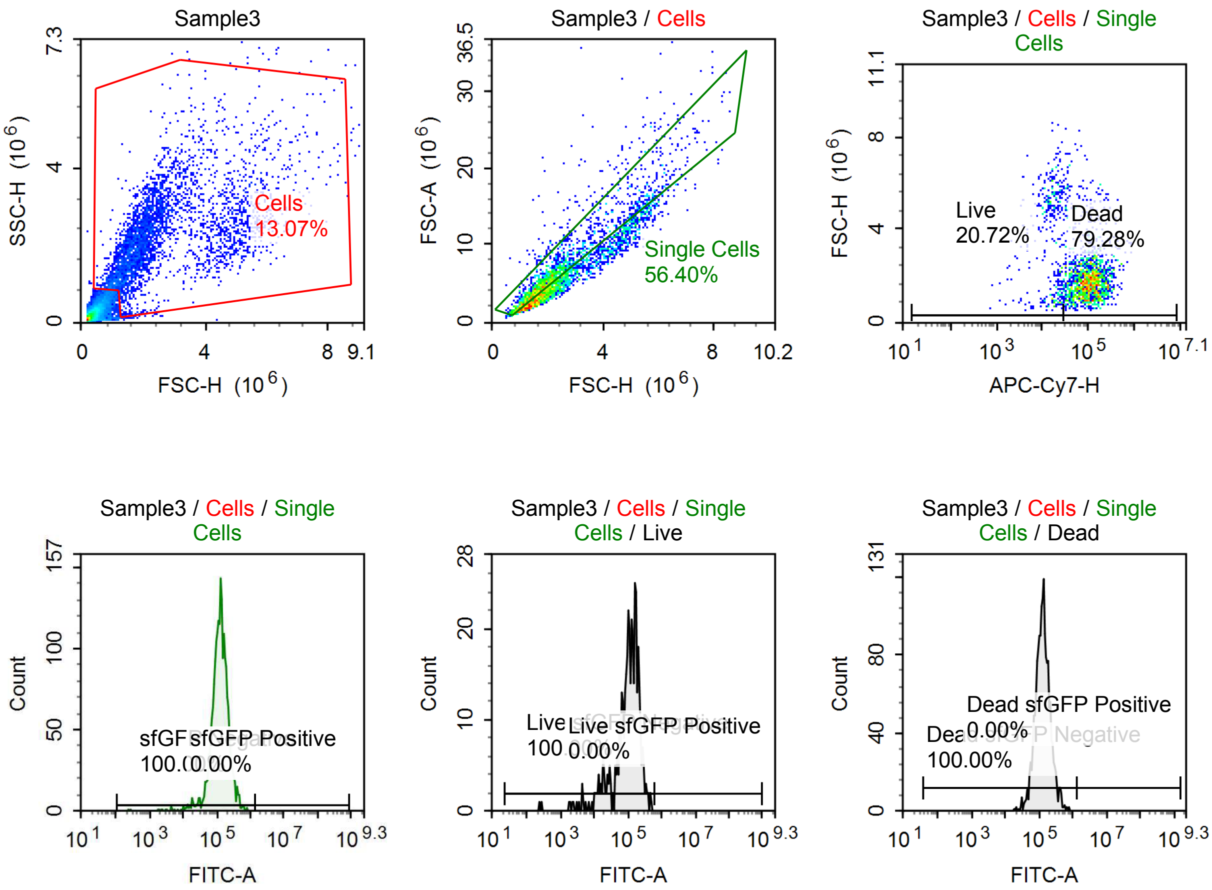

Sample Statistics of Sample3

| Gate | Count | % Parent | X | Y | Median X | Median Y |
|---|---|---|---|---|---|---|
| All | 27,561 | | | | | |
| Cells | 3,603 | 13.07 % | FSC-H | SSC-H | 2,059,658 | 2,132,449 |
| Single Cells | 2,032 | 56.40 % | FSC-H | FSC-A | 1,857,589 | 5,049,247 |
| Live | 421 | 20.72 % | APC-Cy7-H | FSC-H | 17,327 | 3,736,605 |
| Live sfGFP Negative | 421 | 100.00 % | FITC-A | Count | 110,823 | |
| Live sfGFP Positive | 0 | 0.00 % | FITC-A | Count | 0 | |
| Dead | 1,611 | 79.28 % | APC-Cy7-H | FSC-H | 102,056 | 1,770,918 |
| Dead sfGFP Negative | 1,611 | 100.00 % | FITC-A | Count | 128,160 | |
| Dead sfGFP Positive | 0 | 0.00 % | FITC-A | Count | 0 | |
| sfGFP Negative | 2,032 | 100.00 % | FITC-A | Count | 126,081 | |
| sfGFP Positive | 0 | 0.00 % | FITC-A | Count | 0 | |

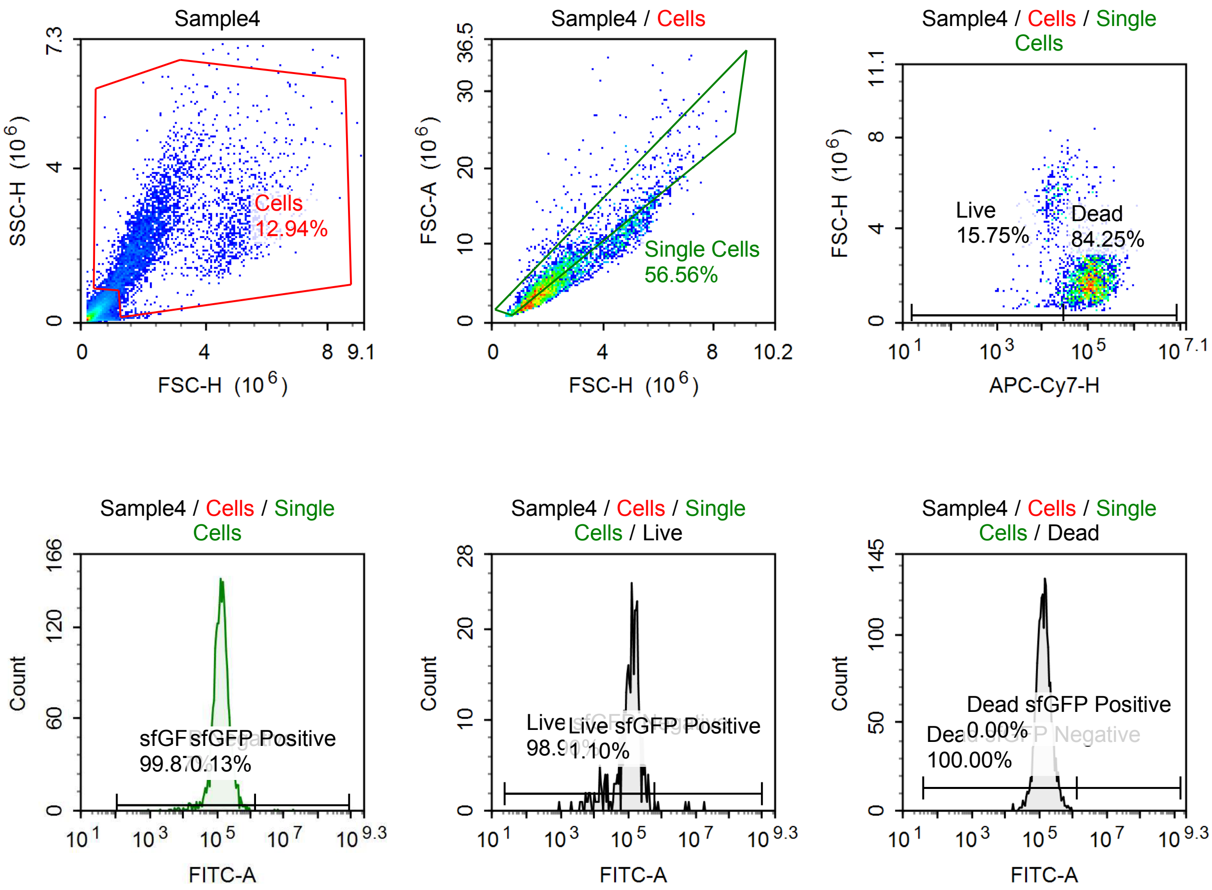

Sample Statistics of Sample4

| Gate | Count | % Parent | X | Y | Median X | Median Y |
|---|---|---|---|---|---|---|
| All | 31,499 | | | | | |
| Cells | 4,075 | 12.94 % | FSC-H | SSC-H | 2,101,251 | 2,093,230 |
| Single Cells | 2,305 | 56.56 % | FSC-H | FSC-A | 1,893,686 | 5,218,020 |
| Live | 363 | 15.75 % | APC-Cy7-H | FSC-H | 15,644 | 4,462,457 |
| Live sfGFP Negative | 359 | 98.90 % | FITC-A | Count | 129,746 | |
| Live sfGFP Positive | 4 | 1.10 % | FITC-A | Count | 5,879,066 | |
| Dead | 1,942 | 84.25 % | APC-Cy7-H | FSC-H | 111,226 | 1,815,442 |
| Dead sfGFP Negative | 1,942 | 100.00 % | FITC-A | Count | 135,404 | |
| Dead sfGFP Positive | 0 | 0.00 % | FITC-A | Count | 0 | |
| sfGFP Negative | 2,302 | 99.87 % | FITC-A | Count | 133,823 | |
| sfGFP Positive | 3 | 0.13 % | FITC-A | Count | 6,679,496 | |

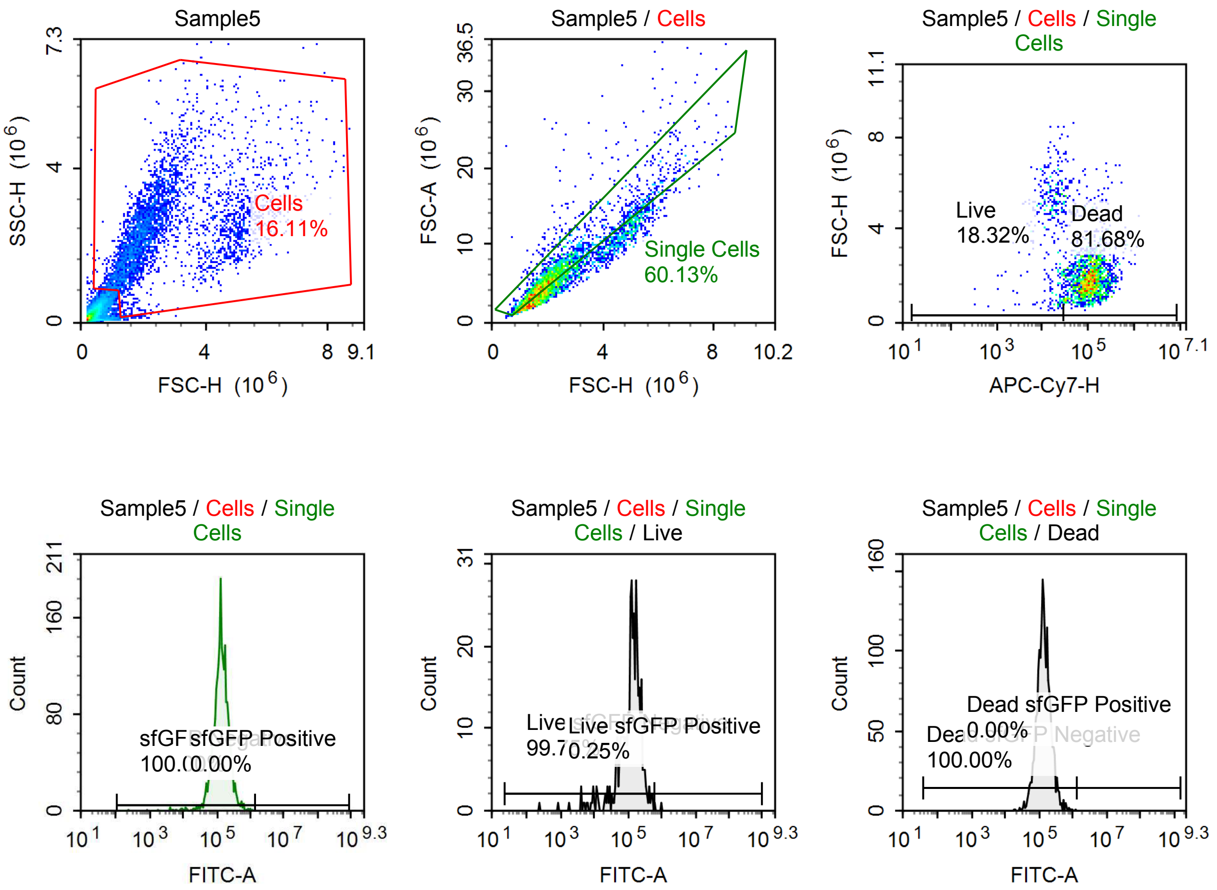

Sample Statistics of Sample5

| Gate | Count | % Parent | X | Y | Median X | Median Y |
|---|---|---|---|---|---|---|
| All | 22,652 | | | | | |
| Cells | 3,649 | 16.11 % | FSC-H | SSC-H | 2,160,568 | 2,314,215 |
| Single Cells | 2,194 | 60.13 % | FSC-H | FSC-A | 1,983,859 | 5,504,986 |
| Live | 402 | 18.32 % | APC-Cy7-H | FSC-H | 17,418 | 4,637,144 |
| Live sfGFP Negative | 401 | 99.75 % | FITC-A | Count | 133,325 | |
| Live sfGFP Positive | 1 | 0.25 % | FITC-A | Count | 977,859 | |
| Dead | 1,792 | 81.68 % | APC-Cy7-H | FSC-H | 107,650 | 1,879,722 |
| Dead sfGFP Negative | 1,792 | 100.00 % | FITC-A | Count | 134,660 | |
| Dead sfGFP Positive | 0 | 0.00 % | FITC-A | Count | 0 | |
| sfGFP Negative | 2,194 | 100.00 % | FITC-A | Count | 134,429 | |
| sfGFP Positive | 0 | 0.00 % | FITC-A | Count | 0 | |

#### Commercial saRNA

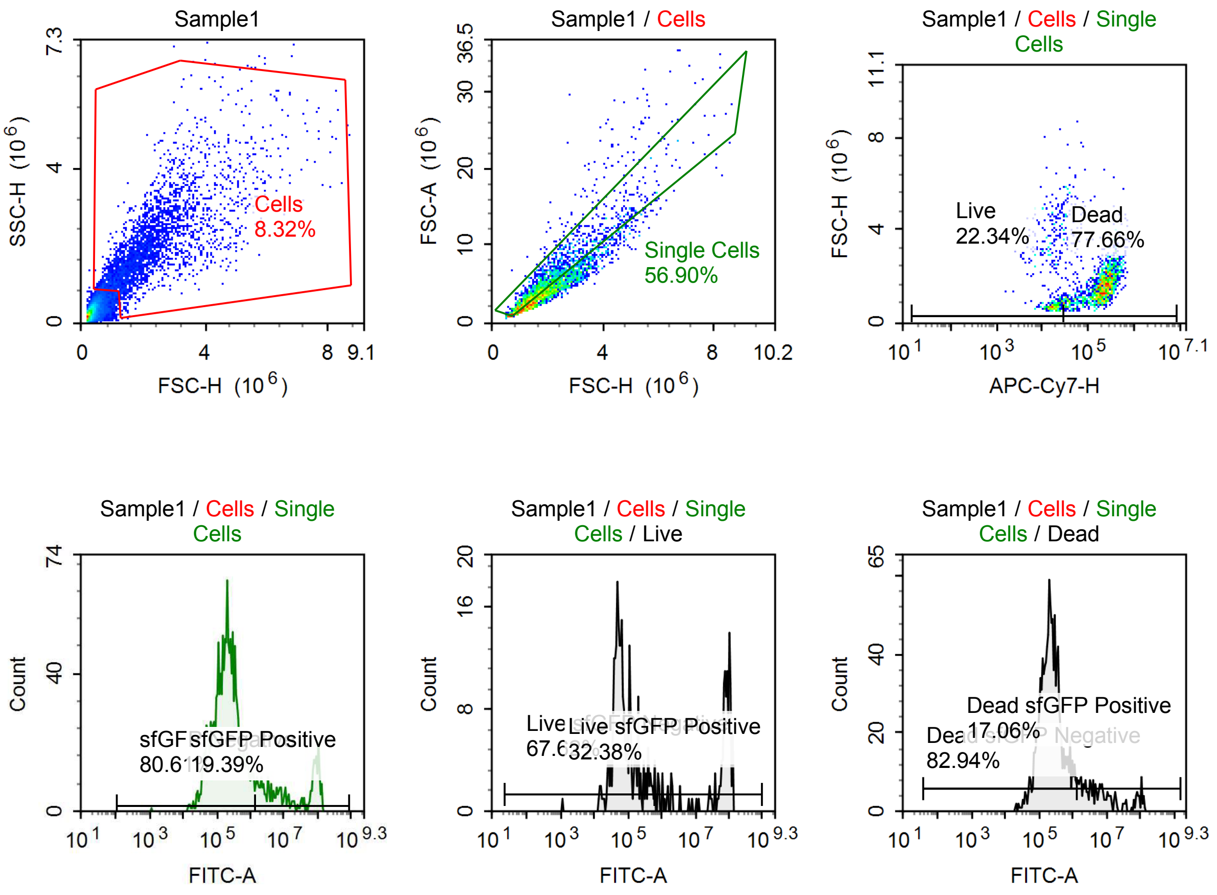

Sample Statistics of Sample1

| Gate | Count | % Parent | X | Y | Median X | Median Y |
|---|---|---|---|---|---|---|
| All | 36,516 | | | | | |
| Cells | 3,037 | 8.32 % | FSC-H | SSC-H | 1,910,343 | 1,763,726 |
| Single Cells | 1,728 | 56.90 % | FSC-H | FSC-A | 1,609,347 | 4,234,626 |
| Live | 386 | 22.34 % | APC-Cy7-H | FSC-H | 16,707 | 1,334,573 |
| Live sfGFP Negative | 261 | 67.62 % | FITC-A | Count | 64,621 | |
| Live sfGFP Positive | 125 | 32.38 % | FITC-A | Count | 78,437,144 | |
| Dead | 1,342 | 77.66 % | APC-Cy7-H | FSC-H | 209,417 | 1,615,604 |
| Dead sfGFP Negative | 1,113 | 82.94 % | FITC-A | Count | 214,372 | |
| Dead sfGFP Positive | 229 | 17.06 % | FITC-A | Count | 5,823,921 | |
| sfGFP Negative | 1,393 | 80.61 % | FITC-A | Count | 194,542 | |
| sfGFP Positive | 335 | 19.39 % | FITC-A | Count | 13,482,767 | |

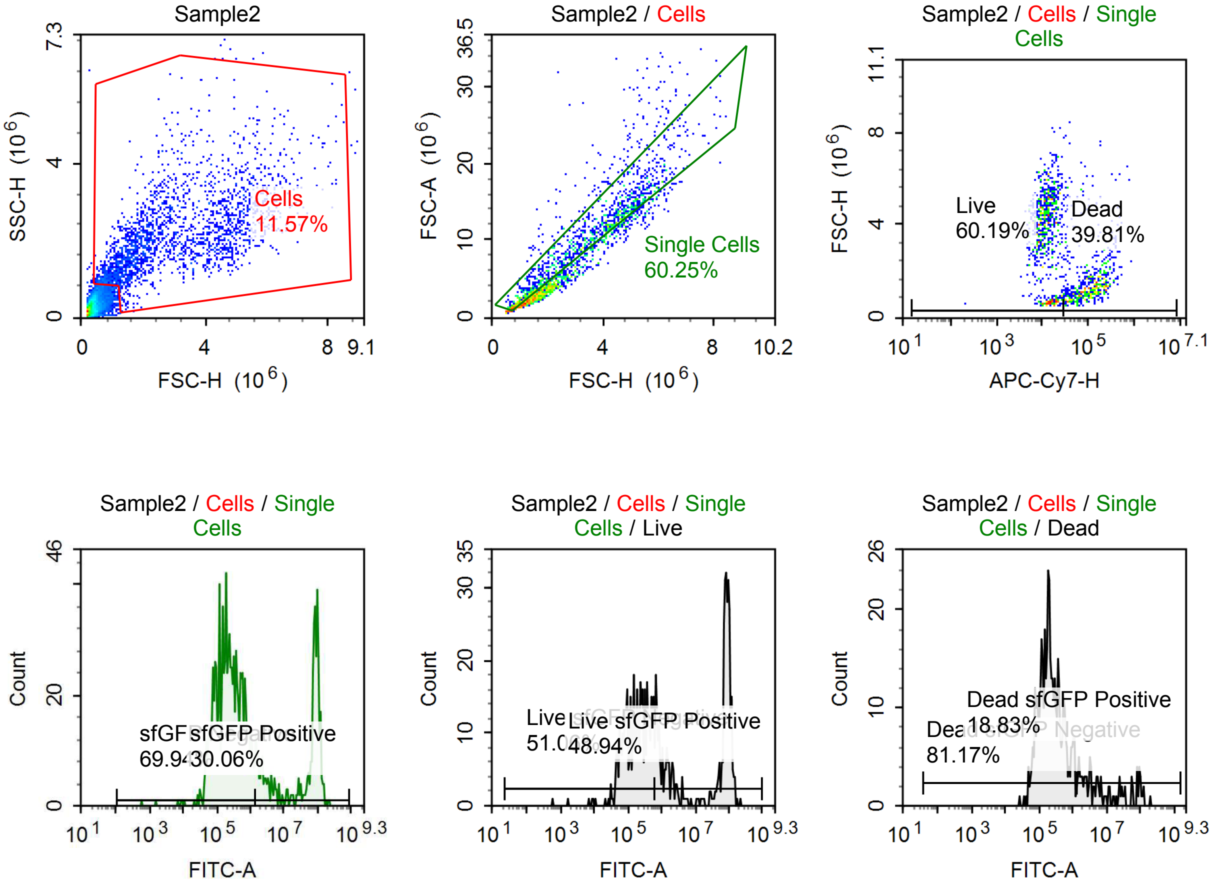

Sample Statistics of Sample2

| Gate | Count | % Parent | X | Y | Median X | Median Y |
|---|---|---|---|---|---|---|
| All | 19,143 | | | | | |
| Cells | 2,214 | 11.57 % | FSC-H | SSC-H | 2,432,589 | 1,946,668 |
| Single Cells | 1,334 | 60.25 % | FSC-H | FSC-A | 2,719,477 | 8,240,370 |
| Live | 803 | 60.19 % | APC-Cy7-H | FSC-H | 12,452 | 4,072,914 |
| Live sfGFP Negative | 410 | 51.06 % | FITC-A | Count | 189,867 | |
| Live sfGFP Positive | 393 | 48.94 % | FITC-A | Count | 72,692,064 | |
| Dead | 531 | 39.81 % | APC-Cy7-H | FSC-H | 119,792 | 1,386,080 |
| Dead sfGFP Negative | 431 | 81.17 % | FITC-A | Count | 203,060 | |
| Dead sfGFP Positive | 100 | 18.83 % | FITC-A | Count | 5,695,007 | |
| sfGFP Negative | 933 | 69.94 % | FITC-A | Count | 224,377 | |
| sfGFP Positive | 401 | 30.06 % | FITC-A | Count | 76,405,792 | |

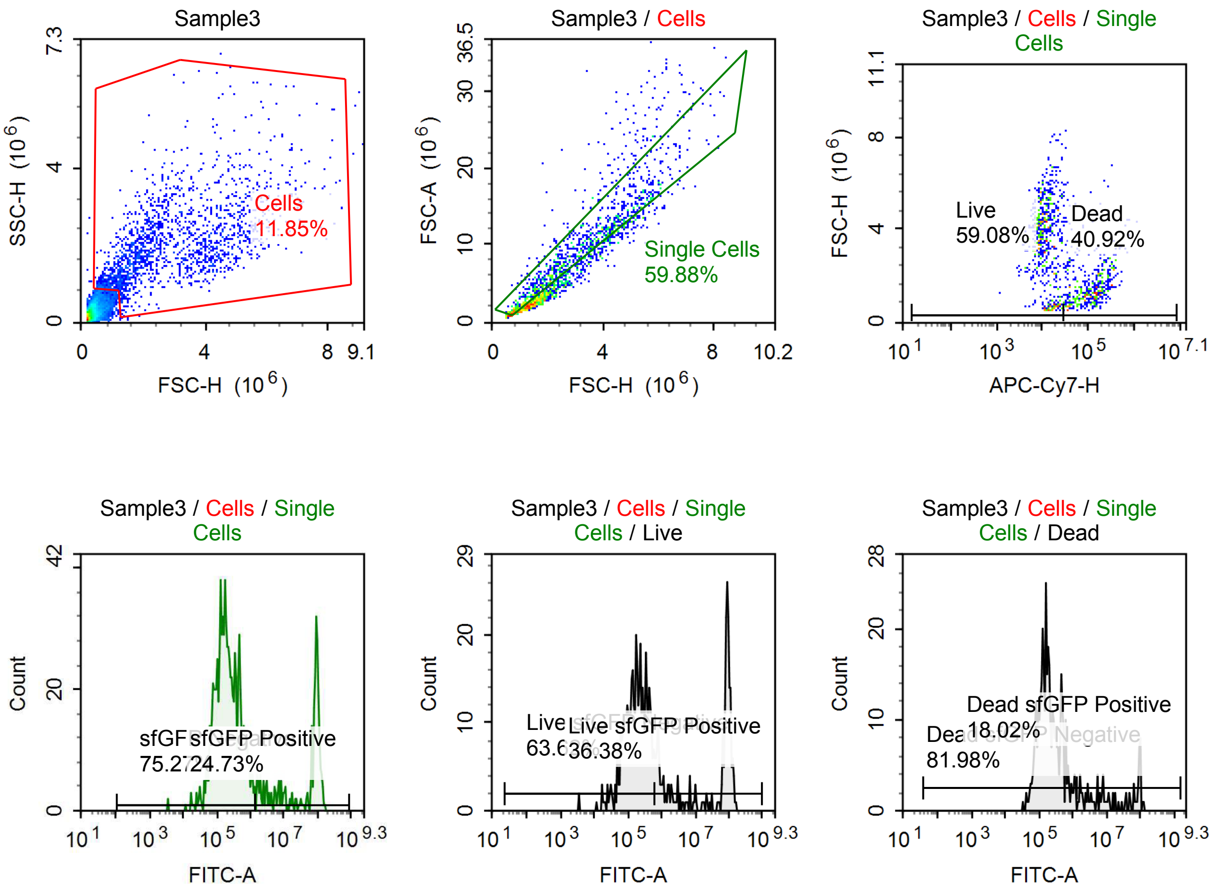

Sample Statistics of Sample3

| Gate | Count | % Parent | X | Y | Median X | Median Y |
|---|---|---|---|---|---|---|
| All | 15,676 | | | | | |
| Cells | 1,857 | 11.85 % | FSC-H | SSC-H | 2,394,862 | 1,839,912 |
| Single Cells | 1,112 | 59.88 % | FSC-H | FSC-A | 2,473,618 | 7,573,765 |
| Live | 657 | 59.08 % | APC-Cy7-H | FSC-H | 11,511 | 3,795,400 |
| Live sfGFP Negative | 418 | 63.62 % | FITC-A | Count | 178,253 | |
| Live sfGFP Positive | 239 | 36.38 % | FITC-A | Count | 77,293,856 | |
| Dead | 455 | 40.92 % | APC-Cy7-H | FSC-H | 126,251 | 1,490,440 |
| Dead sfGFP Negative | 373 | 81.98 % | FITC-A | Count | 181,346 | |
| Dead sfGFP Positive | 82 | 18.02 % | FITC-A | Count | 11,033,487 | |
| sfGFP Negative | 837 | 75.27 % | FITC-A | Count | 189,256 | |
| sfGFP Positive | 275 | 24.73 % | FITC-A | Count | 78,212,312 | |

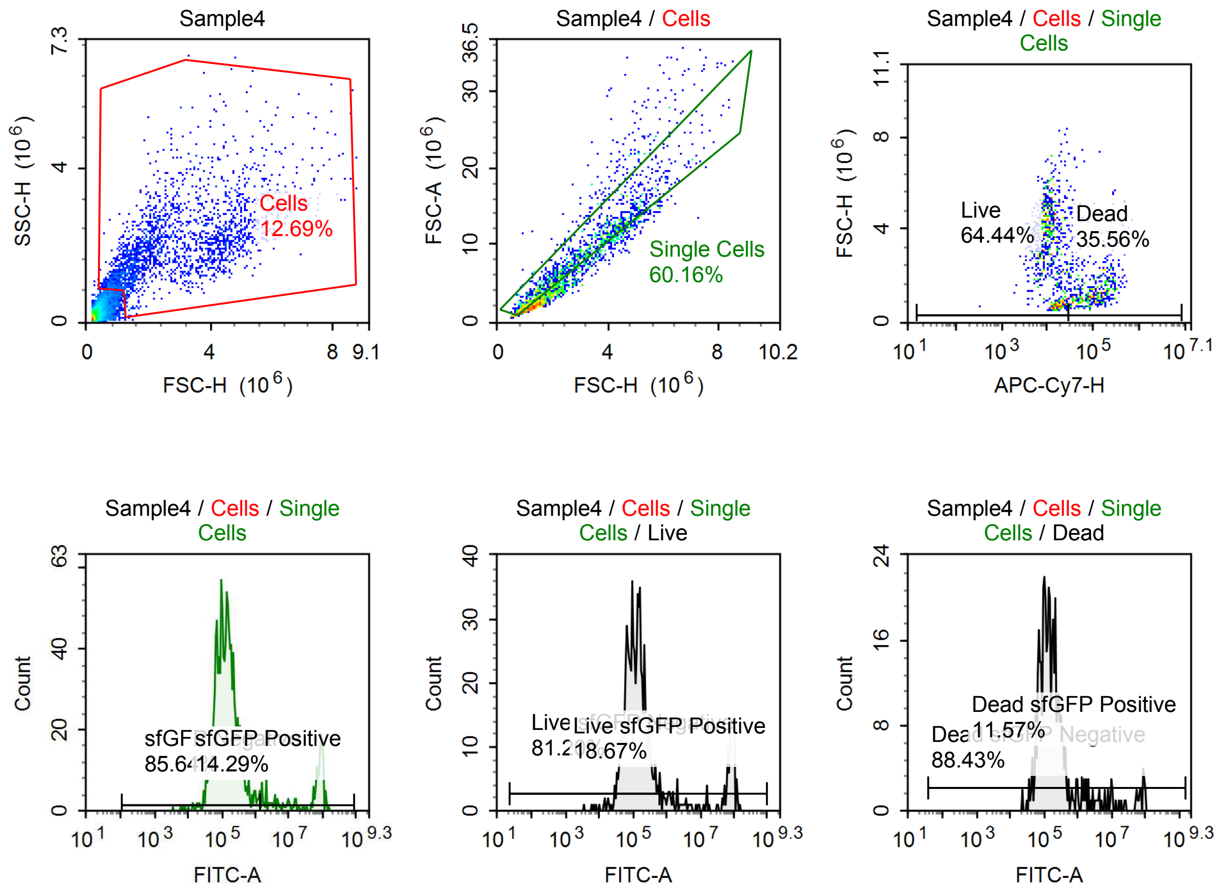

Sample Statistics of Sample4

| Gate | Count | % Parent | X | Y | Median X | Median Y |
|---|---|---|---|---|---|---|
| All | 16,873 | | | | | |
| Cells | 2,141 | 12.69 % | FSC-H | SSC-H | 2,345,896 | 1,879,836 |
| Single Cells | 1,288 | 60.16 % | FSC-H | FSC-A | 2,391,073 | 7,072,637 |
| Live | 830 | 64.44 % | APC-Cy7-H | FSC-H | 11,256 | 3,497,091 |
| Live sfGFP Negative | 674 | 81.20 % | FITC-A | Count | 118,154 | |
| Live sfGFP Positive | 155 | 18.67 % | FITC-A | Count | 69,121,840 | |
| Dead | 458 | 35.56 % | APC-Cy7-H | FSC-H | 105,515 | 1,408,592 |
| Dead sfGFP Negative | 405 | 88.43 % | FITC-A | Count | 137,784 | |
| Dead sfGFP Positive | 53 | 11.57 % | FITC-A | Count | 9,787,914 | |
| sfGFP Negative | 1,103 | 85.64 % | FITC-A | Count | 129,565 | |
| sfGFP Positive | 184 | 14.29 % | FITC-A | Count | 68,500,656 | |

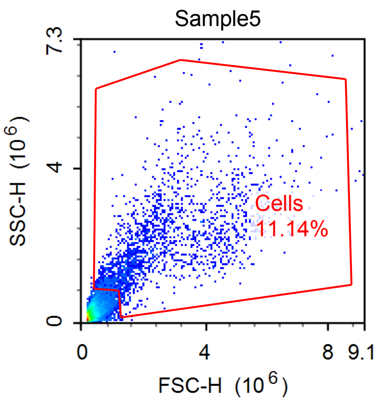


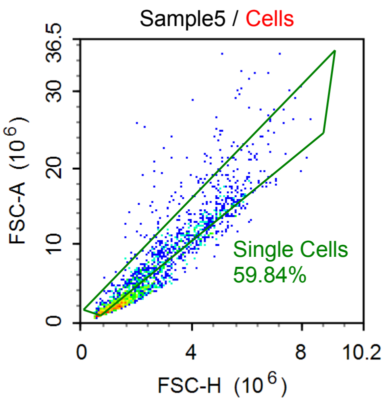


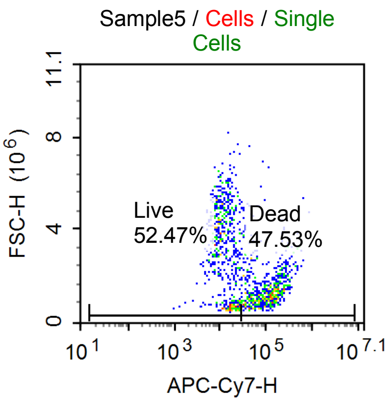


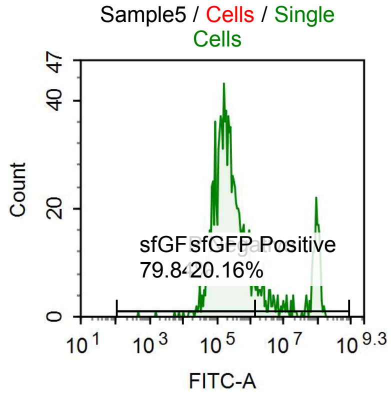


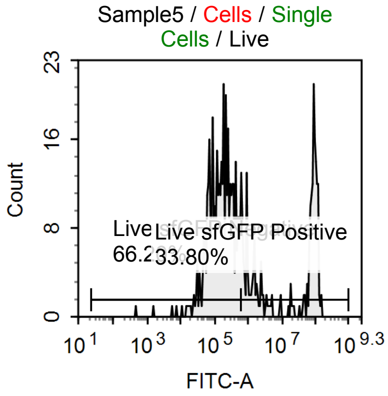


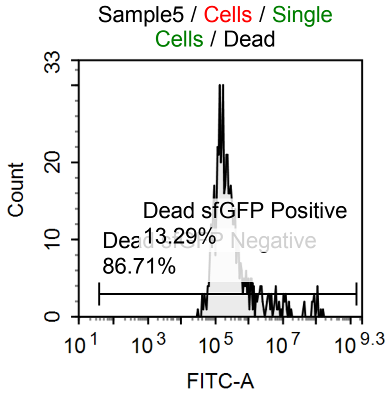

Sample Statistics of Sample5

| Gate | Count | % Parent | X | Y | Median X | Median Y |
|---|---|---|---|---|---|---|
| All | 18,526 | | | | | |
| Cells | 2,064 | 11.14 % | FSC-H | SSC-H | 2,032,641 | 1,814,139 |
| Single Cells | 1,235 | 59.84 % | FSC-H | FSC-A | 1,950,250 | 5,549,091 |
| Live | 648 | 52.47 % | APC-Cy7-H | FSC-H | 12,362 | 3,482,947 |
| Live sfGFP Negative | 429 | 66.20 % | FITC-A | Count | 166,434 | |
| Live sfGFP Positive | 219 | 33.80 % | FITC-A | Count | 69,968,224 | |
| Dead | 587 | 47.53 % | APC-Cy7-H | FSC-H | 113,554 | 1,335,777 |
| Dead sfGFP Negative | 509 | 86.71 % | FITC-A | Count | 187,653 | |
| Dead sfGFP Positive | 78 | 13.29 % | FITC-A | Count | 5,828,750 | |
| sfGFP Negative | 986 | 79.84 % | FITC-A | Count | 189,144 | |
| sfGFP Positive | 249 | 20.16 % | FITC-A | Count | 69,516,112 | |

#### Mock Transfection

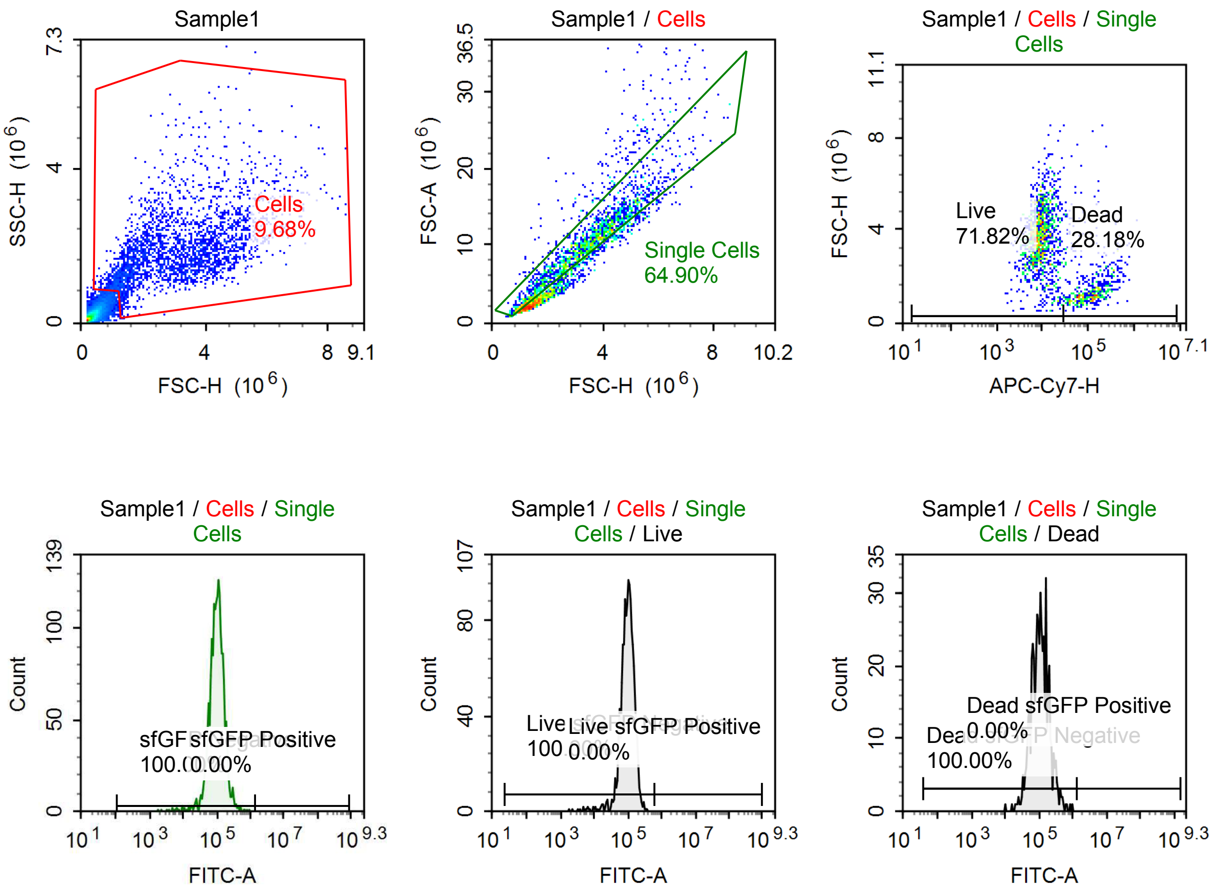

Sample Statistics of Sample1

| Gate | Count | % Parent | X | Y | Median X | Median Y |
|---|---|---|---|---|---|---|
| All | 28,817 | | | | | |
| Cells | 2,789 | 9.68 % | FSC-H | SSC-H | 2,888,119 | 1,761,074 |
| Single Cells | 1,810 | 64.90 % | FSC-H | FSC-A | 3,206,651 | 9,585,478 |
| Live | 1,300 | 71.82 % | APC-Cy7-H | FSC-H | 8,924 | 3,666,945 |
| Live sfGFP Negative | 1,300 | 100.00 % | FITC-A | Count | 100,377 | |
| Live sfGFP Positive | 0 | 0.00 % | FITC-A | Count | 0 | |
| Dead | 510 | 28.18 % | APC-Cy7-H | FSC-H | 120,888 | 1,394,937 |
| Dead sfGFP Negative | 510 | 100.00 % | FITC-A | Count | 108,085 | |
| Dead sfGFP Positive | 0 | 0.00 % | FITC-A | Count | 0 | |
| sfGFP Negative | 1,810 | 100.00 % | FITC-A | Count | 101,517 | |
| sfGFP Positive | 0 | 0.00 % | FITC-A | Count | 0 | |

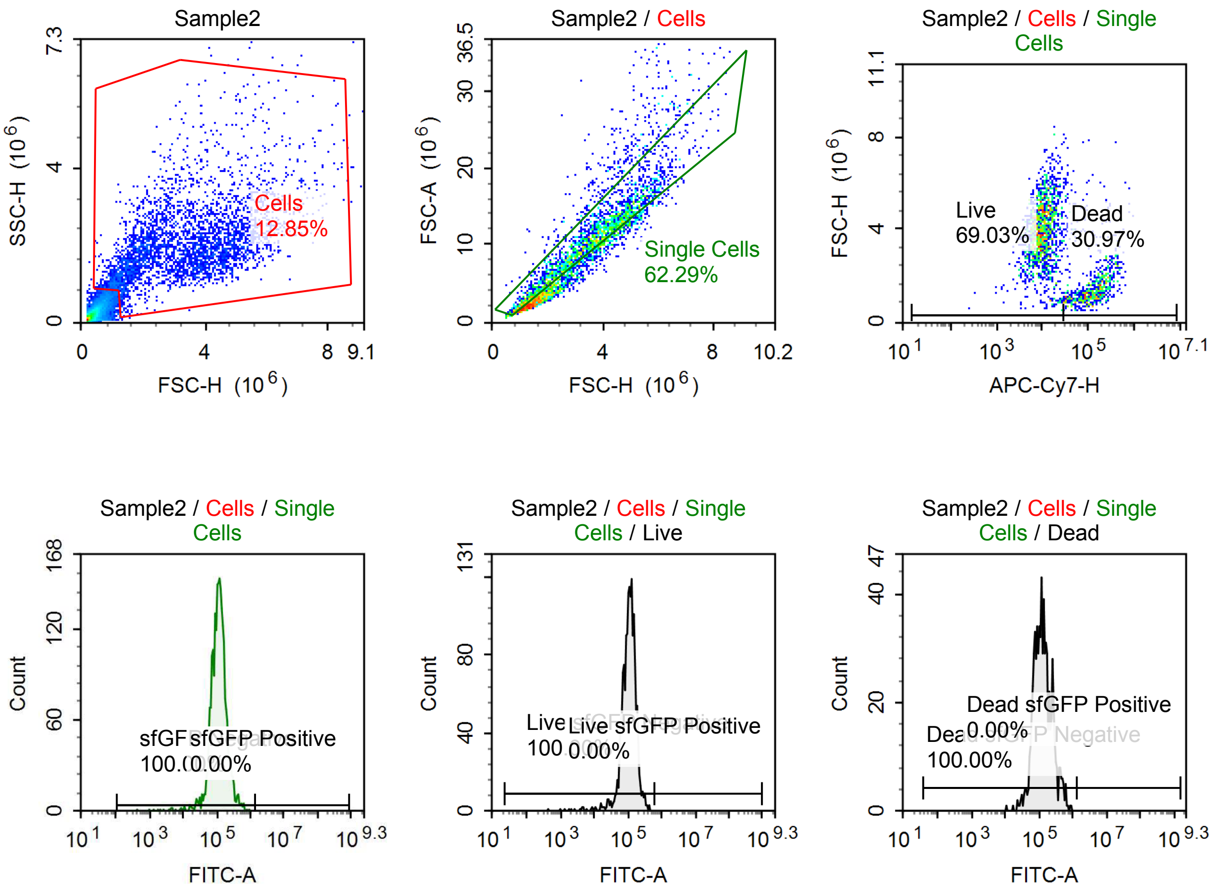

Sample Statistics of Sample2

| Gate | Count | % Parent | X | Y | Median X | Median Y |
|---|---|---|---|---|---|---|
| All | 28,395 | | | | | |
| Cells | 3,649 | 12.85 % | FSC-H | SSC-H | 3,174,841 | 1,883,247 |
| Single Cells | 2,273 | 62.29 % | FSC-H | FSC-A | 3,356,414 | 10,133,886 |
| Live | 1,569 | 69.03 % | APC-Cy7-H | FSC-H | 9,672 | 3,873,604 |
| Live sfGFP Negative | 1,569 | 100.00 % | FITC-A | Count | 110,725 | |
| Live sfGFP Positive | 0 | 0.00 % | FITC-A | Count | 0 | |
| Dead | 704 | 30.97 % | APC-Cy7-H | FSC-H | 138,124 | 1,502,186 |
| Dead sfGFP Negative | 704 | 100.00 % | FITC-A | Count | 122,364 | |
| Dead sfGFP Positive | 0 | 0.00 % | FITC-A | Count | 0 | |
| sfGFP Negative | 2,273 | 100.00 % | FITC-A | Count | 114,142 | |
| sfGFP Positive | 0 | 0.00 % | FITC-A | Count | 0 | |

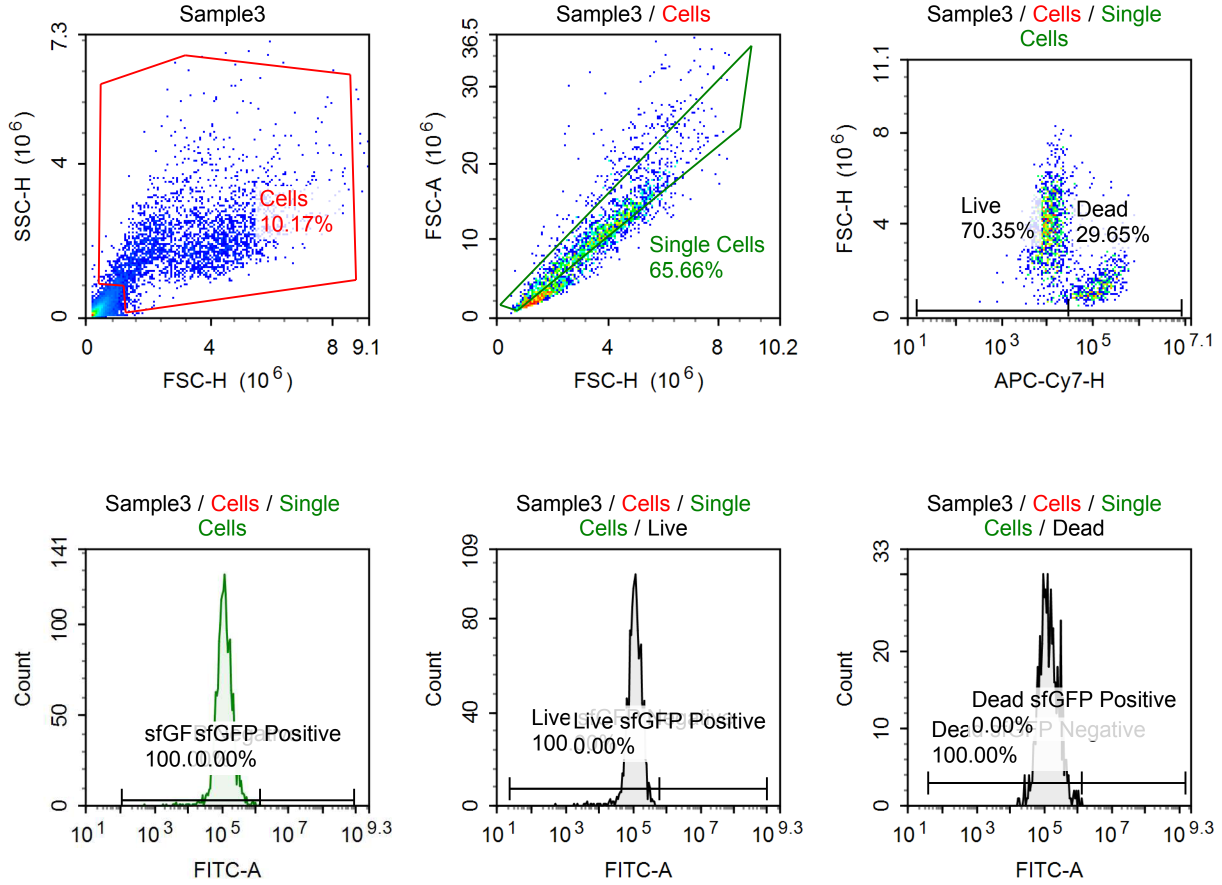

Sample Statistics of Sample3

| Gate | Count | % Parent | X | Y | Median X | Median Y |
|---|---|---|---|---|---|---|
| All | 28,680 | | | | | |
| Cells | 2,918 | 10.17 % | FSC-H | SSC-H | 3,144,628 | 1,873,073 |
| Single Cells | 1,916 | 65.66 % | FSC-H | FSC-A | 3,347,840 | 9,983,034 |
| Live | 1,348 | 70.35 % | APC-Cy7-H | FSC-H | 10,110 | 3,877,887 |
| Live sfGFP Negative | 1,348 | 100.00 % | FITC-A | Count | 114,368 | |
| Live sfGFP Positive | 0 | 0.00 % | FITC-A | Count | 0 | |
| Dead | 568 | 29.65 % | APC-Cy7-H | FSC-H | 130,787 | 1,494,819 |
| Dead sfGFP Negative | 568 | 100.00 % | FITC-A | Count | 130,632 | |
| Dead sfGFP Positive | 0 | 0.00 % | FITC-A | Count | 0 | |
| sfGFP Negative | 1,916 | 100.00 % | FITC-A | Count | 118,610 | |
| sfGFP Positive | 0 | 0.00 % | FITC-A | Count | 0 | |

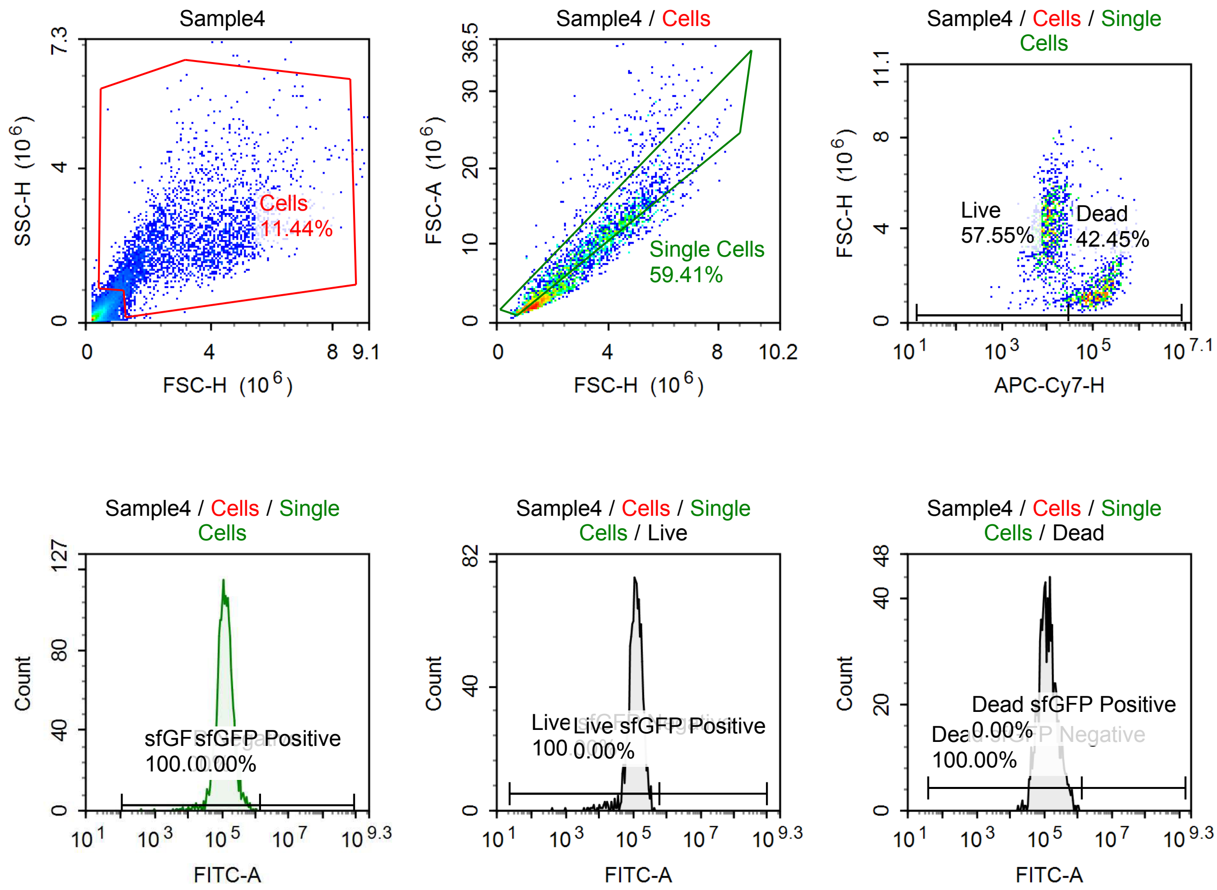

Sample Statistics of Sample4

| Gate | Count | % Parent | X | Y | Median X | Median Y |
|---|---|---|---|---|---|---|
| All | 27,183 | | | | | |
| Cells | 3,109 | 11.44 % | FSC-H | SSC-H | 2,668,673 | 1,902,410 |
| Single Cells | 1,847 | 59.41 % | FSC-H | FSC-A | 2,898,123 | 8,582,055 |
| Live | 1,063 | 57.55 % | APC-Cy7-H | FSC-H | 10,680 | 3,986,770 |
| Live sfGFP Negative | 1,063 | 100.00 % | FITC-A | Count | 121,517 | |
| Live sfGFP Positive | 0 | 0.00 % | FITC-A | Count | 0 | |
| Dead | 784 | 42.45 % | APC-Cy7-H | FSC-H | 130,715 | 1,468,246 |
| Dead sfGFP Negative | 784 | 100.00 % | FITC-A | Count | 124,004 | |
| Dead sfGFP Positive | 0 | 0.00 % | FITC-A | Count | 0 | |
| sfGFP Negative | 1,847 | 100.00 % | FITC-A | Count | 121,897 | |
| sfGFP Positive | 0 | 0.00 % | FITC-A | Count | 0 | |

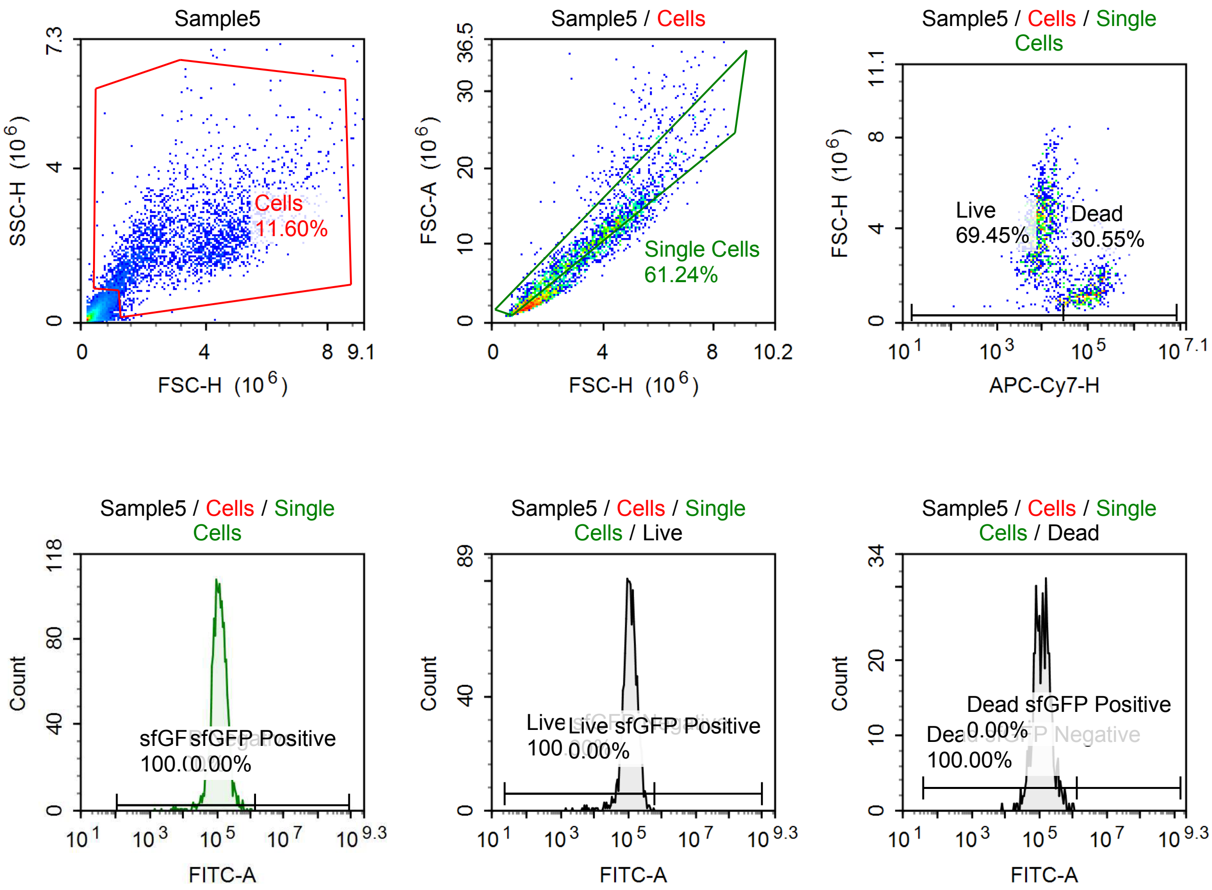

Sample Statistics of Sample5

| Gate | Count | % Parent | X | Y | Median X | Median Y |
|---|---|---|---|---|---|---|
| All | 23,729 | | | | | |
| Cells | 2,753 | 11.60 % | FSC-H | SSC-H | 3,020,252 | 1,973,151 |
| Single Cells | 1,686 | 61.24 % | FSC-H | FSC-A | 3,249,444 | 9,571,130 |
| Live | 1,171 | 69.45 % | APC-Cy7-H | FSC-H | 9,041 | 4,046,325 |
| Live sfGFP Negative | 1,171 | 100.00 % | FITC-A | Count | 116,278 | |
| Live sfGFP Positive | 0 | 0.00 % | FITC-A | Count | 0 | |
| Dead | 515 | 30.55 % | APC-Cy7-H | FSC-H | 110,057 | 1,381,037 |
| Dead sfGFP Negative | 515 | 100.00 % | FITC-A | Count | 124,019 | |
| Dead sfGFP Positive | 0 | 0.00 % | FITC-A | Count | 0 | |
| sfGFP Negative | 1,686 | 100.00 % | FITC-A | Count | 117,541 | |
| sfGFP Positive | 0 | 0.00 % | FITC-A | Count | 0 | |

### Report of 260420_1148

File Name: 260420_1148 Run Time: 4/20/2026 12:00 PM

Cytometer: NovoCyte Quanteon 621190610657 Software: NovoExpress 1.6.1

#### Specimen1

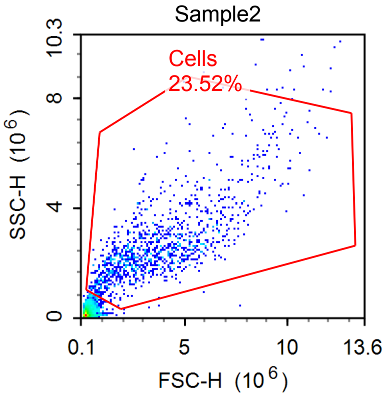


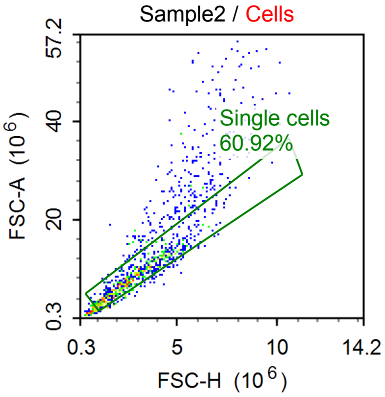


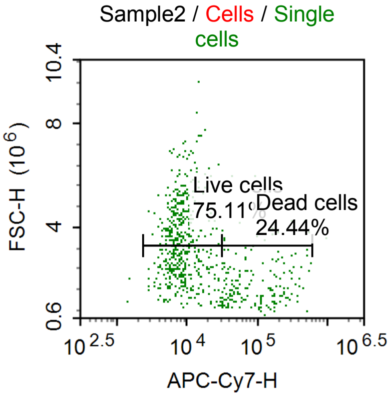


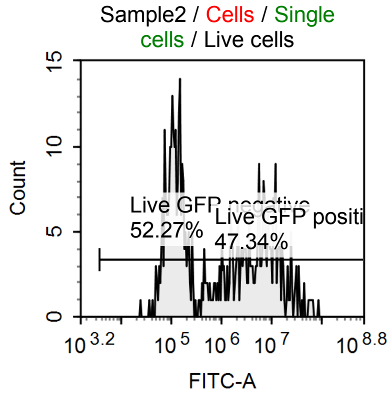


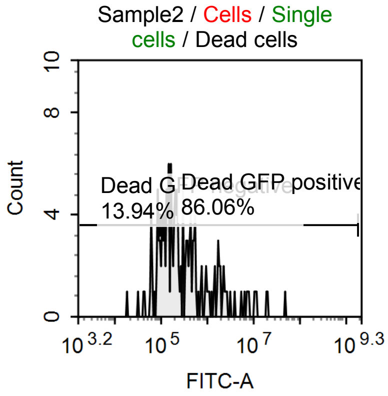


Sample Statistics of Sample2

| Gate | Count | % Parent | X | Y | Median X | Median Y |
|---|---|---|---|---|---|---|
| All | 4,710 | | | | | |
| Cells | 1,108 | 23.52 % | FSC-H | SSC-H | 3,341,335 | 2,412,314 |
| Single cells | 675 | 60.92 % | FSC-H | FSC-A | 2,786,823 | 8,611,563 |
| Live cells | 507 | 75.11 % | APC-Cy7-H | FSC-H | 7,978 | 3,231,350 |
| Live GFP negative | 265 | 52.27 % | FITC-A | Count | 131,520 | |
| Live GFP positive | 240 | 47.34 % | FITC-A | Count | 7,040,231 | |
| Dead cells | 165 | 24.44 % | APC-Cy7-H | FSC-H | 107,804 | 1,661,886 |
| Dead GFP negative | 23 | 13.94 % | FITC-A | Count | 72,675 | |
| Dead GFP positive | 142 | 86.06 % | FITC-A | Count | 309,044 | |

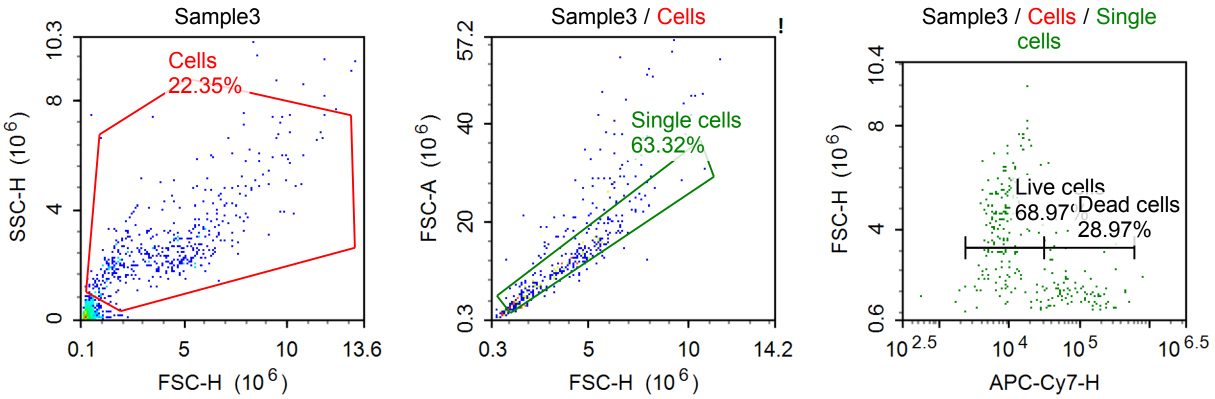


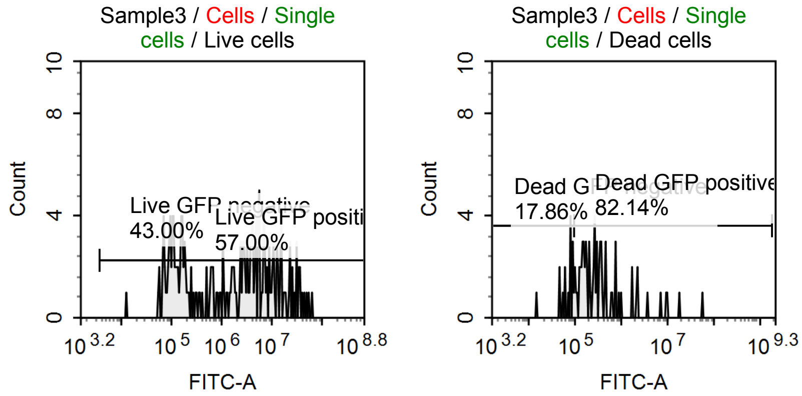


Sample Statistics of Sample3

| Gate | Count | % Parent | X | Y | Median X | Median Y |
|---|---|---|---|---|---|---|
| All | 2,049 | | | | | |
| Cells | 458 | 22.35 % | FSC-H | SSC-H | 3,891,268 | 2,436,090 |
| Single cells | 290 | 63.32 % | FSC-H | FSC-A | 3,295,354 | 9,974,012 |
| Live cells | 200 | 68.97 % | APC-Cy7-H | FSC-H | 8,525 | 4,340,796 |
| Live GFP negative | 86 | 43.00 % | FITC-A | Count | 156,977 | |
| Live GFP positive | 114 | 57.00 % | FITC-A | Count | 5,941,914 | |
| Dead cells | 84 | 28.97 % | APC-Cy7-H | FSC-H | 71,435 | 1,600,527 |
| Dead GFP negative | 15 | 17.86 % | FITC-A | Count | 80,514 | |
| Dead GFP positive | 69 | 82.14 % | FITC-A | Count | 294,234 | |

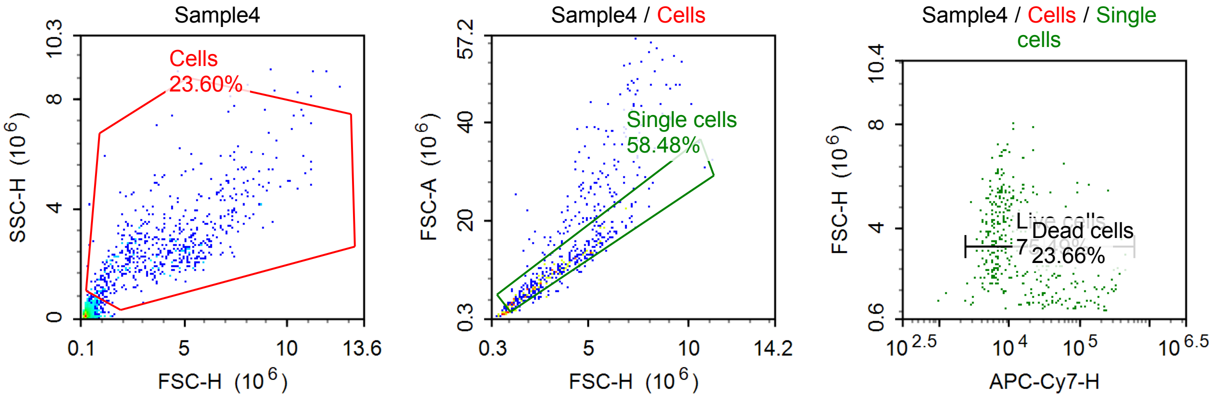

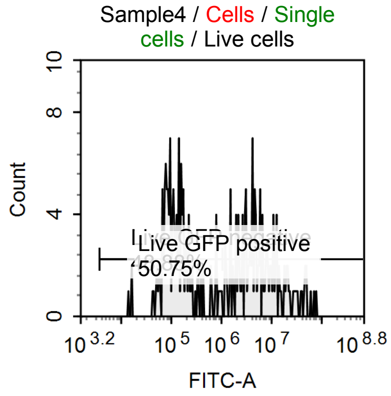


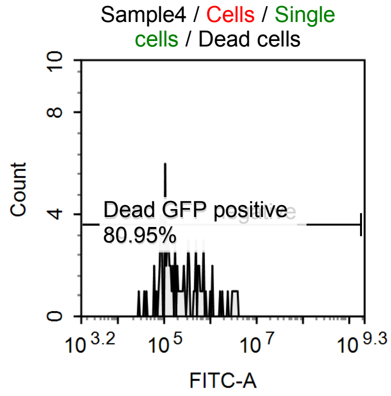


Sample Statistics of Sample4

| Gate | Count | % Parent | X | Y | Median X | Median Y |
|---|---|---|---|---|---|---|
| All | 2,572 | | | | | |
| Cells | 607 | 23.60 % | FSC-H | SSC-H | 3,885,440 | 2,504,523 |
| Single cells | 355 | 58.48 % | FSC-H | FSC-A | 3,111,925 | 9,585,846 |
| Live cells | 268 | 75.49 % | APC-Cy7-H | FSC-H | 7,477 | 3,707,919 |
| Live GFP negative | 131 | 48.88 % | FITC-A | Count | 126,696 | |
| Live GFP positive | 136 | 50.75 % | FITC-A | Count | 5,106,137 | |
| Dead cells | 84 | 23.66 % | APC-Cy7-H | FSC-H | 83,873 | 1,794,502 |
| Dead GFP negative | 16 | 19.05 % | FITC-A | Count | 80,353 | |
| Dead GFP positive | 68 | 80.95 % | FITC-A | Count | 303,495 | |

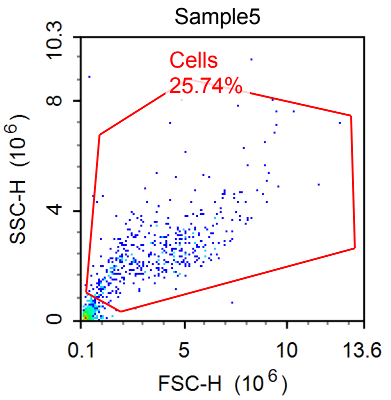


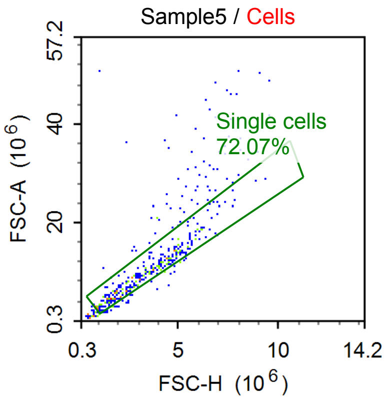


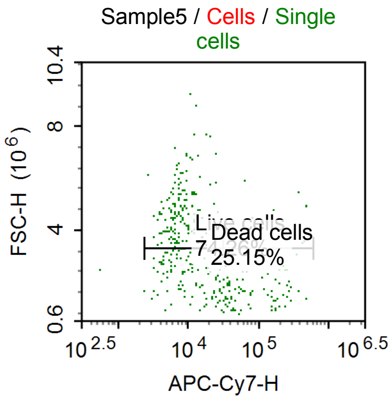


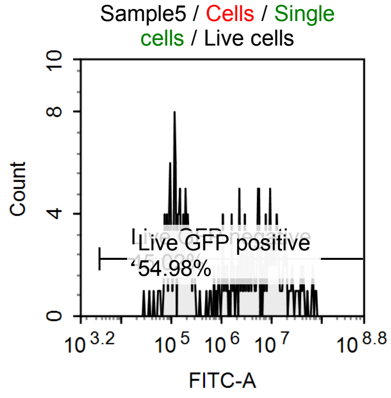


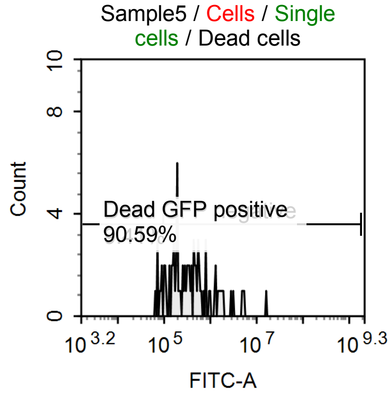

Sample Statistics of Sample5

| Gate | Count | % Parent | X | Y | Median X | Median Y |
|---|---|---|---|---|---|---|
| All | 1,822 | | | | | |
| Cells | 469 | 25.74 % | FSC-H | SSC-H | 3,534,620 | 2,385,663 |
| Single cells | 338 | 72.07 % | FSC-H | FSC-A | 3,153,708 | 9,489,454 |
| Live cells | 251 | 74.26 % | APC-Cy7-H | FSC-H | 7,808 | 3,901,235 |
| Live GFP negative | 113 | 45.02 % | FITC-A | Count | 151,076 | |
| Live GFP positive | 138 | 54.98 % | FITC-A | Count | 7,881,293 | |
| Dead cells | 85 | 25.15 % | APC-Cy7-H | FSC-H | 69,873 | 1,746,045 |
| Dead GFP negative | 8 | 9.41 % | FITC-A | Count | 78,276 | |
| Dead GFP positive | 77 | 90.59 % | FITC-A | Count | 378,915 | |

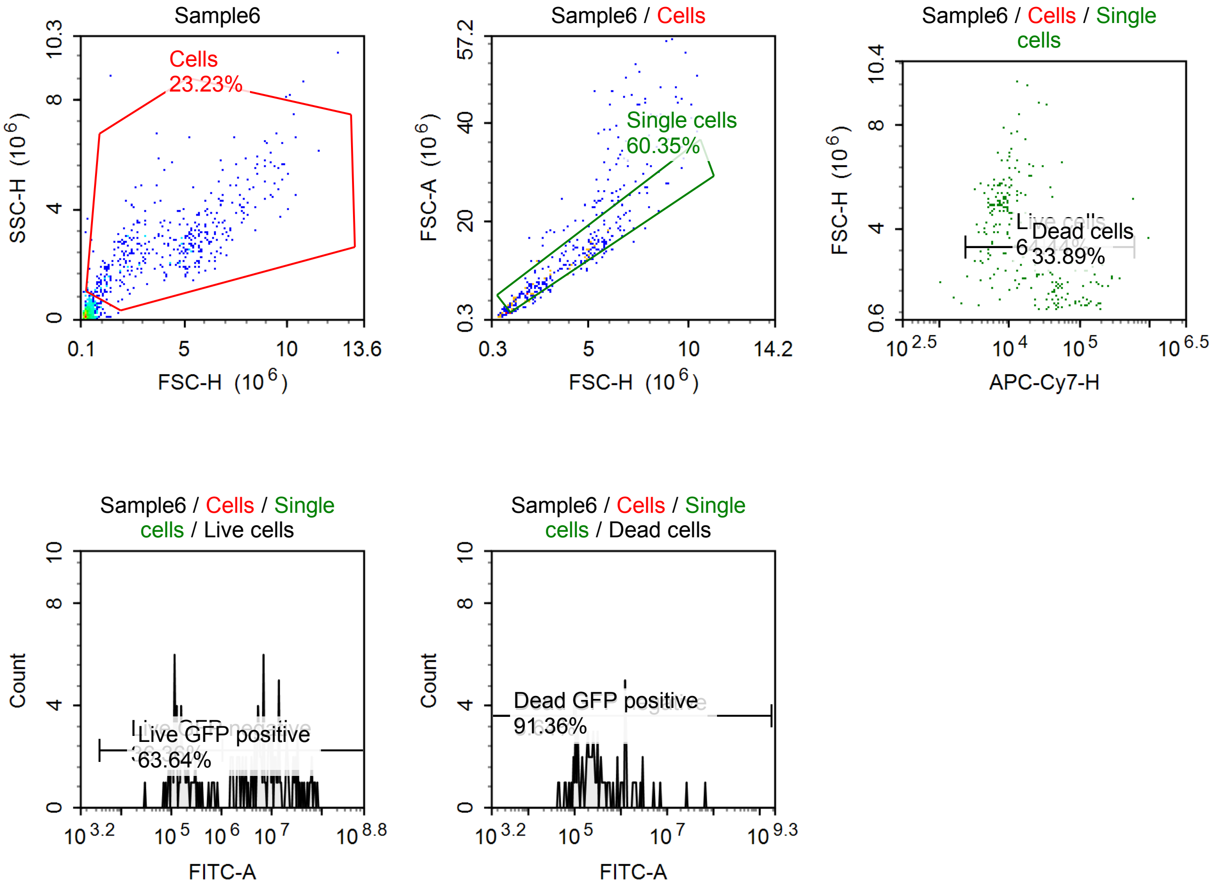


Sample Statistics of Sample6

| Gate | Count | % Parent | X | Y | Median X | Median Y |
|---|---|---|---|---|---|---|
| All | 1,705 | | | | | |
| Cells | 396 | 23.23 % | FSC-H | SSC-H | 4,413,680 | 2,702,367 |
| Single cells | 239 | 60.35 % | FSC-H | FSC-A | 3,519,951 | 10,497,204 |
| Live cells | 154 | 64.44 % | APC-Cy7-H | FSC-H | 8,774 | 4,871,845 |
| Live GFP negative | 56 | 36.36 % | FITC-A | Count | 159,561 | |
| Live GFP positive | 98 | 63.64 % | FITC-A | Count | 8,849,141 | |
| Dead cells | 81 | 33.89 % | APC-Cy7-H | FSC-H | 66,096 | 1,883,003 |
| Dead GFP negative | 7 | 8.64 % | FITC-A | Count | 65,252 | |
| Dead GFP positive | 74 | 91.36 % | FITC-A | Count | 316,340 | |

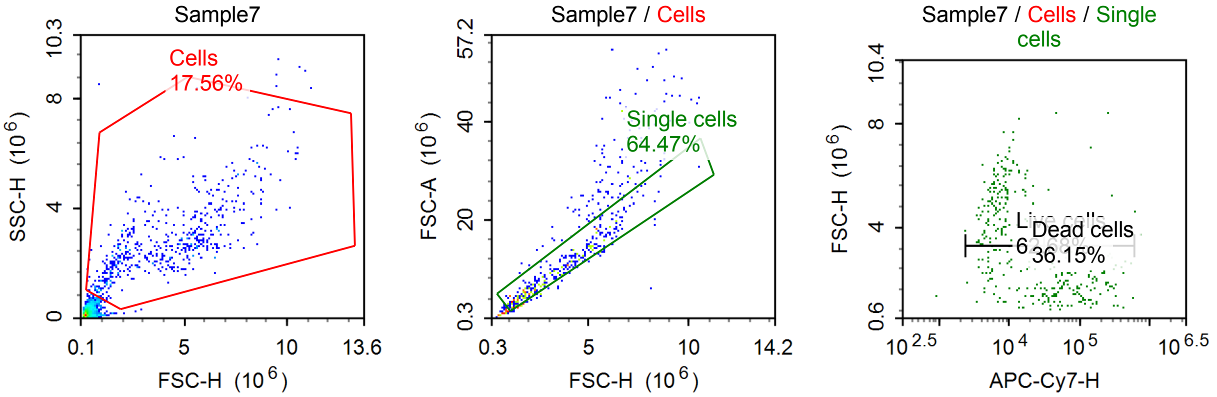


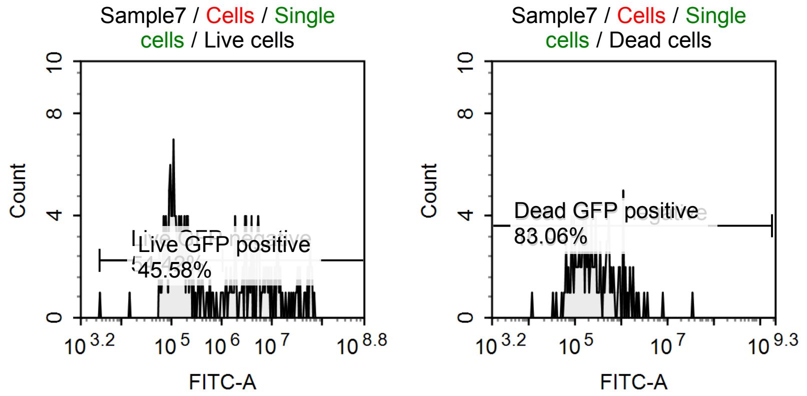


Sample Statistics of Sample7

| Gate | Count | % Parent | X | Y | Median X | Median Y |
|---|---|---|---|---|---|---|
| All | 3,030 | | | | | |
| Cells | 532 | 17.56 % | FSC-H | SSC-H | 3,930,274 | 2,599,425 |
| Single cells | 343 | 64.47 % | FSC-H | FSC-A | 3,048,308 | 9,053,321 |
| Live cells | 215 | 62.68 % | APC-Cy7-H | FSC-H | 7,250 | 4,331,044 |
| Live GFP negative | 117 | 54.42 % | FITC-A | Count | 132,314 | |
| Live GFP positive | 98 | 45.58 % | FITC-A | Count | 5,806,275 | |
| Dead cells | 124 | 36.15 % | APC-Cy7-H | FSC-H | 100,424 | 1,799,234 |
| Dead GFP negative | 21 | 16.94 % | FITC-A | Count | 72,178 | |
| Dead GFP positive | 103 | 83.06 % | FITC-A | Count | 318,928 | |

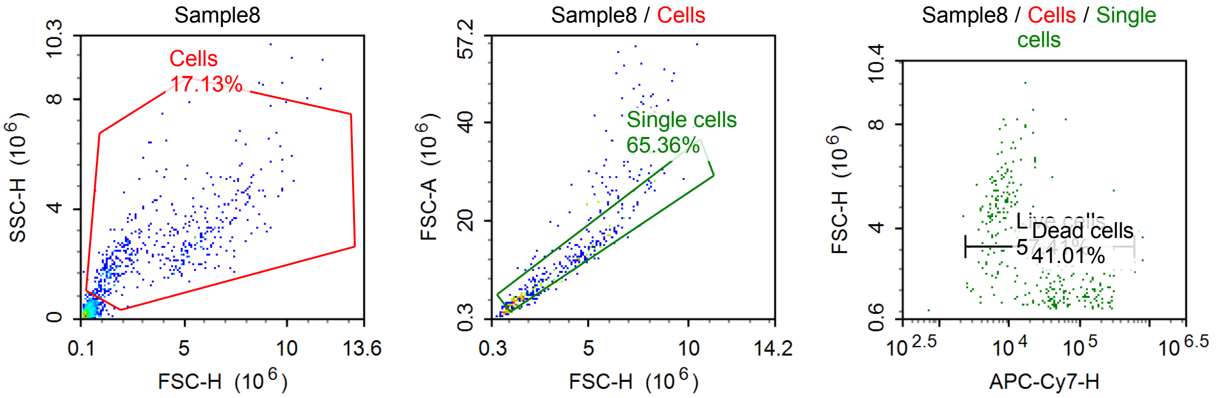

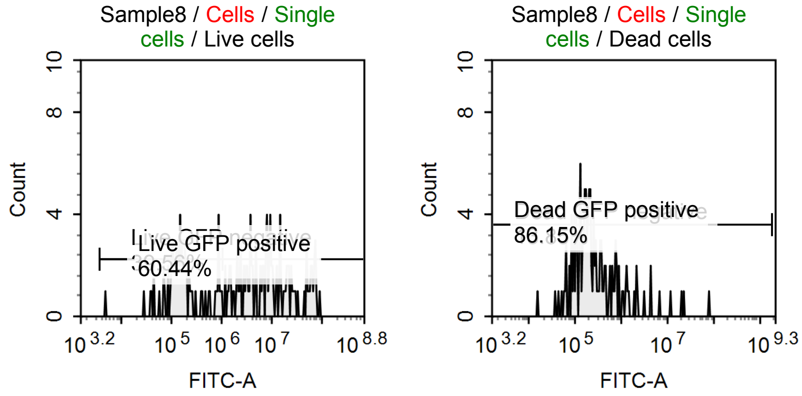


Sample Statistics of Sample8

| Gate | Count | % Parent | X | Y | Median X | Median Y |
|---|---|---|---|---|---|---|
| All | 2,831 | | | | | |
| Cells | 485 | 17.13 % | FSC-H | SSC-H | 3,317,207 | 2,462,777 |
| Single cells | 317 | 65.36 % | FSC-H | FSC-A | 2,724,524 | 8,063,302 |
| Live cells | 182 | 57.41 % | APC-Cy7-H | FSC-H | 8,019 | 4,534,794 |
| Live GFP negative | 72 | 39.56 % | FITC-A | Count | 157,721 | |
| Live GFP positive | 110 | 60.44 % | FITC-A | Count | 8,834,386 | |
| Dead cells | 130 | 41.01 % | APC-Cy7-H | FSC-H | 76,281 | 1,606,253 |
| Dead GFP negative | 18 | 13.85 % | FITC-A | Count | 70,989 | |
| Dead GFP positive | 112 | 86.15 % | FITC-A | Count | 251,994 | |

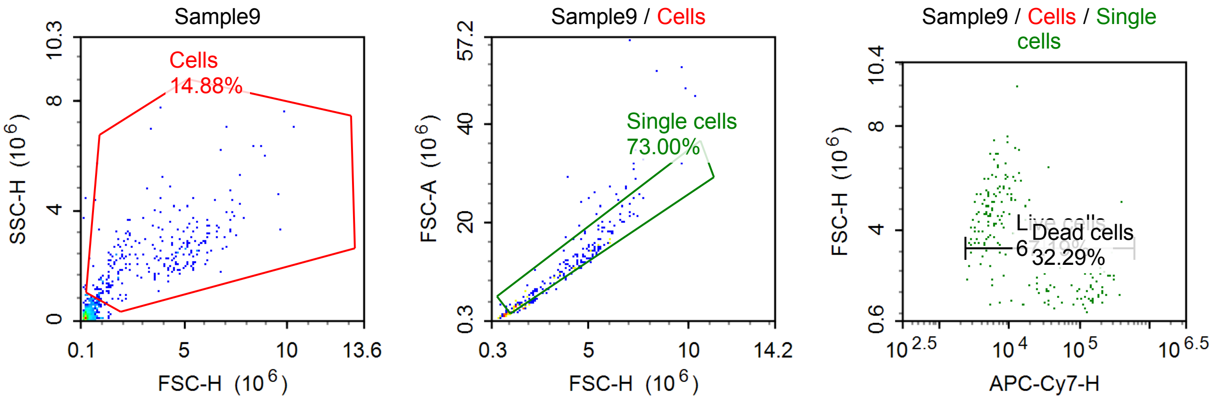


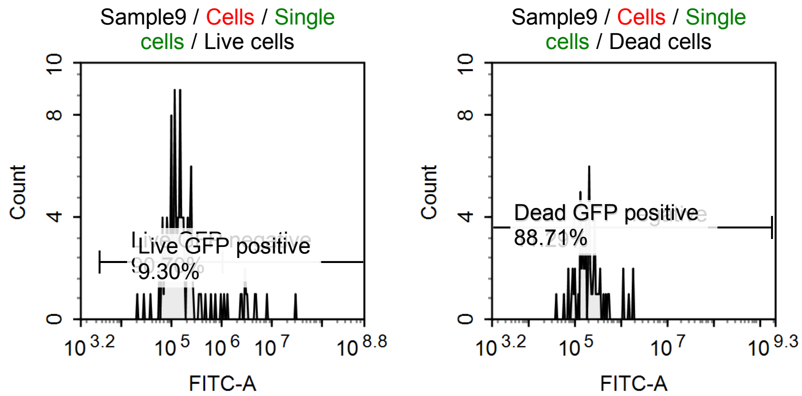

Sample Statistics of Sample9

| Gate | Count | % Parent | X | Y | Median X | Median Y |
|---|---|---|---|---|---|---|
| All | 1,767 | | | | | |
| Cells | 263 | 14.88 % | FSC-H | SSC-H | 3,701,997 | 2,345,079 |
| Single cells | 192 | 73.00 % | FSC-H | FSC-A | 3,803,402 | 10,740,139 |
| Live cells | 129 | 67.19 % | APC-Cy7-H | FSC-H | 5,920 | 4,659,696 |
| Live GFP negative | 117 | 90.70 % | FITC-A | Count | 141,135 | |
| Live GFP positive | 12 | 9.30 % | FITC-A | Count | 3,154,973 | |
| Dead cells | 62 | 32.29 % | APC-Cy7-H | FSC-H | 141,446 | 1,764,605 |
| Dead GFP negative | 7 | 11.29 % | FITC-A | Count | 73,979 | |
| Dead GFP positive | 55 | 88.71 % | FITC-A | Count | 204,291 | |

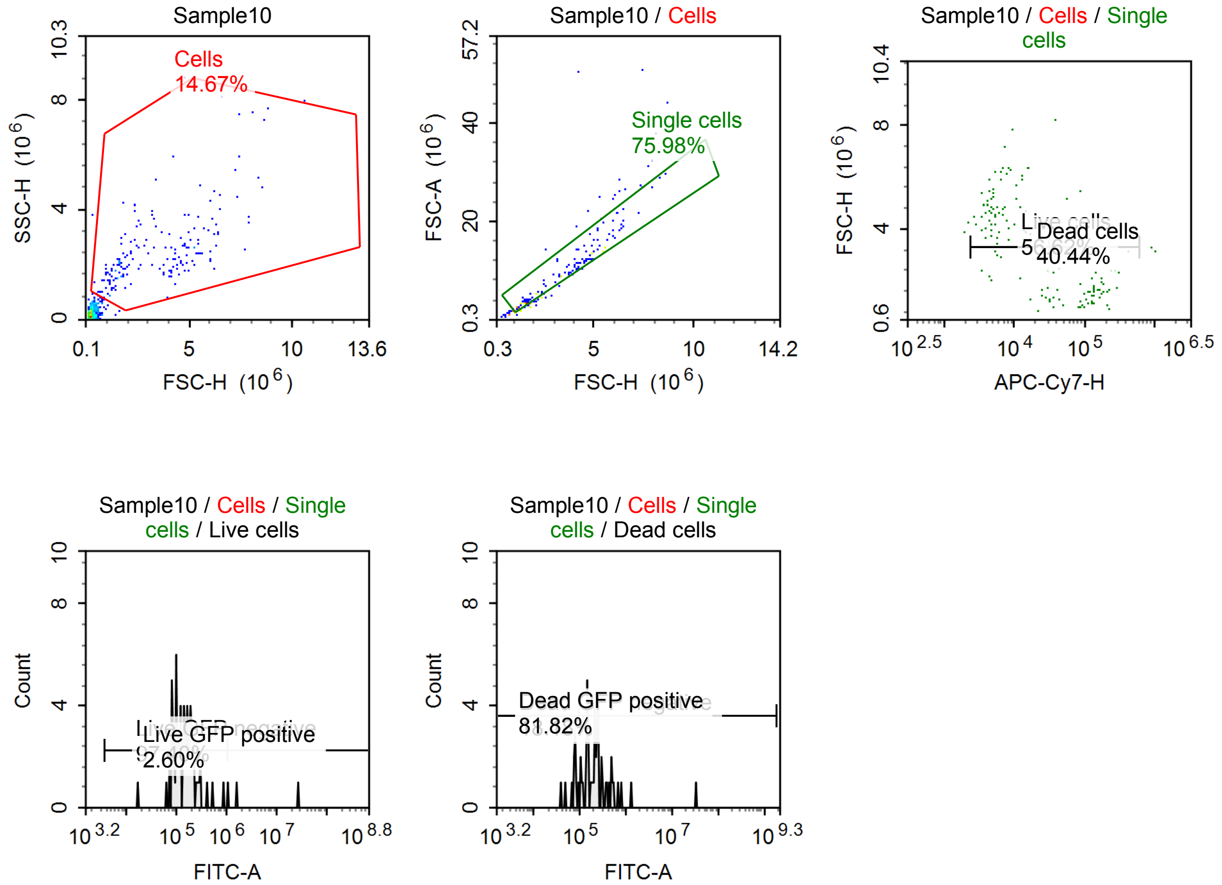


Sample Statistics of Sample10

| Gate | Count | % Parent | X | Y | Median X | Median Y |
|---|---|---|---|---|---|---|
| All | 1,220 | | | | | |
| Cells | 179 | 14.67 % | FSC-H | SSC-H | 3,198,382 | 2,176,726 |
| Single cells | 136 | 75.98 % | FSC-H | FSC-A | 3,266,136 | 9,259,284 |
| Live cells | 77 | 56.62 % | APC-Cy7-H | FSC-H | 5,729 | 4,515,043 |
| Live GFP negative | 75 | 97.40 % | FITC-A | Count | 149,511 | |
| Live GFP positive | 2 | 2.60 % | FITC-A | Count | 14,864,784 | |
| Dead cells | 55 | 40.44 % | APC-Cy7-H | FSC-H | 128,850 | 1,613,294 |
| Dead GFP negative | 10 | 18.18 % | FITC-A | Count | 77,893 | |
| Dead GFP positive | 45 | 81.82 % | FITC-A | Count | 224,135 | |

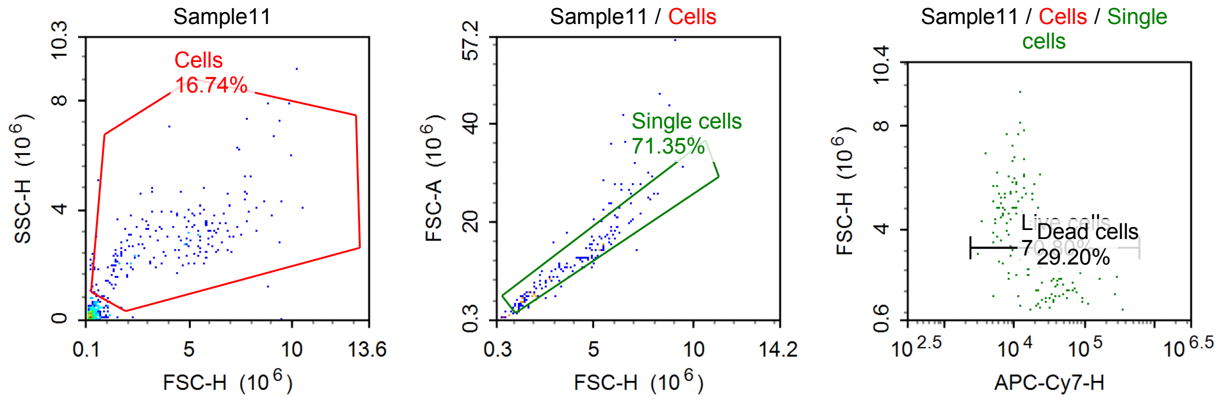


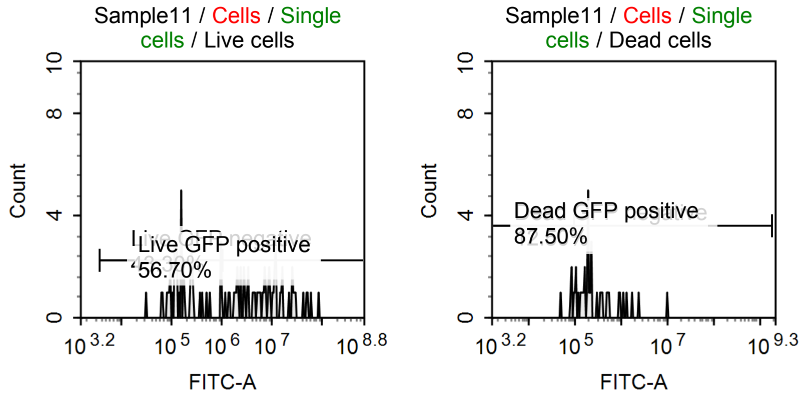


Sample Statistics of Sample11

| Gate | Count | % Parent | X | Y | Median X | Median Y |
|---|---|---|---|---|---|---|
| All | 1,147 | | | | | |
| Cells | 192 | 16.74 % | FSC-H | SSC-H | 4,315,637 | 2,861,498 |
| Single cells | 137 | 71.35 % | FSC-H | FSC-A | 3,441,063 | 10,941,760 |
| Live cells | 97 | 70.80 % | APC-Cy7-H | FSC-H | 9,560 | 4,720,069 |
| Live GFP negative | 42 | 43.30 % | FITC-A | Count | 162,364 | |
| Live GFP positive | 55 | 56.70 % | FITC-A | Count | 7,660,529 | |
| Dead cells | 40 | 29.20 % | APC-Cy7-H | FSC-H | 54,236 | 1,869,138 |
| Dead GFP negative | 5 | 12.50 % | FITC-A | Count | 84,132 | |
| Dead GFP positive | 35 | 87.50 % | FITC-A | Count | 220,875 | |

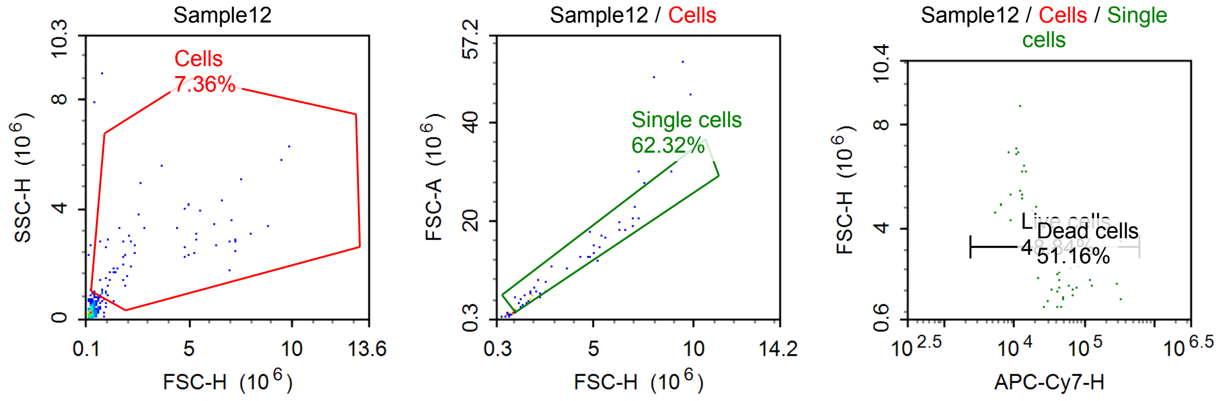

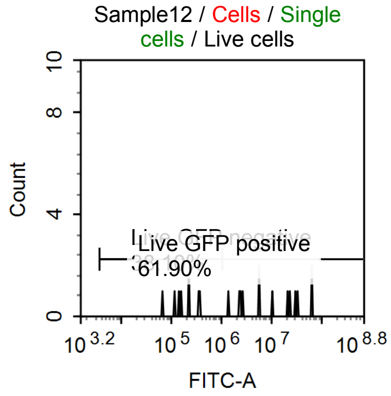


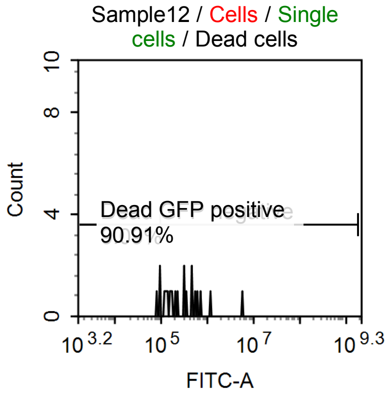


Sample Statistics of Sample12

| Gate | Count | % Parent | X | Y | Median X | Median Y |
|---|---|---|---|---|---|---|
| All | 937 | | | | | |
| Cells | 69 | 7.36 % | FSC-H | SSC-H | 1,866,259 | 2,347,096 |
| Single cells | 43 | 62.32 % | FSC-H | FSC-A | 2,525,891 | 7,809,130 |
| Live cells | 21 | 48.84 % | APC-Cy7-H | FSC-H | 12,793 | 5,282,960 |
| Live GFP negative | 8 | 38.10 % | FITC-A | Count | 193,804 | |
| Live GFP positive | 13 | 61.90 % | FITC-A | Count | 10,691,249 | |
| Dead cells | 22 | 51.16 % | APC-Cy7-H | FSC-H | 59,806 | 1,816,422 |
| Dead GFP negative | 2 | 9.09 % | FITC-A | Count | 86,359 | |
| Dead GFP positive | 20 | 90.91 % | FITC-A | Count | 276,436 | |

#### Specimen2

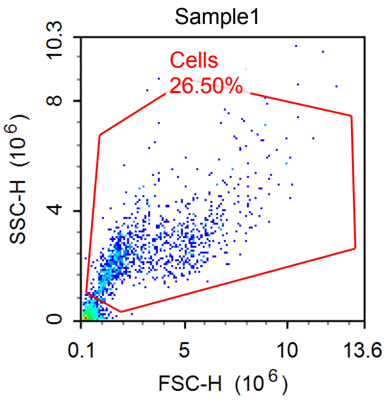


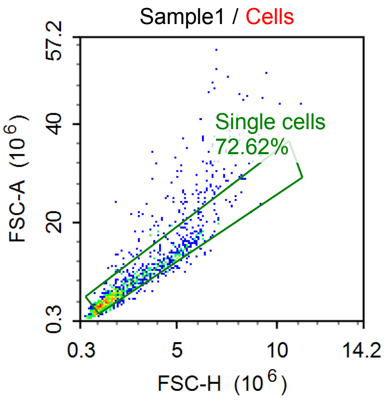


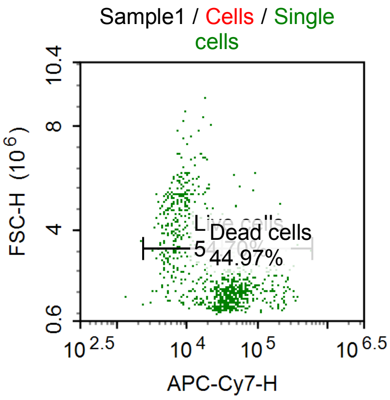


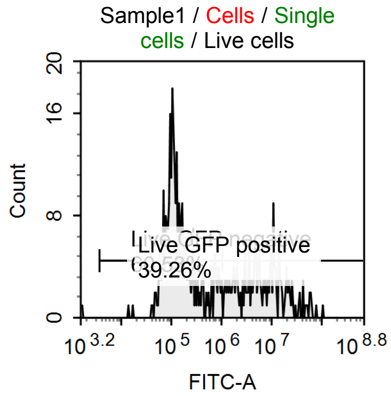


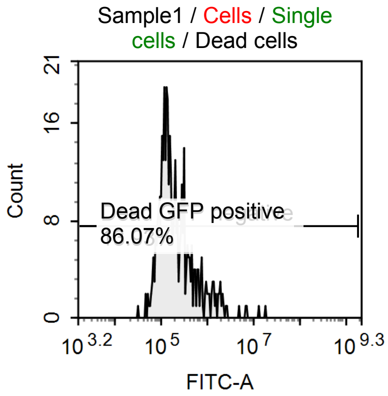

Sample Statistics of Sample1

| Gate | Count | % Parent | X | Y | Median X | Median Y |
|---|---|---|---|---|---|---|
| All | 4,645 | | | | | |
| Cells | 1,231 | 26.50 % | FSC-H | SSC-H | 2,235,793 | 2,351,029 |
| Single cells | 894 | 72.62 % | FSC-H | FSC-A | 2,051,073 | 6,011,934 |
| Live cells | 489 | 54.70 % | APC-Cy7-H | FSC-H | 9,872 | 3,309,194 |
| Live GFP negative | 296 | 60.53 % | FITC-A | Count | 127,035 | |
| Live GFP positive | 192 | 39.26 % | FITC-A | Count | 6,375,480 | |
| Dead cells | 402 | 44.97 % | APC-Cy7-H | FSC-H | 52,486 | 1,616,251 |
| Dead GFP negative | 56 | 13.93 % | FITC-A | Count | 79,580 | |
| Dead GFP positive | 346 | 86.07 % | FITC-A | Count | 201,069 | |

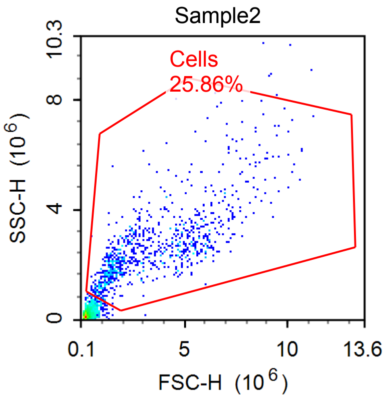


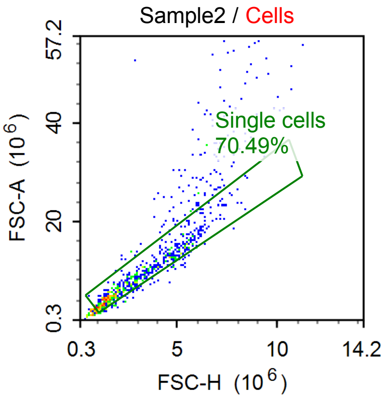


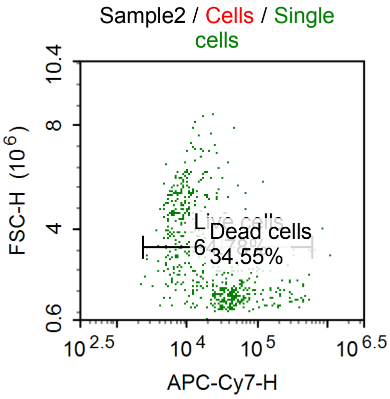


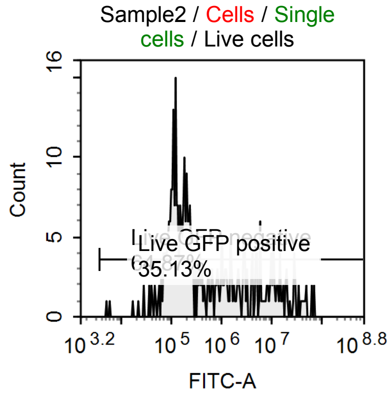


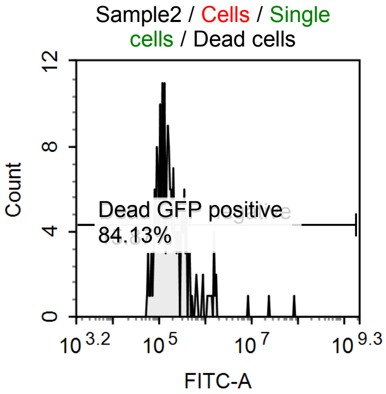


Sample Statistics of Sample2

| Gate | Count | % Parent | X | Y | Median X | Median Y |
|---|---|---|---|---|---|---|
| All | 3,302 | | | | | |
| Cells | 854 | 25.86 % | FSC-H | SSC-H | 2,948,085 | 2,461,422 |
| Single cells | 602 | 70.49 % | FSC-H | FSC-A | 2,604,295 | 7,555,312 |
| Live cells | 390 | 64.78 % | APC-Cy7-H | FSC-H | 10,351 | 3,754,638 |
| Live GFP negative | 253 | 64.87 % | FITC-A | Count | 152,541 | |
| Live GFP positive | 137 | 35.13 % | FITC-A | Count | 6,091,685 | |
| Dead cells | 208 | 34.55 % | APC-Cy7-H | FSC-H | 54,012 | 1,578,431 |
| Dead GFP negative | 33 | 15.87 % | FITC-A | Count | 81,854 | |
| Dead GFP positive | 175 | 84.13 % | FITC-A | Count | 182,811 | |

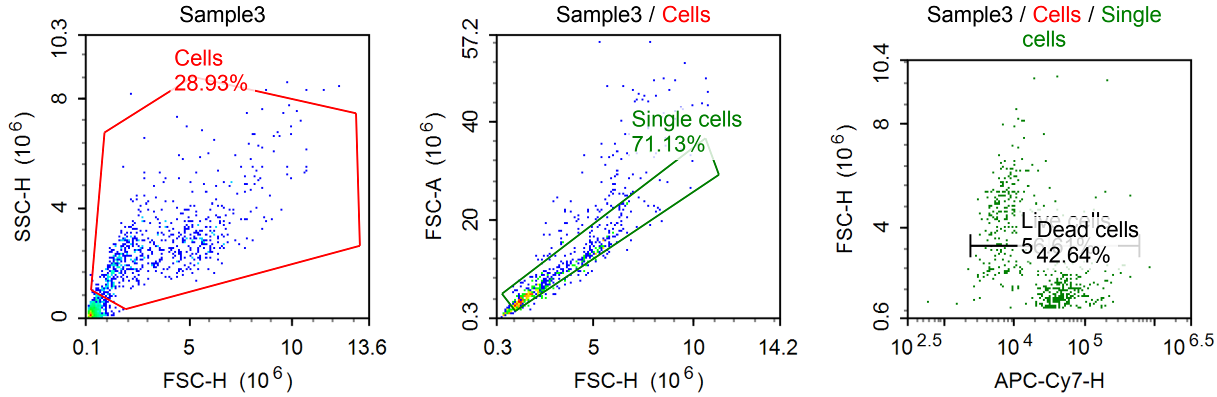


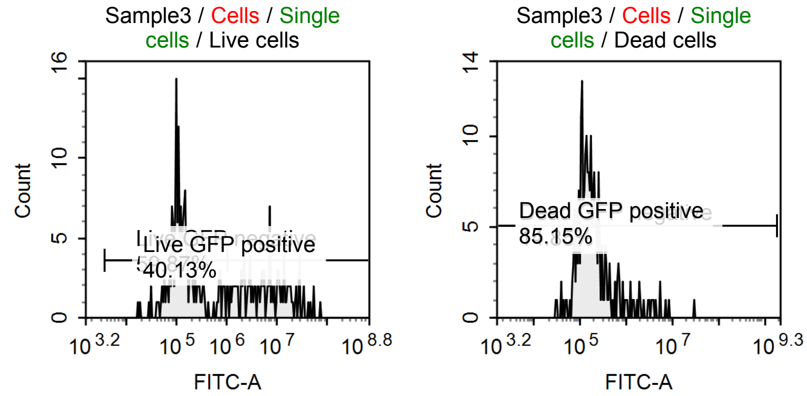


Sample Statistics of Sample3

| Gate | Count | % Parent | X | Y | Median X | Median Y |
|---|---|---|---|---|---|---|
| All | 2,610 | | | | | |
| Cells | 755 | 28.93 % | FSC-H | SSC-H | 2,697,538 | 2,419,795 |
| Single cells | 537 | 71.13 % | FSC-H | FSC-A | 2,281,047 | 6,655,947 |
| Live cells | 304 | 56.61 % | APC-Cy7-H | FSC-H | 8,400 | 3,977,941 |
| Live GFP negative | 182 | 59.87 % | FITC-A | Count | 125,767 | |
| Live GFP positive | 122 | 40.13 % | FITC-A | Count | 6,676,502 | |
| Dead cells | 229 | 42.64 % | APC-Cy7-H | FSC-H | 55,479 | 1,585,339 |
| Dead GFP negative | 34 | 14.85 % | FITC-A | Count | 76,970 | |
| Dead GFP positive | 195 | 85.15 % | FITC-A | Count | 188,594 | |

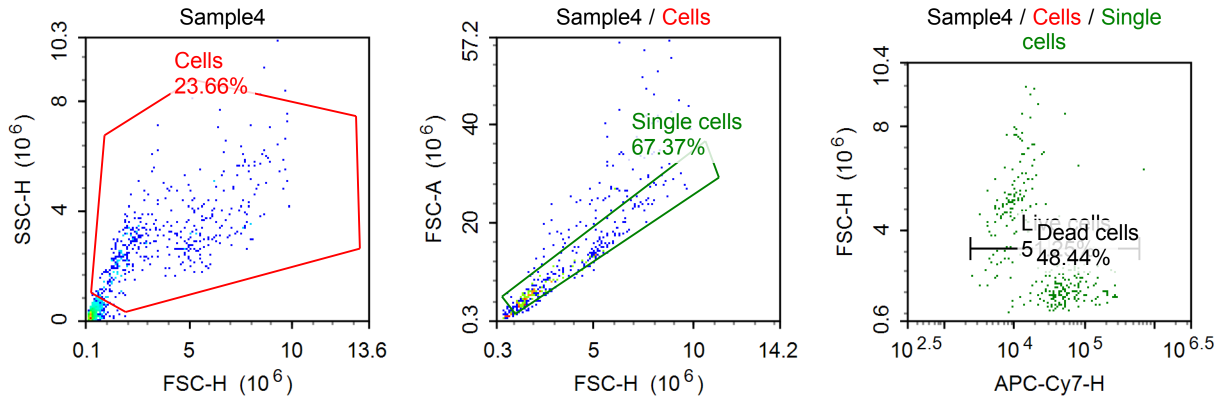

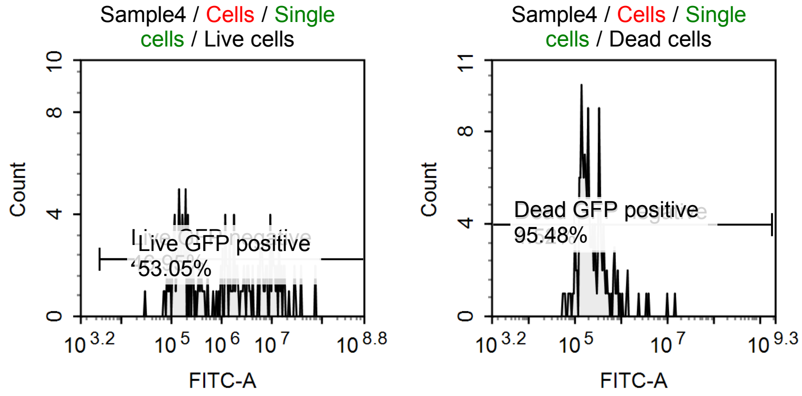


Sample Statistics of Sample4

| Gate | Count | % Parent | X | Y | Median X | Median Y |
|---|---|---|---|---|---|---|
| All | 2,008 | | | | | |
| Cells | 475 | 23.66 % | FSC-H | SSC-H | 2,822,444 | 2,881,358 |
| Single cells | 320 | 67.37 % | FSC-H | FSC-A | 2,499,564 | 7,237,218 |
| Live cells | 164 | 51.25 % | APC-Cy7-H | FSC-H | 10,674 | 4,770,339 |
| Live GFP negative | 77 | 46.95 % | FITC-A | Count | 181,807 | |
| Live GFP positive | 87 | 53.05 % | FITC-A | Count | 6,487,246 | |
| Dead cells | 155 | 48.44 % | APC-Cy7-H | FSC-H | 67,347 | 1,843,644 |
| Dead GFP negative | 7 | 4.52 % | FITC-A | Count | 71,571 | |
| Dead GFP positive | 148 | 95.48 % | FITC-A | Count | 210,523 | |

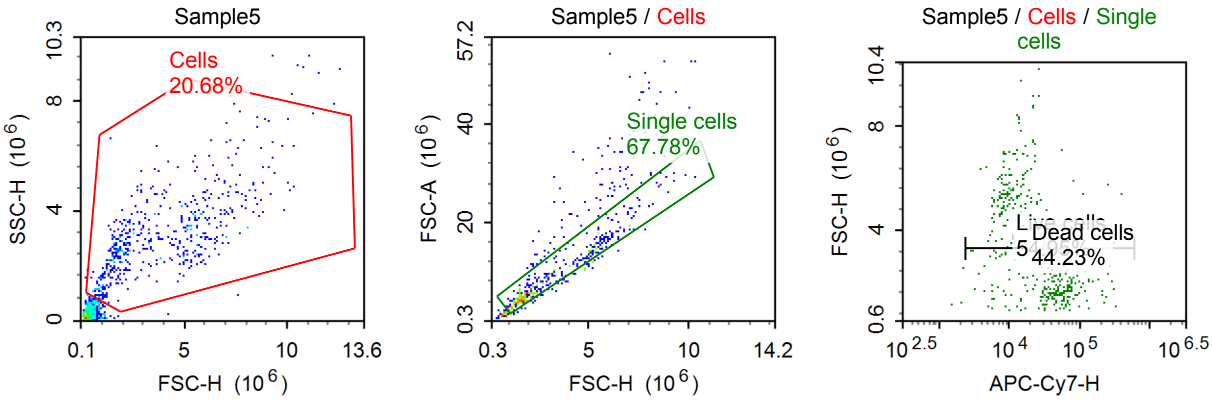


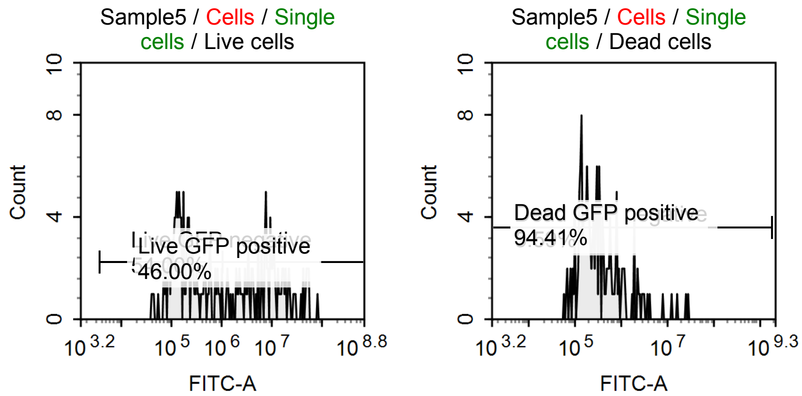

Sample Statistics of Sample5

| Gate | Count | % Parent | X | Y | Median X | Median Y |
|---|---|---|---|---|---|---|
| All | 2,597 | | | | | |
| Cells | 537 | 20.68 % | FSC-H | SSC-H | 3,201,259 | 2,902,834 |
| Single cells | 364 | 67.78 % | FSC-H | FSC-A | 2,634,179 | 7,944,045 |
| Live cells | 200 | 54.95 % | APC-Cy7-H | FSC-H | 10,676 | 4,978,579 |
| Live GFP negative | 108 | 54.00 % | FITC-A | Count | 169,349 | |
| Live GFP positive | 92 | 46.00 % | FITC-A | Count | 8,562,360 | |
| Dead cells | 161 | 44.23 % | APC-Cy7-H | FSC-H | 63,439 | 1,851,952 |
| Dead GFP negative | 9 | 5.59 % | FITC-A | Count | 75,439 | |
| Dead GFP positive | 152 | 94.41 % | FITC-A | Count | 303,496 | |

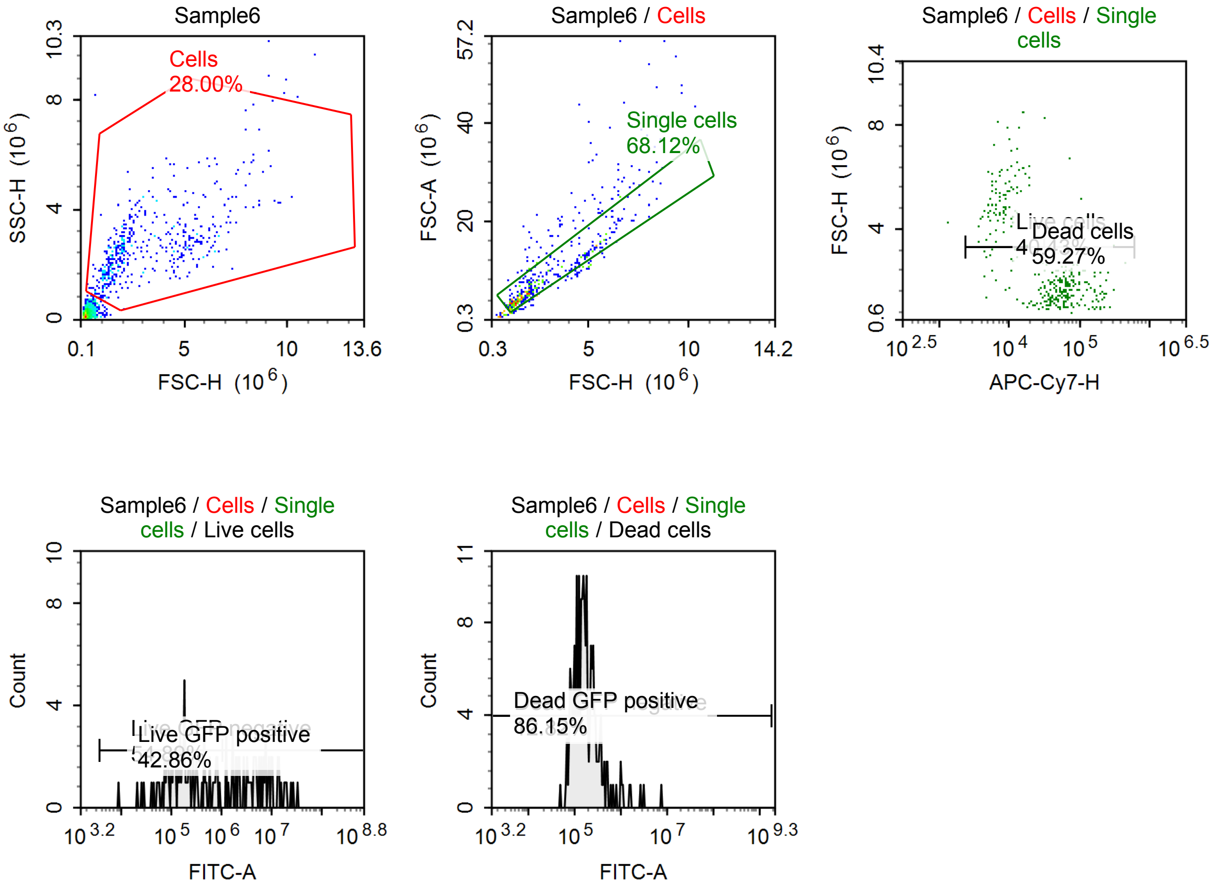


Sample Statistics of Sample6

| Gate | Count | % Parent | X | Y | Median X | Median Y |
|---|---|---|---|---|---|---|
| All | 1,725 | | | | | |
| Cells | 483 | 28.00 % | FSC-H | SSC-H | 2,320,130 | 2,577,508 |
| Single cells | 329 | 68.12 % | FSC-H | FSC-A | 2,041,395 | 6,350,543 |
| Live cells | 133 | 40.43 % | APC-Cy7-H | FSC-H | 9,475 | 4,598,399 |
| Live GFP negative | 73 | 54.89 % | FITC-A | Count | 175,921 | |
| Live GFP positive | 57 | 42.86 % | FITC-A | Count | 5,312,851 | |
| Dead cells | 195 | 59.27 % | APC-Cy7-H | FSC-H | 62,917 | 1,645,976 |
| Dead GFP negative | 25 | 12.82 % | FITC-A | Count | 80,512 | |
| Dead GFP positive | 168 | 86.15 % | FITC-A | Count | 178,264 | |

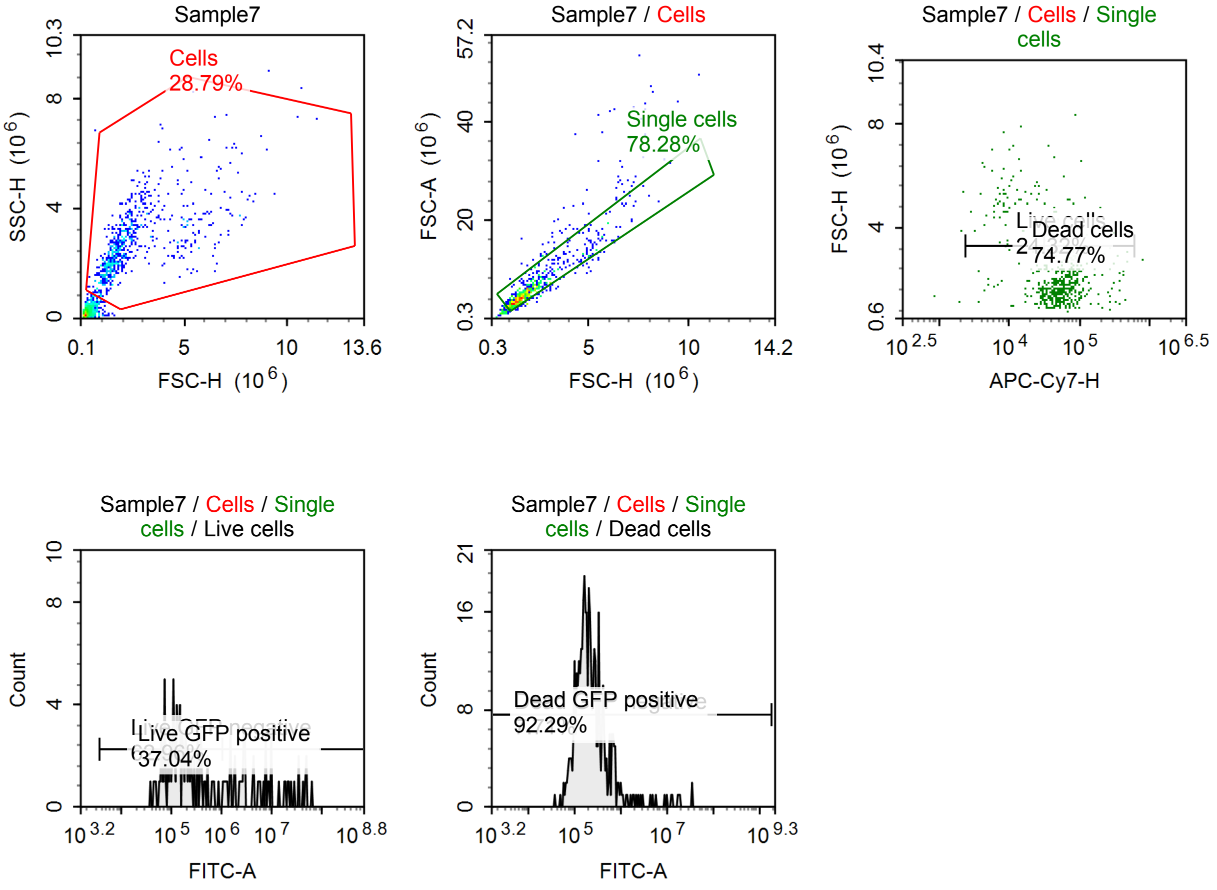


Sample Statistics of Sample7

| Gate | Count | % Parent | X | Y | Median X | Median Y |
|---|---|---|---|---|---|---|
| All | 2,463 | | | | | |
| Cells | 709 | 28.79 % | FSC-H | SSC-H | 1,928,930 | 2,603,590 |
| Single cells | 555 | 78.28 % | FSC-H | FSC-A | 1,861,713 | 5,428,166 |
| Live cells | 135 | 24.32 % | APC-Cy7-H | FSC-H | 16,647 | 2,653,972 |
| Live GFP negative | 85 | 62.96 % | FITC-A | Count | 144,676 | |
| Live GFP positive | 50 | 37.04 % | FITC-A | Count | 5,684,467 | |
| Dead cells | 415 | 74.77 % | APC-Cy7-H | FSC-H | 60,875 | 1,762,465 |
| Dead GFP negative | 32 | 7.71 % | FITC-A | Count | 78,392 | |
| Dead GFP positive | 383 | 92.29 % | FITC-A | Count | 217,543 | |

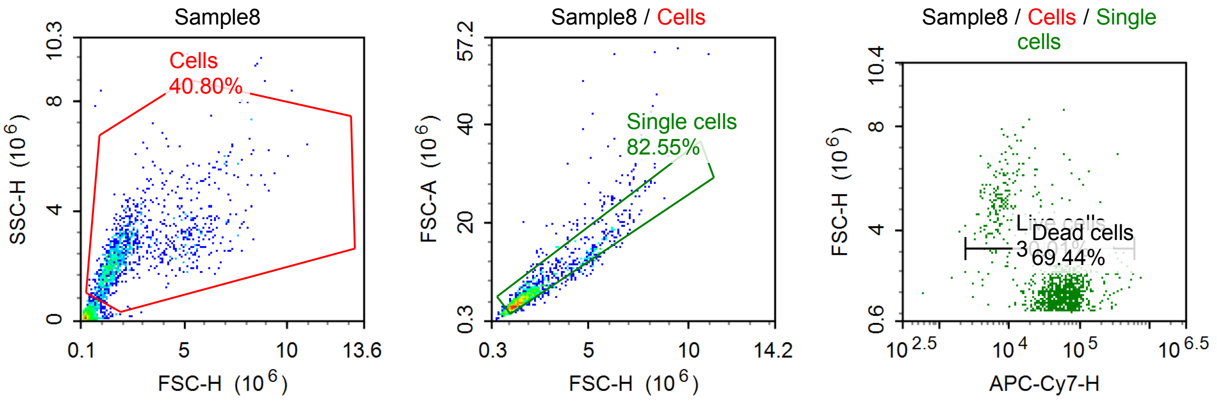

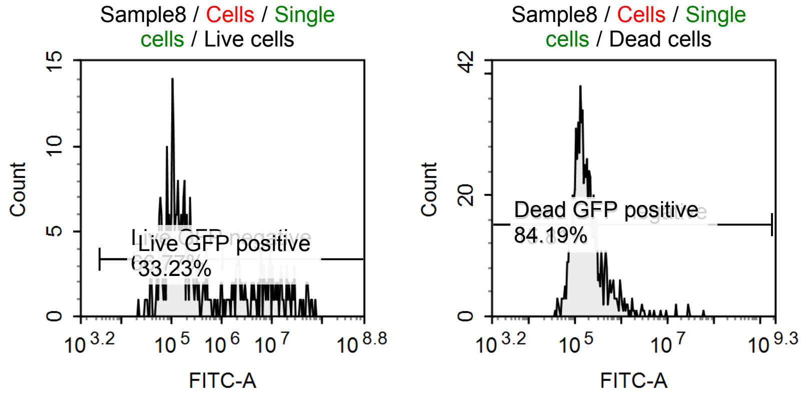


Sample Statistics of Sample8

| Gate | Count | % Parent | X | Y | Median X | Median Y |
|---|---|---|---|---|---|---|
| All | 3,245 | | | | | |
| Cells | 1,324 | 40.80 % | FSC-H | SSC-H | 1,847,640 | 2,384,241 |
| Single cells | 1,093 | 82.55 % | FSC-H | FSC-A | 1,756,710 | 5,037,111 |
| Live cells | 328 | 30.01 % | APC-Cy7-H | FSC-H | 11,733 | 3,928,888 |
| Live GFP negative | 219 | 66.77 % | FITC-A | Count | 128,976 | |
| Live GFP positive | 109 | 33.23 % | FITC-A | Count | 7,402,837 | |
| Dead cells | 759 | 69.44 % | APC-Cy7-H | FSC-H | 63,797 | 1,616,844 |
| Dead GFP negative | 120 | 15.81 % | FITC-A | Count | 79,687 | |
| Dead GFP positive | 639 | 84.19 % | FITC-A | Count | 181,509 | |

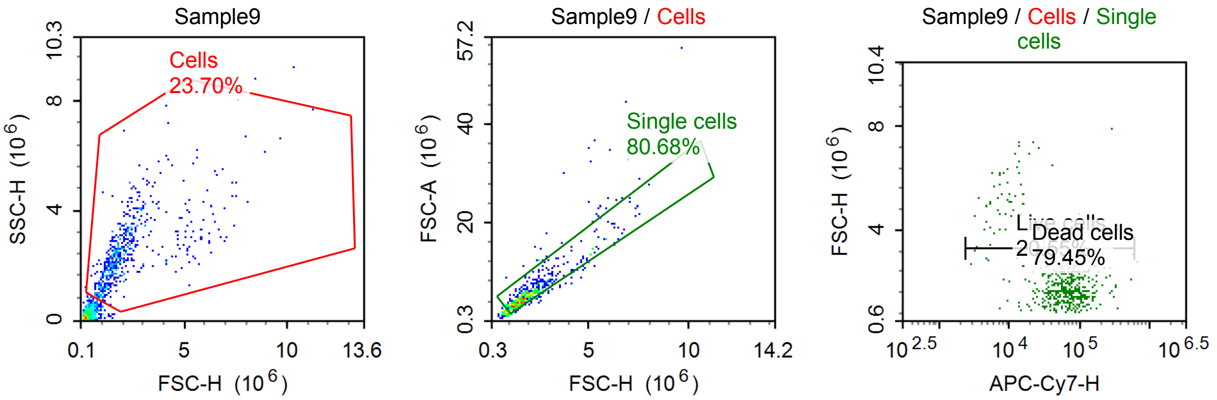


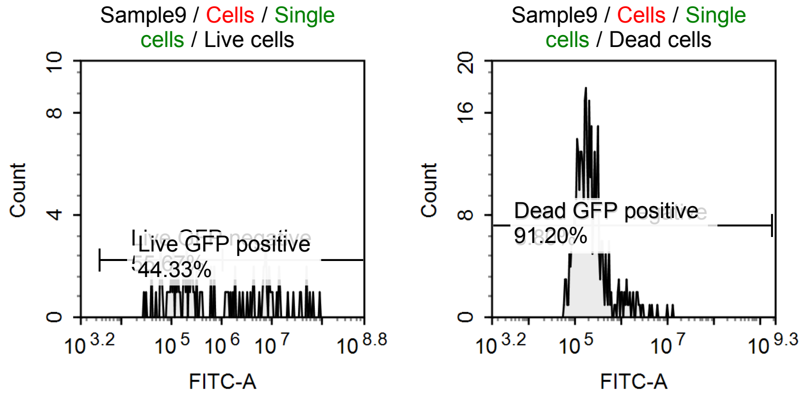

Sample Statistics of Sample9

| Gate | Count | % Parent | X | Y | Median X | Median Y |
|---|---|---|---|---|---|---|
| All | 2,468 | | | | | |
| Cells | 585 | 23.70 % | FSC-H | SSC-H | 1,780,439 | 2,386,910 |
| Single cells | 472 | 80.68 % | FSC-H | FSC-A | 1,801,041 | 5,173,022 |
| Live cells | 97 | 20.55 % | APC-Cy7-H | FSC-H | 13,543 | 3,260,196 |
| Live GFP negative | 54 | 55.67 % | FITC-A | Count | 150,550 | |
| Live GFP positive | 43 | 44.33 % | FITC-A | Count | 7,784,702 | |
| Dead cells | 375 | 79.45 % | APC-Cy7-H | FSC-H | 73,522 | 1,733,445 |
| Dead GFP negative | 33 | 8.80 % | FITC-A | Count | 81,356 | |
| Dead GFP positive | 342 | 91.20 % | FITC-A | Count | 206,177 | |

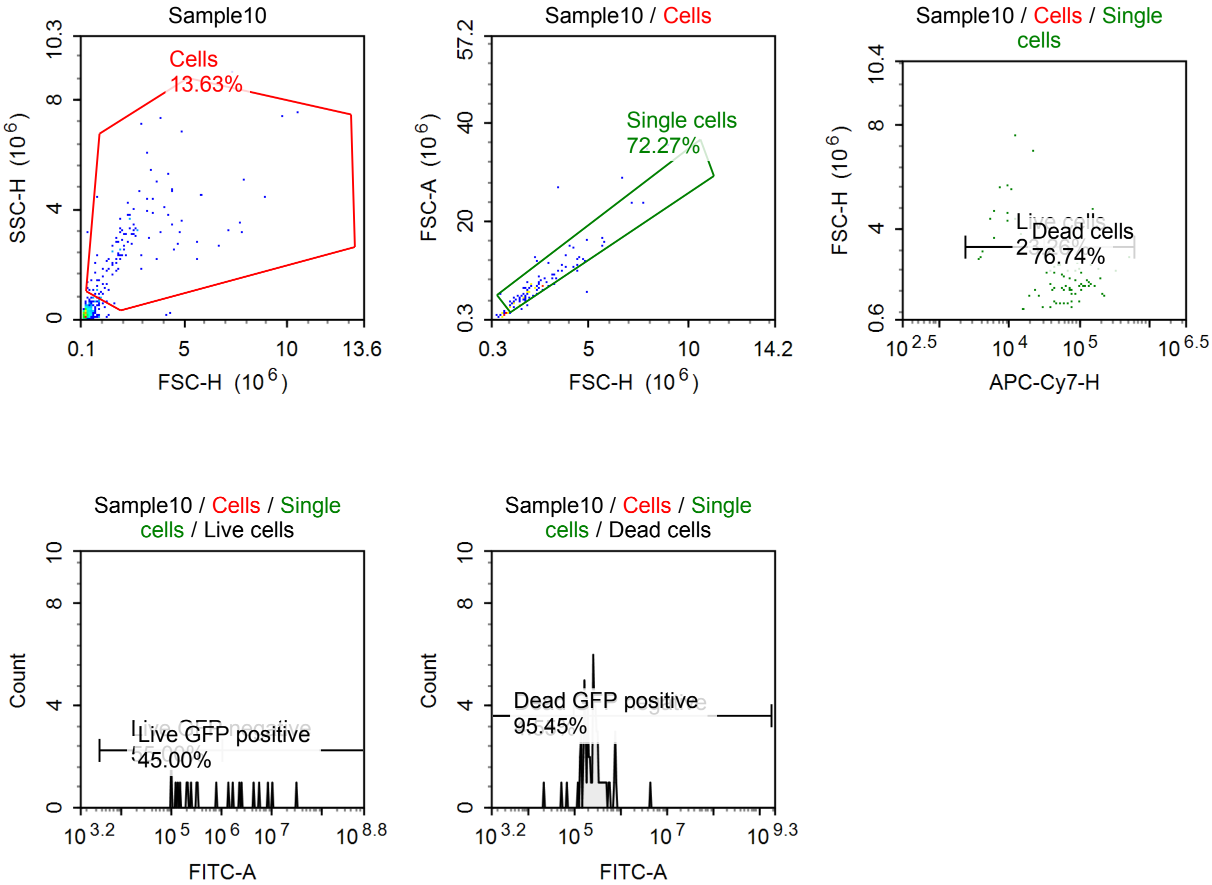


Sample Statistics of Sample10

| Gate | Count | % Parent | X | Y | Median X | Median Y |
|---|---|---|---|---|---|---|
| All | 873 | | | | | |
| Cells | 119 | 13.63 % | FSC-H | SSC-H | 1,951,561 | 2,709,967 |
| Single cells | 86 | 72.27 % | FSC-H | FSC-A | 2,220,467 | 6,592,616 |
| Live cells | 20 | 23.26 % | APC-Cy7-H | FSC-H | 9,872 | 4,100,221 |
| Live GFP negative | 11 | 55.00 % | FITC-A | Count | 205,941 | |
| Live GFP positive | 9 | 45.00 % | FITC-A | Count | 4,450,887 | |
| Dead cells | 66 | 76.74 % | APC-Cy7-H | FSC-H | 74,542 | 2,016,403 |
| Dead GFP negative | 3 | 4.55 % | FITC-A | Count | 53,089 | |
| Dead GFP positive | 63 | 95.45 % | FITC-A | Count | 260,067 | |

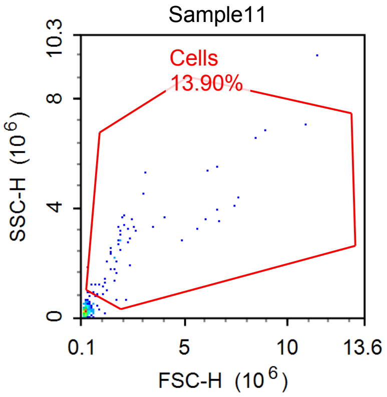


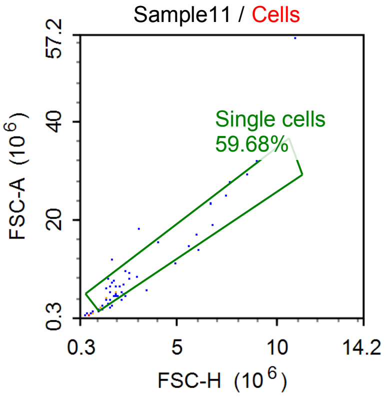


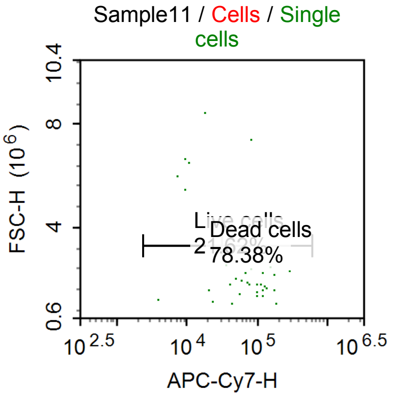


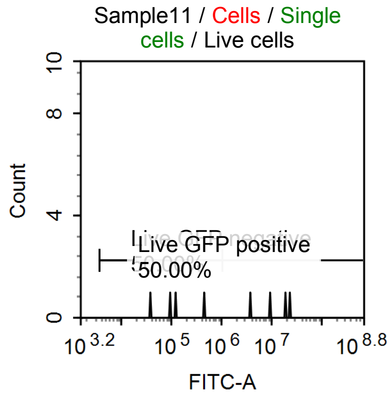


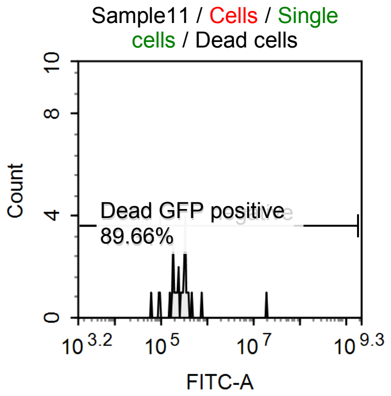


Sample Statistics of Sample11

| Gate | Count | % Parent | X | Y | Median X | Median Y |
|---|---|---|---|---|---|---|
| All | 446 | | | | | |
| Cells | 62 | 13.90 % | FSC-H | SSC-H | 1,840,470 | 2,707,318 |
| Single cells | 37 | 59.68 % | FSC-H | FSC-A | 1,872,078 | 5,574,281 |
| Live cells | 8 | 21.62 % | APC-Cy7-H | FSC-H | 9,875 | 5,757,621 |
| Live GFP negative | 4 | 50.00 % | FITC-A | Count | 110,584 | |
| Live GFP positive | 4 | 50.00 % | FITC-A | Count | 14,384,328 | |
| Dead cells | 29 | 78.38 % | APC-Cy7-H | FSC-H | 93,432 | 1,857,321 |
| Dead GFP negative | 3 | 10.34 % | FITC-A | Count | 91,673 | |
| Dead GFP positive | 26 | 89.66 % | FITC-A | Count | 291,482 | |

#### Specimen3

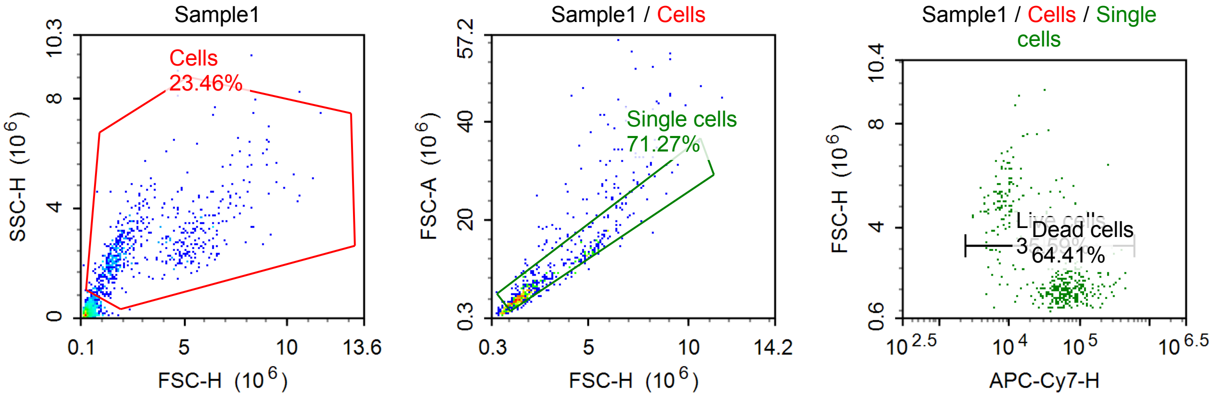


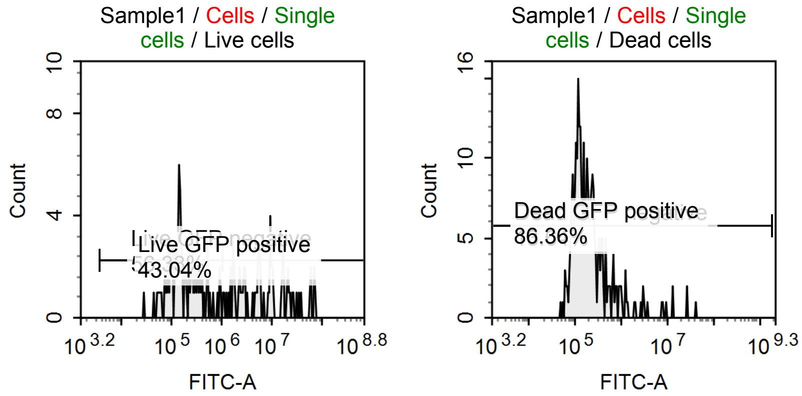


Sample Statistics of Sample1

| Gate | Count | % Parent | X | Y | Median X | Median Y |
|---|---|---|---|---|---|---|
| All | 2,656 | | | | | |
| Cells | 623 | 23.46 % | FSC-H | SSC-H | 2,220,892 | 2,474,678 |
| Single cells | 444 | 71.27 % | FSC-H | FSC-A | 1,998,745 | 6,094,491 |
| Live cells | 158 | 35.59 % | APC-Cy7-H | FSC-H | 9,375 | 4,757,532 |
| Live GFP negative | 89 | 56.33 % | FITC-A | Count | 157,419 | |
| Live GFP positive | 68 | 43.04 % | FITC-A | Count | 6,767,969 | |
| Dead cells | 286 | 64.41 % | APC-Cy7-H | FSC-H | 66,855 | 1,700,242 |
| Dead GFP negative | 39 | 13.64 % | FITC-A | Count | 84,086 | |
| Dead GFP positive | 247 | 86.36 % | FITC-A | Count | 197,976 | |

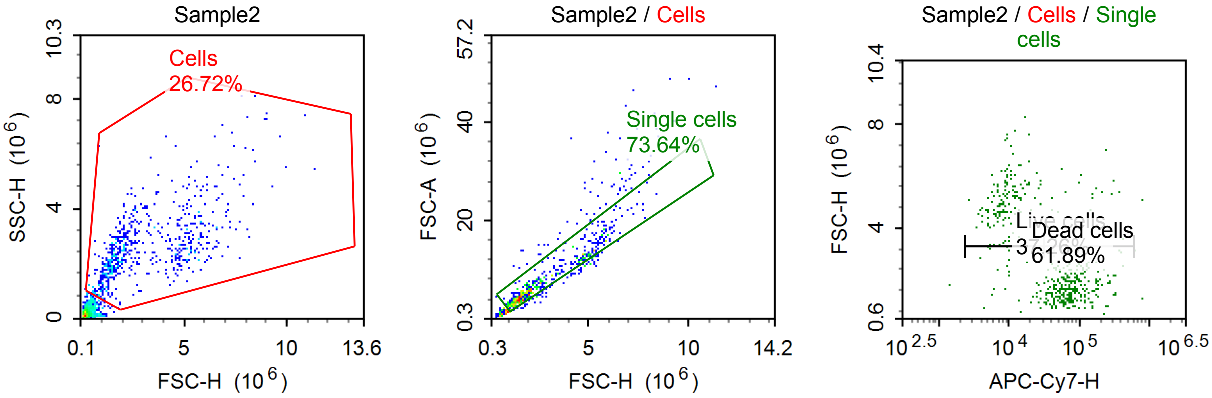

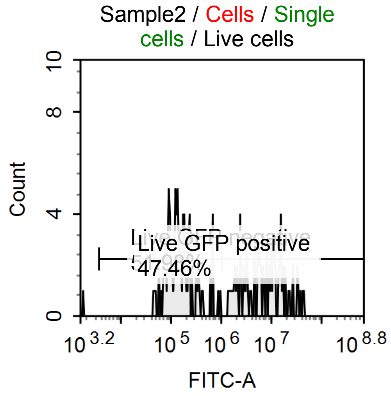


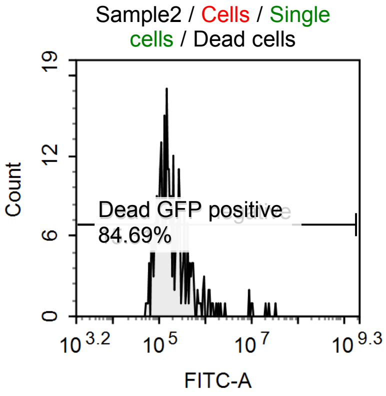


Sample Statistics of Sample2

| Gate | Count | % Parent | X | Y | Median X | Median Y |
|---|---|---|---|---|---|---|
| All | 2,414 | | | | | |
| Cells | 645 | 26.72 % | FSC-H | SSC-H | 2,326,261 | 2,454,853 |
| Single cells | 475 | 73.64 % | FSC-H | FSC-A | 2,136,471 | 6,481,144 |
| Live cells | 177 | 37.26 % | APC-Cy7-H | FSC-H | 9,149 | 4,790,506 |
| Live GFP negative | 92 | 51.98 % | FITC-A | Count | 139,110 | |
| Live GFP positive | 84 | 47.46 % | FITC-A | Count | 7,462,441 | |
| Dead cells | 294 | 61.89 % | APC-Cy7-H | FSC-H | 73,038 | 1,738,345 |
| Dead GFP negative | 45 | 15.31 % | FITC-A | Count | 80,153 | |
| Dead GFP positive | 249 | 84.69 % | FITC-A | Count | 186,905 | |

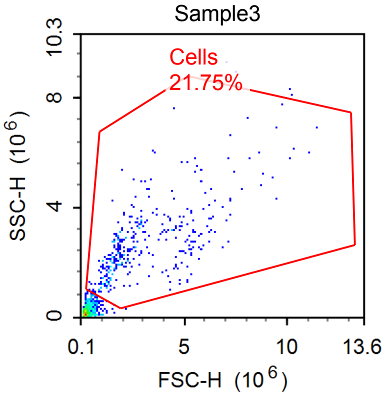


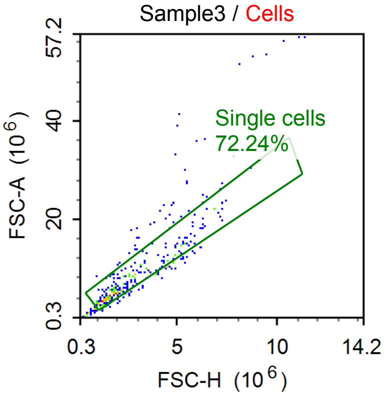


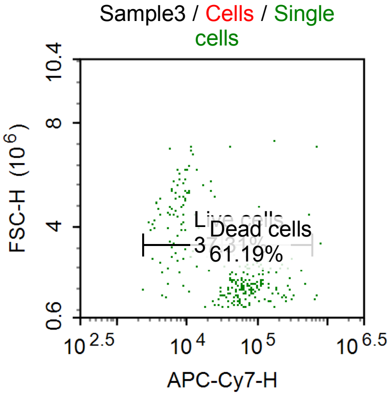


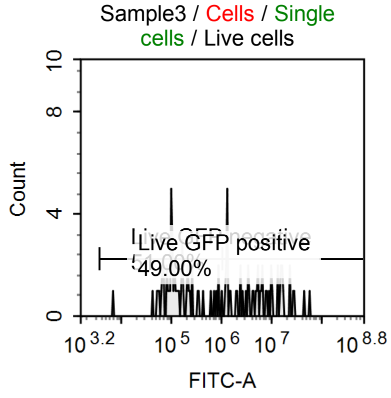


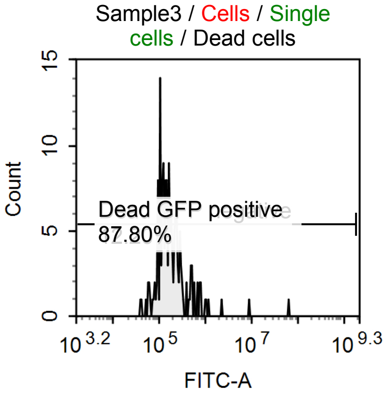

Sample Statistics of Sample3

| Gate | Count | % Parent | X | Y | Median X | Median Y |
|---|---|---|---|---|---|---|
| All | 1,706 | | | | | |
| Cells | 371 | 21.75 % | FSC-H | SSC-H | 2,275,170 | 2,466,035 |
| Single cells | 268 | 72.24 % | FSC-H | FSC-A | 2,063,817 | 5,940,171 |
| Live cells | 100 | 37.31 % | APC-Cy7-H | FSC-H | 8,559 | 4,396,788 |
| Live GFP negative | 51 | 51.00 % | FITC-A | Count | 121,555 | |
| Live GFP positive | 49 | 49.00 % | FITC-A | Count | 5,623,352 | |
| Dead cells | 164 | 61.19 % | APC-Cy7-H | FSC-H | 73,641 | 1,783,696 |
| Dead GFP negative | 20 | 12.20 % | FITC-A | Count | 76,867 | |
| Dead GFP positive | 144 | 87.80 % | FITC-A | Count | 175,368 | |

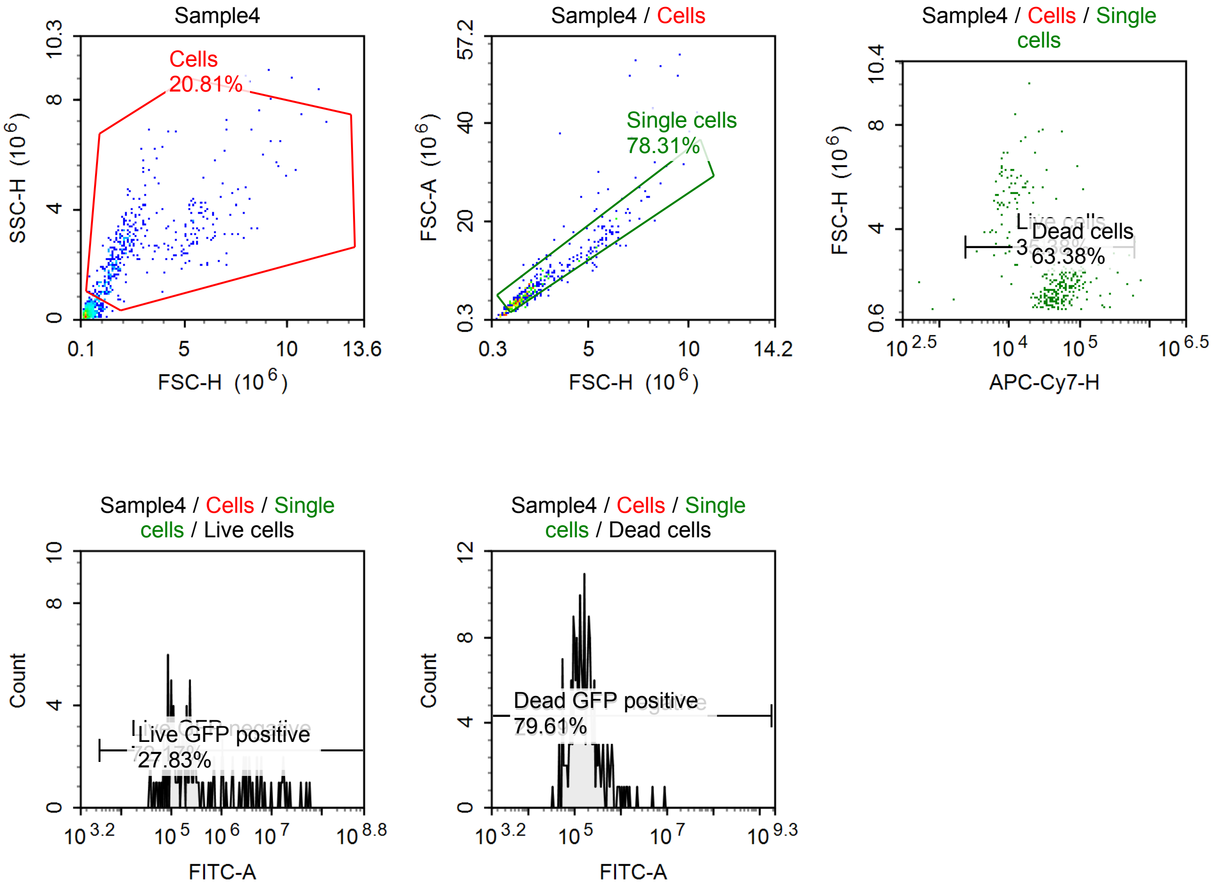


Sample Statistics of Sample4

| Gate | Count | % Parent | X | Y | Median X | Median Y |
|---|---|---|---|---|---|---|
| All | 1,994 | | | | | |
| Cells | 415 | 20.81 % | FSC-H | SSC-H | 2,075,510 | 2,585,179 |
| Single cells | 325 | 78.31 % | FSC-H | FSC-A | 2,067,594 | 6,045,700 |
| Live cells | 115 | 35.38 % | APC-Cy7-H | FSC-H | 12,772 | 4,858,738 |
| Live GFP negative | 83 | 72.17 % | FITC-A | Count | 166,974 | |
| Live GFP positive | 32 | 27.83 % | FITC-A | Count | 5,616,069 | |
| Dead cells | 206 | 63.38 % | APC-Cy7-H | FSC-H | 58,679 | 1,826,595 |
| Dead GFP negative | 42 | 20.39 % | FITC-A | Count | 76,197 | |
| Dead GFP positive | 164 | 79.61 % | FITC-A | Count | 190,842 | |

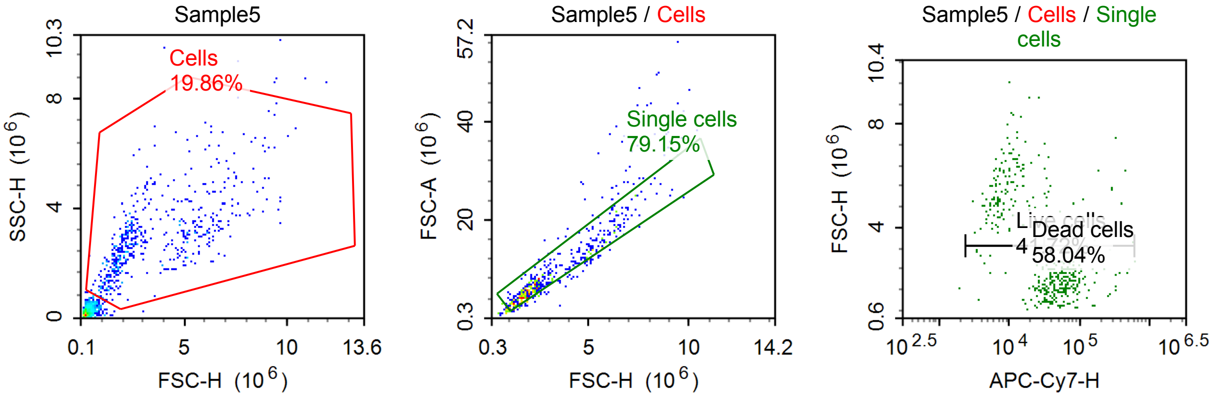


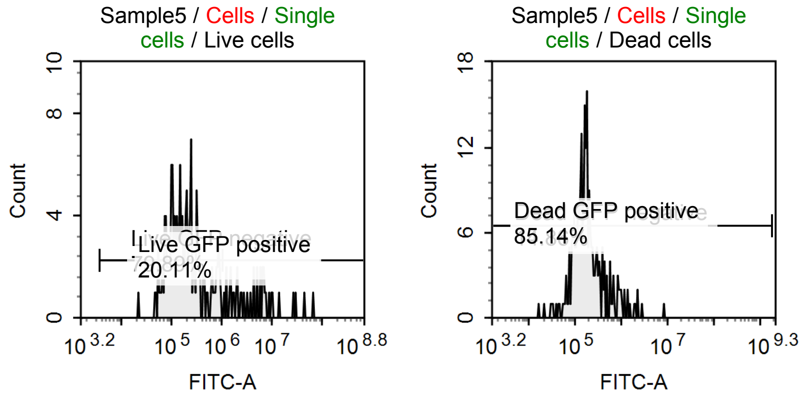


Sample Statistics of Sample5

| Gate | Count | % Parent | X | Y | Median X | Median Y |
|---|---|---|---|---|---|---|
| All | 2,729 | | | | | |
| Cells | 542 | 19.86 % | FSC-H | SSC-H | 2,561,286 | 2,792,714 |
| Single cells | 429 | 79.15 % | FSC-H | FSC-A | 2,401,553 | 7,203,450 |
| Live cells | 179 | 41.72 % | APC-Cy7-H | FSC-H | 10,075 | 4,947,316 |
| Live GFP negative | 143 | 79.89 % | FITC-A | Count | 169,289 | |
| Live GFP positive | 36 | 20.11 % | FITC-A | Count | 4,921,441 | |
| Dead cells | 249 | 58.04 % | APC-Cy7-H | FSC-H | 62,853 | 2,081,255 |
| Dead GFP negative | 37 | 14.86 % | FITC-A | Count | 75,777 | |
| Dead GFP positive | 212 | 85.14 % | FITC-A | Count | 185,423 | |

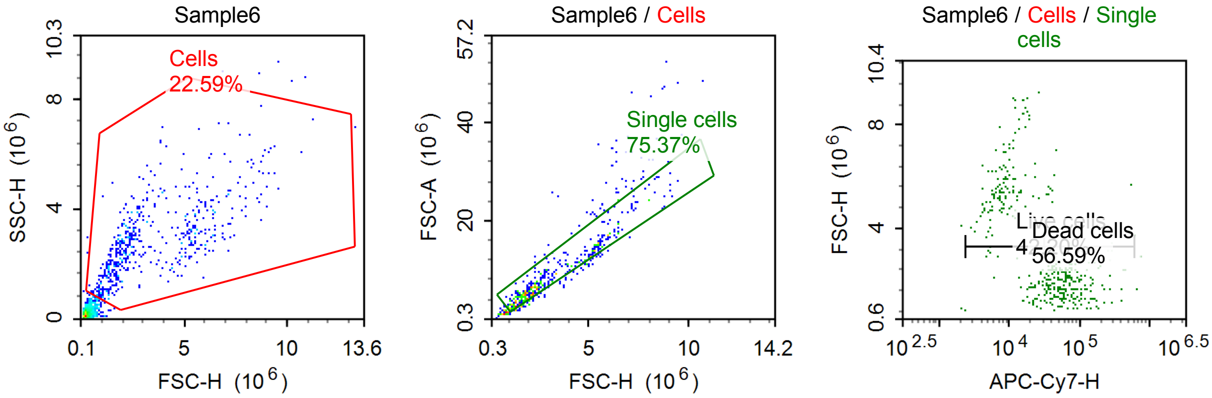

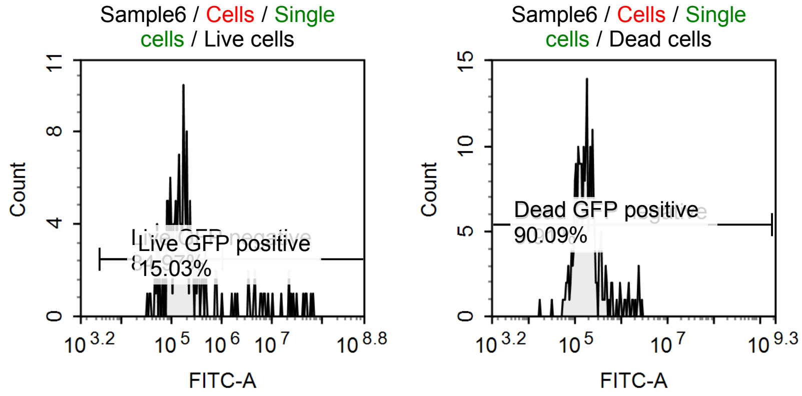


Sample Statistics of Sample6

| Gate | Count | % Parent | X | Y | Median X | Median Y |
|---|---|---|---|---|---|---|
| All | 2,408 | | | | | |
| Cells | 544 | 22.59 % | FSC-H | SSC-H | 2,530,787 | 2,757,219 |
| Single cells | 410 | 75.37 % | FSC-H | FSC-A | 2,402,534 | 6,959,423 |
| Live cells | 173 | 42.20 % | APC-Cy7-H | FSC-H | 9,766 | 5,101,193 |
| Live GFP negative | 147 | 84.97 % | FITC-A | Count | 160,797 | |
| Live GFP positive | 26 | 15.03 % | FITC-A | Count | 8,205,477 | |
| Dead cells | 232 | 56.59 % | APC-Cy7-H | FSC-H | 62,851 | 2,058,191 |
| Dead GFP negative | 23 | 9.91 % | FITC-A | Count | 74,377 | |
| Dead GFP positive | 209 | 90.09 % | FITC-A | Count | 184,396 | |

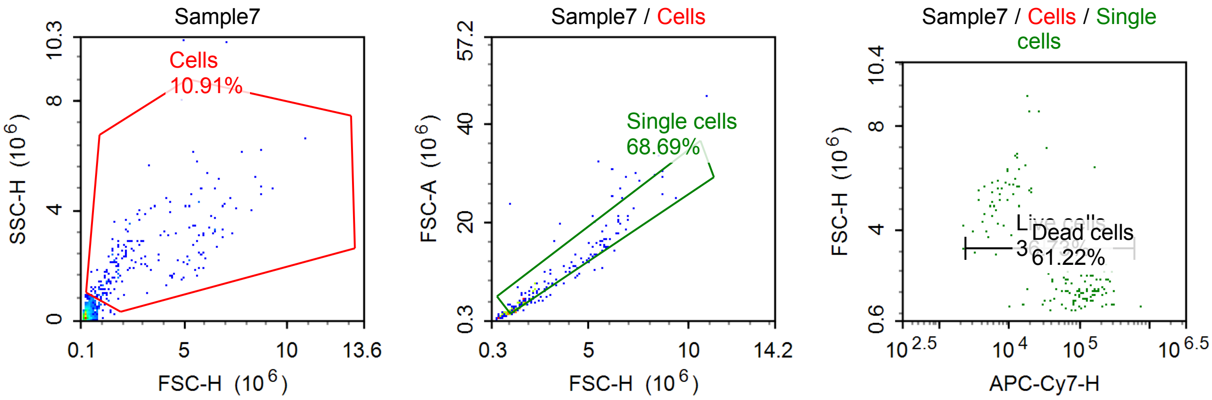


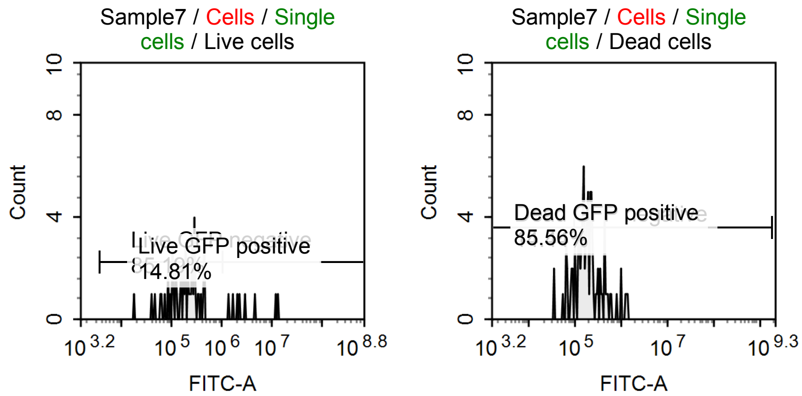

Sample Statistics of Sample7

| Gate | Count | % Parent | X | Y | Median X | Median Y |
|---|---|---|---|---|---|---|
| All | 1,961 | | | | | |
| Cells | 214 | 10.91 % | FSC-H | SSC-H | 2,137,019 | 2,161,369 |
| Single cells | 147 | 68.69 % | FSC-H | FSC-A | 2,157,472 | 6,120,506 |
| Live cells | 54 | 36.73 % | APC-Cy7-H | FSC-H | 8,990 | 5,448,123 |
| Live GFP negative | 46 | 85.19 % | FITC-A | Count | 194,985 | |
| Live GFP positive | 8 | 14.81 % | FITC-A | Count | 2,708,120 | |
| Dead cells | 90 | 61.22 % | APC-Cy7-H | FSC-H | 107,281 | 1,730,000 |
| Dead GFP negative | 13 | 14.44 % | FITC-A | Count | 67,582 | |
| Dead GFP positive | 77 | 85.56 % | FITC-A | Count | 200,166 | |

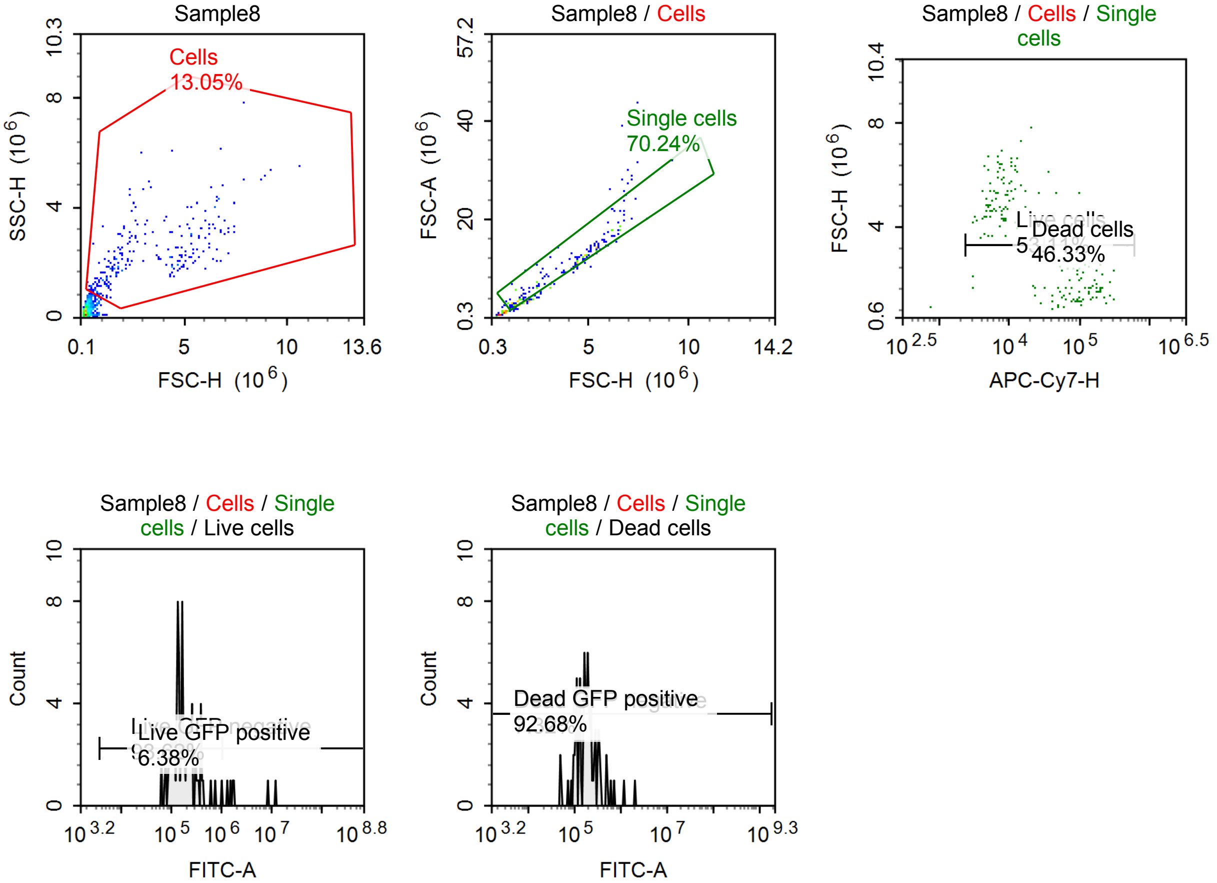


Sample Statistics of Sample8

| Gate | Count | % Parent | X | Y | Median X | Median Y |
|---|---|---|---|---|---|---|
| All | 1,931 | | | | | |
| Cells | 252 | 13.05 % | FSC-H | SSC-H | 2,519,959 | 2,123,730 |
| Single cells | 177 | 70.24 % | FSC-H | FSC-A | 3,680,294 | 10,064,164 |
| Live cells | 94 | 53.11 % | APC-Cy7-H | FSC-H | 7,484 | 4,974,181 |
| Live GFP negative | 88 | 93.62 % | FITC-A | Count | 167,191 | |
| Live GFP positive | 6 | 6.38 % | FITC-A | Count | 1,788,131 | |
| Dead cells | 82 | 46.33 % | APC-Cy7-H | FSC-H | 98,701 | 1,646,040 |
| Dead GFP negative | 6 | 7.32 % | FITC-A | Count | 61,593 | |
| Dead GFP positive | 76 | 92.68 % | FITC-A | Count | 191,637 | |

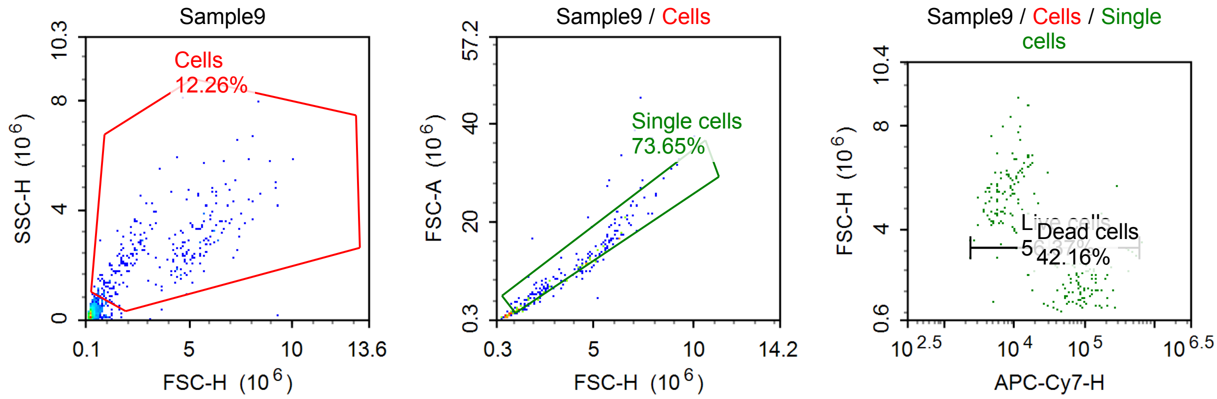


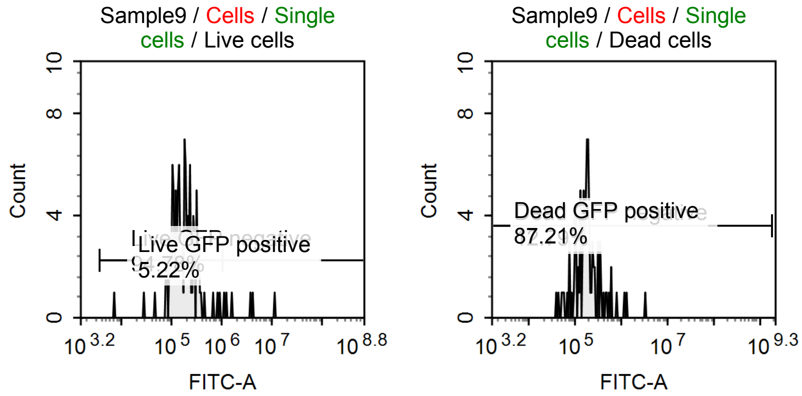


Sample Statistics of Sample9

| Gate | Count | % Parent | X | Y | Median X | Median Y |
|---|---|---|---|---|---|---|
| All | 2,260 | | | | | |
| Cells | 277 | 12.26 % | FSC-H | SSC-H | 3,327,024 | 2,405,044 |
| Single cells | 204 | 73.65 % | FSC-H | FSC-A | 4,148,758 | 10,520,344 |
| Live cells | 115 | 56.37 % | APC-Cy7-H | FSC-H | 7,879 | 5,322,124 |
| Live GFP negative | 109 | 94.78 % | FITC-A | Count | 178,879 | |
| Live GFP positive | 6 | 5.22 % | FITC-A | Count | 2,785,782 | |
| Dead cells | 86 | 42.16 % | APC-Cy7-H | FSC-H | 92,138 | 1,883,958 |
| Dead GFP negative | 11 | 12.79 % | FITC-A | Count | 69,514 | |
| Dead GFP positive | 75 | 87.21 % | FITC-A | Count | 188,760 | |

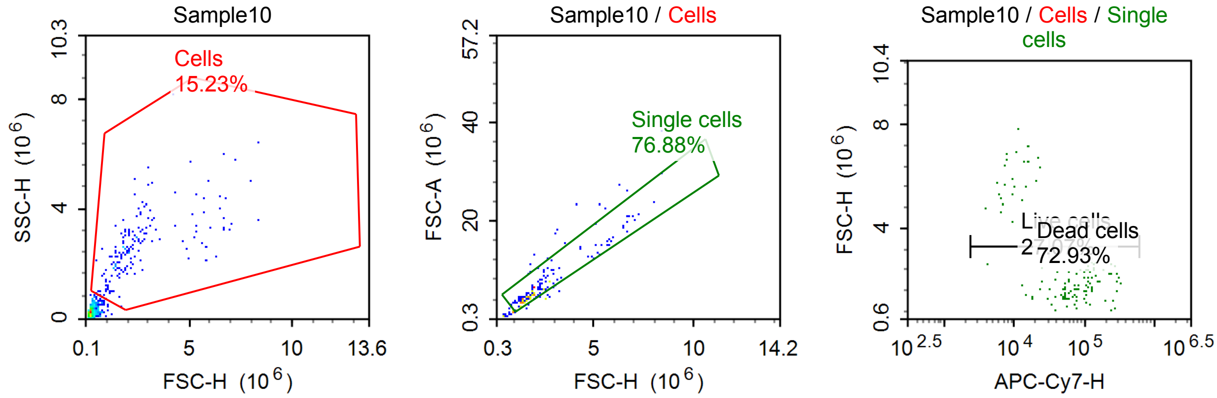

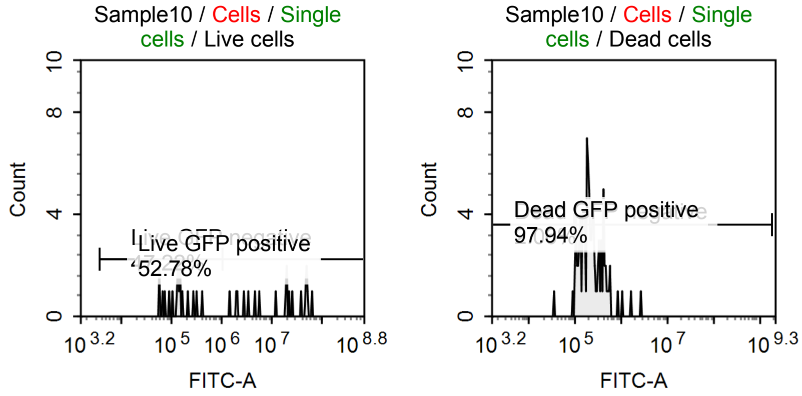


Sample Statistics of Sample10

| Gate | Count | % Parent | X | Y | Median X | Median Y |
|---|---|---|---|---|---|---|
| All | 1,136 | | | | | |
| Cells | 173 | 15.23 % | FSC-H | SSC-H | 2,004,757 | 2,591,084 |
| Single cells | 133 | 76.88 % | FSC-H | FSC-A | 2,020,179 | 5,850,266 |
| Live cells | 36 | 27.07 % | APC-Cy7-H | FSC-H | 12,800 | 5,250,364 |
| Live GFP negative | 17 | 47.22 % | FITC-A | Count | 141,475 | |
| Live GFP positive | 19 | 52.78 % | FITC-A | Count | 20,326,862 | |
| Dead cells | 97 | 72.93 % | APC-Cy7-H | FSC-H | 100,435 | 1,884,425 |
| Dead GFP negative | 2 | 2.06 % | FITC-A | Count | 63,981 | |
| Dead GFP positive | 95 | 97.94 % | FITC-A | Count | 203,555 | |

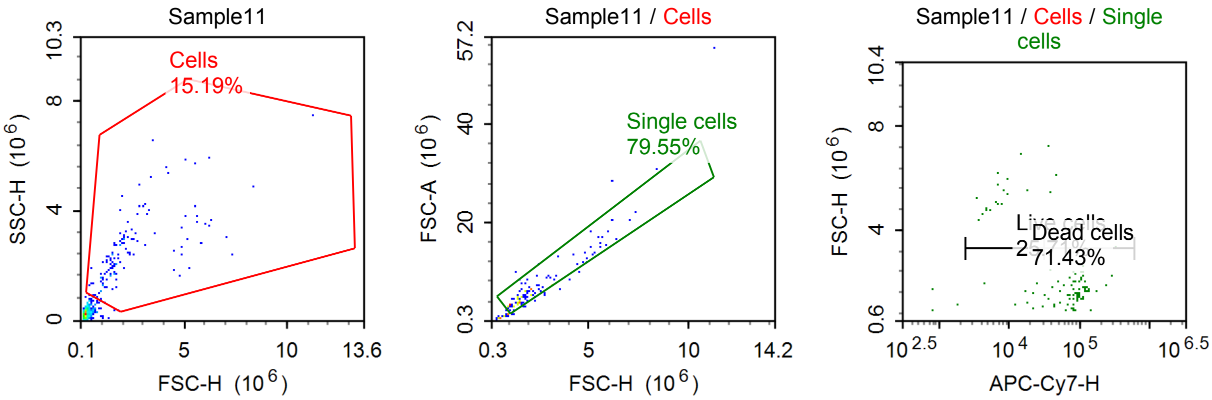


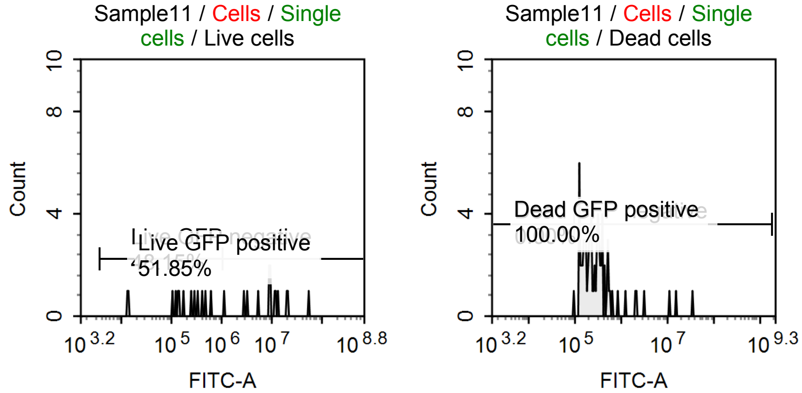

Sample Statistics of Sample11

| Gate | Count | % Parent | X | Y | Median X | Median Y |
|---|---|---|---|---|---|---|
| All | 869 | | | | | |
| Cells | 132 | 15.19 % | FSC-H | SSC-H | 1,763,394 | 2,363,787 |
| Single cells | 105 | 79.55 % | FSC-H | FSC-A | 1,846,499 | 5,612,916 |
| Live cells | 27 | 25.71 % | APC-Cy7-H | FSC-H | 8,878 | 4,819,248 |
| Live GFP negative | 13 | 48.15 % | FITC-A | Count | 179,481 | |
| Live GFP positive | 14 | 51.85 % | FITC-A | Count | 9,777,584 | |
| Dead cells | 75 | 71.43 % | APC-Cy7-H | FSC-H | 89,015 | 1,779,826 |
| Dead GFP negative | 0 | 0.00 % | FITC-A | Count | 0 | |
| Dead GFP positive | 75 | 100.00 % | FITC-A | Count | 268,489 | |

#### Specimen4

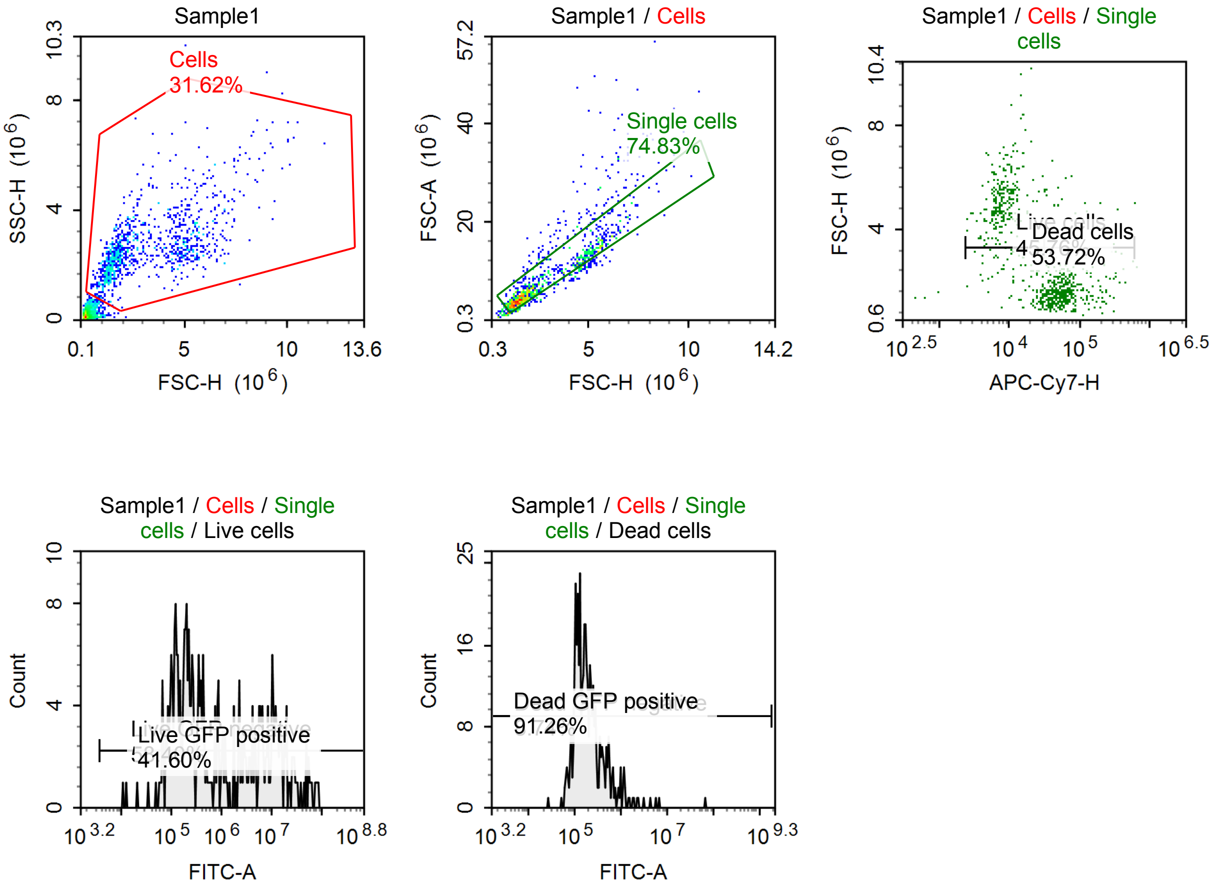

Sample Statistics of Sample1

| Gate | Count | % Parent | X | Y | Median X | Median Y |
|---|---|---|---|---|---|---|
| All | 3,242 | | | | | |
| Cells | 1,025 | 31.62 % | FSC-H | SSC-H | 2,466,165 | 2,487,489 |
| Single cells | 767 | 74.83 % | FSC-H | FSC-A | 2,064,877 | 6,280,777 |
| Live cells | 351 | 45.76 % | APC-Cy7-H | FSC-H | 9,158 | 4,675,378 |
| Live GFP negative | 205 | 58.40 % | FITC-A | Count | 202,245 | |
| Live GFP positive | 146 | 41.60 % | FITC-A | Count | 8,594,970 | |
| Dead cells | 412 | 53.72 % | APC-Cy7-H | FSC-H | 54,970 | 1,589,620 |
| Dead GFP negative | 36 | 8.74 % | FITC-A | Count | 81,602 | |
| Dead GFP positive | 376 | 91.26 % | FITC-A | Count | 182,888 | |

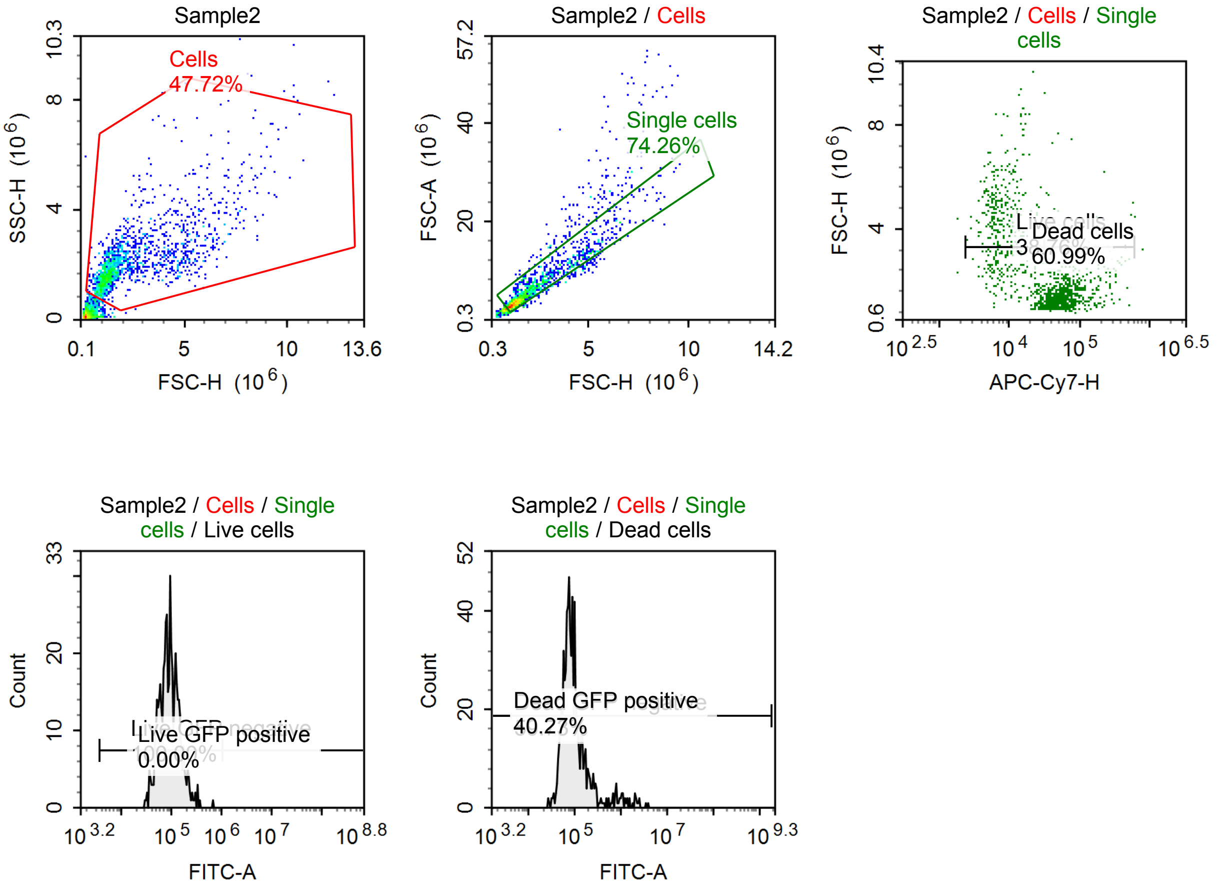


Sample Statistics of Sample2

| Gate | Count | % Parent | X | Y | Median X | Median Y |
|---|---|---|---|---|---|---|
| All | 3,378 | | | | | |
| Cells | 1,612 | 47.72 % | FSC-H | SSC-H | 1,677,340 | 1,943,343 |
| Single cells | 1,197 | 74.26 % | FSC-H | FSC-A | 1,652,904 | 4,903,202 |
| Live cells | 464 | 38.76 % | APC-Cy7-H | FSC-H | 9,215 | 3,508,719 |
| Live GFP negative | 464 | 100.00 % | FITC-A | Count | 93,345 | |
| Live GFP positive | 0 | 0.00 % | FITC-A | Count | 0 | |
| Dead cells | 730 | 60.99 % | APC-Cy7-H | FSC-H | 57,161 | 1,366,095 |
| Dead GFP negative | 436 | 59.73 % | FITC-A | Count | 69,942 | |
| Dead GFP positive | 294 | 40.27 % | FITC-A | Count | 147,354 | |

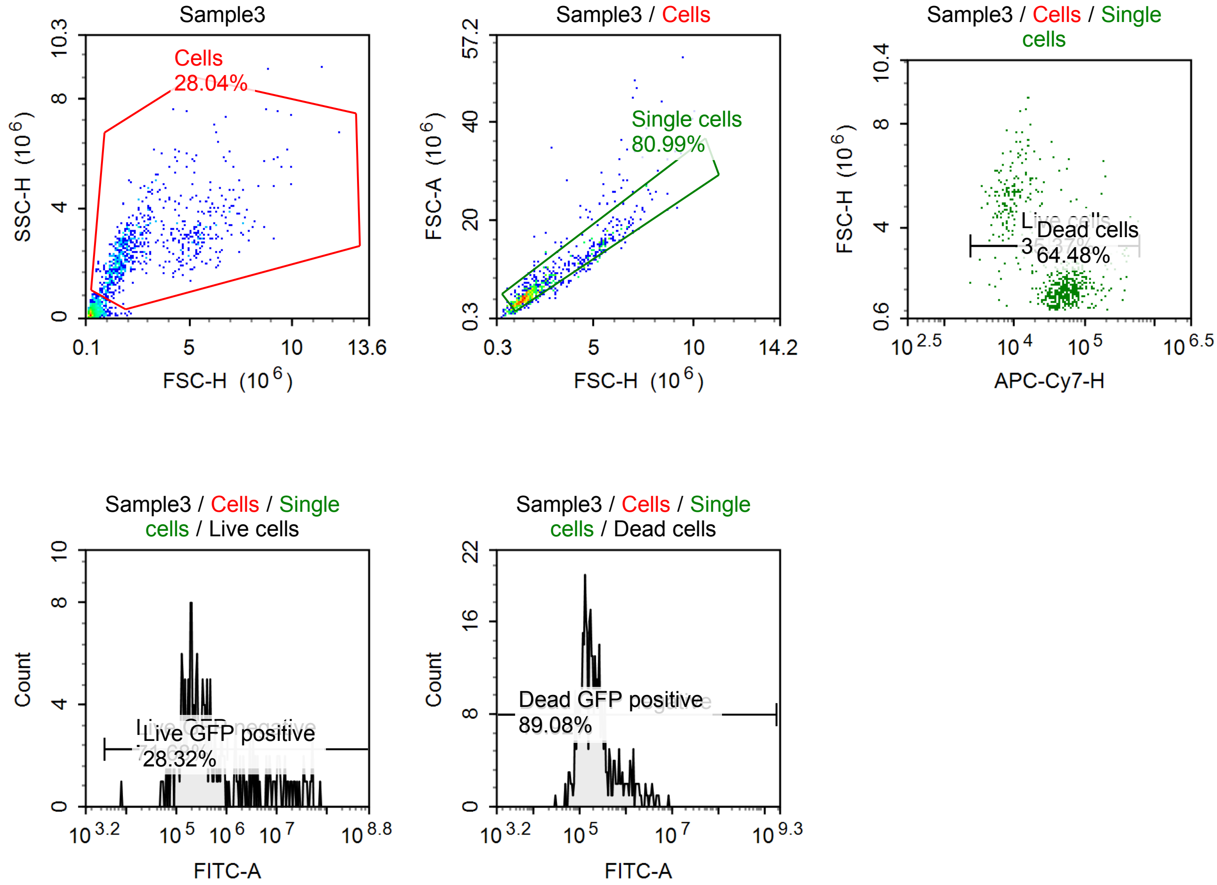


Sample Statistics of Sample3

| Gate | Count | % Parent | X | Y | Median X | Median Y |
|---|---|---|---|---|---|---|
| All | 2,814 | | | | | |
| Cells | 789 | 28.04 % | FSC-H | SSC-H | 2,092,670 | 2,562,636 |
| Single cells | 639 | 80.99 % | FSC-H | FSC-A | 2,004,532 | 5,970,146 |
| Live cells | 226 | 35.37 % | APC-Cy7-H | FSC-H | 10,368 | 4,704,684 |
| Live GFP negative | 162 | 71.68 % | FITC-A | Count | 220,459 | |
| Live GFP positive | 64 | 28.32 % | FITC-A | Count | 8,205,483 | |
| Dead cells | 412 | 64.48 % | APC-Cy7-H | FSC-H | 60,462 | 1,702,651 |
| Dead GFP negative | 45 | 10.92 % | FITC-A | Count | 82,055 | |
| Dead GFP positive | 367 | 89.08 % | FITC-A | Count | 214,227 | |

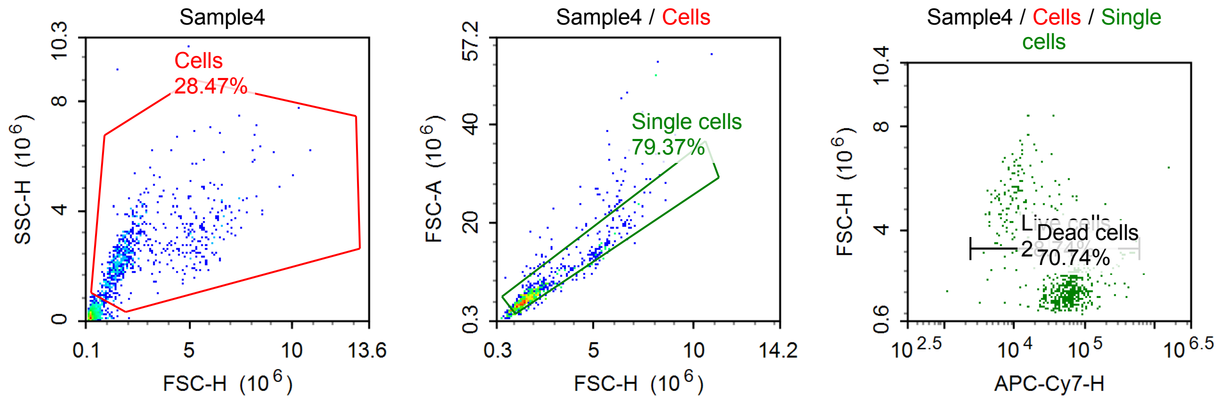

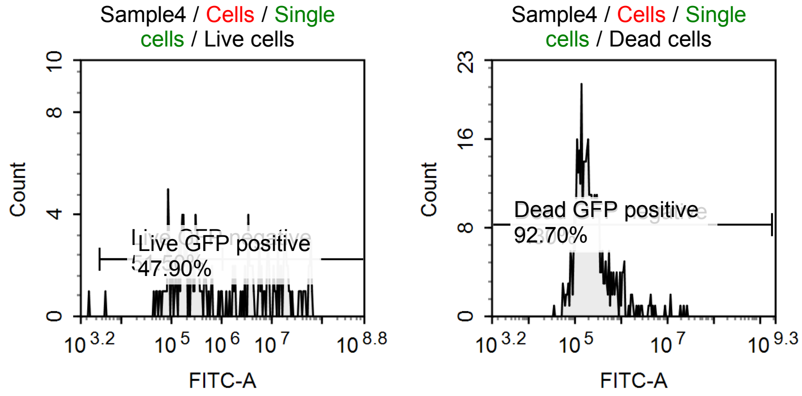


Sample Statistics of Sample4

| Gate | Count | % Parent | X | Y | Median X | Median Y |
|---|---|---|---|---|---|---|
| All | 2,571 | | | | | |
| Cells | 732 | 28.47 % | FSC-H | SSC-H | 1,963,345 | 2,451,842 |
| Single cells | 581 | 79.37 % | FSC-H | FSC-A | 1,907,774 | 5,562,778 |
| Live cells | 167 | 28.74 % | APC-Cy7-H | FSC-H | 11,176 | 4,777,986 |
| Live GFP negative | 86 | 51.50 % | FITC-A | Count | 188,798 | |
| Live GFP positive | 80 | 47.90 % | FITC-A | Count | 10,243,146 | |
| Dead cells | 411 | 70.74 % | APC-Cy7-H | FSC-H | 62,838 | 1,693,181 |
| Dead GFP negative | 30 | 7.30 % | FITC-A | Count | 76,691 | |
| Dead GFP positive | 381 | 92.70 % | FITC-A | Count | 206,875 | |

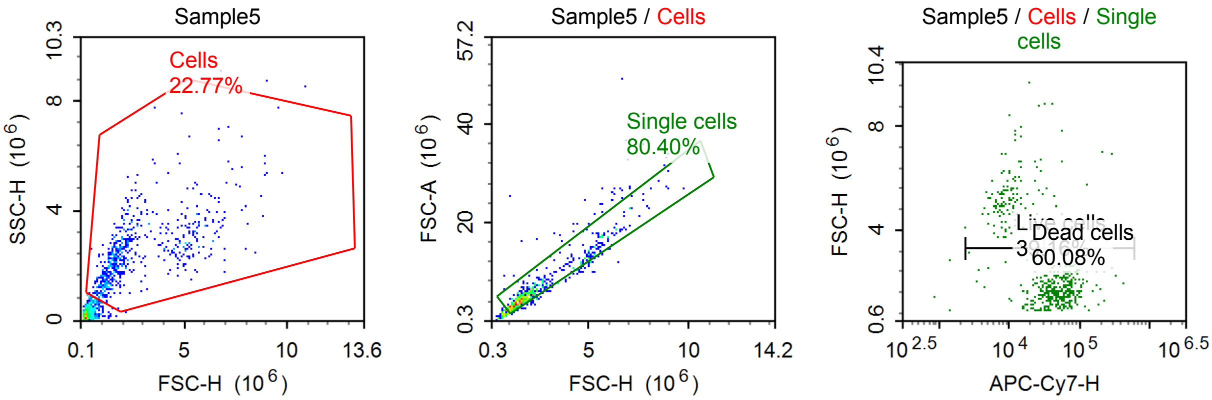


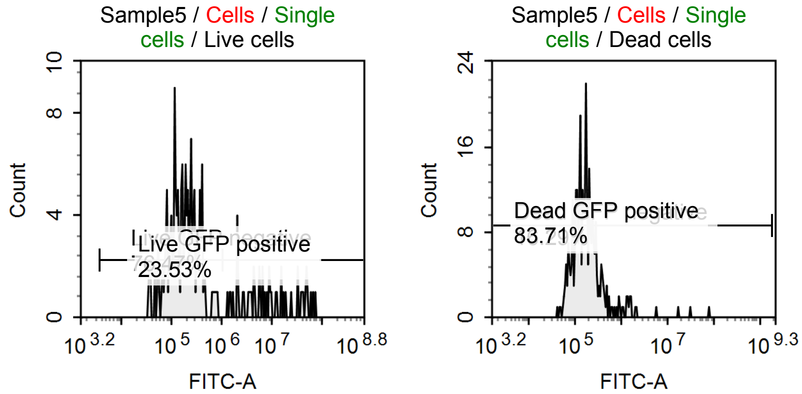

Sample Statistics of Sample5

| Gate | Count | % Parent | X | Y | Median X | Median Y |
|---|---|---|---|---|---|---|
| All | 2,846 | | | | | |
| Cells | 648 | 22.77 % | FSC-H | SSC-H | 1,918,109 | 2,380,604 |
| Single cells | 521 | 80.40 % | FSC-H | FSC-A | 1,949,735 | 5,534,485 |
| Live cells | 204 | 39.16 % | APC-Cy7-H | FSC-H | 13,188 | 4,575,045 |
| Live GFP negative | 156 | 76.47 % | FITC-A | Count | 174,758 | |
| Live GFP positive | 48 | 23.53 % | FITC-A | Count | 7,913,912 | |
| Dead cells | 313 | 60.08 % | APC-Cy7-H | FSC-H | 55,428 | 1,690,816 |
| Dead GFP negative | 51 | 16.29 % | FITC-A | Count | 75,217 | |
| Dead GFP positive | 262 | 83.71 % | FITC-A | Count | 182,743 | |

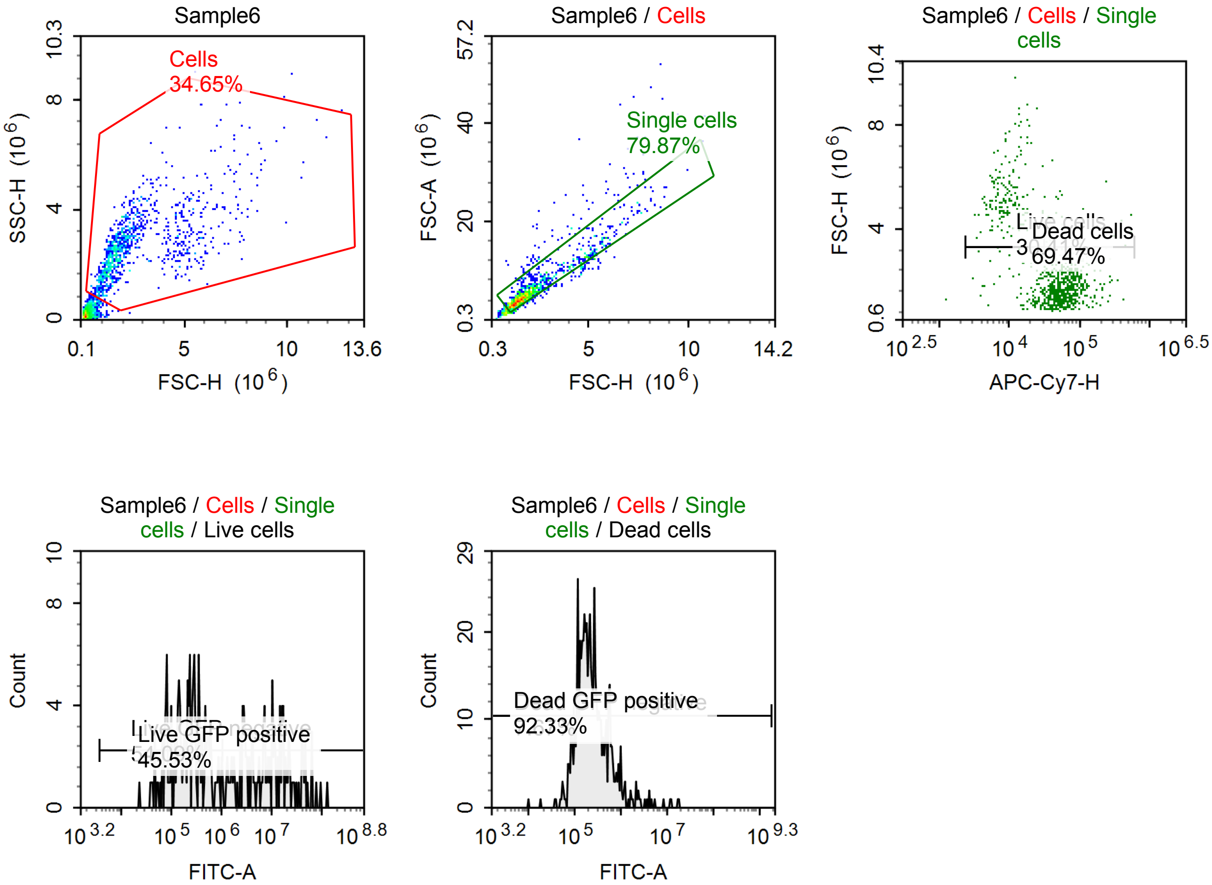


Sample Statistics of Sample6

| Gate | Count | % Parent | X | Y | Median X | Median Y |
|---|---|---|---|---|---|---|
| All | 3,053 | | | | | |
| Cells | 1,058 | 34.65 % | FSC-H | SSC-H | 1,899,722 | 2,574,781 |
| Single cells | 845 | 79.87 % | FSC-H | FSC-A | 1,870,980 | 5,455,655 |
| Live cells | 257 | 30.41 % | APC-Cy7-H | FSC-H | 11,955 | 4,610,942 |
| Live GFP negative | 139 | 54.09 % | FITC-A | Count | 225,727 | |
| Live GFP positive | 117 | 45.53 % | FITC-A | Count | 9,855,760 | |
| Dead cells | 587 | 69.47 % | APC-Cy7-H | FSC-H | 61,166 | 1,692,721 |
| Dead GFP negative | 45 | 7.67 % | FITC-A | Count | 78,657 | |
| Dead GFP positive | 542 | 92.33 % | FITC-A | Count | 233,214 | |

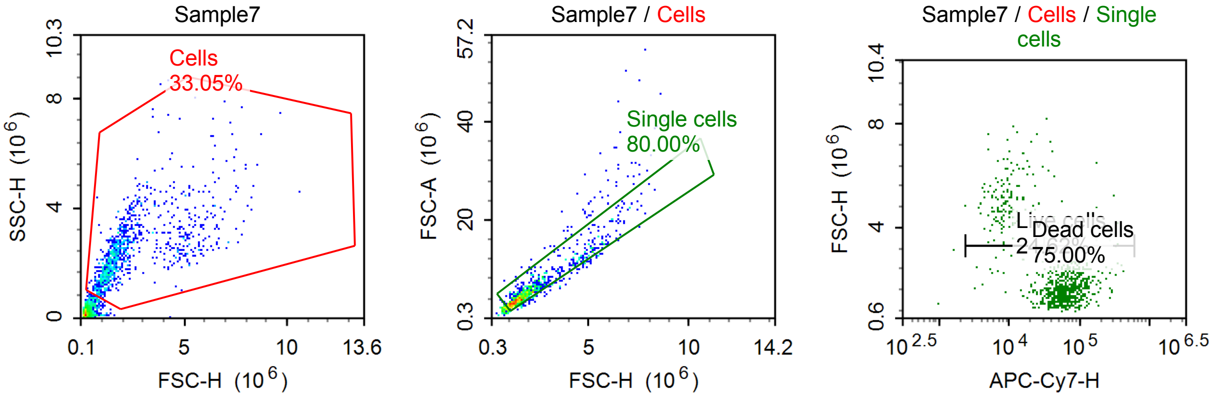


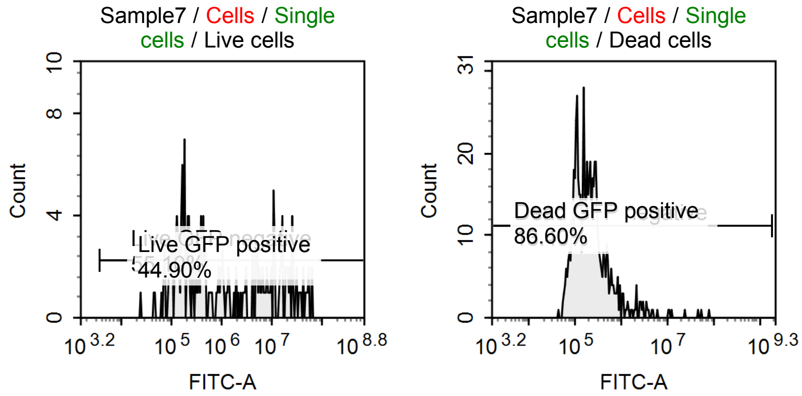


Sample Statistics of Sample7

| Gate | Count | % Parent | X | Y | Median X | Median Y |
|---|---|---|---|---|---|---|
| All | 3,011 | | | | | |
| Cells | 995 | 33.05 % | FSC-H | SSC-H | 1,751,757 | 2,333,581 |
| Single cells | 796 | 80.00 % | FSC-H | FSC-A | 1,764,505 | 5,091,690 |
| Live cells | 196 | 24.62 % | APC-Cy7-H | FSC-H | 10,227 | 4,668,530 |
| Live GFP negative | 108 | 55.10 % | FITC-A | Count | 189,551 | |
| Live GFP positive | 88 | 44.90 % | FITC-A | Count | 11,228,435 | |
| Dead cells | 597 | 75.00 % | APC-Cy7-H | FSC-H | 64,301 | 1,602,774 |
| Dead GFP negative | 80 | 13.40 % | FITC-A | Count | 80,371 | |
| Dead GFP positive | 517 | 86.60 % | FITC-A | Count | 215,995 | |

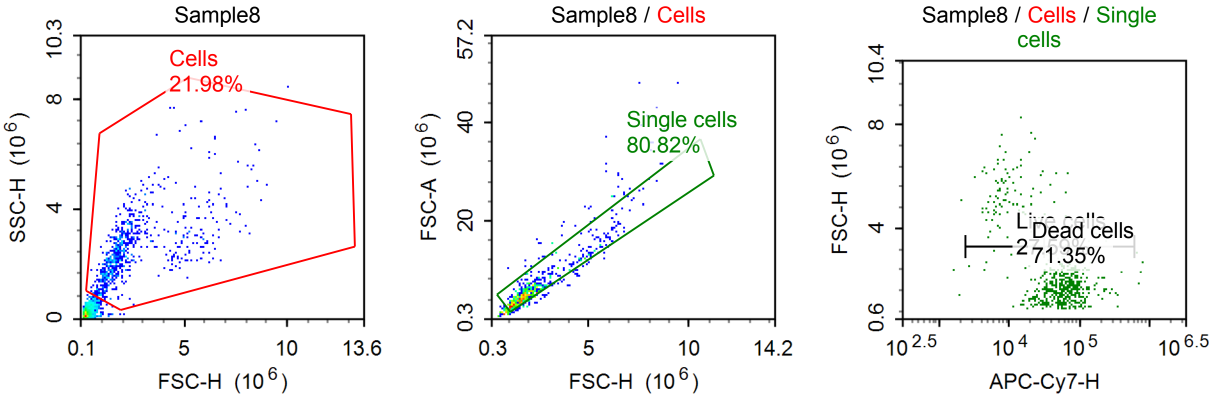

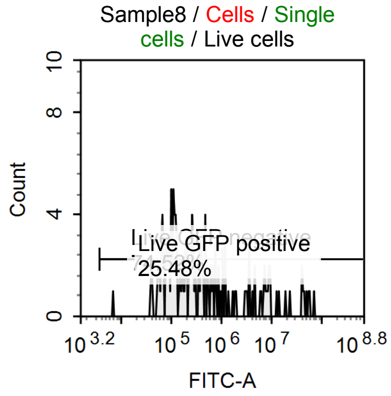


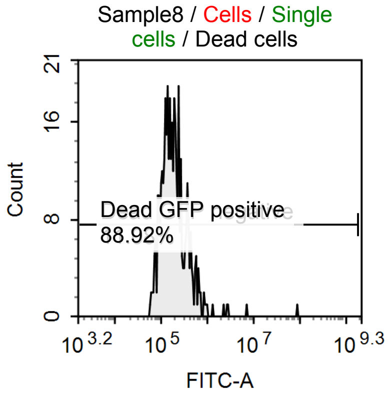


Sample Statistics of Sample8

| Gate | Count | % Parent | X | Y | Median X | Median Y |
|---|---|---|---|---|---|---|
| All | 3,203 | | | | | |
| Cells | 704 | 21.98 % | FSC-H | SSC-H | 1,897,840 | 2,423,931 |
| Single cells | 569 | 80.82 % | FSC-H | FSC-A | 1,888,138 | 5,347,501 |
| Live cells | 157 | 27.59 % | APC-Cy7-H | FSC-H | 15,393 | 4,151,115 |
| Live GFP negative | 117 | 74.52 % | FITC-A | Count | 159,182 | |
| Live GFP positive | 40 | 25.48 % | FITC-A | Count | 7,523,993 | |
| Dead cells | 406 | 71.35 % | APC-Cy7-H | FSC-H | 62,952 | 1,782,250 |
| Dead GFP negative | 45 | 11.08 % | FITC-A | Count | 83,797 | |
| Dead GFP positive | 361 | 88.92 % | FITC-A | Count | 192,133 | |

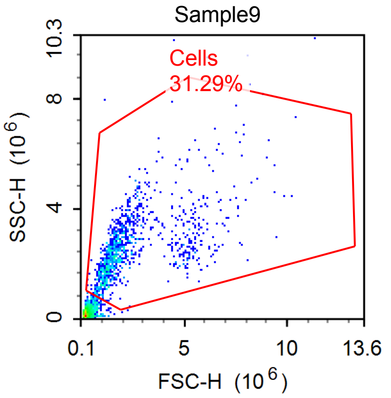


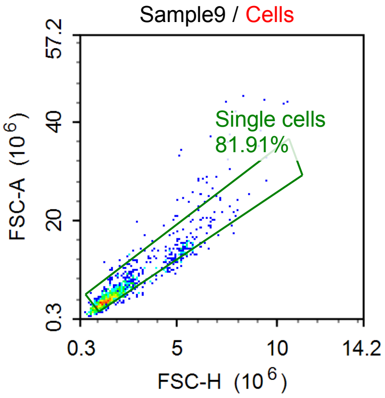


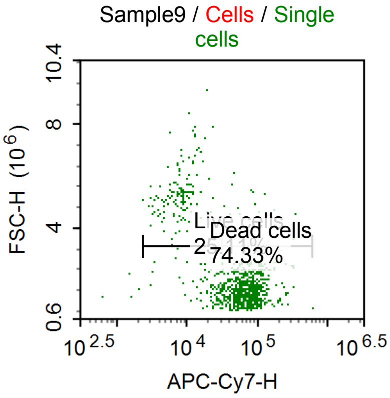


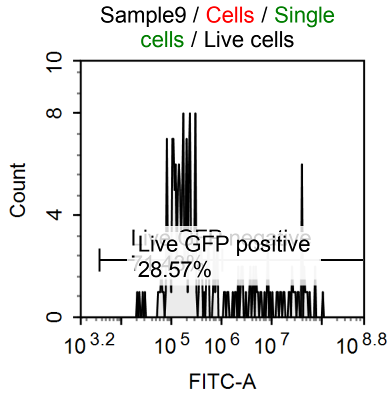


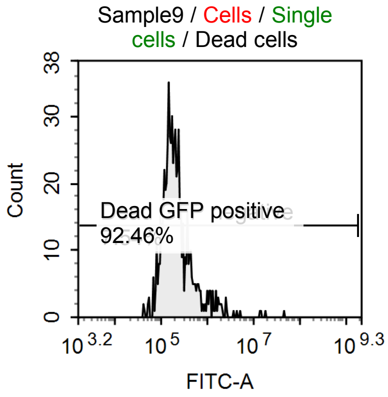

Sample Statistics of Sample9

| Gate | Count | % Parent | X | Y | Median X | Median Y |
|---|---|---|---|---|---|---|
| All | 3,480 | | | | | |
| Cells | 1,089 | 31.29 % | FSC-H | SSC-H | 1,736,916 | 2,435,550 |
| Single cells | 892 | 81.91 % | FSC-H | FSC-A | 1,761,917 | 4,900,651 |
| Live cells | 224 | 25.11 % | APC-Cy7-H | FSC-H | 11,044 | 4,594,960 |
| Live GFP negative | 160 | 71.43 % | FITC-A | Count | 177,055 | |
| Live GFP positive | 64 | 28.57 % | FITC-A | Count | 9,225,052 | |
| Dead cells | 663 | 74.33 % | APC-Cy7-H | FSC-H | 70,697 | 1,603,252 |
| Dead GFP negative | 50 | 7.54 % | FITC-A | Count | 81,662 | |
| Dead GFP positive | 613 | 92.46 % | FITC-A | Count | 203,899 | |

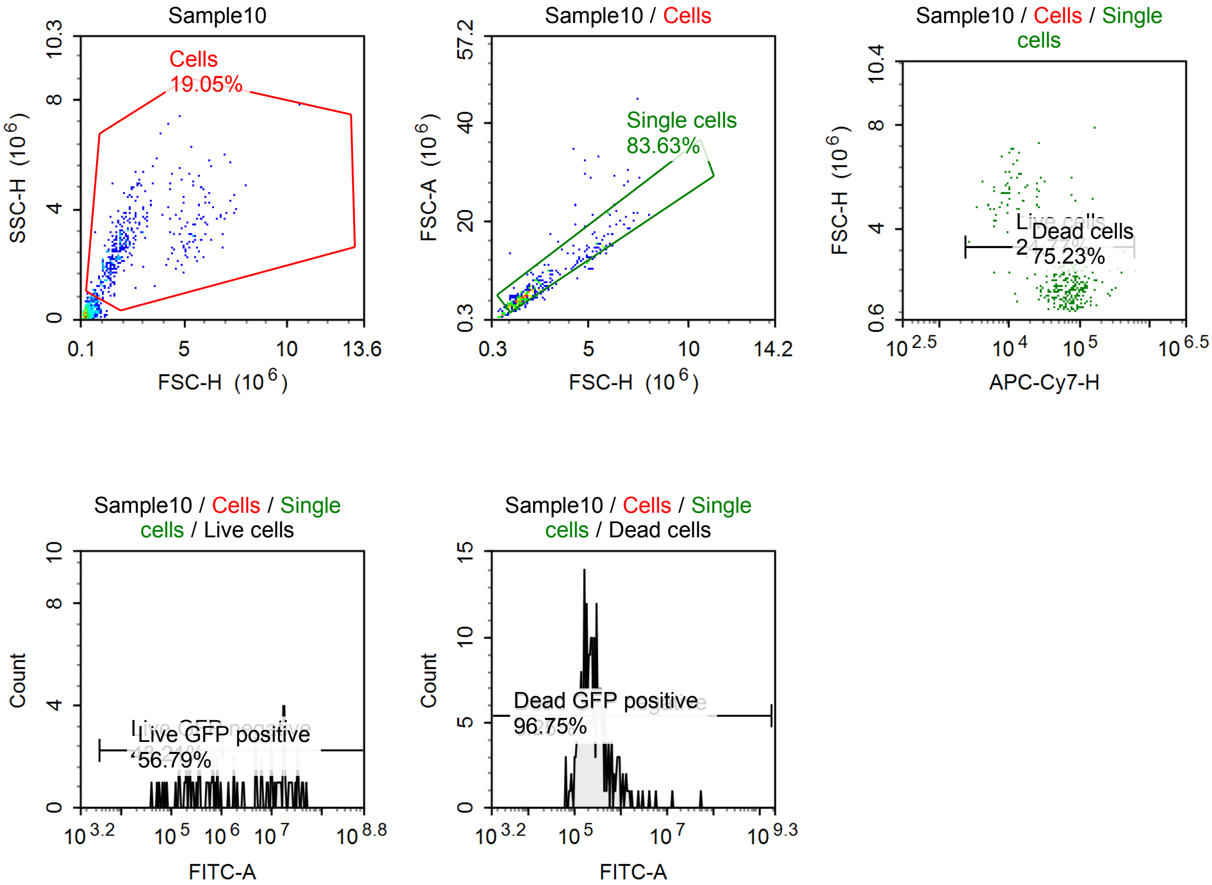


Sample Statistics of Sample10

| Gate | Count | % Parent | X | Y | Median X | Median Y |
|---|---|---|---|---|---|---|
| All | 2,052 | | | | | |
| Cells | 391 | 19.05 % | FSC-H | SSC-H | 1,967,405 | 2,680,200 |
| Single cells | 327 | 83.63 % | FSC-H | FSC-A | 1,967,405 | 5,371,684 |
| Live cells | 81 | 24.77 % | APC-Cy7-H | FSC-H | 12,371 | 4,989,566 |
| Live GFP negative | 35 | 43.21 % | FITC-A | Count | 262,474 | |
| Live GFP positive | 46 | 56.79 % | FITC-A | Count | 14,937,072 | |
| Dead cells | 246 | 75.23 % | APC-Cy7-H | FSC-H | 78,717 | 1,799,475 |
| Dead GFP negative | 8 | 3.25 % | FITC-A | Count | 78,430 | |
| Dead GFP positive | 238 | 96.75 % | FITC-A | Count | 247,537 | |

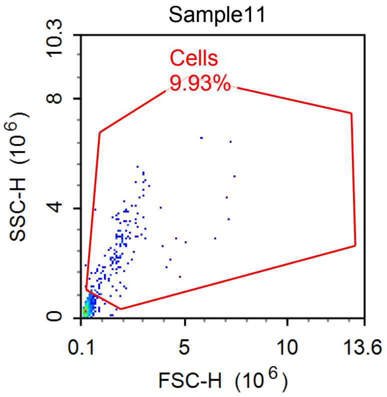


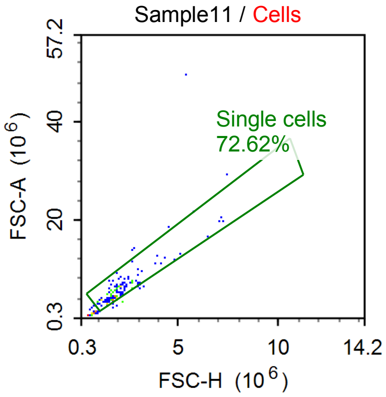


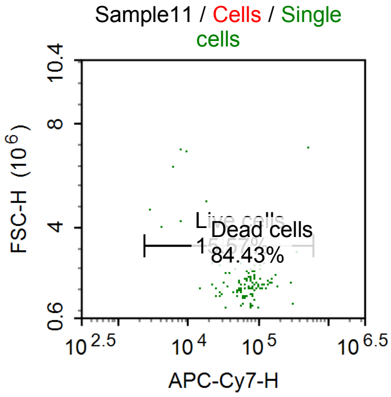


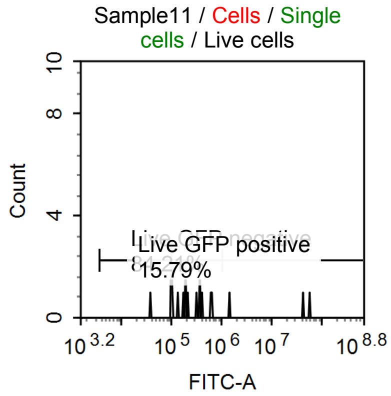


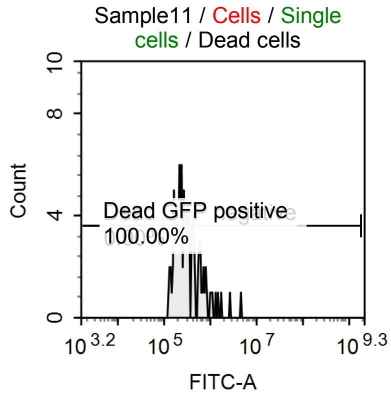


Sample Statistics of Sample11

| Gate | Count | % Parent | X | Y | Median X | Median Y |
|---|---|---|---|---|---|---|
| All | 1,691 | | | | | |
| Cells | 168 | 9.93 % | FSC-H | SSC-H | 1,758,607 | 2,379,646 |
| Single cells | 122 | 72.62 % | FSC-H | FSC-A | 1,848,933 | 5,610,381 |
| Live cells | 19 | 15.57 % | APC-Cy7-H | FSC-H | 21,483 | 2,611,551 |
| Live GFP negative | 16 | 84.21 % | FITC-A | Count | 195,892 | |
| Live GFP positive | 3 | 15.79 % | FITC-A | Count | 42,955,676 | |
| Dead cells | 103 | 84.43 % | APC-Cy7-H | FSC-H | 76,754 | 1,845,084 |
| Dead GFP negative | 0 | 0.00 % | FITC-A | Count | 0 | |
| Dead GFP positive | 103 | 100.00 % | FITC-A | Count | 296,123 | |

#### Specimen5

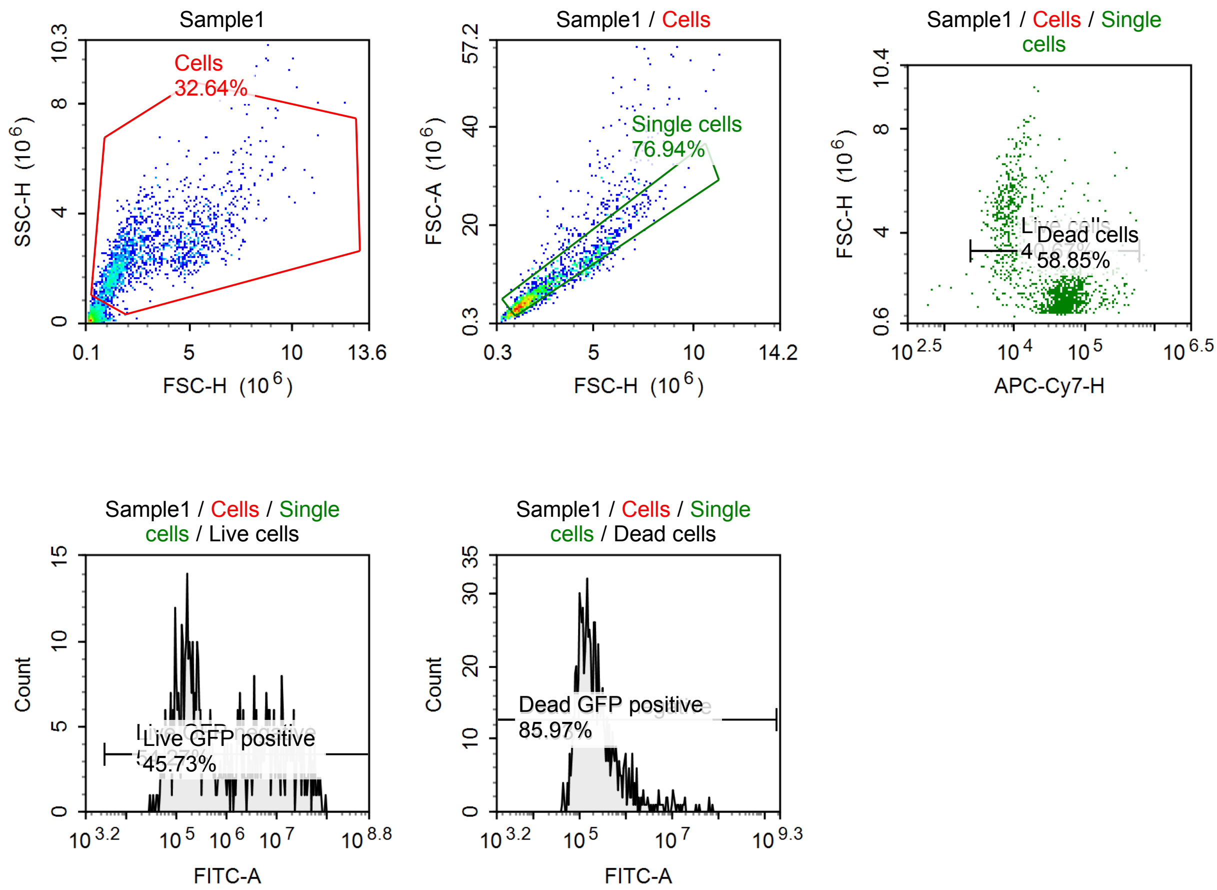


Sample Statistics of Sample1

| Gate | Count | % Parent | X | Y | Median X | Median Y |
|---|---|---|---|---|---|---|
| All | 5,738 | | | | | |
| Cells | 1,873 | 32.64 % | FSC-H | SSC-H | 2,110,983 | 2,370,279 |
| Single cells | 1,441 | 76.94 % | FSC-H | FSC-A | 1,948,394 | 5,776,739 |
| Live cells | 586 | 40.67 % | APC-Cy7-H | FSC-H | 10,201 | 4,062,688 |
| Live GFP negative | 318 | 54.27 % | FITC-A | Count | 178,675 | |
| Live GFP positive | 268 | 45.73 % | FITC-A | Count | 8,386,936 | |
| Dead cells | 848 | 58.85 % | APC-Cy7-H | FSC-H | 55,165 | 1,600,053 |
| Dead GFP negative | 119 | 14.03 % | FITC-A | Count | 79,232 | |
| Dead GFP positive | 729 | 85.97 % | FITC-A | Count | 218,607 | |

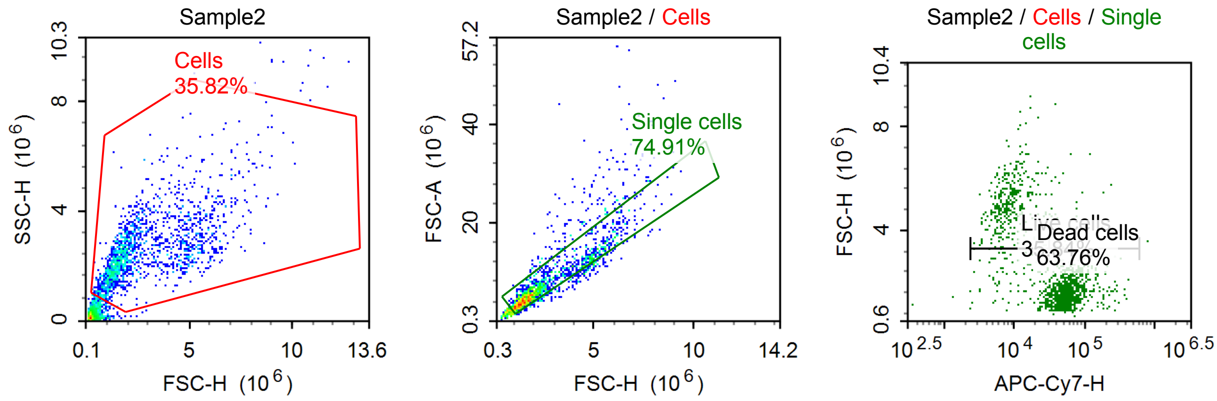

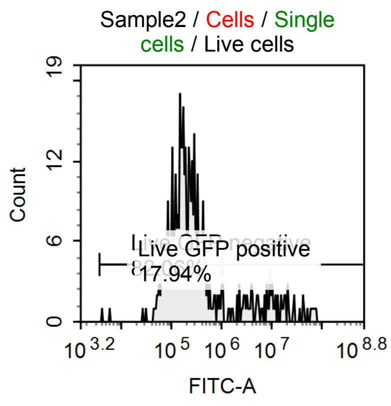


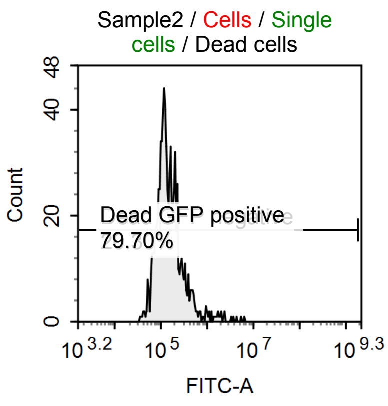


Sample Statistics of Sample2

| Gate | Count | % Parent | X | Y | Median X | Median Y |
|---|---|---|---|---|---|---|
| All | 4,752 | | | | | |
| Cells | 1,702 | 35.82 % | FSC-H | SSC-H | 2,040,148 | 2,416,524 |
| Single cells | 1,275 | 74.91 % | FSC-H | FSC-A | 1,918,927 | 5,775,781 |
| Live cells | 457 | 35.84 % | APC-Cy7-H | FSC-H | 9,322 | 4,553,281 |
| Live GFP negative | 375 | 82.06 % | FITC-A | Count | 187,818 | |
| Live GFP positive | 82 | 17.94 % | FITC-A | Count | 9,277,154 | |
| Dead cells | 813 | 63.76 % | APC-Cy7-H | FSC-H | 55,110 | 1,657,781 |
| Dead GFP negative | 165 | 20.30 % | FITC-A | Count | 77,044 | |
| Dead GFP positive | 648 | 79.70 % | FITC-A | Count | 163,223 | |

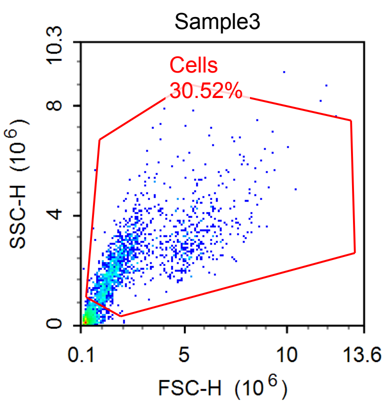


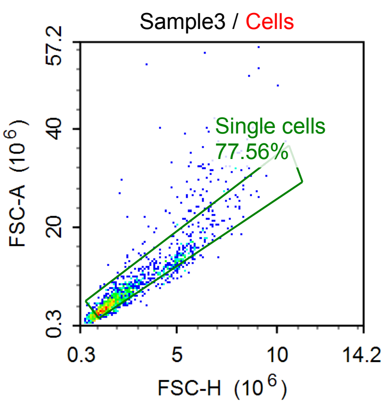


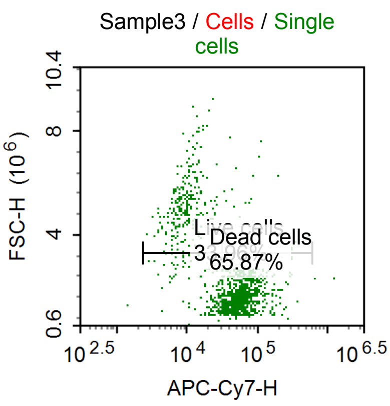


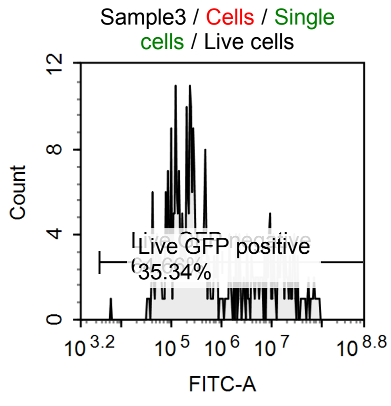


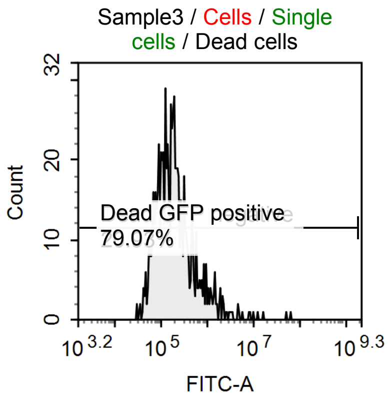

Sample Statistics of Sample3

| Gate | Count | % Parent | X | Y | Median X | Median Y |
|---|---|---|---|---|---|---|
| All | 4,964 | | | | | |
| Cells | 1,515 | 30.52 % | FSC-H | SSC-H | 1,904,357 | 2,327,991 |
| Single cells | 1,175 | 77.56 % | FSC-H | FSC-A | 1,901,866 | 5,545,248 |
| Live cells | 399 | 33.96 % | APC-Cy7-H | FSC-H | 11,624 | 4,337,504 |
| Live GFP negative | 258 | 64.66 % | FITC-A | Count | 203,741 | |
| Live GFP positive | 141 | 35.34 % | FITC-A | Count | 9,874,067 | |
| Dead cells | 774 | 65.87 % | APC-Cy7-H | FSC-H | 60,005 | 1,666,329 |
| Dead GFP negative | 162 | 20.93 % | FITC-A | Count | 71,287 | |
| Dead GFP positive | 612 | 79.07 % | FITC-A | Count | 212,082 | |

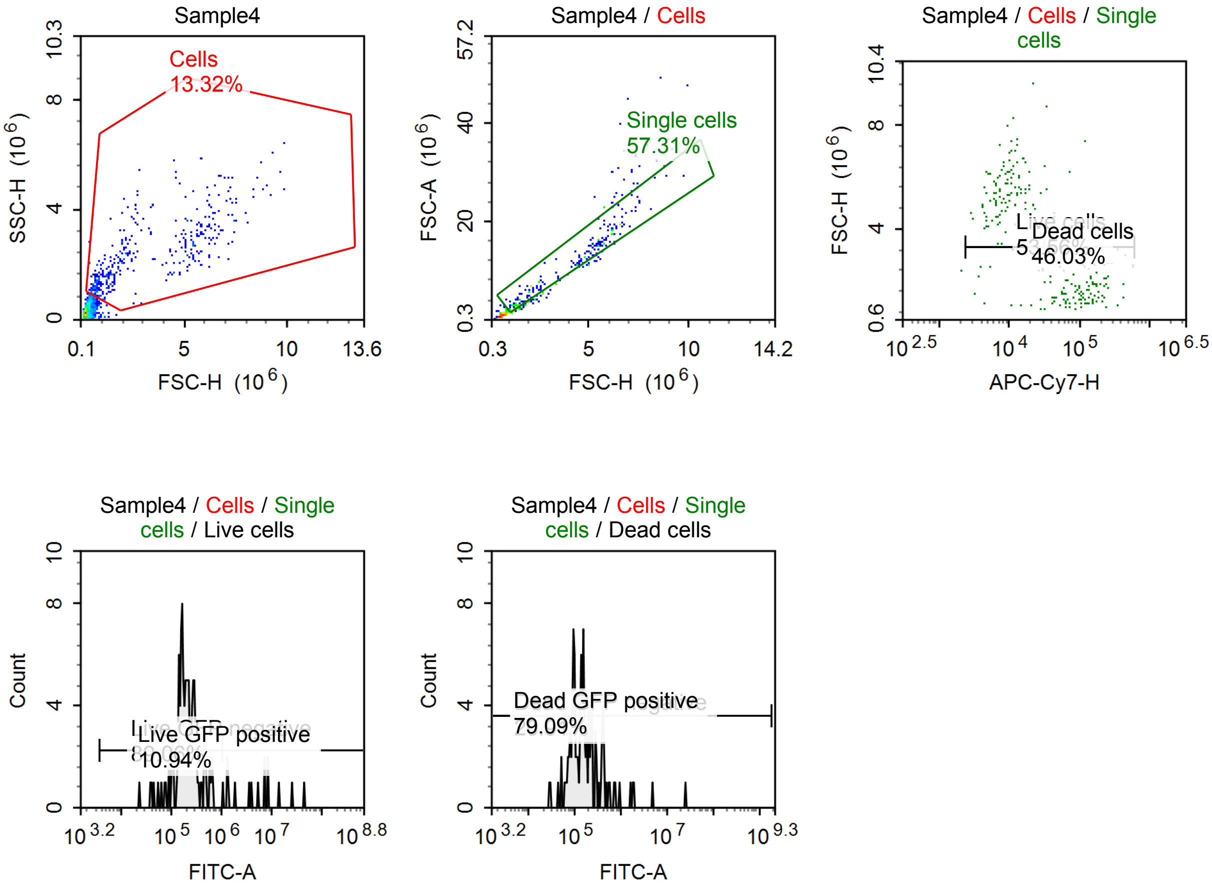


Sample Statistics of Sample4

| Gate | Count | % Parent | X | Y | Median X | Median Y |
|---|---|---|---|---|---|---|
| All | 3,131 | | | | | |
| Cells | 417 | 13.32 % | FSC-H | SSC-H | 1,924,263 | 2,032,148 |
| Single cells | 239 | 57.31 % | FSC-H | FSC-A | 3,297,036 | 8,554,739 |
| Live cells | 128 | 53.56 % | APC-Cy7-H | FSC-H | 9,070 | 5,277,541 |
| Live GFP negative | 114 | 89.06 % | FITC-A | Count | 198,553 | |
| Live GFP positive | 14 | 10.94 % | FITC-A | Count | 6,558,598 | |
| Dead cells | 110 | 46.03 % | APC-Cy7-H | FSC-H | 118,303 | 1,717,850 |
| Dead GFP negative | 23 | 20.91 % | FITC-A | Count | 77,690 | |
| Dead GFP positive | 87 | 79.09 % | FITC-A | Count | 180,679 | |

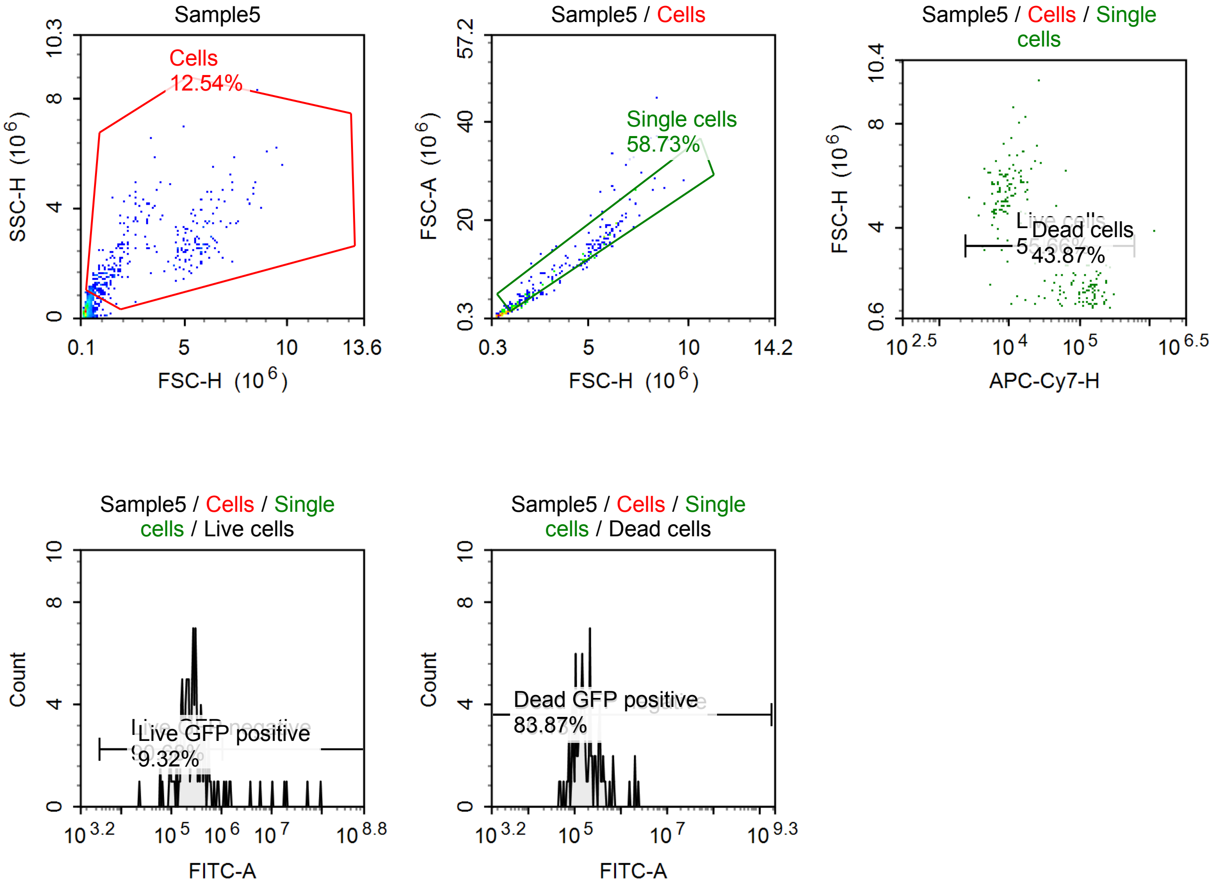


Sample Statistics of Sample5

| Gate | Count | % Parent | X | Y | Median X | Median Y |
|---|---|---|---|---|---|---|
| All | 2,879 | | | | | |
| Cells | 361 | 12.54 % | FSC-H | SSC-H | 1,876,868 | 1,959,085 |
| Single cells | 212 | 58.73 % | FSC-H | FSC-A | 3,585,643 | 10,473,131 |
| Live cells | 118 | 55.66 % | APC-Cy7-H | FSC-H | 9,175 | 5,295,844 |
| Live GFP negative | 107 | 90.68 % | FITC-A | Count | 262,948 | |
| Live GFP positive | 11 | 9.32 % | FITC-A | Count | 6,245,447 | |
| Dead cells | 93 | 43.87 % | APC-Cy7-H | FSC-H | 128,673 | 1,611,976 |
| Dead GFP negative | 15 | 16.13 % | FITC-A | Count | 78,659 | |
| Dead GFP positive | 78 | 83.87 % | FITC-A | Count | 196,160 | |

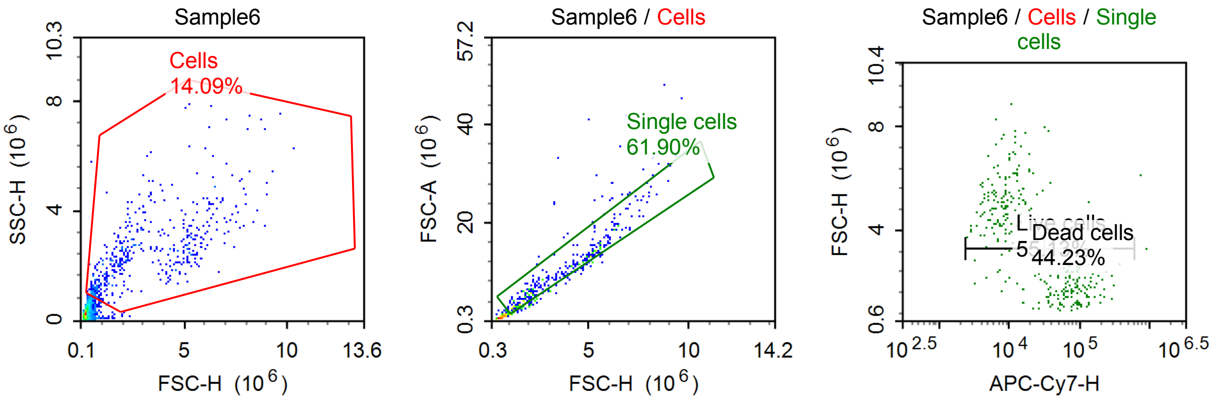

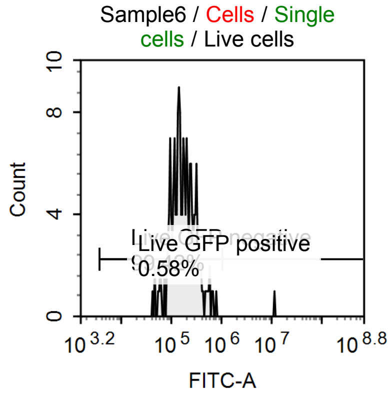


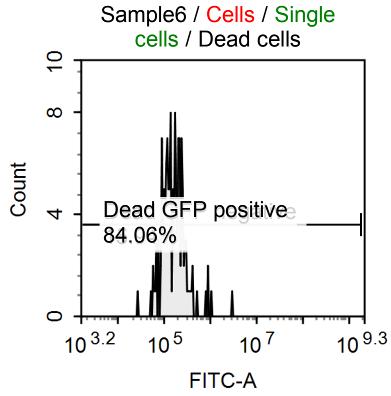


Sample Statistics of Sample6

| Gate | Count | % Parent | X | Y | Median X | Median Y |
|---|---|---|---|---|---|---|
| All | 3,576 | | | | | |
| Cells | 504 | 14.09 % | FSC-H | SSC-H | 2,210,034 | 2,232,496 |
| Single cells | 312 | 61.90 % | FSC-H | FSC-A | 3,357,063 | 9,275,666 |
| Live cells | 172 | 55.13 % | APC-Cy7-H | FSC-H | 8,541 | 5,082,909 |
| Live GFP negative | 171 | 99.42 % | FITC-A | Count | 173,505 | |
| Live GFP positive | 1 | 0.58 % | FITC-A | Count | 12,063,314 | |
| Dead cells | 138 | 44.23 % | APC-Cy7-H | FSC-H | 75,886 | 1,839,479 |
| Dead GFP negative | 22 | 15.94 % | FITC-A | Count | 76,772 | |
| Dead GFP positive | 116 | 84.06 % | FITC-A | Count | 177,739 | |

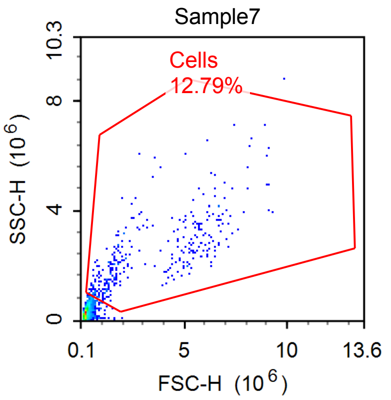


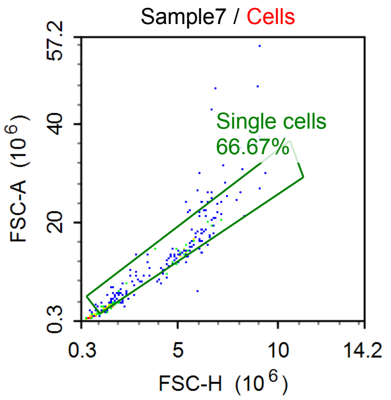


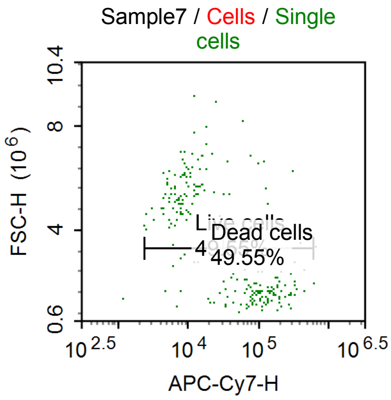


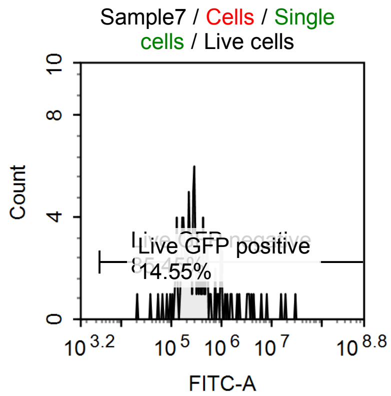


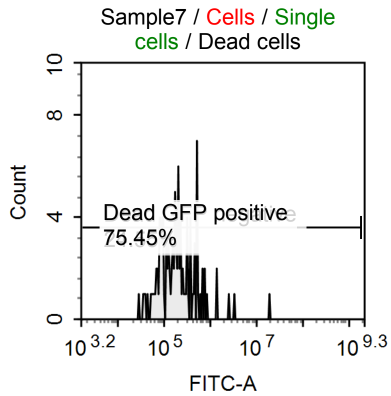

Sample Statistics of Sample7

| Gate | Count | % Parent | X | Y | Median X | Median Y |
|---|---|---|---|---|---|---|
| All | 2,604 | | | | | |
| Cells | 333 | 12.79 % | FSC-H | SSC-H | 2,137,853 | 2,139,350 |
| Single cells | 222 | 66.67 % | FSC-H | FSC-A | 3,102,215 | 9,438,980 |
| Live cells | 110 | 49.55 % | APC-Cy7-H | FSC-H | 8,223 | 5,298,619 |
| Live GFP negative | 94 | 85.45 % | FITC-A | Count | 261,732 | |
| Live GFP positive | 16 | 14.55 % | FITC-A | Count | 3,627,845 | |
| Dead cells | 110 | 49.55 % | APC-Cy7-H | FSC-H | 113,477 | 1,673,356 |
| Dead GFP negative | 27 | 24.55 % | FITC-A | Count | 74,268 | |
| Dead GFP positive | 83 | 75.45 % | FITC-A | Count | 227,491 | |

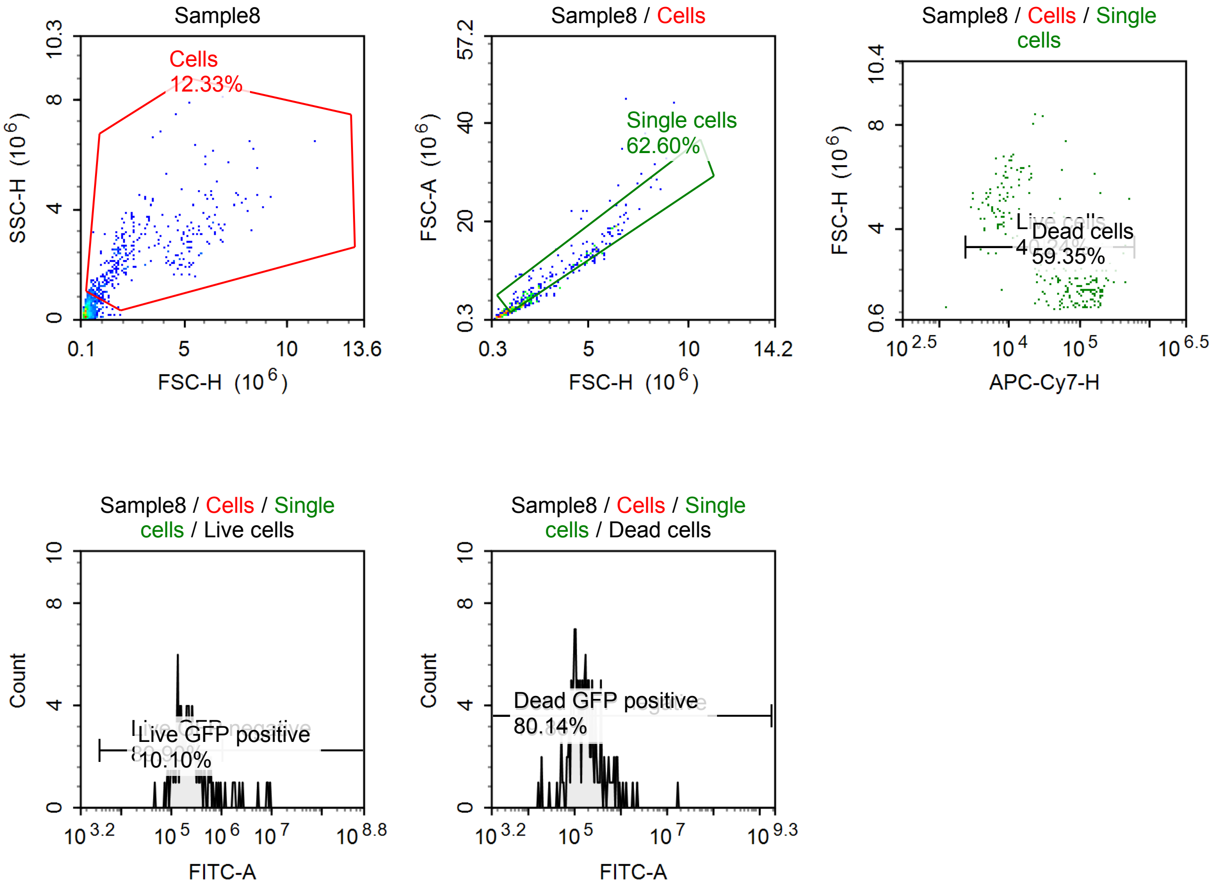


Sample Statistics of Sample8

| Gate | Count | % Parent | X | Y | Median X | Median Y |
|---|---|---|---|---|---|---|
| All | 3,187 | | | | | |
| Cells | 393 | 12.33 % | FSC-H | SSC-H | 1,973,739 | 2,143,088 |
| Single cells | 246 | 62.60 % | FSC-H | FSC-A | 2,403,086 | 6,560,645 |
| Live cells | 99 | 40.24 % | APC-Cy7-H | FSC-H | 8,171 | 5,016,978 |
| Live GFP negative | 89 | 89.90 % | FITC-A | Count | 211,883 | |
| Live GFP positive | 10 | 10.10 % | FITC-A | Count | 2,530,170 | |
| Dead cells | 146 | 59.35 % | APC-Cy7-H | FSC-H | 116,971 | 1,729,198 |
| Dead GFP negative | 29 | 19.86 % | FITC-A | Count | 73,353 | |
| Dead GFP positive | 117 | 80.14 % | FITC-A | Count | 189,871 | |

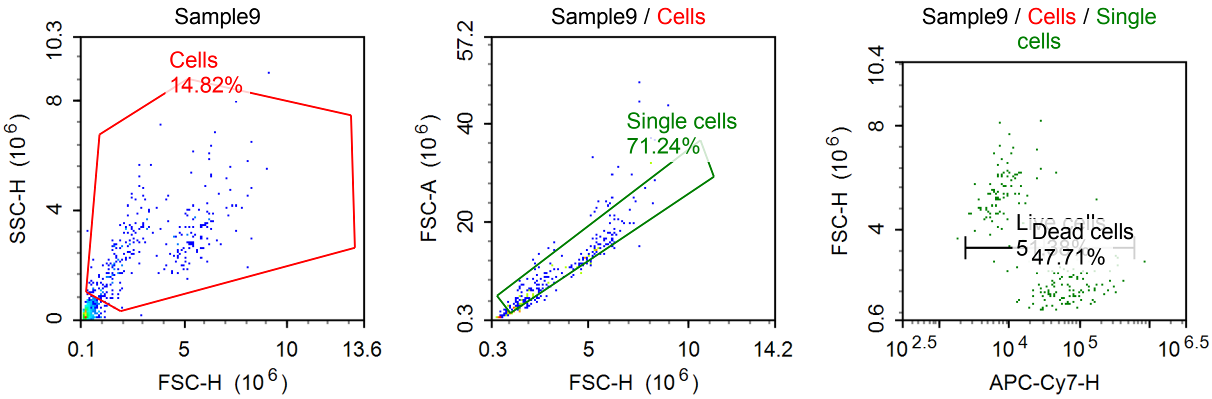


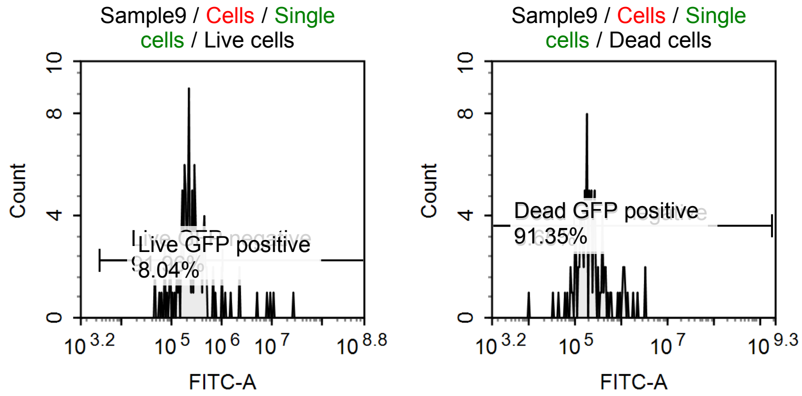


Sample Statistics of Sample9

| Gate | Count | % Parent | X | Y | Median X | Median Y |
|---|---|---|---|---|---|---|
| All | 2,065 | | | | | |
| Cells | 306 | 14.82 % | FSC-H | SSC-H | 2,563,797 | 2,617,578 |
| Single cells | 218 | 71.24 % | FSC-H | FSC-A | 2,860,138 | 8,786,912 |
| Live cells | 112 | 51.38 % | APC-Cy7-H | FSC-H | 8,200 | 5,156,948 |
| Live GFP negative | 103 | 91.96 % | FITC-A | Count | 229,864 | |
| Live GFP positive | 9 | 8.04 % | FITC-A | Count | 5,210,251 | |
| Dead cells | 104 | 47.71 % | APC-Cy7-H | FSC-H | 72,125 | 1,903,171 |
| Dead GFP negative | 9 | 8.65 % | FITC-A | Count | 69,991 | |
| Dead GFP positive | 95 | 91.35 % | FITC-A | Count | 216,377 | |

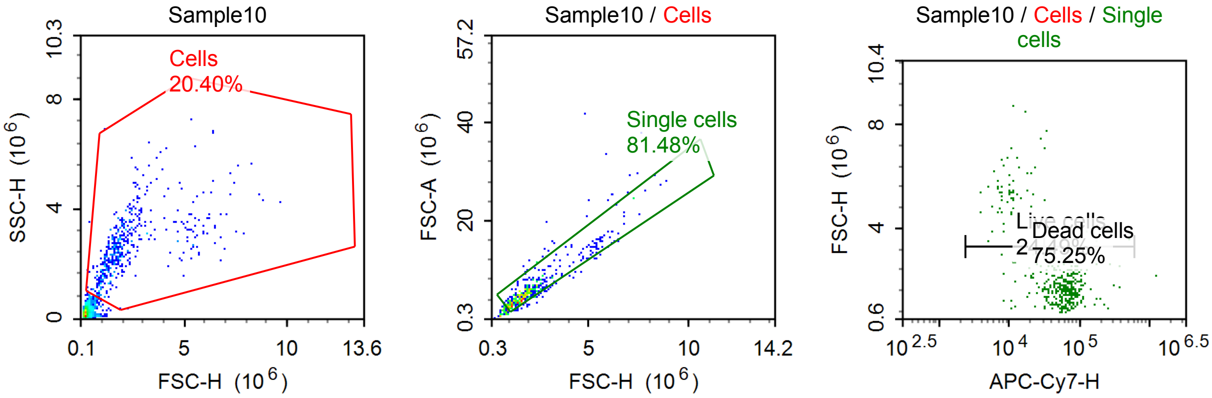

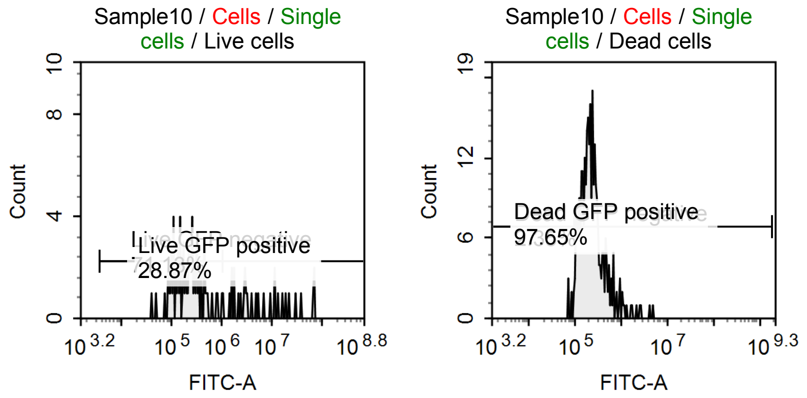


Sample Statistics of Sample10

| Gate | Count | % Parent | X | Y | Median X | Median Y |
|---|---|---|---|---|---|---|
| All | 2,382 | | | | | |
| Cells | 486 | 20.40 % | FSC-H | SSC-H | 1,806,145 | 2,590,474 |
| Single cells | 396 | 81.48 % | FSC-H | FSC-A | 1,828,614 | 5,347,769 |
| Live cells | 97 | 24.49 % | APC-Cy7-H | FSC-H | 11,510 | 4,724,519 |
| Live GFP negative | 69 | 71.13 % | FITC-A | Count | 222,481 | |
| Live GFP positive | 28 | 28.87 % | FITC-A | Count | 5,271,281 | |
| Dead cells | 298 | 75.25 % | APC-Cy7-H | FSC-H | 66,827 | 1,663,152 |
| Dead GFP negative | 7 | 2.35 % | FITC-A | Count | 84,931 | |
| Dead GFP positive | 291 | 97.65 % | FITC-A | Count | 222,380 | |

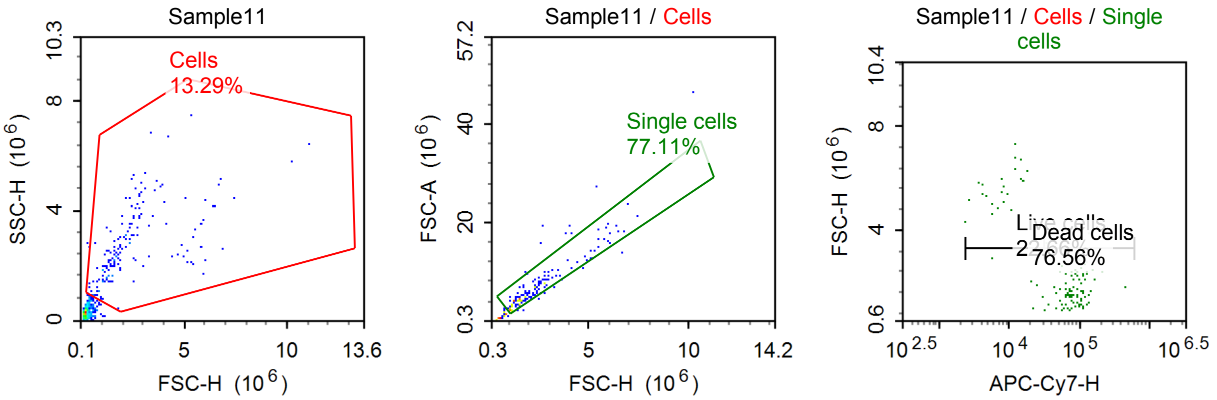


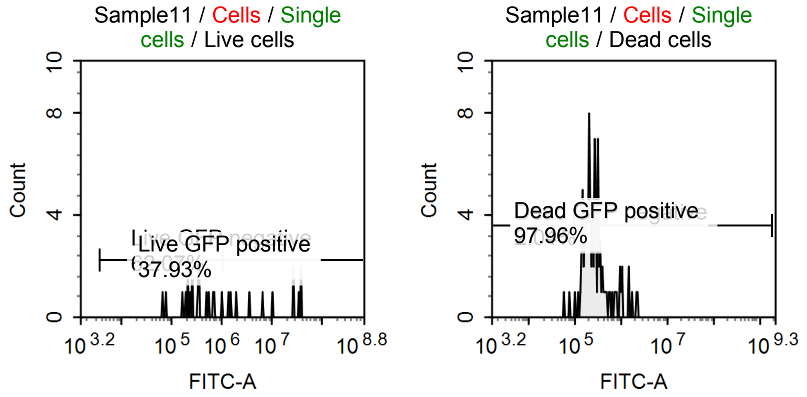

Sample Statistics of Sample11

| Gate | Count | % Parent | X | Y | Median X | Median Y |
|---|---|---|---|---|---|---|
| All | 1,249 | | | | | |
| Cells | 166 | 13.29 % | FSC-H | SSC-H | 1,845,164 | 2,455,996 |
| Single cells | 128 | 77.11 % | FSC-H | FSC-A | 2,115,369 | 5,865,575 |
| Live cells | 29 | 22.66 % | APC-Cy7-H | FSC-H | 10,106 | 5,301,309 |
| Live GFP negative | 18 | 62.07 % | FITC-A | Count | 308,629 | |
| Live GFP positive | 11 | 37.93 % | FITC-A | Count | 10,903,668 | |
| Dead cells | 98 | 76.56 % | APC-Cy7-H | FSC-H | 84,902 | 1,780,054 |
| Dead GFP negative | 2 | 2.04 % | FITC-A | Count | 68,226 | |
| Dead GFP positive | 96 | 97.96 % | FITC-A | Count | 262,411 | |

#### Specimen6

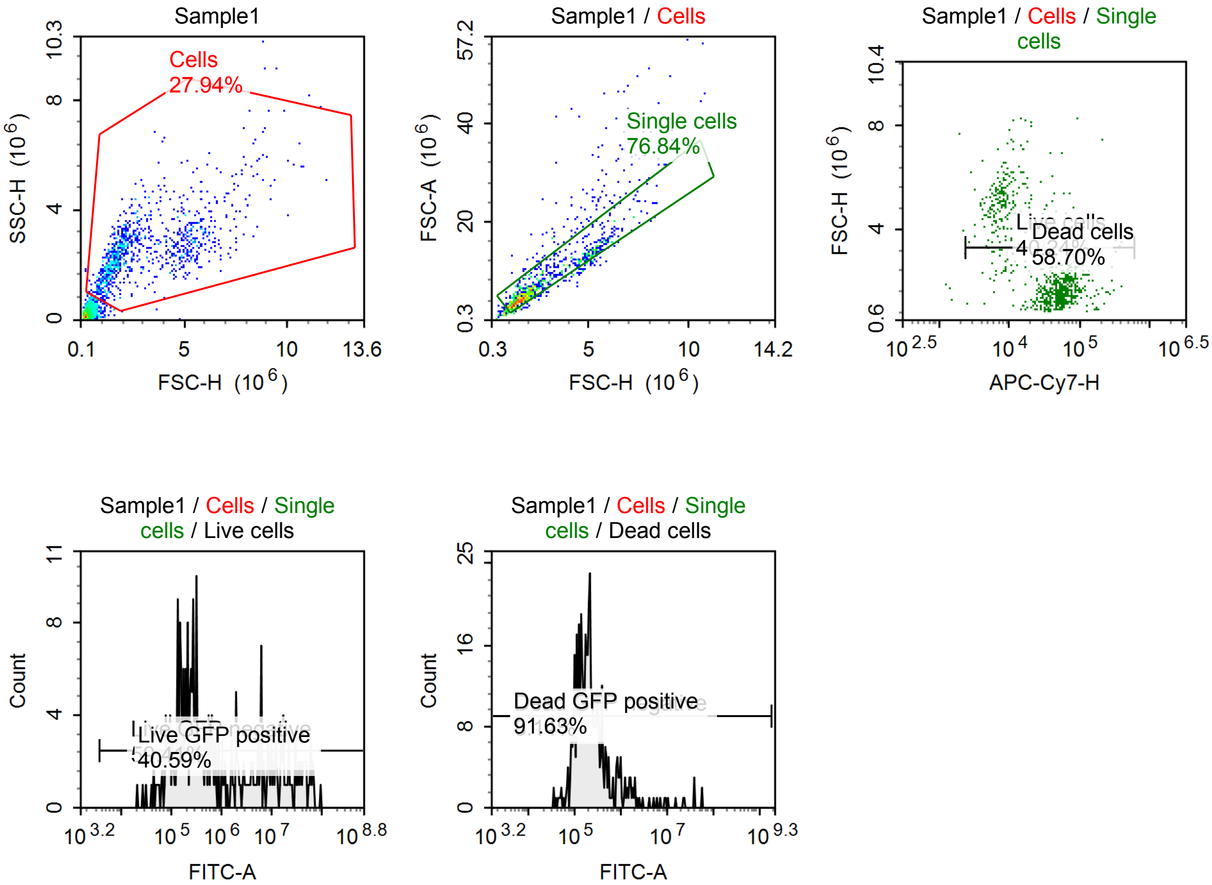

Sample Statistics of Sample1

| Gate | Count | % Parent | X | Y | Median X | Median Y |
|---|---|---|---|---|---|---|
| All | 3,507 | | | | | |
| Cells | 980 | 27.94 % | FSC-H | SSC-H | 2,234,304 | 2,493,010 |
| Single cells | 753 | 76.84 % | FSC-H | FSC-A | 2,041,616 | 6,091,416 |
| Live cells | 303 | 40.24 % | APC-Cy7-H | FSC-H | 9,193 | 4,504,449 |
| Live GFP negative | 180 | 59.41 % | FITC-A | Count | 217,492 | |
| Live GFP positive | 123 | 40.59 % | FITC-A | Count | 9,982,647 | |
| Dead cells | 442 | 58.70 % | APC-Cy7-H | FSC-H | 54,830 | 1,703,837 |
| Dead GFP negative | 36 | 8.14 % | FITC-A | Count | 82,217 | |
| Dead GFP positive | 405 | 91.63 % | FITC-A | Count | 215,447 | |

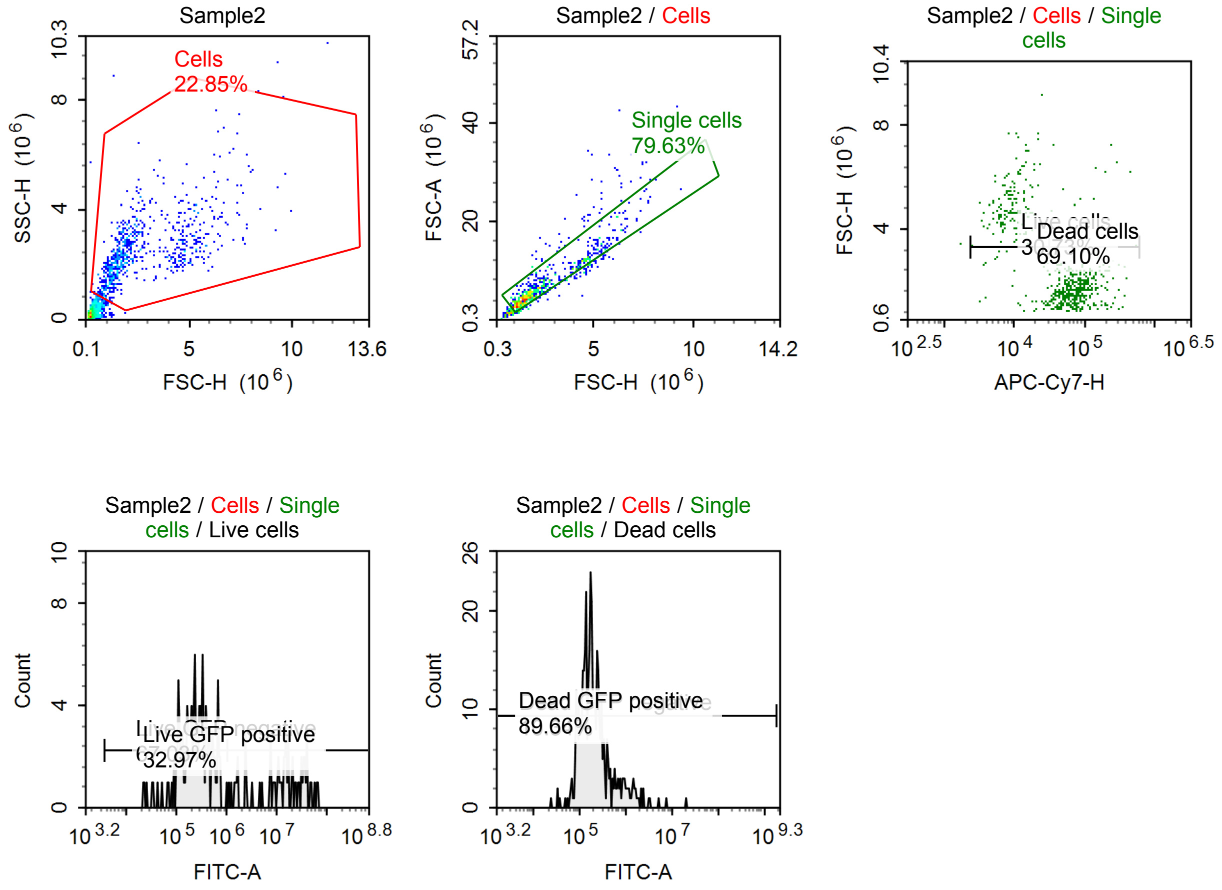


Sample Statistics of Sample2

| Gate | Count | % Parent | X | Y | Median X | Median Y |
|---|---|---|---|---|---|---|
| All | 3,309 | | | | | |
| Cells | 756 | 22.85 % | FSC-H | SSC-H | 1,967,193 | 2,372,077 |
| Single cells | 602 | 79.63 % | FSC-H | FSC-A | 1,926,511 | 5,731,220 |
| Live cells | 185 | 30.73 % | APC-Cy7-H | FSC-H | 9,779 | 4,646,058 |
| Live GFP negative | 124 | 67.03 % | FITC-A | Count | 238,704 | |
| Live GFP positive | 61 | 32.97 % | FITC-A | Count | 15,312,879 | |
| Dead cells | 416 | 69.10 % | APC-Cy7-H | FSC-H | 70,478 | 1,640,776 |
| Dead GFP negative | 43 | 10.34 % | FITC-A | Count | 84,157 | |
| Dead GFP positive | 373 | 89.66 % | FITC-A | Count | 199,467 | |

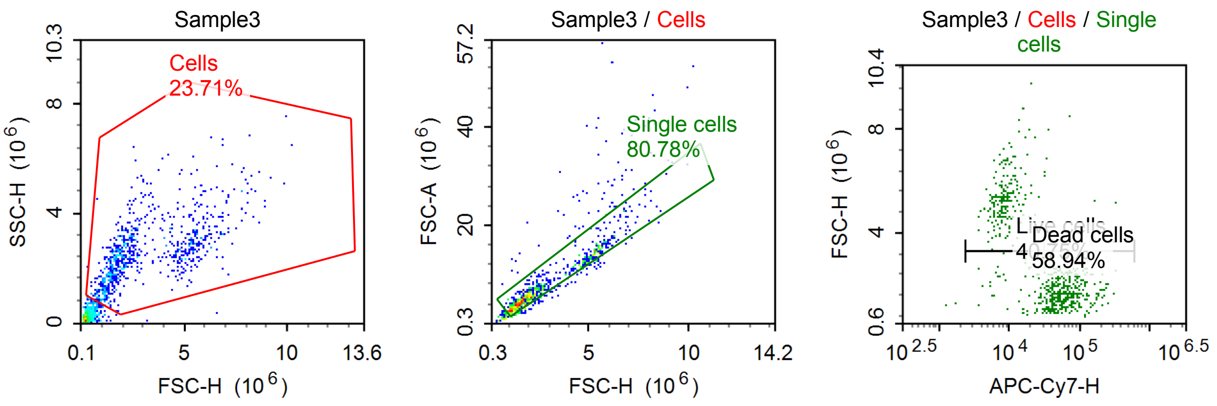


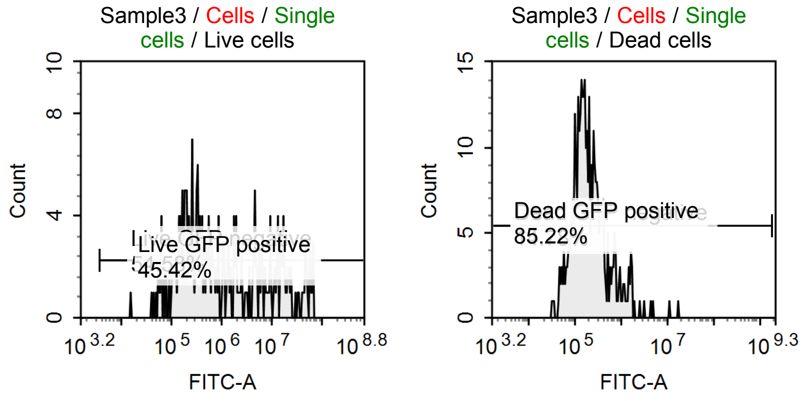


Sample Statistics of Sample3

| Gate | Count | % Parent | X | Y | Median X | Median Y |
|---|---|---|---|---|---|---|
| All | 3,357 | | | | | |
| Cells | 796 | 23.71 % | FSC-H | SSC-H | 2,271,278 | 2,483,864 |
| Single cells | 643 | 80.78 % | FSC-H | FSC-A | 2,180,143 | 6,419,482 |
| Live cells | 262 | 40.75 % | APC-Cy7-H | FSC-H | 8,850 | 5,007,906 |
| Live GFP negative | 143 | 54.58 % | FITC-A | Count | 261,569 | |
| Live GFP positive | 119 | 45.42 % | FITC-A | Count | 8,308,914 | |
| Dead cells | 379 | 58.94 % | APC-Cy7-H | FSC-H | 62,339 | 1,728,485 |
| Dead GFP negative | 56 | 14.78 % | FITC-A | Count | 72,556 | |
| Dead GFP positive | 323 | 85.22 % | FITC-A | Count | 211,494 | |

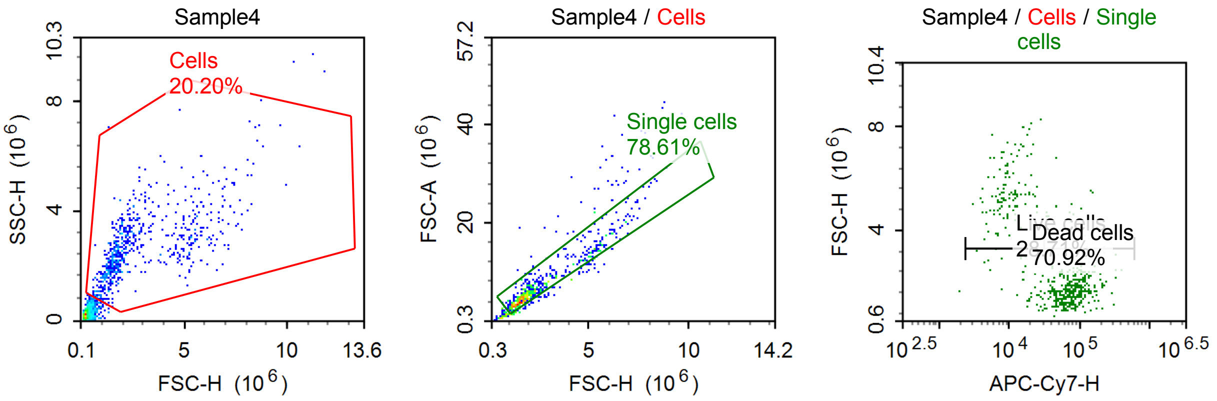

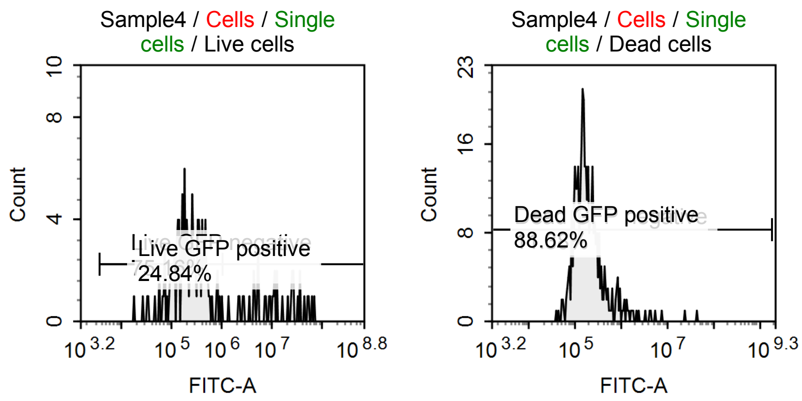


Sample Statistics of Sample4

| Gate | Count | % Parent | X | Y | Median X | Median Y |
|---|---|---|---|---|---|---|
| All | 3,356 | | | | | |
| Cells | 678 | 20.20 % | FSC-H | SSC-H | 1,950,153 | 2,589,745 |
| Single cells | 533 | 78.61 % | FSC-H | FSC-A | 1,941,577 | 5,696,380 |
| Live cells | 153 | 28.71 % | APC-Cy7-H | FSC-H | 10,731 | 5,105,049 |
| Live GFP negative | 115 | 75.16 % | FITC-A | Count | 216,164 | |
| Live GFP positive | 38 | 24.84 % | FITC-A | Count | 11,617,898 | |
| Dead cells | 378 | 70.92 % | APC-Cy7-H | FSC-H | 77,340 | 1,753,234 |
| Dead GFP negative | 43 | 11.38 % | FITC-A | Count | 79,355 | |
| Dead GFP positive | 335 | 88.62 % | FITC-A | Count | 187,060 | |

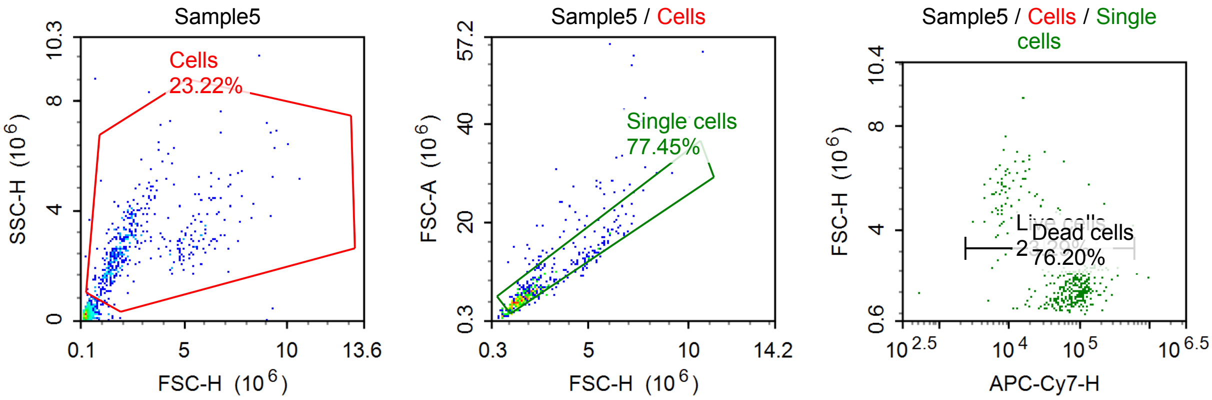


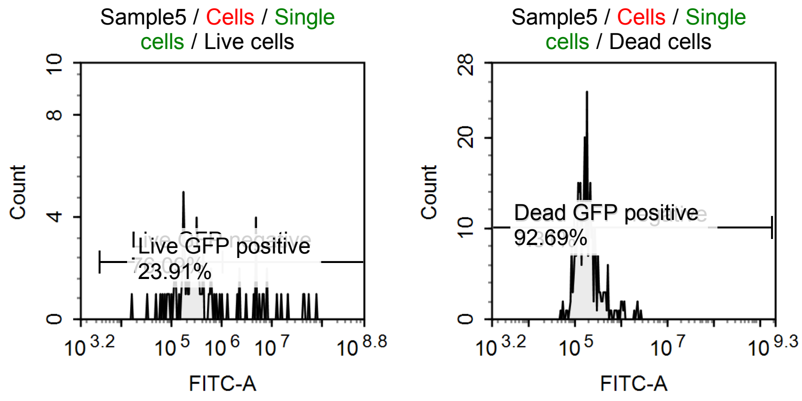

Sample Statistics of Sample5

| Gate | Count | % Parent | X | Y | Median X | Median Y |
|---|---|---|---|---|---|---|
| All | 2,196 | | | | | |
| Cells | 510 | 23.22 % | FSC-H | SSC-H | 2,076,756 | 2,615,490 |
| Single cells | 395 | 77.45 % | FSC-H | FSC-A | 1,937,113 | 5,750,326 |
| Live cells | 92 | 23.29 % | APC-Cy7-H | FSC-H | 9,571 | 5,006,230 |
| Live GFP negative | 70 | 76.09 % | FITC-A | Count | 252,776 | |
| Live GFP positive | 22 | 23.91 % | FITC-A | Count | 6,360,493 | |
| Dead cells | 301 | 76.20 % | APC-Cy7-H | FSC-H | 90,262 | 1,784,976 |
| Dead GFP negative | 22 | 7.31 % | FITC-A | Count | 85,939 | |
| Dead GFP positive | 279 | 92.69 % | FITC-A | Count | 183,156 | |

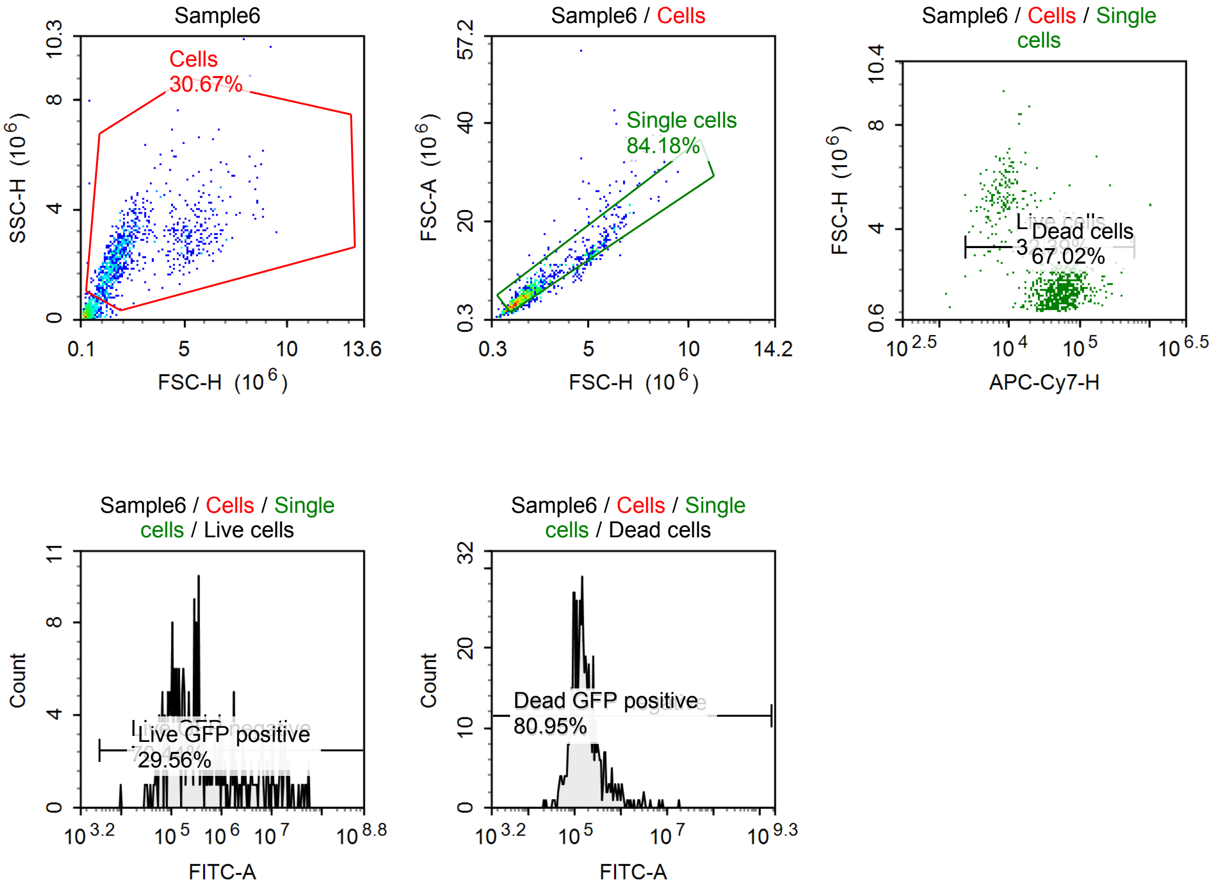


Sample Statistics of Sample6

| Gate | Count | % Parent | X | Y | Median X | Median Y |
|---|---|---|---|---|---|---|
| All | 3,277 | | | | | |
| Cells | 1,005 | 30.67 % | FSC-H | SSC-H | 1,925,501 | 2,399,002 |
| Single cells | 846 | 84.18 % | FSC-H | FSC-A | 1,878,149 | 5,214,278 |
| Live cells | 274 | 32.39 % | APC-Cy7-H | FSC-H | 11,476 | 4,479,698 |
| Live GFP negative | 193 | 70.44 % | FITC-A | Count | 177,616 | |
| Live GFP positive | 81 | 29.56 % | FITC-A | Count | 6,610,078 | |
| Dead cells | 567 | 67.02 % | APC-Cy7-H | FSC-H | 61,548 | 1,679,391 |
| Dead GFP negative | 108 | 19.05 % | FITC-A | Count | 79,360 | |
| Dead GFP positive | 459 | 80.95 % | FITC-A | Count | 172,315 | |

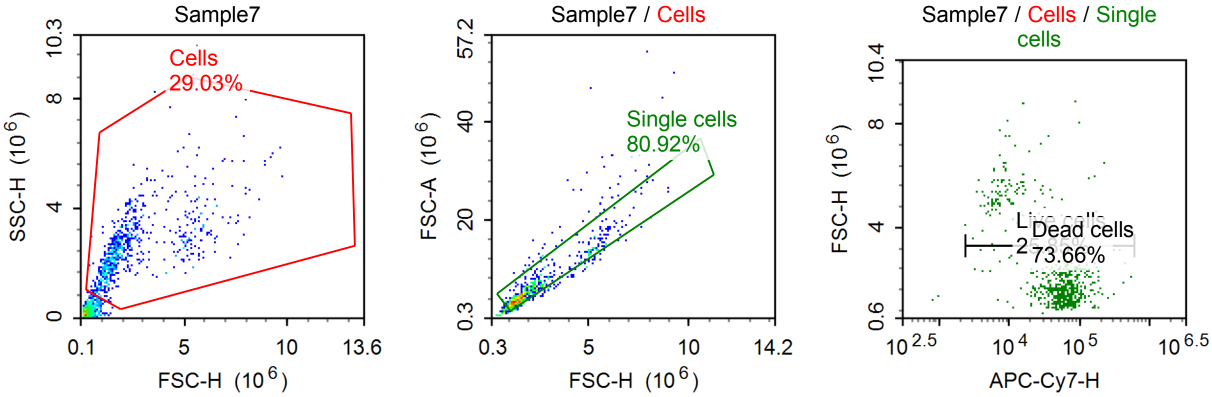


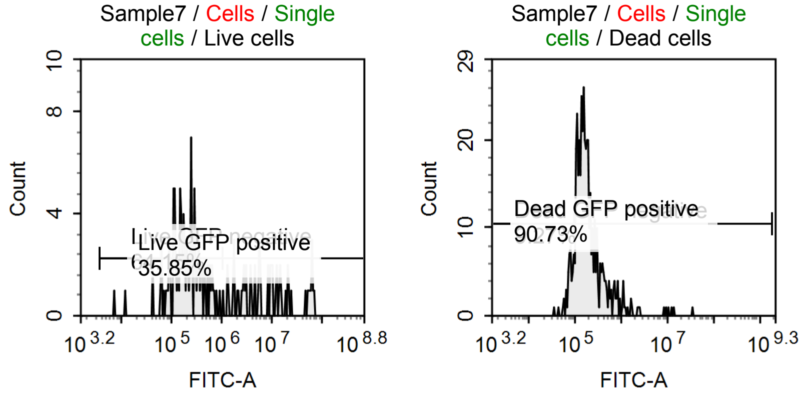


Sample Statistics of Sample7

| Gate | Count | % Parent | X | Y | Median X | Median Y |
|---|---|---|---|---|---|---|
| All | 2,618 | | | | | |
| Cells | 760 | 29.03 % | FSC-H | SSC-H | 1,767,410 | 2,440,495 |
| Single cells | 615 | 80.92 % | FSC-H | FSC-A | 1,736,264 | 5,005,425 |
| Live cells | 159 | 25.85 % | APC-Cy7-H | FSC-H | 11,001 | 4,906,319 |
| Live GFP negative | 102 | 64.15 % | FITC-A | Count | 194,064 | |
| Live GFP positive | 57 | 35.85 % | FITC-A | Count | 8,724,694 | |
| Dead cells | 453 | 73.66 % | APC-Cy7-H | FSC-H | 64,598 | 1,601,614 |
| Dead GFP negative | 42 | 9.27 % | FITC-A | Count | 79,104 | |
| Dead GFP positive | 411 | 90.73 % | FITC-A | Count | 175,439 | |

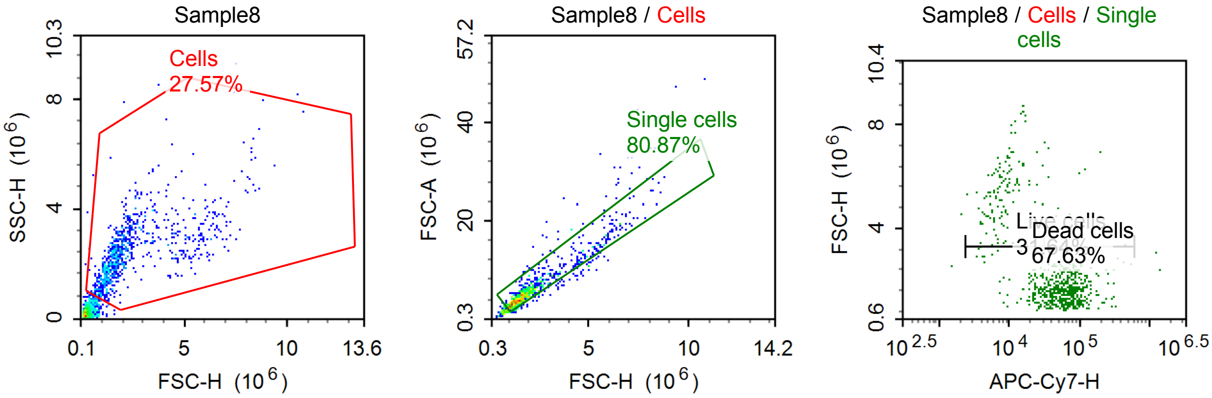

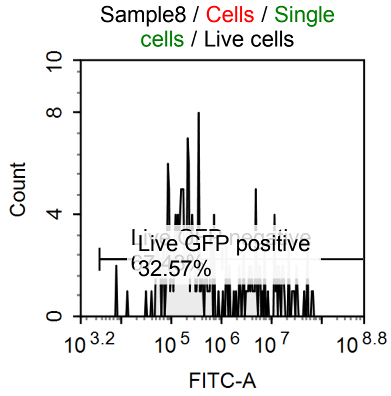


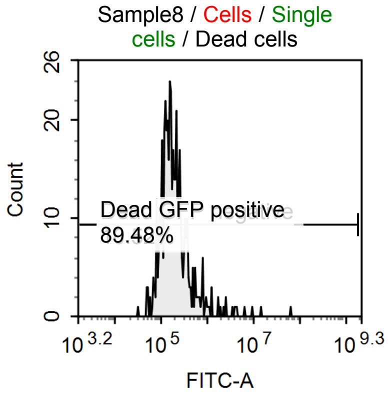


Sample Statistics of Sample8

| Gate | Count | % Parent | X | Y | Median X | Median Y |
|---|---|---|---|---|---|---|
| All | 3,090 | | | | | |
| Cells | 852 | 27.57 % | FSC-H | SSC-H | 1,797,174 | 2,339,079 |
| Single cells | 689 | 80.87 % | FSC-H | FSC-A | 1,797,653 | 5,249,140 |
| Live cells | 218 | 31.64 % | APC-Cy7-H | FSC-H | 12,454 | 3,815,708 |
| Live GFP negative | 147 | 67.43 % | FITC-A | Count | 175,532 | |
| Live GFP positive | 71 | 32.57 % | FITC-A | Count | 6,158,772 | |
| Dead cells | 466 | 67.63 % | APC-Cy7-H | FSC-H | 67,096 | 1,639,464 |
| Dead GFP negative | 49 | 10.52 % | FITC-A | Count | 81,640 | |
| Dead GFP positive | 417 | 89.48 % | FITC-A | Count | 188,209 | |

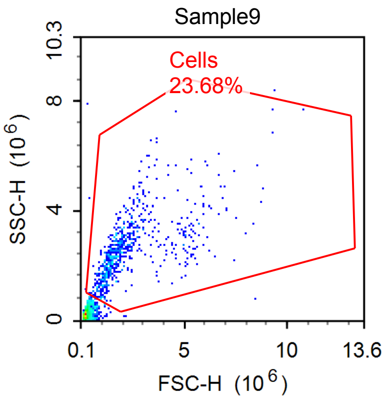


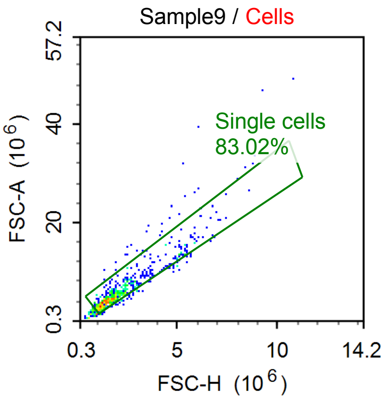


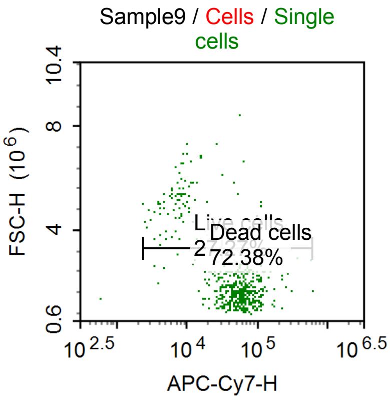


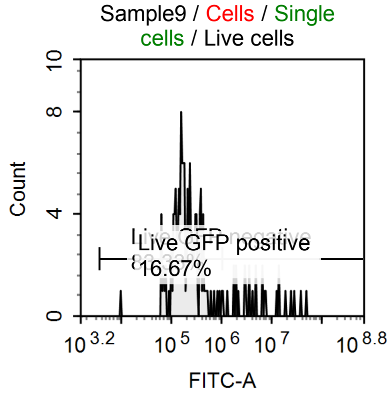


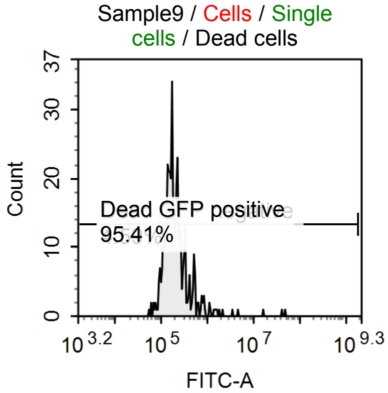

Sample Statistics of Sample9

| Gate | Count | % Parent | X | Y | Median X | Median Y |
|---|---|---|---|---|---|---|
| All | 2,910 | | | | | |
| Cells | 689 | 23.68 % | FSC-H | SSC-H | 1,780,426 | 2,511,538 |
| Single cells | 572 | 83.02 % | FSC-H | FSC-A | 1,832,189 | 5,248,215 |
| Live cells | 156 | 27.27 % | APC-Cy7-H | FSC-H | 13,646 | 3,669,115 |
| Live GFP negative | 130 | 83.33 % | FITC-A | Count | 186,155 | |
| Live GFP positive | 26 | 16.67 % | FITC-A | Count | 5,012,366 | |
| Dead cells | 414 | 72.38 % | APC-Cy7-H | FSC-H | 61,659 | 1,660,964 |
| Dead GFP negative | 19 | 4.59 % | FITC-A | Count | 87,282 | |
| Dead GFP positive | 395 | 95.41 % | FITC-A | Count | 188,986 | |

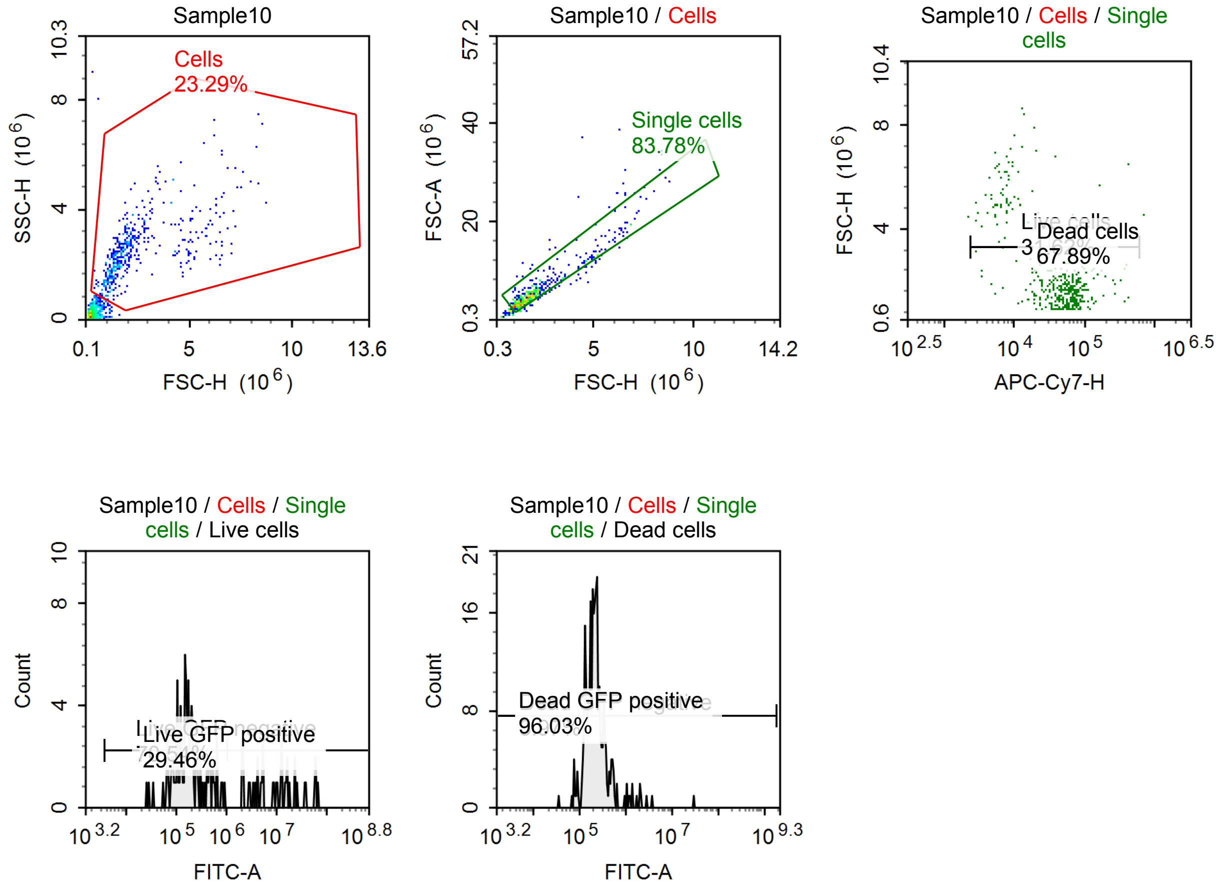


Sample Statistics of Sample10

| Gate | Count | % Parent | X | Y | Median X | Median Y |
|---|---|---|---|---|---|---|
| All | 2,091 | | | | | |
| Cells | 487 | 23.29 % | FSC-H | SSC-H | 1,802,222 | 2,400,458 |
| Single cells | 408 | 83.78 % | FSC-H | FSC-A | 1,816,768 | 5,080,671 |
| Live cells | 129 | 31.62 % | APC-Cy7-H | FSC-H | 12,250 | 3,377,514 |
| Live GFP negative | 91 | 70.54 % | FITC-A | Count | 173,318 | |
| Live GFP positive | 38 | 29.46 % | FITC-A | Count | 10,567,110 | |
| Dead cells | 277 | 67.89 % | APC-Cy7-H | FSC-H | 60,894 | 1,633,714 |
| Dead GFP negative | 11 | 3.97 % | FITC-A | Count | 77,384 | |
| Dead GFP positive | 266 | 96.03 % | FITC-A | Count | 220,802 | |

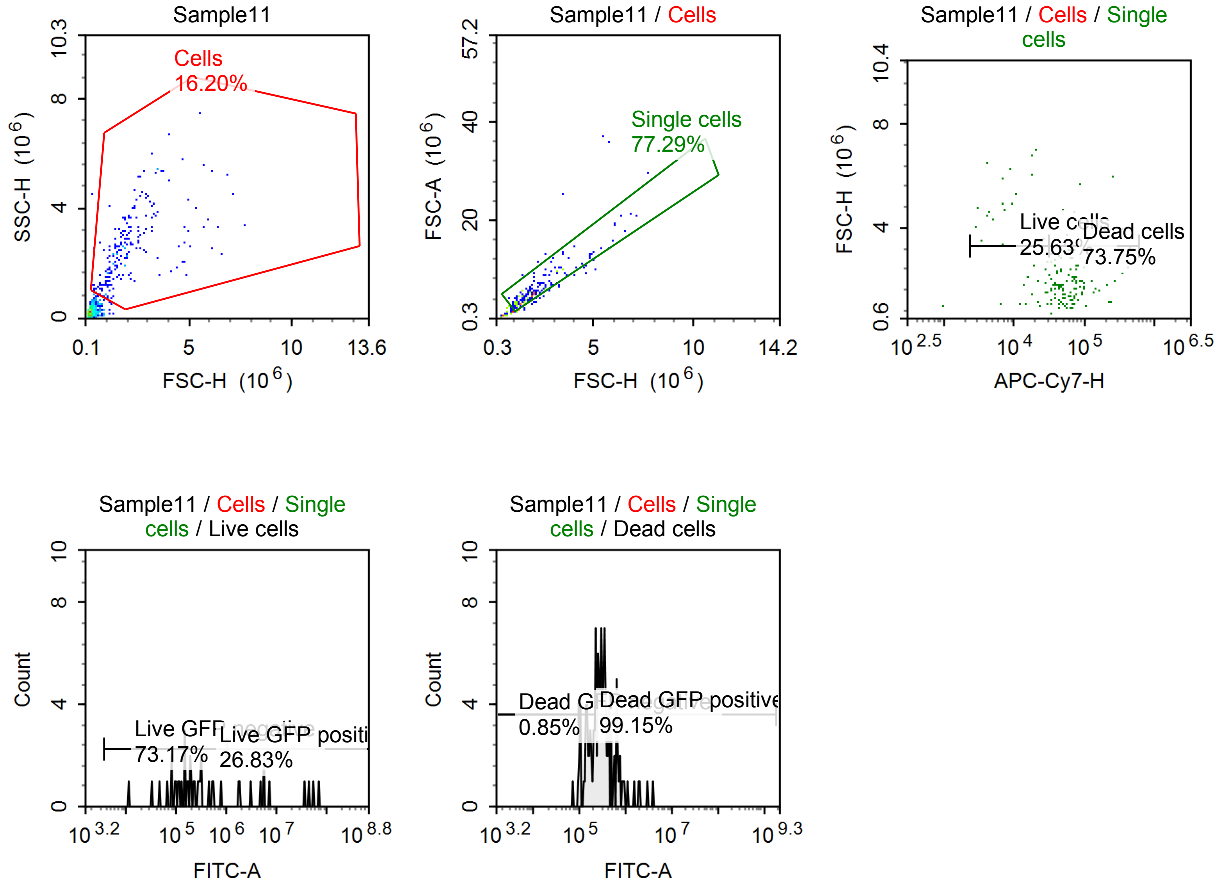


Sample Statistics of Sample11

| Gate | Count | % Parent | X | Y | Median X | Median Y |
|---|---|---|---|---|---|---|
| All | 1,278 | | | | | |
| Cells | 207 | 16.20 % | FSC-H | SSC-H | 1,805,438 | 2,499,856 |
| Single cells | 160 | 77.29 % | FSC-H | FSC-A | 1,908,356 | 5,425,111 |
| Live cells | 41 | 25.63 % | APC-Cy7-H | FSC-H | 14,727 | 2,585,224 |
| Live GFP negative | 30 | 73.17 % | FITC-A | Count | 173,304 | |
| Live GFP positive | 11 | 26.83 % | FITC-A | Count | 5,906,322 | |
| Dead cells | 118 | 73.75 % | APC-Cy7-H | FSC-H | 63,263 | 1,859,620 |
| Dead GFP negative | 1 | 0.85 % | FITC-A | Count | 71,629 | |
| Dead GFP positive | 117 | 99.15 % | FITC-A | Count | 296,609 | |

#### Specimen7

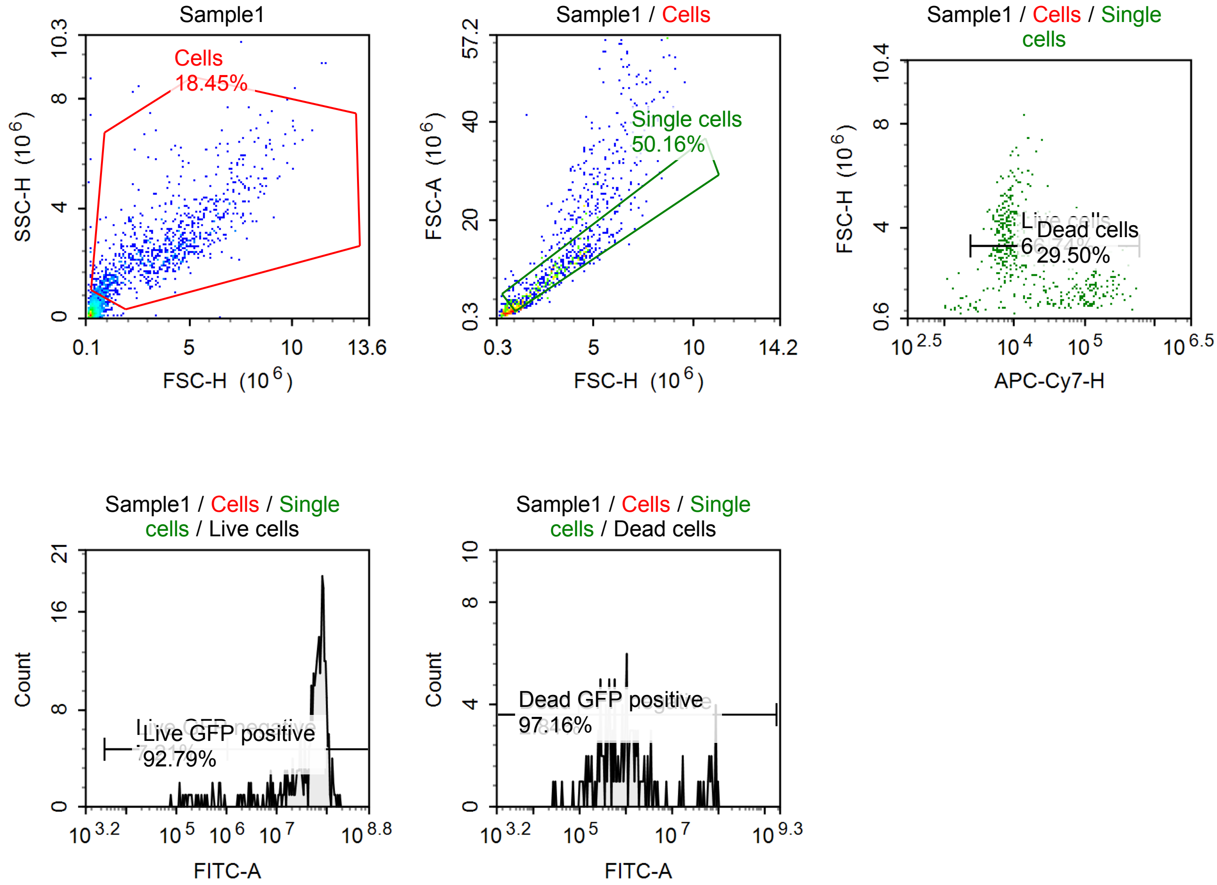


Sample Statistics of Sample1

| Gate | Count | % Parent | X | Y | Median X | Median Y |
|---|---|---|---|---|---|---|
| All | 5,165 | | | | | |
| Cells | 953 | 18.45 % | FSC-H | SSC-H | 3,257,832 | 2,321,088 |
| Single cells | 478 | 50.16 % | FSC-H | FSC-A | 2,837,642 | 8,517,654 |
| Live cells | 319 | 66.74 % | APC-Cy7-H | FSC-H | 8,420 | 3,541,879 |
| Live GFP negative | 23 | 7.21 % | FITC-A | Count | 287,689 | |
| Live GFP positive | 296 | 92.79 % | FITC-A | Count | 66,547,224 | |
| Dead cells | 141 | 29.50 % | APC-Cy7-H | FSC-H | 114,705 | 1,619,714 |
| Dead GFP negative | 4 | 2.84 % | FITC-A | Count | 36,202 | |
| Dead GFP positive | 137 | 97.16 % | FITC-A | Count | 937,194 | |

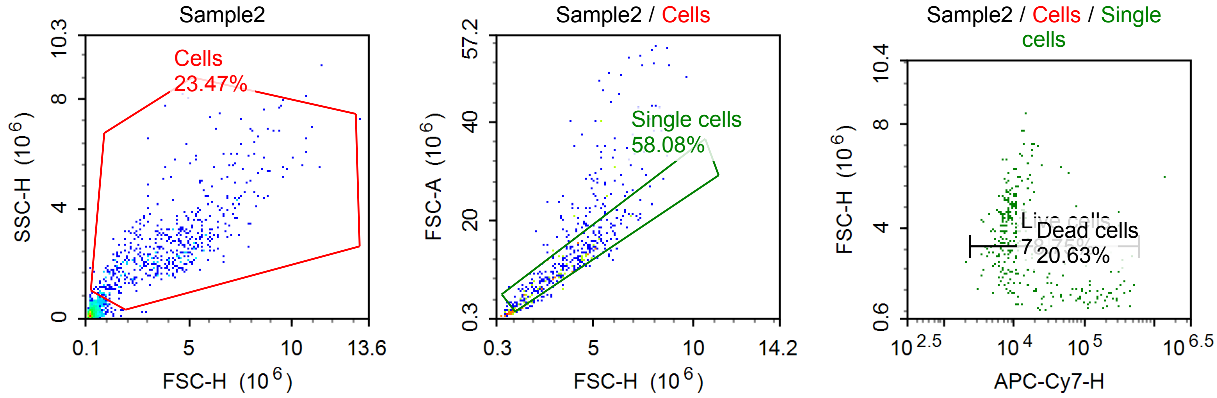

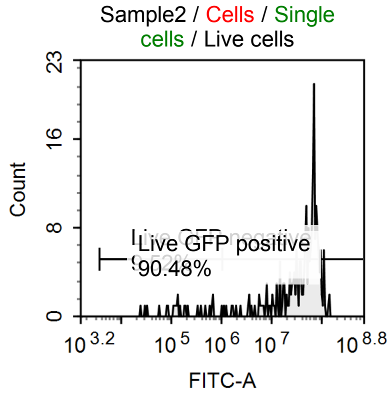


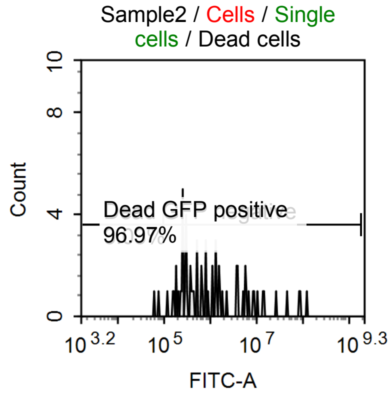


Sample Statistics of Sample2

| Gate | Count | % Parent | X | Y | Median X | Median Y |
|---|---|---|---|---|---|---|
| All | 2,348 | | | | | |
| Cells | 551 | 23.47 % | FSC-H | SSC-H | 3,850,073 | 2,405,677 |
| Single cells | 320 | 58.08 % | FSC-H | FSC-A | 3,414,951 | 10,057,415 |
| Live cells | 252 | 78.75 % | APC-Cy7-H | FSC-H | 8,698 | 3,909,210 |
| Live GFP negative | 24 | 9.52 % | FITC-A | Count | 146,156 | |
| Live GFP positive | 228 | 90.48 % | FITC-A | Count | 55,532,240 | |
| Dead cells | 66 | 20.63 % | APC-Cy7-H | FSC-H | 128,571 | 1,555,842 |
| Dead GFP negative | 2 | 3.03 % | FITC-A | Count | 68,781 | |
| Dead GFP positive | 64 | 96.97 % | FITC-A | Count | 828,502 | |

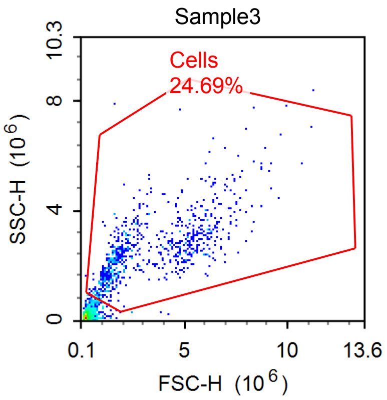


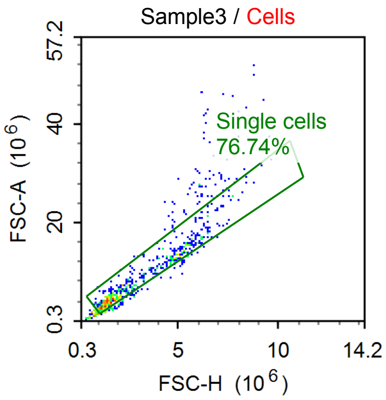


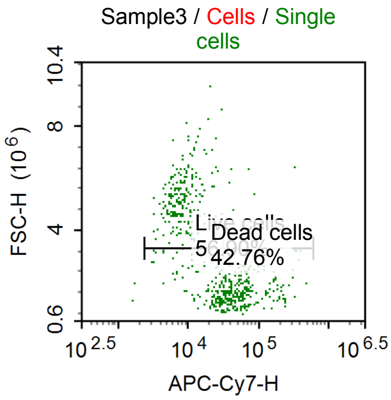


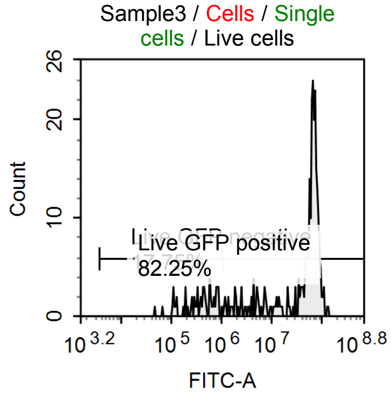


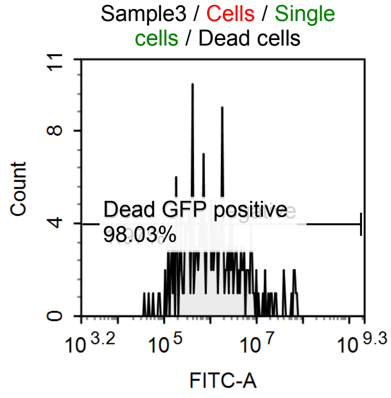

Sample Statistics of Sample3

| Gate | Count | % Parent | X | Y | Median X | Median Y |
|---|---|---|---|---|---|---|
| All | 3,135 | | | | | |
| Cells | 774 | 24.69 % | FSC-H | SSC-H | 2,449,176 | 2,471,055 |
| Single cells | 594 | 76.74 % | FSC-H | FSC-A | 2,214,817 | 6,480,554 |
| Live cells | 338 | 56.90 % | APC-Cy7-H | FSC-H | 10,278 | 4,536,969 |
| Live GFP negative | 60 | 17.75 % | FITC-A | Count | 439,534 | |
| Live GFP positive | 278 | 82.25 % | FITC-A | Count | 65,517,428 | |
| Dead cells | 254 | 42.76 % | APC-Cy7-H | FSC-H | 55,007 | 1,616,739 |
| Dead GFP negative | 5 | 1.97 % | FITC-A | Count | 58,712 | |
| Dead GFP positive | 249 | 98.03 % | FITC-A | Count | 1,061,154 | |

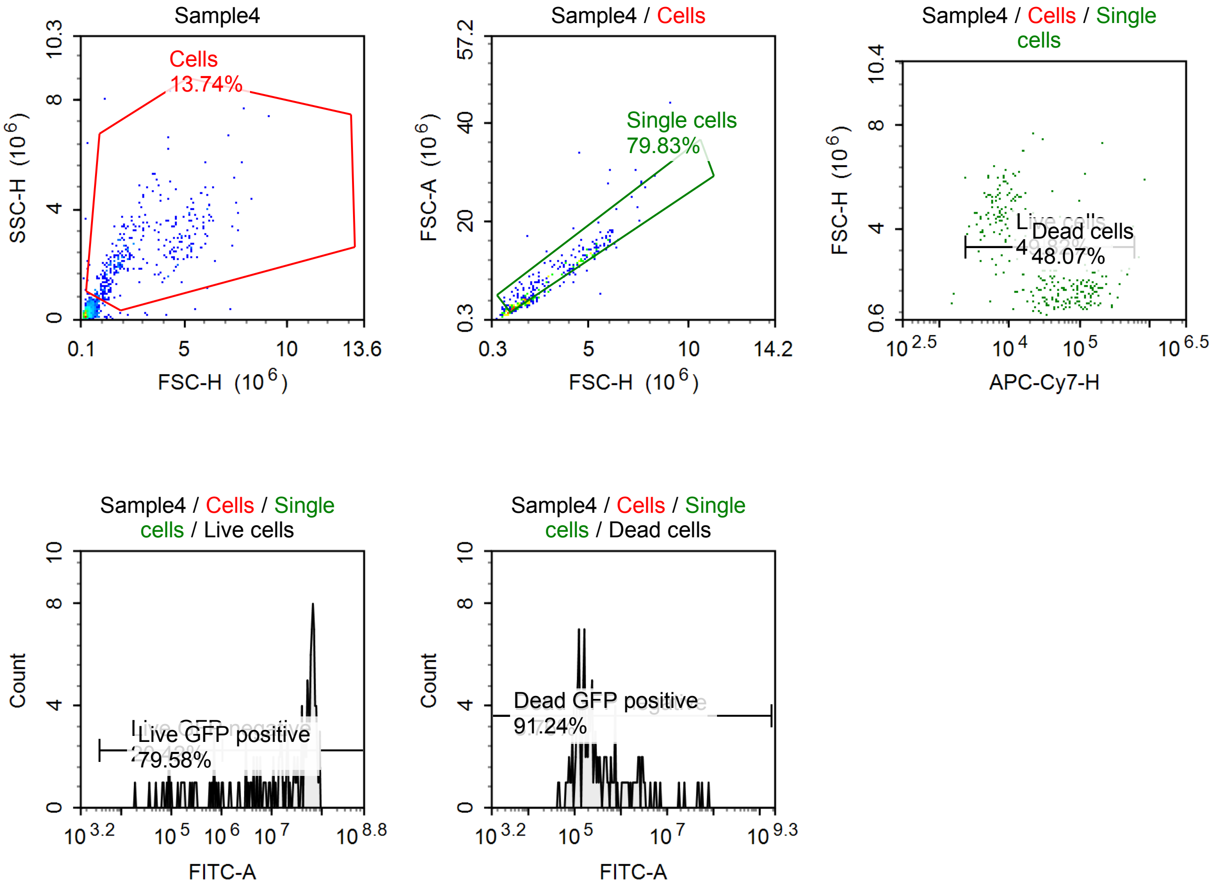


Sample Statistics of Sample4

| Gate | Count | % Parent | X | Y | Median X | Median Y |
|---|---|---|---|---|---|---|
| All | 2,598 | | | | | |
| Cells | 357 | 13.74 % | FSC-H | SSC-H | 2,179,717 | 2,277,932 |
| Single cells | 285 | 79.83 % | FSC-H | FSC-A | 2,240,212 | 6,618,841 |
| Live cells | 142 | 49.82 % | APC-Cy7-H | FSC-H | 8,605 | 4,441,816 |
| Live GFP negative | 29 | 20.42 % | FITC-A | Count | 190,811 | |
| Live GFP positive | 113 | 79.58 % | FITC-A | Count | 49,050,152 | |
| Dead cells | 137 | 48.07 % | APC-Cy7-H | FSC-H | 94,268 | 1,739,482 |
| Dead GFP negative | 12 | 8.76 % | FITC-A | Count | 74,858 | |
| Dead GFP positive | 125 | 91.24 % | FITC-A | Count | 274,522 | |

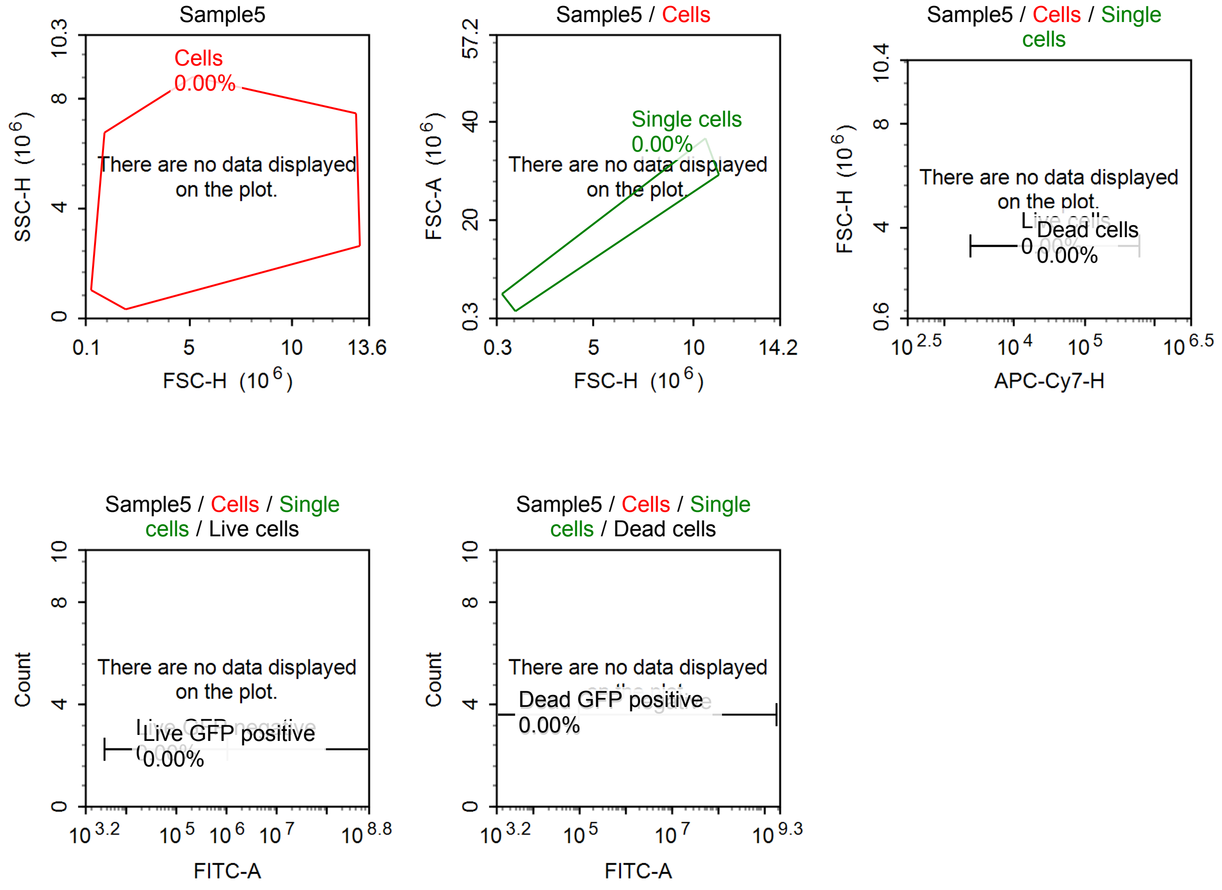


Sample Statistics of Sample5

| Gate | Count | % Parent | X | Y | Median X | Median Y |
|---|---|---|---|---|---|---|
| All | 0 | | | | | |
| Cells | 0 | 0.00 % | FSC-H | SSC-H | 0 | 0 |
| Single cells | 0 | 0.00 % | FSC-H | FSC-A | 0 | 0 |
| Live cells | 0 | 0.00 % | APC-Cy7-H | FSC-H | 0 | 0 |
| Live GFP negative | 0 | 0.00 % | FITC-A | Count | 0 | |
| Live GFP positive | 0 | 0.00 % | FITC-A | Count | 0 | |
| Dead cells | 0 | 0.00 % | APC-Cy7-H | FSC-H | 0 | 0 |
| Dead GFP negative | 0 | 0.00 % | FITC-A | Count | 0 | |
| Dead GFP positive | 0 | 0.00 % | FITC-A | Count | 0 | |

#### Specimen8

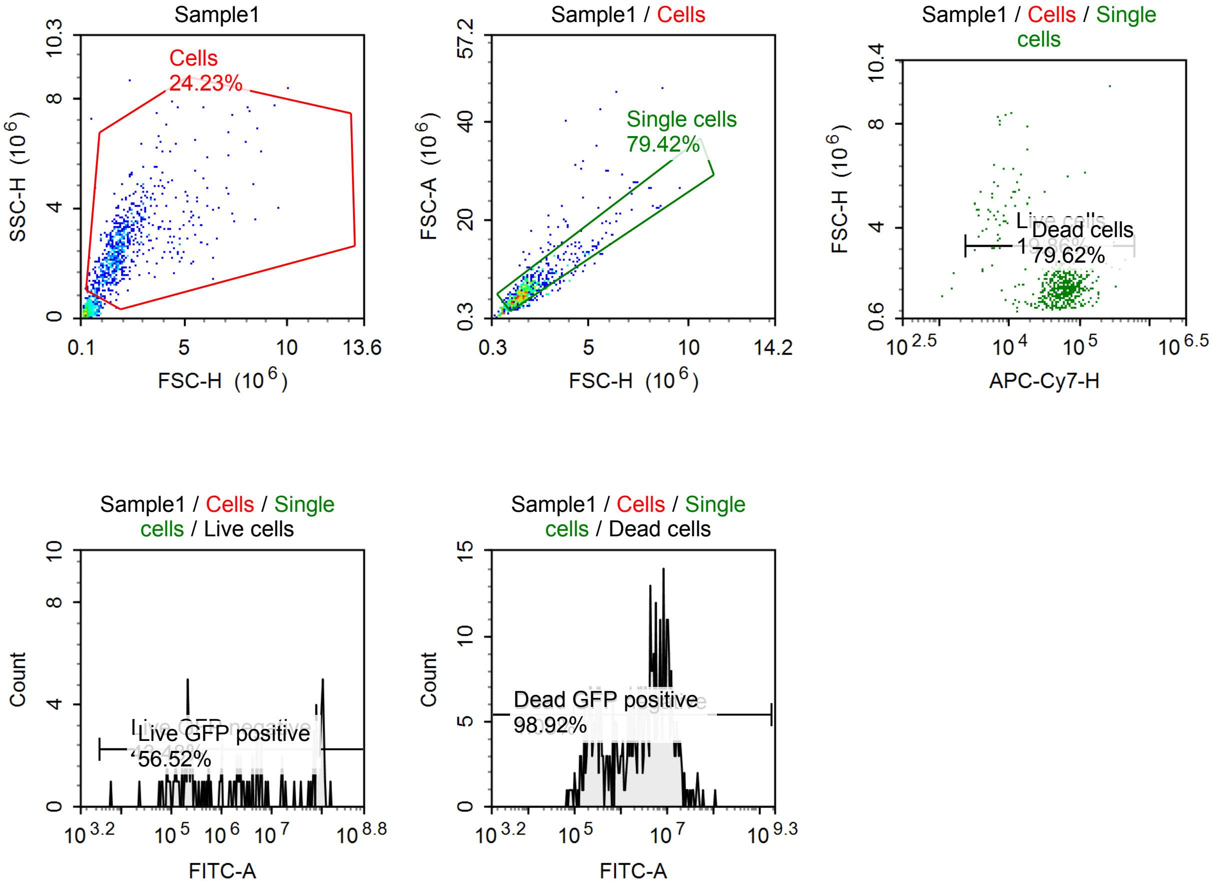


Sample Statistics of Sample1

| Gate | Count | % Parent | X | Y | Median X | Median Y |
|---|---|---|---|---|---|---|
| All | 3,009 | | | | | |
| Cells | 729 | 24.23 % | FSC-H | SSC-H | 1,775,220 | 2,512,638 |
| Single cells | 579 | 79.42 % | FSC-H | FSC-A | 1,766,063 | 5,524,328 |
| Live cells | 115 | 19.86 % | APC-Cy7-H | FSC-H | 13,417 | 3,128,848 |
| Live GFP negative | 50 | 43.48 % | FITC-A | Count | 217,964 | |
| Live GFP positive | 65 | 56.52 % | FITC-A | Count | 55,714,176 | |
| Dead cells | 461 | 79.62 % | APC-Cy7-H | FSC-H | 62,892 | 1,733,614 |
| Dead GFP negative | 5 | 1.08 % | FITC-A | Count | 78,864 | |
| Dead GFP positive | 456 | 98.92 % | FITC-A | Count | 3,592,982 | |

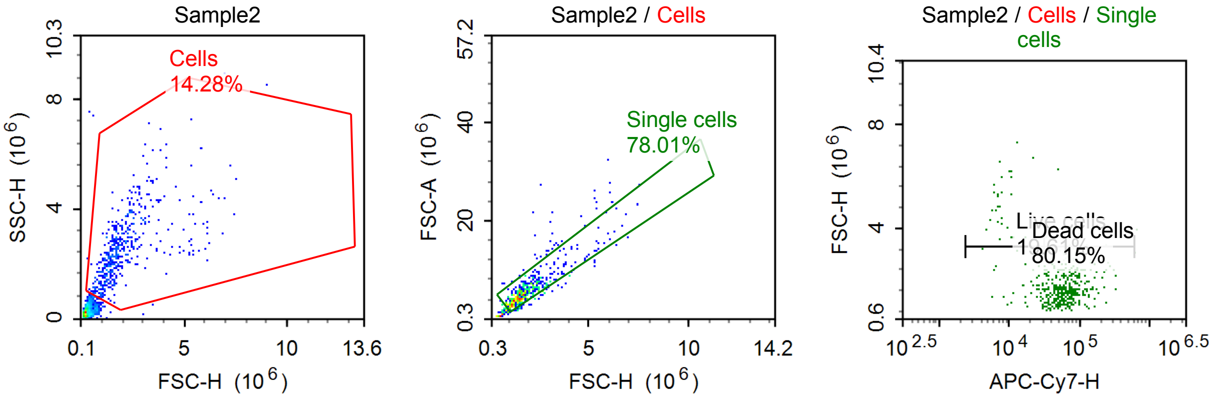

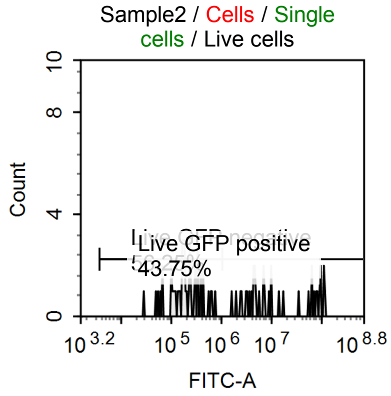


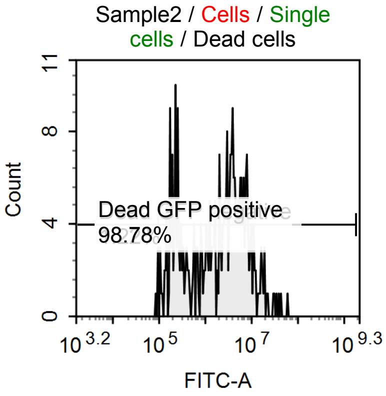


Sample Statistics of Sample2

| Gate | Count | % Parent | X | Y | Median X | Median Y |
|---|---|---|---|---|---|---|
| All | 3,663 | | | | | |
| Cells | 523 | 14.28 % | FSC-H | SSC-H | 1,664,284 | 2,403,450 |
| Single cells | 408 | 78.01 % | FSC-H | FSC-A | 1,673,825 | 5,309,051 |
| Live cells | 80 | 19.61 % | APC-Cy7-H | FSC-H | 17,181 | 2,245,868 |
| Live GFP negative | 45 | 56.25 % | FITC-A | Count | 185,331 | |
| Live GFP positive | 35 | 43.75 % | FITC-A | Count | 8,999,991 | |
| Dead cells | 327 | 80.15 % | APC-Cy7-H | FSC-H | 62,210 | 1,629,654 |
| Dead GFP negative | 4 | 1.22 % | FITC-A | Count | 93,233 | |
| Dead GFP positive | 323 | 98.78 % | FITC-A | Count | 2,463,757 | |

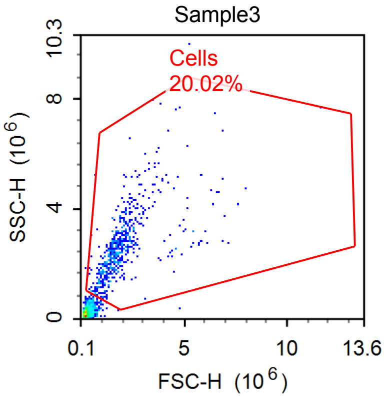


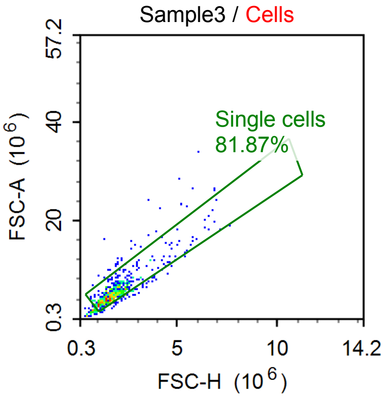


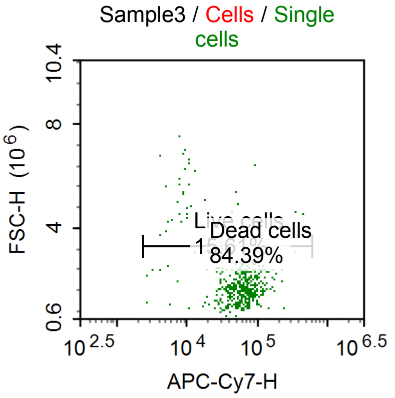


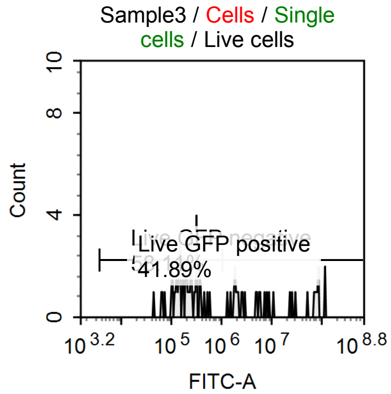


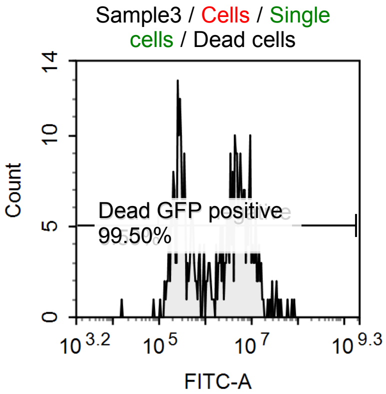

Sample Statistics of Sample3

| Gate | Count | % Parent | X | Y | Median X | Median Y |
|---|---|---|---|---|---|---|
| All | 2,892 | | | | | |
| Cells | 579 | 20.02 % | FSC-H | SSC-H | 1,737,750 | 2,498,506 |
| Single cells | 474 | 81.87 % | FSC-H | FSC-A | 1,753,520 | 5,227,001 |
| Live cells | 74 | 15.61 % | APC-Cy7-H | FSC-H | 17,451 | 2,291,785 |
| Live GFP negative | 43 | 58.11 % | FITC-A | Count | 196,365 | |
| Live GFP positive | 31 | 41.89 % | FITC-A | Count | 7,563,451 | |
| Dead cells | 400 | 84.39 % | APC-Cy7-H | FSC-H | 66,761 | 1,732,468 |
| Dead GFP negative | 2 | 0.50 % | FITC-A | Count | 45,623 | |
| Dead GFP positive | 398 | 99.50 % | FITC-A | Count | 2,439,382 | |

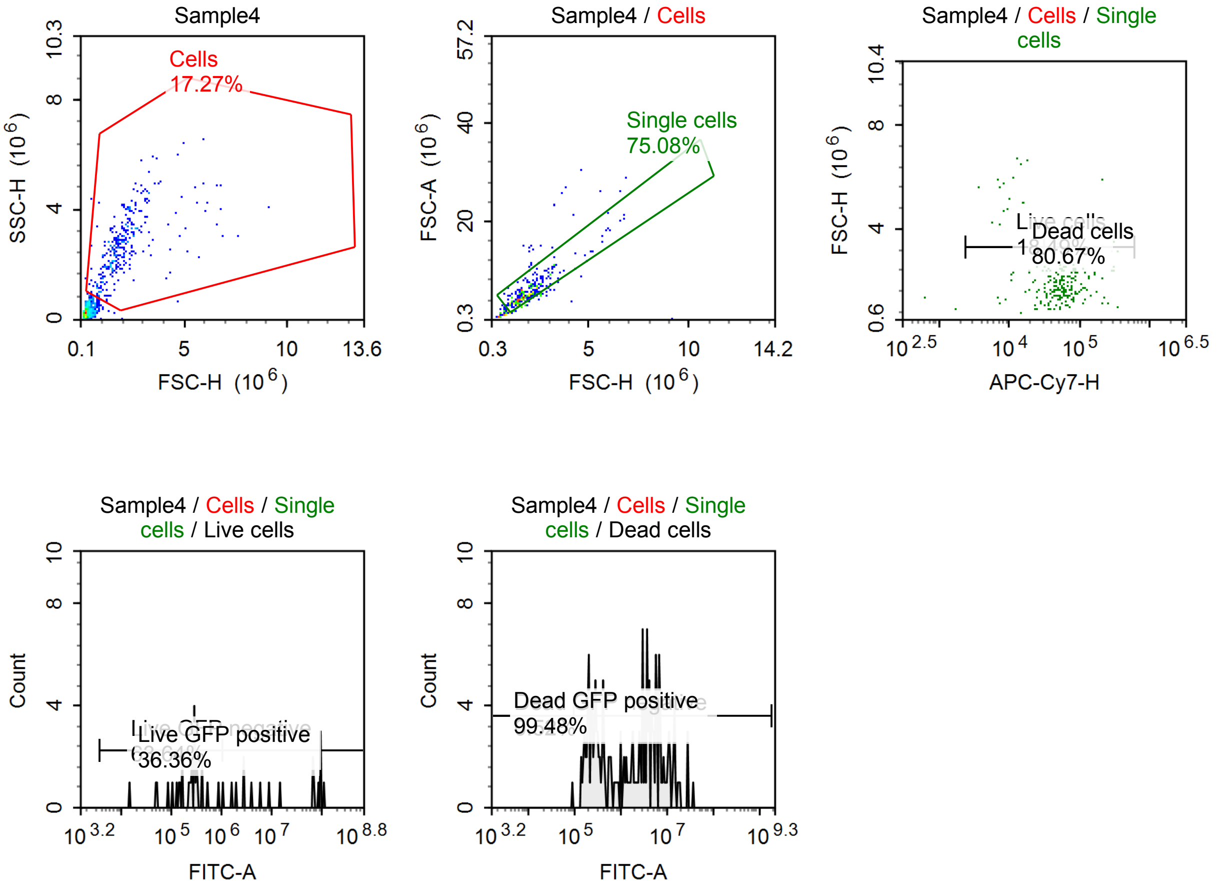


Sample Statistics of Sample4

| Gate | Count | % Parent | X | Y | Median X | Median Y |
|---|---|---|---|---|---|---|
| All | 1,836 | | | | | |
| Cells | 317 | 17.27 % | FSC-H | SSC-H | 1,797,306 | 2,640,579 |
| Single cells | 238 | 75.08 % | FSC-H | FSC-A | 1,874,521 | 5,384,051 |
| Live cells | 44 | 18.49 % | APC-Cy7-H | FSC-H | 17,505 | 2,175,405 |
| Live GFP negative | 28 | 63.64 % | FITC-A | Count | 270,355 | |
| Live GFP positive | 16 | 36.36 % | FITC-A | Count | 41,873,132 | |
| Dead cells | 192 | 80.67 % | APC-Cy7-H | FSC-H | 60,262 | 1,849,861 |
| Dead GFP negative | 1 | 0.52 % | FITC-A | Count | 92,305 | |
| Dead GFP positive | 191 | 99.48 % | FITC-A | Count | 2,340,583 | |

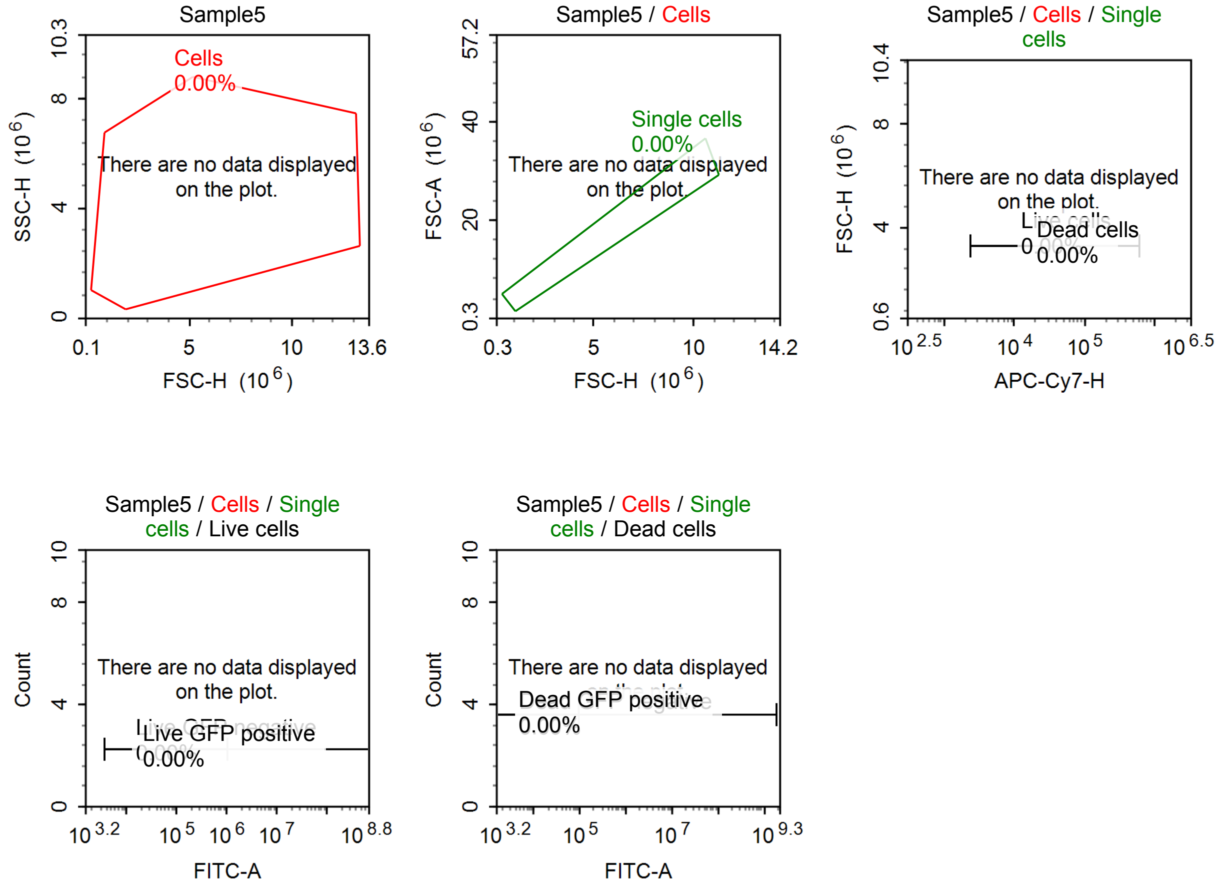


Sample Statistics of Sample5

| Gate | Count | % Parent | X | Y | Median X | Median Y |
|---|---|---|---|---|---|---|
| All | 0 | | | | | |
| Cells | 0 | 0.00 % | FSC-H | SSC-H | 0 | 0 |
| Single cells | 0 | 0.00 % | FSC-H | FSC-A | 0 | 0 |
| Live cells | 0 | 0.00 % | APC-Cy7-H | FSC-H | 0 | 0 |
| Live GFP negative | 0 | 0.00 % | FITC-A | Count | 0 | |
| Live GFP positive | 0 | 0.00 % | FITC-A | Count | 0 | |
| Dead cells | 0 | 0.00 % | APC-Cy7-H | FSC-H | 0 | 0 |
| Dead GFP negative | 0 | 0.00 % | FITC-A | Count | 0 | |
| Dead GFP positive | 0 | 0.00 % | FITC-A | Count | 0 | |

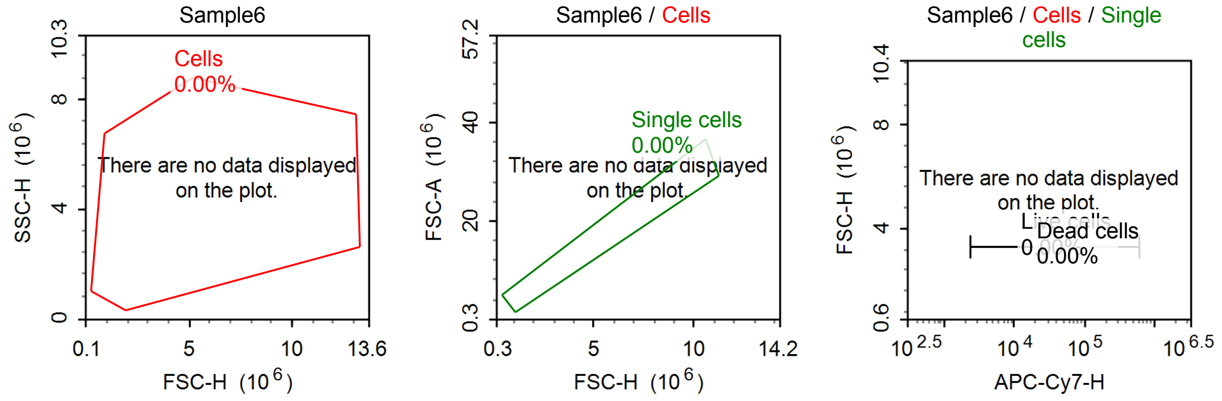

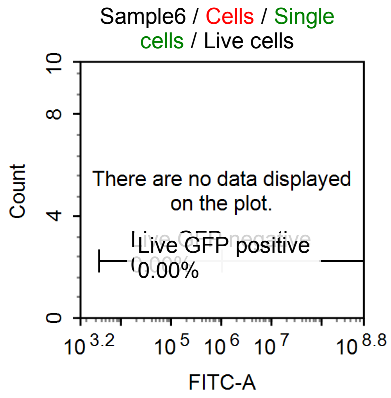


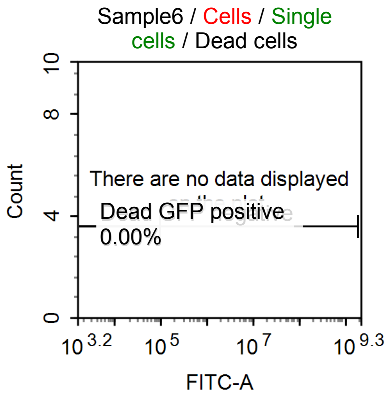


Sample Statistics of Sample6

| Gate | Count | % Parent | X | Y | Median X | Median Y |
|---|---|---|---|---|---|---|
| All | 0 | | | | | |
| Cells | 0 | 0.00 % | FSC-H | SSC-H | 0 | 0 |
| Single cells | 0 | 0.00 % | FSC-H | FSC-A | 0 | 0 |
| Live cells | 0 | 0.00 % | APC-Cy7-H | FSC-H | 0 | 0 |
| Live GFP negative | 0 | 0.00 % | FITC-A | Count | 0 | |
| Live GFP positive | 0 | 0.00 % | FITC-A | Count | 0 | |
| Dead cells | 0 | 0.00 % | APC-Cy7-H | FSC-H | 0 | 0 |
| Dead GFP negative | 0 | 0.00 % | FITC-A | Count | 0 | |
| Dead GFP positive | 0 | 0.00 % | FITC-A | Count | 0 | |

#### Specimen9

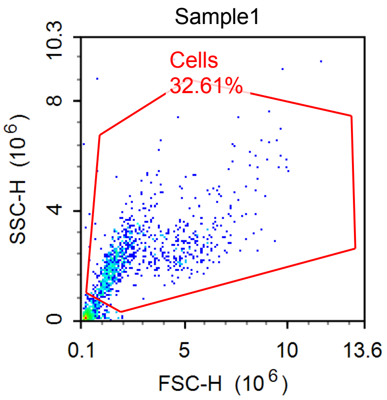


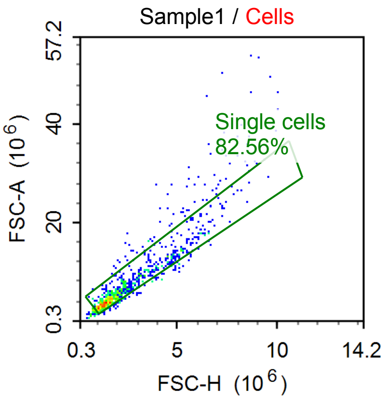


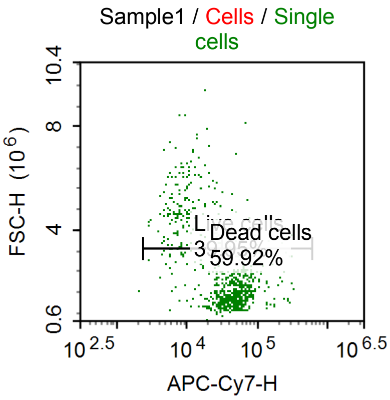


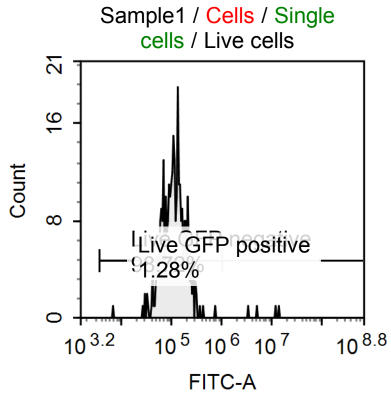


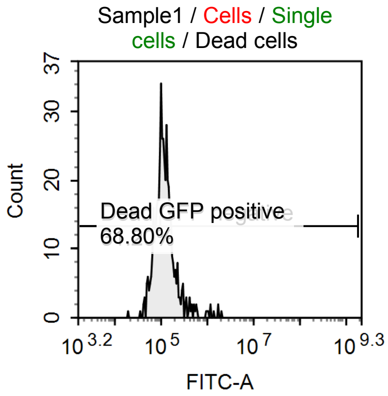

Sample Statistics of Sample1

| Gate | Count | % Parent | X | Y | Median X | Median Y |
|---|---|---|---|---|---|---|
| All | 2,901 | | | | | |
| Cells | 946 | 32.61 % | FSC-H | SSC-H | 1,927,831 | 2,257,260 |
| Single cells | 781 | 82.56 % | FSC-H | FSC-A | 1,787,472 | 5,088,749 |
| Live cells | 312 | 39.95 % | APC-Cy7-H | FSC-H | 13,390 | 3,520,610 |
| Live GFP negative | 308 | 98.72 % | FITC-A | Count | 115,664 | |
| Live GFP positive | 4 | 1.28 % | FITC-A | Count | 8,815,436 | |
| Dead cells | 468 | 59.92 % | APC-Cy7-H | FSC-H | 52,691 | 1,609,919 |
| Dead GFP negative | 146 | 31.20 % | FITC-A | Count | 75,492 | |
| Dead GFP positive | 322 | 68.80 % | FITC-A | Count | 133,395 | |

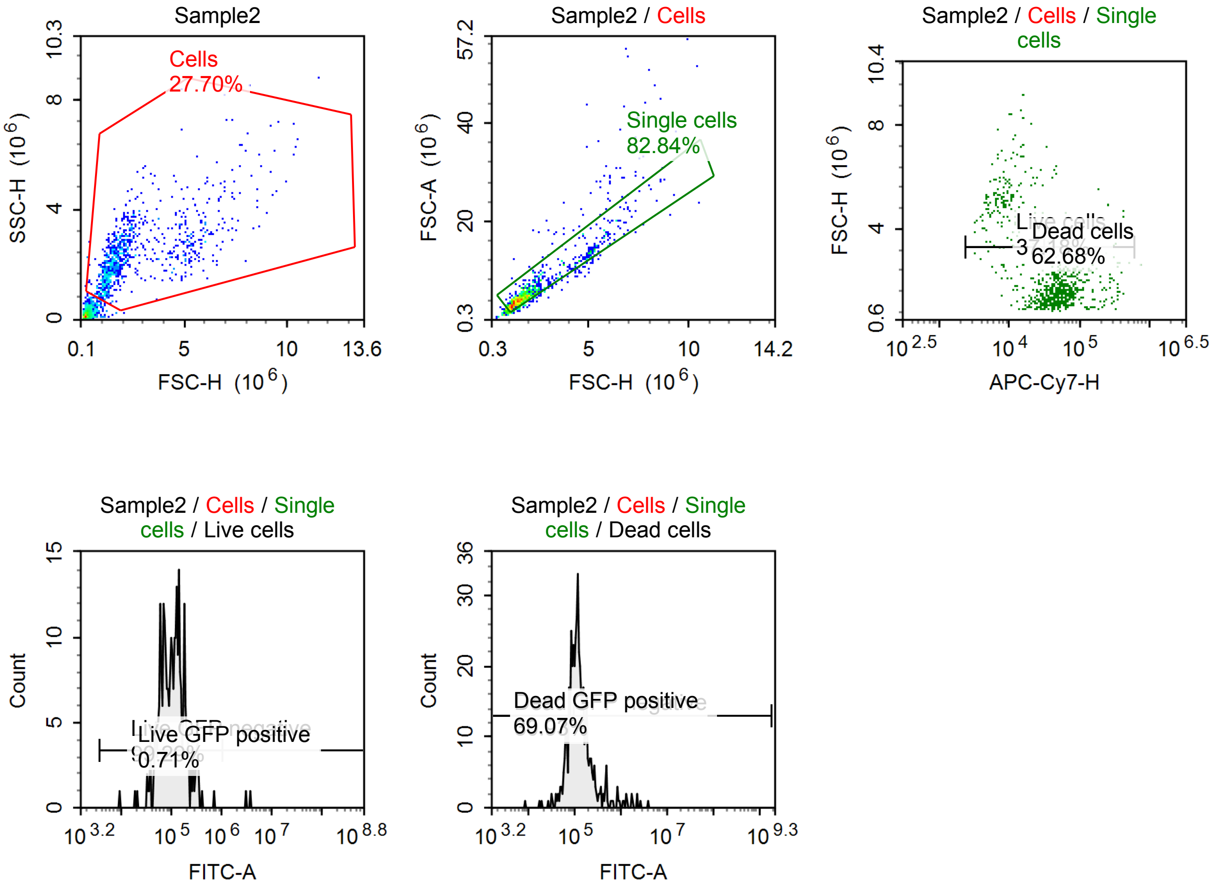


Sample Statistics of Sample2

| Gate | Count | % Parent | X | Y | Median X | Median Y |
|---|---|---|---|---|---|---|
| All | 3,281 | | | | | |
| Cells | 909 | 27.70 % | FSC-H | SSC-H | 1,835,778 | 2,251,630 |
| Single cells | 753 | 82.84 % | FSC-H | FSC-A | 1,792,022 | 5,019,991 |
| Live cells | 280 | 37.18 % | APC-Cy7-H | FSC-H | 12,861 | 3,877,384 |
| Live GFP negative | 278 | 99.29 % | FITC-A | Count | 109,091 | |
| Live GFP positive | 2 | 0.71 % | FITC-A | Count | 3,543,074 | |
| Dead cells | 472 | 62.68 % | APC-Cy7-H | FSC-H | 57,053 | 1,628,033 |
| Dead GFP negative | 146 | 30.93 % | FITC-A | Count | 77,858 | |
| Dead GFP positive | 326 | 69.07 % | FITC-A | Count | 137,702 | |

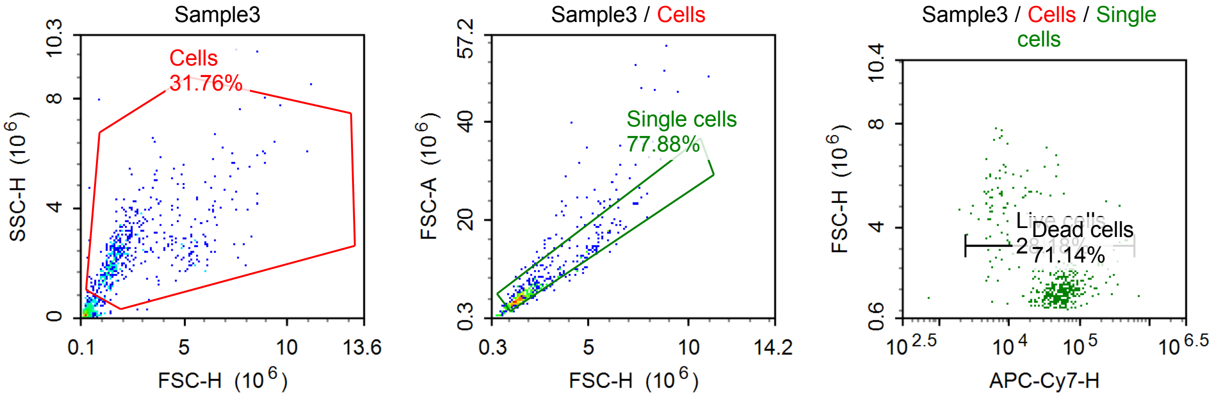


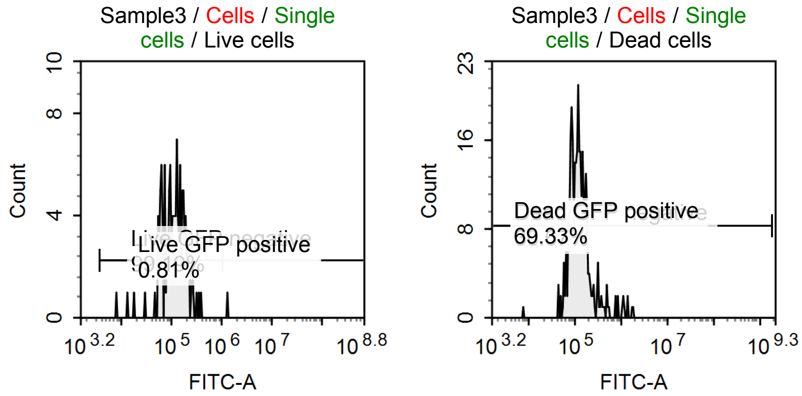


Sample Statistics of Sample3

| Gate | Count | % Parent | X | Y | Median X | Median Y |
|---|---|---|---|---|---|---|
| All | 1,779 | | | | | |
| Cells | 565 | 31.76 % | FSC-H | SSC-H | 1,864,056 | 2,352,266 |
| Single cells | 440 | 77.88 % | FSC-H | FSC-A | 1,725,366 | 4,996,357 |
| Live cells | 124 | 28.18 % | APC-Cy7-H | FSC-H | 9,580 | 3,739,335 |
| Live GFP negative | 123 | 99.19 % | FITC-A | Count | 117,435 | |
| Live GFP positive | 1 | 0.81 % | FITC-A | Count | 1,365,767 | |
| Dead cells | 313 | 71.14 % | APC-Cy7-H | FSC-H | 59,784 | 1,628,874 |
| Dead GFP negative | 96 | 30.67 % | FITC-A | Count | 81,528 | |
| Dead GFP positive | 217 | 69.33 % | FITC-A | Count | 140,343 | |

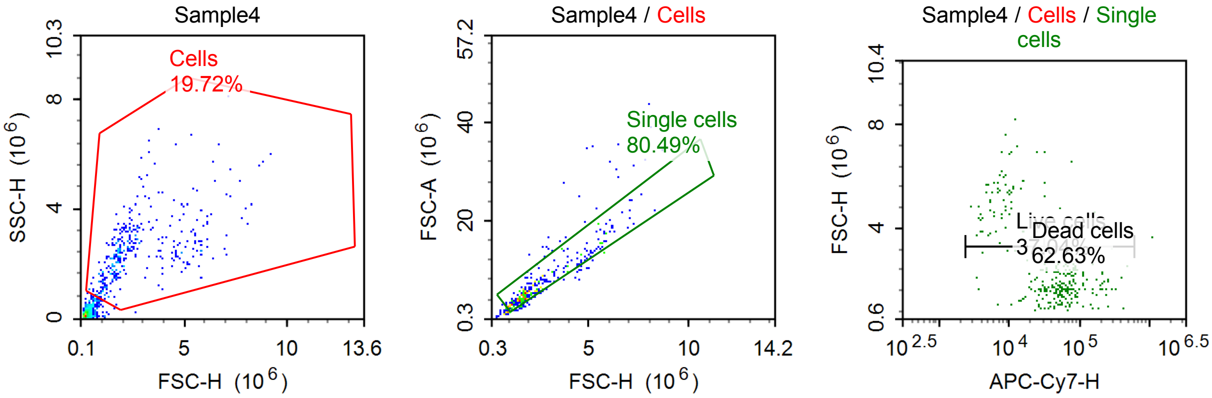

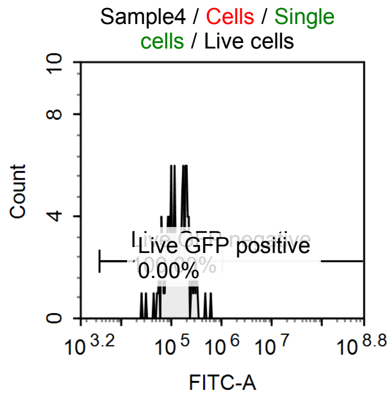


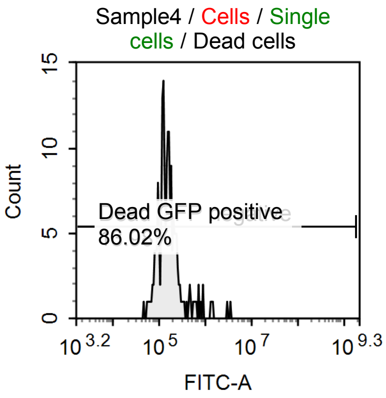


Sample Statistics of Sample4

| Gate | Count | % Parent | X | Y | Median X | Median Y |
|---|---|---|---|---|---|---|
| All | 1,871 | | | | | |
| Cells | 369 | 19.72 % | FSC-H | SSC-H | 2,018,155 | 2,419,907 |
| Single cells | 297 | 80.49 % | FSC-H | FSC-A | 2,021,587 | 5,793,119 |
| Live cells | 110 | 37.04 % | APC-Cy7-H | FSC-H | 10,235 | 4,213,149 |
| Live GFP negative | 110 | 100.00 % | FITC-A | Count | 139,770 | |
| Live GFP positive | 0 | 0.00 % | FITC-A | Count | 0 | |
| Dead cells | 186 | 62.63 % | APC-Cy7-H | FSC-H | 65,056 | 1,811,818 |
| Dead GFP negative | 26 | 13.98 % | FITC-A | Count | 81,076 | |
| Dead GFP positive | 160 | 86.02 % | FITC-A | Count | 163,198 | |

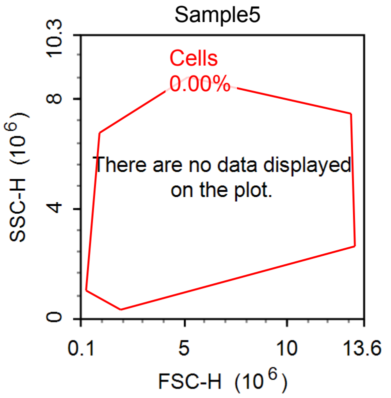


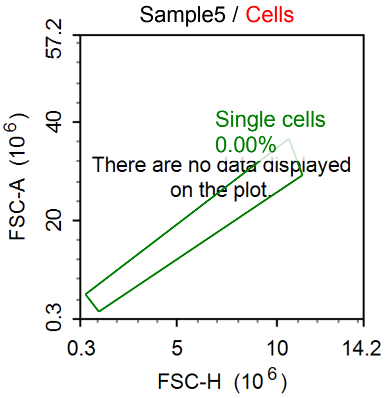


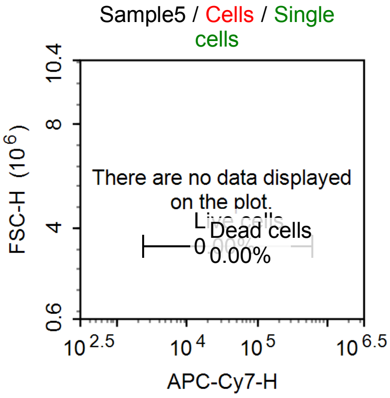


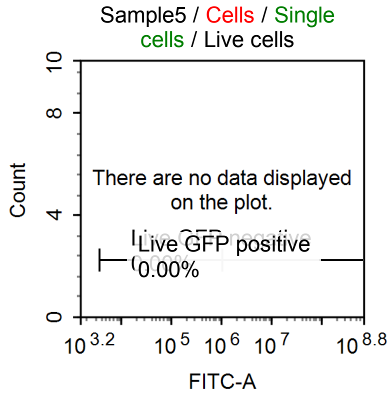


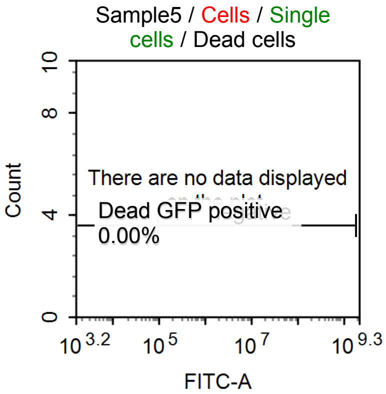

Sample Statistics of Sample5

| Gate | Count | % Parent | X | Y | Median X | Median Y |
|---|---|---|---|---|---|---|
| All | 0 | | | | | |
| Cells | 0 | 0.00 % | FSC-H | SSC-H | 0 | 0 |
| Single cells | 0 | 0.00 % | FSC-H | FSC-A | 0 | 0 |
| Live cells | 0 | 0.00 % | APC-Cy7-H | FSC-H | 0 | 0 |
| Live GFP negative | 0 | 0.00 % | FITC-A | Count | 0 | |
| Live GFP positive | 0 | 0.00 % | FITC-A | Count | 0 | |
| Dead cells | 0 | 0.00 % | APC-Cy7-H | FSC-H | 0 | 0 |
| Dead GFP negative | 0 | 0.00 % | FITC-A | Count | 0 | |
| Dead GFP positive | 0 | 0.00 % | FITC-A | Count | 0 | |
